# Modulation of calcium signaling on demand to decipher the molecular mechanisms of primary aldosteronism

**DOI:** 10.1101/2024.04.29.591306

**Authors:** Bakhta Fedlaoui, Teresa Cosentino, Zeina R. Al Sayed, Isabelle Giscos-Douriez, Nicolo Faedda, May Fayad, Jean-Sebastien Hulot, Chris Magnus, Scott M Sternson, Simon Travers-Allard, Stephanie Baron, Fabio L Fernandes-Rosa, Maria-Christina Zennaro, Sheerazed Boulkroun

**Affiliations:** Université Paris Cité, Inserm, PARCC, F-75015 Paris France; CIC1418 and DMU CARTE, Assistance Publique Hôpitaux de Paris (AP-HP), Hôpital Européen Georges Pompidou, F-75015 Paris, France; Howard Hughes Medical Institute & Department of Neurosciences, University of California, San Diego, San Diego, United States; Service de Physiologie, Assistance Publique Hôpitaux de Paris (AP-HP), Hôpital Européen Georges Pompidou, F-75015 Paris, France; Service de Génétique, Assistance Publique Hôpitaux de Paris (AP-HP), Hôpital Européen Georges Pompidou, F-75015 Paris, France

**Keywords:** primary aldosteronism, adrenal cortex, aldosterone biosynthesis, calcium signaling, cell proliferation

## Abstract

Primary aldosteronism (PA) is the most common form of secondary hypertension. Major advances have been made in our understanding of PA with the identification of germline and somatic mutations in ion pumps and channels. These mutations lead to the activation of calcium signalling, the major trigger of aldosterone biosynthesis.

To elucidate the molecular mechanisms underlying the development of PA, we established an adrenocortical cell model in which we can modulate sodium entry into the cells “on demand” leading to calcium signalling activation. These cells recapitulated the major features of *KCNJ5* mutations, the most frequent genetic alteration identified in Aldosterone-Producing-Adenoma. Activation of calcium signalling was associated with increased aldosterone biosynthesis and decreased cell proliferation. RNA sequencing and steroidome analyses revealed unique profiles associated with Na^+^ entry. Altogether, this work offers valuable insights into the role of sodium-induced calcium signalling in PA development and paves the way for developing new therapeutic strategies.

## Introduction

The adrenal gland, a crucial endocrine organ located on top of each kidney, consists of two distinct regions: the outer adrenal cortex and the inner adrenal medulla. The adrenal cortex is further divided into three zones: the *zona glomerulosa* (ZG), *zona fasciculata*, and *zona reticularis*, each specialized in hormone production due to expression of specific enzyme. Among these hormones, aldosterone, produced by the adrenal ZG, plays an important role in regulating salt and fluid balance, thereby controlling arterial blood pressure. As a key component of the renin-angiotensin-aldosterone system (RAAS), its production is mainly stimulated by angiotensin II (Ang II), which increases in response to volume depletion, and elevated plasma potassium (K^+^) levels^1,2^. The stimulation by AngII or K^+^ results in ZG cell membrane depolarization and opening of voltage-gated calcium (Ca^2+^) channels, leading to an increased intracellular Ca^2+^ concentration. AngII also signals through its type 1 receptor (AT1R) to stimulate Ca^2+^ release from the endoplasmic reticulum. The activation of calcium signaling triggers a phosphorylation cascade that leads to increased transcription of *CYP11B2*, coding for aldosterone synthase, and aldosterone biosynthesis^1^.

Deregulation of the mechanisms regulating adrenal aldosterone biosynthesis results in primary aldosteronism (PA). PA is the leading cause of secondary hypertension affecting approximately 5-10% of hypertensive patients and up to 20% of those with treatment resistant hypertension^3–6^. It is characterized by hypertension with elevated aldosterone levels, low plasma renin concentration, increased aldosterone to renin ratio and often associated with hypokalemia. The main subtypes of PA are bilateral adrenal hyperplasia and aldosterone-producing adenomas (APA), accounting together for 95% of cases. PA is associated with an increased risk of cardiometabolic and renal complications^7^ due to the major adverse effects of aldosterone excess; therefore, early diagnosis and appropriate treatment of PA are essential.

In the past decade, research has uncovered mutations in ion channels (*KCNJ5*^8^, *CACNA1D*^9,10^, *CACNA1H*^11,12^, *CLCN2*^13,14^, *SLC30A1*^15^) and pumps (*ATP1A1*^9,16^, *ATP2B3*^16^), as principal causes of APA and familial forms of PA. These mutations enhance either directly or indirectly intracellular calcium concentrations, the main trigger for aldosterone biosynthesis. Notably, in the case of *KCNJ5,* which encodes the G protein-coupled inwardly rectifying K^+^ channel (GIRK4), the majority of mutations cluster near the channel’s ion-selective filter, leading to a loss of selectivity for K^+^ ions in favour of an intracellular sodium (Na^+^) influx^8^. These mutations lead to cell membrane depolarization, the opening of voltage-gated calcium channels, an increase of intracellular calcium concentration, activation of calcium signalling pathways, and ultimately an increase of *CYP11B2* expression and aldosterone biosynthesis^8^.

While the role of *KCNJ5* mutations in autonomous aldosterone production has been well established, their role in modulating cell proliferation is still under debate^17–20^. Chemogenetic tools allow to manipulate specific ion fluxes via modified ion channels with pharmacologically selective properties, known as Pharmacologically Selective Actuator Modules (PSAM). PSAM, comprising mutated ligand-binding domains and selective ionic pore domains respond to Pharmacologically Selective Effector Molecules (PSEM), inducing channel opening and allowing specific ions to flow through the activated channel^21,22^. Here we employed second-generation chemogenetic tools to established a human adrenal cell model in which we could modulate sodium entry on demand, by introducing the α7-5HT3 chimeric receptor. Activation of this receptor by uPSEM-817 was used to study the effect of Na^+^ entry into the cells, mimicking molecular abnormalities observed in the presence of *KCNJ5* mutations. We assessed membrane depolarization, intracellular calcium concentration, and impact on cell proliferation. Additionally, we conducted RNA sequencing and steroidomic analyses following treatment with uPSEM-817, AngII, and K^+^.

## Material and Methods

### Cell culture and electroporation

The H295R-S2 cell line, a subclone of the H295R human adrenocortical carcinoma cell, was kindly provided by Dr. William E. Rainey^23^ and cultured in complete medium containing DMEM/Eagle’s F12 medium (1:1) (GIBCO, Life Technologies, Carlsbad, CA) supplemented with 2% Ultroser G (Sartorius, France), 1% insulin/transferrin/selenium premix (BD Bioscineces), 7.5 mM HEPES (GIBCO), 1% penicillin and streptomycin (GIBCO) and 20 mg/mL G418 (Thermo Fisher Scientific). Cells were maintained at 37 °C under a humid atmosphere of 95% air and 5% CO_2_.

The α7-5HT3 chimeric receptor consists of the ligand binding domain of the nicotinic receptor fused with the ionic pore domain of the serotonin receptor type 3. The sequence has been inserted into pcDNA3.1 vector^21^.

5 million of H295R-S2 cells were seeded into a 100 mm tissue culture dish. After 24 hours, cells were tripsinized, counted and 3.10^6^ cells resuspended in 100 μl Nucleofactor R solution and electroporated with 3 µg of plasmid (pcDNA3 containing or not α7-5HT3 cDNA) using the Amaxa nucleofactor kit R (Lonza) according to the manufacturer instructions. After electroporation, the mixed populations were amplified in under selection pressure with G418. Pure clones were isolated by picking clones after limited dilution of the cells in order to isolate one cell at an optimum distance from the other in a 200 mm diameter plate. Colonies were, afterwards, isolated in wells and amplified for further characterization.

### Functional studies in α7-5HT3 expressing cells

For functional experiments, H295R-S2 cells were seeded at 400,000 cells in 12-wells plates in complete medium. After 24h, cells were serum-deprived in DMEM/F12 containing 0.1% Ultroser G for 24h and then incubated for 8h and 24h with fresh serum-deprived medium with no secretagogue (Basal) or with 10^-8^ M AngII (Sigma-Aldrich), 12 mM K^+^ (Sigma-Aldrich) or uPSEM-817 (Tocris Biosciences) from a range of concentration going from 10^-9^ to 10^-5^ M. At the end of the incubation period, supernatants were collected for steroid profiling and cells were washed with PBS and frozen for protein or RNA extraction. For cell cycle investigation, cells were not serum-deprived and incubated 24h after seeding with no secretagogue (Basal) or with 10^-8^ M AngII (Sigma-Aldrich), 12 mM K^+^ (Sigma-Aldrich) or uPSEM-817 (Tocris Biosciences) from a range of concentration going from 10^-9^ to 10^-5^ M. For calcium channel inhibition investigation cells were pre-treated for 2h with 10^-6^ M of nifedipine or 10^-6^M of mibefradil for calcium measure or co-treated with 10^-9^, 10^-7^ or 10^-5^ M u-PSEM-817, 10^-8^ M AngII or 12mM K^+^.

### RNA extraction and RT-qPCR

Total RNA was extracted in TRIzol reagent (Ambion, Life Technologies) according to the manufacturer’s recommendations. After DNase I treatment (Life Technologies), 500 ng of total RNA was reverse transcribed (iScript reverse transcriptase, Bio-Rad). The quantitative PCR was performed using SYBR Green (Sso Advanced Universal SYBR Green Supermix, Bio-Rad) on a C1000 touch thermal cycler from Bio-Rad (CFX96 Real-Time System), according to the manufacturer’s instructions. Controls without template were included to verify that fluorescence was not overestimated from primer dimer formation or PCR contaminations. The primers used for qPCR are described in Supplementary Table S1. Normalization for RNA quantity and reverse transcriptase efficiency was performed against three reference genes (geometric mean of the expression of *18S* rRNA, *HPRT1*, and *GAPDH*), in accordance with the MIQE guidelines^24^. Quantification was performed using the standard curve method. Standard curves were generated using serial dilutions from a cDNA pool of all samples of each experiment, yielding a correlation coefficient of at least 0.98 in all experiments. Data are reported as fold change over control.

### Immunofluorescence

H295R-S2 cells (400 000 cells/mL) were plated 24 hours in poly-L-lysine coated coverslips. Cells were then incubated with 1 µg/mL of Alexa 594 conjugated α-Bungarotoxin solution in complete medium (Invitrogen) in complete medium for 15 minutes at 37°C. Cells were fixed for 30 minutes with PFA 4% at room temperature, washed with PBS and nuclei counterstained with DAPI. For Dab-2 and phalloidin staining, treated cells were fixed in 4% cold PFA during 15 minutes at room temperature, washed twice with PBS and permeabilized with PBS-Triton 0.5% during 10 minutes. Cells were afterwards washed with PBS-tween 0.1% once and blocked in PBS-BSA 1% during 30 minutes. Antibodies were diluted in PBS-BSA 1% at 1/2000 for phalloidin-TRITC (Sigma #P1951) and 1/200 for dab-2. For Dab-2 staining, primary antibody incubation was followed by secondary antibody incubation at 1/400.

All microscopic examinations were done with a x40 and x63 objectives lens using Leica SP8 confocal microscope.

### Measurement of intracellular Ca^2+^ levels

Cytosolic free Ca^2+^ activity was measured using the ratiometric fluorescent Ca^2+^ sensitive dye Fura-2-AM (Thermo Fisher). Cells were loaded at 37°C for 60 minutes with 4 μM Fura-2-AM in the presence of 1X Power Load permeabilizing reagent (Thermo Fisher). Injection of AngII (10^-8^M), K^+^(12 mM) and uPSEM-817 (from 10^-9^ M to 10^-5^ M) occurs 18 sec after the beginning of fluorescence record. Mean fluorescence ratios of emission at 490 to 530 nm after excitation at 340 nm and 380 nm were calculated

### Aldosterone and protein assay

Aldosterone concentrations were measured in cell culture supernatants after treatments using indirect ELISA. Aldosterone antibody (AB A2E11) and aldosterone-3-CMO biotin were kindly provided by Dr Celso Gomez-Sanchez^25^. Results were normalized by cell protein concentration (determined using Bradford protein assay).

### Perforated patch-clamp

Electrophysiological measurements by whole-cell patch-clamp were performed on H295R-S2 control cells and clones 17 and 42 using an Axopatch 200B amplifier (Molecular Devices) controlled by Axon pClamp 11 software through an A/D converter (Digidata 1550B; Molecular Devices). Borosilicate glass pipettes with 5 to 10 MΩ were used for the recordings. Cellular resting membrane potential (Em) was acquired in a perforated-patch configuration using 0.22 mM amphotericin-B. The patch pipette was back-filled with a solution containing in mM: 100 K-gluconate, 30 KCl, 4 NaCl, 1 EGTA, 3 MgATP, 1 MgCl2, and 10 HEPES (pH 7.2 adjusted with KOH). The cells bathed in a Tyrode solution containing (in mM): 140 NaCl, 5 KCl, 10 glucose, 1.8 MgCl2, 1.8 CaCl2, and 10 HEPES (pH 7.4 adjusted with NaOH). All recordings were performed at 37°C. Data were collected from at least three independent passages from each cell line and analysed using Clampfit 11 software (Molecular Devices). Nifedipine and mibefradil were solubilized in dimethyl sulfoxide (DMSO) and water respectively and used at 10 µM concentration in a solution containing 140 mM NaCl, 5 mM KCl, 1.8 mM CaCl_2_, 1.8 mM MgCl_2_, 30 mM mannitol, 10 mM HEPES; (pH 7.4 adjusted with NaOH).

### LC-MS/MS steroid profiling

Nineteen steroids were measured simultaneously by Liquid Chromatography coupled to tandem Mass Spectrometry: 18-oxocortisol, 18-hydroxycortisol, aldosterone, cortisone, cortisol, 11-deoxycortisol, 21-deoxycortisol, 18-hydroxy-11-deoxycorticosterone, 11-deoxycorticosterone, 18-hydroxycorticosterone, corticosterone, 17-hydroxyprogesterone, delta-4-androstenedione in a 13-minute run. The complete steroid profiling procedure is described in ^26^.

### Cell proliferation

CellTiter 96® Aqueous One solution Cell Proliferation Assay was used accordingly to manufacturer instruction (Promega). Briefly, 50 000 cells were seeded into 96 well plates in complete medium. After 24 hours, cells were treated with different concentrations of uPSEM-817 or with potassium. Cell viability was evaluated after 24hours and 72hours of treatment.

### Flow cytometry analysis of the cell cycle

Cell suspensions (1×10^6^ cells/mL) were treated during 8, 24, 48 or 72 hours with different concentrations of uPSEM817, AngII or K^+^, prepared by trypsinization and washed once with phosphate buffered saline (PBS). The cells were then fixed with 70% ethanol at - 20°C and resuspended in PBS containing 50µg/mL of propidium iodide (PI) and 100µg/mL of RNAse A. Cells were appropriately gated in DIVA recording 100 000 events and further analyzed in FlowJo.

### RNAseq experiments

Gene expression profiles was determined for cells treated or not for 8 hours and 24 hours with AngII (10-8 M), K^+^ (12 mM) and uPSEM-817 (10-7 M). After RNA extraction with Trizol, RNA concentrations were obtained using nanodrop or a fluorometric Qubit RNA assay (Life Technologies, Grand Island, New York, USA). The quality of the RNA (RNA integrity number) was determined on the Agilent 2100 Bioanalyzer (Agilent Technologies, Palo Alto, CA, USA) as per the manufacturer’s instructions.

To construct the libraries, 250 ng of high-quality total RNA sample (RIN range between 6.6 and 9.68.8) was processed using Stranded mRNA Prep kit (Illumina) according to manufacturer instructions. Briefly, after purification of poly-A containing mRNA molecules, mRNA molecules are fragmented and reverse-transcribed using random primers. Replacement of dTTP by dUTP during the second strand synthesis will permit to achieve the strand specificity. Addition of a single A base to the cDNA is followed by ligation of Illumina adapters.

Libraries were quantified by Q bit and profiles were assessed using the DNA High Sensitivity LabChip kit on an Agilent Bioanalyzer. Libraries were sequenced on an Illumina Nextseq 500 instrument using 75 base-lengths read V2 chemistry in a paired-end mode.

After sequencing, a primary analysis based on AOZAN software (ENS, Paris) was applied to demultiplex and control the quality of the raw data (based of FastQC modules / version 0.11.5). Fastq files were aligned using STAR algorithm (version 2.7.6a), on the Ensembl release 101 reference. Reads were then count using RSEM (v1.3.1) and the statistical analyses on the read counts were performed with R (version 3.6.3) and the DESeq2 package (DESeq2_1.26.0) to determine the proportion of differentially expressed genes between two conditions. We used the standard DESeq2 normalization method (DESeq2’s median of ratios with the DESeq function), with a pre-filter of reads and genes (reads uniquely mapped on the genome, or up to 10 different loci with a count adjustement, and genes with at least 10 reads in at least 3 different samples). Following the package recommendations, we used the Wald test with the contrast function and the Benjamini-Hochberg FDR control procedure to identify the differentially expressed genes. R scripts and parameters are availables on the plateform, https://github.com/GENOM-IC-Cochin/RNA-Seq_analysis.

Analyses were performed using the Radish application (https://github.com/GENOM-IC-Cochin/Radish). Genes with adjusted P value <0.05 and log2 fold changes <-0.5 or >0.5 for treatment with uPSEM-817 and log2 fold changes <-1 or >1 for treatments with AngII and K+ were considered to be down- or up-regulated.

Functional annotation clustering and pathway enrichment analyses were performed using DAVID (Database for Annotation, Visualization, and Integrated Discovery) Functional Annotation Tool^27,28^. For the analyses, we used the list of differentially expressed genes in 8h or 24h uPSEM-817 vs basal, 8h or 24h AngII vs basal and 8h or 24h K^+^ vs basal.

### Statistics

The number of independent experiments (n) refer to the number of cells or dishes studied to calculate mean values ± SEM or medians. The measurements were carried out at different days and from different cell preparations using different cell passages to ensure the reproducibility of the experiments. For patch-clamp experiments, each cell was analyzed separately.

Data were analysed in Prism10 software (GraphPad, San Diego, CA) using the appropriate statistical tests as indicated in the text. The normal distribution of the data was checked by the ShapiroWilk test. Quantitative variables were reported as means ±s.e.m. when a Gaussian distribution was present or as medians and interquartile ranges when no Gaussian distribution was present. Pairwise comparisons were done with unpaired t tests and Mann–Whitney tests, respectively; multiple comparisons were done with the ANOVA test followed by a test for pairwise comparison of subgroups according to Bonferroni when a Gaussian distribution was present or Kruskal–Wallis followed by Dunn’s test when no Gaussian distribution was present. A P value of < 0.05 was considered significant.

## Results

### Characterization of a cell model expressing the α7-5HT3 receptor

To assess the impact of modulating intracellular Na^+^ concentration on adrenal cell function, we established stable expression of the chimeric α7-5HT3 receptor in H295R-S2 cells. This receptor was formed by combining the extracellular ligand-binding domain of the α7 nicotinic acetylcholine receptor and the ion pore domain of the serotonin receptor 5HT3^21^. Expression of the α7-5HT3 receptor was investigated by RT-qPCR and found exclusively in cells transfected with the vector containing its genetic sequence (Figure S1A), but not in control cells transfected with an empty vector. Upon treatment with 12 mM of K^+^ both cell lines exhibited a rapid increase in intracellular Ca^2+^ concentration (Figure S1B). However, treatment with different concentrations of varenicline, a nicotinic receptor partial agonist and cholinergic agonist, resulted in a dose-dependent increase in intracellular Ca^2+^ concentration in cells expressing the α7-5HT3 receptor only (Figure S1B), indicating a specific effect on the chimeric receptor. This was associated with an increase in *CYP11B2* mRNA expression for varenicline concentrations ranging from 10^-8^ to 10^-5^ M (Figure S1C).

To further characterize the cells expressing the α7-5HT3 receptor, monoclonal cell populations were obtained via limiting dilution under antibiotic selection pressure. Among the 19 monoclonal cell populations selected, six showed α7-5HT3 receptor mRNA expression (Figure S1D), and in-depth characterization was carried out on three of them, designated as clones 17, 24, and 42 (Figure 1). The expression of α7-5HT3 receptor mRNA was not statistically different between clones 17, 24, and 42 (Figure 1A).

**Figure 1.**
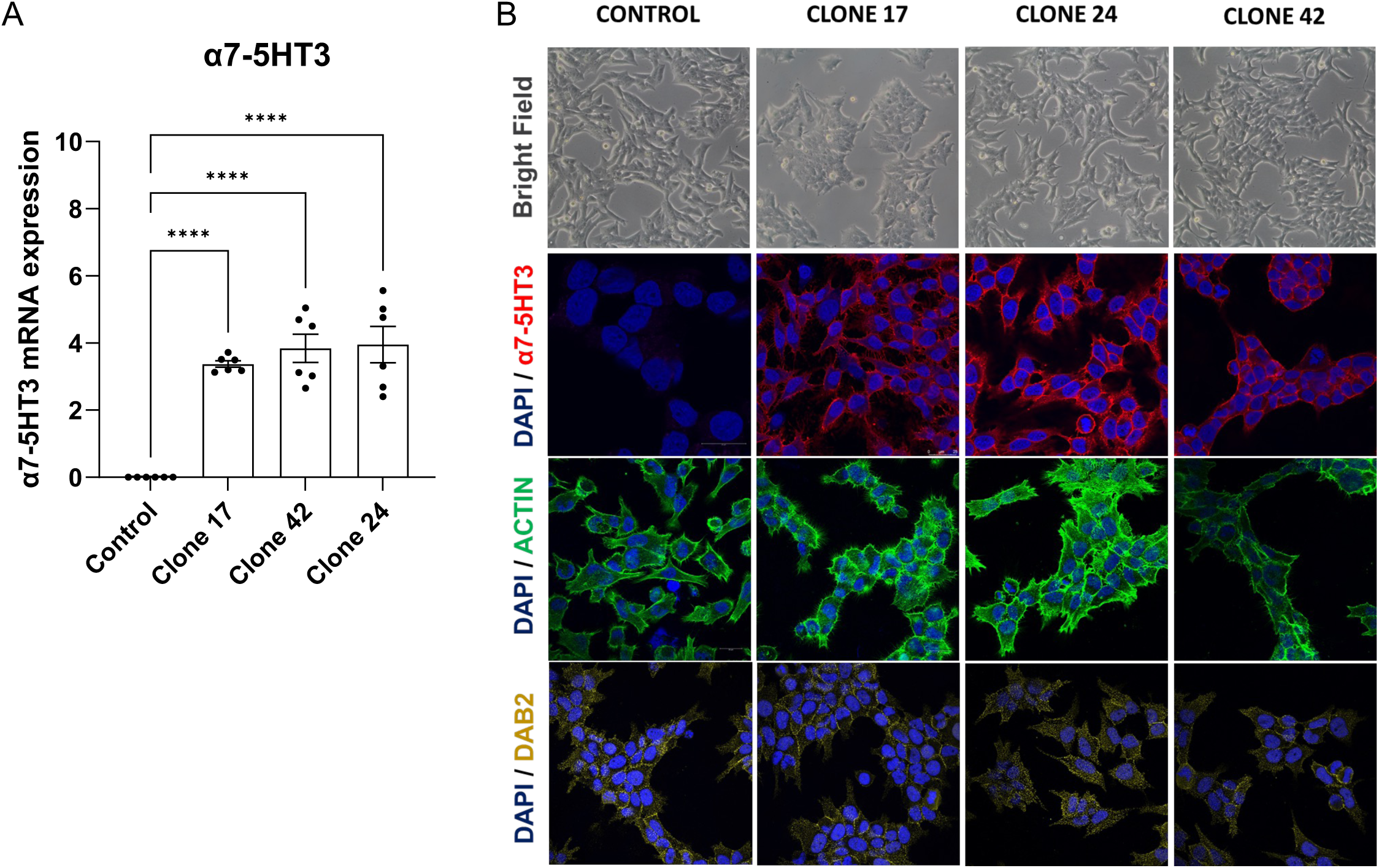
Morphological characterization of α7-5HT3 expressing cells. The expression of α7-5HT3 was investigated in control cells, expressing an empty vector, and in three selected clones, 17, 24 and 42. **(A)** mRNA expression of α7-5HT3 receptor was investigated by RT-qPCR. **(B)** Morphological characteristics of control and α7-5HT3 expressing cells were evaluated by immunofluorescence. Bright field images revealed no structural difference between cells. Staining with α-bungarotoxin, a peptide presenting high affinity for the α subunit of the nicotinic acetylcholine receptor (nAChR), showed expression of α7-5HT3 receptor at the plasmatic membrane only of cells expressing it. Phalloidin and Dab2 stainings were similar between control cells and cells expressing the α7-5HT3 receptor. DAPI nuclear staining is shown in blue, α-bungarotoxin (α7-5HT3) in red, Phalloidin (actin) in green and dab2 (yellow). ****, p<0,0001, Bar represents 20μm

Brightfield images revealed no discernible variations in cell morphology between control cells and the three selected clones (Figure 1B). Examination of the cellular localization of the α7-5HT3 receptor using α-bungarotoxin, a peptide binding to the α subunit of the nicotinic acetylcholine receptor, showed its presence on the cell surface of clones 17, 24, and 42 but not in control cells (Figure 1B). To ensure that the expression of the α7-5HT3 receptor did not induce structural changes, we studied the expression of markers of cytoskeletal organisation and of ZG cells. Phalloidin staining revealed similar actin organization in cells expressing an empty vector (control cells) or expressing the α7-5HT3 receptor (Figure 1B). A similar staining was observed for Disabled-2 (Dab2) (Figure 1B), a protein marker of ZG cells expressed at the cell membrane.

Varenicline is a nicotinic receptor partial agonist and a cholinergic agonist, which may non-specific effects in our cell model. To avoid this, we compared the effect of varenicline to those of a drug, uPSEM-817, specifically designed to bind to the α7-5HT3 receptor. Similar responses were obtained in response to PSEM and varenicline treatment in intracellular Ca^2+^ responses, both as traces (Figure S2A and C) and maximum of activation (Figure S2B and D).

### α7-5HT3 receptor activation by PSEM-817 induces cell membrane depolarization and increases intracellular Ca^2+^ content

Perforated patch-clamp recordings were conducted to assess the electrophysiological properties of two of the three clones expressing the α7-5HT3 receptor and selected for further investigations (clones 17 and 42) in comparison to control cells. Control cells expressing an empty vector, as well as clones 17 and 42, displayed similar membrane hyperpolarization with membrane potentials of -60.83±1.2, -61.73±1.16, and -63.27±1.26, respectively (Figure 2A), indicating that the α7-5HT3 receptor was not leaking ions.

**Figure 2.**
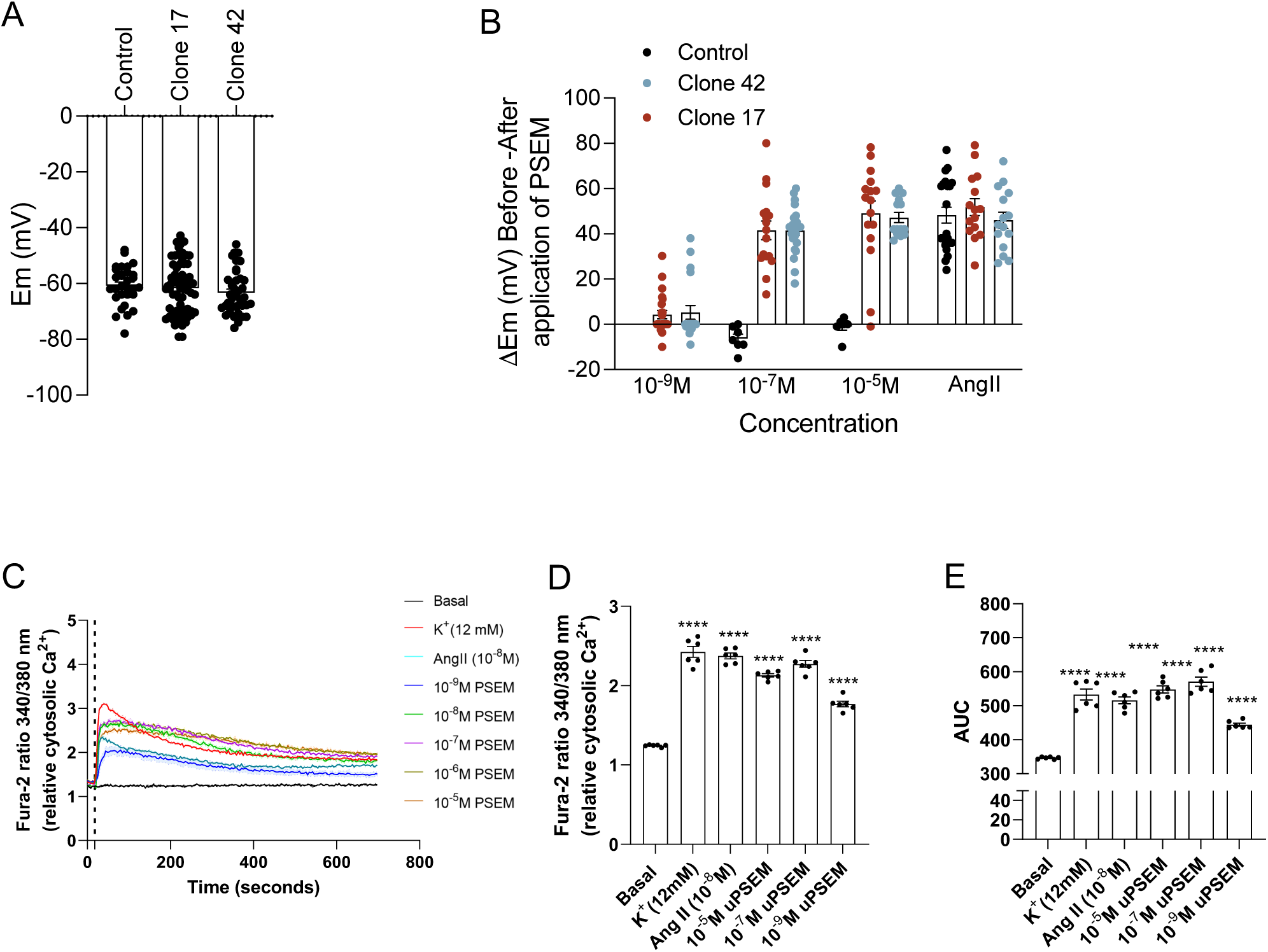
Functional characterization of α7-5HT3 expressing cells. **(A and B)** Perforated patch clamp experiments were performed to determine the membrane potential of cells in basal condition **(A)** or after stimulation with different concentrations of uPSEM-817 (10^-9^, 10^-7^ and 10^-5^ M) and AngII (10^-8^ M) **(B)**. **(C)** Calcium entry into the cells was evaluated using Fura-2 AM assay. Representative traces of intracellular Ca^2+^ responses to 10^-8^ M AngII, 12 mM K^+^ and 10^-9^ to 10^-5^ M uPSEM-817. **(D)** Determination of the maximum Fura-2 ratio 340/380nm in response to 10^-8^ M AngII, 12 mM K^+^ and 10^-9^, 10^-7^ and 10^-5^ M uPSEM-817. **(E)** Area under the curve (AUC) was determined to assess whether cell depolarization led to a calcium entry in the intracellular compartment. n=6, ****, p<0,0001

All the clones responded to induction by AngII, exhibiting membrane depolarization upon the application of 10^-8^M AngII (ýEm = 48.26±3.52, 51.81±3.73, 46.00±3,51 for control cells, clones 17 and 42 respectively) (Figure 2B). However, only clones 17 and 42 were responsive to 10^-7^ (ýEm = 41.58±4.12 and 41.49±2.13 for clones 17 and 42 respectively) and 10^-5^M of uPSEM-817 (ýEm = 49.03±5.51 and 47.19±2.24 for clones 17 and 42 respectively). The lower concentration of 10^-9^ M uPSEM-817 did not consistently induce membrane depolarization (Figure 2B).

To determine whether the membrane depolarization triggered by uPSEM-817 correlated with an increase in intracellular Ca^2+^ concentration, α7-5HT3 expressing cells were exposed to concentrations of uPSEM-817 ranging from 10^-9^ to 10^-5^M, as well as 12 mM K^+^ and 10^-8^ M AngII as positive controls (Figure 2C-E). The results were presented for clone 17 and similar results were obtained for clone 42 (Figure S3). Application of AngII or K^+^ led to a rapid increase in intracellular Ca^2+^ levels followed by a rapid decline without returning to baseline levels (Figure 2C), occurring after similar response latency (data not shown). The maximum of activation (Figure 2D) and the area under the curve (AUC) (Figure 2E) were significantly increased when cells were treated with AngII or K^+^. Treatment with uPSEM-817 ranging from 10^-9^ to 10^-5^ M also resulted in a similar rapid increase in intracellular Ca^2+^ content; this increase was more modest at 10^-9^ M uPSEM-817 compared to higher concentrations of the compound. Interestingly, the decline following the peak was slower with uPSEM-817 compared to AngII or K^+^ (Figure 2C). Maximum activation (Figure 2D) and AUC (Figure 2E) were lower in response to 10^-9^M of uPSEM-817 than with 10^-7^ and 10^-5^M uPSEM-817. Despite variations in Ca^2+^ content, the AUC and maximum activation were comparable when cells were treated with AngII, K^+^, 10^-7^ and 10^-5^M uPSEM-817.

### α7-5HT3 receptor activation by uPSEM-817 activates steroidogenesis

The steroidome of three cell clones expressing the α7-5HT3 receptor, clone 17, 24 and 42 was determined using the cell supernatant after 8h (Figure 3A and B) or 24h (Figure 3C and D) of treatment with 10^-8^M AngII or 12mM K^+^ as positive controls (Figure 3A and C), and uPSEM-817 ranging from 10^-9^ to 10^-5^M (Figure 3B and D). Interestingly, despite similar modification in intracellular Ca^2+^ concentration, different profiles were obtained for the three different clones in response to AngII, K^+^ and uPSEM-817. However, they converge all towards a stimulation of steroid biosynthesis in response to uPSEM-817. Among the 19 steroids measured, only 14 were detectable after 8h and 24h of treatment. After 8h of treatment, an overall activation of steroid biosynthesis was observed in response to AngII and K^+^ (Figure 3A and Table 1). Similarly, uPSEM-817 at concentrations from 10^-7^ to 10^-5^M led to an increase in steroid biosynthesis, with no significant changes observed at 10^-9^ and 10^-8^ M of uPSEM-817 (Figure 3B and Table 1). Interestingly, the increases in certain steroids due to uPSEM-817 were less pronounced compared to those induced by AngII and K^+^. After 24h of treatment, a significant decrease in the concentration of two steroid precursors, pregnenolone and progesterone, was observed in response to AngII and K^+^, while DOC, 17-hydroxyprogesterone, 17-hydroxypregnenolone, pregnenolone and progesterone concentrations remained elevated in response to uPSEM-817 (Table 1). Aldosterone biosynthesis was highly stimulated by AngII and K^+^ and to a lesser extent by uPSEM-817. While no effect was observed after 8 hours of treatment, after 24 hours the concentration of 18-oxocortisol was significantly increased in response to K^+^, and a trend was also observed in response to AngII and for all the concentrations of uPSEM-817. The concentrations of 18-hydroxycortisol were undetectable in all the conditions tested.

**Figure 3.**
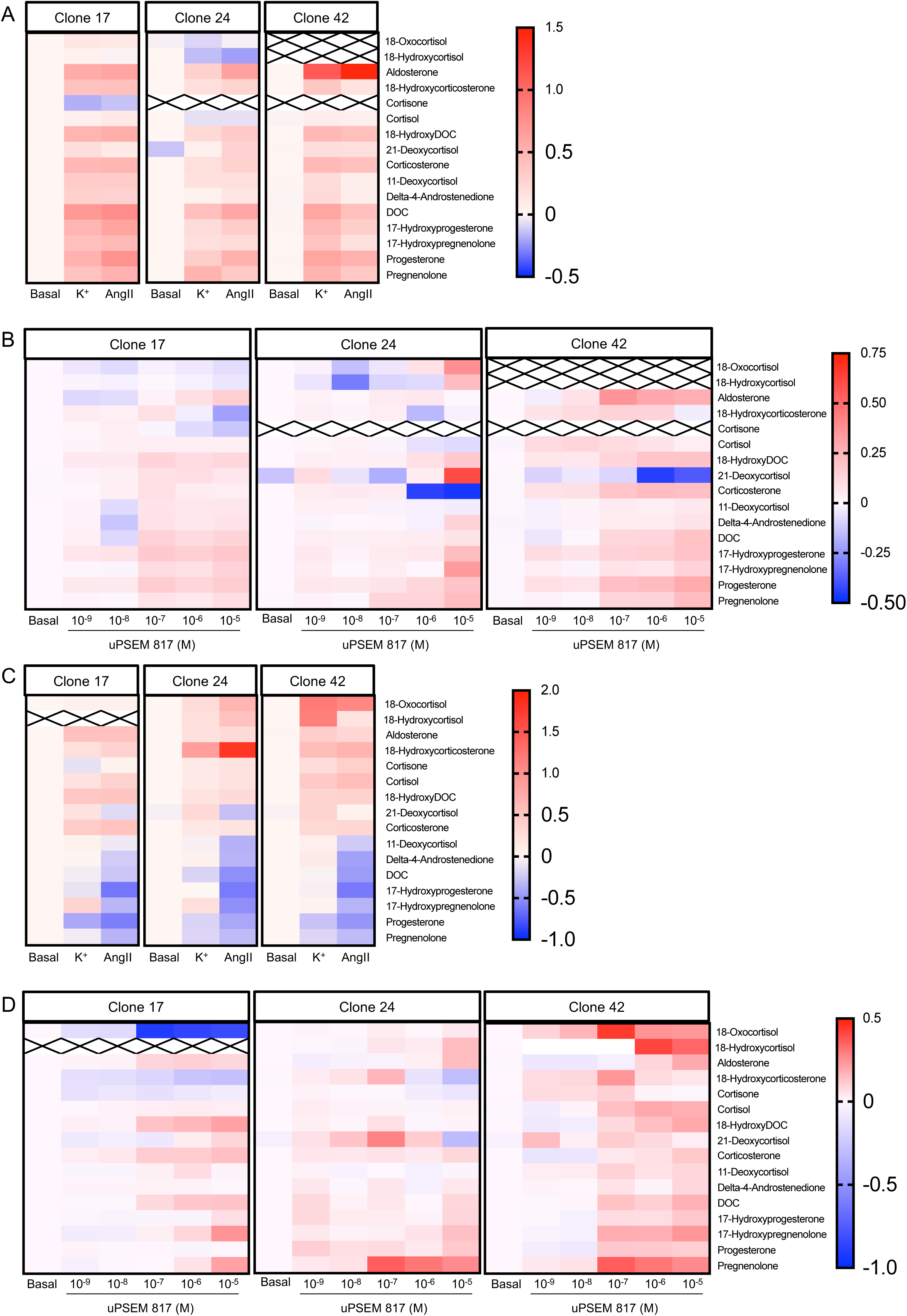
Steroid profiles of α7-5HT3-expressing cells in response to uPSEM-817, AngII and K^+^. **(A-D)** The steroid profiles were performed in three different clones of α7-5HT3-expressing cells, clone 17, clone 24 and clone 42. Hierarchical clustering of steroid fold induction compared to untreated cells (Basal) after 8h **(A-B)** or 24 hours **(C-D)** of treatment with 10^-8^M AngII **(A, C)**, 12mM K^+^ **(A, C)** or uPSEM-817 from 10^-9^ to 10^-5^ M **(B, D)**. n=6-9

**Table 1.**
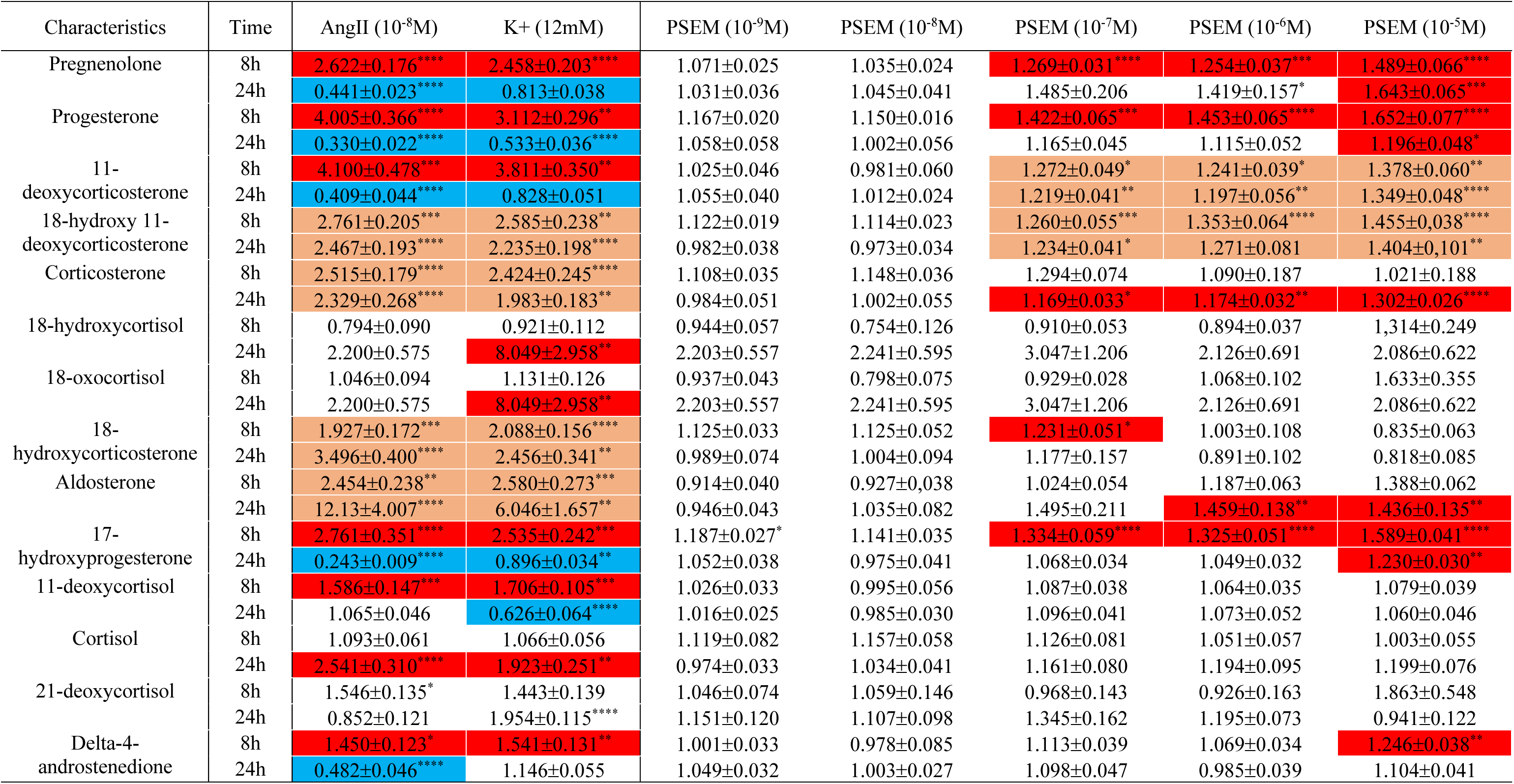
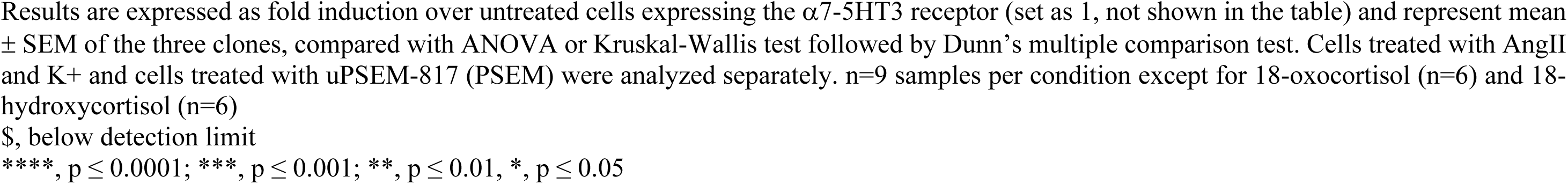
Steroid profiles in H295R_S2 cells expressing the α7-5HT3 receptor in response to AngII (10^-8^M), K^+^ (12mM) and uPSEM-817 (10^-9^ to 10^-5^M).

### T-type and L-type channels are both involved in calcium entry into the cells in response to uPSEM-817

Both T-type and L-type voltage-gated calcium channels have been shown to be involved in aldosterone biosynthesis regulation by modulating intracellular Ca^2+^ concentrations^29,30^. We evaluated the effect of a 2h pre-treatment with 10^-6^ M of the L-type calcium channel blocker nifedipine, or 10^-6^ M of the T-type calcium channel blocker mibefradil on the modulation of intracellular Ca^2+^ content in response to 10^-8^ M AngII, 12 mM K^+^ and increasing concentrations of uPSEM-817 (Figure 4A-C). Interestingly, pre-treatment of the cells with nifedipine or mibefradil partially abolished Ca^2+^ entry into the cells after treatment with K^+^ and uPSEM-817, as revealed by lower peak (Figure 4A-B) and decreased AUC (representing the modifications of intracellular Ca^2+^ concentrations and thus illustrating activation of Ca^2+^ signaling) (Figure 4C). In contrast, nifedipine and mibefradil had no effect on the initial entry of Ca^2+^ into the cell in response to AngII (Figure 4B), and only nifedipine pre-treatment led to a decrease in the AUC (Figure 4A-C). These results reveal that while only L-type Ca^2+^ channels are involved in Ca^2+^ entry into the cells in response to AngII, both T- and L-type Ca^2+^ channel are mobilized to allow Ca^2+^ entry into the cells in response to K^+^ and uPSEM-817.

**Figure 4.**
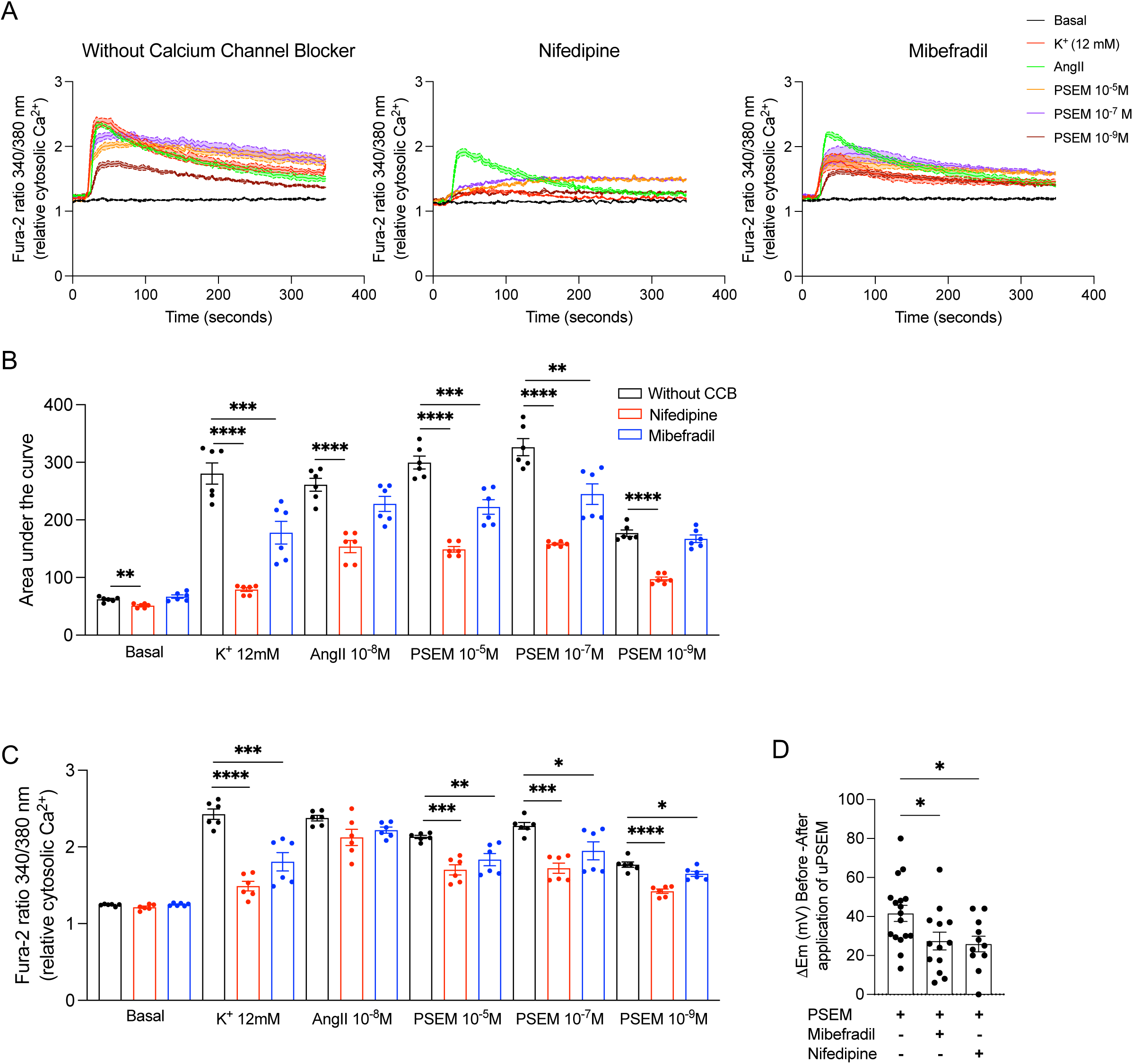
Inhibition of calcium entry using Nifedipine or Mibefradil. **(A)** Representative traces of intracellular Ca^2+^ responses to 10^-8^ M AngII, 12 mM K^+^ and 10^-9^, 10^-7^ and 10^-5^ M uPSEM-817 after pre-treatment with Nifedipine (left), or Mibefradil (right). (B) Determination of the maximum Fura-2 ratio 340/380nm in response to uPSEM-817, AngII and K^+^ after pre-treatment with nifedipine (red) or mibefradil (blue) or without pre-treatment (black). **(C)** Illustration of the [Ca^2+^]_i_ signaling, illustrated by the determination of the AUC assessed for 360 sec, in response to uPSEM-817, AngII and K^+^ after pre-treatment with nifedipine (red) or mibefradil (blue) or without pre-treatment (black). **(D)** Path clamp recordings of cells treated with uPSEM-817 and/or nifedipine or mibefradil. n=6, *, p<0.05; **, p<0.01; ***, p<0.001; ****, p<0.0001

To determine if the differences observed in the modulation of intracellular Ca^2+^ concentration in response to calcium channel blockers and uPSEM-817 cotreatment were due to differences in cell membrane depolarization, we performed patch-clamp analyses (Figure 4D). Prior to the administration of 10^-7^M uPSEM-817, we treated each patched cell with 10^-6^ M mibefradil, or with 10^-6^ M Nifedipine. These blockers were applied locally using an ejector solution. The results displayed a range of responses among cells: in certain instances, the blockers effectively suppressed depolarization, returning the membrane potential to a state near its baseline (ΔEM≈0). However, in other cases, the membrane remained depolarized, albeit to a lesser degree than when exposed exclusively to 10^-7^ M uPSEM-817 (Figure 4D).

### α7-5HT3 receptor activation by uPSEM-817 led to a decrease in cell proliferation and an increase of apoptosis

To evaluate the impact of elevated intracellular Ca^2+^ concentration on cell proliferation, cells were treated with 12 mM K^+^ or 10^-9^ to 10^-5^ M uPSEM-817 for 24h and 72h. The number of viable proliferating cells was determined using a colorimetric method. Whereas following 24 hours of treatment, cell proliferation remained unaffected by K^+^ or uPSEM-817 treatment, a decrease in cell proliferation was observed after 72 hours of treatment with the higher concentration of uPSEM-817 (Figure 5A).

**Figure 5.**
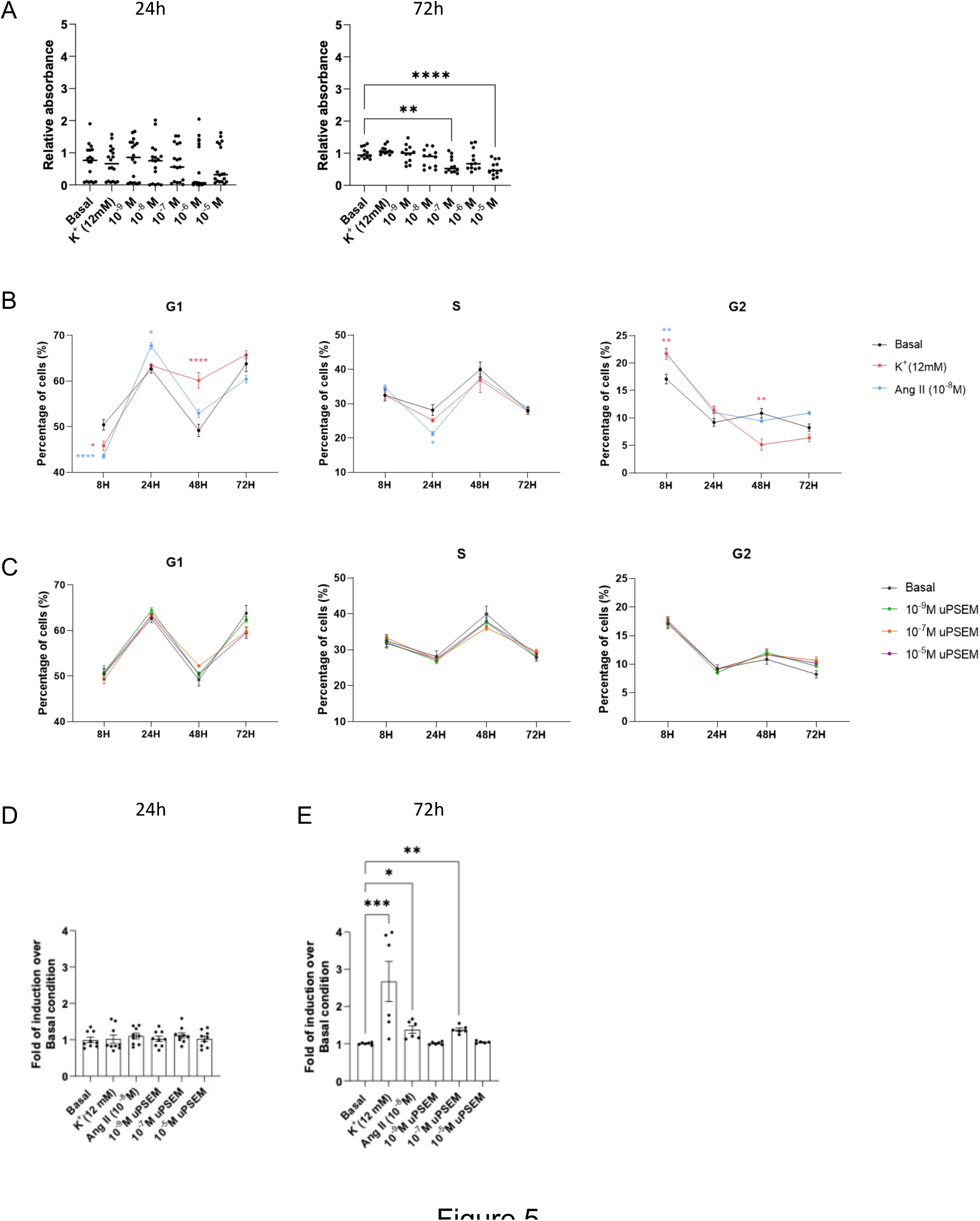
Effect of uPSEM-817 on cell proliferation and apoptosis. (**A**) Cell viability was measured on cells treated for 24 or 72 hours with 12 mM K+ or uPSEM-817 by MTS assay using the CellTiter 96 Aqueous One Solution Cell Proliferation Assay. (**B**) Cell cycle distribution measured by FACS using propidium iodide in response to 10^-8^M AngII and 12mM K^+^ (upper panel) and 10^-9^, 10^-7^ and 10^-5^M uPSEM-817 (lower panel). (C) Apoptosis determined by the proportion of cells in sub-G1 phase. n=6 (cell cycle) – 12 (proliferation), *, p<0.05; **, p<0.01; ****, p<0.0001

To explore the impact on cell cycle progression of AngII, K^+^ and uPSEM-817 treatment, we assessed the cell cycle phase distribution using propidium iodide staining and cytometric analysis at 8h, 24h, 48h, and 72h. Treatment with AngII for 8h resulted in a reduction in the percentage of cells in the G1 phase and an increase in the percentage of cells in G2 compared to untreated cells suggesting an increase in cell proliferation (Figure 5B). After 24h of AngII treatment, an increase in the percentage of cells in the G1 phase and a decrease in the S phase was also observed. Longer exposure to AngII had no effect on the distribution of cells across different phases of the cell cycle. In response to K^+^ treatment, a similar pattern was observed with a decrease in the percentage of cells in G1, and an increase in G2 after 8h. While no alterations in cell distribution were noted after 24h of K+ treatment, an increase in the percentage of cells in G1 and a decrease in G2 was observed (Figure 5B). Conversely, treatment with uPSEM-817 did not affect cell distribution among the different phases compared to untreated cells (Figure 5C). We assessed the effect of uPSEM-817 on apoptosis by determining the proportion of cells in the sub-G1 phase. The sub-G1 phase reflects DNA fragmentation which occurs in the late stage of apoptosis. Similar to what was observed for cell proliferation results, no effect was observed when cells were treated with uPSEM-817, AngII and K^+^ for 24h (Figure 5D). At 72h, a significant increase in the percentage of cells in sub-G1 phase was observed in response to K^+^, AngII and uPSEM-817 (10^-7^M) (Figure 5E), indicating an increase in cell apoptosis.

### α7-5HT3 receptor activation by uPSEM-817 results in the activation of specific pathways

To gain insight into the impact of modulating intracellular Na^+^ concentration in cells expressing the α7-5HT3, we conducted RNA sequencing analysis on these cells (clones 17, 24, and 42) following treatment with 10^-7^M uPSEM-817 for 8h (Figure S4 and S6) and 24h (Figure 6 and S7). Hierarchical clustering effectively segregated cells treated for 8h and 24h from untreated cells (Figure S4A, S6A and S7A and Figure 6A). After 8h of treatment, we identified 28 differentially expressed genes with a fold change of at least 0.5; 18 upregulated genes (64.29%) and 10 downregulated genes (35.71%) (Table S2). Gene ontology analyses unveiled 18 specific enriched biological processes in uPSEM-817 treated cells (Figure S4B and Table 2), primarily related to calcium ion transport and signalling pathways, cell adhesion, and aldosterone synthesis and secretion. Following 24h of treatment, 30 genes were differentially expressed with a fold change of at least 0.5, with 22 genes (73.33%) upregulated and 8 (26.67%) downregulated (Figure 6B and Table S3). Enriched pathways included protein regulation of GTPase activity, Ras protein signal transduction, and aldosterone synthesis and regulation (Figure 6B and Table 3). These analyses were completed by using Gene Set Enrichment Analysis (GSEA) and Hallmark database that provides information on most universal cellular mechanisms (Figures S5 and S6). Using a FDR <25% we identified 6 differentially regulated pathways after 8h of treatment with PSEM and 7 after 24h. After 8h of treatment, TNFα signaling via NFκB and coagulation pathway were found enriched in basal condition (Figure S5B) whereas Kras signaling, heme metabolism, UV response and Interferon alpha response were found to be enriched in response to uPSEM-817 (Figure S5C). After 24h of treatment (Figure S6), only TNFα signaling via NFκB pathway (Figure S6B) was enriched in basal condition and Myc targets, oxidative phosphorylation, DNA repair Apical surface, unfolded protein response and G2M checkpoint pathways were enriched after uPSEM-817 treatment.

**Figure 6.**
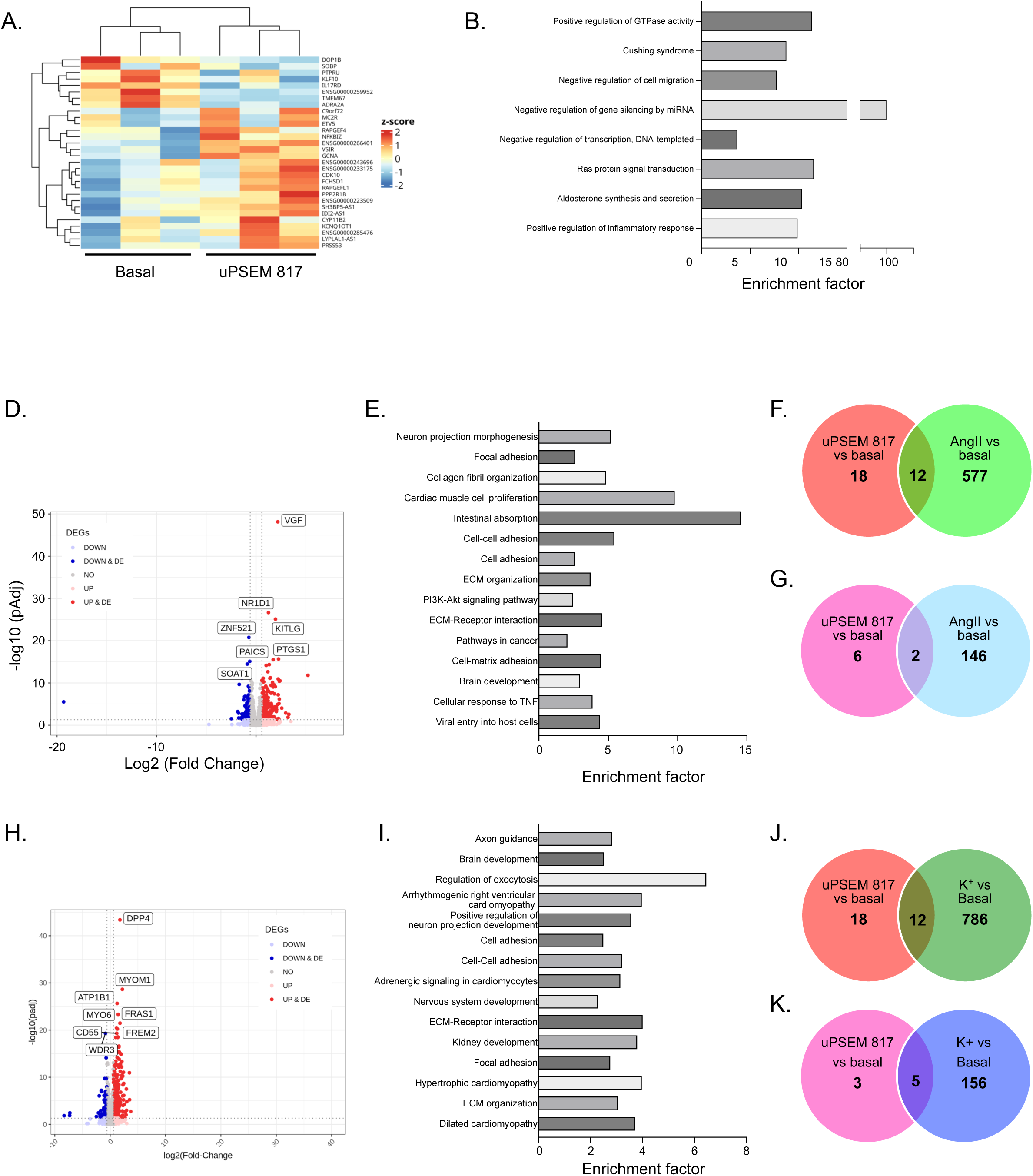
Gene expression profiles of α7-5HT3-expressing H295R-S2 cells treated 24 hours with 10^-7^M uPSEM-817, 10^-8^M AngII or 12mM K^+^. (**A) Hierarchical clustering of** samples using the 28 differentially expressed genes in cells treated or not with 10^-7^M uPSEM-817. (**B**) Biological process enrichments determined using the list of differentially expressed genes in cells treated or not with 10^-7^M uPSEM-817. (**C**) Volcano plot showing the 4932 differentially in response to 10^-8^M AngII. Differential expressed genes are highlighted as blue (down-regulated) or red (up-regulated) dots. (**D**) Biological process enrichments determined using the list of differentially expressed genes in cells treated or not with 10^-8^M AngII. (**E**) Venn diagram representing the common and different genes differentially expressed in response to 10^-8^M AngII and 10^-7^M uPSEM-817. (F) Venn diagram representing the common and different enriched biological process in response to 10^-8^M AngII and 10^-7^M uPSEM-817. (**G**) Volcano plos showing the 728 differentially in response to 12mM K^+^. The x-axis is the Log2 fold change between the two conditions; the adjusted p value based on –log_10_ is reported on the y-axis. Genes significantly different are highlighted as blue (down-regulated in cells treated with 12mM K^+^) or red (up-regulated in cells treated with 12mM K^+^) dots. (**H**) Biological process enrichments determined using the list of differentially expressed genes in cells treated or not with 12mM K^+^. (**I**) Venn diagram representing the common and different genes differentially expressed in response to 12mM K^+^ and 10^-7^M uPSEM-817. (J) Venn diagram representing the common and different enriched biological process in response to 12mM K^+^ and 10^-7^M uPSEM-817.

**Table 2.** List of significantly enriched pathways in H295R_S2 cells expressing the α7-5HT3 receptor in response to 8h treatment with10^-7^M uPSEM-817.

| **Pathway** | **Fold Enrichment** | **pValue** |
| --- | --- | --- |
| Calcium ion transport | 20.29 | 9.39e-4 |
| Focal adhesion | 7.72 | 0.00317 |
| cAMP signaling pathway | 6.97 | 0.00459 |
| Calcium signaling pathway | 6.20 | 0.00694 |
| Cellular response to amino acid stimulus | 22.96 | 0.00723 |
| Cell-cell adhesion | 9.26 | 0.00859 |
| Human papillomavirus infection | 4.74 | 0.01738 |
| Fatty acid metabolic process | 13.96 | 0.01868 |
| PI3K-Akt signaling pathway | 4.43 | 0.02169 |
| Biosynthesis of amino acids | 12.54 | 0.02194 |
| Calcium ion transmembrane transport | 12.09 | 0.02444 |
| Myofibril assembly | 64.50 | 0.02988 |
| Transmembrane receptor protein tyrosine kinase signaling pathway | 10.03 | 0.03448 |
| Prostate cancer | 9.70 | 0.03534 |
| Aldosterone synthesis and secretion | 9.60 | 0.03600 |
| Canonical glycolysis | 50.17 | 0.03825 |
| HIF-1 signaling pathway | 8.63 | 0.04368 |
| Regulation peptidyl-tyrosine phosphorylation | 39.26 | 0.04862 |

**Table 3.** List of significantly enriched pathways in H295R_S2 cells expressing the α7-5HT3 receptor in response to 24h treatment with10^-7^M uPSEM-817.

| **Pathway** | **Fold Enrichment** | **pValue** |
| --- | --- | --- |
| Positive regulation of GTPase activity | 11.41 | 1.63e-4 |
| Cushing syndrome | 8.74 | 0.00919 |
| Negative regulation of cell migration | 7.77 | 0.01407 |
| Negative regulation of gene silencing by miRNA | 89.88 | 0.02164 |
| Negative regulation of transcription, DNA-templated | 3.67 | 0.02178 |
| Ras protein signal transduction | 11.60 | 0.02671 |
| Aldosterone synthesis and secretion | 10.37 | 0.03103 |
| Positive regulation of inflammatory response | 9.90 | 0.03575 |

To identify genes and pathways specifically regulated by modulation of intracellular Na^+^ concentration, we also performed RNA sequencing on α7-5HT3 receptor expressing cells treated with 12mM K^+^ and 10^-8^M AngII for 8h and 24h (Figure S4 and Figure 6). After 8h of treatment, 4932 genes were differentially expressed in response to AngII and 728 in response to K^+^ (Figure S4C and S4G and Table S4 and S5). Gene ontology analyses for 8h AngII treatment revealed enrichment in 390 different pathways (Table S6), with significant involvement in the regulation of reticulum activity, negative regulation of cell growth, and aldosterone, cortisol and parathyroid hormone synthesis and secretion (Figure 6D). Eight hours of treatment with K^+^ (Figure S4H) resulted in enrichment of 211 pathways (Table S7), including membrane depolarization during action potential, calcium and sodium ion transport, and circadian rhythm (Figure S4H). Interestingly, both AngII and K^+^ treatment led to enrichment in positive regulation of transcription and MAPK signaling pathways. Among the 28 differentially expressed genes in response to uPSEM-817, 17 were common to AngII (*OLFML2B*, *RGPD6*, *LMOD1*, *MAT1A*, *MC2R*, *CA2*, *CACNA1C*, *EGFR*, *ETV5*, *TNXB*, *PFKP*, antisense to *WASF3*, antisense to *ERVW-1* and *PEX1*, *PPP2R1B*, *LYPLAL1-AS1*, *LONRF2* and *ST6GALNAC6*) (Figure S4E) and 12 to K^+^ (*OLFML2B*, *LMOD1*, *MAT1A*, *MC2R*, *CA2*, *CACNA1C*, *ATP8B4*, *ETV5 PFKP*, *PPP2R1B*, *LYPLAL1-AS1* and *LONRF2* (Figure S4I).

Among the 18 enriched pathways in response to uPSEM-817, 9 were commonly enriched in response to AngII (Figure S4F) and 5 in response to K^+^ (Figure S4J). Most importantly, 11 genes were specifically regulated by uPSEM-817 when compared with AngII (*HPX*, *EPHA10*, *ATP8B4*, *HHIP*, *MICAL1*, *RBMS2*, *E9PAM4*, *H7C2Y5*, *TP53TG3B*, *DUXAP8* and *RNVU1-7*) and 16 when compared to K^+^ (*HPX*, antisense to *WASF3*, *EPHA10*, *HHIP*, *ST6GALNAC6*, antisense to *ERVW-1* and *PEX1*, *EGFR*, *MICAL1*, *TNXB*, *RGPD6*, *RBMS2*, *E9PAM4*, *H7C2Y5*, *TP53TG3B*, *DUXAP8* and *RNVU1-7*). This allowed us to define a specific Na^+^-induced specific signature composed of 10 genes (*HPX*, *EPHA10*, *HHIP*, *MICAL1*, *RBMS2*, *E9PAM4*, *H7C2Y5*, *TP53TG3B, DUXAP8* and *RNVU1-7*) (Figure S7).

After 24h of treatment, 589 genes were statistically differentially expressed in response to AngII (Figure 6D, 6F and Table S8). Among them, 12 were common to uPSEM-817 (*SOBP*, *ADRA2A*, *ETV5*, *antisense to DLGAP1*, *VSIR*, *GCNA*, *FCHSD1*, *RAPGEFL1*, *PPP2R1B*, *CYP11B2*, *LYPLAL1-AS1* and *PRSS53*); 146 pathways were specifically enriched in response to AngII and 2 (Cushing syndrome and Aldosterone synthesis and secretion) were commonly enriched in response to AngII and uPSEM-817 (Figure 6G and Table S9). Interestingly, the most significant enriched pathways were involved in cell adhesion and extracellular matrix organization (Figure 6E). K^+^ treatment for 24h led to the regulation of 798 genes (Figure 6H, 6J and Table S10), among them 12 were common to uPSEM-817 (*KLF10*, *ADRA2A*, *C9ORF72*, *MC2R*, *ETV5*, *RAPGEF4*, *VSIR*, *GCNA*, *FCHSD1*, *PPP2R1B*, *CYP11B2* and *LYPLAL1-AS1*) (Figure 6J). Similarly, enriched pathways included those associated with cell adhesion and extracellular matrix organization (Figure 6I and Table S11). Five out of the eight enriched pathways identified in response to uPSEM-817 treatment were also commonly enriched in response to K^+^ (Figure 6K). These common pathways involved positive regulation of GTPase activity, Cushing Syndrome, Negative regulation of cell migration, Ras protein signal transduction, and Aldosterone synthesis and secretion. Notably, Cushing syndrome and aldosterone biosynthesis and secretion pathways were also commonly enriched between uPSEM-817 and AngII. We were able to define a 24h sodium-specific signature composed by 14 genes (*FMNL1-AS1*, *IDI2-AS1*, *WHAMMP1*, *MUSTN1-ITIH4 Readthrough*, *NFKBIZ*, *KCNQ1OT1*, *SH3BP5-AS1*, CDK10, *PTPRU*, *IL17RD*, *TMEM67*, *MAZ-AS*, *DOP1B* and one unknown sequence) (Figure S8).

The list of differentially expressed genes was compared to their expression in 11 control adrenal and 123 APA retrieved from transcriptomic data^31^ (Table S12 and S13). Of these 123 APA, 50 carried a mutation in the *KCNJ5* gene and 73 in another APA driver gene; the comparison was therefore made between the 11 control adrenals, the 50 APA with a mutation in the *KCNJ5* gene and the 73 APA without a mutation in the *KCNJ5* gene. After 8h of treatment with 10^-7^M of uPSEM-817, among the 55 identified genes that were significatively differentially expressed without consideration of the fold change, the expression of 49 were found in our transcriptomic data from control adrenals and APA and could be retrieved. In these data, no expression was detected for five genes, and out of the remaining 44 genes, 13 were also found to be differentially expressed in APA compared to control adrenal (Table S12), two only in APA with a KCNJ5 mutation (*HPX* and *WHRN*) and three in APA without KCNJ5 mutation (ETV5, *GPAM* and *OLFML2B*). Interestingly, among the 13 genes differentially expressed in APA independently of the mutational status, 4 were found to be up-regulated (*CACNA1C*, *ATP2B2*, *VDR* and *EPHA10*) and 1 downregulated (*RGPD6)* in both APA and α7-5HT3 cells, whereas 6 genes were found to be upregulated in α7-5HT3 cells but downregulated in APA (*PPP2R1B*, *LONRF2*, *ATP8B4*, *TNXB*, *ITGA7* and *HSPA12A*), independently of the mutational status and 2 down-regulated in response to uPSEM-817 but up-regulated in all APA (*TMEM200A* and *ELL2*). Similarly, after 24h of treatment, among the 73 genes identified genes that were significantly differentially expressed regardless of the fold change, the expression of 64 genes could be retrieved from our transcriptomic data. No expression was found for 10 genes, and among the remaining 54 genes, 15 were also found to be differentially expressed in APA compared to control adrenal (Table S13). Among these 15 genes, three were found to be commonly upregulated in both α7-5HT3 cells and APA (*FCHSD1*, *CACNA1C* and *TTN1*) and one downregulated (*KLF10*) whereas two were found to upregulated in α7-5HT3 cells and downregulated in APA, independently of the mutational status (*PPP2R1B* and *NEAT1*) and four downregulated in α7-5HT3 cells and upregulated in all APA (*STMN3*, *FIBCD1*, *FSCN1* and *CCDC71L*). Interestingly, three genes were found to be significantly regulated only in APA without KCNJ5 mutations, two downregulated in α7-5HT3 cells but upregulated in APA (*ETV5* and *CEP170*) and one downregulated in both (*NR2F2*); two genes were found to be significantly regulated only in APA with KCNJ5 mutations, one downregulated in α7-5HT3 cells but upregulated in APA (*NFKBI2*) and one downregulated in both (*ABHD2*). Interestingly, none of the genes commonly regulated in α7-5HT3 cells and APA belongs to the Na^+^ signature defined after 8h and 24h of treatment suggesting that they are probably involved in the early phase of APA development.

## Discussion

Somatic *KCNJ5* mutations are the most frequent genetic abnormalities found in APA with a prevalence between 43% and 75% of cases. APA with KCNJ5 mutations are more frequent in women and at younger age^31,32^ and are characterized by hybrid steroid production and larger adenoma size^33^. Expression of mutated KCNJ5 in adrenocortical cells leads to increased aldosterone production without increasing cell proliferation^17,19^, raising the question as to whether those mutations are sufficient to lead to both increased aldosterone production and adenoma formation. Here we describe the development of an adrenocortical cell model that recapitulates the main features of KCNJ5 mutations by modulating “on demand” sodium entry into the cells, using chemogenetic tools. Increased sodium entry through the chimeric α7-5HT3 receptor upon stimulation with uPSEM-817 led to cell membrane depolarization, opening of voltage-gated Ca^2+^ channels and increased intracellular Ca^2+^ concentrations, resulting in the stimulation of *CYP11B2* expression and increased aldosterone biosynthesis.

The steroid response of H295R-S2 α7-5HT3 cells to AngII is similar to that described in H295R cells^34^, with an early and transitory increase of pregnenolone, progesterone and 11-deoxycorticosterone and a late and sustained production of aldosterone and corticosterone^34^. In contrast, PSEM induced sodium entry into the cells led to a specific steroid profile only partially overlapping with that induced by AngII and K^+^. In particular, we observed a prolonged induction of early steroid precursors and DOC, suggesting an action on both early and late steps of aldosterone biosynthesis, as well as a delayed and smaller induction of aldosterone biosynthesis; there was no induction of glucocorticoid biosynthesis H295R-S2 α7-5HT3 cells. An increase of 18-oxocortisol and 18-hydroxycortisol was also observed in response to uPSEM-817 in two of the three clones, suggesting production of hybrid steroids in response to increased intracellular Na^+^ concentration. This is consistent with results reported for the overexpression of KCNJ5 harboring the p.Tyr158Ala mutation in HAC cells resulting in a significant increase of both 18-hydroxycortisol and 18-oxocortisol^18,35^. However, despite similar depolarization of the cells in response to uPSEM-817 or AngII, our data suggest a delayed activation of mineralocorticoid biosynthesis in response to the modulation of intracellular Na^+^ concentration compared to AngII and K^+^ that could be due to the activation of specific signaling pathways. This hypothesis is supported by our finding of a specific gene expression signature associated with Na^+^ influx into the cells. Interestingly, gene expression analysis revealed expression of *KCNJ5* in H295R-S2 α7-5HT3 cells (Figure S9A), which remains unchanged after uPSEM-817 treatment. In this context, Na^+^ may act as an activator for the KCNJ5 channel^36^ by binding to the C-terminal part of the channel, near a region also sensitive to the PtdIns(4,5)P_2_, enhancing the sensitivity of the channels to PtdIns(4,5)P_2_^37^. The interaction of PtdIns(4,5)P_2_ with the C-terminal part of GIRK4 has been shown to regulate its opening by stabilizing the structure of the pore^37^. The resulting K^+^ extrusion may attenuate the activation of Ca^2+^ signaling and explain the delayed increased in aldosterone biosynthesis observed in response to uPSEM817.

If the role of *KCNJ5* mutations in inducing aldosterone biosynthesis has been clearly established, their role in promoting abnormal cell proliferation is still a matter of debate. In our model, we demonstrated that Na^+^ entry into the cells lead to decreased cell proliferation and increased apoptosis, confirming previous *in vitro* data. This is in accordance with previous studies showing that expression of *KCNJ5* mutants in adrenocortical cells resulted in decreased cell proliferation associated in some cases with increased apoptosis^18,20^. *Choi et al* postulated that the activation of calcium signaling induced by *KCNJ5* mutations is responsible for both^8^ increased aldosterone biosynthesis and cell proliferation, and indeed some studies also reported an association between *KCNJ5* mutations and larger adenoma size^31,32^ and a positive correlation between expression of Ki67, a marker of cell proliferation, and the diameter of APA harboring *KCNJ5* mutations^20^. Specific factors overexpressed in APA harboring a *KCNJ5* mutations^38,39^ have been suggested to possibly counteract the pro-apoptotic effect of these mutations^20^ suggesting, in the long term, the existence of compensatory mechanisms maintaining cell proliferation. In particular, expression of Teratocarcinoma-Derived Growth Factor-1 (*TDGF-1*) and Vinsinin-like 1 (*VSNL1*), two genes with anti-apoptotic properties, were found to be upregulated in *KCNJ5* mutated APA^40,41^. However, in our cell model the expression of *VSNL1* was not modified (Figure S9B) and *TDGF1* was not expressed at all. It is possible that increased expression the modulation of these genes is a later event secondary to the development of an APA to compensate for Na^+^-induced cell death. Moreover, signaling pathway analyses revealed downregulation of genes involved in the G2/M checkpoint pathway and in progression through the cell division cycle, probably contributing to the decrease of cell proliferation observed in our model. Alternatively, our data suggest that other mechanisms may be responsible for abnormal cell proliferation, leading to the development of APA. These include the presence of two hits, one responsible for cell proliferation and the other for autonomous aldosterone production or the development of APA from APCCs. Recently, we and others identified risk alleles associated to PA in genome wide association studies. Within the identified genetic loci, some genes appear to modulate adrenal cortex homeostasis and may affect cell proliferation eventually generating a propitious environment leading to the occurrence of somatic mutations^42,43^. The three models are not mutually exclusive and may be linked together by mechanisms that remain to be identified. Finally, in FH-III it cannot be ruled out that Na^+^ entry indued by *KCNJ5* mutations may have an impact on cell proliferation during adrenal development, which could explain the bilateral hyperplasia observed in severe cases^8,19^.

Gene expression profiles allowed us to identify a sodium-induced gene signature composed of 10 genes after 8h of treatment with uPSEM-817 (Figure S7) and 14 genes after 24h of treatment (Figure S8). Interestingly, among these lists of genes, some are involved in cell cycle regulation, proliferation and apoptosis. Hence, the expression of *RBMS2* and *PTPRU*, two genes involved in the control of cell cycle progression^44^ and cell proliferation^45,46^ respectively, was decreased after 8h or 24h while the expression of *CDK10*, a Cdc2-related kinase involved in the regulation of the G2/M phase of the cell cycle, was increased. However, the role of CDK10 in regulating cell proliferation is not clear, as some studies suggest a role of CDK10 in cell proliferation activation^47–49^, while others report a tumor suppressor role for CDK10 through inhibition of cell proliferation^50,51^. The identification of these genes in the Na^+^-signature supports our results showing an inhibitory effect of uPSEM-817 on cell proliferation. Among the genes belonging to the Na^+^ signature after 8h of treatment with uPSEM-817, *HPX* expression was found to be upregulated. *HPX* encodes hemopexin, a protein with high binding affinity for heme. Interestingly, CYP11B2 and other cytochromes P450 enzymes use heme as cofactor required for their activity. The availability of heme has been shown to affect aldosterone and corticosterone biosynthesis in rats^52^. Our transcriptomic data revealed a significant increase of *HPX* expression in APA harboring a *KCNJ5* mutation^31^ and a recent study reported also increased expression of *HPX* associated with CpG hypomethylation^53^ in APA compared to adjacent adrenal gland suggesting a role of *HPX* in APA development potentially through the regulation of aldosterone biosynthesis. Similarly, the expression of *MC2R*, the melanocortin 2 receptor, was found to be increased in APA associated with CpG hypomethylation^53^. Gene expression profiling of H295R-S2 α7-5HT3 cells treated with either AngII, K^+^ or uPSEM-817 revealed also a significant increase of *MC2R* expression; and a previous study reported higher expression of *MC2R* in APA compared to control adrenals^54^. Finally, a recent work of our laboratory shows that expression of MC2R in APA was more frequently found in regions expressing CYP11B2^55^, suggesting a potential role in regulating aldosterone biosynthesis. Moreover, the existence of a continuum of PA and dysregulated aldosterone production, prominently influenced by ACTH has recently been described^56^.

In conclusion, we demonstrate that H295R-S2 α7-5HT3 cells are a new cell model in which intracellular Ca^2+^ concentration can be modulated on “demand”, reproducing the major features of cells harbouring *KCNJ5* mutations. The stimulation of Na^+^ entry into the cells leads to cell membrane depolarization, calcium entry into the cells, activation of calcium signaling, increased expression of *CYP11B2* and stimulation of the mineralocorticoid biosynthesis pathway. This is associated with a decrease of cell proliferation and an increase of apoptosis, indicating that additional events may be required for the development of APA. RNA sequencing revealed that Na^+^ entry into the cells is responsible for a specific transcriptomic signature that may explain some of the features of APA carrying a *KCNJ5* mutations. This cellular model thus provides important new insight into the mechanisms leading to PA development, uncovering new mechanisms involved in the disease, thereby paving the way for new therapeutic approaches in primary aldosteronism.

## Acknowledgments

This work was funded through institutional support from INSERM, by the Agence Nationale pour la Recherche (ANR-18-CE93-0003-01), the Fondation pour la Recherche Médicale (EQU201903007864), the European Union’s Horizon 2020 research and innovation programme under the Marie Sklodowska-Curie grant agreement No. 954798 (MINDSHIFT ITN), for which M.-C.Z. is principal investigator, and by a grant from the Leducq Foundation (18CVD05).

