## Supplemental information for "Modulation of calcium signaling on demand to decipher the molecular mechanisms of primary aldosteronism"

**Running Title**: Molecular mechanisms responsible for primary aldosteronism

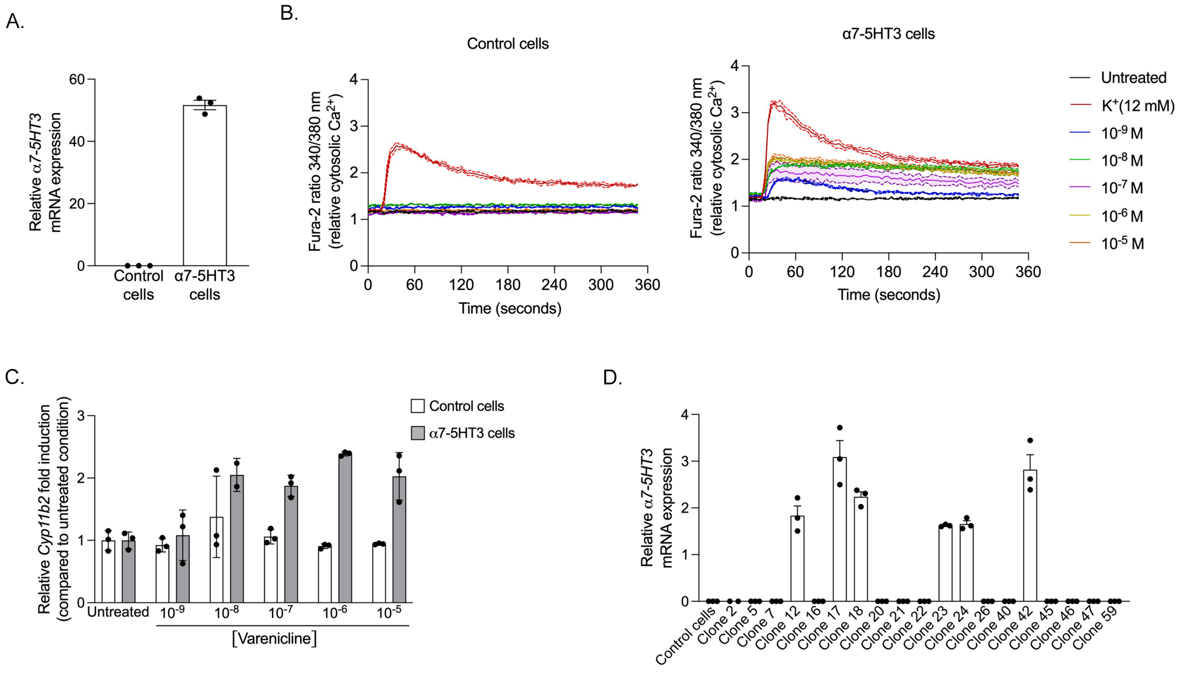


**Figure S1. Characterization of a cell model stably expressing the α7-5HT3 receptor in response to varenicline.** (**A**) mRNA expression, by RT-qPCR, of α7-5HT3 receptor was investigated in cells transfected with an empty vector (control cells) or a vector containing the α7-5HT3 receptor sequence (α7-5HT3 cells). (**B**) Calcium entry into the cells was evaluated using Fura-2 AM assay. Representative traces of intracellular $Ca^{2+}$ responses to 12 mM $K^+$ and $10^{-9}$ to $10^{-5}$ M Varenicline, a uPSEM-817 analog, of control cells and α7-5HT3 cells. (**C**) mRNA expression of Cyp11b2 in control and α7-5HT3 cells in response to $10^{-9}$ to $10^{-5}$ M Varenicline**.** (**D**) Characterization of monoclonal cell populations for the expression of α7-5HT3 mRNA by RT-qPCR.

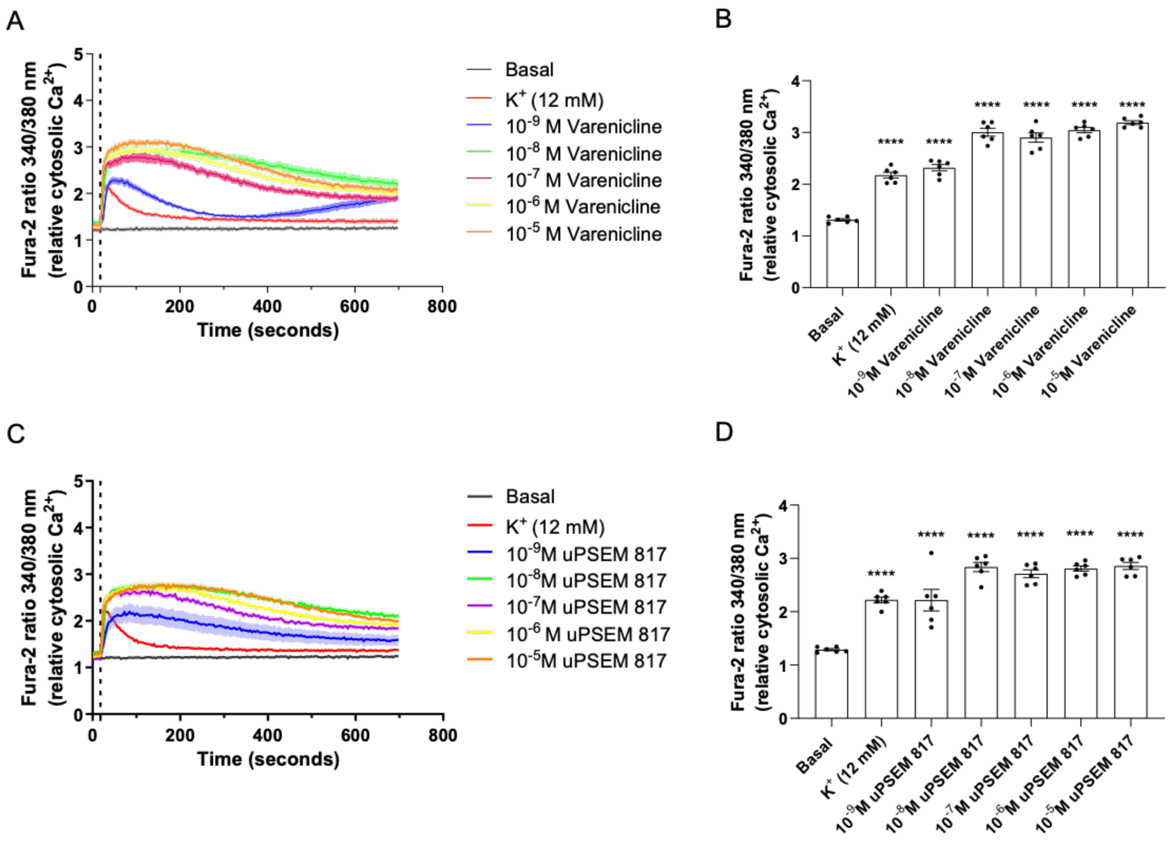


**Figure S2. Comparison of varenicline and uPSEM-817 effect on intracellular $Ca^{2+}$ entry in α7-5HT3 expressing cells.** (**A**) Calcium entry into the cells was evaluated using Fura-2 AM assay. Representative traces of intracellular $Ca^{2+}$ responses to $10^{-8}$ M AngII, 12 mM $K^{+}$ and $10^{-9}$ to $10^{-5}$ M varenicline. (**B**) Determination of the maximum Fura-2 ratio 340/380nm in response to $10^{-8}$ M AngII, 12 mM $K^{+}$ and $10^{-9}$ to $10^{-5}$ M varenicline. (**C**) Calcium entry into the cells was evaluated using Fura-2 AM assay. Representative traces of intracellular $Ca^{2+}$ responses to $10^{-8}$ M AngII, 12 mM $K^{+}$ and $10^{-9}$ to $10^{-5}$ M uPSEM-817. (**D**) Determination of the maximum Fura-2 ratio 340/380nm in response to $10^{-8}$ M AngII, 12 mM $K^{+}$ and $10^{-9}$ to $10^{-5}$ M uPSEM-817.

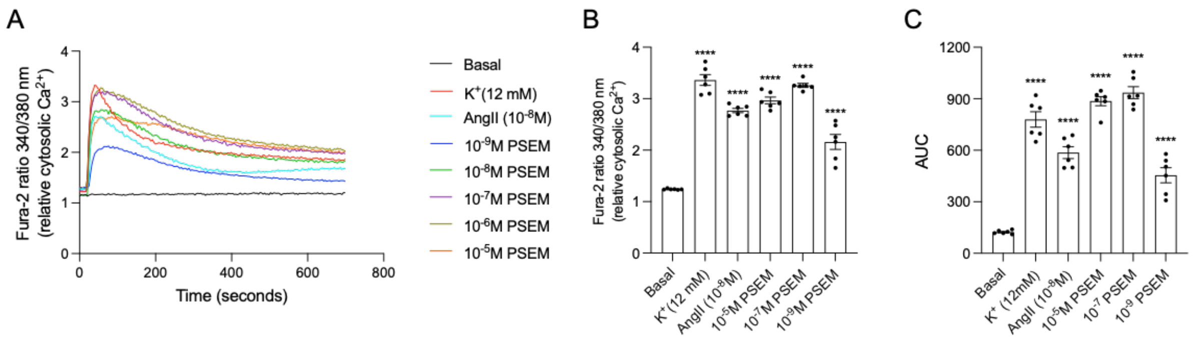


**Figure S3. Functional characterization of α7-5HT3 expressing cells (Clone 42). (A)** Calcium entry into the cells was evaluated using Fura-2 AM assay. Representative traces of intracellular $Ca^{2+}$ responses to $10^{-8}$ M AngII, 12 mM $K^{+}$ and $10^{-9}$ to $10^{-5}$ M uPSEM-817. **(B)** Determination of the maximum Fura-2 ratio 340/380nm in response to $10^{-8}$ M AngII, 12 mM $K^{+}$ and $10^{-9}$, $10^{-7}$ and $10^{-5}$ M uPSEM-817. **(C)** Area under the curve (AUC) was determined to assess whether cell depolarization led to a calcium entry in the intracellular compartment. n=6, ****, p<0,0001

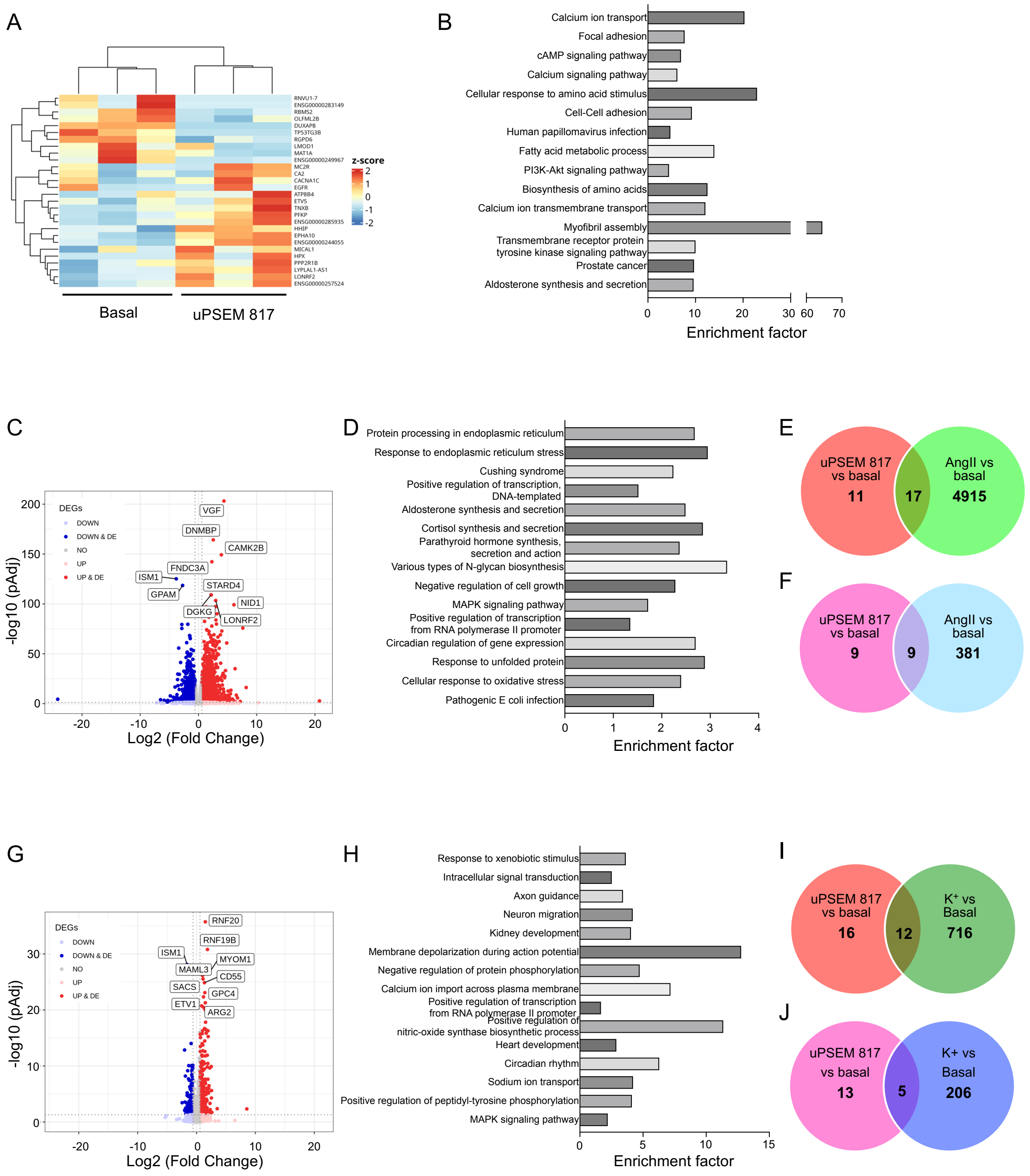

**Figure S4. Gene expression profiles of α7-5HT3-expressing H295R-S2 cells treated 8 hours with $10^{-7}$M uPSEM-817, $10^{-8}$M AngII or 12mM $K^{+}$.** (**A**) Hierarchical clustering of samples using the 28 differentially expressed genes in cells treated or not with $10^{-7}$M uPSEM-817. (**B**) Biological process enrichments determined using the list of differentially expressed genes in cells treated or not with $10^{-7}$M uPSEM-817. (**C**) Volcano plot showing the 4932 differentially in response to $10^{-8}$M AngII. Differentially expressed genes are highlighted as blue (down-regulated) or red (up-regulated) dots. (**D**) Biological process enrichments determined using the list of differentially expressed genes in cells treated or not with $10^{-8}$M AngII. (**E**) Venn diagram representing the common and different genes differentially expressed in response to $10^{-8}$M AngII and $10^{-7}$M uPSEM-817. (F) Venn diagram representing the common and different enriched biological process in response to $10^{-8}$M AngII and $10^{-7}$M uPSEM-817. (**G**) Volcano plot showing the 728 differentially in response to 12mM $K^+$. The x-axis is the Log2 fold change between the two conditions; the adjusted p value based on $-\log_{10}$ is reported on the y-axis. Genes significantly different are highlighted as blue (down-regulated in cells treated with 12mM $K^+$) or red (up-regulated in cells treated with 12mM $K^+$) dots. (**H**) Biological process enrichments determined using the list of differentially expressed genes in cells treated or not with 12mM $K^+$. (**I**) Venn diagram representing the common and different genes differentially expressed in response to 12mM $K^+$ and $10^{-7}$M uPSEM-817. (J) Venn diagram representing the common and different enriched biological process in response to 12mM $K^+$ and $10^{-7}$M uPSEM-817.

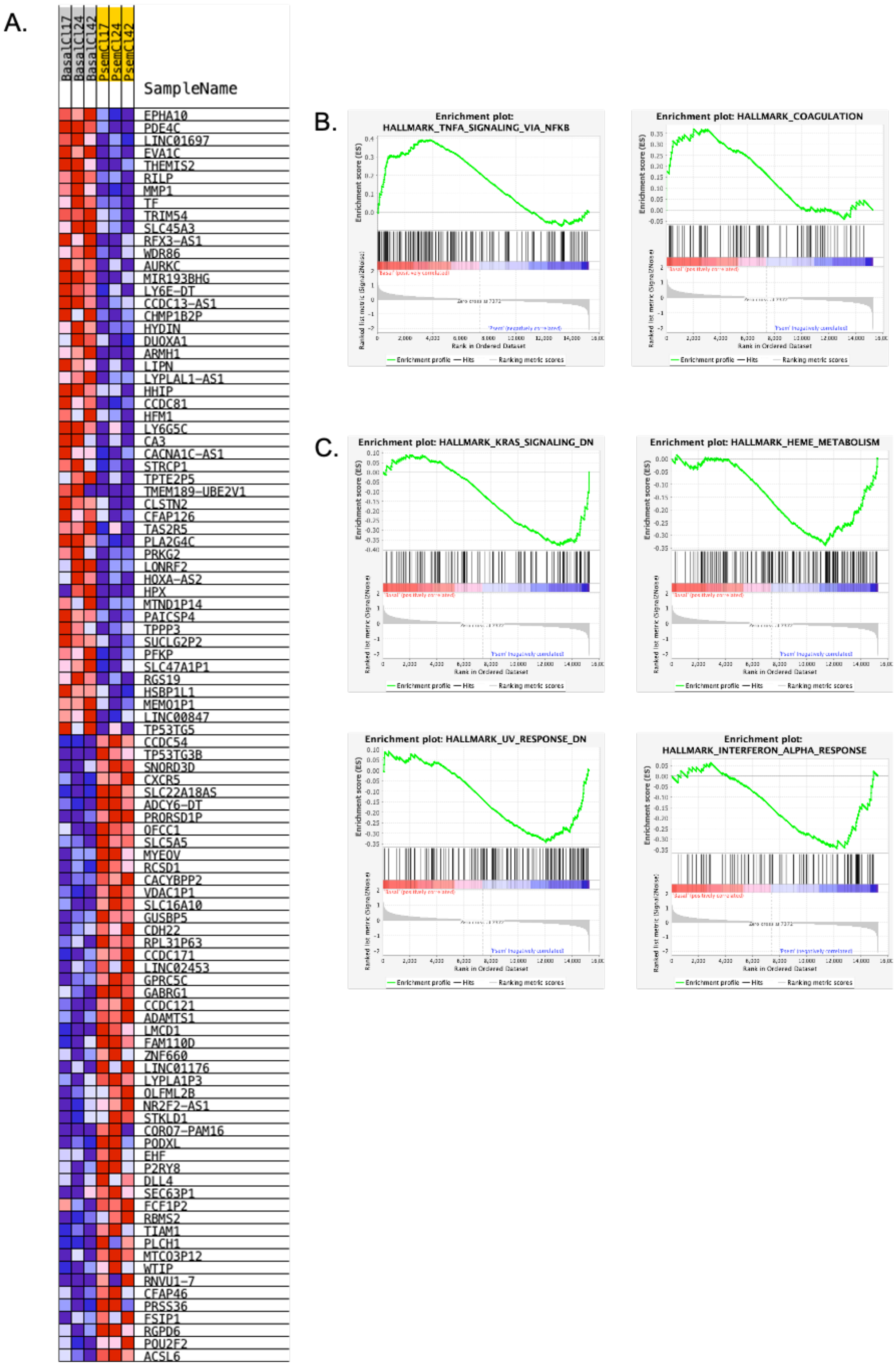


**Figure S5. RNA sequencing of α7-5HT3-expressing H295R-S2 cells treated 8 hours with $10^{-7}$M uPSEM-817 using GSEA.** A. Top 50 upregulated and downregulated genes in basal condition in comparison to uPSEM-817 treated cells. B. Pathways upregulated in basal condition. C. pathways upregulated in uPSEM-817 treated cells.

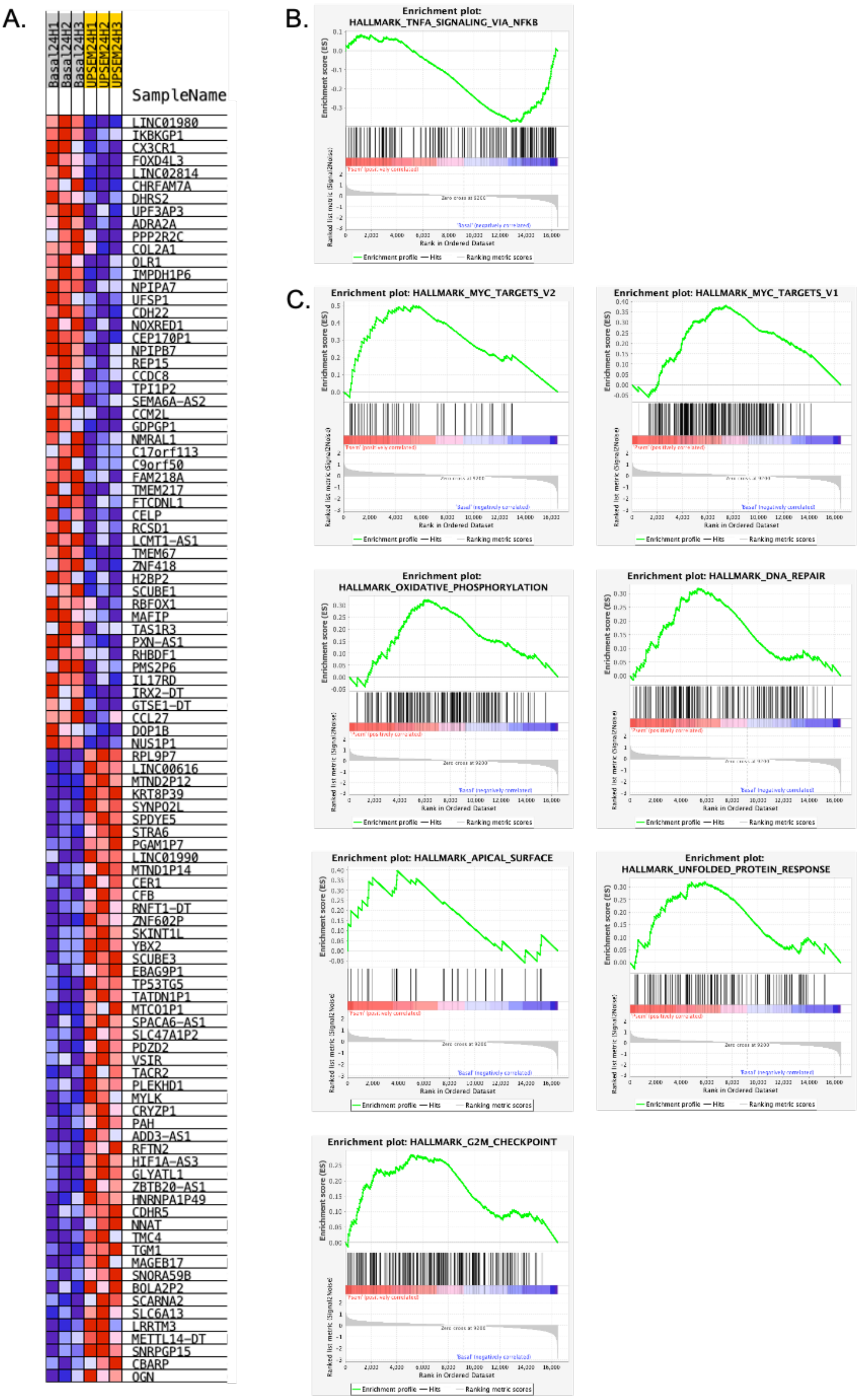


**Figure S6. RNA sequencing of α7-5HT3-expressing H295R-S2 cells treated 24 hours with $10^{-7}$M uPSEM-817 using GSEA.** A. Top 50 upregulated and downregulated genes in basal condition in comparison to uPSEM-817 treated cells. B. Pathways upregulated in basal condition. C. pathways upregulated in uPSEM-817 treated cells.

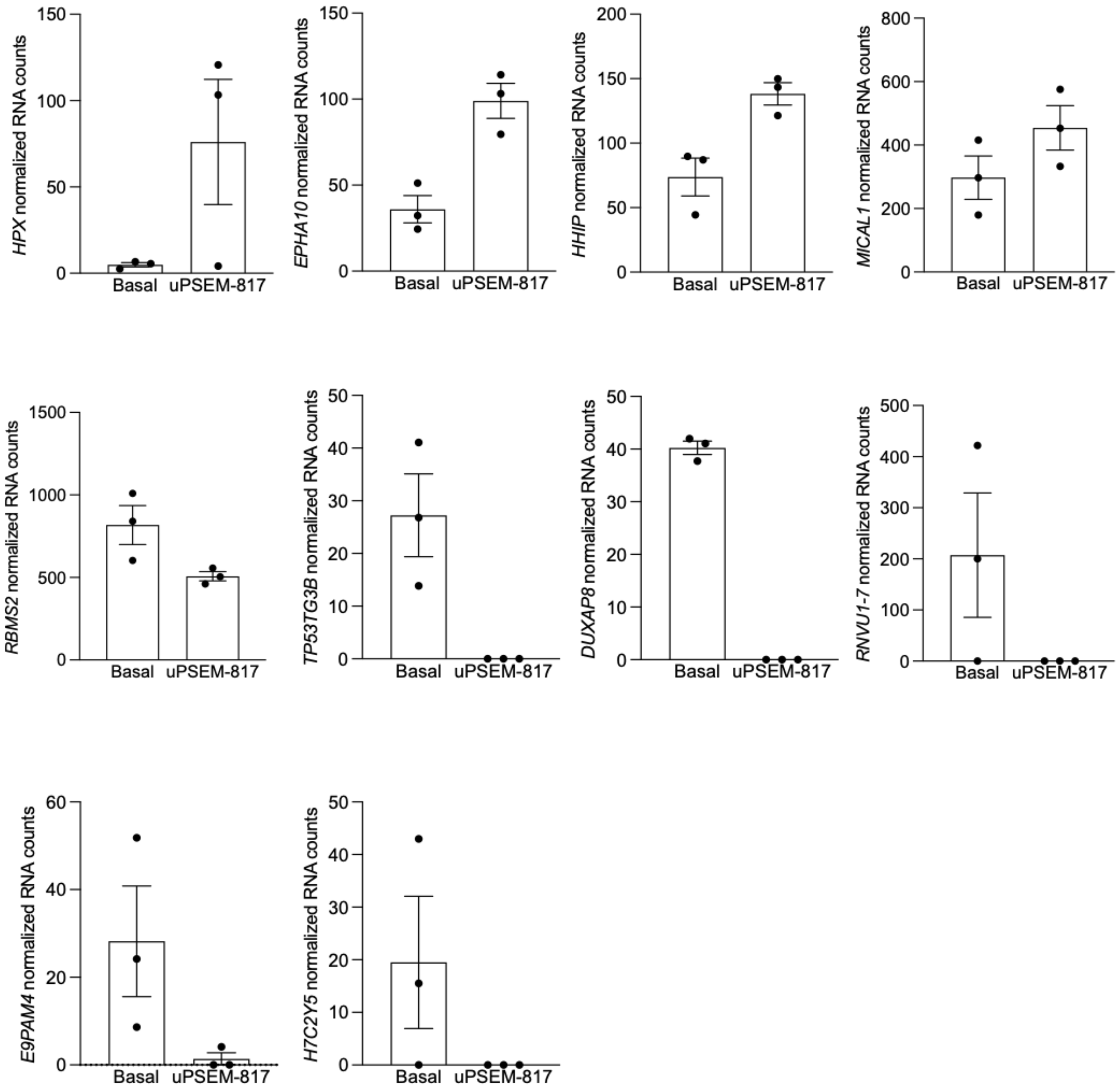


**Figure S7. Normalized RNA counts for the 10 genes from the $Na^+$ signature at 8h**

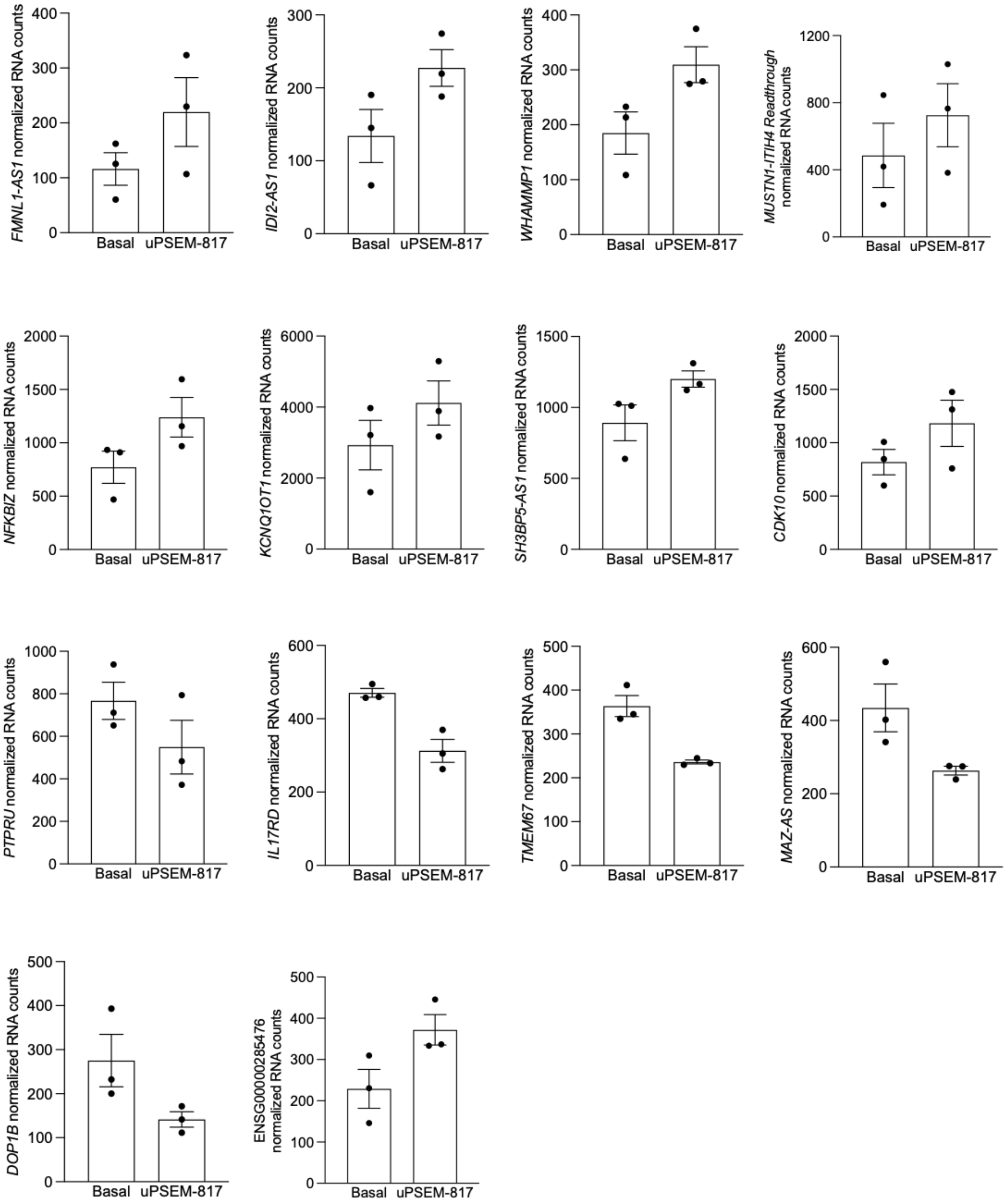


**Figure S8. Normalized RNA counts for the 10 genes from the $Na^+$ signature at 24h**

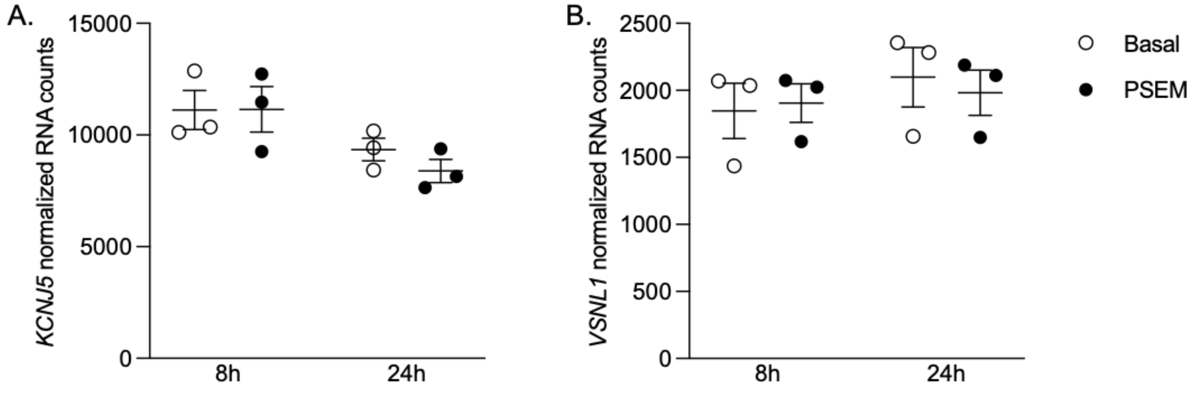


**Figure S9. Normalized RNA counts for KCNJ5 (A) and VSNL1 (B)**

### Table S1. Primers used for RT-qPCR

| Gene symbol | Forward primer | Reverse primer |
|---|---|---|
| *18S* | CCCTGCCTTTGTACACACC | CGATCCGAGGGCCTCACTA |
| *HPRT* | CTCAACTTTAACTGGAAAGAATGTC | TCCTTTTCACCAGCAAGCT |
| *GAPDH* | TGCACCACCAACTGCTTAGC | GGCATGGACTGTGGTCATGAG |
| *α7-5HT3 receptor* | CCAACGTGGGAGTCAATTCT | TTGAGTTTGCAGTGCTGGAC |

**Table S2. List of differentially expressed genes in H295R_S2 cells expressing the α7-5HT3 receptor in response to 8h treatment with $10^{-7}$M uPSEM-817.**

| Ensembl Gene ID | Gene Symbol | Gene Name | Log2 FC | pAdj |
|---|---|---|---|---|
| **ENSG00000110169** | **HPX** | **Hemopexin** | **3.4957** | **0.00432** |
| **ENSG00000285935** | **NA** | **Novel transcript, antisense to WASF3** | **1.7757** | **0.01352** |
| **ENSG00000183317** | **EPHA10** | **EPH receptor A10** | **1.4789** | **0.03196** |
| **ENSG00000104043** | **ATP8B4** | **ATPase phospholipid transporting 8B4** | **1.0694** | **0.00674** |
| **ENSG00000185231** | **MC2R** | **Melanocortin 2 receptor** | **0.9767** | **4.198e-05** |
| **ENSG00000164161** | **HHIP** | **Hedgehog interacting protein** | **0.9651** | **0.01800** |
| **ENSG00000257524** | **NA** | **Novel protein** | **0.9099** | **0.00063** |
| **ENSG00000244405** | **ETV5** | **ETS variant transcription factor 5** | **0.842** | **7.871e-05** |
| **ENSG00000244055** | **NA** | **Novel transcript, antisense to ERVW-1 and PEX1** | **0.8221** | **0.04555** |
| **ENSG00000146648** | **EGFR** | **Epidermal growth factor receptor** | **0.7711** | **0.02590** |
| **ENSG00000228536** | **LYPLAL1-AS1** | **LYPLAL1 antisense RNA 1** | **0.7645** | **0.00067** |
| **ENSG00000170500** | **LONRF2** | **LON peptidase N-terminal domain and ring finger 2** | **0.7204** | **4.198e-05** |
| **ENSG00000104267** | **CA2** | **Carbonic anhydrase 2** | **0.6865** | **0.02242** |
| **ENSG00000135596** | **MICAL1** | **Microtubule associated monooxygenase, calponin and LIM domain containing 1** | **0.6473** | **0.01159** |
| **ENSG00000067057** | **PFKP** | **Phosphofructokinase, platelet** | **0.6213** | **0.00191** |
| **ENSG00000151067** | **CACNA1C** | **Calcium voltage-gated channel subunit alpha1 C** | **0.5847** | **7.096e-07** |
| **ENSG00000168477** | **TNXB** | **Tenascin XB** | **0.5803** | **0.00747** |
| **ENSG00000137713** | **PPP2R1B** | **Protein phosphatase 2 scaffold subunit Abeta** | **0.513** | **4.198e-05** |
| ENSG00000111424 | VDR | Vitamin D receptor | 0.4996 | 0.03117 |
| ENSG00000128564 | VGF | VGF nerve growth factor inducible | 0.4384 | 0.00747 |
| ENSG00000157087 | ATP2B2 | ATPase plasma membrane Ca2+ transporting 2 | 0.4296 | 0.02590 |
| ENSG00000070061 | ELP1 | Elongator complex protein 1 | 0.4257 | 0.02357 |
| ENSG00000165868 | HSPA12A | Heat shock protein family A (Hsp70) member 12A | 0.4186 | 0.03657 |
| ENSG00000108515 | ENO3 | Enolase 3 | 0.4082 | 0.00141 |
| ENSG00000095397 | WHRN | Whirlin | 0.4015 | 0.02769 |
| ENSG00000235431 | NA | Novel transcript | 0.3981 | 0.02829 |
| ENSG00000173262 | SLC2A14 | Solute carrier family 2 member 14 | 0.393 | 0.02242 |
| ENSG00000196782 | MAML3 | Mastermind like transcriptional coactivator 3 | 0.3709 | 0.02829 |
| ENSG00000102531 | FNDC3A | Fibronectin type III domain containing 3A | 0.3251 | 0.03544 |
| ENSG00000162909 | CAPN2 | Calpain 2 | 0.3029 | 0.03117 |
| ENSG00000025772 | TOMM34 | Translocase of outer mitochondrial membrane 34 | 0.2986 | 0.00747 |
| ENSG00000170873 | MTSS1 | MTSS I-BAR domain containing 1 | 0.2905 | 0.02232 |
| ENSG00000085733 | CTTN | Cortactin | 0.2894 | 0.02590 |
| ENSG00000135424 | ITGA7 | Integrin subunit alpha 7 | 0.2778 | 0.02829 |
| ENSG00000118985 | ELL2 | Elongation factor for RNA polymerase II 2 | -0.3122 | 0.02829 |
| ENSG00000166224 | SGPL1 | Sphingosine-1-phosphate lyase 1 | -0.323 | 0.01800 |
| ENSG00000065183 | WDR3 | WD repeat domain 3 | -0.3486 | 0.01480 |
| ENSG00000150593 | PDCD4 | Programmed cell death 4 | -0.3704 | 0.03149 |
| ENSG00000119927 | GPAM | Glycerol-3-phosphate acyltransferase, mitochondrial | -0.3834 | 0.00539 |
| ENSG00000154330 | PGM5 | Phosphoglucomutase 5 | -0.4277 | 0.00141 |
| ENSG00000170962 | PDGFD | Platelet derived growth factor D | -0.429 | 0.01762 |
| ENSG00000164484 | TMEM200A | Transmembrane protein 200A | -0.4402 | 0.00484 |

| | | | | |
|---|---|---|---|---|
| ENSG00000101230 | ISM1 | Isthmin 1 | -0.4849 | 0.00432 |
| ENSG00000198785 | GRIN3A | Glutamate ionotropic receptor NMDA type subunit 3A | -0.4935 | 2.088e-06 |
| ENSG00000183044 | ABAT | 4-aminobutyrate aminotransferase | -0.4943 | 0.02195 |
| ENSG00000183054 | RGPD6 | RANBP2 like and GRIP domain containing 6 | -0.5431 | 0.02829 |
| **ENSG00000076067** | **RBMS2** | **RNA binding motif single stranded interacting protein 2** | **-0.6762** | **0.01013** |
| **ENSG00000162745** | **OLFML2B** | **Olfactomedin like 2B** | **-0.7848** | **0.03077** |
| **ENSG00000163431** | **LMOD1** | **Leiomodin 1** | **-0.8481** | **0.00491** |
| **ENSG00000151224** | **MAT1A** | **Methionine adenosyltransferase 1A** | **-2.1222** | **0.00674** |
| **ENSG00000249967** | **NA** | **Novel protein** | **-4.1038** | **0.01762** |
| **ENSG00000283149** | **NA** | **Novel protein** | **-5.9724** | **0.01159** |
| **ENSG00000261509** | **TP53TG3B** | **TP53 target 3B** | **-7.1268** | **0.00078** |
| **ENSG00000271672** | **DUXAP8** | **Double homeobox A pseudogene 8** | **-7.8953** | **8.305e-05** |
| **ENSG00000206585** | **RNVU1-7** | **RNA, variant U1 small nuclear 7** | **-27.6958** | **7.096e-07** |

Differentially expressed genes with a log2 fold changes <-0.5 or >0.5 and an adjusted P-value <0.5 are highlighted in bold.

**Table S3. List of differentially expressed genes in H295R_S2 cells expressing the α7-5HT3 receptor in response to 24h treatment with $10^{-7}$M uPSEM-817.**

| Ensembl Gene ID | Gene Symbol | Gene Name | Log2 FC | pAdj |
|---|---|---|---|---|
| **ENSG00000107738** | **VSIR** | **V-set immunoregulatory receptor** | **1.7690** | **0.02216** |
| **ENSG00000266401** | **NA** | **novel transcript, antisense to DLGAP1** | **1.2938** | **0.02340** |
| **ENSG00000179142** | **CYP11B2** | **Cytochrome P450 family 11 subfamily B member 2** | **1.2735** | **0.01902** |
| **ENSG00000151006** | **PRSS53** | **Serine protease 53** | **1.0692** | **0.02137** |
| **ENSG00000147174** | **GCNA** | **germ cell nuclear acidic peptidase** | **0.9139** | **0.03037** |
| **ENSG00000233175** | **NA** | **Novel transcript, antisense to FMNL1** | **0.9047** | **0.00399** |
| **ENSG00000232656** | **IDI2-AS1** | **IDI2 antisense RNA 1** | **0.8578** | **0.03037** |
| **ENSG00000185231** | **MC2R** | **Melanocortin 2 receptor** | **0.8273** | **0.00127** |
| **ENSG00000223509** | **NA** | **WAS protein homolog associated with actin, golgi membranes and microtubules (WHAMM) pseudogene** | **0.8026** | **0.03613** |
| **ENSG00000285476** | **NA** | **Novel transcript** | **0.7226** | **0.04326** |
| **ENSG00000243696** | **NA** | **Novel MUSTN1-ITIH4 readthrough** | **0.7216** | **0.02137** |
| **ENSG00000144802** | **NFKBIZ** | **NFKB inhibitor zeta** | **0.7207** | **0.01244** |
| **ENSG00000244405** | **ETV5** | **ETS variant transcription factor 5** | **0.7151** | **0.00378** |
| **ENSG00000228536** | **LYPLAL1-AS1** | **LYPLAL1 antisense RNA 1** | **0.6532** | **0.01145** |
| **ENSG00000091428** | **RAPGEF4** | **Rap guanine nucleotide exchange factor 4** | **0.6075** | **0.00378** |
| **ENSG00000197948** | **FCHSD1** | **FCH and double SH3 domains 1** | **0.5923** | **0.00407** |
| **ENSG00000269821** | **KCNQ1OT1** | **KCNQ1 opposite strand/antisense transcript 1** | **0.5729** | **0.02697** |
| **ENSG00000137713** | **PPP2R1B** | **Protein phosphatase 2 scaffold subunit Abeta** | **0.5585** | **1.00e-5** |
| **ENSG00000108352** | **RAPGEFL1** | **Rap guanine nucleotide exchange factor like 1** | **0.5524** | **0.03804** |
| **ENSG00000224660** | **SH3BP5-AS1** | **SH3BP5 antisense RNA 1** | **0.5411** | **0.01859** |
| **ENSG00000147894** | **C9orf72** | **C9orf72-SMCR8 complex subunit** | **0.5271** | **0.00074** |
| **ENSG00000185324** | **CDK10** | **Cyclin dependent kinase 10** | **0.5058** | **0.02137** |
| ENSG00000070061 | ELP1 | Elongator complex protein 1 | 0.4966 | 0.00378 |
| ENSG00000268350 | FAM156A | Family with sequence similarity 156 member A | 0.4918 | 0.02216 |
| ENSG00000164050 | PLXNB1 | Plexin B1 | 0.4914 | 0.02502 |
| ENSG00000117586 | TNFSF4 | TNF superfamily member 4 | 0.4906 | 0.04201 |
| ENSG00000151067 | CACNA1C | Calcium voltage-gated channel subunit alpha1 C | 0.4829 | 0.00022 |
| ENSG00000155265 | GOLGA7B | Golgin A7 family member B | 0.4794 | 0.00489 |
| ENSG00000245532 | NEAT1 | Nuclear paraspeckle assembly transcript 1 | 0.4739 | 0.01859 |
| ENSG00000224086 | PPM1F-AS1 | PPM1F antisense RNA 1 | 0.4681 | 0.04716 |
| ENSG00000249087 | ZNF436-AS1 | ZNF436 antisense RNA 1 | 0.4581 | 0.03257 |
| ENSG00000111674 | ENO2 | Enolase 2 | 0.4489 | 0.00378 |
| ENSG00000197635 | DPP4 | Dipeptidyl peptidase 4 | 0.4332 | 0.04360 |
| ENSG00000250159 | NA | Novel transcript | 0,4296 | 0.03037 |
| ENSG00000246859 | STARD4-AS1 | STARD4 antisense RNA 1 | 0.4237 | 0.03525 |
| ENSG00000155657 | TTN | Titin | 0.4168 | 0.02418 |
| ENSG00000095397 | WHRN | Whirlin | 0.4087 | 0.02137 |
| ENSG00000183426 | NPIPA1 | Nuclear pore complex interacting protein family member A1 | 0.4027 | 0.03804 |
| ENSG00000235431 | NA | Novel transcript | 0.3881 | 0.02697 |
| ENSG00000143702 | CEP170 | Centrosomal protein 170 | 0.3733 | 0.01859 |
| ENSG00000259848 | NA | Pseudogene similar to part of asparagine synthetase (ASNS) | 0.3576 | 0.04350 |

| | | | | |
|---|---|---|---|---|
| ENSG00000085733 | CTTN | Cortactin | 0.3205 | 0.00798 |
| ENSG00000162552 | WNT4 | Wnt family member 4 | -0.2488 | 0.04716 |
| ENSG00000168159 | RNF187 | ring finger protein 187 | -0.2507 | 0.02697 |
| ENSG00000140526 | ABHD2 | Abhydrolase domain containing 2, acylglycerol lipase | -0.2548 | 0.04716 |
| ENSG00000198795 | ZNF521 | zinc finger protein 521 | -0.2646 | 0.03037 |
| ENSG00000185551 | NR2F2 | Nuclear receptor subfamily 2 group F member 2 | -0.2732 | 0.02137 |
| ENSG00000197457 | STMN3 | Stathmin 3 | -0.2737 | 0.04787 |
| ENSG00000177733 | HNRNPA0 | Heterogeneous nuclear ribonucleoprotein A0 | -0.2762 | 0.01244 |
| ENSG00000183741 | CBX6 | Chromobox 6 | -0.2782 | 0.049556 |
| ENSG00000124422 | USP22 | ubiquitin specific peptidase 22 | -0.2846 | 0.02502 |
| ENSG00000166224 | SGPL1 | sphingosine-1-phosphate lyase 1 | -0.3051 | 0.02473 |
| ENSG00000130720 | FIBCD1 | Fibrinogen C domain containing 1 | -0.3250 | 0.03233 |
| ENSG00000121653 | MAPK8IP1 | Mitogen-activated protein kinase 8 interacting protein 1 | -0.3274 | 0.04011 |
| ENSG00000132970 | WASF3 | WASP family member 3 | -0.3311 | 0.02924 |
| ENSG00000085741 | WNT11 | Wnt family member 11 | -0.3369 | 0.02502 |
| ENSG00000075618 | FSCN1 | Fascin actin-bundling protein 1 | -0.3375 | 0.00167 |
| ENSG00000130749 | ZC3H4 | Zinc finger CCCH-type containing 4 | -0.3511 | 0.02202 |
| ENSG00000277443 | MARCKS | Myristoylated alanine rich protein kinase C substrate | -0.3648 | 0.00022 |
| ENSG00000177666 | PNPLA2 | Patatin like phospholipase domain containing 2 | -0.3744 | 0.04053 |
| ENSG00000115255 | REEP6 | Receptor accessory protein 6 | -0.4093 | 0.01529 |
| ENSG00000253276 | CCDC71L | Coiled-coil domain containing 71 like | -0.4101 | 0.01244 |
| ENSG00000157600 | TMEM164 | Transmembrane protein 164 | -0.4159 | 0.00378 |
| ENSG00000101230 | ISM1 | Isthmin 1 | -0.4235 | 0.02216 |
| ENSG00000083812 | ZNF324 | Zinc finger protein 324 | -0.4967 | 0.01244 |
| **ENSG00000060656** | **PTPRU** | **Protein tyrosine phosphatase receptor type U** | **-0.5225** | **0.01244** |
| **ENSG00000155090** | **KLF10** | **Kruppel like factor 10** | **-0.5225** | **0.04760** |
| **ENSG00000144730** | **IL17RD** | **Interleukin 17 receptor D** | **-0.6023** | **0.01244** |
| **ENSG00000112320** | **SOBP** | **Sine oculis binding protein homolog** | **-0.6111** | **0.03232** |
| **ENSG00000164953** | **TMEM167** | **Transmembrane protein 167** | **-0.6136** | **0.01859** |
| **ENSG00000259952** | **NA** | **novel transcript, antisense to MAZ** | **-0.6749** | **0.02697** |
| **ENSG00000150594** | **ADRA2A** | **Adrenoceptor alpha 2A** | **-0.8037** | **0.03350** |
| **ENSG00000142197** | **DOP1B** | **DOP1 leucine zipper like protein B** | **-0.9376** | **0.02924** |

Differentially expressed genes with a log2 fold changes <-0.5 or >0.5 and an adjusted P-value <0.5 are highlighted in bold.

**Table S4: List of differentially expressed genes in H295R_S2 cells expressing the α7-5HT3 receptor in response to 8h treatment with AngII.**

**Table S5: List of differentially expressed genes in H295R_S2 cells expressing the α7-5HT3 receptor in response to 8h treatment with $K^+$.**

**Table S6: List of significantly enriched pathways in H295R_S2 cells expressing the α7-5HT3 receptor in response to 8h treatment with $10^{-8}$M AngII.**

**Table S7: List of significantly enriched pathways in H295R_S2 cells expressing the α7-5HT3 receptor in response to 8h treatment with 12mM $K^+$.**

**Table S8: List of differentially expressed genes in H295R_S2 cells expressing the α7-5HT3 receptor in response to 24h treatment with $10^{-8}$M AngII.**

**Table S9: List of significantly enriched pathways in H295R_S2 cells expressing the α7-5HT3 receptor in response to 24h treatment with $10^{-8}$M AngII.**

**Table S10: List of differentially expressed genes in H295R_S2 cells expressing the α7-5HT3 receptor in response to 24h treatment with 12mM $K^+$.**

**Table S11: List of significantly enriched pathways in H295R_S2 cells expressing the α7-5HT3 receptor in response to 24h treatment with 12mM $K^+$.**

**Table S12. Comparison of genes differentially expressed after 8h of treatment with uPSEM-817 with that obtained in control adrenal and APA extracted from transcriptomic data**

| Gene name | Control Adrenal (n=11) | APA without KCNJ5 mutation (n=73) | P value | APA with KCNJ5 Mutation (n=50) | P value | Gene name | Log2 FC In cells | Padj In cells |
|---|---|---|---|---|---|---|---|---|
| *CACNA1C* Probe 1 | 7.182±1.387 | 23.28±0.309*** | 0.0002 | 18.79±1.969** | 0.0068 | *CACNA1C* | 0.5847 | 7.09e-07 |
| *CACNA1C* Probe 2 | 2.491±0.3533 | 5.363±0.236**** | <0,0001 | 5.161±0,309*** | 0.0002 | | | |
| *RNVU1-7* | *N.A.* | *N.A.* | | *N.A.* | *N.A.* | *RNVU1-7* | -27.6957 | 7.09e-07 |
| *GRIN3A* | 0.484±0.063 | 0.539±0.042 | >0.9999 | 0.558±0.068 | >0.9999 | *GRIN3A* | -0.4935 | 2.09e-06 |
| *PPP2R1B* | 3.749±0.377 | 2.871±0.182* | 0.0499 | 2.650±0.164* | 0.0375 | *PPP2R1B* | 0.5129 | 4.19e-05 |
| *LONRF2* Probe 1 | 6.666±0.781 | 3.459±0.220** | 0.0015 | 3.808±0.403** | 0.0022 | *LONRF2* | 0.7204 | 4.19e-05 |
| *LONRF2* Probe 2 | 1.398±0.142 | 0.998±0.057* | 0.0310 | 1.079±0.097* | 0.0490 | | | |
| *LONRF2* Probe 3 | 4.805±0.639 | 2.428±0.143** | 0.0013 | 2.482±0.233** | 0.0011 | | | |
| *LONRF2* Probe 4 | 0.980±0.132 | 0.621±0.070** | 0.0042 | 0.698±0.078 | 0.0634 | | | |
| *MC2R* | 9.003±1.263 | 14.76±1.907 | 0.6319 | 14.73±1.790 | 0.8617 | *MC2R* | 0.9767 | 4.19e-05 |
| *ETV5* Probe 1 | 12.51±1.491 | 16.69±1.153 | 0.3827 | 17.35±1.452 | 0.2815 | *ETV5* Probe 1 | 0.8420 | 7.87e-05 |
| *ETV5* Probe 2 | 13.35±1.831 | 8.588±0.796* | 0.0209 | 9.059±0.897 | 0.0670 | *ETV5* Probe 2 | | |
| *DUXAP8* | *N.A.* | *N.A.* | *N.A.* | *N.A.* | *N.A.* | *DUXAP8* | -7.8953 | 8.31e-05 |
| *LYPLAL1-AS1* | *N.A.* | *N.A.* | *N.A.* | *N.A.* | *N.A.* | *LYPLAL1-AS1* | 0.7644 | 0.0007 |
| *TP53TG3B* | *N.A.* | *N.A.* | *N.A.* | *N.A.* | *N.A.* | *TP53TG3B* | -7.1268 | 0.0008 |
| *ENO3* | 0.484±0.077 | 0.838±0.117 | 0.4299 | 0.910±0.130 | 0.1886 | *ENO3* | 0.4083 | 0.0014 |
| *PGM5* Probe 1 | 1.632±0.236 | 1.608±0.195 | 0.3469 | 1.824±0.243 | 0.8781 | *PGM5* | -0.4277 | 0.0014 |
| *PGM5* Probe 2 | 2.718±0.312 | 2.761±0.375 | 0.1848 | 3.139±0.432 | 0.4021 | | | |
| *PGM5* Probe 3 | 1.663±0.148 | 1.648±0.189 | 0.4054 | 1.859±0.196 | >0.9999 | | | |
| *PFKP* | 0.977±0.136 | 0.853±0.070 | 0.4602 | 0.839±0.010 | 0.2831 | *PFKP* | 0.6213 | 0.0020 |
| *ISM1* | 0.467±0.044 | 0.516±0.036 | >0.9999 | 0.617±0.064 | >0.9999 | *ISM1* | -0.4889 | 0.0043 |
| *HPX* | 0.684±0.120 | 1.027±0.078 | 0.0789 | 1.070±0.073* | 0.0225 | *HPX* | 3.4957 | 0.0043 |
| *TMEM200A* | 13.86±1.74 | 44.11±5.402** | 0.0023 | 34.47±4.164** | 0.0085 | *TMEM200A* | -0.4402 | 0.0048 |
| *LMOD1* | *N.A.* | *N.A.* | *N.A.* | *N.A.* | *N.A.* | *LMOD1* | -0.8481 | 0.0049 |
| *GPAM* | 3.490±0.514 | 2.050±0.153** | 0.0044 | 2.481±0.212 | 0.1049 | *GPAM* | -0,3833 | 0.0054 |

| | | | | | | | | |
|---|---|---|---|---|---|---|---|---|
| *ATP8B4* Probe 1 | 0.375±0.052 | 0.381±0.032 | >0.9999 | 0.584±0.095 | 0.9362 | *ATP8B4* | 1.0694 | 0.0067 |
| *ATP8B4* Probe 2 | 1.452±0.206 | 0.935±0.127** | 0.0051 | 0.833±0.082** | 0.0055 | | | |
| *MAT1A* | 0.784±0.144 | 1.005±0.138 | >0.9999 | 0.878±0.067 | 0.9774 | *MAT1A* | -2.1222 | 0.0067 |
| *TOMM34* | 5.632±0.812 | 7.625±0.815 | >0.9999 | 7.002±0.937 | >0.9999 | *TOMM34* | 0.2986 | 0.0075 |
| *VGF* | 1.213±0.347 | 1.415±0.428 | 0.6481 | 0.738±0.126 | 0.2161 | *VGF* | 0.4384 | 0.0075 |
| *TNXB* Probe 1 | 32.73±7.049 | 17.87±1.49* | 0.0219 | 17.68±1.965* | 0.0148 | *TNXB* | 0.5803 | 0.0075 |
| *TNXB* Probe 2 | 6.434±1.258 | 9.889±0.628 | 0.1291 | 9.389±0.628 | 0.1221 | | | |
| *TNXB* Probe 3 | 0.324±0.033 | 0.488±0.037 | 0.1505 | 0.686±0.167 | 0.1607 | | | |
| *RBMS2* | 0.604±0.156 | 0.877±0.092 | 0.4378 | 0.952±0.118 | 0.2479 | *RBMS2* | -0.6762 | 0.0101 |
| *MICAL1* | 0.755±0.086 | 1.150±0.240 | >0.9999 | 1.036±0.104 | >0.9999 | *MICAL1* | 0.6473 | 0.0116 |
| *WDR3* | 4.629±0.894 | 5.597±0.493 | 0.7235 | 3.718±0.275 | >0.9999 | *WDR3* | -0.3486 | 0.0148 |
| *PDGFD* | 180.3±13.29 | 146.9±9.59 | 0.0551 | 154.1±13.22 | 0.3079 | *PDGFD* | -0.4290 | 0.0176 |
| *HHIP* | 0.438±0.045 | 0.527±0.038 | >0.9999 | 0.448±0.031 | >0.9999 | *HHIP* | 0.9651 | 0.0180 |
| *SGPL1* | 0.734±0.099 | 0.639±0.049 | 0.3380 | 0.836±0.111 | >0.9999 | *SGPL1* | -0.3230 | 0.0180 |
| *ABAT* Probe 1 | 0.703±0.122 | 0.788±0.061 | >0.9999 | 0.963±0.077 | 0.3532 | *ABAT* | -0.4943 | 0.0220 |
| *ABAT* Probe 2 | 2.784±0.392 | 2.969±0.301 | >0.9999 | 3.125±0.379 | >0.9999 | | | |
| *MTSS1* | 17.06±1.279 | 18.98±1.317 | >0.9999 | 16.41±1.690 | 0.6468 | *MTSS1* | 0.2905 | 0.0223 |
| *CA2* | 0.893±0.144 | 3.736±2.046 | 0.1678 | 1.231±0.151 | >0.9999 | *CA2* | 0.6865 | 0.0224 |
| *SLC2A14* | 3.231±0.525 | 3.164±0.151 | >0.9999 | 2.745±0.155 | >0.9999 | *SLC2A14* | 0.3930 | 0.0224 |
| *ELP1* | 116.0±15.54 | 149.8±10.71 | 0.5960 | 131.6±12.90 | >0.9999 | *ELP1* | 0.4257 | 0.0236 |
| *CTTN* | 134.4±9.24 | 152.7±7.578 | 0.9233 | 146.3±9.167 | 0.8119 | *CTTN* | 0.2894 | 0.0259 |
| *EGFR* | 2.950±0.443 | 3.525±0.238 | 0.9516 | 3.591±0.273 | 0.6857 | *EGFR* | 0.7711 | 0.0259 |
| *ATP2B2* | 0.556±0.067 | 1.292±0.110** | 0.0018 | 0.955±0.092* | 0.0387 | *ATP2B2* | 0.4296 | 0.0259 |
| *WHRN* | 9.832±1.618 | 7.042±0.798 | 0.0984 | 5.658±0.992* | 0.0114 | *WHRN* | 0.4015 | 0.0277 |
| *ELL2* | 5.401±0.437 | 12.99±1.020**** | <0.0001 | 11.42±0.839*** | 0.0002 | *ELL2* | -0.3121 | 0.0283 |
| *ITGA7* | 2.537±0.452 | 1.598±0.139* | 0.0258 | 1.186±0.089** | 0.0024 | *ITGA7* | 0.2778 | 0.0283 |
| *RGPD6* | 2.140±0.365 | 0.897±0.071** | 0.0021 | 1.088±0.118* | 0.0215 | *RGPD6* | -0.5431 | 0.0283 |
| *MAML3* | 0.643±0.100 | 0.659±0.077 | >0.9999 | 0.615±0.057 | 0.8855 | *MAML3* | 0.3709 | 0.0283 |
| *OLFML2B* | 2.745±0.362 | 4.029±0.189* | 0.0282 | 3.774±0.283 | 0.1035 | *OLFML2B* | -0.7848 | 0.0308 |
| *VDR* | 18.37±5.70 | 54.16±6.75** | 0.0066 | 45.01±4.95** | 0.0085 | *VDR* | 0.4996 | 0.0312 |

| | | | | | | | | |
|---|---|---|---|---|---|---|---|---|
| *CAPN2* | 94.77±6.37 | 97.04±5.49 | >0.9999 | 101,9±6,83 | >0.9999 | *CAPN2* | 0.3029 | 0.0315 |
| *PDCD4* | 4.324±0.657 | 4.820±0.398 | >0.9999 | 4.691±0,477 | >0.9999 | *PDCD4* | -0.3704 | 0.0315 |
| *EPHA10* Probe 1 | 0,908±0.119 | 0.774±0.048 | 0.4132 | 0.880±0,090 | 0.6644 | *EPHA10* | 1.4789 | 0.0320 |
| *EPHA10* Probe 2 | 0.514±0.072 | 0,686±0.077 | 0.6054 | 0.731±0.061 | 0.1850 | | | |
| *EPHA10* Probe 3 | 0.530±0.124 | 0.395±0.035 | 0.4312 | 0.577±0.562 | >0.9999 | | | |
| *EPHA10* Probe 4 | 0.938±0.191 | 1.570±0.162* | 0.0458 | 1.469±0,115* | 0.0431 | | | |
| *EPHA10* Probe 5 | 0.620±0.157 | 0.529±0.050 | >0.9999 | 0.489±0,051 | 0.9241 | | | |
| *FNDC3A* | 26.85±2.39 | 27.31±1.96 | >0.9999 | 32.00±3.622 | >0.9999 | *FNDC3A* | 0.3251 | 0.0354 |
| *HSPA12A* Probe 1 | 12.82±10.16 | 8.087±0.785*** | 0.0031 | 6.501±0,802*** | 0.0002 | *HSPA12A* | 0.4186 | 0.0367 |
| *HSPA12A* Probe 2 | 0.578±0.043 | 0.527±0.027 | 0.6508 | 0.521±0.034 | 0.4631 | | | |

**Table S13. Comparison of genes differentially expressed after 24h of treatment with uPSEM-817 with that obtained in control adrenal and APA extracted from transcriptomic data**

| Gene name | Control Adrenal (n=11) | APA without KCNJ5 mutation (n=73) | P value | APA with KCNJ5 Mutation (n=50) | P value | Gene name | Log2 FC In cells | Padj in cells |
|---|---|---|---|---|---|---|---|---|
| *VSIR* | 3.940±0.390 | 4.701±0.336 | >0.9999 | 3.990±0.314 | >0.9999 | *VSIR* | 1.7690 | 0.0222 |
| *CYP11B2* | 123.2±18.55 | 244.4±25.57 | 0.3037 | 196.0±23.79 | 0.6421 | *CYP11B2* | 1.2735 | 0.0190 |
| *PRSS53* | *N.A.* | *N.A.* | | *N.A.* | | *PRSS53* | 1.0692 | 0.0214 |
| *GCNA* | 6.113±0.866 | 9.243±1.371 | >0.9999 | 8.274±1.736 | >0.9999 | *GCNA* | 0.9139 | 0.0304 |
| *MC2R* | 9.003±1.263 | 14.76± 1.907 | 0.8617 | 14.73±1.790 | 0.6319 | *MC2R* | 0.8723 | 0.0013 |
| *IDI2-AS1* | 0.457±0.095 | 0.496±0.049 | >0.9999 | 0.504±0.045 | >0.9999 | *IDI2-AS1* | 0.8578 | 0.0304 |
| *NFKBIZ* | 13.20±2.575 | 9.390±0.952 | 0.1062 | 7.981±0.970* | 0.0220 | *NFKBIZ* | 0.7207 | 0.0124 |
| *ETV5* Probe 1 | 12.51±1.491 | 16.69±0.901 | 0.3827 | 17.35±1.452 | 0.2815 | *ETV5* | 0.7151 | 0.0038 |
| *ETV5* Probe 2 | 13.35±1.831 | 8.588±0.796* | 0.0209 | 9.059±0.879 | 0.0670 | | | |
| *LYPLAL1-AS1* | *N.A.* | *N.A.* | | *N.A.* | | *LYPLAL1-AS1* | 0.6532 | 0.0115 |
| *RAPGEF4* | 11.66±1.013 | 12.56±1.052 | >0.9999 | 12.38±1.280 | >0.9999 | *RAPGEF4* | 0.6075 | 0.0038 |
| *FCHSD1* | 4.668±0.560 | 6.986±0.397* | 0.0280 | 6.508±0.366* | 0.0408 | *FCHSD1* | 0.5923 | 0.0041 |
| *KCNQ1OT1* | *N.A.* | *N.A.* | | *N.A.* | | *KCNQ1OT1* | 0.5729 | 0.0270 |
| *PPP2R1B* | 3.749±0.377 | 2.871±0.182* | 0.0499 | 2.650±0.164* | 0.0375 | *PPP2R1B* | 0.5585 | 1.00$^{e}$-05 |
| *RAPGEFL1* | 1.416±0.324 | 0.960±0.112 | 0.0840 | 0.812±0.065 | 0.0766 | *RAPGEFL1* | 0.5524 | 0.0380 |
| *SH3BP5-AS1* | *N.A.* | *N.A.* | | *N.A.* | | *SH3BP5-AS1* | 0.5411 | 0.0186 |
| *C9orf72* | *N.A.* | *N.A.* | | *N.A.* | | *C9orf72* | 0.5271 | 0.0007 |
| *CDK10* | 26.63±2.976 | 32.04±1.875 | 0.4963 | 30.37±2.314 | 0.4963 | *CDK10* | 0.5058 | 0.0214 |
| *ELP1* | 116.0±15.54 | 149.8±10.71 | 0.5960 | 131.6±12,90 | >0.9999 | *ELP1* | 0.4966 | 0.0038 |
| *FAM156A* | 24.08±1.232 | 30.75±1.835 | 0.5123 | 24.45±1.467 | >0.9999 | *FAM156A* | 0.4918 | 0.0222 |
| *PLXNB1* | 1.308±0.144 | 1.149±0.090 | 0.3101 | 1.190±0.127 | 0.2783 | *PLXNB1* | 0.4914 | 0.0250 |
| *TNFSF4* | 1.376±0.178 | 1.999±0.247 | >0.9999 | 2.087±0.256 | >0.9999 | *TNFSF4* | 0.4906 | 0.0420 |
| *CACNA1C* Probe 1 | 7.182±1.387 | 23.28±1.846*** | 0.0002 | 18.79±1.969** | 0.0068 | *CACNA1C* | 0.4829 | 0.0002 |
| *CACNA1C* Probe 2 | 2.491±0.353 | 5.161±0.309**** | <0.0001 | 5.161±0,309*** | 0.0002 | | | |
| *GOLGA7B* | 1.359±0.143 | 2.011±0.128 | 0.0657 | 1.840±0.114 | 0.2235 | *GOLGA7B* | 0.4794 | 0.0049 |

| | | | | | | | | |
|---|---|---|---|---|---|---|---|---|
| *NEAT1* | 4.536± 0.801 | 2.053 ± 0.302*** | 0.0002 | 2.004±0.281*** | 0.004 | *NEAT1* | 0.4739 | 0.0186 |
| *PPM1F-AS1* | *N.A.* | *N.A.* | | *N.A.* | | *PPM1F-AS1* | 0.4681 | 0.0472 |
| *ZNF436-AS1* | *N.A.* | *N.A.* | | *N.A.* | | *ZNF436-AS1* | 0.4581 | 0.0326 |
| *ENO2* | 6.728± 1.346 | 4.982 ± 0.509 | 0.1549 | 6.848±0.875 | 0.7798 | *ENO2* | 0.4489 | 0.0038 |
| *DPP4* | 0.575± 0.096 | 1.283 ± 0.258 | 0.2577 | 0.945±0.109 | 0.5216 | *DPP4* | 0.4332 | 0.0436 |
| *STARD4-AS1* | *N.A.* | *N.A.* | | *N.A.* | | *STARD4-AS1* | 0.4237 | 0.0353 |
| *TTN* | 0.696±0.106 | 2.211±0.300** | 0.0032 | 1.658±0.264* | 0.0400 | *TTN* | 0.4168 | 0.0242 |
| *WHRN* | 9.832±1.618 | 7.042±0.800 | 0.0984 | 5,658±0.992 | 0.0114 | *WHRN* | 0.4087 | 0.0214 |
| *NPIPA1* | 4.961±0.606 | 4.074±0.258 | 0.3230 | 4.212±0.330 | 0.5382 | *NPIPA1* | 0.4027 | 0.0380 |
| *CEP170* | 7.076±0.427 | 5.862±0.347* | 0.0407 | 5.755±0.395 | 0.0766 | *CEP170* | 0.3733 | 0.0186 |
| *CTTN* | 134.4±9.24 | 152.7±7.58 | 0.9233 | 146,3±9.167 | 0.8119 | *CTTN* | 0.3205 | 0.0080 |
| *WNT4* | 0.596±0.101 | 1.099±0.144 | 0.2637 | 0.963±0.106 | 0.1863 | *WNT4* | -0.2488 | 0.0472 |
| *RNF187* | 35.06±5.152 | 37.51±1.904 | 0.9193 | 40.56±3.672 | 0.8172 | *RNF187* | -0.2507 | 0.0270 |
| *ABHD2* | 20.54±3.327 | 14.25±1.205 | 0.0736 | 11.90±1.090* | 0.0227 | *ABHD2* | -0.2548 | 0.0472 |
| *ZNF521* | 18.14±2.741 | 17.70±1.397 | >0.9999 | 22.11±2.194 | >0.9999 | *ZNF521* | -0.2646 | 0.0304 |
| *NR2F2* | 18.63±2.451 | 13.33±0.798* | 0.0353 | 13.76±1.036 | 0.0631 | *NR2F2* | -0.2732 | 0.0214 |
| *STMN3* Probe 1 | 46.93±4.413 | 51.13±3.364 | >0.9999 | 36.30±2.268 | 0.1031 | *STMN3* | -0.2737 | 0.0479 |
| *STMN3* Probe 2 | 7.931±1.456 | 14.15±0.651*** | 0.0002 | 12.29±0.596** | 0.0070 | | | |
| *STMN3* Probe 3 | 2.456±0.300 | 2.708±0.179 | >0.9999 | 2.740±0.243 | >0.9999 | | | |
| *HNRNPA0* | 3.553±0.745 | 2.624±0.303 | 0.1612 | 3.067±0.491 | 0.2538 | *HNRNPA0* | -0.2762 | 0.0124 |
| *CBX6* | 17.86±1.348 | 22.18±1.348 | 0.5004 | 18.33±1.445 | >0.9999 | *CBX6* | -0.2782 | 0.0496 |
| *USP22* | 0.770±0.124 | 0.721±0.081 | 0.3230 | 0.615±0.097 | 0.1410 | *USP22* | -0.2846 | 0.0250 |
| *SGPL1* | 0.734±0.099 | 0.639±0.049 | 0.3293 | 0.828±0.109 | >0.9999 | *SGPL1* | -0.3051 | 0.0247 |
| *FIBCD1* | 7.464±0.681 | 11.25±0.535* | 0.0125 | 10.68±0.715* | 0.0376 | *FIBCD1* | -0.3250 | 0.0323 |
| *MAPK8IP1* | 5.110±0.571 | 6.363±0.695 | >0.9999 | 4.972±0.718 | 0.3834 | *MAPK8IP1* | -0.3274 | 0.0401 |
| *WASF3* | 0.561±0.114 | 0.637±0.045 | 0.7265 | 0.732±0.100 | 0.5580 | *WASF3* | -0.3311 | 0.0292 |
| *WNT11* | 2.974±0.510 | 2.517±0.210 | 0.6392 | 2.687±0.333 | 0.6072 | *WNT11* | -0.3369 | 0.0250 |
| *FSCN1* | 131.5±11.89 | 217.1±9.16*** | 0.0007 | 198.7±12.56** | 0.0049 | *FSCN1* | -0.3375 | 0.0017 |
| *ZC3H4* | *N.A.* | *N.A.* | | *N.A.* | | *ZC3H4* | -0.3511 | 0.0220 |
| *MARCKS* | 1.449±0.342 | 1.970±0.185 | 0.5649 | 1.731±0.181 | 0.8882 | *MARCKS* | -0.3648 | 0.0002 |
| *PNPLA2* | 1.347±0.284 | 2.062±0.269 | >0.9999 | 2.616±0.431 | >0.9999 | *PNPLA2* | -0.3744 | 0.0405 |

| | | | | | | | | |
|---|---|---|---|---|---|---|---|---|
| *REEP6* | *N.A.* | *N.A.* | | *N.A.* | | *REEP6* | -0.4093 | 0.0153 |
| *CCDC71L* | 5.008±0.629 | 13.85±1.530**** | 0.0002 | 12.53±1.053*** | 0.0003 | *CCDC71L* | -0.4101 | 0.0124 |
| *TMEM164* | *N.A.* | *N.A.* | | *N.A.* | | *TMEM164* | -0.4159 | 0.0038 |
| *ISM1* | 0.467±0.044 | 0.516±0.036 | >0.9999 | 0.617±0.064 | >0.9999 | *ISM1* | -0.4235 | 0.0222 |
| *ZNF324* | 1.171±0.227 | 1.550±0.154 | 0.5998 | 1.332±0.129 | >0.9999 | *ZNF324* | -0.4967 | 0.0124 |
| *PTPRU* | 0.668±0.101 | 0.564±0.038 | 0.4499 | 0.671±0.071 | >0.9999 | *PTPRU* | -0.5225 | 0.0124 |
| *KLF10* Probe 1 | 9.260±0.935 | 7.061±0.570* | 0.0432 | 6.566±0.891** | 0.0035 | *KLF10* | -0.5225 | 0.0476 |
| *KLF10* Probe 2 | 5.779±0.433 | 3.393±0.462*** | 0.0007 | 2.891±0.332**** | <0.0001 | | | |
| *IL17RD* | 0.574±0.104 | 0.523±0.045 | 0.9648 | 0.619±0.070 | >0.9999 | *IL17RD* | -0.6023 | 0.0124 |
| *SOBP* | 10.03±1.440 | 13.75±1.229 | 0.6458 | 10.39±0.884 | >0.9999 | *SOBP* | -0.6111 | 0.0323 |
| *TMEM67* | 0.460±0.048 | 0.585±0.052 | >0.9999 | 0.605±0.070 | >0.9999 | *TMEM67* | -0.6136 | 0.0186 |
| *ADRA2A* | 0.452±0.079 | 0.377±0.035 | 0.4655 | 0.420±0.061 | 0.7753 | *ADRA2A* | -0.8037 | 0.0335 |
| *DOP1B* | 0.816±0.147 | 0.902±0.129 | 0.8568 | 0.763±0.086 | >0.9999 | *DOP1B* | -0.9376 | 0.0292 |
