## Supplemental Table 4 for "Modulation of calcium signaling on demand to decipher the molecular mechanisms of primary aldosteronism"

**Supplementary Table 4. List of differentially expressed genes in H295R_S2 cells expressing the α7-5HT3 receptor in response to 8h treatment with 10-8M AngII**

| Ensembl Gene ID | Gene Symbol | Gene Name | Log2 Fold Change | pAdj |
|---|---|---|---|---|
| ENSG00000128564 | VGF | VGF nerve growth factor inducible [Source:HGNC Symbol;Acc:HGNC:12684] | 4.3740 | 6.98E-204 |
| ENSG00000107554 | DNMBP | dynamin binding protein [Source:HGNC Symbol;Acc:HGNC:30373] | 2.5415 | 5.95E-165 |
| ENSG00000058404 | CAMK2B | calcium/calmodulin dependent protein kinase II beta [Source:HGNC Symbol;Acc:HGNC:1461] | 3.9307 | 5.63E-150 |
| ENSG00000102531 | FNDC3A | fibronectin type III domain containing 3A [Source:HGNC Symbol;Acc:HGNC:20296] | 2.2942 | 5.51E-143 |
| ENSG00000101230 | ISM1 | isthmin 1 [Source:HGNC Symbol;Acc:HGNC:16213] | -3.8125 | 7.00E-126 |
| ENSG00000119927 | GPAM | glycerol-3-phosphate acyltransferase, mitochondrial [Source:HGNC Symbol;Acc:HGNC:24865] | -2.7262 | 3.08E-119 |
| ENSG00000058866 | DGKG | diacylglycerol kinase gamma [Source:HGNC Symbol;Acc:HGNC:2853] | 2.1636 | 7.73E-110 |
| ENSG00000164211 | STARD4 | StAR related lipid transfer domain containing 4 [Source:HGNC Symbol;Acc:HGNC:18058] | 2.2001 | 9.79E-110 |
| ENSG00000170500 | LONRF2 | LON peptidase N-terminal domain and ring finger 2 [Source:HGNC Symbol;Acc:HGNC:24788] | 2.9559 | 3.65E-104 |
| ENSG00000116962 | NID1 | nidogen 1 [Source:HGNC Symbol;Acc:HGNC:7821] | 6.0922 | 7.19E-100 |
| ENSG00000139197 | PEX5 | peroxisomal biogenesis factor 5 [Source:HGNC Symbol;Acc:HGNC:9719] | 2.9380 | 5.33E-98 |
| ENSG00000006453 | BAIAP2L1 | BAR/IMD domain containing adaptor protein 2 like 1 [Source:HGNC Symbol;Acc:HGNC:21649] | 1.3201 | 1.03E-97 |
| ENSG00000143384 | MCL1 | MCL1 apoptosis regulator, BCL2 family member [Source:HGNC Symbol;Acc:HGNC:6943] | 1.8395 | 1.84E-91 |
| ENSG00000138166 | DUSP5 | dual specificity phosphatase 5 [Source:HGNC Symbol;Acc:HGNC:3071] | 3.1790 | 7.86E-91 |
| ENSG00000136560 | TANK | TRAF family member associated NFKB activator [Source:HGNC Symbol;Acc:HGNC:11562] | 1.7941 | 1.65E-87 |
| ENSG00000006327 | TNFRSF12A | TNF receptor superfamily member 12A [Source:HGNC Symbol;Acc:HGNC:18152] | 5.5144 | 1.40E-86 |
| ENSG00000147162 | OGT | O-linked N-acetylglucosamine (GlcNAc) transferase [Source:HGNC Symbol;Acc:HGNC:8127] | 2.9981 | 1.31E-84 |
| ENSG00000087263 | OGFOD1 | 2-oxoglutarate and iron dependent oxygenase domain containing 1 [Source:HGNC Symbol;Acc:HGNC:25585] | 1.0437 | 3.32E-83 |
| ENSG00000116717 | GADD45A | growth arrest and DNA damage inducible alpha [Source:HGNC Symbol;Acc:HGNC:4095] | 3.0865 | 1.54E-80 |
| ENSG00000154330 | PGM5 | phosphoglucomutase 5 [Source:HGNC Symbol;Acc:HGNC:8908] | -1.8315 | 2.46E-80 |
| ENSG00000198785 | GRIN3A | glutamate ionotropic receptor NMDA type subunit 3A [Source:HGNC Symbol;Acc:HGNC:16767] | -2.8579 | 4.25E-80 |
| ENSG00000090530 | P3H2 | prolyl 3-hydroxylase 2 [Source:HGNC Symbol;Acc:HGNC:19317] | 2.2871 | 1.57E-78 |
| ENSG00000147174 | GCNA | germ cell nuclear acidic peptidase [Source:HGNC Symbol;Acc:HGNC:15805] | 7.6138 | 1.92E-76 |
| ENSG00000161640 | SIGLEC11 | sialic acid binding Ig like lectin 11 [Source:HGNC Symbol;Acc:HGNC:15622] | -2.8572 | 3.84E-76 |
| ENSG00000168917 | SLC35G2 | solute carrier family 35 member G2 [Source:HGNC Symbol;Acc:HGNC:28480] | 1.4082 | 7.33E-75 |
| ENSG00000100644 | HIF1A | hypoxia inducible factor 1 subunit alpha [Source:HGNC Symbol;Acc:HGNC:4910] | 1.4397 | 1.81E-74 |
| ENSG00000133800 | LYVE1 | lymphatic vessel endothelial hyaluronan receptor 1 [Source:HGNC Symbol;Acc:HGNC:14687] | 4.0074 | 1.60E-73 |
| ENSG00000151835 | SACS | sacsin molecular chaperone [Source:HGNC Symbol;Acc:HGNC:10519] | 1.8012 | 3.09E-73 |
| ENSG00000181026 | AEN | apoptosis enhancing nuclease [Source:HGNC Symbol;Acc:HGNC:25722] | 1.8257 | 1.62E-70 |
| ENSG00000125148 | MT2A | metallothionein 2A [Source:HGNC Symbol;Acc:HGNC:7406] | 2.6540 | 8.53E-70 |
| ENSG00000176697 | BDNF | brain derived neurotrophic factor [Source:HGNC Symbol;Acc:HGNC:1033] | 3.7633 | 8.53E-70 |
| ENSG00000186350 | RXRA | retinoid X receptor alpha [Source:HGNC Symbol;Acc:HGNC:10477] | -1.5227 | 5.13E-69 |
| ENSG00000157851 | DPYSL5 | dihydropyrimidinase like 5 [Source:HGNC Symbol;Acc:HGNC:20637] | -1.4890 | 9.86E-69 |
| ENSG00000124762 | CDKN1A | cyclin dependent kinase inhibitor 1A [Source:HGNC Symbol;Acc:HGNC:1784] | 2.1301 | 1.00E-68 |
| ENSG00000164949 | GEM | GTP binding protein overexpressed in skeletal muscle [Source:HGNC Symbol;Acc:HGNC:4234] | 3.3342 | 1.68E-67 |
| ENSG00000196352 | CD55 | CD55 molecule (Cromer blood group) [Source:HGNC Symbol;Acc:HGNC:2665] | 2.1100 | 4.51E-67 |
| ENSG00000185432 | METTL7A | methyltransferase like 7A [Source:HGNC Symbol;Acc:HGNC:24550] | -1.4488 | 8.71E-67 |
| ENSG00000126351 | THRA | thyroid hormone receptor alpha [Source:HGNC Symbol;Acc:HGNC:11796] | -1.4072 | 5.65E-66 |
| ENSG00000150782 | IL18 | interleukin 18 [Source:HGNC Symbol;Acc:HGNC:5986] | 2.3035 | 3.76E-65 |
| ENSG00000139182 | CLSTN3 | calsyntenin 3 [Source:HGNC Symbol;Acc:HGNC:18371] | 2.5791 | 1.24E-64 |
| ENSG00000144655 | CSRNP1 | cysteine and serine rich nuclear protein 1 [Source:HGNC Symbol;Acc:HGNC:14300] | 2.9486 | 1.99E-63 |

| | | | | |
|---|---|---|---|---|
| ENSG00000166886 | NAB2 | NGFI-A binding protein 2 [Source:HGNC Symbol;Acc:HGNC:7627] | 3.5509 | 4.90E-63 |
| ENSG00000117899 | MESD | mesoderm development LRP chaperone [Source:HGNC Symbol;Acc:HGNC:13520] | 0.9552 | 6.20E-63 |
| ENSG00000112245 | PTP4A1 | protein tyrosine phosphatase 4A1 [Source:HGNC Symbol;Acc:HGNC:9634] | 1.9797 | 1.56E-62 |
| ENSG00000215218 | UBE2QL1 | ubiquitin conjugating enzyme E2 Q family like 1 [Source:HGNC Symbol;Acc:HGNC:37269] | 2.5109 | 1.77E-62 |
| ENSG00000173221 | GLRX | glutaredoxin [Source:HGNC Symbol;Acc:HGNC:4330] | 3.3173 | 2.06E-62 |
| ENSG00000111424 | VDR | vitamin D receptor [Source:HGNC Symbol;Acc:HGNC:12679] | 2.3888 | 7.14E-62 |
| ENSG00000154856 | APCDD1 | APC down-regulated 1 [Source:HGNC Symbol;Acc:HGNC:15718] | -2.0238 | 7.40E-62 |
| ENSG00000134686 | PHC2 | polyhomeotic homolog 2 [Source:HGNC Symbol;Acc:HGNC:3183] | 1.5549 | 1.42E-61 |
| ENSG00000233581 | NA | novel transcript | 4.8513 | 1.62E-61 |
| ENSG00000137713 | PPP2R1B | protein phosphatase 2 scaffold subunit Abeta [Source:HGNC Symbol;Acc:HGNC:9303] | 2.5386 | 8.36E-61 |
| ENSG00000168477 | TNXB | tenascin XB [Source:HGNC Symbol;Acc:HGNC:11976] | 3.1029 | 5.99E-60 |
| ENSG00000116514 | RNF19B | ring finger protein 19B [Source:HGNC Symbol;Acc:HGNC:26886] | 2.4814 | 2.03E-59 |
| ENSG00000166025 | AMOTL1 | angiomotin like 1 [Source:HGNC Symbol;Acc:HGNC:17811] | 1.1400 | 2.81E-59 |
| ENSG00000170873 | MTSS1 | MTSS I-BAR domain containing 1 [Source:HGNC Symbol;Acc:HGNC:20443] | 1.4846 | 4.31E-59 |
| ENSG00000176406 | RIMS2 | regulating synaptic membrane exocytosis 2 [Source:HGNC Symbol;Acc:HGNC:17283] | 1.4510 | 8.41E-59 |
| ENSG00000228536 | LYPLAL1-AS1 | LYPLAL1 antisense RNA 1 [Source:HGNC Symbol;Acc:HGNC:54054] | 2.8850 | 4.20E-58 |
| ENSG00000101421 | CHMP4B | charged multivesicular body protein 4B [Source:HGNC Symbol;Acc:HGNC:16171] | 1.1761 | 6.52E-57 |
| ENSG00000108984 | MAP2K6 | mitogen-activated protein kinase kinase 6 [Source:HGNC Symbol;Acc:HGNC:6846] | -3.3954 | 1.89E-56 |
| ENSG00000168243 | GNG4 | G protein subunit gamma 4 [Source:HGNC Symbol;Acc:HGNC:4407] | 3.5678 | 3.36E-56 |
| ENSG00000139289 | PHLDA1 | pleckstrin homology like domain family A member 1 [Source:HGNC Symbol;Acc:HGNC:8933] | 1.6568 | 4.33E-56 |
| ENSG00000197170 | PSMD12 | proteasome 26S subunit, non-ATPase 12 [Source:HGNC Symbol;Acc:HGNC:9557] | 1.0658 | 1.46E-55 |
| ENSG00000112419 | PHACTR2 | phosphatase and actin regulator 2 [Source:HGNC Symbol;Acc:HGNC:20956] | -1.2523 | 4.40E-55 |
| ENSG00000070061 | ELP1 | elongator complex protein 1 [Source:HGNC Symbol;Acc:HGNC:5959] | 2.0445 | 5.06E-55 |
| ENSG00000117586 | TNFSF4 | TNF superfamily member 4 [Source:HGNC Symbol;Acc:HGNC:11934] | 2.3537 | 1.99E-54 |
| ENSG00000138767 | CNOT6L | CCR4-NOT transcription complex subunit 6 like [Source:HGNC Symbol;Acc:HGNC:18042] | 1.5136 | 3.30E-54 |
| ENSG00000089639 | GMIP | GEM interacting protein [Source:HGNC Symbol;Acc:HGNC:24852] | 2.9162 | 3.55E-54 |
| ENSG00000075618 | FSCN1 | fascin actin-bundling protein 1 [Source:HGNC Symbol;Acc:HGNC:11148] | -0.8826 | 5.48E-54 |
| ENSG00000122012 | SV2C | synaptic vesicle glycoprotein 2C [Source:HGNC Symbol;Acc:HGNC:30670] | 2.5063 | 1.65E-52 |
| ENSG00000090776 | EFNB1 | ephrin B1 [Source:HGNC Symbol;Acc:HGNC:3226] | -1.9351 | 1.90E-52 |
| ENSG00000109065 | NAT9 | N-acetyltransferase 9 (putative) [Source:HGNC Symbol;Acc:HGNC:23133] | 1.2947 | 4.42E-52 |
| ENSG00000170915 | PAQR8 | progestin and adipoQ receptor family member 8 [Source:HGNC Symbol;Acc:HGNC:15708] | -1.9712 | 1.17E-51 |
| ENSG00000109654 | TRIM2 | tripartite motif containing 2 [Source:HGNC Symbol;Acc:HGNC:15974] | 1.8718 | 1.56E-51 |
| ENSG00000034152 | MAP2K3 | mitogen-activated protein kinase kinase 3 [Source:HGNC Symbol;Acc:HGNC:6843] | 2.5325 | 1.91E-51 |
| ENSG00000064547 | LPAR2 | lysophosphatidic acid receptor 2 [Source:HGNC Symbol;Acc:HGNC:3168] | 3.2309 | 6.53E-51 |
| ENSG00000160326 | SLC2A6 | solute carrier family 2 member 6 [Source:HGNC Symbol;Acc:HGNC:11011] | -1.4577 | 1.63E-50 |
| ENSG00000167693 | NXN | nucleoredoxin [Source:HGNC Symbol;Acc:HGNC:18008] | 3.1950 | 1.83E-50 |
| ENSG00000178947 | SMIM10L2A | small integral membrane protein 10 like 2A [Source:HGNC Symbol;Acc:HGNC:34499] | -1.2860 | 1.84E-50 |
| ENSG00000105717 | PBX4 | PBX homeobox 4 [Source:HGNC Symbol;Acc:HGNC:13403] | 3.4257 | 5.00E-50 |
| ENSG00000069020 | MAST4 | microtubule associated serine/threonine kinase family member 4 [Source:HGNC Symbol;Acc:HGNC:19037] | 1.9395 | 2.75E-49 |
| ENSG00000118257 | NRP2 | neuropilin 2 [Source:HGNC Symbol;Acc:HGNC:8005] | 2.5199 | 3.16E-49 |
| ENSG00000164080 | RAD54L2 | RAD54 like 2 [Source:HGNC Symbol;Acc:HGNC:29123] | 1.0419 | 4.66E-49 |
| ENSG00000213719 | CLIC1 | chloride intracellular channel 1 [Source:HGNC Symbol;Acc:HGNC:2062] | 1.1524 | 7.05E-49 |
| ENSG00000162889 | MAPKAPK2 | MAPK activated protein kinase 2 [Source:HGNC Symbol;Acc:HGNC:6887] | 1.1734 | 1.12E-48 |

| | | | | |
|---|---|---|---|---|
| ENSG00000060656 | PTPRU | protein tyrosine phosphatase receptor type U [Source:HGNC Symbol;Acc:HGNC:9683] | -1.7784 | 1.15E-48 |
| ENSG00000132970 | WASF3 | WASP family member 3 [Source:HGNC Symbol;Acc:HGNC:12734] | -1.4627 | 1.83E-48 |
| ENSG00000166401 | SERPINB8 | serpin family B member 8 [Source:HGNC Symbol;Acc:HGNC:8952] | 3.0762 | 2.34E-48 |
| ENSG00000116729 | WLS | Wnt ligand secretion mediator [Source:HGNC Symbol;Acc:HGNC:30238] | 1.5951 | 7.07E-48 |
| ENSG00000033627 | ATP6V0A1 | ATPase H+ transporting V0 subunit a1 [Source:HGNC Symbol;Acc:HGNC:865] | 1.2542 | 7.70E-48 |
| ENSG00000064651 | SLC12A2 | solute carrier family 12 member 2 [Source:HGNC Symbol;Acc:HGNC:10911] | 1.9431 | 1.13E-47 |
| ENSG00000169756 | LIMS1 | LIM zinc finger domain containing 1 [Source:HGNC Symbol;Acc:HGNC:6616] | 1.4225 | 2.45E-47 |
| ENSG00000119326 | CTNNAL1 | catenin alpha like 1 [Source:HGNC Symbol;Acc:HGNC:2512] | 0.9981 | 4.76E-47 |
| ENSG00000020181 | ADGRA2 | adhesion G protein-coupled receptor A2 [Source:HGNC Symbol;Acc:HGNC:17849] | -2.1377 | 5.14E-47 |
| ENSG00000090520 | DNAJB11 | DnaJ heat shock protein family (Hsp40) member B11 [Source:HGNC Symbol;Acc:HGNC:14889] | 2.0135 | 5.69E-47 |
| ENSG00000044574 | HSPA5 | heat shock protein family A (Hsp70) member 5 [Source:HGNC Symbol;Acc:HGNC:5238] | 2.5921 | 7.01E-47 |
| ENSG00000118503 | TNFAIP3 | TNF alpha induced protein 3 [Source:HGNC Symbol;Acc:HGNC:11896] | 2.1259 | 7.06E-47 |
| ENSG00000101986 | ABCD1 | ATP binding cassette subfamily D member 1 [Source:HGNC Symbol;Acc:HGNC:61] | 1.3610 | 1.67E-46 |
| ENSG00000148841 | ITPRIP | inositol 1,4,5-trisphosphate receptor interacting protein [Source:HGNC Symbol;Acc:HGNC:29370] | 2.1813 | 1.71E-46 |
| ENSG00000164741 | DLC1 | DLC1 Rho GTPase activating protein [Source:HGNC Symbol;Acc:HGNC:2897] | 1.6656 | 2.45E-46 |
| ENSG00000137714 | FDX1 | ferredoxin 1 [Source:HGNC Symbol;Acc:HGNC:3638] | 2.1564 | 3.19E-46 |
| ENSG00000171724 | VAT1L | vesicle amine transport 1 like [Source:HGNC Symbol;Acc:HGNC:29315] | 2.9539 | 5.09E-46 |
| ENSG00000145623 | OSMR | oncostatin M receptor [Source:HGNC Symbol;Acc:HGNC:8507] | 2.1074 | 9.89E-46 |
| ENSG00000010818 | HIVEP2 | HIVEP zinc finger 2 [Source:HGNC Symbol;Acc:HGNC:4921] | 1.8456 | 1.05E-45 |
| ENSG00000176595 | KBTBD11 | kelch repeat and BTB domain containing 11 [Source:HGNC Symbol;Acc:HGNC:29104] | 1.6719 | 1.49E-45 |
| ENSG00000273749 | CYFIP1 | cytoplasmic FMR1 interacting protein 1 [Source:HGNC Symbol;Acc:HGNC:13759] | 0.7506 | 1.63E-45 |
| ENSG00000120875 | DUSP4 | dual specificity phosphatase 4 [Source:HGNC Symbol;Acc:HGNC:3070] | 2.8263 | 1.64E-45 |
| ENSG00000104885 | DOT1L | DOT1 like histone lysine methyltransferase [Source:HGNC Symbol;Acc:HGNC:24948] | 1.6954 | 2.24E-45 |
| ENSG00000104973 | MED25 | mediator complex subunit 25 [Source:HGNC Symbol;Acc:HGNC:28845] | -0.9734 | 2.93E-45 |
| ENSG00000166598 | HSP90B1 | heat shock protein 90 beta family member 1 [Source:HGNC Symbol;Acc:HGNC:12028] | 1.4646 | 8.73E-45 |
| ENSG00000152684 | PELO | pelota mRNA surveillance and ribosome rescue factor [Source:HGNC Symbol;Acc:HGNC:8829] | 1.6129 | 1.03E-44 |
| ENSG00000006468 | ETV1 | ETS variant transcription factor 1 [Source:HGNC Symbol;Acc:HGNC:3490] | 1.5992 | 1.44E-44 |
| ENSG00000159840 | ZYX | zyxin [Source:HGNC Symbol;Acc:HGNC:13200] | 1.4089 | 2.01E-44 |
| ENSG00000196782 | MAML3 | mastermind like transcriptional coactivator 3 [Source:HGNC Symbol;Acc:HGNC:16272] | 1.2776 | 2.68E-44 |
| ENSG00000134775 | FHOD3 | formin homology 2 domain containing 3 [Source:HGNC Symbol;Acc:HGNC:26178] | -1.7670 | 1.19E-43 |
| ENSG00000166780 | BMERB1 | bMERB domain containing 1 [Source:HGNC Symbol;Acc:HGNC:19213] | 1.9034 | 1.29E-43 |
| ENSG00000204054 | LINC00963 | long intergenic non-protein coding RNA 963 [Source:HGNC Symbol;Acc:HGNC:48716] | 1.1271 | 4.07E-43 |
| ENSG00000169372 | CRADD | CASP2 and RIPK1 domain containing adaptor with death domain [Source:HGNC Symbol;Acc:HGNC:2340] | 2.2609 | 6.46E-43 |
| ENSG00000006210 | CX3CL1 | C-X3-C motif chemokine ligand 1 [Source:HGNC Symbol;Acc:HGNC:10647] | 1.5374 | 1.49E-42 |
| ENSG00000133083 | DCLK1 | doublecortin like kinase 1 [Source:HGNC Symbol;Acc:HGNC:2700] | -1.4458 | 1.72E-42 |
| ENSG00000175093 | SPSB4 | splA/ryanodine receptor domain and SOCS box containing 4 [Source:HGNC Symbol;Acc:HGNC:30630] | -3.0704 | 5.30E-42 |
| ENSG00000198363 | ASPH | aspartate beta-hydroxylase [Source:HGNC Symbol;Acc:HGNC:757] | 1.4247 | 7.56E-42 |
| ENSG00000110841 | PPFIBP1 | PPFIA binding protein 1 [Source:HGNC Symbol;Acc:HGNC:9249] | 1.7103 | 1.35E-41 |
| ENSG00000072422 | RHOBTB1 | Rho related BTB domain containing 1 [Source:HGNC Symbol;Acc:HGNC:18738] | -1.5755 | 1.52E-41 |
| ENSG00000087074 | PPP1R15A | protein phosphatase 1 regulatory subunit 15A [Source:HGNC Symbol;Acc:HGNC:14375] | 1.5034 | 2.46E-41 |
| ENSG00000105825 | TFPI2 | tissue factor pathway inhibitor 2 [Source:HGNC Symbol;Acc:HGNC:11761] | 3.8052 | 2.85E-41 |
| ENSG00000124374 | PAIP2B | poly(A) binding protein interacting protein 2B [Source:HGNC Symbol;Acc:HGNC:29200] | -1.9375 | 2.88E-41 |

| | | | | |
|---|---|---|---|---|
| ENSG00000174099 | MSRB3 | methionine sulfoxide reductase B3 [Source:HGNC Symbol;Acc:HGNC:27375] | 0.7598 | 1.33E-40 |
| ENSG00000167695 | TLCD3A | TLC domain containing 3A [Source:HGNC Symbol;Acc:HGNC:29646] | 3.0266 | 1.82E-40 |
| ENSG00000076770 | MBNL3 | muscleblind like splicing regulator 3 [Source:HGNC Symbol;Acc:HGNC:20564] | -1.3119 | 3.57E-40 |
| ENSG00000196814 | MVB12B | multivesicular body subunit 12B [Source:HGNC Symbol;Acc:HGNC:23368] | -1.4968 | 3.65E-40 |
| ENSG00000067057 | PFKP | phosphofructokinase, platelet [Source:HGNC Symbol;Acc:HGNC:8878] | 2.4064 | 4.34E-40 |
| ENSG00000143067 | ZNF697 | zinc finger protein 697 [Source:HGNC Symbol;Acc:HGNC:32034] | 2.3045 | 5.74E-40 |
| ENSG00000184840 | TMED9 | transmembrane p24 trafficking protein 9 [Source:HGNC Symbol;Acc:HGNC:24878] | 0.8235 | 7.85E-40 |
| ENSG00000152661 | GJA1 | gap junction protein alpha 1 [Source:HGNC Symbol;Acc:HGNC:4274] | 2.1377 | 8.69E-40 |
| ENSG00000153208 | MERTK | MER proto-oncogene, tyrosine kinase [Source:HGNC Symbol;Acc:HGNC:7027] | -1.4543 | 1.97E-39 |
| ENSG00000272168 | CASC15 | cancer susceptibility 15 [Source:HGNC Symbol;Acc:HGNC:28245] | 2.4063 | 2.00E-39 |
| ENSG00000057019 | DCBLD2 | discoidin, CUB and LCCL domain containing 2 [Source:HGNC Symbol;Acc:HGNC:24627] | 1.7726 | 2.19E-39 |
| ENSG00000172965 | MIR4435-2HG | MIR4435-2 host gene [Source:HGNC Symbol;Acc:HGNC:35163] | 1.8139 | 3.26E-39 |
| ENSG00000186806 | VSIG10L | V-set and immunoglobulin domain containing 10 like [Source:HGNC Symbol;Acc:HGNC:27111] | -1.9019 | 3.87E-39 |
| ENSG00000158186 | MRAS | muscle RAS oncogene homolog [Source:HGNC Symbol;Acc:HGNC:7227] | 1.5206 | 6.06E-39 |
| ENSG00000077150 | NFKB2 | nuclear factor kappa B subunit 2 [Source:HGNC Symbol;Acc:HGNC:7795] | 1.4705 | 7.20E-39 |
| ENSG00000136044 | APPL2 | adaptor protein, phosphotyrosine interacting with PH domain and leucine zipper 2 [Source:HGNC Symbol;Acc:HGNC:18242] | 1.3408 | 8.16E-39 |
| ENSG00000138134 | STAMBPL1 | STAM binding protein like 1 [Source:HGNC Symbol;Acc:HGNC:24105] | 1.9658 | 9.96E-39 |
| ENSG00000204219 | TCEA3 | transcription elongation factor A3 [Source:HGNC Symbol;Acc:HGNC:11615] | -0.7929 | 1.53E-38 |
| ENSG00000137727 | ARHGAP20 | Rho GTPase activating protein 20 [Source:HGNC Symbol;Acc:HGNC:18357] | 2.2982 | 2.43E-38 |
| ENSG00000172159 | FRMD3 | FERM domain containing 3 [Source:HGNC Symbol;Acc:HGNC:24125] | 2.2617 | 2.43E-38 |
| ENSG00000082482 | KCNK2 | potassium two pore domain channel subfamily K member 2 [Source:HGNC Symbol;Acc:HGNC:6277] | -1.3238 | 2.76E-38 |
| ENSG00000145703 | IQGAP2 | IQ motif containing GTPase activating protein 2 [Source:HGNC Symbol;Acc:HGNC:6111] | -1.8179 | 2.83E-38 |
| ENSG00000055118 | KCNH2 | potassium voltage-gated channel subfamily H member 2 [Source:HGNC Symbol;Acc:HGNC:6251] | -1.9753 | 4.71E-38 |
| ENSG00000071127 | WDR1 | WD repeat domain 1 [Source:HGNC Symbol;Acc:HGNC:12754] | 1.2951 | 5.97E-38 |
| ENSG00000114698 | PLSCR4 | phospholipid scramblase 4 [Source:HGNC Symbol;Acc:HGNC:16497] | -1.3381 | 6.81E-38 |
| ENSG00000064225 | ST3GAL6 | ST3 beta-galactoside alpha-2,3-sialyltransferase 6 [Source:HGNC Symbol;Acc:HGNC:18080] | -1.6847 | 8.28E-38 |
| ENSG00000179604 | CDC42EP4 | CDC42 effector protein 4 [Source:HGNC Symbol;Acc:HGNC:17147] | -1.3774 | 9.27E-38 |
| ENSG00000132963 | POMP | proteasome maturation protein [Source:HGNC Symbol;Acc:HGNC:20330] | 0.9665 | 9.56E-38 |
| ENSG00000166557 | TMED3 | transmembrane p24 trafficking protein 3 [Source:HGNC Symbol;Acc:HGNC:28889] | 1.5516 | 1.04E-37 |
| ENSG00000095303 | PTGS1 | prostaglandin-endoperoxide synthase 1 [Source:HGNC Symbol;Acc:HGNC:9604] | 3.2442 | 1.77E-37 |
| ENSG00000155252 | PI4K2A | phosphatidylinositol 4-kinase type 2 alpha [Source:HGNC Symbol;Acc:HGNC:30031] | 0.9740 | 2.51E-37 |
| ENSG00000140105 | WARS1 | tryptophanyl-tRNA synthetase 1 [Source:HGNC Symbol;Acc:HGNC:12729] | 1.2283 | 3.06E-37 |
| ENSG00000143845 | ETNK2 | ethanolamine kinase 2 [Source:HGNC Symbol;Acc:HGNC:25575] | -1.8423 | 3.26E-37 |
| ENSG00000204217 | BMPR2 | bone morphogenetic protein receptor type 2 [Source:HGNC Symbol;Acc:HGNC:1078] | 1.8541 | 3.44E-37 |
| ENSG00000184349 | EFNA5 | ephrin A5 [Source:HGNC Symbol;Acc:HGNC:3225] | 2.9252 | 3.50E-37 |
| ENSG00000222041 | CYTOR | cytoskeleton regulator RNA [Source:HGNC Symbol;Acc:HGNC:28717] | 1.9877 | 4.59E-37 |
| ENSG00000244405 | ETV5 | ETS variant transcription factor 5 [Source:HGNC Symbol;Acc:HGNC:3494] | 3.2996 | 4.96E-37 |
| ENSG00000135750 | KCNK1 | potassium two pore domain channel subfamily K member 1 [Source:HGNC Symbol;Acc:HGNC:6272] | 1.5390 | 7.08E-37 |
| ENSG00000261037 | NA | novel transcript | 3.0566 | 7.57E-37 |
| ENSG00000106789 | CORO2A | coronin 2A [Source:HGNC Symbol;Acc:HGNC:2255] | -1.9171 | 1.20E-36 |
| ENSG00000196923 | PDLIM7 | PDZ and LIM domain 7 [Source:HGNC Symbol;Acc:HGNC:22958] | 1.4472 | 1.34E-36 |
| ENSG00000140406 | TLNRD1 | talin rod domain containing 1 [Source:HGNC Symbol;Acc:HGNC:13519] | 2.1118 | 1.36E-36 |

| | | | | |
|---|---|---|---|---|
| ENSG00000185630 | PBX1 | PBX homeobox 1 [Source:HGNC Symbol;Acc:HGNC:8632] | 1.4164 | 1.59E-36 |
| ENSG00000164442 | CITED2 | Cbp/p300 interacting transactivator with Glu/Asp rich carboxy-terminal domain 2 [Source:HGNC Symbol;Acc:HGNC:1987] | 2.0335 | 1.85E-36 |
| ENSG00000150893 | FREM2 | FRAS1 related extracellular matrix 2 [Source:HGNC Symbol;Acc:HGNC:25396] | 1.8577 | 3.24E-36 |
| ENSG00000119408 | NEK6 | NIMA related kinase 6 [Source:HGNC Symbol;Acc:HGNC:7749] | 2.4421 | 4.13E-36 |
| ENSG00000175344 | CHRNA7 | cholinergic receptor nicotinic alpha 7 subunit [Source:HGNC Symbol;Acc:HGNC:1960] | 2.7347 | 5.86E-36 |
| ENSG00000160161 | CILP2 | cartilage intermediate layer protein 2 [Source:HGNC Symbol;Acc:HGNC:24213] | 2.2841 | 6.08E-36 |
| ENSG00000120686 | UFM1 | ubiquitin fold modifier 1 [Source:HGNC Symbol;Acc:HGNC:20597] | 1.0157 | 8.46E-36 |
| ENSG00000122641 | INHBA | inhibin subunit beta A [Source:HGNC Symbol;Acc:HGNC:6066] | 6.2806 | 8.67E-36 |
| ENSG00000165997 | ARL5B | ADP ribosylation factor like GTPase 5B [Source:HGNC Symbol;Acc:HGNC:23052] | 1.4690 | 2.60E-35 |
| ENSG00000198597 | ZNF536 | zinc finger protein 536 [Source:HGNC Symbol;Acc:HGNC:29025] | 1.5940 | 2.85E-35 |
| ENSG00000005249 | PRKAR2B | protein kinase cAMP-dependent type II regulatory subunit beta [Source:HGNC Symbol;Acc:HGNC:9392] | 1.6934 | 2.96E-35 |
| ENSG00000070159 | PTPN3 | protein tyrosine phosphatase non-receptor type 3 [Source:HGNC Symbol;Acc:HGNC:9655] | 1.9655 | 3.00E-35 |
| ENSG00000167528 | ZNF641 | zinc finger protein 641 [Source:HGNC Symbol;Acc:HGNC:31834] | 1.9881 | 3.12E-35 |
| ENSG00000087250 | MT3 | metallothionein 3 [Source:HGNC Symbol;Acc:HGNC:7408] | 1.3123 | 4.73E-35 |
| ENSG00000074842 | MYDGF | myeloid derived growth factor [Source:HGNC Symbol;Acc:HGNC:16948] | 1.0193 | 5.73E-35 |
| ENSG00000127824 | TUBA4A | tubulin alpha 4a [Source:HGNC Symbol;Acc:HGNC:12407] | 2.5424 | 6.07E-35 |
| ENSG00000138449 | SLC40A1 | solute carrier family 40 member 1 [Source:HGNC Symbol;Acc:HGNC:10909] | -1.9611 | 9.60E-35 |
| ENSG00000110921 | MVK | mevalonate kinase [Source:HGNC Symbol;Acc:HGNC:7530] | -1.2918 | 9.65E-35 |
| ENSG00000168280 | KIF5C | kinesin family member 5C [Source:HGNC Symbol;Acc:HGNC:6325] | 1.7610 | 1.47E-34 |
| ENSG00000228594 | FNDC10 | fibronectin type III domain containing 10 [Source:HGNC Symbol;Acc:HGNC:42951] | -1.1865 | 1.94E-34 |
| ENSG00000135919 | SERPINE2 | serpin family E member 2 [Source:HGNC Symbol;Acc:HGNC:8951] | 2.3655 | 2.05E-34 |
| ENSG00000123933 | MXD4 | MAX dimerization protein 4 [Source:HGNC Symbol;Acc:HGNC:13906] | -1.4029 | 2.26E-34 |
| ENSG00000154217 | PITPNC1 | phosphatidylinositol transfer protein cytoplasmic 1 [Source:HGNC Symbol;Acc:HGNC:21045] | 2.1500 | 2.88E-34 |
| ENSG00000181773 | GPR3 | G protein-coupled receptor 3 [Source:HGNC Symbol;Acc:HGNC:4484] | 4.6403 | 6.73E-34 |
| ENSG00000185924 | RTN4RL1 | reticulon 4 receptor like 1 [Source:HGNC Symbol;Acc:HGNC:21329] | -1.7000 | 6.98E-34 |
| ENSG00000162073 | PAQR4 | progestin and adipoQ receptor family member 4 [Source:HGNC Symbol;Acc:HGNC:26386] | -1.4740 | 7.03E-34 |
| ENSG00000135241 | PNPLA8 | patatin like phospholipase domain containing 8 [Source:HGNC Symbol;Acc:HGNC:28900] | 1.1897 | 8.55E-34 |
| ENSG00000117016 | RIMS3 | regulating synaptic membrane exocytosis 3 [Source:HGNC Symbol;Acc:HGNC:21292] | -1.9394 | 8.72E-34 |
| ENSG00000107099 | DOCK8 | dedicator of cytokinesis 8 [Source:HGNC Symbol;Acc:HGNC:19191] | 1.2873 | 1.77E-33 |
| ENSG00000198624 | CCDC69 | coiled-coil domain containing 69 [Source:HGNC Symbol;Acc:HGNC:24487] | -1.0359 | 2.40E-33 |
| ENSG00000114480 | GBE1 | 1,4-alpha-glucan branching enzyme 1 [Source:HGNC Symbol;Acc:HGNC:4180] | 1.4375 | 2.47E-33 |
| ENSG00000250986 | LINC02600 | long intergenic non-protein coding RNA 2600 [Source:HGNC Symbol;Acc:HGNC:53177] | -2.5154 | 2.94E-33 |
| ENSG00000161681 | SHANK1 | SH3 and multiple ankyrin repeat domains 1 [Source:HGNC Symbol;Acc:HGNC:15474] | -1.2464 | 3.08E-33 |
| ENSG00000149428 | HYOU1 | hypoxia up-regulated 1 [Source:HGNC Symbol;Acc:HGNC:16931] | 1.7400 | 3.52E-33 |
| ENSG00000118939 | UCHL3 | ubiquitin C-terminal hydrolase L3 [Source:HGNC Symbol;Acc:HGNC:12515] | 1.1412 | 4.70E-33 |
| ENSG00000145050 | MANF | mesencephalic astrocyte derived neurotrophic factor [Source:HGNC Symbol;Acc:HGNC:15461] | 1.9519 | 5.78E-33 |
| ENSG00000095794 | CREM | cAMP responsive element modulator [Source:HGNC Symbol;Acc:HGNC:2352] | 1.4816 | 5.82E-33 |
| ENSG00000216775 | NA | heterogeneous nuclear ribonucleoprotein A/B (HNRNPAB) pseudogene | -1.8983 | 6.55E-33 |
| ENSG00000162437 | RAVER2 | ribonucleoprotein, PTB binding 2 [Source:HGNC Symbol;Acc:HGNC:25577] | -0.7264 | 1.01E-32 |
| ENSG00000182534 | MXRA7 | matrix remodeling associated 7 [Source:HGNC Symbol;Acc:HGNC:7541] | 1.4840 | 1.13E-32 |
| ENSG00000068878 | PSME4 | proteasome activator subunit 4 [Source:HGNC Symbol;Acc:HGNC:20635] | 0.9329 | 1.22E-32 |
| ENSG00000151689 | INPP1 | inositol polyphosphate-1-phosphatase [Source:HGNC Symbol;Acc:HGNC:6071] | 2.1250 | 1.41E-32 |

| | | | | |
|---|---|---|---|---|
| ENSG00000107175 | CREB3 | cAMP responsive element binding protein 3 [Source:HGNC Symbol;Acc:HGNC:2347] | 0.6825 | 1.45E-32 |
| ENSG00000101109 | STK4 | serine/threonine kinase 4 [Source:HGNC Symbol;Acc:HGNC:11408] | 0.8971 | 1.70E-32 |
| ENSG00000221968 | FADS3 | fatty acid desaturase 3 [Source:HGNC Symbol;Acc:HGNC:3576] | 1.4311 | 1.87E-32 |
| ENSG00000280061 | NA | TEC | 1.7447 | 1.90E-32 |
| ENSG00000153930 | ANKFN1 | ankyrin repeat and fibronectin type III domain containing 1 [Source:HGNC Symbol;Acc:HGNC:26766] | -2.3110 | 1.93E-32 |
| ENSG00000060709 | RIMBP2 | RIMS binding protein 2 [Source:HGNC Symbol;Acc:HGNC:30339] | 2.8234 | 2.22E-32 |
| ENSG00000152620 | NADK2 | NAD kinase 2, mitochondrial [Source:HGNC Symbol;Acc:HGNC:26404] | -1.2296 | 2.38E-32 |
| ENSG00000173599 | PC | pyruvate carboxylase [Source:HGNC Symbol;Acc:HGNC:8636] | 1.0536 | 2.76E-32 |
| ENSG00000176658 | MYO1D | myosin ID [Source:HGNC Symbol;Acc:HGNC:7598] | 1.4100 | 3.22E-32 |
| ENSG00000156471 | PTDSS1 | phosphatidylserine synthase 1 [Source:HGNC Symbol;Acc:HGNC:9587] | 0.8051 | 3.41E-32 |
| ENSG00000185070 | FLRT2 | fibronectin leucine rich transmembrane protein 2 [Source:HGNC Symbol;Acc:HGNC:3761] | 1.6577 | 4.41E-32 |
| ENSG00000215301 | DDX3X | DEAD-box helicase 3 X-linked [Source:HGNC Symbol;Acc:HGNC:2745] | 0.7589 | 5.13E-32 |
| ENSG00000033100 | CHPF2 | chondroitin polymerizing factor 2 [Source:HGNC Symbol;Acc:HGNC:29270] | 1.3028 | 7.44E-32 |
| ENSG00000118680 | MYL12B | myosin light chain 12B [Source:HGNC Symbol;Acc:HGNC:29827] | 1.2561 | 7.75E-32 |
| ENSG00000026103 | FAS | Fas cell surface death receptor [Source:HGNC Symbol;Acc:HGNC:11920] | 1.7455 | 1.22E-31 |
| ENSG00000137309 | HMGA1 | high mobility group AT-hook 1 [Source:HGNC Symbol;Acc:HGNC:5010] | 2.3128 | 1.22E-31 |
| ENSG00000120696 | KBTBD7 | kelch repeat and BTB domain containing 7 [Source:HGNC Symbol;Acc:HGNC:25266] | -1.1132 | 1.30E-31 |
| ENSG00000023171 | GRAMD1B | GRAM domain containing 1B [Source:HGNC Symbol;Acc:HGNC:29214] | 1.8018 | 1.43E-31 |
| ENSG00000114850 | SSR3 | signal sequence receptor subunit 3 [Source:HGNC Symbol;Acc:HGNC:11325] | 1.1117 | 1.47E-31 |
| ENSG00000143590 | EFNA3 | ephrin A3 [Source:HGNC Symbol;Acc:HGNC:3223] | -1.9600 | 1.57E-31 |
| ENSG00000111057 | KRT18 | keratin 18 [Source:HGNC Symbol;Acc:HGNC:6430] | 1.2369 | 1.79E-31 |
| ENSG00000162407 | PLPP3 | phospholipid phosphatase 3 [Source:HGNC Symbol;Acc:HGNC:9229] | 1.9357 | 1.98E-31 |
| ENSG00000133872 | SARAF | store-operated calcium entry associated regulatory factor [Source:HGNC Symbol;Acc:HGNC:28789] | 0.8146 | 2.04E-31 |
| ENSG00000023318 | ERP44 | endoplasmic reticulum protein 44 [Source:HGNC Symbol;Acc:HGNC:18311] | 0.8977 | 2.12E-31 |
| ENSG00000181291 | TMEM132E | transmembrane protein 132E [Source:HGNC Symbol;Acc:HGNC:26991] | 1.5481 | 3.15E-31 |
| ENSG00000177370 | TIMM22 | translocase of inner mitochondrial membrane 22 [Source:HGNC Symbol;Acc:HGNC:17317] | 1.0203 | 3.27E-31 |
| ENSG00000108244 | KRT23 | keratin 23 [Source:HGNC Symbol;Acc:HGNC:6438] | 4.2224 | 3.71E-31 |
| ENSG00000248323 | LUCAT1 | lung cancer associated transcript 1 [Source:HGNC Symbol;Acc:HGNC:48498] | 1.4724 | 4.33E-31 |
| ENSG00000134769 | DTNA | dystrobrevin alpha [Source:HGNC Symbol;Acc:HGNC:3057] | -1.2310 | 6.11E-31 |
| ENSG00000078814 | MYH7B | myosin heavy chain 7B [Source:HGNC Symbol;Acc:HGNC:15906] | -2.1658 | 7.23E-31 |
| ENSG00000100219 | XBP1 | X-box binding protein 1 [Source:HGNC Symbol;Acc:HGNC:12801] | 1.7640 | 8.96E-31 |
| ENSG00000086619 | ERO1B | endoplasmic reticulum oxidoreductase 1 beta [Source:HGNC Symbol;Acc:HGNC:14355] | 2.6315 | 9.96E-31 |
| ENSG00000177971 | IMP3 | IMP U3 small nucleolar ribonucleoprotein 3 [Source:HGNC Symbol;Acc:HGNC:14497] | 0.7249 | 1.03E-30 |
| ENSG00000239264 | TXNDC5 | thioredoxin domain containing 5 [Source:HGNC Symbol;Acc:HGNC:21073] | 0.8391 | 1.48E-30 |
| ENSG00000167861 | HID1 | HID1 domain containing [Source:HGNC Symbol;Acc:HGNC:15736] | 1.9382 | 1.52E-30 |
| ENSG00000179862 | CITED4 | Cbp/p300 interacting transactivator with Glu/Asp rich carboxy-terminal domain 4 [Source:HGNC Symbol;Acc:HGNC:18696] | 2.0153 | 1.59E-30 |
| ENSG00000136928 | GABBR2 | gamma-aminobutyric acid type B receptor subunit 2 [Source:HGNC Symbol;Acc:HGNC:4507] | -1.2973 | 1.87E-30 |
| ENSG00000153214 | TMEM87B | transmembrane protein 87B [Source:HGNC Symbol;Acc:HGNC:25913] | 2.3025 | 2.02E-30 |
| ENSG00000162745 | OLFML2B | olfactomedin like 2B [Source:HGNC Symbol;Acc:HGNC:24558] | -3.5530 | 2.20E-30 |
| ENSG00000125520 | SLC2A4RG | SLC2A4 regulator [Source:HGNC Symbol;Acc:HGNC:15930] | -1.3308 | 2.25E-30 |
| ENSG00000075420 | FNDC3B | fibronectin type III domain containing 3B [Source:HGNC Symbol;Acc:HGNC:24670] | 1.1741 | 2.26E-30 |
| ENSG00000144285 | SCN1A | sodium voltage-gated channel alpha subunit 1 [Source:HGNC Symbol;Acc:HGNC:10585] | -3.2830 | 4.26E-30 |

| | | | | |
|---|---|---|---|---|
| ENSG00000260920 | NA | novel transcript, sense overlapping with ZNF643 | 2.4410 | 4.26E-30 |
| ENSG00000017797 | RALBP1 | ralA binding protein 1 [Source:HGNC Symbol;Acc:HGNC:9841] | -0.9212 | 5.02E-30 |
| ENSG00000151012 | SLC7A11 | solute carrier family 7 member 11 [Source:HGNC Symbol;Acc:HGNC:11059] | 1.5096 | 6.27E-30 |
| ENSG00000150961 | SEC24D | SEC24 homolog D, COPII coat complex component [Source:HGNC Symbol;Acc:HGNC:10706] | 2.1688 | 6.97E-30 |
| ENSG00000154639 | CXADR | CXADR Ig-like cell adhesion molecule [Source:HGNC Symbol;Acc:HGNC:2559] | 1.7446 | 8.17E-30 |
| ENSG00000114315 | HES1 | hes family bHLH transcription factor 1 [Source:HGNC Symbol;Acc:HGNC:5192] | 2.2431 | 8.17E-30 |
| ENSG00000170145 | SIK2 | salt inducible kinase 2 [Source:HGNC Symbol;Acc:HGNC:21680] | 1.3542 | 8.17E-30 |
| ENSG00000139636 | LMBR1L | limb development membrane protein 1 like [Source:HGNC Symbol;Acc:HGNC:18268] | 1.4739 | 1.03E-29 |
| ENSG00000278530 | CHMP1B2P | charged multivesicular body protein 1B2, pseudogene [Source:HGNC Symbol;Acc:HGNC:49380] | 1.9896 | 1.03E-29 |
| ENSG00000106803 | SEC61B | SEC61 translocon subunit beta [Source:HGNC Symbol;Acc:HGNC:16993] | 0.9076 | 1.25E-29 |
| ENSG00000173812 | EIF1 | eukaryotic translation initiation factor 1 [Source:HGNC Symbol;Acc:HGNC:3249] | 0.9235 | 1.36E-29 |
| ENSG00000132906 | CASP9 | caspase 9 [Source:HGNC Symbol;Acc:HGNC:1511] | 1.1634 | 1.40E-29 |
| ENSG00000150093 | ITGB1 | integrin subunit beta 1 [Source:HGNC Symbol;Acc:HGNC:6153] | 1.0123 | 1.63E-29 |
| ENSG00000165434 | PGM2L1 | phosphoglucomutase 2 like 1 [Source:HGNC Symbol;Acc:HGNC:20898] | 2.2061 | 1.88E-29 |
| ENSG00000107290 | SETX | senataxin [Source:HGNC Symbol;Acc:HGNC:445] | 0.6299 | 2.38E-29 |
| ENSG00000144802 | NFKBIZ | NFKB inhibitor zeta [Source:HGNC Symbol;Acc:HGNC:29805] | 2.2811 | 2.50E-29 |
| ENSG00000166794 | PPIB | peptidylprolyl isomerase B [Source:HGNC Symbol;Acc:HGNC:9255] | 0.8472 | 2.51E-29 |
| ENSG00000106537 | TSPAN13 | tetraspanin 13 [Source:HGNC Symbol;Acc:HGNC:21643] | 1.1941 | 2.58E-29 |
| ENSG00000148468 | FAM171A1 | family with sequence similarity 171 member A1 [Source:HGNC Symbol;Acc:HGNC:23522] | -1.0207 | 2.66E-29 |
| ENSG00000176018 | LYSMD3 | LysM domain containing 3 [Source:HGNC Symbol;Acc:HGNC:26969] | 0.8853 | 3.02E-29 |
| ENSG00000150593 | PDCD4 | programmed cell death 4 [Source:HGNC Symbol;Acc:HGNC:8763] | -1.2652 | 3.66E-29 |
| ENSG00000165102 | HGSNAT | heparan-alpha-glucosaminide N-acetyltransferase [Source:HGNC Symbol;Acc:HGNC:26527] | 1.1842 | 4.22E-29 |
| ENSG00000171385 | KCND3 | potassium voltage-gated channel subfamily D member 3 [Source:HGNC Symbol;Acc:HGNC:6239] | -1.1737 | 4.43E-29 |
| ENSG00000120833 | SOCS2 | suppressor of cytokine signaling 2 [Source:HGNC Symbol;Acc:HGNC:19382] | 4.0993 | 4.65E-29 |
| ENSG00000197147 | LRRC8B | leucine rich repeat containing 8 VRAC subunit B [Source:HGNC Symbol;Acc:HGNC:30692] | 0.9170 | 4.95E-29 |
| ENSG00000151276 | MAGI1 | membrane associated guanylate kinase, WW and PDZ domain containing 1 [Source:HGNC Symbol;Acc:HGNC:946] | 0.7951 | 6.38E-29 |
| ENSG00000113742 | CPEB4 | cytoplasmic polyadenylation element binding protein 4 [Source:HGNC Symbol;Acc:HGNC:21747] | 1.4818 | 6.99E-29 |
| ENSG00000130720 | FIBCD1 | fibrinogen C domain containing 1 [Source:HGNC Symbol;Acc:HGNC:25922] | -1.2258 | 8.49E-29 |
| ENSG00000139173 | TMEM117 | transmembrane protein 117 [Source:HGNC Symbol;Acc:HGNC:25308] | 1.2142 | 9.44E-29 |
| ENSG00000179218 | CALR | calreticulin [Source:HGNC Symbol;Acc:HGNC:1455] | 1.1033 | 1.14E-28 |
| ENSG00000149294 | NCAM1 | neural cell adhesion molecule 1 [Source:HGNC Symbol;Acc:HGNC:7656] | 0.9261 | 1.25E-28 |
| ENSG00000151067 | CACNA1C | calcium voltage-gated channel subunit alpha1 C [Source:HGNC Symbol;Acc:HGNC:1390] | 1.8640 | 1.32E-28 |
| ENSG00000106799 | TGFBR1 | transforming growth factor beta receptor 1 [Source:HGNC Symbol;Acc:HGNC:11772] | 1.2569 | 1.47E-28 |
| ENSG00000245105 | A2M-AS1 | A2M antisense RNA 1 [Source:HGNC Symbol;Acc:HGNC:27057] | -1.3602 | 1.59E-28 |
| ENSG00000108389 | MTMR4 | myotubularin related protein 4 [Source:HGNC Symbol;Acc:HGNC:7452] | -0.7931 | 1.79E-28 |
| ENSG00000181754 | AMIGO1 | adhesion molecule with Ig like domain 1 [Source:HGNC Symbol;Acc:HGNC:20824] | -1.0707 | 2.04E-28 |
| ENSG00000178966 | RMI1 | RecQ mediated genome instability 1 [Source:HGNC Symbol;Acc:HGNC:25764] | -0.8762 | 2.27E-28 |
| ENSG00000136720 | HS6ST1 | heparan sulfate 6-O-sulfotransferase 1 [Source:HGNC Symbol;Acc:HGNC:5201] | -1.3558 | 2.45E-28 |
| ENSG00000146648 | EGFR | epidermal growth factor receptor [Source:HGNC Symbol;Acc:HGNC:3236] | 4.5145 | 2.55E-28 |
| ENSG00000169359 | SLC33A1 | solute carrier family 33 member 1 [Source:HGNC Symbol;Acc:HGNC:95] | 0.8109 | 2.65E-28 |
| ENSG00000101255 | TRIB3 | tribbles pseudokinase 3 [Source:HGNC Symbol;Acc:HGNC:16228] | 2.0835 | 3.84E-28 |
| ENSG00000174574 | AKIRIN1 | akirin 1 [Source:HGNC Symbol;Acc:HGNC:25744] | 0.6650 | 4.44E-28 |
| ENSG00000184009 | ACTG1 | actin gamma 1 [Source:HGNC Symbol;Acc:HGNC:144] | 1.5115 | 4.96E-28 |

| | | | | |
|---|---|---|---|---|
| ENSG00000105329 | TGFB1 | transforming growth factor beta 1 [Source:HGNC Symbol;Acc:HGNC:11766] | 0.9950 | 5.33E-28 |
| ENSG00000117707 | PROX1 | prospero homeobox 1 [Source:HGNC Symbol;Acc:HGNC:9459] | 1.9863 | 5.69E-28 |
| ENSG00000134001 | EIF2S1 | eukaryotic translation initiation factor 2 subunit alpha [Source:HGNC Symbol;Acc:HGNC:3265] | 0.6968 | 7.40E-28 |
| ENSG00000104043 | ATP8B4 | ATPase phospholipid transporting 8B4 (putative) [Source:HGNC Symbol;Acc:HGNC:13536] | 2.5667 | 7.50E-28 |
| ENSG00000058668 | ATP2B4 | ATPase plasma membrane Ca2+ transporting 4 [Source:HGNC Symbol;Acc:HGNC:817] | -1.4998 | 8.83E-28 |
| ENSG00000139793 | MBNL2 | muscleblind like splicing regulator 2 [Source:HGNC Symbol;Acc:HGNC:16746] | 1.1492 | 8.83E-28 |
| ENSG00000185813 | PCYT2 | phosphate cytidylyltransferase 2, ethanolamine [Source:HGNC Symbol;Acc:HGNC:8756] | -1.3584 | 9.02E-28 |
| ENSG00000147027 | TMEM47 | transmembrane protein 47 [Source:HGNC Symbol;Acc:HGNC:18515] | 1.5613 | 9.19E-28 |
| ENSG00000223947 | NA | novel transcript | 3.2485 | 1.00E-27 |
| ENSG00000169330 | MINAR1 | membrane integral NOTCH2 associated receptor 1 [Source:HGNC Symbol;Acc:HGNC:29172] | 1.7953 | 1.26E-27 |
| ENSG00000170485 | NPAS2 | neuronal PAS domain protein 2 [Source:HGNC Symbol;Acc:HGNC:7895] | 1.9988 | 1.27E-27 |
| ENSG00000196557 | CACNA1H | calcium voltage-gated channel subunit alpha1 H [Source:HGNC Symbol;Acc:HGNC:1395] | 1.2026 | 1.32E-27 |
| ENSG00000161960 | EIF4A1 | eukaryotic translation initiation factor 4A1 [Source:HGNC Symbol;Acc:HGNC:3282] | 0.7439 | 1.65E-27 |
| ENSG00000157870 | PRXL2B | peroxiredoxin like 2B [Source:HGNC Symbol;Acc:HGNC:28390] | -1.3681 | 1.76E-27 |
| ENSG00000108176 | DNAJC12 | DnaJ heat shock protein family (Hsp40) member C12 [Source:HGNC Symbol;Acc:HGNC:28908] | 1.9713 | 2.04E-27 |
| ENSG00000133398 | MED10 | mediator complex subunit 10 [Source:HGNC Symbol;Acc:HGNC:28760] | 0.9871 | 2.06E-27 |
| ENSG00000107731 | UNC5B | unc-5 netrin receptor B [Source:HGNC Symbol;Acc:HGNC:12568] | -2.3240 | 2.77E-27 |
| ENSG00000078804 | TP53INP2 | tumor protein p53 inducible nuclear protein 2 [Source:HGNC Symbol;Acc:HGNC:16104] | 1.4747 | 2.96E-27 |
| ENSG00000197019 | SERTAD1 | SERTA domain containing 1 [Source:HGNC Symbol;Acc:HGNC:17932] | 1.9787 | 3.17E-27 |
| ENSG00000120705 | ETF1 | eukaryotic translation termination factor 1 [Source:HGNC Symbol;Acc:HGNC:3477] | 0.8030 | 3.71E-27 |
| ENSG00000171867 | PRNP | prion protein [Source:HGNC Symbol;Acc:HGNC:9449] | 1.4616 | 3.98E-27 |
| ENSG00000275202 | NA | novel transcript, antisense to FNDC3A | 2.2178 | 4.18E-27 |
| ENSG00000139998 | RAB15 | RAB15, member RAS oncogene family [Source:HGNC Symbol;Acc:HGNC:20150] | 1.3615 | 4.44E-27 |
| ENSG00000186480 | INSIG1 | insulin induced gene 1 [Source:HGNC Symbol;Acc:HGNC:6083] | -1.2900 | 4.94E-27 |
| ENSG00000071575 | TRIB2 | tribbles pseudokinase 2 [Source:HGNC Symbol;Acc:HGNC:30809] | 1.5637 | 5.06E-27 |
| ENSG00000111711 | GOLT1B | golgi transport 1B [Source:HGNC Symbol;Acc:HGNC:20175] | 0.9627 | 6.33E-27 |
| ENSG00000186897 | C1QL4 | complement C1q like 4 [Source:HGNC Symbol;Acc:HGNC:31416] | -2.0407 | 6.62E-27 |
| ENSG00000113811 | SELENOK | selenoprotein K [Source:HGNC Symbol;Acc:HGNC:30394] | 1.2509 | 7.34E-27 |
| ENSG00000143761 | ARF1 | ADP ribosylation factor 1 [Source:HGNC Symbol;Acc:HGNC:652] | 0.5570 | 9.58E-27 |
| ENSG00000183688 | RFLNB | refilin B [Source:HGNC Symbol;Acc:HGNC:28705] | 2.8863 | 1.00E-26 |
| ENSG00000164237 | CMBL | carboxymethylenebutenolidase homolog [Source:HGNC Symbol;Acc:HGNC:25090] | -1.4017 | 1.02E-26 |
| ENSG00000165272 | AQP3 | aquaporin 3 (Gill blood group) [Source:HGNC Symbol;Acc:HGNC:636] | 4.4832 | 1.08E-26 |
| ENSG00000008405 | CRY1 | cryptochrome circadian regulator 1 [Source:HGNC Symbol;Acc:HGNC:2384] | 1.1846 | 1.10E-26 |
| ENSG00000166292 | TMEM100 | transmembrane protein 100 [Source:HGNC Symbol;Acc:HGNC:25607] | 1.3998 | 1.48E-26 |
| ENSG00000266401 | NA | novel transcript, antisense to DLGAP1 | 3.0135 | 1.54E-26 |
| ENSG00000236013 | NA | novel transcript | 2.1197 | 1.87E-26 |
| ENSG00000185231 | MC2R | melanocortin 2 receptor [Source:HGNC Symbol;Acc:HGNC:6930] | 3.8048 | 1.96E-26 |
| ENSG00000102996 | MMP15 | matrix metallopeptidase 15 [Source:HGNC Symbol;Acc:HGNC:7161] | -1.6182 | 2.07E-26 |
| ENSG00000067167 | TRAM1 | translocation associated membrane protein 1 [Source:HGNC Symbol;Acc:HGNC:20568] | 0.8114 | 2.25E-26 |
| ENSG00000113712 | CSNK1A1 | casein kinase 1 alpha 1 [Source:HGNC Symbol;Acc:HGNC:2451] | 0.9040 | 2.27E-26 |
| ENSG00000107140 | TESK1 | testis associated actin remodelling kinase 1 [Source:HGNC Symbol;Acc:HGNC:11731] | 1.0616 | 2.30E-26 |
| ENSG00000152465 | NMT2 | N-myristoyltransferase 2 [Source:HGNC Symbol;Acc:HGNC:7858] | -1.2662 | 2.40E-26 |
| ENSG00000203727 | SAMD5 | sterile alpha motif domain containing 5 [Source:HGNC Symbol;Acc:HGNC:21180] | -1.1406 | 3.09E-26 |

| | | | | |
|---|---|---|---|---|
| ENSG00000112144 | CILK1 | ciliogenesis associated kinase 1 [Source:HGNC Symbol;Acc:HGNC:21219] | -0.8661 | 3.37E-26 |
| ENSG00000115211 | EIF2B4 | eukaryotic translation initiation factor 2B subunit delta [Source:HGNC Symbol;Acc:HGNC:3260] | 0.8253 | 3.38E-26 |
| ENSG00000105849 | TWISTNB | TWIST neighbor [Source:HGNC Symbol;Acc:HGNC:18027] | 1.7892 | 3.41E-26 |
| ENSG00000158669 | GPAT4 | glycerol-3-phosphate acyltransferase 4 [Source:HGNC Symbol;Acc:HGNC:20880] | 1.1766 | 3.73E-26 |
| ENSG00000186594 | MIR22HG | MIR22 host gene [Source:HGNC Symbol;Acc:HGNC:28219] | 1.8246 | 3.75E-26 |
| ENSG00000214595 | EML6 | EMAP like 6 [Source:HGNC Symbol;Acc:HGNC:35412] | 1.7615 | 4.07E-26 |
| ENSG00000150787 | PTS | 6-pyruvoyltetrahydropterin synthase [Source:HGNC Symbol;Acc:HGNC:9689] | 1.3811 | 4.48E-26 |
| ENSG00000048140 | TSPAN17 | tetraspanin 17 [Source:HGNC Symbol;Acc:HGNC:13594] | -0.7093 | 6.80E-26 |
| ENSG00000179314 | WSCD1 | WSC domain containing 1 [Source:HGNC Symbol;Acc:HGNC:29060] | 3.0251 | 7.09E-26 |
| ENSG00000083857 | FAT1 | FAT atypical cadherin 1 [Source:HGNC Symbol;Acc:HGNC:3595] | 1.9483 | 9.10E-26 |
| ENSG00000244005 | NFS1 | NFS1 cysteine desulfurase [Source:HGNC Symbol;Acc:HGNC:15910] | -1.3688 | 1.14E-25 |
| ENSG00000108829 | LRRC59 | leucine rich repeat containing 59 [Source:HGNC Symbol;Acc:HGNC:28817] | 0.8663 | 1.28E-25 |
| ENSG00000174021 | GNG5 | G protein subunit gamma 5 [Source:HGNC Symbol;Acc:HGNC:4408] | 0.8113 | 1.34E-25 |
| ENSG00000165502 | RPL36AL | ribosomal protein L36a like [Source:HGNC Symbol;Acc:HGNC:10346] | 0.7877 | 1.51E-25 |
| ENSG00000165733 | BMS1 | BMS1 ribosome biogenesis factor [Source:HGNC Symbol;Acc:HGNC:23505] | 0.6716 | 1.51E-25 |
| ENSG00000139194 | RBP5 | retinol binding protein 5 [Source:HGNC Symbol;Acc:HGNC:15847] | 1.6776 | 1.63E-25 |
| ENSG00000102471 | NDFIP2 | Nedd4 family interacting protein 2 [Source:HGNC Symbol;Acc:HGNC:18537] | 1.2755 | 1.89E-25 |
| ENSG00000171951 | SCG2 | secretogranin II [Source:HGNC Symbol;Acc:HGNC:10575] | 3.6784 | 1.92E-25 |
| ENSG00000167460 | TPM4 | tropomyosin 4 [Source:HGNC Symbol;Acc:HGNC:12013] | 1.0607 | 2.29E-25 |
| ENSG00000004660 | CAMKK1 | calcium/calmodulin dependent protein kinase kinase 1 [Source:HGNC Symbol;Acc:HGNC:1469] | -1.8819 | 2.84E-25 |
| ENSG00000143870 | PDIA6 | protein disulfide isomerase family A member 6 [Source:HGNC Symbol;Acc:HGNC:30168] | 1.0462 | 2.87E-25 |
| ENSG00000106031 | HOXA13 | homeobox A13 [Source:HGNC Symbol;Acc:HGNC:5102] | 1.5840 | 3.26E-25 |
| ENSG00000116604 | MEF2D | myocyte enhancer factor 2D [Source:HGNC Symbol;Acc:HGNC:6997] | 0.8585 | 3.68E-25 |
| ENSG00000124006 | OBSL1 | obscurin like cytoskeletal adaptor 1 [Source:HGNC Symbol;Acc:HGNC:29092] | -0.7831 | 3.68E-25 |
| ENSG00000155660 | PDIA4 | protein disulfide isomerase family A member 4 [Source:HGNC Symbol;Acc:HGNC:30167] | 1.5793 | 3.81E-25 |
| ENSG00000100221 | JOSD1 | Josephin domain containing 1 [Source:HGNC Symbol;Acc:HGNC:28953] | 1.0973 | 4.22E-25 |
| ENSG00000166938 | DIS3L | DIS3 like exosome 3'-5' exoribonuclease [Source:HGNC Symbol;Acc:HGNC:28698] | -0.7511 | 4.23E-25 |
| ENSG00000130477 | UNC13A | unc-13 homolog A [Source:HGNC Symbol;Acc:HGNC:23150] | 1.6088 | 4.27E-25 |
| ENSG00000073060 | SCARB1 | scavenger receptor class B member 1 [Source:HGNC Symbol;Acc:HGNC:1664] | 1.7054 | 5.77E-25 |
| ENSG00000154511 | DIPK1A | divergent protein kinase domain 1A [Source:HGNC Symbol;Acc:HGNC:32213] | -1.1092 | 6.06E-25 |
| ENSG00000259070 | LINC00639 | long intergenic non-protein coding RNA 639 [Source:HGNC Symbol;Acc:HGNC:27502] | 2.3626 | 6.19E-25 |
| ENSG00000008513 | ST3GAL1 | ST3 beta-galactoside alpha-2,3-sialyltransferase 1 [Source:HGNC Symbol;Acc:HGNC:10862] | 1.1266 | 6.54E-25 |
| ENSG00000080822 | CLDND1 | claudin domain containing 1 [Source:HGNC Symbol;Acc:HGNC:1322] | 0.8791 | 6.76E-25 |
| ENSG00000150594 | ADRA2A | adrenoceptor alpha 2A [Source:HGNC Symbol;Acc:HGNC:281] | -3.5882 | 6.79E-25 |
| ENSG00000004478 | FKBP4 | FKBP prolyl isomerase 4 [Source:HGNC Symbol;Acc:HGNC:3720] | 0.6861 | 8.62E-25 |
| ENSG00000187801 | ZFP69B | ZFP69 zinc finger protein B [Source:HGNC Symbol;Acc:HGNC:28053] | 2.4421 | 9.06E-25 |
| ENSG00000132405 | TBC1D14 | TBC1 domain family member 14 [Source:HGNC Symbol;Acc:HGNC:29246] | -0.6153 | 1.05E-24 |
| ENSG00000079257 | LXN | latexin [Source:HGNC Symbol;Acc:HGNC:13347] | 1.8126 | 1.13E-24 |
| ENSG00000285976 | NA | novel protein | 1.8835 | 1.22E-24 |
| ENSG00000160285 | LSS | lanosterol synthase [Source:HGNC Symbol;Acc:HGNC:6708] | -1.0743 | 1.22E-24 |
| ENSG00000157657 | ZNF618 | zinc finger protein 618 [Source:HGNC Symbol;Acc:HGNC:29416] | -0.6619 | 1.28E-24 |
| ENSG00000163902 | RPN1 | ribophorin I [Source:HGNC Symbol;Acc:HGNC:10381] | 0.6875 | 1.35E-24 |
| ENSG00000123104 | ITPR2 | inositol 1,4,5-trisphosphate receptor type 2 [Source:HGNC Symbol;Acc:HGNC:6181] | -1.1056 | 1.38E-24 |

| | | | | |
|---|---|---|---|---|
| ENSG00000165527 | ARF6 | ADP ribosylation factor 6 [Source:HGNC Symbol;Acc:HGNC:659] | 0.6333 | 1.44E-24 |
| ENSG00000022840 | RNF10 | ring finger protein 10 [Source:HGNC Symbol;Acc:HGNC:10055] | 0.7996 | 1.47E-24 |
| ENSG00000101294 | HM13 | histocompatibility minor 13 [Source:HGNC Symbol;Acc:HGNC:16435] | 0.8316 | 1.64E-24 |
| ENSG00000160882 | CYP11B1 | cytochrome P450 family 11 subfamily B member 1 [Source:HGNC Symbol;Acc:HGNC:2591] | 3.1542 | 1.98E-24 |
| ENSG00000112773 | TENT5A | terminal nucleotidyltransferase 5A [Source:HGNC Symbol;Acc:HGNC:18345] | 3.0078 | 2.02E-24 |
| ENSG00000204060 | FOXO6 | forkhead box O6 [Source:HGNC Symbol;Acc:HGNC:24814] | -2.7873 | 2.07E-24 |
| ENSG00000135409 | AMHR2 | anti-Mullerian hormone receptor type 2 [Source:HGNC Symbol;Acc:HGNC:465] | -1.3573 | 2.50E-24 |
| ENSG00000116741 | RGS2 | regulator of G protein signaling 2 [Source:HGNC Symbol;Acc:HGNC:9998] | 1.4459 | 2.63E-24 |
| ENSG00000129990 | SYT5 | synaptotagmin 5 [Source:HGNC Symbol;Acc:HGNC:11513] | 1.3559 | 3.00E-24 |
| ENSG00000143889 | HNRNPLL | heterogeneous nuclear ribonucleoprotein L like [Source:HGNC Symbol;Acc:HGNC:25127] | 1.2752 | 3.08E-24 |
| ENSG00000103742 | IGDCC4 | immunoglobulin superfamily DCC subclass member 4 [Source:HGNC Symbol;Acc:HGNC:13770] | -0.8257 | 3.52E-24 |
| ENSG00000115226 | FNDC4 | fibronectin type III domain containing 4 [Source:HGNC Symbol;Acc:HGNC:20239] | 1.8418 | 3.64E-24 |
| ENSG00000148660 | CAMK2G | calcium/calmodulin dependent protein kinase II gamma [Source:HGNC Symbol;Acc:HGNC:1463] | -0.8619 | 3.95E-24 |
| ENSG00000070214 | SLC44A1 | solute carrier family 44 member 1 [Source:HGNC Symbol;Acc:HGNC:18798] | -0.9797 | 4.66E-24 |
| ENSG00000104419 | NDRG1 | N-myc downstream regulated 1 [Source:HGNC Symbol;Acc:HGNC:7679] | 2.4874 | 4.88E-24 |
| ENSG00000179981 | TSHZ1 | teashirt zinc finger homeobox 1 [Source:HGNC Symbol;Acc:HGNC:10669] | -1.0984 | 5.06E-24 |
| ENSG00000173275 | ZNF449 | zinc finger protein 449 [Source:HGNC Symbol;Acc:HGNC:21039] | -1.2929 | 5.54E-24 |
| ENSG00000095397 | WHRN | whirlin [Source:HGNC Symbol;Acc:HGNC:16361] | 1.3380 | 5.69E-24 |
| ENSG00000102780 | DGKH | diacylglycerol kinase eta [Source:HGNC Symbol;Acc:HGNC:2854] | 1.4535 | 5.79E-24 |
| ENSG00000148339 | SLC25A25 | solute carrier family 25 member 25 [Source:HGNC Symbol;Acc:HGNC:20663] | 1.4639 | 5.90E-24 |
| ENSG00000124641 | MED20 | mediator complex subunit 20 [Source:HGNC Symbol;Acc:HGNC:16840] | -1.1466 | 6.17E-24 |
| ENSG00000132341 | RAN | RAN, member RAS oncogene family [Source:HGNC Symbol;Acc:HGNC:9846] | 1.2454 | 6.40E-24 |
| ENSG00000102580 | DNAJC3 | DnaJ heat shock protein family (Hsp40) member C3 [Source:HGNC Symbol;Acc:HGNC:9439] | 1.5311 | 6.66E-24 |
| ENSG00000112541 | PDE10A | phosphodiesterase 10A [Source:HGNC Symbol;Acc:HGNC:8772] | 2.0603 | 6.84E-24 |
| ENSG00000095321 | CRAT | carnitine O-acetyltransferase [Source:HGNC Symbol;Acc:HGNC:2342] | -0.7858 | 6.87E-24 |
| ENSG00000185340 | GAS2L1 | growth arrest specific 2 like 1 [Source:HGNC Symbol;Acc:HGNC:16955] | -1.4415 | 7.22E-24 |
| ENSG00000128228 | SDF2L1 | stromal cell derived factor 2 like 1 [Source:HGNC Symbol;Acc:HGNC:10676] | 2.3552 | 7.52E-24 |
| ENSG00000185022 | MAFF | MAF bZIP transcription factor F [Source:HGNC Symbol;Acc:HGNC:6780] | 4.0248 | 8.25E-24 |
| ENSG00000096384 | HSP90AB1 | heat shock protein 90 alpha family class B member 1 [Source:HGNC Symbol;Acc:HGNC:5258] | 0.8248 | 9.28E-24 |
| ENSG00000077063 | CTTNBP2 | cortactin binding protein 2 [Source:HGNC Symbol;Acc:HGNC:15679] | -1.8381 | 1.01E-23 |
| ENSG00000256235 | SMIM3 | small integral membrane protein 3 [Source:HGNC Symbol;Acc:HGNC:30248] | 1.6279 | 1.02E-23 |
| ENSG00000120742 | SERP1 | stress associated endoplasmic reticulum protein 1 [Source:HGNC Symbol;Acc:HGNC:10759] | 0.9258 | 1.15E-23 |
| ENSG00000167004 | PDIA3 | protein disulfide isomerase family A member 3 [Source:HGNC Symbol;Acc:HGNC:4606] | 0.8719 | 1.25E-23 |
| ENSG00000106105 | GARS1 | glycyl-tRNA synthetase 1 [Source:HGNC Symbol;Acc:HGNC:4162] | 0.8833 | 1.43E-23 |
| ENSG00000103479 | RBL2 | RB transcriptional corepressor like 2 [Source:HGNC Symbol;Acc:HGNC:9894] | -0.5899 | 1.47E-23 |
| ENSG00000054356 | PTPRN | protein tyrosine phosphatase receptor type N [Source:HGNC Symbol;Acc:HGNC:9676] | 3.3181 | 1.47E-23 |
| ENSG00000057757 | PITHD1 | PITH domain containing 1 [Source:HGNC Symbol;Acc:HGNC:25022] | 0.7871 | 1.62E-23 |
| ENSG00000129292 | PHF20L1 | PHD finger protein 20 like 1 [Source:HGNC Symbol;Acc:HGNC:24280] | 0.7018 | 1.62E-23 |
| ENSG00000122694 | GLIPR2 | GLI pathogenesis related 2 [Source:HGNC Symbol;Acc:HGNC:18007] | -1.1832 | 1.81E-23 |
| ENSG00000155304 | HSPA13 | heat shock protein family A (Hsp70) member 13 [Source:HGNC Symbol;Acc:HGNC:11375] | 0.7113 | 1.93E-23 |
| ENSG00000108010 | GLRX3 | glutaredoxin 3 [Source:HGNC Symbol;Acc:HGNC:15987] | 0.6948 | 2.01E-23 |
| ENSG00000159023 | EPB41 | erythrocyte membrane protein band 4.1 [Source:HGNC Symbol;Acc:HGNC:3377] | -0.8928 | 2.08E-23 |
| ENSG00000128011 | LRFN1 | leucine rich repeat and fibronectin type III domain containing 1 [Source:HGNC Symbol;Acc:HGNC:29290] | -1.2403 | 2.28E-23 |

| | | | | |
|---|---|---|---|---|
| ENSG00000243335 | KCTD7 | potassium channel tetramerization domain containing 7 [Source:HGNC Symbol;Acc:HGNC:21957] | -1.2819 | 2.42E-23 |
| ENSG00000185269 | NOTUM | notum, palmitoleoyl-protein carboxylesterase [Source:HGNC Symbol;Acc:HGNC:27106] | -2.3864 | 2.62E-23 |
| ENSG00000081181 | ARG2 | arginase 2 [Source:HGNC Symbol;Acc:HGNC:664] | 1.4809 | 2.72E-23 |
| ENSG00000127022 | CANX | calnexin [Source:HGNC Symbol;Acc:HGNC:1473] | 0.6784 | 2.72E-23 |
| ENSG00000113407 | TARS1 | threonyl-tRNA synthetase 1 [Source:HGNC Symbol;Acc:HGNC:11572] | 0.5731 | 3.56E-23 |
| ENSG00000111266 | DUSP16 | dual specificity phosphatase 16 [Source:HGNC Symbol;Acc:HGNC:17909] | 0.8528 | 3.92E-23 |
| ENSG00000127955 | GNAI1 | G protein subunit alpha i1 [Source:HGNC Symbol;Acc:HGNC:4384] | 1.4171 | 4.20E-23 |
| ENSG00000167291 | TBC1D16 | TBC1 domain family member 16 [Source:HGNC Symbol;Acc:HGNC:28356] | -0.8311 | 5.12E-23 |
| ENSG00000163866 | SMIM12 | small integral membrane protein 12 [Source:HGNC Symbol;Acc:HGNC:25154] | 0.6378 | 5.26E-23 |
| ENSG00000068383 | INPP5A | inositol polyphosphate-5-phosphatase A [Source:HGNC Symbol;Acc:HGNC:6076] | 1.3698 | 5.43E-23 |
| ENSG00000109265 | CRACD | capping protein inhibiting regulator of actin dynamics [Source:HGNC Symbol;Acc:HGNC:29219] | -1.0092 | 5.46E-23 |
| ENSG00000138709 | LARP1B | La ribonucleoprotein 1B [Source:HGNC Symbol;Acc:HGNC:24704] | 1.3465 | 5.46E-23 |
| ENSG00000113575 | PPP2CA | protein phosphatase 2 catalytic subunit alpha [Source:HGNC Symbol;Acc:HGNC:9299] | 0.5742 | 5.67E-23 |
| ENSG00000168288 | MMADHC | metabolism of cobalamin associated D [Source:HGNC Symbol;Acc:HGNC:25221] | 0.6807 | 5.89E-23 |
| ENSG00000118971 | CCND2 | cyclin D2 [Source:HGNC Symbol;Acc:HGNC:1583] | -1.5231 | 6.51E-23 |
| ENSG00000272325 | NUDT3 | nudix hydrolase 3 [Source:HGNC Symbol;Acc:HGNC:8050] | -0.8260 | 7.18E-23 |
| ENSG00000177311 | ZBTB38 | zinc finger and BTB domain containing 38 [Source:HGNC Symbol;Acc:HGNC:26636] | 0.8458 | 7.55E-23 |
| ENSG00000110697 | PITPNM1 | phosphatidylinositol transfer protein membrane associated 1 [Source:HGNC Symbol;Acc:HGNC:9003] | 1.0948 | 7.94E-23 |
| ENSG00000112062 | MAPK14 | mitogen-activated protein kinase 14 [Source:HGNC Symbol;Acc:HGNC:6876] | -0.7561 | 7.98E-23 |
| ENSG00000160211 | G6PD | glucose-6-phosphate dehydrogenase [Source:HGNC Symbol;Acc:HGNC:4057] | 1.1259 | 8.13E-23 |
| ENSG00000163431 | LMOD1 | leiomodin 1 [Source:HGNC Symbol;Acc:HGNC:6647] | -1.9502 | 9.52E-23 |
| ENSG00000173548 | SNX33 | sorting nexin 33 [Source:HGNC Symbol;Acc:HGNC:28468] | 1.9595 | 1.05E-22 |
| ENSG00000165802 | NSMF | NMDA receptor synaptonuclear signaling and neuronal migration factor [Source:HGNC Symbol;Acc:HGNC:29843] | 0.7282 | 1.08E-22 |
| ENSG00000143061 | IGSF3 | immunoglobulin superfamily member 3 [Source:HGNC Symbol;Acc:HGNC:5950] | -1.2328 | 1.10E-22 |
| ENSG00000065057 | NTHL1 | nth like DNA glycosylase 1 [Source:HGNC Symbol;Acc:HGNC:8028] | -1.5461 | 1.18E-22 |
| ENSG00000122378 | PRXL2A | peroxiredoxin like 2A [Source:HGNC Symbol;Acc:HGNC:28651] | -0.6591 | 1.29E-22 |
| ENSG00000100075 | SLC25A1 | solute carrier family 25 member 1 [Source:HGNC Symbol;Acc:HGNC:10979] | -0.8629 | 1.40E-22 |
| ENSG00000135679 | MDM2 | MDM2 proto-oncogene [Source:HGNC Symbol;Acc:HGNC:6973] | 1.0028 | 1.54E-22 |
| ENSG00000076864 | RAP1GAP | RAP1 GTPase activating protein [Source:HGNC Symbol;Acc:HGNC:9858] | -1.5109 | 1.58E-22 |
| ENSG00000240694 | PNMA2 | PNMA family member 2 [Source:HGNC Symbol;Acc:HGNC:9159] | -1.0817 | 1.63E-22 |
| ENSG00000164506 | STXBP5 | syntaxin binding protein 5 [Source:HGNC Symbol;Acc:HGNC:19665] | 1.2095 | 1.69E-22 |
| ENSG00000164484 | TMEM200A | transmembrane protein 200A [Source:HGNC Symbol;Acc:HGNC:21075] | 1.7328 | 1.74E-22 |
| ENSG00000139372 | TDG | thymine DNA glycosylase [Source:HGNC Symbol;Acc:HGNC:11700] | 0.8598 | 1.88E-22 |
| ENSG00000144597 | EAF1 | ELL associated factor 1 [Source:HGNC Symbol;Acc:HGNC:20907] | 0.7352 | 2.07E-22 |
| ENSG00000162885 | B3GALNT2 | beta-1,3-N-acetylgalactosaminyltransferase 2 [Source:HGNC Symbol;Acc:HGNC:28596] | 2.8036 | 2.07E-22 |
| ENSG00000143621 | ILF2 | interleukin enhancer binding factor 2 [Source:HGNC Symbol;Acc:HGNC:6037] | 0.6924 | 2.14E-22 |
| ENSG00000131791 | PRKAB2 | protein kinase AMP-activated non-catalytic subunit beta 2 [Source:HGNC Symbol;Acc:HGNC:9379] | -0.9959 | 2.20E-22 |
| ENSG00000178996 | SNX18 | sorting nexin 18 [Source:HGNC Symbol;Acc:HGNC:19245] | -0.8881 | 2.51E-22 |
| ENSG00000114529 | C3orf52 | chromosome 3 open reading frame 52 [Source:HGNC Symbol;Acc:HGNC:26255] | 1.3018 | 2.60E-22 |
| ENSG00000140859 | KIFC3 | kinesin family member C3 [Source:HGNC Symbol;Acc:HGNC:6326] | 1.4218 | 2.66E-22 |
| ENSG00000179029 | TMEM107 | transmembrane protein 107 [Source:HGNC Symbol;Acc:HGNC:28128] | -1.2194 | 3.33E-22 |
| ENSG00000171408 | PDE7B | phosphodiesterase 7B [Source:HGNC Symbol;Acc:HGNC:8792] | 1.1780 | 3.33E-22 |
| ENSG00000184164 | CRELD2 | cysteine rich with EGF like domains 2 [Source:HGNC Symbol;Acc:HGNC:28150] | 1.7254 | 3.35E-22 |

| | | | | |
|---|---|---|---|---|
| ENSG00000138413 | IDH1 | isocitrate dehydrogenase (NADP(+)) 1 [Source:HGNC Symbol;Acc:HGNC:5382] | -0.8462 | 3.57E-22 |
| ENSG00000165476 | REEP3 | receptor accessory protein 3 [Source:HGNC Symbol;Acc:HGNC:23711] | 1.2471 | 4.26E-22 |
| ENSG00000103160 | HSDL1 | hydroxysteroid dehydrogenase like 1 [Source:HGNC Symbol;Acc:HGNC:16475] | -0.8415 | 4.51E-22 |
| ENSG00000006283 | CACNA1G | calcium voltage-gated channel subunit alpha1 G [Source:HGNC Symbol;Acc:HGNC:1394] | -1.4460 | 4.97E-22 |
| ENSG00000234956 | LINC02539 | long intergenic non-protein coding RNA 2539 [Source:HGNC Symbol;Acc:HGNC:53572] | 1.9867 | 5.32E-22 |
| ENSG00000142677 | IL22RA1 | interleukin 22 receptor subunit alpha 1 [Source:HGNC Symbol;Acc:HGNC:13700] | -2.4811 | 5.55E-22 |
| ENSG00000013275 | PSMC4 | proteasome 26S subunit, ATPase 4 [Source:HGNC Symbol;Acc:HGNC:9551] | 0.5960 | 5.74E-22 |
| ENSG00000169083 | AR | androgen receptor [Source:HGNC Symbol;Acc:HGNC:644] | -1.2174 | 6.37E-22 |
| ENSG00000127603 | MACF1 | microtubule actin crosslinking factor 1 [Source:HGNC Symbol;Acc:HGNC:13664] | 0.6656 | 6.53E-22 |
| ENSG00000085415 | SEH1L | SEH1 like nucleoporin [Source:HGNC Symbol;Acc:HGNC:30379] | 1.0664 | 7.69E-22 |
| ENSG00000075426 | FOSL2 | FOS like 2, AP-1 transcription factor subunit [Source:HGNC Symbol;Acc:HGNC:3798] | 1.4145 | 8.91E-22 |
| ENSG00000110076 | NRXN2 | neurexin 2 [Source:HGNC Symbol;Acc:HGNC:8009] | -0.9724 | 9.23E-22 |
| ENSG00000081189 | MEF2C | myocyte enhancer factor 2C [Source:HGNC Symbol;Acc:HGNC:6996] | 1.5055 | 9.43E-22 |
| ENSG00000184216 | IRAK1 | interleukin 1 receptor associated kinase 1 [Source:HGNC Symbol;Acc:HGNC:6112] | 0.8253 | 9.85E-22 |
| ENSG00000181090 | EHMT1 | euchromatic histone lysine methyltransferase 1 [Source:HGNC Symbol;Acc:HGNC:24650] | -0.7094 | 1.03E-21 |
| ENSG00000198795 | ZNF521 | zinc finger protein 521 [Source:HGNC Symbol;Acc:HGNC:24605] | -0.7023 | 1.09E-21 |
| ENSG00000025772 | TOMM34 | translocase of outer mitochondrial membrane 34 [Source:HGNC Symbol;Acc:HGNC:15746] | 0.7799 | 1.20E-21 |
| ENSG00000080031 | PTPRH | protein tyrosine phosphatase receptor type H [Source:HGNC Symbol;Acc:HGNC:9672] | 1.7774 | 1.26E-21 |
| ENSG00000067955 | CBFB | core-binding factor subunit beta [Source:HGNC Symbol;Acc:HGNC:1539] | 1.1787 | 1.37E-21 |
| ENSG00000135249 | RINT1 | RAD50 interactor 1 [Source:HGNC Symbol;Acc:HGNC:21876] | 1.0959 | 1.44E-21 |
| ENSG00000196273 | LINC00523 | long intergenic non-protein coding RNA 523 [Source:HGNC Symbol;Acc:HGNC:20117] | -1.6854 | 1.44E-21 |
| ENSG00000145220 | LYAR | Ly1 antibody reactive [Source:HGNC Symbol;Acc:HGNC:26021] | 0.8240 | 1.50E-21 |
| ENSG00000136158 | SPRY2 | sprouty RTK signaling antagonist 2 [Source:HGNC Symbol;Acc:HGNC:11270] | 2.1732 | 1.54E-21 |
| ENSG00000089057 | SLC23A2 | solute carrier family 23 member 2 [Source:HGNC Symbol;Acc:HGNC:10973] | 1.1856 | 1.56E-21 |
| ENSG00000164733 | CTSB | cathepsin B [Source:HGNC Symbol;Acc:HGNC:2527] | 0.9129 | 1.58E-21 |
| ENSG00000164951 | PDP1 | pyruvate dehyrogenase phosphatase catalytic subunit 1 [Source:HGNC Symbol;Acc:HGNC:9279] | 1.8017 | 1.61E-21 |
| ENSG00000198853 | RUSC2 | RUN and SH3 domain containing 2 [Source:HGNC Symbol;Acc:HGNC:23625] | 1.1739 | 1.65E-21 |
| ENSG00000134363 | FST | follistatin [Source:HGNC Symbol;Acc:HGNC:3971] | 2.5892 | 1.80E-21 |
| ENSG00000092068 | SLC7A8 | solute carrier family 7 member 8 [Source:HGNC Symbol;Acc:HGNC:11066] | -1.1178 | 1.81E-21 |
| ENSG00000171067 | C11orf24 | chromosome 11 open reading frame 24 [Source:HGNC Symbol;Acc:HGNC:1174] | 1.1007 | 2.15E-21 |
| ENSG00000088280 | ASAP3 | ArfGAP with SH3 domain, ankyrin repeat and PH domain 3 [Source:HGNC Symbol;Acc:HGNC:14987] | -0.9548 | 2.39E-21 |
| ENSG00000170439 | METTL7B | methyltransferase like 7B [Source:HGNC Symbol;Acc:HGNC:28276] | 1.5747 | 2.57E-21 |
| ENSG00000100629 | CEP128 | centrosomal protein 128 [Source:HGNC Symbol;Acc:HGNC:20359] | 1.1924 | 2.70E-21 |
| ENSG00000175183 | CSRP2 | cysteine and glycine rich protein 2 [Source:HGNC Symbol;Acc:HGNC:2470] | -1.1549 | 2.71E-21 |
| ENSG00000133657 | ATP13A3 | ATPase 13A3 [Source:HGNC Symbol;Acc:HGNC:24113] | 0.7963 | 2.80E-21 |
| ENSG00000137575 | SDCBP | syndecan binding protein [Source:HGNC Symbol;Acc:HGNC:10662] | 1.0865 | 3.24E-21 |
| ENSG00000204291 | COL15A1 | collagen type XV alpha 1 chain [Source:HGNC Symbol;Acc:HGNC:2192] | 1.5575 | 3.31E-21 |
| ENSG00000148248 | SURF4 | surfeit 4 [Source:HGNC Symbol;Acc:HGNC:11476] | 0.7162 | 3.88E-21 |
| ENSG00000124766 | SOX4 | SRY-box transcription factor 4 [Source:HGNC Symbol;Acc:HGNC:11200] | 1.9655 | 3.97E-21 |
| ENSG00000145721 | LIX1 | limb and CNS expressed 1 [Source:HGNC Symbol;Acc:HGNC:18581] | -1.5172 | 4.18E-21 |
| ENSG00000162909 | CAPN2 | calpain 2 [Source:HGNC Symbol;Acc:HGNC:1479] | 1.0478 | 4.50E-21 |
| ENSG00000122515 | ZMIZ2 | zinc finger MIZ-type containing 2 [Source:HGNC Symbol;Acc:HGNC:22229] | 0.7290 | 6.04E-21 |
| ENSG00000169213 | RAB3B | RAB3B, member RAS oncogene family [Source:HGNC Symbol;Acc:HGNC:9778] | -1.5091 | 6.73E-21 |

| | | | | |
|---|---|---|---|---|
| ENSG00000111674 | ENO2 | enolase 2 [Source:HGNC Symbol;Acc:HGNC:3353] | 1.2782 | 6.95E-21 |
| ENSG00000120802 | TMPO | thymopoietin [Source:HGNC Symbol;Acc:HGNC:11875] | 0.8527 | 6.97E-21 |
| ENSG00000131871 | SELENOS | selenoprotein S [Source:HGNC Symbol;Acc:HGNC:30396] | 0.9631 | 7.46E-21 |
| ENSG00000185033 | SEMA4B | semaphorin 4B [Source:HGNC Symbol;Acc:HGNC:10730] | -1.1425 | 8.63E-21 |
| ENSG00000113070 | HBEGF | heparin binding EGF like growth factor [Source:HGNC Symbol;Acc:HGNC:3059] | 1.8205 | 9.19E-21 |
| ENSG00000139278 | GLIPR1 | GLI pathogenesis related 1 [Source:HGNC Symbol;Acc:HGNC:17001] | 1.8280 | 9.76E-21 |
| ENSG00000141452 | RMC1 | regulator of MON1-CCZ1 [Source:HGNC Symbol;Acc:HGNC:24326] | 0.8042 | 9.85E-21 |
| ENSG00000137878 | GCOM1 | GRINL1A complex locus 1 [Source:HGNC Symbol;Acc:HGNC:26424] | 1.5679 | 1.01E-20 |
| ENSG00000167972 | ABCA3 | ATP binding cassette subfamily A member 3 [Source:HGNC Symbol;Acc:HGNC:33] | -0.8865 | 1.05E-20 |
| ENSG00000175727 | MLXIP | MLX interacting protein [Source:HGNC Symbol;Acc:HGNC:17055] | 1.1385 | 1.08E-20 |
| ENSG00000155256 | ZFYVE27 | zinc finger FYVE-type containing 27 [Source:HGNC Symbol;Acc:HGNC:26559] | 1.0846 | 1.09E-20 |
| ENSG00000235314 | LINC00957 | long intergenic non-protein coding RNA 957 [Source:HGNC Symbol;Acc:HGNC:22332] | -0.8341 | 1.14E-20 |
| ENSG00000121957 | GPSM2 | G protein signaling modulator 2 [Source:HGNC Symbol;Acc:HGNC:29501] | -0.8080 | 1.16E-20 |
| ENSG00000072135 | PTPN18 | protein tyrosine phosphatase non-receptor type 18 [Source:HGNC Symbol;Acc:HGNC:9649] | -1.2900 | 1.25E-20 |
| ENSG00000113384 | GOLPH3 | golgi phosphoprotein 3 [Source:HGNC Symbol;Acc:HGNC:15452] | 0.6955 | 1.29E-20 |
| ENSG00000231852 | CYP21A2 | cytochrome P450 family 21 subfamily A member 2 [Source:HGNC Symbol;Acc:HGNC:2600] | 1.6071 | 1.38E-20 |
| ENSG00000141965 | FEM1A | fem-1 homolog A [Source:HGNC Symbol;Acc:HGNC:16934] | 0.9663 | 1.59E-20 |
| ENSG00000132613 | MTSS2 | MTSS I-BAR domain containing 2 [Source:HGNC Symbol;Acc:HGNC:25094] | -0.7184 | 1.75E-20 |
| ENSG00000145284 | SCD5 | stearoyl-CoA desaturase 5 [Source:HGNC Symbol;Acc:HGNC:21088] | -0.6403 | 2.11E-20 |
| ENSG00000136710 | CCDC115 | coiled-coil domain containing 115 [Source:HGNC Symbol;Acc:HGNC:28178] | -0.8495 | 2.12E-20 |
| ENSG00000101558 | VAPA | VAMP associated protein A [Source:HGNC Symbol;Acc:HGNC:12648] | 0.8067 | 2.32E-20 |
| ENSG00000133112 | TPT1 | tumor protein, translationally-controlled 1 [Source:HGNC Symbol;Acc:HGNC:12022] | 0.6943 | 2.46E-20 |
| ENSG00000149218 | ENDOD1 | endonuclease domain containing 1 [Source:HGNC Symbol;Acc:HGNC:29129] | 1.1133 | 2.47E-20 |
| ENSG00000101608 | MYL12A | myosin light chain 12A [Source:HGNC Symbol;Acc:HGNC:16701] | 1.0264 | 2.55E-20 |
| ENSG00000241058 | NSUN6 | NOP2/Sun RNA methyltransferase 6 [Source:HGNC Symbol;Acc:HGNC:23529] | 1.0657 | 2.69E-20 |
| ENSG00000117650 | NEK2 | NIMA related kinase 2 [Source:HGNC Symbol;Acc:HGNC:7745] | -0.6747 | 2.71E-20 |
| ENSG00000119915 | ELOVL3 | ELOVL fatty acid elongase 3 [Source:HGNC Symbol;Acc:HGNC:18047] | -2.2532 | 2.85E-20 |
| ENSG00000175556 | LONRF3 | LON peptidase N-terminal domain and ring finger 3 [Source:HGNC Symbol;Acc:HGNC:21152] | 1.9896 | 2.98E-20 |
| ENSG00000162695 | SLC30A7 | solute carrier family 30 member 7 [Source:HGNC Symbol;Acc:HGNC:19306] | 0.8427 | 3.12E-20 |
| ENSG00000144891 | AGTR1 | angiotensin II receptor type 1 [Source:HGNC Symbol;Acc:HGNC:336] | -1.0096 | 3.12E-20 |
| ENSG00000100228 | RAB36 | RAB36, member RAS oncogene family [Source:HGNC Symbol;Acc:HGNC:9775] | 0.8331 | 3.13E-20 |
| ENSG00000137177 | KIF13A | kinesin family member 13A [Source:HGNC Symbol;Acc:HGNC:14566] | -1.5524 | 3.16E-20 |
| ENSG00000065833 | ME1 | malic enzyme 1 [Source:HGNC Symbol;Acc:HGNC:6983] | 0.6500 | 3.70E-20 |
| ENSG00000160131 | VMA21 | vacuolar ATPase assembly factor VMA21 [Source:HGNC Symbol;Acc:HGNC:22082] | 1.0586 | 3.80E-20 |
| ENSG00000127920 | GNG11 | G protein subunit gamma 11 [Source:HGNC Symbol;Acc:HGNC:4403] | 1.6376 | 3.97E-20 |
| ENSG00000179409 | GEMIN4 | gem nuclear organelle associated protein 4 [Source:HGNC Symbol;Acc:HGNC:15717] | 1.3220 | 4.01E-20 |
| ENSG00000186868 | MAPT | microtubule associated protein tau [Source:HGNC Symbol;Acc:HGNC:6893] | -0.9701 | 4.03E-20 |
| ENSG00000166847 | DCTN5 | dynactin subunit 5 [Source:HGNC Symbol;Acc:HGNC:24594] | 0.6387 | 4.07E-20 |
| ENSG00000174437 | ATP2A2 | ATPase sarcoplasmic/endoplasmic reticulum Ca2+ transporting 2 [Source:HGNC Symbol;Acc:HGNC:812] | 1.0382 | 4.07E-20 |
| ENSG00000165698 | SPACA9 | sperm acrosome associated 9 [Source:HGNC Symbol;Acc:HGNC:1367] | -1.4749 | 4.12E-20 |
| ENSG00000146733 | PSPH | phosphoserine phosphatase [Source:HGNC Symbol;Acc:HGNC:9577] | -0.9270 | 4.26E-20 |
| ENSG00000160209 | PDXK | pyridoxal kinase [Source:HGNC Symbol;Acc:HGNC:8819] | 1.0323 | 4.64E-20 |
| ENSG00000174738 | NR1D2 | nuclear receptor subfamily 1 group D member 2 [Source:HGNC Symbol;Acc:HGNC:7963] | -1.4595 | 5.02E-20 |

| | | | | |
|---|---|---|---|---|
| ENSG00000163637 | PRICKLE2 | prickle planar cell polarity protein 2 [Source:HGNC Symbol;Acc:HGNC:20340] | 2.1229 | 5.42E-20 |
| ENSG00000060138 | YBX3 | Y-box binding protein 3 [Source:HGNC Symbol;Acc:HGNC:2428] | 1.0173 | 5.56E-20 |
| ENSG00000167114 | SLC27A4 | solute carrier family 27 member 4 [Source:HGNC Symbol;Acc:HGNC:10998] | -1.1406 | 6.34E-20 |
| ENSG00000101605 | MYOM1 | myomesin 1 [Source:HGNC Symbol;Acc:HGNC:7613] | 1.7431 | 6.87E-20 |
| ENSG00000005700 | IBTK | inhibitor of Bruton tyrosine kinase [Source:HGNC Symbol;Acc:HGNC:17853] | 0.8334 | 7.89E-20 |
| ENSG00000076043 | REXO2 | RNA exonuclease 2 [Source:HGNC Symbol;Acc:HGNC:17851] | 0.7997 | 8.24E-20 |
| ENSG00000109519 | GRPEL1 | GrpE like 1, mitochondrial [Source:HGNC Symbol;Acc:HGNC:19696] | 0.8037 | 8.24E-20 |
| ENSG00000272763 | NA | novel transcript | 0.9466 | 8.25E-20 |
| ENSG00000198546 | ZNF511 | zinc finger protein 511 [Source:HGNC Symbol;Acc:HGNC:28445] | 1.0289 | 8.47E-20 |
| ENSG00000176871 | WSB2 | WD repeat and SOCS box containing 2 [Source:HGNC Symbol;Acc:HGNC:19222] | 0.7602 | 8.86E-20 |
| ENSG00000143756 | FBXO28 | F-box protein 28 [Source:HGNC Symbol;Acc:HGNC:29046] | 1.0590 | 9.34E-20 |
| ENSG00000170445 | HARS1 | histidyl-tRNA synthetase 1 [Source:HGNC Symbol;Acc:HGNC:4816] | 0.6620 | 9.36E-20 |
| ENSG00000163938 | GNL3 | G protein nucleolar 3 [Source:HGNC Symbol;Acc:HGNC:29931] | 0.8620 | 9.59E-20 |
| ENSG00000142459 | EVI5L | ecotropic viral integration site 5 like [Source:HGNC Symbol;Acc:HGNC:30464] | -0.7494 | 9.60E-20 |
| ENSG00000242498 | ARPIN | actin related protein 2/3 complex inhibitor [Source:HGNC Symbol;Acc:HGNC:28782] | -0.8527 | 9.71E-20 |
| ENSG00000176788 | BASP1 | brain abundant membrane attached signal protein 1 [Source:HGNC Symbol;Acc:HGNC:957] | -1.2708 | 1.03E-19 |
| ENSG00000088970 | KIZ | kizuna centrosomal protein [Source:HGNC Symbol;Acc:HGNC:15865] | 1.4609 | 1.05E-19 |
| ENSG00000175416 | CLTB | clathrin light chain B [Source:HGNC Symbol;Acc:HGNC:2091] | 0.9687 | 1.09E-19 |
| ENSG00000225465 | RFPL1S | RFPL1 antisense RNA 1 [Source:HGNC Symbol;Acc:HGNC:9978] | -2.9243 | 1.12E-19 |
| ENSG00000198380 | GFPT1 | glutamine--fructose-6-phosphate transaminase 1 [Source:HGNC Symbol;Acc:HGNC:4241] | 0.7798 | 1.16E-19 |
| ENSG00000112343 | TRIM38 | tripartite motif containing 38 [Source:HGNC Symbol;Acc:HGNC:10059] | 0.9580 | 1.19E-19 |
| ENSG00000171877 | FRMD5 | FERM domain containing 5 [Source:HGNC Symbol;Acc:HGNC:28214] | 2.8287 | 1.32E-19 |
| ENSG00000172738 | TMEM217 | transmembrane protein 217 [Source:HGNC Symbol;Acc:HGNC:21238] | 2.2152 | 1.39E-19 |
| ENSG00000275880 | NA | novel transcript, antisense to CARKD | 2.5981 | 1.41E-19 |
| ENSG00000142669 | SH3BGRL3 | SH3 domain binding glutamate rich protein like 3 [Source:HGNC Symbol;Acc:HGNC:15568] | 0.8860 | 1.43E-19 |
| ENSG00000184205 | TSPYL2 | TSPY like 2 [Source:HGNC Symbol;Acc:HGNC:24358] | 1.2074 | 1.54E-19 |
| ENSG00000188322 | SBK1 | SH3 domain binding kinase 1 [Source:HGNC Symbol;Acc:HGNC:17699] | -1.0376 | 1.65E-19 |
| ENSG00000185634 | SHC4 | SHC adaptor protein 4 [Source:HGNC Symbol;Acc:HGNC:16743] | 3.3751 | 1.69E-19 |
| ENSG00000165732 | DDX21 | DExD-box helicase 21 [Source:HGNC Symbol;Acc:HGNC:2744] | 0.7545 | 1.96E-19 |
| ENSG00000169032 | MAP2K1 | mitogen-activated protein kinase kinase 1 [Source:HGNC Symbol;Acc:HGNC:6840] | 0.7805 | 1.97E-19 |
| ENSG00000104064 | GABPB1 | GA binding protein transcription factor subunit beta 1 [Source:HGNC Symbol;Acc:HGNC:4074] | 0.9113 | 2.03E-19 |
| ENSG00000147649 | MTDH | metadherin [Source:HGNC Symbol;Acc:HGNC:29608] | 0.7358 | 2.08E-19 |
| ENSG00000231784 | DBIL5P | diazepam binding inhibitor-like 5, pseudogene [Source:HGNC Symbol;Acc:HGNC:38519] | 4.0962 | 2.15E-19 |
| ENSG00000142207 | URB1 | URB1 ribosome biogenesis homolog [Source:HGNC Symbol;Acc:HGNC:17344] | 0.9258 | 2.26E-19 |
| ENSG00000029639 | TFB1M | transcription factor B1, mitochondrial [Source:HGNC Symbol;Acc:HGNC:17037] | 0.9910 | 2.28E-19 |
| ENSG00000134109 | EDEM1 | ER degradation enhancing alpha-mannosidase like protein 1 [Source:HGNC Symbol;Acc:HGNC:18967] | 1.3910 | 2.30E-19 |
| ENSG00000160360 | GPSM1 | G protein signaling modulator 1 [Source:HGNC Symbol;Acc:HGNC:17858] | 0.8447 | 2.46E-19 |
| ENSG00000137409 | MTCH1 | mitochondrial carrier 1 [Source:HGNC Symbol;Acc:HGNC:17586] | 0.5979 | 2.73E-19 |
| ENSG00000072778 | ACADVL | acyl-CoA dehydrogenase very long chain [Source:HGNC Symbol;Acc:HGNC:92] | 0.5987 | 2.73E-19 |
| ENSG00000110107 | PRPF19 | pre-mRNA processing factor 19 [Source:HGNC Symbol;Acc:HGNC:17896] | 0.5048 | 2.73E-19 |
| ENSG00000160867 | FGFR4 | fibroblast growth factor receptor 4 [Source:HGNC Symbol;Acc:HGNC:3691] | -1.4012 | 2.74E-19 |
| ENSG00000175643 | RMI2 | RecQ mediated genome instability 2 [Source:HGNC Symbol;Acc:HGNC:28349] | -1.0173 | 3.14E-19 |
| ENSG00000270112 | NA | novel transcript, antisense to ST8SIA5 | -1.5064 | 3.15E-19 |

| | | | | |
|---|---|---|---|---|
| ENSG00000079332 | SAR1A | secretion associated Ras related GTPase 1A [Source:HGNC Symbol;Acc:HGNC:10534] | 0.6138 | 3.29E-19 |
| ENSG00000131669 | NINJ1 | ninjurin 1 [Source:HGNC Symbol;Acc:HGNC:7824] | -0.9869 | 3.92E-19 |
| ENSG00000146242 | TPBG | trophoblast glycoprotein [Source:HGNC Symbol;Acc:HGNC:12004] | 0.9401 | 4.12E-19 |
| ENSG00000104131 | EIF3J | eukaryotic translation initiation factor 3 subunit J [Source:HGNC Symbol;Acc:HGNC:3270] | 0.7565 | 4.14E-19 |
| ENSG00000102158 | MAGT1 | magnesium transporter 1 [Source:HGNC Symbol;Acc:HGNC:28880] | 0.9959 | 4.58E-19 |
| ENSG00000111885 | MAN1A1 | mannosidase alpha class 1A member 1 [Source:HGNC Symbol;Acc:HGNC:6821] | 3.5442 | 5.03E-19 |
| ENSG00000145604 | SKP2 | S-phase kinase associated protein 2 [Source:HGNC Symbol;Acc:HGNC:10901] | -0.6739 | 5.03E-19 |
| ENSG00000118432 | CNR1 | cannabinoid receptor 1 [Source:HGNC Symbol;Acc:HGNC:2159] | -1.1256 | 5.14E-19 |
| ENSG00000115970 | THADA | THADA armadillo repeat containing [Source:HGNC Symbol;Acc:HGNC:19217] | 0.9884 | 5.86E-19 |
| ENSG00000104722 | NEFM | neurofilament medium [Source:HGNC Symbol;Acc:HGNC:7734] | -1.1962 | 6.02E-19 |
| ENSG00000144746 | ARL6IP5 | ADP ribosylation factor like GTPase 6 interacting protein 5 [Source:HGNC Symbol;Acc:HGNC:16937] | -0.7365 | 6.04E-19 |
| ENSG00000088305 | DNMT3B | DNA methyltransferase 3 beta [Source:HGNC Symbol;Acc:HGNC:2979] | -2.1099 | 6.05E-19 |
| ENSG00000142867 | BCL10 | BCL10 immune signaling adaptor [Source:HGNC Symbol;Acc:HGNC:989] | 0.8983 | 6.07E-19 |
| ENSG00000109956 | B3GAT1 | beta-1,3-glucuronyltransferase 1 [Source:HGNC Symbol;Acc:HGNC:921] | 1.7409 | 6.17E-19 |
| ENSG00000096092 | TMEM14A | transmembrane protein 14A [Source:HGNC Symbol;Acc:HGNC:21076] | -0.9037 | 7.38E-19 |
| ENSG00000185909 | KLHDC8B | kelch domain containing 8B [Source:HGNC Symbol;Acc:HGNC:28557] | 0.8963 | 7.68E-19 |
| ENSG00000170271 | FAXDC2 | fatty acid hydroxylase domain containing 2 [Source:HGNC Symbol;Acc:HGNC:1334] | -1.8389 | 7.87E-19 |
| ENSG00000099814 | CEP170B | centrosomal protein 170B [Source:HGNC Symbol;Acc:HGNC:20362] | 1.2102 | 8.09E-19 |
| ENSG00000177034 | MTX3 | metaxin 3 [Source:HGNC Symbol;Acc:HGNC:24812] | 0.7414 | 8.12E-19 |
| ENSG00000118816 | CCNI | cyclin I [Source:HGNC Symbol;Acc:HGNC:1595] | 0.6120 | 8.14E-19 |
| ENSG00000095380 | NANS | N-acetylneuraminate synthase [Source:HGNC Symbol;Acc:HGNC:19237] | 0.7811 | 9.18E-19 |
| ENSG00000279606 | NA | TEC | -3.5226 | 9.21E-19 |
| ENSG00000096070 | BRPF3 | bromodomain and PHD finger containing 3 [Source:HGNC Symbol;Acc:HGNC:14256] | -0.7301 | 9.69E-19 |
| ENSG00000092330 | TINF2 | TERF1 interacting nuclear factor 2 [Source:HGNC Symbol;Acc:HGNC:11824] | 0.7218 | 9.83E-19 |
| ENSG00000141510 | TP53 | tumor protein p53 [Source:HGNC Symbol;Acc:HGNC:11998] | -0.7479 | 1.14E-18 |
| ENSG00000286696 | NA | novel transcript, antisense to SEMA7A | 1.8806 | 1.16E-18 |
| ENSG00000116406 | EDEM3 | ER degradation enhancing alpha-mannosidase like protein 3 [Source:HGNC Symbol;Acc:HGNC:16787] | 0.7154 | 1.35E-18 |
| ENSG00000142549 | IGLON5 | IgLON family member 5 [Source:HGNC Symbol;Acc:HGNC:34550] | -0.9071 | 1.35E-18 |
| ENSG00000213190 | MLLT11 | MLLT11 transcription factor 7 cofactor [Source:HGNC Symbol;Acc:HGNC:16997] | 0.8651 | 2.02E-18 |
| ENSG00000249378 | LINC01060 | long intergenic non-protein coding RNA 1060 [Source:HGNC Symbol;Acc:HGNC:49081] | 2.3295 | 2.03E-18 |
| ENSG00000226950 | DANCR | differentiation antagonizing non-protein coding RNA [Source:HGNC Symbol;Acc:HGNC:28964] | 0.8807 | 2.24E-18 |
| ENSG00000185338 | SOCS1 | suppressor of cytokine signaling 1 [Source:HGNC Symbol;Acc:HGNC:19383] | 1.5654 | 2.37E-18 |
| ENSG00000184602 | SNN | stannin [Source:HGNC Symbol;Acc:HGNC:11149] | -1.0646 | 2.42E-18 |
| ENSG00000150630 | VEGFC | vascular endothelial growth factor C [Source:HGNC Symbol;Acc:HGNC:12682] | 1.9426 | 2.47E-18 |
| ENSG00000103381 | CPPED1 | calcineurin like phosphoesterase domain containing 1 [Source:HGNC Symbol;Acc:HGNC:25632] | -1.1152 | 2.49E-18 |
| ENSG00000005175 | RPAP3 | RNA polymerase II associated protein 3 [Source:HGNC Symbol;Acc:HGNC:26151] | 0.8458 | 2.57E-18 |
| ENSG00000189337 | KAZN | kazrin, periplakin interacting protein [Source:HGNC Symbol;Acc:HGNC:29173] | 1.2578 | 2.62E-18 |
| ENSG00000022267 | FHL1 | four and a half LIM domains 1 [Source:HGNC Symbol;Acc:HGNC:3702] | 1.0755 | 2.65E-18 |
| ENSG00000168675 | LDLRAD4 | low density lipoprotein receptor class A domain containing 4 [Source:HGNC Symbol;Acc:HGNC:1224] | 1.8875 | 2.97E-18 |
| ENSG00000244509 | APOBEC3C | apolipoprotein B mRNA editing enzyme catalytic subunit 3C [Source:HGNC Symbol;Acc:HGNC:17353] | -0.9953 | 3.12E-18 |
| ENSG00000065548 | ZC3H15 | zinc finger CCCH-type containing 15 [Source:HGNC Symbol;Acc:HGNC:29528] | 0.7126 | 3.24E-18 |
| ENSG00000185359 | HGS | hepatocyte growth factor-regulated tyrosine kinase substrate [Source:HGNC Symbol;Acc:HGNC:4897] | 0.7613 | 3.34E-18 |
| ENSG00000243766 | HOTTIP | HOXA distal transcript antisense RNA [Source:HGNC Symbol;Acc:HGNC:37461] | 1.4291 | 3.63E-18 |

| | | | | |
|---|---|---|---|---|
| ENSG00000116455 | WDR77 | WD repeat domain 77 [Source:HGNC Symbol;Acc:HGNC:29652] | 0.6697 | 3.88E-18 |
| ENSG00000131374 | TBC1D5 | TBC1 domain family member 5 [Source:HGNC Symbol;Acc:HGNC:19166] | -0.9321 | 3.93E-18 |
| ENSG00000182168 | UNC5C | unc-5 netrin receptor C [Source:HGNC Symbol;Acc:HGNC:12569] | -1.0760 | 3.93E-18 |
| ENSG00000080845 | DLGAP4 | DLG associated protein 4 [Source:HGNC Symbol;Acc:HGNC:24476] | 0.8000 | 3.93E-18 |
| ENSG00000247796 | NA | novel transcript, antisense to MOCS2 | -1.5711 | 3.96E-18 |
| ENSG00000116815 | CD58 | CD58 molecule [Source:HGNC Symbol;Acc:HGNC:1688] | 1.3798 | 4.03E-18 |
| ENSG00000104886 | PLEKHJ1 | pleckstrin homology domain containing J1 [Source:HGNC Symbol;Acc:HGNC:18211] | -0.6947 | 4.03E-18 |
| ENSG00000103994 | ZNF106 | zinc finger protein 106 [Source:HGNC Symbol;Acc:HGNC:12886] | 0.8190 | 4.05E-18 |
| ENSG00000088543 | C3orf18 | chromosome 3 open reading frame 18 [Source:HGNC Symbol;Acc:HGNC:24837] | -1.1419 | 4.10E-18 |
| ENSG00000076248 | UNG | uracil DNA glycosylase [Source:HGNC Symbol;Acc:HGNC:12572] | -0.6514 | 4.75E-18 |
| ENSG00000198887 | SMC5 | structural maintenance of chromosomes 5 [Source:HGNC Symbol;Acc:HGNC:20465] | 0.7655 | 5.03E-18 |
| ENSG00000125648 | SLC25A23 | solute carrier family 25 member 23 [Source:HGNC Symbol;Acc:HGNC:19375] | -1.0779 | 5.52E-18 |
| ENSG00000135655 | USP15 | ubiquitin specific peptidase 15 [Source:HGNC Symbol;Acc:HGNC:12613] | 0.7151 | 5.52E-18 |
| ENSG00000127314 | RAP1B | RAP1B, member of RAS oncogene family [Source:HGNC Symbol;Acc:HGNC:9857] | 0.6169 | 5.92E-18 |
| ENSG00000105398 | SULT2A1 | sulfotransferase family 2A member 1 [Source:HGNC Symbol;Acc:HGNC:11458] | -1.3659 | 6.01E-18 |
| ENSG00000177854 | TMEM187 | transmembrane protein 187 [Source:HGNC Symbol;Acc:HGNC:13705] | -1.2165 | 6.85E-18 |
| ENSG00000162636 | FAM102B | family with sequence similarity 102 member B [Source:HGNC Symbol;Acc:HGNC:27637] | -1.2167 | 6.86E-18 |
| ENSG00000138639 | ARHGAP24 | Rho GTPase activating protein 24 [Source:HGNC Symbol;Acc:HGNC:25361] | 1.1511 | 6.97E-18 |
| ENSG00000011009 | LYPLA2 | lysophospholipase 2 [Source:HGNC Symbol;Acc:HGNC:6738] | 0.8241 | 7.28E-18 |
| ENSG00000122965 | RBM19 | RNA binding motif protein 19 [Source:HGNC Symbol;Acc:HGNC:29098] | 0.7855 | 7.34E-18 |
| ENSG00000146021 | KLHL3 | kelch like family member 3 [Source:HGNC Symbol;Acc:HGNC:6354] | 1.2298 | 7.49E-18 |
| ENSG00000197157 | SND1 | staphylococcal nuclease and tudor domain containing 1 [Source:HGNC Symbol;Acc:HGNC:30646] | 0.6711 | 7.54E-18 |
| ENSG00000183654 | MARCHF11 | membrane associated ring-CH-type finger 11 [Source:HGNC Symbol;Acc:HGNC:33609] | 1.3650 | 8.84E-18 |
| ENSG00000168610 | STAT3 | signal transducer and activator of transcription 3 [Source:HGNC Symbol;Acc:HGNC:11364] | 1.4096 | 9.67E-18 |
| ENSG00000105516 | DBP | D-box binding PAR bZIP transcription factor [Source:HGNC Symbol;Acc:HGNC:2697] | -1.6355 | 9.95E-18 |
| ENSG00000129562 | DAD1 | defender against cell death 1 [Source:HGNC Symbol;Acc:HGNC:2664] | 0.5977 | 1.10E-17 |
| ENSG00000185950 | IRS2 | insulin receptor substrate 2 [Source:HGNC Symbol;Acc:HGNC:6126] | 1.0322 | 1.11E-17 |
| ENSG00000196924 | FLNA | filamin A [Source:HGNC Symbol;Acc:HGNC:3754] | 1.0552 | 1.15E-17 |
| ENSG00000137869 | CYP19A1 | cytochrome P450 family 19 subfamily A member 1 [Source:HGNC Symbol;Acc:HGNC:2594] | 3.6037 | 1.15E-17 |
| ENSG00000136828 | RALGPS1 | Ral GEF with PH domain and SH3 binding motif 1 [Source:HGNC Symbol;Acc:HGNC:16851] | -0.9785 | 1.19E-17 |
| ENSG00000086991 | NOX4 | NADPH oxidase 4 [Source:HGNC Symbol;Acc:HGNC:7891] | 2.4244 | 1.19E-17 |
| ENSG00000167705 | RILP | Rab interacting lysosomal protein [Source:HGNC Symbol;Acc:HGNC:30266] | 2.9864 | 1.19E-17 |
| ENSG00000277561 | GOLGA8IP | golgin A8 family member I, pseudogene [Source:HGNC Symbol;Acc:HGNC:26660] | -1.7753 | 1.21E-17 |
| ENSG00000154640 | BTG3 | BTG anti-proliferation factor 3 [Source:HGNC Symbol;Acc:HGNC:1132] | 1.3526 | 1.25E-17 |
| ENSG00000162415 | ZSWIM5 | zinc finger SWIM-type containing 5 [Source:HGNC Symbol;Acc:HGNC:29299] | 0.6359 | 1.25E-17 |
| ENSG00000118200 | CAMSAP2 | calmodulin regulated spectrin associated protein family member 2 [Source:HGNC Symbol;Acc:HGNC:29188] | 0.5715 | 1.26E-17 |
| ENSG00000204590 | GNL1 | G protein nucleolar 1 (putative) [Source:HGNC Symbol;Acc:HGNC:4413] | 0.7613 | 1.28E-17 |
| ENSG00000179431 | FJX1 | four-jointed box kinase 1 [Source:HGNC Symbol;Acc:HGNC:17166] | 1.7577 | 1.33E-17 |
| ENSG00000063587 | ZNF275 | zinc finger protein 275 [Source:HGNC Symbol;Acc:HGNC:13069] | 0.5215 | 1.43E-17 |
| ENSG00000182796 | TMEM198B | transmembrane protein 198B (pseudogene) [Source:HGNC Symbol;Acc:HGNC:43629] | -0.9186 | 1.43E-17 |
| ENSG00000231890 | DARS-AS1 | DARS antisense RNA 1 [Source:HGNC Symbol;Acc:HGNC:40170] | 1.4524 | 1.50E-17 |
| ENSG00000138768 | USO1 | USO1 vesicle transport factor [Source:HGNC Symbol;Acc:HGNC:30904] | 0.9785 | 1.57E-17 |
| ENSG00000124588 | NQO2 | N-ribosyldihydronicotinamide:quinone reductase 2 [Source:HGNC Symbol;Acc:HGNC:7856] | 1.1890 | 1.63E-17 |

| | | | | |
|---|---|---|---|---|
| ENSG00000173744 | AGFG1 | ArfGAP with FG repeats 1 [Source:HGNC Symbol;Acc:HGNC:5175] | 0.6835 | 1.66E-17 |
| ENSG00000168389 | MFSD2A | major facilitator superfamily domain containing 2A [Source:HGNC Symbol;Acc:HGNC:25897] | 2.8677 | 1.68E-17 |
| ENSG00000189050 | RNFT1 | ring finger protein, transmembrane 1 [Source:HGNC Symbol;Acc:HGNC:30206] | 1.5517 | 1.68E-17 |
| ENSG00000164761 | TNFRSF11B | TNF receptor superfamily member 11b [Source:HGNC Symbol;Acc:HGNC:11909] | 1.4614 | 1.71E-17 |
| ENSG00000145779 | TNFAIP8 | TNF alpha induced protein 8 [Source:HGNC Symbol;Acc:HGNC:17260] | 1.6227 | 1.71E-17 |
| ENSG00000148498 | PARD3 | par-3 family cell polarity regulator [Source:HGNC Symbol;Acc:HGNC:16051] | 1.0184 | 1.76E-17 |
| ENSG00000198369 | SPRED2 | sprouty related EVH1 domain containing 2 [Source:HGNC Symbol;Acc:HGNC:17722] | 1.7509 | 1.90E-17 |
| ENSG00000135404 | CD63 | CD63 molecule [Source:HGNC Symbol;Acc:HGNC:1692] | 0.8242 | 1.93E-17 |
| ENSG00000121440 | PDZRN3 | PDZ domain containing ring finger 3 [Source:HGNC Symbol;Acc:HGNC:17704] | -0.9769 | 1.96E-17 |
| ENSG00000184897 | H1-10 | H1.10 linker histone [Source:HGNC Symbol;Acc:HGNC:4722] | -0.9391 | 2.00E-17 |
| ENSG00000160767 | FAM189B | family with sequence similarity 189 member B [Source:HGNC Symbol;Acc:HGNC:1233] | -0.6810 | 2.09E-17 |
| ENSG00000151790 | TDO2 | tryptophan 2,3-dioxygenase [Source:HGNC Symbol;Acc:HGNC:11708] | 1.6341 | 2.11E-17 |
| ENSG00000165702 | GFI1B | growth factor independent 1B transcriptional repressor [Source:HGNC Symbol;Acc:HGNC:4238] | -4.0252 | 2.19E-17 |
| ENSG00000256967 | NA | novel transcript | 2.5921 | 2.24E-17 |
| ENSG00000095139 | ARCN1 | archain 1 [Source:HGNC Symbol;Acc:HGNC:649] | 0.5912 | 2.36E-17 |
| ENSG00000176654 | NANOGP1 | Nanog homeobox pseudogene 1 [Source:HGNC Symbol;Acc:HGNC:23099] | -2.2431 | 2.36E-17 |
| ENSG00000086598 | TMED2 | transmembrane p24 trafficking protein 2 [Source:HGNC Symbol;Acc:HGNC:16996] | 0.6068 | 2.57E-17 |
| ENSG00000064309 | CDON | cell adhesion associated, oncogene regulated [Source:HGNC Symbol;Acc:HGNC:17104] | -1.8407 | 2.59E-17 |
| ENSG00000115738 | ID2 | inhibitor of DNA binding 2 [Source:HGNC Symbol;Acc:HGNC:5361] | -1.0067 | 2.66E-17 |
| ENSG00000164828 | SUN1 | Sad1 and UNC84 domain containing 1 [Source:HGNC Symbol;Acc:HGNC:18587] | 0.6599 | 2.92E-17 |
| ENSG00000135541 | AHI1 | Abelson helper integration site 1 [Source:HGNC Symbol;Acc:HGNC:21575] | 1.0348 | 2.99E-17 |
| ENSG00000077585 | GPR137B | G protein-coupled receptor 137B [Source:HGNC Symbol;Acc:HGNC:11862] | 1.6611 | 3.10E-17 |
| ENSG00000058262 | SEC61A1 | SEC61 translocon subunit alpha 1 [Source:HGNC Symbol;Acc:HGNC:18276] | 0.8069 | 3.33E-17 |
| ENSG00000188766 | SPRED3 | sprouty related EVH1 domain containing 3 [Source:HGNC Symbol;Acc:HGNC:31041] | 2.0030 | 3.33E-17 |
| ENSG00000196937 | FAM3C | FAM3 metabolism regulating signaling molecule C [Source:HGNC Symbol;Acc:HGNC:18664] | 0.7280 | 3.33E-17 |
| ENSG00000186469 | GNG2 | G protein subunit gamma 2 [Source:HGNC Symbol;Acc:HGNC:4404] | -1.1743 | 3.76E-17 |
| ENSG00000137404 | NRM | nurim [Source:HGNC Symbol;Acc:HGNC:8003] | -0.9392 | 3.77E-17 |
| ENSG00000112096 | SOD2 | superoxide dismutase 2 [Source:HGNC Symbol;Acc:HGNC:11180] | -0.7331 | 3.98E-17 |
| ENSG00000122884 | P4HA1 | prolyl 4-hydroxylase subunit alpha 1 [Source:HGNC Symbol;Acc:HGNC:8546] | 1.3632 | 4.36E-17 |
| ENSG00000113163 | CERT1 | ceramide transporter 1 [Source:HGNC Symbol;Acc:HGNC:2205] | -0.7713 | 4.59E-17 |
| ENSG00000188158 | NHS | NHS actin remodeling regulator [Source:HGNC Symbol;Acc:HGNC:7820] | -0.9610 | 5.65E-17 |
| ENSG00000168282 | MGAT2 | alpha-1,6-mannosyl-glycoprotein 2-beta-N-acetylglucosaminyltransferase [Source:HGNC Symbol;Acc:HGNC:7045] | 0.7171 | 6.04E-17 |
| ENSG00000165475 | CRYL1 | crystallin lambda 1 [Source:HGNC Symbol;Acc:HGNC:18246] | -1.3736 | 6.22E-17 |
| ENSG00000137955 | RABGGTB | Rab geranylgeranyltransferase subunit beta [Source:HGNC Symbol;Acc:HGNC:9796] | 0.8265 | 6.42E-17 |
| ENSG00000204876 | NA | uncharacterized LOC389602 [Source:NCBI gene (formerly Entrezgene);Acc:389602] | 1.8032 | 6.45E-17 |
| ENSG00000006128 | TAC1 | tachykinin precursor 1 [Source:HGNC Symbol;Acc:HGNC:11517] | 8.1892 | 6.54E-17 |
| ENSG00000165406 | MARCHF8 | membrane associated ring-CH-type finger 8 [Source:HGNC Symbol;Acc:HGNC:23356] | -0.7974 | 6.64E-17 |
| ENSG00000152527 | PLEKHH2 | pleckstrin homology, MyTH4 and FERM domain containing H2 [Source:HGNC Symbol;Acc:HGNC:30506] | 1.1592 | 6.70E-17 |
| ENSG00000052749 | RRP12 | ribosomal RNA processing 12 homolog [Source:HGNC Symbol;Acc:HGNC:29100] | 1.0991 | 6.72E-17 |
| ENSG00000150990 | DHX37 | DEAH-box helicase 37 [Source:HGNC Symbol;Acc:HGNC:17210] | 0.6917 | 6.76E-17 |
| ENSG00000269416 | LINC01224 | long intergenic non-protein coding RNA 1224 [Source:HGNC Symbol;Acc:HGNC:49676] | -1.9174 | 6.83E-17 |
| ENSG00000073849 | ST6GAL1 | ST6 beta-galactoside alpha-2,6-sialyltransferase 1 [Source:HGNC Symbol;Acc:HGNC:10860] | -1.4216 | 7.22E-17 |
| ENSG00000163833 | FBXO40 | F-box protein 40 [Source:HGNC Symbol;Acc:HGNC:29816] | -1.8941 | 7.62E-17 |

| | | | | |
|---|---|---|---|---|
| ENSG00000027075 | PRKCH | protein kinase C eta [Source:HGNC Symbol;Acc:HGNC:9403] | 2.1541 | 7.98E-17 |
| ENSG00000111450 | STX2 | syntaxin 2 [Source:HGNC Symbol;Acc:HGNC:3403] | 1.1196 | 7.99E-17 |
| ENSG00000108021 | TASOR2 | transcription activation suppressor family member 2 [Source:HGNC Symbol;Acc:HGNC:23484] | 0.7995 | 8.07E-17 |
| ENSG00000260285 | NA | novel transcript, antisense to TRMT61A | -1.1834 | 8.23E-17 |
| ENSG00000105642 | KCNN1 | potassium calcium-activated channel subfamily N member 1 [Source:HGNC Symbol;Acc:HGNC:6290] | -1.7487 | 8.34E-17 |
| ENSG00000155966 | AFF2 | AF4/FMR2 family member 2 [Source:HGNC Symbol;Acc:HGNC:3776] | 1.1920 | 8.34E-17 |
| ENSG00000130402 | ACTN4 | actinin alpha 4 [Source:HGNC Symbol;Acc:HGNC:166] | 0.8284 | 8.54E-17 |
| ENSG00000087085 | ACHE | acetylcholinesterase (Cartwright blood group) [Source:HGNC Symbol;Acc:HGNC:108] | 2.0834 | 8.72E-17 |
| ENSG00000107679 | PLEKHA1 | pleckstrin homology domain containing A1 [Source:HGNC Symbol;Acc:HGNC:14335] | 0.9781 | 8.98E-17 |
| ENSG00000145029 | NICN1 | nicolin 1 [Source:HGNC Symbol;Acc:HGNC:18317] | -1.3238 | 9.00E-17 |
| ENSG00000143624 | INTS3 | integrator complex subunit 3 [Source:HGNC Symbol;Acc:HGNC:26153] | 0.9638 | 9.39E-17 |
| ENSG00000128595 | CALU | calumenin [Source:HGNC Symbol;Acc:HGNC:1458] | 0.7547 | 9.69E-17 |
| ENSG00000139637 | MYG1 | MYG1 exonuclease [Source:HGNC Symbol;Acc:HGNC:17590] | 0.7085 | 9.71E-17 |
| ENSG00000136930 | PSMB7 | proteasome 20S subunit beta 7 [Source:HGNC Symbol;Acc:HGNC:9544] | 0.5876 | 1.05E-16 |
| ENSG00000198464 | ZNF480 | zinc finger protein 480 [Source:HGNC Symbol;Acc:HGNC:23305] | -1.9435 | 1.07E-16 |
| ENSG00000163377 | TAFA4 | TAFA chemokine like family member 4 [Source:HGNC Symbol;Acc:HGNC:21591] | -1.4201 | 1.10E-16 |
| ENSG00000103342 | GSPT1 | G1 to S phase transition 1 [Source:HGNC Symbol;Acc:HGNC:4621] | 0.6499 | 1.10E-16 |
| ENSG00000197322 | C17orf102 | chromosome 17 open reading frame 102 [Source:HGNC Symbol;Acc:HGNC:34412] | 1.2991 | 1.14E-16 |
| ENSG00000197077 | KIAA1671 | KIAA1671 [Source:HGNC Symbol;Acc:HGNC:29345] | 0.6758 | 1.16E-16 |
| ENSG00000173786 | CNP | 2',3'-cyclic nucleotide 3' phosphodiesterase [Source:HGNC Symbol;Acc:HGNC:2158] | -0.6627 | 1.18E-16 |
| ENSG00000160233 | LRRC3 | leucine rich repeat containing 3 [Source:HGNC Symbol;Acc:HGNC:14965] | -1.3684 | 1.19E-16 |
| ENSG00000141448 | GATA6 | GATA binding protein 6 [Source:HGNC Symbol;Acc:HGNC:4174] | 1.5146 | 1.21E-16 |
| ENSG00000256664 | NA | ribosomal L24 domain containing 1 (RSL24D1) pseudogene | 1.0590 | 1.21E-16 |
| ENSG00000257365 | FNTB | farnesyltransferase, CAAX box, beta [Source:HGNC Symbol;Acc:HGNC:3785] | 0.8474 | 1.22E-16 |
| ENSG00000143630 | HCN3 | hyperpolarization activated cyclic nucleotide gated potassium channel 3 [Source:HGNC Symbol;Acc:HGNC:19183] | -2.0828 | 1.23E-16 |
| ENSG00000123080 | CDKN2C | cyclin dependent kinase inhibitor 2C [Source:HGNC Symbol;Acc:HGNC:1789] | -0.9322 | 1.26E-16 |
| ENSG00000135338 | LCA5 | lebercilin LCA5 [Source:HGNC Symbol;Acc:HGNC:31923] | 0.9905 | 1.33E-16 |
| ENSG00000150347 | ARID5B | AT-rich interaction domain 5B [Source:HGNC Symbol;Acc:HGNC:17362] | 0.9227 | 1.39E-16 |
| ENSG00000177732 | SOX12 | SRY-box transcription factor 12 [Source:HGNC Symbol;Acc:HGNC:11198] | -1.0574 | 1.39E-16 |
| ENSG00000243364 | EFNA4 | ephrin A4 [Source:HGNC Symbol;Acc:HGNC:3224] | -1.2265 | 1.39E-16 |
| ENSG00000126733 | DACH2 | dachshund family transcription factor 2 [Source:HGNC Symbol;Acc:HGNC:16814] | 1.8317 | 1.51E-16 |
| ENSG00000137312 | FLOT1 | flotillin 1 [Source:HGNC Symbol;Acc:HGNC:3757] | 0.8543 | 1.58E-16 |
| ENSG00000106236 | NPTX2 | neuronal pentraxin 2 [Source:HGNC Symbol;Acc:HGNC:7953] | 1.3815 | 1.78E-16 |
| ENSG00000113790 | EHHADH | enoyl-CoA hydratase and 3-hydroxyacyl CoA dehydrogenase [Source:HGNC Symbol;Acc:HGNC:3247] | -1.1178 | 1.83E-16 |
| ENSG00000134480 | CCNH | cyclin H [Source:HGNC Symbol;Acc:HGNC:1594] | 0.7207 | 1.89E-16 |
| ENSG00000139318 | DUSP6 | dual specificity phosphatase 6 [Source:HGNC Symbol;Acc:HGNC:3072] | 2.1505 | 1.96E-16 |
| ENSG00000089041 | P2RX7 | purinergic receptor P2X 7 [Source:HGNC Symbol;Acc:HGNC:8537] | -1.7915 | 1.97E-16 |
| ENSG00000205309 | NT5M | 5',3'-nucleotidase, mitochondrial [Source:HGNC Symbol;Acc:HGNC:15769] | -1.7246 | 2.10E-16 |
| ENSG00000167994 | RAB3IL1 | RAB3A interacting protein like 1 [Source:HGNC Symbol;Acc:HGNC:9780] | -1.4781 | 2.15E-16 |
| ENSG00000135678 | CPM | carboxypeptidase M [Source:HGNC Symbol;Acc:HGNC:2311] | -1.5942 | 2.20E-16 |
| ENSG00000170390 | DCLK2 | doublecortin like kinase 2 [Source:HGNC Symbol;Acc:HGNC:19002] | -0.9191 | 2.25E-16 |
| ENSG00000133059 | DSTYK | dual serine/threonine and tyrosine protein kinase [Source:HGNC Symbol;Acc:HGNC:29043] | -0.7138 | 2.40E-16 |
| ENSG00000182667 | NTM | neurotrimin [Source:HGNC Symbol;Acc:HGNC:17941] | -1.0494 | 2.40E-16 |

| | | | | |
|---|---|---|---|---|
| ENSG00000177119 | ANO6 | anoctamin 6 [Source:HGNC Symbol;Acc:HGNC:25240] | 0.6654 | 2.68E-16 |
| ENSG00000198431 | TXNRD1 | thioredoxin reductase 1 [Source:HGNC Symbol;Acc:HGNC:12437] | 0.9731 | 2.94E-16 |
| ENSG00000159387 | IRX6 | iroquois homeobox 6 [Source:HGNC Symbol;Acc:HGNC:14675] | -1.3920 | 3.09E-16 |
| ENSG00000184588 | PDE4B | phosphodiesterase 4B [Source:HGNC Symbol;Acc:HGNC:8781] | 1.2499 | 3.24E-16 |
| ENSG00000185238 | PRMT3 | protein arginine methyltransferase 3 [Source:HGNC Symbol;Acc:HGNC:30163] | 1.2462 | 3.45E-16 |
| ENSG00000197296 | FITM2 | fat storage inducing transmembrane protein 2 [Source:HGNC Symbol;Acc:HGNC:16135] | -0.8136 | 3.46E-16 |
| ENSG00000091428 | RAPGEF4 | Rap guanine nucleotide exchange factor 4 [Source:HGNC Symbol;Acc:HGNC:16626] | 2.2074 | 3.47E-16 |
| ENSG00000105647 | PIK3R2 | phosphoinositide-3-kinase regulatory subunit 2 [Source:HGNC Symbol;Acc:HGNC:8980] | -0.8497 | 3.84E-16 |
| ENSG00000128791 | TWSG1 | twisted gastrulation BMP signaling modulator 1 [Source:HGNC Symbol;Acc:HGNC:12429] | -0.7860 | 3.94E-16 |
| ENSG00000140948 | ZCCHC14 | zinc finger CCHC-type containing 14 [Source:HGNC Symbol;Acc:HGNC:24134] | -0.7263 | 4.04E-16 |
| ENSG00000008441 | NFIX | nuclear factor I X [Source:HGNC Symbol;Acc:HGNC:7788] | -0.8959 | 4.82E-16 |
| ENSG00000158792 | SPATA2L | spermatogenesis associated 2 like [Source:HGNC Symbol;Acc:HGNC:28393] | 1.0760 | 5.09E-16 |
| ENSG00000110321 | EIF4G2 | eukaryotic translation initiation factor 4 gamma 2 [Source:HGNC Symbol;Acc:HGNC:3297] | 0.6212 | 5.39E-16 |
| ENSG00000109689 | STIM2 | stromal interaction molecule 2 [Source:HGNC Symbol;Acc:HGNC:19205] | 1.2530 | 5.40E-16 |
| ENSG00000148154 | UGCG | UDP-glucose ceramide glucosyltransferase [Source:HGNC Symbol;Acc:HGNC:12524] | 0.9677 | 5.40E-16 |
| ENSG00000137815 | RTF1 | RTF1 homolog, Paf1/RNA polymerase II complex component [Source:HGNC Symbol;Acc:HGNC:28996] | -0.5169 | 5.54E-16 |
| ENSG00000080503 | SMARCA2 | SWI/SNF related, matrix associated, actin dependent regulator of chromatin, subfamily a, member 2 [Source:HGNC Symbol;Acc:HGNC:11098] | -0.9341 | 5.93E-16 |
| ENSG00000168040 | FADD | Fas associated via death domain [Source:HGNC Symbol;Acc:HGNC:3573] | 0.6769 | 5.93E-16 |
| ENSG00000026950 | BTN3A1 | butyrophilin subfamily 3 member A1 [Source:HGNC Symbol;Acc:HGNC:1138] | -1.0183 | 6.09E-16 |
| ENSG00000142599 | RERE | arginine-glutamic acid dipeptide repeats [Source:HGNC Symbol;Acc:HGNC:9965] | -0.8197 | 6.14E-16 |
| ENSG00000141542 | RAB40B | RAB40B, member RAS oncogene family [Source:HGNC Symbol;Acc:HGNC:18284] | -1.7030 | 6.41E-16 |
| ENSG00000198598 | MMP17 | matrix metallopeptidase 17 [Source:HGNC Symbol;Acc:HGNC:7163] | 2.9405 | 7.04E-16 |
| ENSG00000141367 | CLTC | clathrin heavy chain [Source:HGNC Symbol;Acc:HGNC:2092] | 0.5638 | 7.11E-16 |
| ENSG00000186212 | SOWAHB | sosondowah ankyrin repeat domain family member B [Source:HGNC Symbol;Acc:HGNC:32958] | 2.1070 | 7.17E-16 |
| ENSG00000185274 | GALNT17 | polypeptide N-acetylgalactosaminyltransferase 17 [Source:HGNC Symbol;Acc:HGNC:16347] | -0.9731 | 7.18E-16 |
| ENSG00000086848 | ALG9 | ALG9 alpha-1,2-mannosyltransferase [Source:HGNC Symbol;Acc:HGNC:15672] | 1.2966 | 7.52E-16 |
| ENSG00000100350 | FOXRED2 | FAD dependent oxidoreductase domain containing 2 [Source:HGNC Symbol;Acc:HGNC:26264] | -0.6539 | 7.61E-16 |
| ENSG00000102572 | STK24 | serine/threonine kinase 24 [Source:HGNC Symbol;Acc:HGNC:11403] | 0.8374 | 7.83E-16 |
| ENSG00000085563 | ABCB1 | ATP binding cassette subfamily B member 1 [Source:HGNC Symbol;Acc:HGNC:40] | 0.9228 | 8.08E-16 |
| ENSG00000160310 | PRMT2 | protein arginine methyltransferase 2 [Source:HGNC Symbol;Acc:HGNC:5186] | -0.5263 | 8.20E-16 |
| ENSG00000149932 | TMEM219 | transmembrane protein 219 [Source:HGNC Symbol;Acc:HGNC:25201] | -0.6064 | 8.49E-16 |
| ENSG00000254615 | NA | novel transcript | 2.6373 | 8.91E-16 |
| ENSG00000267432 | DNAH17-AS1 | DNAH17 antisense RNA 1 [Source:HGNC Symbol;Acc:HGNC:48594] | 2.2037 | 9.12E-16 |
| ENSG00000057608 | GDI2 | GDP dissociation inhibitor 2 [Source:HGNC Symbol;Acc:HGNC:4227] | 0.8126 | 9.39E-16 |
| ENSG00000068912 | ERLEC1 | endoplasmic reticulum lectin 1 [Source:HGNC Symbol;Acc:HGNC:25222] | 0.6922 | 9.54E-16 |
| ENSG00000175264 | CHST1 | carbohydrate sulfotransferase 1 [Source:HGNC Symbol;Acc:HGNC:1969] | 0.7865 | 9.85E-16 |
| ENSG00000100027 | YPEL1 | yippee like 1 [Source:HGNC Symbol;Acc:HGNC:12845] | -1.9737 | 9.88E-16 |
| ENSG00000176532 | PRR15 | proline rich 15 [Source:HGNC Symbol;Acc:HGNC:22310] | -1.4432 | 1.01E-15 |
| ENSG00000085741 | WNT11 | Wnt family member 11 [Source:HGNC Symbol;Acc:HGNC:12776] | -0.8748 | 1.11E-15 |
| ENSG00000100528 | CNIH1 | cornichon family AMPA receptor auxiliary protein 1 [Source:HGNC Symbol;Acc:HGNC:19431] | 0.6245 | 1.12E-15 |
| ENSG00000176170 | SPHK1 | sphingosine kinase 1 [Source:HGNC Symbol;Acc:HGNC:11240] | 3.0424 | 1.16E-15 |
| ENSG00000213523 | SRA1 | steroid receptor RNA activator 1 [Source:HGNC Symbol;Acc:HGNC:11281] | 0.6373 | 1.17E-15 |

| | | | | |
|---|---|---|---|---|
| ENSG00000004975 | DVL2 | dishevelled segment polarity protein 2 [Source:HGNC Symbol;Acc:HGNC:3086] | -0.6791 | 1.22E-15 |
| ENSG00000244968 | LIFR-AS1 | LIFR antisense RNA 1 [Source:HGNC Symbol;Acc:HGNC:43600] | -1.3685 | 1.34E-15 |
| ENSG00000251493 | FOXD1 | forkhead box D1 [Source:HGNC Symbol;Acc:HGNC:3802] | 1.2836 | 1.49E-15 |
| ENSG00000124145 | SDC4 | syndecan 4 [Source:HGNC Symbol;Acc:HGNC:10661] | 1.2801 | 1.59E-15 |
| ENSG00000176845 | METRNL | meteorin like, glial cell differentiation regulator [Source:HGNC Symbol;Acc:HGNC:27584] | 2.2326 | 1.60E-15 |
| ENSG00000154997 | SEPTIN14 | septin 14 [Source:HGNC Symbol;Acc:HGNC:33280] | -1.5612 | 1.66E-15 |
| ENSG00000050030 | NEXMIF | neurite extension and migration factor [Source:HGNC Symbol;Acc:HGNC:29433] | -2.2047 | 1.69E-15 |
| ENSG00000179142 | CYP11B2 | cytochrome P450 family 11 subfamily B member 2 [Source:HGNC Symbol;Acc:HGNC:2592] | 3.8198 | 1.71E-15 |
| ENSG00000145882 | PCYOX1L | prenylcysteine oxidase 1 like [Source:HGNC Symbol;Acc:HGNC:28477] | -0.7875 | 1.74E-15 |
| ENSG00000196376 | SLC35F1 | solute carrier family 35 member F1 [Source:HGNC Symbol;Acc:HGNC:21483] | 0.7272 | 1.76E-15 |
| ENSG00000111679 | PTPN6 | protein tyrosine phosphatase non-receptor type 6 [Source:HGNC Symbol;Acc:HGNC:9658] | -1.2303 | 1.78E-15 |
| ENSG00000154380 | ENAH | ENAH actin regulator [Source:HGNC Symbol;Acc:HGNC:18271] | 0.8922 | 1.85E-15 |
| ENSG00000153234 | NR4A2 | nuclear receptor subfamily 4 group A member 2 [Source:HGNC Symbol;Acc:HGNC:7981] | 4.2700 | 2.05E-15 |
| ENSG00000088538 | DOCK3 | dedicator of cytokinesis 3 [Source:HGNC Symbol;Acc:HGNC:2989] | -1.0657 | 2.06E-15 |
| ENSG00000197780 | TAF13 | TATA-box binding protein associated factor 13 [Source:HGNC Symbol;Acc:HGNC:11546] | 0.8648 | 2.08E-15 |
| ENSG00000138944 | SHISAL1 | shisa like 1 [Source:HGNC Symbol;Acc:HGNC:29335] | -1.1805 | 2.11E-15 |
| ENSG00000117133 | RPF1 | ribosome production factor 1 homolog [Source:HGNC Symbol;Acc:HGNC:30350] | 0.5234 | 2.17E-15 |
| ENSG00000112851 | ERBIN | erbb2 interacting protein [Source:HGNC Symbol;Acc:HGNC:15842] | 0.5200 | 2.19E-15 |
| ENSG00000023330 | ALAS1 | 5'-aminolevulinate synthase 1 [Source:HGNC Symbol;Acc:HGNC:396] | 0.6587 | 2.38E-15 |
| ENSG00000140545 | MFGE8 | milk fat globule EGF and factor V/VIII domain containing [Source:HGNC Symbol;Acc:HGNC:7036] | 0.8234 | 2.41E-15 |
| ENSG00000125835 | SNRPB | small nuclear ribonucleoprotein polypeptides B and B1 [Source:HGNC Symbol;Acc:HGNC:11153] | 0.5438 | 2.54E-15 |
| ENSG00000196151 | WDSUB1 | WD repeat, sterile alpha motif and U-box domain containing 1 [Source:HGNC Symbol;Acc:HGNC:26697] | -1.0447 | 2.54E-15 |
| ENSG00000127863 | TNFRSF19 | TNF receptor superfamily member 19 [Source:HGNC Symbol;Acc:HGNC:11915] | 0.8429 | 2.70E-15 |
| ENSG00000166979 | EVA1C | eva-1 homolog C [Source:HGNC Symbol;Acc:HGNC:13239] | 2.0414 | 2.97E-15 |
| ENSG00000144867 | SRPRB | SRP receptor subunit beta [Source:HGNC Symbol;Acc:HGNC:24085] | 0.8639 | 3.14E-15 |
| ENSG00000284691 | NA | novel zinc finger protein | -1.6120 | 3.16E-15 |
| ENSG00000080824 | HSP90AA1 | heat shock protein 90 alpha family class A member 1 [Source:HGNC Symbol;Acc:HGNC:5253] | 0.5011 | 3.38E-15 |
| ENSG00000134375 | TIMM17A | translocase of inner mitochondrial membrane 17A [Source:HGNC Symbol;Acc:HGNC:17315] | 0.5652 | 3.53E-15 |
| ENSG00000181751 | MACIR | macrophage immunometabolism regulator [Source:HGNC Symbol;Acc:HGNC:25052] | -0.8825 | 3.74E-15 |
| ENSG00000143153 | ATP1B1 | ATPase Na+/K+ transporting subunit beta 1 [Source:HGNC Symbol;Acc:HGNC:804] | 0.9158 | 3.99E-15 |
| ENSG00000112237 | CCNC | cyclin C [Source:HGNC Symbol;Acc:HGNC:1581] | 0.6999 | 4.07E-15 |
| ENSG00000239857 | GET4 | guided entry of tail-anchored proteins factor 4 [Source:HGNC Symbol;Acc:HGNC:21690] | 0.7575 | 4.32E-15 |
| ENSG00000117592 | PRDX6 | peroxiredoxin 6 [Source:HGNC Symbol;Acc:HGNC:16753] | 0.5688 | 4.33E-15 |
| ENSG00000147642 | SYBU | syntabulin [Source:HGNC Symbol;Acc:HGNC:26011] | 1.0901 | 4.39E-15 |
| ENSG00000221125 | NA | NA | -2.7560 | 4.51E-15 |
| ENSG00000164164 | OTUD4 | OTU deubiquitinase 4 [Source:HGNC Symbol;Acc:HGNC:24949] | 1.0145 | 4.94E-15 |
| ENSG00000137872 | SEMA6D | semaphorin 6D [Source:HGNC Symbol;Acc:HGNC:16770] | 1.6430 | 4.97E-15 |
| ENSG00000136052 | SLC41A2 | solute carrier family 41 member 2 [Source:HGNC Symbol;Acc:HGNC:31045] | 1.5831 | 4.98E-15 |
| ENSG00000232956 | SNHG15 | small nucleolar RNA host gene 15 [Source:HGNC Symbol;Acc:HGNC:27797] | 0.6822 | 5.05E-15 |
| ENSG00000103496 | STX4 | syntaxin 4 [Source:HGNC Symbol;Acc:HGNC:11439] | 0.7184 | 5.06E-15 |
| ENSG00000164081 | TEX264 | testis expressed 264, ER-phagy receptor [Source:HGNC Symbol;Acc:HGNC:30247] | -0.5354 | 5.20E-15 |
| ENSG00000171223 | JUNB | JunB proto-oncogene, AP-1 transcription factor subunit [Source:HGNC Symbol;Acc:HGNC:6205] | 1.8815 | 5.53E-15 |
| ENSG00000103018 | CYB5B | cytochrome b5 type B [Source:HGNC Symbol;Acc:HGNC:24374] | 0.7397 | 5.62E-15 |

| | | | | |
|---|---|---|---|---|
| ENSG00000124126 | PREX1 | phosphatidylinositol-3,4,5-trisphosphate dependent Rac exchange factor 1 [Source:HGNC Symbol;Acc:HGNC:32594] | -1.2010 | 5.90E-15 |
| ENSG00000147454 | SLC25A37 | solute carrier family 25 member 37 [Source:HGNC Symbol;Acc:HGNC:29786] | 0.7591 | 6.26E-15 |
| ENSG00000125753 | VASP | vasodilator stimulated phosphoprotein [Source:HGNC Symbol;Acc:HGNC:12652] | 0.9404 | 6.46E-15 |
| ENSG00000111653 | ING4 | inhibitor of growth family member 4 [Source:HGNC Symbol;Acc:HGNC:19423] | -0.8392 | 6.50E-15 |
| ENSG00000198856 | OSTC | oligosaccharyltransferase complex non-catalytic subunit [Source:HGNC Symbol;Acc:HGNC:24448] | 0.7334 | 6.52E-15 |
| ENSG00000136155 | SCEL | sciellin [Source:HGNC Symbol;Acc:HGNC:10573] | -3.3034 | 6.56E-15 |
| ENSG00000130429 | ARPC1B | actin related protein 2/3 complex subunit 1B [Source:HGNC Symbol;Acc:HGNC:704] | 1.1617 | 6.56E-15 |
| ENSG00000164125 | GASK1B | golgi associated kinase 1B [Source:HGNC Symbol;Acc:HGNC:25312] | -1.0066 | 6.84E-15 |
| ENSG00000198900 | 1,00 TOP | DNA topoisomerase I [Source:HGNC Symbol;Acc:HGNC:11986] | 0.6525 | 6.96E-15 |
| ENSG00000143409 | MINDY1 | MINDY lysine 48 deubiquitinase 1 [Source:HGNC Symbol;Acc:HGNC:25648] | -0.9910 | 7.97E-15 |
| ENSG00000066827 | ZFAT | zinc finger and AT-hook domain containing [Source:HGNC Symbol;Acc:HGNC:19899] | 1.3903 | 8.17E-15 |
| ENSG00000126457 | PRMT1 | protein arginine methyltransferase 1 [Source:HGNC Symbol;Acc:HGNC:5187] | 0.5387 | 8.64E-15 |
| ENSG00000147180 | ZNF711 | zinc finger protein 711 [Source:HGNC Symbol;Acc:HGNC:13128] | -0.7267 | 8.88E-15 |
| ENSG00000151320 | AKAP6 | A-kinase anchoring protein 6 [Source:HGNC Symbol;Acc:HGNC:376] | -1.6656 | 8.88E-15 |
| ENSG00000169925 | BRD3 | bromodomain containing 3 [Source:HGNC Symbol;Acc:HGNC:1104] | -0.9605 | 8.93E-15 |
| ENSG00000168404 | MLKL | mixed lineage kinase domain like pseudokinase [Source:HGNC Symbol;Acc:HGNC:26617] | 2.9752 | 9.77E-15 |
| ENSG00000102172 | SMS | spermine synthase [Source:HGNC Symbol;Acc:HGNC:11123] | -0.6240 | 9.84E-15 |
| ENSG00000134970 | TMED7 | transmembrane p24 trafficking protein 7 [Source:HGNC Symbol;Acc:HGNC:24253] | 0.7801 | 1.17E-14 |
| ENSG00000106689 | LHX2 | LIM homeobox 2 [Source:HGNC Symbol;Acc:HGNC:6594] | 1.8383 | 1.18E-14 |
| ENSG00000182934 | SRPRA | SRP receptor subunit alpha [Source:HGNC Symbol;Acc:HGNC:11307] | 0.5554 | 1.21E-14 |
| ENSG00000269893 | SNHG8 | small nucleolar RNA host gene 8 [Source:HGNC Symbol;Acc:HGNC:33098] | 0.6539 | 1.22E-14 |
| ENSG00000115233 | PSMD14 | proteasome 26S subunit, non-ATPase 14 [Source:HGNC Symbol;Acc:HGNC:16889] | 0.5346 | 1.24E-14 |
| ENSG00000174791 | RIN1 | Ras and Rab interactor 1 [Source:HGNC Symbol;Acc:HGNC:18749] | 2.4239 | 1.25E-14 |
| ENSG00000262001 | DLGAP1-AS2 | DLGAP1 antisense RNA 2 [Source:HGNC Symbol;Acc:HGNC:28146] | 1.1823 | 1.27E-14 |
| ENSG00000276023 | DUSP14 | dual specificity phosphatase 14 [Source:HGNC Symbol;Acc:HGNC:17007] | 0.8582 | 1.35E-14 |
| ENSG00000162409 | PRKAA2 | protein kinase AMP-activated catalytic subunit alpha 2 [Source:HGNC Symbol;Acc:HGNC:9377] | 1.2099 | 1.37E-14 |
| ENSG00000023902 | PLEKHO1 | pleckstrin homology domain containing O1 [Source:HGNC Symbol;Acc:HGNC:24310] | 1.5696 | 1.39E-14 |
| ENSG00000178764 | ZHX2 | zinc fingers and homeoboxes 2 [Source:HGNC Symbol;Acc:HGNC:18513] | 1.0580 | 1.43E-14 |
| ENSG00000204386 | NEU1 | neuraminidase 1 [Source:HGNC Symbol;Acc:HGNC:7758] | 0.9859 | 1.44E-14 |
| ENSG00000048162 | NOP16 | NOP16 nucleolar protein [Source:HGNC Symbol;Acc:HGNC:26934] | 0.7588 | 1.45E-14 |
| ENSG00000067177 | PHKA1 | phosphorylase kinase regulatory subunit alpha 1 [Source:HGNC Symbol;Acc:HGNC:8925] | 0.8259 | 1.47E-14 |
| ENSG00000102743 | SLC25A15 | solute carrier family 25 member 15 [Source:HGNC Symbol;Acc:HGNC:10985] | -0.7403 | 1.47E-14 |
| ENSG00000140650 | PMM2 | phosphomannomutase 2 [Source:HGNC Symbol;Acc:HGNC:9115] | 0.8965 | 1.50E-14 |
| ENSG00000160404 | TOR2A | torsin family 2 member A [Source:HGNC Symbol;Acc:HGNC:11996] | -0.6427 | 1.51E-14 |
| ENSG00000113369 | ARRDC3 | arrestin domain containing 3 [Source:HGNC Symbol;Acc:HGNC:29263] | 1.1530 | 1.52E-14 |
| ENSG00000101417 | PXMP4 | peroxisomal membrane protein 4 [Source:HGNC Symbol;Acc:HGNC:15920] | -1.3840 | 1.58E-14 |
| ENSG00000237943 | PRKCQ-AS1 | PRKCQ antisense RNA 1 [Source:HGNC Symbol;Acc:HGNC:44689] | 0.9001 | 1.58E-14 |
| ENSG00000185010 | 8,00 F | coagulation factor VIII [Source:HGNC Symbol;Acc:HGNC:3546] | -1.1015 | 1.59E-14 |
| ENSG00000086062 | B4GALT1 | beta-1,4-galactosyltransferase 1 [Source:HGNC Symbol;Acc:HGNC:924] | -0.8327 | 1.67E-14 |
| ENSG00000142065 | ZFP14 | ZFP14 zinc finger protein [Source:HGNC Symbol;Acc:HGNC:29312] | -1.4835 | 1.77E-14 |
| ENSG00000233485 | FHAD1-AS1 | FHAD1 antisense RNA 1 [Source:HGNC Symbol;Acc:HGNC:41241] | -1.7320 | 1.78E-14 |
| ENSG00000170689 | HOXB9 | homeobox B9 [Source:HGNC Symbol;Acc:HGNC:5120] | 1.0362 | 1.80E-14 |
| ENSG00000196482 | ESRRG | estrogen related receptor gamma [Source:HGNC Symbol;Acc:HGNC:3474] | 0.8216 | 2.00E-14 |

| | | | | |
|---|---|---|---|---|
| ENSG00000156515 | HK1 | hexokinase 1 [Source:HGNC Symbol;Acc:HGNC:4922] | 0.5973 | 2.04E-14 |
| ENSG00000019144 | PHLDB1 | pleckstrin homology like domain family B member 1 [Source:HGNC Symbol;Acc:HGNC:23697] | -0.7763 | 2.07E-14 |
| ENSG00000204682 | MIR1915HG | MIR1915 host gene [Source:HGNC Symbol;Acc:HGNC:31448] | 1.1267 | 2.08E-14 |
| ENSG00000254685 | FPGT | fucose-1-phosphate guanylyltransferase [Source:HGNC Symbol;Acc:HGNC:3825] | -0.9555 | 2.08E-14 |
| ENSG00000168003 | SLC3A2 | solute carrier family 3 member 2 [Source:HGNC Symbol;Acc:HGNC:11026] | 0.8416 | 2.13E-14 |
| ENSG00000223891 | OSER1-DT | OSER1 divergent transcript [Source:HGNC Symbol;Acc:HGNC:48585] | -1.7962 | 2.13E-14 |
| ENSG00000135424 | ITGA7 | integrin subunit alpha 7 [Source:HGNC Symbol;Acc:HGNC:6143] | 0.6369 | 2.19E-14 |
| ENSG00000106546 | AHR | aryl hydrocarbon receptor [Source:HGNC Symbol;Acc:HGNC:348] | 1.4582 | 2.32E-14 |
| ENSG00000166595 | CIAO2B | cytosolic iron-sulfur assembly component 2B [Source:HGNC Symbol;Acc:HGNC:24261] | 0.6148 | 2.32E-14 |
| ENSG00000140396 | NCOA2 | nuclear receptor coactivator 2 [Source:HGNC Symbol;Acc:HGNC:7669] | -0.6552 | 2.36E-14 |
| ENSG00000154229 | PRKCA | protein kinase C alpha [Source:HGNC Symbol;Acc:HGNC:9393] | 2.1875 | 2.43E-14 |
| ENSG00000196305 | IARS1 | isoleucyl-tRNA synthetase 1 [Source:HGNC Symbol;Acc:HGNC:5330] | 0.5180 | 2.43E-14 |
| ENSG00000168743 | NPNT | nephronectin [Source:HGNC Symbol;Acc:HGNC:27405] | -0.9001 | 2.48E-14 |
| ENSG00000272405 | NA | novel transcript, antisense to BCAN | 1.4027 | 2.50E-14 |
| ENSG00000182197 | EXT1 | exostosin glycosyltransferase 1 [Source:HGNC Symbol;Acc:HGNC:3512] | 1.2351 | 2.81E-14 |
| ENSG00000196535 | MYO18A | myosin XVIIIA [Source:HGNC Symbol;Acc:HGNC:31104] | -0.9577 | 2.86E-14 |
| ENSG00000104825 | NFKBIB | NFKB inhibitor beta [Source:HGNC Symbol;Acc:HGNC:7798] | 0.8485 | 2.94E-14 |
| ENSG00000102181 | CD99L2 | CD99 molecule like 2 [Source:HGNC Symbol;Acc:HGNC:18237] | -0.6504 | 2.96E-14 |
| ENSG00000158987 | RAPGEF6 | Rap guanine nucleotide exchange factor 6 [Source:HGNC Symbol;Acc:HGNC:20655] | 0.7808 | 3.03E-14 |
| ENSG00000100138 | SNU13 | small nuclear ribonucleoprotein 13 [Source:HGNC Symbol;Acc:HGNC:7819] | 0.6108 | 3.22E-14 |
| ENSG00000133739 | LRRCC1 | leucine rich repeat and coiled-coil centrosomal protein 1 [Source:HGNC Symbol;Acc:HGNC:29373] | -0.6579 | 3.25E-14 |
| ENSG00000139514 | SLC7A1 | solute carrier family 7 member 1 [Source:HGNC Symbol;Acc:HGNC:11057] | 0.7183 | 3.30E-14 |
| ENSG00000197106 | SLC6A17 | solute carrier family 6 member 17 [Source:HGNC Symbol;Acc:HGNC:31399] | 5.5702 | 3.32E-14 |
| ENSG00000115255 | REEP6 | receptor accessory protein 6 [Source:HGNC Symbol;Acc:HGNC:30078] | -0.7300 | 3.40E-14 |
| ENSG00000175048 | ZDHHC14 | zinc finger DHHC-type palmitoyltransferase 14 [Source:HGNC Symbol;Acc:HGNC:20341] | 1.1205 | 3.42E-14 |
| ENSG00000127220 | ABHD8 | abhydrolase domain containing 8 [Source:HGNC Symbol;Acc:HGNC:23759] | -0.8863 | 3.51E-14 |
| ENSG00000075391 | RASAL2 | RAS protein activator like 2 [Source:HGNC Symbol;Acc:HGNC:9874] | 0.6752 | 3.58E-14 |
| ENSG00000143842 | SOX13 | SRY-box transcription factor 13 [Source:HGNC Symbol;Acc:HGNC:11192] | -1.8289 | 3.71E-14 |
| ENSG00000183741 | CBX6 | chromobox 6 [Source:HGNC Symbol;Acc:HGNC:1556] | -0.5778 | 3.76E-14 |
| ENSG00000166257 | SCN3B | sodium voltage-gated channel beta subunit 3 [Source:HGNC Symbol;Acc:HGNC:20665] | 1.3605 | 4.08E-14 |
| ENSG00000103855 | CD276 | CD276 molecule [Source:HGNC Symbol;Acc:HGNC:19137] | 0.7747 | 4.49E-14 |
| ENSG00000100211 | CBY1 | chibby family member 1, beta catenin antagonist [Source:HGNC Symbol;Acc:HGNC:1307] | -0.6797 | 4.54E-14 |
| ENSG00000166166 | TRMT61A | tRNA methyltransferase 61A [Source:HGNC Symbol;Acc:HGNC:23790] | 0.7345 | 4.65E-14 |
| ENSG00000188010 | MORN2 | MORN repeat containing 2 [Source:HGNC Symbol;Acc:HGNC:30166] | 1.2543 | 4.71E-14 |
| ENSG00000179163 | FUCA1 | alpha-L-fucosidase 1 [Source:HGNC Symbol;Acc:HGNC:4006] | -1.1212 | 4.74E-14 |
| ENSG00000176853 | FAM91A1 | family with sequence similarity 91 member A1 [Source:HGNC Symbol;Acc:HGNC:26306] | 0.5055 | 5.00E-14 |
| ENSG00000096696 | DSP | desmoplakin [Source:HGNC Symbol;Acc:HGNC:3052] | -0.8133 | 5.06E-14 |
| ENSG00000165905 | LARGE2 | LARGE xylosyl- and glucuronyltransferase 2 [Source:HGNC Symbol;Acc:HGNC:16522] | -1.3083 | 5.08E-14 |
| ENSG00000158715 | SLC45A3 | solute carrier family 45 member 3 [Source:HGNC Symbol;Acc:HGNC:8642] | 2.6685 | 5.13E-14 |
| ENSG00000177875 | CCDC184 | coiled-coil domain containing 184 [Source:HGNC Symbol;Acc:HGNC:33749] | 1.2724 | 5.24E-14 |
| ENSG00000082781 | ITGB5 | integrin subunit beta 5 [Source:HGNC Symbol;Acc:HGNC:6160] | 1.7247 | 5.43E-14 |
| ENSG00000164163 | ABCE1 | ATP binding cassette subfamily E member 1 [Source:HGNC Symbol;Acc:HGNC:69] | 0.7618 | 5.46E-14 |
| ENSG00000174405 | LIG4 | DNA ligase 4 [Source:HGNC Symbol;Acc:HGNC:6601] | 0.5368 | 5.48E-14 |

| | | | | |
|---|---|---|---|---|
| ENSG00000101079 | NDRG3 | NDRG family member 3 [Source:HGNC Symbol;Acc:HGNC:14462] | -0.8361 | 5.57E-14 |
| ENSG00000172731 | LRRC20 | leucine rich repeat containing 20 [Source:HGNC Symbol;Acc:HGNC:23421] | -1.4192 | 5.62E-14 |
| ENSG00000168394 | TAP1 | transporter 1, ATP binding cassette subfamily B member [Source:HGNC Symbol;Acc:HGNC:43] | -1.1520 | 5.83E-14 |
| ENSG00000168374 | ARF4 | ADP ribosylation factor 4 [Source:HGNC Symbol;Acc:HGNC:655] | 0.5744 | 5.86E-14 |
| ENSG00000160014 | CALM3 | calmodulin 3 [Source:HGNC Symbol;Acc:HGNC:1449] | -0.5149 | 5.90E-14 |
| ENSG00000170677 | SOCS6 | suppressor of cytokine signaling 6 [Source:HGNC Symbol;Acc:HGNC:16833] | 1.0121 | 5.91E-14 |
| ENSG00000105429 | MEGF8 | multiple EGF like domains 8 [Source:HGNC Symbol;Acc:HGNC:3233] | -0.6630 | 6.10E-14 |
| ENSG00000161813 | LARP4 | La ribonucleoprotein 4 [Source:HGNC Symbol;Acc:HGNC:24320] | 0.5655 | 6.10E-14 |
| ENSG00000181192 | DHTKD1 | dehydrogenase E1 and transketolase domain containing 1 [Source:HGNC Symbol;Acc:HGNC:23537] | -0.8453 | 6.13E-14 |
| ENSG00000031698 | SARS1 | seryl-tRNA synthetase 1 [Source:HGNC Symbol;Acc:HGNC:10537] | 0.6213 | 6.14E-14 |
| ENSG00000280734 | LINC01232 | long intergenic non-protein coding RNA 1232 [Source:HGNC Symbol;Acc:HGNC:49755] | -1.1310 | 6.18E-14 |
| ENSG00000023734 | STRAP | serine/threonine kinase receptor associated protein [Source:HGNC Symbol;Acc:HGNC:30796] | 0.5240 | 6.21E-14 |
| ENSG00000148429 | USP6NL | USP6 N-terminal like [Source:HGNC Symbol;Acc:HGNC:16858] | 0.6698 | 6.81E-14 |
| ENSG00000173846 | PLK3 | polo like kinase 3 [Source:HGNC Symbol;Acc:HGNC:2154] | 1.7898 | 6.99E-14 |
| ENSG00000123358 | NR4A1 | nuclear receptor subfamily 4 group A member 1 [Source:HGNC Symbol;Acc:HGNC:7980] | 2.2994 | 7.12E-14 |
| ENSG00000167580 | AQP2 | aquaporin 2 [Source:HGNC Symbol;Acc:HGNC:634] | 1.5272 | 7.62E-14 |
| ENSG00000173641 | HSPB7 | heat shock protein family B (small) member 7 [Source:HGNC Symbol;Acc:HGNC:5249] | 0.8452 | 7.64E-14 |
| ENSG00000104332 | SFRP1 | secreted frizzled related protein 1 [Source:HGNC Symbol;Acc:HGNC:10776] | 3.5240 | 7.67E-14 |
| ENSG00000198805 | PNP | purine nucleoside phosphorylase [Source:HGNC Symbol;Acc:HGNC:7892] | 0.8922 | 7.67E-14 |
| ENSG00000197448 | GSTK1 | glutathione S-transferase kappa 1 [Source:HGNC Symbol;Acc:HGNC:16906] | -0.5570 | 8.19E-14 |
| ENSG00000105821 | DNAJC2 | DnaJ heat shock protein family (Hsp40) member C2 [Source:HGNC Symbol;Acc:HGNC:13192] | 0.9977 | 8.42E-14 |
| ENSG00000010318 | PHF7 | PHD finger protein 7 [Source:HGNC Symbol;Acc:HGNC:18458] | -1.8103 | 8.59E-14 |
| ENSG00000008283 | CYB561 | cytochrome b561 [Source:HGNC Symbol;Acc:HGNC:2571] | 1.0696 | 8.74E-14 |
| ENSG00000090621 | PABPC4 | poly(A) binding protein cytoplasmic 4 [Source:HGNC Symbol;Acc:HGNC:8557] | 0.5477 | 9.32E-14 |
| ENSG00000164403 | SHROOM1 | shroom family member 1 [Source:HGNC Symbol;Acc:HGNC:24084] | 1.4038 | 9.42E-14 |
| ENSG00000092964 | DPYSL2 | dihydropyrimidinase like 2 [Source:HGNC Symbol;Acc:HGNC:3014] | -0.7932 | 9.81E-14 |
| ENSG00000064666 | CNN2 | calponin 2 [Source:HGNC Symbol;Acc:HGNC:2156] | 1.3569 | 1.02E-13 |
| ENSG00000154813 | DPH3 | diphthamide biosynthesis 3 [Source:HGNC Symbol;Acc:HGNC:27717] | 0.7073 | 1.02E-13 |
| ENSG00000117643 | MAN1C1 | mannosidase alpha class 1C member 1 [Source:HGNC Symbol;Acc:HGNC:19080] | -1.8446 | 1.04E-13 |
| ENSG00000134684 | YARS1 | tyrosyl-tRNA synthetase 1 [Source:HGNC Symbol;Acc:HGNC:12840] | 0.8063 | 1.06E-13 |
| ENSG00000170927 | PKHD1 | PKHD1 ciliary IPT domain containing fibrocystin/polyductin [Source:HGNC Symbol;Acc:HGNC:9016] | -0.9081 | 1.09E-13 |
| ENSG00000148053 | NTRK2 | neurotrophic receptor tyrosine kinase 2 [Source:HGNC Symbol;Acc:HGNC:8032] | -2.1799 | 1.19E-13 |
| ENSG00000104964 | TLE5 | TLE family member 5, transcriptional modulator [Source:HGNC Symbol;Acc:HGNC:307] | 0.5663 | 1.20E-13 |
| ENSG00000102265 | TIMP1 | TIMP metallopeptidase inhibitor 1 [Source:HGNC Symbol;Acc:HGNC:11820] | 0.8167 | 1.20E-13 |
| ENSG00000137692 | DCUN1D5 | defective in cullin neddylation 1 domain containing 5 [Source:HGNC Symbol;Acc:HGNC:28409] | 0.6736 | 1.24E-13 |
| ENSG00000177728 | TMEM94 | transmembrane protein 94 [Source:HGNC Symbol;Acc:HGNC:28983] | -0.7226 | 1.28E-13 |
| ENSG00000151468 | CCDC3 | coiled-coil domain containing 3 [Source:HGNC Symbol;Acc:HGNC:23813] | 2.7594 | 1.29E-13 |
| ENSG00000277200 | NA | novel transcript | -1.7538 | 1.30E-13 |
| ENSG00000092421 | SEMA6A | semaphorin 6A [Source:HGNC Symbol;Acc:HGNC:10738] | -1.6080 | 1.33E-13 |
| ENSG00000069869 | NEDD4 | NEDD4 E3 ubiquitin protein ligase [Source:HGNC Symbol;Acc:HGNC:7727] | 0.7720 | 1.33E-13 |
| ENSG00000102226 | USP11 | ubiquitin specific peptidase 11 [Source:HGNC Symbol;Acc:HGNC:12609] | 0.5010 | 1.34E-13 |
| ENSG00000132256 | TRIM5 | tripartite motif containing 5 [Source:HGNC Symbol;Acc:HGNC:16276] | 1.7770 | 1.39E-13 |
| ENSG00000175899 | A2M | alpha-2-macroglobulin [Source:HGNC Symbol;Acc:HGNC:7] | 0.7206 | 1.41E-13 |

| | | | | |
|---|---|---|---|---|
| ENSG00000205269 | TMEM170B | transmembrane protein 170B [Source:HGNC Symbol;Acc:HGNC:34244] | -0.8587 | 1.43E-13 |
| ENSG00000139160 | ETFBKMT | electron transfer flavoprotein subunit beta lysine methyltransferase [Source:HGNC Symbol;Acc:HGNC:28739] | -1.1583 | 1.46E-13 |
| ENSG00000172551 | MUCL1 | mucin like 1 [Source:HGNC Symbol;Acc:HGNC:30588] | -1.0713 | 1.51E-13 |
| ENSG00000106948 | AKNA | AT-hook transcription factor [Source:HGNC Symbol;Acc:HGNC:24108] | 1.7998 | 1.53E-13 |
| ENSG00000149782 | PLCB3 | phospholipase C beta 3 [Source:HGNC Symbol;Acc:HGNC:9056] | -0.9258 | 1.58E-13 |
| ENSG00000232859 | LYRM9 | LYR motif containing 9 [Source:HGNC Symbol;Acc:HGNC:27314] | -1.5343 | 1.63E-13 |
| ENSG00000139146 | SINHCAF | SIN3-HDAC complex associated factor [Source:HGNC Symbol;Acc:HGNC:30702] | 0.7183 | 1.72E-13 |
| ENSG00000157020 | SEC13 | SEC13 homolog, nuclear pore and COPII coat complex component [Source:HGNC Symbol;Acc:HGNC:10697] | 0.5404 | 1.76E-13 |
| ENSG00000077147 | TM9SF3 | transmembrane 9 superfamily member 3 [Source:HGNC Symbol;Acc:HGNC:21529] | 0.5972 | 1.80E-13 |
| ENSG00000072210 | ALDH3A2 | aldehyde dehydrogenase 3 family member A2 [Source:HGNC Symbol;Acc:HGNC:403] | 0.9338 | 1.86E-13 |
| ENSG00000111670 | GNPTAB | N-acetylglucosamine-1-phosphate transferase subunits alpha and beta [Source:HGNC Symbol;Acc:HGNC:29670] | -0.6481 | 1.86E-13 |
| ENSG00000139973 | SYT16 | synaptotagmin 16 [Source:HGNC Symbol;Acc:HGNC:23142] | 2.4050 | 1.89E-13 |
| ENSG00000108559 | NUP88 | nucleoporin 88 [Source:HGNC Symbol;Acc:HGNC:8067] | 0.6774 | 1.95E-13 |
| ENSG00000109066 | TMEM104 | transmembrane protein 104 [Source:HGNC Symbol;Acc:HGNC:25984] | 0.8976 | 2.00E-13 |
| ENSG00000148019 | CEP78 | centrosomal protein 78 [Source:HGNC Symbol;Acc:HGNC:25740] | -0.9037 | 2.04E-13 |
| ENSG00000171791 | BCL2 | BCL2 apoptosis regulator [Source:HGNC Symbol;Acc:HGNC:990] | -1.9629 | 2.04E-13 |
| ENSG00000165533 | TTC8 | tetratricopeptide repeat domain 8 [Source:HGNC Symbol;Acc:HGNC:20087] | -0.6693 | 2.05E-13 |
| ENSG00000104979 | C19orf53 | chromosome 19 open reading frame 53 [Source:HGNC Symbol;Acc:HGNC:24991] | 0.6398 | 2.14E-13 |
| ENSG00000135709 | KIAA0513 | KIAA0513 [Source:HGNC Symbol;Acc:HGNC:29058] | 1.4131 | 2.14E-13 |
| ENSG00000139988 | RDH12 | retinol dehydrogenase 12 [Source:HGNC Symbol;Acc:HGNC:19977] | 1.5966 | 2.18E-13 |
| ENSG00000120306 | CYSTM1 | cysteine rich transmembrane module containing 1 [Source:HGNC Symbol;Acc:HGNC:30239] | 0.8616 | 2.26E-13 |
| ENSG00000185883 | ATP6V0C | ATPase H+ transporting V0 subunit c [Source:HGNC Symbol;Acc:HGNC:855] | 0.5752 | 2.26E-13 |
| ENSG00000159079 | CFAP298 | cilia and flagella associated protein 298 [Source:HGNC Symbol;Acc:HGNC:1301] | 0.6863 | 2.26E-13 |
| ENSG00000147955 | SIGMAR1 | sigma non-opioid intracellular receptor 1 [Source:HGNC Symbol;Acc:HGNC:8157] | -0.6694 | 2.27E-13 |
| ENSG00000170293 | CMTM8 | CKLF like MARVEL transmembrane domain containing 8 [Source:HGNC Symbol;Acc:HGNC:19179] | -0.8889 | 2.28E-13 |
| ENSG00000134987 | WDR36 | WD repeat domain 36 [Source:HGNC Symbol;Acc:HGNC:30696] | 0.5989 | 2.32E-13 |
| ENSG00000187678 | SPRY4 | sprouty RTK signaling antagonist 4 [Source:HGNC Symbol;Acc:HGNC:15533] | 5.0780 | 2.33E-13 |
| ENSG00000093183 | SEC22C | SEC22 homolog C, vesicle trafficking protein [Source:HGNC Symbol;Acc:HGNC:16828] | -0.5611 | 2.35E-13 |
| ENSG00000121621 | KIF18A | kinesin family member 18A [Source:HGNC Symbol;Acc:HGNC:29441] | 0.7503 | 2.36E-13 |
| ENSG00000161013 | MGAT4B | alpha-1,3-mannosyl-glycoprotein 4-beta-N-acetylglucosaminyltransferase B [Source:HGNC Symbol;Acc:HGNC:7048] | -0.6940 | 2.46E-13 |
| ENSG00000105514 | RAB3D | RAB3D, member RAS oncogene family [Source:HGNC Symbol;Acc:HGNC:9779] | -0.9214 | 2.58E-13 |
| ENSG00000145730 | PAM | peptidylglycine alpha-amidating monooxygenase [Source:HGNC Symbol;Acc:HGNC:8596] | 1.2965 | 2.61E-13 |
| ENSG00000270412 | NA | novel transcript, antisense to COL15A1 | 1.7845 | 2.72E-13 |
| ENSG00000213533 | STIMATE | STIM activating enhancer [Source:HGNC Symbol;Acc:HGNC:30526] | -0.9724 | 2.75E-13 |
| ENSG00000062282 | DGAT2 | diacylglycerol O-acyltransferase 2 [Source:HGNC Symbol;Acc:HGNC:16940] | -2.2648 | 2.86E-13 |
| ENSG00000160963 | COL26A1 | collagen type XXVI alpha 1 chain [Source:HGNC Symbol;Acc:HGNC:18038] | -2.7002 | 2.86E-13 |
| ENSG00000167797 | CDK2AP2 | cyclin dependent kinase 2 associated protein 2 [Source:HGNC Symbol;Acc:HGNC:30833] | 1.4623 | 2.86E-13 |
| ENSG00000072849 | DERL2 | derlin 2 [Source:HGNC Symbol;Acc:HGNC:17943] | 0.6223 | 3.05E-13 |
| ENSG00000140262 | TCF12 | transcription factor 12 [Source:HGNC Symbol;Acc:HGNC:11623] | 0.6986 | 3.06E-13 |
| ENSG00000111432 | FZD10 | frizzled class receptor 10 [Source:HGNC Symbol;Acc:HGNC:4039] | 2.8847 | 3.09E-13 |
| ENSG00000127946 | HIP1 | huntingtin interacting protein 1 [Source:HGNC Symbol;Acc:HGNC:4913] | -0.8387 | 3.15E-13 |
| ENSG00000158615 | PPP1R15B | protein phosphatase 1 regulatory subunit 15B [Source:HGNC Symbol;Acc:HGNC:14951] | 0.5966 | 3.16E-13 |
| ENSG00000164180 | TMEM161B | transmembrane protein 161B [Source:HGNC Symbol;Acc:HGNC:28483] | 0.5570 | 3.21E-13 |

| | | | | |
|---|---|---|---|---|
| ENSG00000136144 | RCBTB1 | RCC1 and BTB domain containing protein 1 [Source:HGNC Symbol;Acc:HGNC:18243] | -0.8086 | 3.33E-13 |
| ENSG00000100246 | DNAL4 | dynein axonemal light chain 4 [Source:HGNC Symbol;Acc:HGNC:2955] | -0.9766 | 3.49E-13 |
| ENSG00000162620 | LRRIQ3 | leucine rich repeats and IQ motif containing 3 [Source:HGNC Symbol;Acc:HGNC:28318] | -1.4470 | 3.49E-13 |
| ENSG00000111911 | HINT3 | histidine triad nucleotide binding protein 3 [Source:HGNC Symbol;Acc:HGNC:18468] | -0.7815 | 3.56E-13 |
| ENSG00000169302 | STK32A | serine/threonine kinase 32A [Source:HGNC Symbol;Acc:HGNC:28317] | -1.2740 | 3.60E-13 |
| ENSG00000106624 | AEBP1 | AE binding protein 1 [Source:HGNC Symbol;Acc:HGNC:303] | 0.6890 | 3.60E-13 |
| ENSG00000118513 | MYB | MYB proto-oncogene, transcription factor [Source:HGNC Symbol;Acc:HGNC:7545] | 1.6251 | 3.62E-13 |
| ENSG00000133169 | BEX1 | brain expressed X-linked 1 [Source:HGNC Symbol;Acc:HGNC:1036] | 0.7603 | 3.65E-13 |
| ENSG00000171928 | TVP23B | trans-golgi network vesicle protein 23 homolog B [Source:HGNC Symbol;Acc:HGNC:20399] | 0.6129 | 3.68E-13 |
| ENSG00000052126 | PLEKHA5 | pleckstrin homology domain containing A5 [Source:HGNC Symbol;Acc:HGNC:30036] | -0.6418 | 3.69E-13 |
| ENSG00000115946 | PNO1 | partner of NOB1 homolog [Source:HGNC Symbol;Acc:HGNC:32790] | 0.9199 | 3.79E-13 |
| ENSG00000090238 | YPEL3 | yippee like 3 [Source:HGNC Symbol;Acc:HGNC:18327] | -1.2524 | 3.79E-13 |
| ENSG00000112031 | MTRF1L | mitochondrial translational release factor 1 like [Source:HGNC Symbol;Acc:HGNC:21051] | 0.7668 | 3.92E-13 |
| ENSG00000181481 | RNF135 | ring finger protein 135 [Source:HGNC Symbol;Acc:HGNC:21158] | -1.1445 | 3.97E-13 |
| ENSG00000119508 | NR4A3 | nuclear receptor subfamily 4 group A member 3 [Source:HGNC Symbol;Acc:HGNC:7982] | 2.0884 | 4.04E-13 |
| ENSG00000170175 | CHRNB1 | cholinergic receptor nicotinic beta 1 subunit [Source:HGNC Symbol;Acc:HGNC:1961] | -1.0662 | 4.21E-13 |
| ENSG00000082153 | BZW1 | basic leucine zipper and W2 domains 1 [Source:HGNC Symbol;Acc:HGNC:18380] | 0.7788 | 4.43E-13 |
| ENSG00000113269 | RNF130 | ring finger protein 130 [Source:HGNC Symbol;Acc:HGNC:18280] | -0.6476 | 4.72E-13 |
| ENSG00000138193 | PLCE1 | phospholipase C epsilon 1 [Source:HGNC Symbol;Acc:HGNC:17175] | -1.5035 | 4.72E-13 |
| ENSG00000111452 | ADGRD1 | adhesion G protein-coupled receptor D1 [Source:HGNC Symbol;Acc:HGNC:19893] | 2.5829 | 4.74E-13 |
| ENSG00000184203 | PPP1R2 | protein phosphatase 1 regulatory inhibitor subunit 2 [Source:HGNC Symbol;Acc:HGNC:9288] | 0.5945 | 4.94E-13 |
| ENSG00000135414 | GDF11 | growth differentiation factor 11 [Source:HGNC Symbol;Acc:HGNC:4216] | 0.5105 | 4.95E-13 |
| ENSG00000273274 | ZBTB8B | zinc finger and BTB domain containing 8B [Source:HGNC Symbol;Acc:HGNC:37057] | -2.0367 | 4.97E-13 |
| ENSG00000179889 | PDXDC1 | pyridoxal dependent decarboxylase domain containing 1 [Source:HGNC Symbol;Acc:HGNC:28995] | -0.6123 | 5.15E-13 |
| ENSG00000167674 | HDGFL2 | HDGF like 2 [Source:HGNC Symbol;Acc:HGNC:14680] | -0.5966 | 5.24E-13 |
| ENSG00000126858 | RHOT1 | ras homolog family member T1 [Source:HGNC Symbol;Acc:HGNC:21168] | -0.6473 | 5.26E-13 |
| ENSG00000235711 | ANKRD34C | ankyrin repeat domain 34C [Source:HGNC Symbol;Acc:HGNC:33888] | 3.5514 | 5.63E-13 |
| ENSG00000105355 | PLIN3 | perilipin 3 [Source:HGNC Symbol;Acc:HGNC:16893] | 0.7884 | 5.64E-13 |
| ENSG00000164933 | SLC25A32 | solute carrier family 25 member 32 [Source:HGNC Symbol;Acc:HGNC:29683] | 0.6667 | 5.67E-13 |
| ENSG00000143578 | CREB3L4 | cAMP responsive element binding protein 3 like 4 [Source:HGNC Symbol;Acc:HGNC:18854] | -1.2103 | 5.81E-13 |
| ENSG00000186648 | CARMIL3 | capping protein regulator and myosin 1 linker 3 [Source:HGNC Symbol;Acc:HGNC:20272] | -1.5730 | 6.23E-13 |
| ENSG00000075624 | ACTB | actin beta [Source:HGNC Symbol;Acc:HGNC:132] | 0.9193 | 6.40E-13 |
| ENSG00000123496 | IL13RA2 | interleukin 13 receptor subunit alpha 2 [Source:HGNC Symbol;Acc:HGNC:5975] | 0.6756 | 6.44E-13 |
| ENSG00000237399 | PITRM1-AS1 | PITRM1 antisense RNA 1 [Source:HGNC Symbol;Acc:HGNC:44675] | 1.8923 | 6.52E-13 |
| ENSG00000253982 | NA | novel transcript | -0.9679 | 6.52E-13 |
| ENSG00000146250 | PRSS35 | serine protease 35 [Source:HGNC Symbol;Acc:HGNC:21387] | 1.4861 | 6.53E-13 |
| ENSG00000183580 | FBXL7 | F-box and leucine rich repeat protein 7 [Source:HGNC Symbol;Acc:HGNC:13604] | 0.8451 | 6.55E-13 |
| ENSG00000155760 | FZD7 | frizzled class receptor 7 [Source:HGNC Symbol;Acc:HGNC:4045] | -0.7972 | 7.26E-13 |
| ENSG00000112874 | NUDT12 | nudix hydrolase 12 [Source:HGNC Symbol;Acc:HGNC:18826] | -0.6976 | 7.51E-13 |
| ENSG00000154767 | XPC | XPC complex subunit, DNA damage recognition and repair factor [Source:HGNC Symbol;Acc:HGNC:12816] | -0.7568 | 7.62E-13 |
| ENSG00000147535 | PLPP5 | phospholipid phosphatase 5 [Source:HGNC Symbol;Acc:HGNC:25026] | 1.2259 | 7.74E-13 |
| ENSG00000065559 | MAP2K4 | mitogen-activated protein kinase kinase 4 [Source:HGNC Symbol;Acc:HGNC:6844] | 0.5456 | 7.85E-13 |

| | | | | |
|---|---|---|---|---|
| ENSG00000136770 | DNAJC1 | DnaJ heat shock protein family (Hsp40) member C1 [Source:HGNC Symbol;Acc:HGNC:20090] | 0.6746 | 7.85E-13 |
| ENSG00000132475 | H3-3B | H3.3 histone B [Source:HGNC Symbol;Acc:HGNC:4765] | 0.5286 | 7.85E-13 |
| ENSG00000115419 | GLS | glutaminase [Source:HGNC Symbol;Acc:HGNC:4331] | 1.0886 | 8.15E-13 |
| ENSG00000112984 | KIF20A | kinesin family member 20A [Source:HGNC Symbol;Acc:HGNC:9787] | -0.8182 | 8.37E-13 |
| ENSG00000171045 | TSNARE1 | t-SNARE domain containing 1 [Source:HGNC Symbol;Acc:HGNC:26437] | -1.7691 | 8.37E-13 |
| ENSG00000162980 | ARL5A | ADP ribosylation factor like GTPase 5A [Source:HGNC Symbol;Acc:HGNC:696] | 0.9383 | 8.77E-13 |
| ENSG00000189339 | SLC35E2B | solute carrier family 35 member E2B [Source:HGNC Symbol;Acc:HGNC:33941] | -0.5514 | 9.06E-13 |
| ENSG00000277443 | MARCKS | myristoylated alanine rich protein kinase C substrate [Source:HGNC Symbol;Acc:HGNC:6759] | -0.8335 | 9.06E-13 |
| ENSG00000181704 | YIPF6 | Yip1 domain family member 6 [Source:HGNC Symbol;Acc:HGNC:28304] | 0.6605 | 9.38E-13 |
| ENSG00000173145 | NOC3L | NOC3 like DNA replication regulator [Source:HGNC Symbol;Acc:HGNC:24034] | 0.5105 | 9.51E-13 |
| ENSG00000166012 | TAF1D | TATA-box binding protein associated factor, RNA polymerase I subunit D [Source:HGNC Symbol;Acc:HGNC:28759] | 0.8172 | 1.01E-12 |
| ENSG00000137261 | KIAA0319 | KIAA0319 [Source:HGNC Symbol;Acc:HGNC:21580] | -1.4991 | 1.02E-12 |
| ENSG00000127585 | FBXL16 | F-box and leucine rich repeat protein 16 [Source:HGNC Symbol;Acc:HGNC:14150] | -0.6867 | 1.02E-12 |
| ENSG00000115540 | MOB4 | MOB family member 4, phocein [Source:HGNC Symbol;Acc:HGNC:17261] | 1.0992 | 1.04E-12 |
| ENSG00000155366 | RHOC | ras homolog family member C [Source:HGNC Symbol;Acc:HGNC:669] | 0.6766 | 1.06E-12 |
| ENSG00000142686 | C1orf216 | chromosome 1 open reading frame 216 [Source:HGNC Symbol;Acc:HGNC:26800] | 0.8577 | 1.07E-12 |
| ENSG00000188522 | FAM83G | family with sequence similarity 83 member G [Source:HGNC Symbol;Acc:HGNC:32554] | 1.3125 | 1.08E-12 |
| ENSG00000236753 | MKLN1-AS | MKLN1 antisense RNA [Source:HGNC Symbol;Acc:HGNC:40374] | 1.4358 | 1.11E-12 |
| ENSG00000188517 | COL25A1 | collagen type XXV alpha 1 chain [Source:HGNC Symbol;Acc:HGNC:18603] | 0.8676 | 1.13E-12 |
| ENSG00000241360 | PDXP | pyridoxal phosphatase [Source:HGNC Symbol;Acc:HGNC:30259] | 0.6070 | 1.15E-12 |
| ENSG00000113460 | BRIX1 | biogenesis of ribosomes BRX1 [Source:HGNC Symbol;Acc:HGNC:24170] | 0.5875 | 1.19E-12 |
| ENSG00000229425 | NA | novel transcript | 1.1882 | 1.21E-12 |
| ENSG00000101347 | SAMHD1 | SAM and HD domain containing deoxynucleoside triphosphate triphosphohydrolase 1 [Source:HGNC Symbol;Acc:HGNC:15925] | -0.9332 | 1.22E-12 |
| ENSG00000117395 | EBNA1BP2 | EBNA1 binding protein 2 [Source:HGNC Symbol;Acc:HGNC:15531] | 0.5030 | 1.23E-12 |
| ENSG00000166006 | KCNC2 | potassium voltage-gated channel subfamily C member 2 [Source:HGNC Symbol;Acc:HGNC:6234] | -1.7305 | 1.26E-12 |
| ENSG00000177706 | FAM20C | FAM20C golgi associated secretory pathway kinase [Source:HGNC Symbol;Acc:HGNC:22140] | -1.3924 | 1.27E-12 |
| ENSG00000106665 | CLIP2 | CAP-Gly domain containing linker protein 2 [Source:HGNC Symbol;Acc:HGNC:2586] | 1.0919 | 1.34E-12 |
| ENSG00000122218 | COPA | COPI coat complex subunit alpha [Source:HGNC Symbol;Acc:HGNC:2230] | 0.6208 | 1.42E-12 |
| ENSG00000250447 | LINC02105 | long intergenic non-protein coding RNA 2105 [Source:HGNC Symbol;Acc:HGNC:52960] | 4.3452 | 1.49E-12 |
| ENSG00000038358 | EDC4 | enhancer of mRNA decapping 4 [Source:HGNC Symbol;Acc:HGNC:17157] | 0.5327 | 1.52E-12 |
| ENSG00000132507 | EIF5A | eukaryotic translation initiation factor 5A [Source:HGNC Symbol;Acc:HGNC:3300] | 0.5057 | 1.54E-12 |
| ENSG00000205885 | C1RL-AS1 | C1RL antisense RNA 1 [Source:HGNC Symbol;Acc:HGNC:27461] | 2.4609 | 1.54E-12 |
| ENSG00000124802 | EEF1E1 | eukaryotic translation elongation factor 1 epsilon 1 [Source:HGNC Symbol;Acc:HGNC:3212] | 0.7530 | 1.55E-12 |
| ENSG00000169446 | MMGT1 | membrane magnesium transporter 1 [Source:HGNC Symbol;Acc:HGNC:28100] | 0.5593 | 1.55E-12 |
| ENSG00000165238 | WNK2 | WNK lysine deficient protein kinase 2 [Source:HGNC Symbol;Acc:HGNC:14542] | -1.1209 | 1.61E-12 |
| ENSG00000167658 | EEF2 | eukaryotic translation elongation factor 2 [Source:HGNC Symbol;Acc:HGNC:3214] | 0.5287 | 1.64E-12 |
| ENSG00000073969 | NSF | N-ethylmaleimide sensitive factor, vesicle fusing ATPase [Source:HGNC Symbol;Acc:HGNC:8016] | 0.8355 | 1.67E-12 |
| ENSG00000109790 | KLHL5 | kelch like family member 5 [Source:HGNC Symbol;Acc:HGNC:6356] | -0.9434 | 1.74E-12 |
| ENSG00000106153 | CHCHD2 | coiled-coil-helix-coiled-coil-helix domain containing 2 [Source:HGNC Symbol;Acc:HGNC:21645] | 0.5643 | 1.74E-12 |
| ENSG00000271601 | LIX1L | limb and CNS expressed 1 like [Source:HGNC Symbol;Acc:HGNC:28715] | -0.7680 | 1.75E-12 |
| ENSG00000256646 | NA | novel PSMA2 and C7orf25 readthrough | -0.5664 | 1.80E-12 |
| ENSG00000197594 | ENPP1 | ectonucleotide pyrophosphatase/phosphodiesterase 1 [Source:HGNC Symbol;Acc:HGNC:3356] | 1.7067 | 1.84E-12 |

| | | | | |
|---|---|---|---|---|
| ENSG00000006007 | GDE1 | glycerophosphodiester phosphodiesterase 1 [Source:HGNC Symbol;Acc:HGNC:29644] | 0.7785 | 1.91E-12 |
| ENSG00000154845 | PPP4R1 | protein phosphatase 4 regulatory subunit 1 [Source:HGNC Symbol;Acc:HGNC:9320] | 0.8058 | 1.95E-12 |
| ENSG00000258334 | NA | novel transcript, antisense to PRPH | -1.9660 | 2.01E-12 |
| ENSG00000128585 | MKLN1 | muskelin 1 [Source:HGNC Symbol;Acc:HGNC:7109] | 0.6057 | 2.11E-12 |
| ENSG00000135299 | ANKRD6 | ankyrin repeat domain 6 [Source:HGNC Symbol;Acc:HGNC:17280] | -1.0851 | 2.11E-12 |
| ENSG00000120832 | MTERF2 | mitochondrial transcription termination factor 2 [Source:HGNC Symbol;Acc:HGNC:30779] | -1.3040 | 2.13E-12 |
| ENSG00000126012 | KDM5C | lysine demethylase 5C [Source:HGNC Symbol;Acc:HGNC:11114] | 0.6572 | 2.16E-12 |
| ENSG00000131504 | DIAPH1 | diaphanous related formin 1 [Source:HGNC Symbol;Acc:HGNC:2876] | 0.6977 | 2.17E-12 |
| ENSG00000108518 | PFN1 | profilin 1 [Source:HGNC Symbol;Acc:HGNC:8881] | 0.5159 | 2.19E-12 |
| ENSG00000166402 | TUB | TUB bipartite transcription factor [Source:HGNC Symbol;Acc:HGNC:12406] | -1.0111 | 2.24E-12 |
| ENSG00000129128 | SPCS3 | signal peptidase complex subunit 3 [Source:HGNC Symbol;Acc:HGNC:26212] | 0.9545 | 2.27E-12 |
| ENSG00000197256 | KANK2 | KN motif and ankyrin repeat domains 2 [Source:HGNC Symbol;Acc:HGNC:29300] | -1.1335 | 2.29E-12 |
| ENSG00000182118 | FAM89A | family with sequence similarity 89 member A [Source:HGNC Symbol;Acc:HGNC:25057] | 1.2063 | 2.35E-12 |
| ENSG00000286129 | NA | novel transcript | 0.9330 | 2.42E-12 |
| ENSG00000132463 | GRSF1 | G-rich RNA sequence binding factor 1 [Source:HGNC Symbol;Acc:HGNC:4610] | 0.6258 | 2.47E-12 |
| ENSG00000272157 | NA | novel transcript, antisense to NEFL | -1.7736 | 2.48E-12 |
| ENSG00000180891 | CUEDC1 | CUE domain containing 1 [Source:HGNC Symbol;Acc:HGNC:31350] | -0.8795 | 2.51E-12 |
| ENSG00000133612 | AGAP3 | ArfGAP with GTPase domain, ankyrin repeat and PH domain 3 [Source:HGNC Symbol;Acc:HGNC:16923] | 0.7490 | 2.56E-12 |
| ENSG00000172115 | CYCS | cytochrome c, somatic [Source:HGNC Symbol;Acc:HGNC:19986] | 0.5461 | 2.56E-12 |
| ENSG00000117500 | TMED5 | transmembrane p24 trafficking protein 5 [Source:HGNC Symbol;Acc:HGNC:24251] | 0.6476 | 2.59E-12 |
| ENSG00000152782 | PANK1 | pantothenate kinase 1 [Source:HGNC Symbol;Acc:HGNC:8598] | -0.9536 | 2.60E-12 |
| ENSG00000249042 | NA | novel transcript | -0.8789 | 2.63E-12 |
| ENSG00000115520 | COQ10B | coenzyme Q10B [Source:HGNC Symbol;Acc:HGNC:25819] | 0.8389 | 2.65E-12 |
| ENSG00000115762 | PLEKHB2 | pleckstrin homology domain containing B2 [Source:HGNC Symbol;Acc:HGNC:19236] | 0.6048 | 2.73E-12 |
| ENSG00000100292 | HMOX1 | heme oxygenase 1 [Source:HGNC Symbol;Acc:HGNC:5013] | -0.8661 | 2.73E-12 |
| ENSG00000171159 | C9orf16 | chromosome 9 open reading frame 16 [Source:HGNC Symbol;Acc:HGNC:17823] | 0.8383 | 2.76E-12 |
| ENSG00000137825 | ITPKA | inositol-trisphosphate 3-kinase A [Source:HGNC Symbol;Acc:HGNC:6178] | 1.3706 | 2.77E-12 |
| ENSG00000181789 | COPG1 | COPI coat complex subunit gamma 1 [Source:HGNC Symbol;Acc:HGNC:2236] | 0.5659 | 2.77E-12 |
| ENSG00000214517 | PPME1 | protein phosphatase methylesterase 1 [Source:HGNC Symbol;Acc:HGNC:30178] | 0.6921 | 2.83E-12 |
| ENSG00000196547 | MAN2A2 | mannosidase alpha class 2A member 2 [Source:HGNC Symbol;Acc:HGNC:6825] | 0.5989 | 3.03E-12 |
| ENSG00000099910 | KLHL22 | kelch like family member 22 [Source:HGNC Symbol;Acc:HGNC:25888] | -0.7039 | 3.05E-12 |
| ENSG00000121691 | CAT | catalase [Source:HGNC Symbol;Acc:HGNC:1516] | -0.6204 | 3.13E-12 |
| ENSG00000112378 | PERP | p53 apoptosis effector related to PMP22 [Source:HGNC Symbol;Acc:HGNC:17637] | 1.0986 | 3.22E-12 |
| ENSG00000174238 | PITPNA | phosphatidylinositol transfer protein alpha [Source:HGNC Symbol;Acc:HGNC:9001] | 0.6691 | 3.25E-12 |
| ENSG00000197879 | MYO1C | myosin IC [Source:HGNC Symbol;Acc:HGNC:7597] | 0.9114 | 3.32E-12 |
| ENSG00000166562 | SEC11C | SEC11 homolog C, signal peptidase complex subunit [Source:HGNC Symbol;Acc:HGNC:23400] | 0.7505 | 3.37E-12 |
| ENSG00000162298 | SYVN1 | synoviolin 1 [Source:HGNC Symbol;Acc:HGNC:20738] | 0.7599 | 3.42E-12 |
| ENSG00000109534 | GAR1 | GAR1 ribonucleoprotein [Source:HGNC Symbol;Acc:HGNC:14264] | 0.6514 | 3.48E-12 |
| ENSG00000153179 | RASSF3 | Ras association domain family member 3 [Source:HGNC Symbol;Acc:HGNC:14271] | -0.8216 | 3.67E-12 |
| ENSG00000196873 | CBWD3 | COBW domain containing 3 [Source:HGNC Symbol;Acc:HGNC:18519] | 0.5640 | 3.68E-12 |
| ENSG00000128266 | GNAZ | G protein subunit alpha z [Source:HGNC Symbol;Acc:HGNC:4395] | -0.7962 | 3.77E-12 |
| ENSG00000109832 | DDX25 | DEAD-box helicase 25 [Source:HGNC Symbol;Acc:HGNC:18698] | -1.7792 | 3.78E-12 |
| ENSG00000134780 | DAGLA | diacylglycerol lipase alpha [Source:HGNC Symbol;Acc:HGNC:1165] | -1.1531 | 3.87E-12 |

| | | | | |
|---|---|---|---|---|
| ENSG00000240086 | NA | novel transcript, antisense to EPHB1 | -1.5553 | 3.98E-12 |
| ENSG00000085733 | CTTN | cortactin [Source:HGNC Symbol;Acc:HGNC:3338] | 0.5238 | 4.00E-12 |
| ENSG00000169045 | HNRNPH1 | heterogeneous nuclear ribonucleoprotein H1 [Source:HGNC Symbol;Acc:HGNC:5041] | 0.5108 | 4.14E-12 |
| ENSG00000284879 | NA | novel transcript | 1.3437 | 4.16E-12 |
| ENSG00000010404 | IDS | iduronate 2-sulfatase [Source:HGNC Symbol;Acc:HGNC:5389] | 0.7248 | 4.31E-12 |
| ENSG00000197056 | ZMYM1 | zinc finger MYM-type containing 1 [Source:HGNC Symbol;Acc:HGNC:26253] | -0.7371 | 4.39E-12 |
| ENSG00000137103 | TMEM8B | transmembrane protein 8B [Source:HGNC Symbol;Acc:HGNC:21427] | -0.8637 | 4.51E-12 |
| ENSG00000150907 | FOXO1 | forkhead box O1 [Source:HGNC Symbol;Acc:HGNC:3819] | 0.9308 | 4.51E-12 |
| ENSG00000011523 | CEP68 | centrosomal protein 68 [Source:HGNC Symbol;Acc:HGNC:29076] | -0.9143 | 4.77E-12 |
| ENSG00000262769 | NA | novel transcript, antisense to SLC47A1 | 1.6277 | 4.77E-12 |
| ENSG00000070413 | DGCR2 | DiGeorge syndrome critical region gene 2 [Source:HGNC Symbol;Acc:HGNC:2845] | -0.5371 | 4.79E-12 |
| ENSG00000044090 | CUL7 | cullin 7 [Source:HGNC Symbol;Acc:HGNC:21024] | -0.6807 | 4.97E-12 |
| ENSG00000012963 | UBR7 | ubiquitin protein ligase E3 component n-recognin 7 [Source:HGNC Symbol;Acc:HGNC:20344] | 0.5985 | 5.45E-12 |
| ENSG00000072110 | ACTN1 | actinin alpha 1 [Source:HGNC Symbol;Acc:HGNC:163] | 0.8906 | 5.67E-12 |
| ENSG00000198863 | RUNDC1 | RUN domain containing 1 [Source:HGNC Symbol;Acc:HGNC:25418] | 0.8092 | 5.78E-12 |
| ENSG00000053372 | MRTO4 | MRT4 homolog, ribosome maturation factor [Source:HGNC Symbol;Acc:HGNC:18477] | 0.6288 | 5.84E-12 |
| ENSG00000111962 | UST | uronyl 2-sulfotransferase [Source:HGNC Symbol;Acc:HGNC:17223] | 1.3521 | 6.14E-12 |
| ENSG00000143126 | CELSR2 | cadherin EGF LAG seven-pass G-type receptor 2 [Source:HGNC Symbol;Acc:HGNC:3231] | -0.7169 | 6.38E-12 |
| ENSG00000157570 | TSPAN18 | tetraspanin 18 [Source:HGNC Symbol;Acc:HGNC:20660] | 1.5248 | 6.41E-12 |
| ENSG00000269964 | MEI4 | meiotic double-stranded break formation protein 4 [Source:HGNC Symbol;Acc:HGNC:43638] | -0.9760 | 6.55E-12 |
| ENSG00000250722 | SELENOP | selenoprotein P [Source:HGNC Symbol;Acc:HGNC:10751] | -0.6279 | 6.59E-12 |
| ENSG00000133135 | RNF128 | ring finger protein 128 [Source:HGNC Symbol;Acc:HGNC:21153] | 0.8193 | 6.60E-12 |
| ENSG00000128272 | ATF4 | activating transcription factor 4 [Source:HGNC Symbol;Acc:HGNC:786] | 0.7127 | 6.60E-12 |
| ENSG00000069011 | PITX1 | paired like homeodomain 1 [Source:HGNC Symbol;Acc:HGNC:9004] | -0.9272 | 6.67E-12 |
| ENSG00000151746 | BICD1 | BICD cargo adaptor 1 [Source:HGNC Symbol;Acc:HGNC:1049] | 1.0250 | 6.71E-12 |
| ENSG00000180901 | KCTD2 | potassium channel tetramerization domain containing 2 [Source:HGNC Symbol;Acc:HGNC:21294] | -0.6370 | 6.79E-12 |
| ENSG00000177432 | NAP1L5 | nucleosome assembly protein 1 like 5 [Source:HGNC Symbol;Acc:HGNC:19968] | 0.7559 | 7.05E-12 |
| ENSG00000120251 | GRIA2 | glutamate ionotropic receptor AMPA type subunit 2 [Source:HGNC Symbol;Acc:HGNC:4572] | -0.9039 | 7.57E-12 |
| ENSG00000197498 | RPF2 | ribosome production factor 2 homolog [Source:HGNC Symbol;Acc:HGNC:20870] | 1.0805 | 7.82E-12 |
| ENSG00000175592 | FOSL1 | FOS like 1, AP-1 transcription factor subunit [Source:HGNC Symbol;Acc:HGNC:13718] | 3.5519 | 8.20E-12 |
| ENSG00000124942 | AHNAK | AHNAK nucleoprotein [Source:HGNC Symbol;Acc:HGNC:347] | 2.9802 | 8.31E-12 |
| ENSG00000186472 | PCLO | piccolo presynaptic cytomatrix protein [Source:HGNC Symbol;Acc:HGNC:13406] | -0.9879 | 8.36E-12 |
| ENSG00000146729 | NIPSNAP2 | nipsnap homolog 2 [Source:HGNC Symbol;Acc:HGNC:4179] | -0.7213 | 8.42E-12 |
| ENSG00000183044 | ABAT | 4-aminobutyrate aminotransferase [Source:HGNC Symbol;Acc:HGNC:23] | -1.1206 | 8.52E-12 |
| ENSG00000138756 | BMP2K | BMP2 inducible kinase [Source:HGNC Symbol;Acc:HGNC:18041] | 0.8324 | 8.81E-12 |
| ENSG00000166411 | IDH3A | isocitrate dehydrogenase (NAD(+)) 3 catalytic subunit alpha [Source:HGNC Symbol;Acc:HGNC:5384] | 0.6470 | 8.89E-12 |
| ENSG00000169047 | IRS1 | insulin receptor substrate 1 [Source:HGNC Symbol;Acc:HGNC:6125] | 2.2385 | 8.94E-12 |
| ENSG00000197694 | SPTAN1 | spectrin alpha, non-erythrocytic 1 [Source:HGNC Symbol;Acc:HGNC:11273] | 0.5376 | 8.99E-12 |
| ENSG00000107262 | BAG1 | BAG cochaperone 1 [Source:HGNC Symbol;Acc:HGNC:937] | 0.6793 | 9.11E-12 |
| ENSG00000186376 | ZNF75D | zinc finger protein 75D [Source:HGNC Symbol;Acc:HGNC:13145] | -0.7367 | 9.40E-12 |
| ENSG00000177192 | PUS1 | pseudouridine synthase 1 [Source:HGNC Symbol;Acc:HGNC:15508] | 0.6976 | 9.52E-12 |
| ENSG00000157224 | CLDN12 | claudin 12 [Source:HGNC Symbol;Acc:HGNC:2034] | 0.5641 | 9.54E-12 |
| ENSG00000235106 | BRD3OS | BRD3 opposite strand [Source:HGNC Symbol;Acc:HGNC:24742] | -0.6858 | 9.65E-12 |

| | | | | |
|---|---|---|---|---|
| ENSG00000169783 | LINGO1 | leucine rich repeat and Ig domain containing 1 [Source:HGNC Symbol;Acc:HGNC:21205] | -1.5275 | 9.70E-12 |
| ENSG00000163346 | PBXIP1 | PBX homeobox interacting protein 1 [Source:HGNC Symbol;Acc:HGNC:21199] | -0.7568 | 9.84E-12 |
| ENSG00000109819 | PPARGC1A | PPARG coactivator 1 alpha [Source:HGNC Symbol;Acc:HGNC:9237] | -1.3056 | 1.09E-11 |
| ENSG00000126106 | TMEM53 | transmembrane protein 53 [Source:HGNC Symbol;Acc:HGNC:26186] | 1.0020 | 1.10E-11 |
| ENSG00000149639 | SOGA1 | suppressor of glucose, autophagy associated 1 [Source:HGNC Symbol;Acc:HGNC:16111] | -0.9696 | 1.11E-11 |
| ENSG00000124191 | TOX2 | TOX high mobility group box family member 2 [Source:HGNC Symbol;Acc:HGNC:16095] | 1.3813 | 1.14E-11 |
| ENSG00000137996 | RTCA | RNA 3'-terminal phosphate cyclase [Source:HGNC Symbol;Acc:HGNC:17981] | 0.5339 | 1.19E-11 |
| ENSG00000131831 | RAI2 | retinoic acid induced 2 [Source:HGNC Symbol;Acc:HGNC:9835] | -1.1501 | 1.22E-11 |
| ENSG00000134294 | SLC38A2 | solute carrier family 38 member 2 [Source:HGNC Symbol;Acc:HGNC:13448] | 0.5994 | 1.27E-11 |
| ENSG00000101346 | POFUT1 | protein O-fucosyltransferase 1 [Source:HGNC Symbol;Acc:HGNC:14988] | -0.5760 | 1.27E-11 |
| ENSG00000138386 | NAB1 | NGFI-A binding protein 1 [Source:HGNC Symbol;Acc:HGNC:7626] | 0.5371 | 1.29E-11 |
| ENSG00000145632 | PLK2 | polo like kinase 2 [Source:HGNC Symbol;Acc:HGNC:19699] | 2.3937 | 1.29E-11 |
| ENSG00000196611 | MMP1 | matrix metallopeptidase 1 [Source:HGNC Symbol;Acc:HGNC:7155] | 4.2925 | 1.30E-11 |
| ENSG00000196843 | ARID5A | AT-rich interaction domain 5A [Source:HGNC Symbol;Acc:HGNC:17361] | 2.4363 | 1.31E-11 |
| ENSG00000127870 | RNF6 | ring finger protein 6 [Source:HGNC Symbol;Acc:HGNC:10069] | 0.7076 | 1.33E-11 |
| ENSG00000204789 | ZNF204P | zinc finger protein 204, pseudogene [Source:HGNC Symbol;Acc:HGNC:12995] | -0.8608 | 1.34E-11 |
| ENSG00000136425 | CIB2 | calcium and integrin binding family member 2 [Source:HGNC Symbol;Acc:HGNC:24579] | -0.9762 | 1.37E-11 |
| ENSG00000072364 | AFF4 | AF4/FMR2 family member 4 [Source:HGNC Symbol;Acc:HGNC:17869] | 0.5093 | 1.38E-11 |
| ENSG00000198730 | CTR9 | CTR9 homolog, Paf1/RNA polymerase II complex component [Source:HGNC Symbol;Acc:HGNC:16850] | 0.5288 | 1.38E-11 |
| ENSG00000158258 | CLSTN2 | calsyntenin 2 [Source:HGNC Symbol;Acc:HGNC:17448] | 2.0533 | 1.39E-11 |
| ENSG00000171303 | KCNK3 | potassium two pore domain channel subfamily K member 3 [Source:HGNC Symbol;Acc:HGNC:6278] | 0.9082 | 1.43E-11 |
| ENSG00000114796 | KLHL24 | kelch like family member 24 [Source:HGNC Symbol;Acc:HGNC:25947] | -1.1641 | 1.48E-11 |
| ENSG00000163527 | STT3B | STT3 oligosaccharyltransferase complex catalytic subunit B [Source:HGNC Symbol;Acc:HGNC:30611] | 0.5914 | 1.49E-11 |
| ENSG00000092036 | HAUS4 | HAUS augmin like complex subunit 4 [Source:HGNC Symbol;Acc:HGNC:20163] | -0.8604 | 1.52E-11 |
| ENSG00000163617 | CCDC191 | coiled-coil domain containing 191 [Source:HGNC Symbol;Acc:HGNC:29272] | -1.5376 | 1.67E-11 |
| ENSG00000151729 | SLC25A4 | solute carrier family 25 member 4 [Source:HGNC Symbol;Acc:HGNC:10990] | 0.8685 | 1.69E-11 |
| ENSG00000174744 | BRMS1 | BRMS1 transcriptional repressor and anoikis regulator [Source:HGNC Symbol;Acc:HGNC:17262] | 0.6156 | 1.71E-11 |
| ENSG00000215256 | DHRS4-AS1 | DHRS4 antisense RNA 1 [Source:HGNC Symbol;Acc:HGNC:23175] | -0.5891 | 1.71E-11 |
| ENSG00000107897 | ACBD5 | acyl-CoA binding domain containing 5 [Source:HGNC Symbol;Acc:HGNC:23338] | -0.5245 | 1.72E-11 |
| ENSG00000131019 | ULBP3 | UL16 binding protein 3 [Source:HGNC Symbol;Acc:HGNC:14895] | -0.6431 | 1.77E-11 |
| ENSG00000157927 | RADIL | Rap associating with DIL domain [Source:HGNC Symbol;Acc:HGNC:22226] | -1.7599 | 1.78E-11 |
| ENSG00000148356 | LRSAM1 | leucine rich repeat and sterile alpha motif containing 1 [Source:HGNC Symbol;Acc:HGNC:25135] | -0.5788 | 1.83E-11 |
| ENSG00000171813 | PWWP2B | PWWP domain containing 2B [Source:HGNC Symbol;Acc:HGNC:25150] | -0.9263 | 1.86E-11 |
| ENSG00000019549 | SNAI2 | snail family transcriptional repressor 2 [Source:HGNC Symbol;Acc:HGNC:11094] | -0.8811 | 1.97E-11 |
| ENSG00000132879 | FBXO44 | F-box protein 44 [Source:HGNC Symbol;Acc:HGNC:24847] | -0.8478 | 1.97E-11 |
| ENSG00000101400 | SNTA1 | syntrophin alpha 1 [Source:HGNC Symbol;Acc:HGNC:11167] | -0.8494 | 1.98E-11 |
| ENSG00000148120 | AOPEP | aminopeptidase O (putative) [Source:HGNC Symbol;Acc:HGNC:1361] | -1.1416 | 1.98E-11 |
| ENSG00000139117 | CPNE8 | copine 8 [Source:HGNC Symbol;Acc:HGNC:23498] | -0.7178 | 1.99E-11 |
| ENSG00000188582 | PAQR9 | progestin and adipoQ receptor family member 9 [Source:HGNC Symbol;Acc:HGNC:30131] | 1.9946 | 2.08E-11 |
| ENSG00000156521 | TYSND1 | trypsin domain containing 1 [Source:HGNC Symbol;Acc:HGNC:28531] | -0.7447 | 2.08E-11 |
| ENSG00000134369 | NAV1 | neuron navigator 1 [Source:HGNC Symbol;Acc:HGNC:15989] | -0.8484 | 2.10E-11 |
| ENSG00000110852 | CLEC2B | C-type lectin domain family 2 member B [Source:HGNC Symbol;Acc:HGNC:2053] | 0.8558 | 2.11E-11 |
| ENSG00000269335 | IKBKG | inhibitor of nuclear factor kappa B kinase regulatory subunit gamma [Source:HGNC Symbol;Acc:HGNC:5961] | 0.5017 | 2.16E-11 |

| | | | | |
|---|---|---|---|---|
| ENSG00000107796 | ACTA2 | actin alpha 2, smooth muscle [Source:HGNC Symbol;Acc:HGNC:130] | 1.6024 | 2.18E-11 |
| ENSG00000170921 | TANC2 | tetratricopeptide repeat, ankyrin repeat and coiled-coil containing 2 [Source:HGNC Symbol;Acc:HGNC:30212] | 0.7134 | 2.21E-11 |
| ENSG00000262664 | OVCA2 | OVCA2 serine hydrolase domain containing [Source:HGNC Symbol;Acc:HGNC:24203] | 0.7189 | 2.26E-11 |
| ENSG00000164751 | PEX2 | peroxisomal biogenesis factor 2 [Source:HGNC Symbol;Acc:HGNC:9717] | -0.7136 | 2.28E-11 |
| ENSG00000135999 | EPC2 | enhancer of polycomb homolog 2 [Source:HGNC Symbol;Acc:HGNC:24543] | 0.5864 | 2.28E-11 |
| ENSG00000137040 | RANBP6 | RAN binding protein 6 [Source:HGNC Symbol;Acc:HGNC:9851] | 0.6465 | 2.28E-11 |
| ENSG00000104833 | TUBB4A | tubulin beta 4A class IVa [Source:HGNC Symbol;Acc:HGNC:20774] | -1.1626 | 2.30E-11 |
| ENSG00000270757 | HSPE1-MOB4 | HSPE1-MOB4 readthrough [Source:HGNC Symbol;Acc:HGNC:49184] | 0.9300 | 2.31E-11 |
| ENSG00000102978 | POLR2C | RNA polymerase II subunit C [Source:HGNC Symbol;Acc:HGNC:9189] | 0.5012 | 2.33E-11 |
| ENSG00000160818 | GPATCH4 | G-patch domain containing 4 [Source:HGNC Symbol;Acc:HGNC:25982] | 0.8177 | 2.37E-11 |
| ENSG00000130066 | SAT1 | spermidine/spermine N1-acetyltransferase 1 [Source:HGNC Symbol;Acc:HGNC:10540] | 1.2975 | 2.46E-11 |
| ENSG00000153896 | ZNF599 | zinc finger protein 599 [Source:HGNC Symbol;Acc:HGNC:26408] | -0.8031 | 2.52E-11 |
| ENSG00000163001 | CFAP36 | cilia and flagella associated protein 36 [Source:HGNC Symbol;Acc:HGNC:30540] | 0.7842 | 2.54E-11 |
| ENSG00000169855 | ROBO1 | roundabout guidance receptor 1 [Source:HGNC Symbol;Acc:HGNC:10249] | 1.1554 | 2.57E-11 |
| ENSG00000152049 | KCNE4 | potassium voltage-gated channel subfamily E regulatory subunit 4 [Source:HGNC Symbol;Acc:HGNC:6244] | 1.2979 | 2.57E-11 |
| ENSG00000141971 | MVB12A | multivesicular body subunit 12A [Source:HGNC Symbol;Acc:HGNC:25153] | -0.7525 | 2.58E-11 |
| ENSG00000152409 | JMY | junction mediating and regulatory protein, p53 cofactor [Source:HGNC Symbol;Acc:HGNC:28916] | -0.9186 | 2.64E-11 |
| ENSG00000175575 | PAAF1 | proteasomal ATPase associated factor 1 [Source:HGNC Symbol;Acc:HGNC:25687] | -0.8347 | 2.71E-11 |
| ENSG00000165617 | DACT1 | dishevelled binding antagonist of beta catenin 1 [Source:HGNC Symbol;Acc:HGNC:17748] | -1.3361 | 2.74E-11 |
| ENSG00000105991 | HOXA1 | homeobox A1 [Source:HGNC Symbol;Acc:HGNC:5099] | 1.4337 | 2.86E-11 |
| ENSG00000261377 | PDCD6IPP2 | PDCD6IP pseudogene 2 [Source:HGNC Symbol;Acc:HGNC:49873] | -1.0972 | 2.87E-11 |
| ENSG00000106025 | TSPAN12 | tetraspanin 12 [Source:HGNC Symbol;Acc:HGNC:21641] | 0.8610 | 3.00E-11 |
| ENSG00000183722 | LHFPL6 | LHFPL tetraspan subfamily member 6 [Source:HGNC Symbol;Acc:HGNC:6586] | -0.7705 | 3.04E-11 |
| ENSG00000101236 | RNF24 | ring finger protein 24 [Source:HGNC Symbol;Acc:HGNC:13779] | 0.8189 | 3.05E-11 |
| ENSG00000121966 | CXCR4 | C-X-C motif chemokine receptor 4 [Source:HGNC Symbol;Acc:HGNC:2561] | 0.8893 | 3.14E-11 |
| ENSG00000162804 | SNED1 | sushi, nidogen and EGF like domains 1 [Source:HGNC Symbol;Acc:HGNC:24696] | 1.4500 | 3.22E-11 |
| ENSG00000197712 | FAM114A1 | family with sequence similarity 114 member A1 [Source:HGNC Symbol;Acc:HGNC:25087] | -0.7774 | 3.23E-11 |
| ENSG00000198053 | SIRPA | signal regulatory protein alpha [Source:HGNC Symbol;Acc:HGNC:9662] | -0.5117 | 3.27E-11 |
| ENSG00000121073 | SLC35B1 | solute carrier family 35 member B1 [Source:HGNC Symbol;Acc:HGNC:20798] | 0.7738 | 3.27E-11 |
| ENSG00000073008 | PVR | PVR cell adhesion molecule [Source:HGNC Symbol;Acc:HGNC:9705] | 0.7649 | 3.29E-11 |
| ENSG00000128655 | PDE11A | phosphodiesterase 11A [Source:HGNC Symbol;Acc:HGNC:8773] | -1.7545 | 3.36E-11 |
| ENSG00000205413 | SAMD9 | sterile alpha motif domain containing 9 [Source:HGNC Symbol;Acc:HGNC:1348] | -0.6092 | 3.39E-11 |
| ENSG00000138018 | SELENOI | selenoprotein I [Source:HGNC Symbol;Acc:HGNC:29361] | 0.5081 | 3.58E-11 |
| ENSG00000197562 | RAB40C | RAB40C, member RAS oncogene family [Source:HGNC Symbol;Acc:HGNC:18285] | -0.8272 | 3.58E-11 |
| ENSG00000183655 | KLHL25 | kelch like family member 25 [Source:HGNC Symbol;Acc:HGNC:25732] | 0.9278 | 3.86E-11 |
| ENSG00000005073 | HOXA11 | homeobox A11 [Source:HGNC Symbol;Acc:HGNC:5101] | 0.8784 | 3.93E-11 |
| ENSG00000130560 | UBAC1 | UBA domain containing 1 [Source:HGNC Symbol;Acc:HGNC:30221] | -0.6237 | 3.98E-11 |
| ENSG00000125434 | SLC25A35 | solute carrier family 25 member 35 [Source:HGNC Symbol;Acc:HGNC:31921] | -0.9417 | 4.04E-11 |
| ENSG00000139549 | DHH | desert hedgehog signaling molecule [Source:HGNC Symbol;Acc:HGNC:2865] | 2.3223 | 4.10E-11 |
| ENSG00000070756 | PABPC1 | poly(A) binding protein cytoplasmic 1 [Source:HGNC Symbol;Acc:HGNC:8554] | 0.7669 | 4.16E-11 |
| ENSG00000147676 | MAL2 | mal, T cell differentiation protein 2 (gene/pseudogene) [Source:HGNC Symbol;Acc:HGNC:13634] | 0.6477 | 4.26E-11 |
| ENSG00000183117 | CSMD1 | CUB and Sushi multiple domains 1 [Source:HGNC Symbol;Acc:HGNC:14026] | -1.4323 | 4.47E-11 |

| | | | | |
|---|---|---|---|---|
| ENSG00000153989 | NUS1 | NUS1 dehydrodolichyl diphosphate synthase subunit [Source:HGNC Symbol;Acc:HGNC:21042] | 0.5340 | 4.65E-11 |
| ENSG00000104946 | TBC1D17 | TBC1 domain family member 17 [Source:HGNC Symbol;Acc:HGNC:25699] | -0.6324 | 4.69E-11 |
| ENSG00000102962 | CCL22 | C-C motif chemokine ligand 22 [Source:HGNC Symbol;Acc:HGNC:10621] | 0.9688 | 4.82E-11 |
| ENSG00000106591 | MRPL32 | mitochondrial ribosomal protein L32 [Source:HGNC Symbol;Acc:HGNC:14035] | 0.5916 | 4.83E-11 |
| ENSG00000138769 | CDKL2 | cyclin dependent kinase like 2 [Source:HGNC Symbol;Acc:HGNC:1782] | -1.8539 | 4.90E-11 |
| ENSG00000050748 | MAPK9 | mitogen-activated protein kinase 9 [Source:HGNC Symbol;Acc:HGNC:6886] | -0.6198 | 4.95E-11 |
| ENSG00000273270 | NA | novel transcript | -0.7723 | 5.10E-11 |
| ENSG00000267278 | MAP3K14-AS1 | MAP3K14 antisense RNA 1 [Source:HGNC Symbol;Acc:HGNC:44359] | 1.0798 | 5.20E-11 |
| ENSG00000162430 | SELENON | selenoprotein N [Source:HGNC Symbol;Acc:HGNC:15999] | -0.5173 | 5.22E-11 |
| ENSG00000092969 | TGFB2 | transforming growth factor beta 2 [Source:HGNC Symbol;Acc:HGNC:11768] | 1.4746 | 5.34E-11 |
| ENSG00000239887 | C1orf226 | chromosome 1 open reading frame 226 [Source:HGNC Symbol;Acc:HGNC:34351] | -0.9580 | 5.37E-11 |
| ENSG00000132432 | SEC61G | SEC61 translocon subunit gamma [Source:HGNC Symbol;Acc:HGNC:18277] | 0.6381 | 5.39E-11 |
| ENSG00000163479 | SSR2 | signal sequence receptor subunit 2 [Source:HGNC Symbol;Acc:HGNC:11324] | 0.6593 | 5.49E-11 |
| ENSG00000101596 | SMCHD1 | structural maintenance of chromosomes flexible hinge domain containing 1 [Source:HGNC Symbol;Acc:HGNC:29090] | 0.5589 | 5.52E-11 |
| ENSG00000139344 | AMDHD1 | amidohydrolase domain containing 1 [Source:HGNC Symbol;Acc:HGNC:28577] | 1.1447 | 5.52E-11 |
| ENSG00000180628 | PCGF5 | polycomb group ring finger 5 [Source:HGNC Symbol;Acc:HGNC:28264] | 0.6482 | 5.64E-11 |
| ENSG00000104691 | UBXN8 | UBX domain protein 8 [Source:HGNC Symbol;Acc:HGNC:30307] | 1.2301 | 5.70E-11 |
| ENSG00000141252 | VPS53 | VPS53 subunit of GARP complex [Source:HGNC Symbol;Acc:HGNC:25608] | 0.9937 | 5.70E-11 |
| ENSG00000047365 | ARAP2 | ArfGAP with RhoGAP domain, ankyrin repeat and PH domain 2 [Source:HGNC Symbol;Acc:HGNC:16924] | 1.0728 | 5.95E-11 |
| ENSG00000102908 | NFAT5 | nuclear factor of activated T cells 5 [Source:HGNC Symbol;Acc:HGNC:7774] | 0.6062 | 5.95E-11 |
| ENSG00000176915 | ANKLE2 | ankyrin repeat and LEM domain containing 2 [Source:HGNC Symbol;Acc:HGNC:29101] | 0.5488 | 6.12E-11 |
| ENSG00000038532 | CLEC16A | C-type lectin domain containing 16A [Source:HGNC Symbol;Acc:HGNC:29013] | 0.6944 | 6.16E-11 |
| ENSG00000124783 | SSR1 | signal sequence receptor subunit 1 [Source:HGNC Symbol;Acc:HGNC:11323] | 0.6613 | 6.30E-11 |
| ENSG00000132204 | LINC00470 | long intergenic non-protein coding RNA 470 [Source:HGNC Symbol;Acc:HGNC:1225] | -1.8844 | 6.54E-11 |
| ENSG00000166068 | SPRED1 | sprouty related EVH1 domain containing 1 [Source:HGNC Symbol;Acc:HGNC:20249] | 1.3133 | 6.54E-11 |
| ENSG00000263155 | MYZAP | myocardial zonula adherens protein [Source:HGNC Symbol;Acc:HGNC:43444] | 0.9866 | 6.71E-11 |
| ENSG00000148219 | ASTN2 | astrotactin 2 [Source:HGNC Symbol;Acc:HGNC:17021] | -1.4141 | 6.77E-11 |
| ENSG00000153902 | LGI4 | leucine rich repeat LGI family member 4 [Source:HGNC Symbol;Acc:HGNC:18712] | -0.9580 | 6.80E-11 |
| ENSG00000127084 | FGD3 | FYVE, RhoGEF and PH domain containing 3 [Source:HGNC Symbol;Acc:HGNC:16027] | -1.6224 | 6.90E-11 |
| ENSG00000224389 | C4B | complement C4B (Chido blood group) [Source:HGNC Symbol;Acc:HGNC:1324] | 1.4300 | 6.90E-11 |
| ENSG00000074695 | LMAN1 | lectin, mannose binding 1 [Source:HGNC Symbol;Acc:HGNC:6631] | 0.5690 | 6.93E-11 |
| ENSG00000168765 | GSTM4 | glutathione S-transferase mu 4 [Source:HGNC Symbol;Acc:HGNC:4636] | -0.9134 | 6.95E-11 |
| ENSG00000152669 | CCNO | cyclin O [Source:HGNC Symbol;Acc:HGNC:18576] | 1.4689 | 7.16E-11 |
| ENSG00000196576 | PLXNB2 | plexin B2 [Source:HGNC Symbol;Acc:HGNC:9104] | 0.6616 | 7.19E-11 |
| ENSG00000008256 | CYTH3 | cytohesin 3 [Source:HGNC Symbol;Acc:HGNC:9504] | -0.5444 | 7.25E-11 |
| ENSG00000103485 | QPRT | quinolinate phosphoribosyltransferase [Source:HGNC Symbol;Acc:HGNC:9755] | -0.8540 | 7.29E-11 |
| ENSG00000119737 | GPR75 | G protein-coupled receptor 75 [Source:HGNC Symbol;Acc:HGNC:4526] | 1.2373 | 7.50E-11 |
| ENSG00000107372 | ZFAND5 | zinc finger AN1-type containing 5 [Source:HGNC Symbol;Acc:HGNC:13008] | 0.7715 | 7.55E-11 |
| ENSG00000105784 | RUNDC3B | RUN domain containing 3B [Source:HGNC Symbol;Acc:HGNC:30286] | -0.8386 | 7.68E-11 |
| ENSG00000107833 | NPM3 | nucleophosmin/nucleoplasmin 3 [Source:HGNC Symbol;Acc:HGNC:7931] | 0.7487 | 7.73E-11 |
| ENSG00000242207 | HOXB-AS4 | HOXB cluster antisense RNA 4 [Source:HGNC Symbol;Acc:HGNC:40285] | 0.9381 | 7.76E-11 |
| ENSG00000240891 | PLCXD2 | phosphatidylinositol specific phospholipase C X domain containing 2 [Source:HGNC Symbol;Acc:HGNC:26462] | 1.0294 | 7.94E-11 |

| | | | | |
|---|---|---|---|---|
| ENSG00000189195 | BTBD8 | BTB domain containing 8 [Source:HGNC Symbol;Acc:HGNC:21019] | -1.1640 | 7.95E-11 |
| ENSG00000178585 | CTNNBIP1 | catenin beta interacting protein 1 [Source:HGNC Symbol;Acc:HGNC:16913] | -1.0100 | 8.04E-11 |
| ENSG00000129991 | TNNI3 | troponin I3, cardiac type [Source:HGNC Symbol;Acc:HGNC:11947] | -0.8032 | 8.23E-11 |
| ENSG00000236008 | LINC01814 | long intergenic non-protein coding RNA 1814 [Source:HGNC Symbol;Acc:HGNC:52618] | -0.9443 | 8.25E-11 |
| ENSG00000108840 | HDAC5 | histone deacetylase 5 [Source:HGNC Symbol;Acc:HGNC:14068] | 1.1324 | 8.29E-11 |
| ENSG00000152133 | GPATCH11 | G-patch domain containing 11 [Source:HGNC Symbol;Acc:HGNC:26768] | 0.6350 | 8.32E-11 |
| ENSG00000118495 | PLAGL1 | PLAG1 like zinc finger 1 [Source:HGNC Symbol;Acc:HGNC:9046] | -0.6158 | 8.64E-11 |
| ENSG00000119922 | IFIT2 | interferon induced protein with tetratricopeptide repeats 2 [Source:HGNC Symbol;Acc:HGNC:5409] | 1.1668 | 8.95E-11 |
| ENSG00000116649 | SRM | spermidine synthase [Source:HGNC Symbol;Acc:HGNC:11296] | 0.8423 | 8.97E-11 |
| ENSG00000166917 | MIR202HG | MIR202 host gene [Source:HGNC Symbol;Acc:HGNC:49402] | -2.1649 | 9.28E-11 |
| ENSG00000138030 | KHK | ketohexokinase [Source:HGNC Symbol;Acc:HGNC:6315] | -0.9011 | 9.37E-11 |
| ENSG00000115159 | GPD2 | glycerol-3-phosphate dehydrogenase 2 [Source:HGNC Symbol;Acc:HGNC:4456] | 0.9667 | 1.02E-10 |
| ENSG00000167749 | KLK4 | kallikrein related peptidase 4 [Source:HGNC Symbol;Acc:HGNC:6365] | -2.5132 | 1.02E-10 |
| ENSG00000043143 | JADE2 | jade family PHD finger 2 [Source:HGNC Symbol;Acc:HGNC:22984] | -0.5961 | 1.04E-10 |
| ENSG00000150281 | CTF1 | cardiotrophin 1 [Source:HGNC Symbol;Acc:HGNC:2499] | -1.3265 | 1.05E-10 |
| ENSG00000108604 | SMARCD2 | SWI/SNF related, matrix associated, actin dependent regulator of chromatin, subfamily d, member 2 [Source:HGNC Symbol;Acc:HGNC:11107] | -0.9118 | 1.06E-10 |
| ENSG00000132623 | ANKEF1 | ankyrin repeat and EF-hand domain containing 1 [Source:HGNC Symbol;Acc:HGNC:15803] | -0.9208 | 1.07E-10 |
| ENSG00000077232 | DNAJC10 | DnaJ heat shock protein family (Hsp40) member C10 [Source:HGNC Symbol;Acc:HGNC:24637] | 0.7459 | 1.09E-10 |
| ENSG00000128590 | DNAJB9 | DnaJ heat shock protein family (Hsp40) member B9 [Source:HGNC Symbol;Acc:HGNC:6968] | 0.9922 | 1.10E-10 |
| ENSG00000186871 | ERCC6L | ERCC excision repair 6 like, spindle assembly checkpoint helicase [Source:HGNC Symbol;Acc:HGNC:20794] | -0.7561 | 1.13E-10 |
| ENSG00000166200 | COPS2 | COP9 signalosome subunit 2 [Source:HGNC Symbol;Acc:HGNC:30747] | 0.5170 | 1.14E-10 |
| ENSG00000138434 | ITPRID2 | ITPR interacting domain containing 2 [Source:HGNC Symbol;Acc:HGNC:11319] | 0.7397 | 1.14E-10 |
| ENSG00000119283 | TRIM67 | tripartite motif containing 67 [Source:HGNC Symbol;Acc:HGNC:31859] | 2.9949 | 1.15E-10 |
| ENSG00000126785 | RHOJ | ras homolog family member J [Source:HGNC Symbol;Acc:HGNC:688] | 1.2514 | 1.17E-10 |
| ENSG00000105538 | RASIP1 | Ras interacting protein 1 [Source:HGNC Symbol;Acc:HGNC:24716] | -1.1283 | 1.22E-10 |
| ENSG00000148834 | GSTO1 | glutathione S-transferase omega 1 [Source:HGNC Symbol;Acc:HGNC:13312] | 0.5534 | 1.27E-10 |
| ENSG00000154734 | ADAMTS1 | ADAM metallopeptidase with thrombospondin type 1 motif 1 [Source:HGNC Symbol;Acc:HGNC:217] | 2.2753 | 1.27E-10 |
| ENSG00000178234 | GALNT11 | polypeptide N-acetylgalactosaminyltransferase 11 [Source:HGNC Symbol;Acc:HGNC:19875] | -0.7789 | 1.29E-10 |
| ENSG00000109133 | TMEM33 | transmembrane protein 33 [Source:HGNC Symbol;Acc:HGNC:25541] | 0.5492 | 1.30E-10 |
| ENSG00000049130 | KITLG | KIT ligand [Source:HGNC Symbol;Acc:HGNC:6343] | 1.1732 | 1.32E-10 |
| ENSG00000024862 | CCDC28A | coiled-coil domain containing 28A [Source:HGNC Symbol;Acc:HGNC:21098] | -0.7451 | 1.34E-10 |
| ENSG00000112685 | EXOC2 | exocyst complex component 2 [Source:HGNC Symbol;Acc:HGNC:24968] | -0.5476 | 1.34E-10 |
| ENSG00000149679 | CABLES2 | Cdk5 and Abl enzyme substrate 2 [Source:HGNC Symbol;Acc:HGNC:16143] | -1.2120 | 1.40E-10 |
| ENSG00000164300 | SERINC5 | serine incorporator 5 [Source:HGNC Symbol;Acc:HGNC:18825] | -0.8690 | 1.41E-10 |
| ENSG00000102755 | FLT1 | fms related receptor tyrosine kinase 1 [Source:HGNC Symbol;Acc:HGNC:3763] | 5.4892 | 1.42E-10 |
| ENSG00000111907 | TPD52L1 | TPD52 like 1 [Source:HGNC Symbol;Acc:HGNC:12006] | 0.5673 | 1.42E-10 |
| ENSG00000166963 | MAP1A | microtubule associated protein 1A [Source:HGNC Symbol;Acc:HGNC:6835] | -1.1153 | 1.42E-10 |
| ENSG00000272779 | NA | BMS1, ribosome biogenesis factor (BSM1) pseudogene | 0.9101 | 1.55E-10 |
| ENSG00000163811 | WDR43 | WD repeat domain 43 [Source:HGNC Symbol;Acc:HGNC:28945] | 0.8552 | 1.56E-10 |
| ENSG00000213585 | VDAC1 | voltage dependent anion channel 1 [Source:HGNC Symbol;Acc:HGNC:12669] | 0.5919 | 1.58E-10 |
| ENSG00000168079 | SCARA5 | scavenger receptor class A member 5 [Source:HGNC Symbol;Acc:HGNC:28701] | -2.0421 | 1.62E-10 |
| ENSG00000196449 | YRDC | yrdC N6-threonylcarbamoyltransferase domain containing [Source:HGNC Symbol;Acc:HGNC:28905] | 0.9599 | 1.64E-10 |

| | | | | |
|---|---|---|---|---|
| ENSG00000133958 | UNC79 | unc-79 homolog, NALCN channel complex subunit [Source:HGNC Symbol;Acc:HGNC:19966] | 1.1632 | 1.64E-10 |
| ENSG00000156097 | GPR61 | G protein-coupled receptor 61 [Source:HGNC Symbol;Acc:HGNC:13300] | -1.4932 | 1.67E-10 |
| ENSG00000149948 | HMGA2 | high mobility group AT-hook 2 [Source:HGNC Symbol;Acc:HGNC:5009] | 1.3602 | 1.68E-10 |
| ENSG00000142632 | ARHGEF19 | Rho guanine nucleotide exchange factor 19 [Source:HGNC Symbol;Acc:HGNC:26604] | 0.7996 | 1.71E-10 |
| ENSG00000122335 | SERAC1 | serine active site containing 1 [Source:HGNC Symbol;Acc:HGNC:21061] | -0.7419 | 1.71E-10 |
| ENSG00000140548 | ZNF710 | zinc finger protein 710 [Source:HGNC Symbol;Acc:HGNC:25352] | -1.0093 | 1.74E-10 |
| ENSG00000074855 | ANO8 | anoctamin 8 [Source:HGNC Symbol;Acc:HGNC:29329] | -0.9351 | 1.80E-10 |
| ENSG00000117724 | CENPF | centromere protein F [Source:HGNC Symbol;Acc:HGNC:1857] | -0.5484 | 1.84E-10 |
| ENSG00000251003 | ZFPM2-AS1 | ZFPM2 antisense RNA 1 [Source:HGNC Symbol;Acc:HGNC:50698] | 0.9409 | 1.91E-10 |
| ENSG00000132467 | UTP3 | UTP3 small subunit processome component [Source:HGNC Symbol;Acc:HGNC:24477] | 0.7318 | 1.96E-10 |
| ENSG00000278291 | NA | novel transcript, antisense to IL17D | 1.0627 | 2.05E-10 |
| ENSG00000162545 | CAMK2N1 | calcium/calmodulin dependent protein kinase II inhibitor 1 [Source:HGNC Symbol;Acc:HGNC:24190] | 1.3932 | 2.07E-10 |
| ENSG00000136628 | EPRS1 | glutamyl-prolyl-tRNA synthetase 1 [Source:HGNC Symbol;Acc:HGNC:3418] | -0.9364 | 2.07E-10 |
| ENSG00000174953 | DHX36 | DEAH-box helicase 36 [Source:HGNC Symbol;Acc:HGNC:14410] | 0.5766 | 2.30E-10 |
| ENSG00000131650 | KREMEN2 | kringle containing transmembrane protein 2 [Source:HGNC Symbol;Acc:HGNC:18797] | -1.1373 | 2.33E-10 |
| ENSG00000138162 | TACC2 | transforming acidic coiled-coil containing protein 2 [Source:HGNC Symbol;Acc:HGNC:11523] | 2.1655 | 2.34E-10 |
| ENSG00000124098 | FAM210B | family with sequence similarity 210 member B [Source:HGNC Symbol;Acc:HGNC:16102] | -0.5706 | 2.47E-10 |
| ENSG00000088205 | DDX18 | DEAD-box helicase 18 [Source:HGNC Symbol;Acc:HGNC:2741] | 0.6142 | 2.49E-10 |
| ENSG00000251350 | LINC02475 | long intergenic non-protein coding RNA 2475 [Source:HGNC Symbol;Acc:HGNC:53418] | -2.5244 | 2.51E-10 |
| ENSG00000157985 | AGAP1 | ArfGAP with GTPase domain, ankyrin repeat and PH domain 1 [Source:HGNC Symbol;Acc:HGNC:16922] | 0.7943 | 2.53E-10 |
| ENSG00000144331 | ZNF385B | zinc finger protein 385B [Source:HGNC Symbol;Acc:HGNC:26332] | -1.1261 | 2.57E-10 |
| ENSG00000104907 | TRMT1 | tRNA methyltransferase 1 [Source:HGNC Symbol;Acc:HGNC:25980] | 0.6743 | 2.60E-10 |
| ENSG00000091592 | NLRP1 | NLR family pyrin domain containing 1 [Source:HGNC Symbol;Acc:HGNC:14374] | 0.8515 | 2.66E-10 |
| ENSG00000189184 | PCDH18 | protocadherin 18 [Source:HGNC Symbol;Acc:HGNC:14268] | -1.2174 | 2.66E-10 |
| ENSG00000120738 | EGR1 | early growth response 1 [Source:HGNC Symbol;Acc:HGNC:3238] | 4.0713 | 2.71E-10 |
| ENSG00000162627 | SNX7 | sorting nexin 7 [Source:HGNC Symbol;Acc:HGNC:14971] | -0.5840 | 2.71E-10 |
| ENSG00000163412 | EIF4E3 | eukaryotic translation initiation factor 4E family member 3 [Source:HGNC Symbol;Acc:HGNC:31837] | -1.0270 | 2.88E-10 |
| ENSG00000115935 | WIPF1 | WAS/WASL interacting protein family member 1 [Source:HGNC Symbol;Acc:HGNC:12736] | 0.6940 | 2.88E-10 |
| ENSG00000146072 | TNFRSF21 | TNF receptor superfamily member 21 [Source:HGNC Symbol;Acc:HGNC:13469] | 2.6402 | 2.98E-10 |
| ENSG00000165209 | STRBP | spermatid perinuclear RNA binding protein [Source:HGNC Symbol;Acc:HGNC:16462] | -0.7029 | 3.00E-10 |
| ENSG00000110047 | EHD1 | EH domain containing 1 [Source:HGNC Symbol;Acc:HGNC:3242] | -0.9634 | 3.02E-10 |
| ENSG00000115840 | SLC25A12 | solute carrier family 25 member 12 [Source:HGNC Symbol;Acc:HGNC:10982] | -0.5240 | 3.04E-10 |
| ENSG00000139044 | B4GALNT3 | beta-1,4-N-acetyl-galactosaminyltransferase 3 [Source:HGNC Symbol;Acc:HGNC:24137] | -0.5523 | 3.09E-10 |
| ENSG00000197021 | EOLA2 | endothelium and lymphocyte associated ASCH domain 2 [Source:HGNC Symbol;Acc:HGNC:17402] | 0.6269 | 3.12E-10 |
| ENSG00000127419 | TMEM175 | transmembrane protein 175 [Source:HGNC Symbol;Acc:HGNC:28709] | -0.9658 | 3.21E-10 |
| ENSG00000151292 | CSNK1G3 | casein kinase 1 gamma 3 [Source:HGNC Symbol;Acc:HGNC:2456] | 0.5150 | 3.29E-10 |
| ENSG00000140564 | FURIN | furin, paired basic amino acid cleaving enzyme [Source:HGNC Symbol;Acc:HGNC:8568] | 0.6980 | 3.39E-10 |
| ENSG00000186063 | AIDA | axin interactor, dorsalization associated [Source:HGNC Symbol;Acc:HGNC:25761] | 0.5321 | 3.40E-10 |
| ENSG00000128309 | MPST | mercaptopyruvate sulfurtransferase [Source:HGNC Symbol;Acc:HGNC:7223] | -0.6926 | 3.42E-10 |
| ENSG00000162755 | KLHDC9 | kelch domain containing 9 [Source:HGNC Symbol;Acc:HGNC:28489] | -1.8102 | 3.44E-10 |
| ENSG00000112578 | BYSL | bystin like [Source:HGNC Symbol;Acc:HGNC:1157] | 0.7798 | 3.46E-10 |
| ENSG00000177239 | MAN1B1 | mannosidase alpha class 1B member 1 [Source:HGNC Symbol;Acc:HGNC:6823] | -0.6584 | 3.71E-10 |
| ENSG00000152484 | USP12 | ubiquitin specific peptidase 12 [Source:HGNC Symbol;Acc:HGNC:20485] | 0.5310 | 3.81E-10 |

| | | | | |
|---|---|---|---|---|
| ENSG00000130881 | LRP3 | LDL receptor related protein 3 [Source:HGNC Symbol;Acc:HGNC:6695] | -0.6958 | 3.83E-10 |
| ENSG00000117410 | ATP6V0B | ATPase H+ transporting V0 subunit b [Source:HGNC Symbol;Acc:HGNC:861] | 0.5172 | 3.84E-10 |
| ENSG00000165175 | MID1IP1 | MID1 interacting protein 1 [Source:HGNC Symbol;Acc:HGNC:20715] | -0.5877 | 3.97E-10 |
| ENSG00000198780 | FAM169A | family with sequence similarity 169 member A [Source:HGNC Symbol;Acc:HGNC:29138] | 0.8974 | 4.01E-10 |
| ENSG00000137693 | YAP1 | Yes1 associated transcriptional regulator [Source:HGNC Symbol;Acc:HGNC:16262] | 0.8052 | 4.03E-10 |
| ENSG00000091136 | LAMB1 | laminin subunit beta 1 [Source:HGNC Symbol;Acc:HGNC:6486] | -0.5713 | 4.03E-10 |
| ENSG00000287059 | NA | novel transcript | -1.1362 | 4.18E-10 |
| ENSG00000173486 | FKBP2 | FKBP prolyl isomerase 2 [Source:HGNC Symbol;Acc:HGNC:3718] | 0.5324 | 4.24E-10 |
| ENSG00000177868 | SVBP | small vasohibin binding protein [Source:HGNC Symbol;Acc:HGNC:29204] | -1.0066 | 4.41E-10 |
| ENSG00000100142 | POLR2F | RNA polymerase II subunit F [Source:HGNC Symbol;Acc:HGNC:9193] | 0.5176 | 4.54E-10 |
| ENSG00000099331 | MYO9B | myosin IXB [Source:HGNC Symbol;Acc:HGNC:7609] | 0.7602 | 4.62E-10 |
| ENSG00000145147 | SLIT2 | slit guidance ligand 2 [Source:HGNC Symbol;Acc:HGNC:11086] | 1.0798 | 4.70E-10 |
| ENSG00000233429 | HOTAIRM1 | HOXA transcript antisense RNA, myeloid-specific 1 [Source:HGNC Symbol;Acc:HGNC:37117] | 1.1430 | 4.95E-10 |
| ENSG00000143641 | GALNT2 | polypeptide N-acetylgalactosaminyltransferase 2 [Source:HGNC Symbol;Acc:HGNC:4124] | 0.6995 | 4.95E-10 |
| ENSG00000176749 | CDK5R1 | cyclin dependent kinase 5 regulatory subunit 1 [Source:HGNC Symbol;Acc:HGNC:1775] | -0.9832 | 4.97E-10 |
| ENSG00000071537 | SEL1L | SEL1L adaptor subunit of ERAD E3 ubiquitin ligase [Source:HGNC Symbol;Acc:HGNC:10717] | 0.7701 | 5.03E-10 |
| ENSG00000162032 | SPSB3 | splA/ryanodine receptor domain and SOCS box containing 3 [Source:HGNC Symbol;Acc:HGNC:30629] | -0.6743 | 5.12E-10 |
| ENSG00000170892 | TSEN34 | tRNA splicing endonuclease subunit 34 [Source:HGNC Symbol;Acc:HGNC:15506] | -0.5102 | 5.13E-10 |
| ENSG00000134107 | BHLHE40 | basic helix-loop-helix family member e40 [Source:HGNC Symbol;Acc:HGNC:1046] | 2.7080 | 5.23E-10 |
| ENSG00000136297 | MMD2 | monocyte to macrophage differentiation associated 2 [Source:HGNC Symbol;Acc:HGNC:30133] | -1.4130 | 5.27E-10 |
| ENSG00000170558 | CDH2 | cadherin 2 [Source:HGNC Symbol;Acc:HGNC:1759] | 0.8491 | 5.33E-10 |
| ENSG00000146476 | ARMT1 | acidic residue methyltransferase 1 [Source:HGNC Symbol;Acc:HGNC:17872] | -0.6066 | 5.36E-10 |
| ENSG00000170264 | FAM161A | FAM161 centrosomal protein A [Source:HGNC Symbol;Acc:HGNC:25808] | -0.8931 | 5.51E-10 |
| ENSG00000071243 | ING3 | inhibitor of growth family member 3 [Source:HGNC Symbol;Acc:HGNC:14587] | 0.6400 | 5.57E-10 |
| ENSG00000165716 | DIPK1B | divergent protein kinase domain 1B [Source:HGNC Symbol;Acc:HGNC:28290] | -0.5758 | 5.68E-10 |
| ENSG00000165152 | PGAP4 | post-GPI attachment to proteins GalNAc transferase 4 [Source:HGNC Symbol;Acc:HGNC:28180] | -0.5504 | 5.73E-10 |
| ENSG00000138069 | RAB1A | RAB1A, member RAS oncogene family [Source:HGNC Symbol;Acc:HGNC:9758] | 0.5630 | 5.83E-10 |
| ENSG00000136783 | NIPSNAP3A | nipsnap homolog 3A [Source:HGNC Symbol;Acc:HGNC:23619] | -0.5263 | 5.87E-10 |
| ENSG00000229334 | NA | novel transcript | 2.0161 | 5.92E-10 |
| ENSG00000228300 | FAM174C | family with sequence similarity 174 member C [Source:HGNC Symbol;Acc:HGNC:26073] | 0.5826 | 5.93E-10 |
| ENSG00000084676 | NCOA1 | nuclear receptor coactivator 1 [Source:HGNC Symbol;Acc:HGNC:7668] | -0.5212 | 6.11E-10 |
| ENSG00000179918 | SEPHS2 | selenophosphate synthetase 2 [Source:HGNC Symbol;Acc:HGNC:19686] | 0.5348 | 6.35E-10 |
| ENSG00000173281 | PPP1R3B | protein phosphatase 1 regulatory subunit 3B [Source:HGNC Symbol;Acc:HGNC:14942] | -1.4931 | 6.55E-10 |
| ENSG00000080854 | IGSF9B | immunoglobulin superfamily member 9B [Source:HGNC Symbol;Acc:HGNC:32326] | 2.3927 | 6.67E-10 |
| ENSG00000198855 | FICD | FIC domain protein adenylyltransferase [Source:HGNC Symbol;Acc:HGNC:18416] | 0.7937 | 6.70E-10 |
| ENSG00000128342 | LIF | LIF interleukin 6 family cytokine [Source:HGNC Symbol;Acc:HGNC:6596] | 2.1878 | 6.78E-10 |
| ENSG00000144730 | IL17RD | interleukin 17 receptor D [Source:HGNC Symbol;Acc:HGNC:17616] | -1.3978 | 7.18E-10 |
| ENSG00000182103 | FAM181B | family with sequence similarity 181 member B [Source:HGNC Symbol;Acc:HGNC:28512] | -1.7412 | 7.40E-10 |
| ENSG00000130449 | ZSWIM6 | zinc finger SWIM-type containing 6 [Source:HGNC Symbol;Acc:HGNC:29316] | 0.7340 | 7.45E-10 |
| ENSG00000128656 | CHN1 | chimerin 1 [Source:HGNC Symbol;Acc:HGNC:1943] | -0.8006 | 7.49E-10 |
| ENSG00000164684 | ZNF704 | zinc finger protein 704 [Source:HGNC Symbol;Acc:HGNC:32291] | -0.9688 | 7.79E-10 |
| ENSG00000100503 | NIN | ninein [Source:HGNC Symbol;Acc:HGNC:14906] | 0.5308 | 7.81E-10 |
| ENSG00000107614 | TRDMT1 | tRNA aspartic acid methyltransferase 1 [Source:HGNC Symbol;Acc:HGNC:2977] | -1.0503 | 8.05E-10 |

| | | | | |
|---|---|---|---|---|
| ENSG00000105186 | ANKRD27 | ankyrin repeat domain 27 [Source:HGNC Symbol;Acc:HGNC:25310] | -0.7011 | 8.15E-10 |
| ENSG00000039560 | RAI14 | retinoic acid induced 14 [Source:HGNC Symbol;Acc:HGNC:14873] | 0.7652 | 8.31E-10 |
| ENSG00000182095 | TNRC18 | trinucleotide repeat containing 18 [Source:HGNC Symbol;Acc:HGNC:11962] | -1.1755 | 8.38E-10 |
| ENSG00000104714 | ERICH1 | glutamate rich 1 [Source:HGNC Symbol;Acc:HGNC:27234] | 0.6758 | 8.44E-10 |
| ENSG00000089063 | TMEM230 | transmembrane protein 230 [Source:HGNC Symbol;Acc:HGNC:15876] | -0.7090 | 8.80E-10 |
| ENSG00000162076 | FLYWCH2 | FLYWCH family member 2 [Source:HGNC Symbol;Acc:HGNC:25178] | -0.7678 | 8.80E-10 |
| ENSG00000198728 | LDB1 | LIM domain binding 1 [Source:HGNC Symbol;Acc:HGNC:6532] | -0.7844 | 8.88E-10 |
| ENSG00000141506 | PIK3R5 | phosphoinositide-3-kinase regulatory subunit 5 [Source:HGNC Symbol;Acc:HGNC:30035] | -3.6940 | 8.95E-10 |
| ENSG00000133789 | SWAP70 | switching B cell complex subunit SWAP70 [Source:HGNC Symbol;Acc:HGNC:17070] | -0.5903 | 9.06E-10 |
| ENSG00000213347 | MXD3 | MAX dimerization protein 3 [Source:HGNC Symbol;Acc:HGNC:14008] | -1.2662 | 9.19E-10 |
| ENSG00000197457 | STMN3 | stathmin 3 [Source:HGNC Symbol;Acc:HGNC:15926] | -0.5823 | 9.22E-10 |
| ENSG00000117597 | UTP25 | UTP25 small subunit processor component [Source:HGNC Symbol;Acc:HGNC:28440] | 0.5741 | 9.38E-10 |
| ENSG00000047230 | CTPS2 | CTP synthase 2 [Source:HGNC Symbol;Acc:HGNC:2520] | -1.0325 | 9.41E-10 |
| ENSG00000163888 | CAMK2N2 | calcium/calmodulin dependent protein kinase II inhibitor 2 [Source:HGNC Symbol;Acc:HGNC:24197] | 0.7027 | 9.53E-10 |
| ENSG00000138395 | CDK15 | cyclin dependent kinase 15 [Source:HGNC Symbol;Acc:HGNC:14434] | -1.1989 | 9.59E-10 |
| ENSG00000169683 | LRRC45 | leucine rich repeat containing 45 [Source:HGNC Symbol;Acc:HGNC:28302] | -1.3303 | 9.63E-10 |
| ENSG00000271474 | NA | novel transcript, antisense to UNC5C | -1.3762 | 9.66E-10 |
| ENSG00000286282 | NA | novel transcript, antisense to NOV | -1.6071 | 9.68E-10 |
| ENSG00000074047 | GLI2 | GLI family zinc finger 2 [Source:HGNC Symbol;Acc:HGNC:4318] | -0.9463 | 9.71E-10 |
| ENSG00000108370 | RGS9 | regulator of G protein signaling 9 [Source:HGNC Symbol;Acc:HGNC:10004] | -1.2446 | 9.81E-10 |
| ENSG00000136842 | TMOD1 | tropomodulin 1 [Source:HGNC Symbol;Acc:HGNC:11871] | 1.2600 | 9.89E-10 |
| ENSG00000153094 | BCL2L11 | BCL2 like 11 [Source:HGNC Symbol;Acc:HGNC:994] | -1.1140 | 9.92E-10 |
| ENSG00000149646 | CNBD2 | cyclic nucleotide binding domain containing 2 [Source:HGNC Symbol;Acc:HGNC:16145] | -0.9057 | 9.92E-10 |
| ENSG00000149257 | SERPINH1 | serpin family H member 1 [Source:HGNC Symbol;Acc:HGNC:1546] | 0.5871 | 9.92E-10 |
| ENSG00000117595 | IRF6 | interferon regulatory factor 6 [Source:HGNC Symbol;Acc:HGNC:6121] | -1.1974 | 9.95E-10 |
| ENSG00000145861 | C1QTNF2 | C1q and TNF related 2 [Source:HGNC Symbol;Acc:HGNC:14325] | -1.4155 | 1.02E-09 |
| ENSG00000138646 | HERC5 | HECT and RLD domain containing E3 ubiquitin protein ligase 5 [Source:HGNC Symbol;Acc:HGNC:24368] | -0.7919 | 1.03E-09 |
| ENSG00000173269 | MMRN2 | multimerin 2 [Source:HGNC Symbol;Acc:HGNC:19888] | -1.8749 | 1.04E-09 |
| ENSG00000250132 | NA | novel transcript | -1.3244 | 1.05E-09 |
| ENSG00000277778 | PGM5P2 | phosphoglucomutase 5 pseudogene 2 [Source:HGNC Symbol;Acc:HGNC:18965] | -1.3591 | 1.08E-09 |
| ENSG00000169891 | REPS2 | RALBP1 associated Eps domain containing 2 [Source:HGNC Symbol;Acc:HGNC:9963] | -1.1020 | 1.09E-09 |
| ENSG00000160305 | DIP2A | disco interacting protein 2 homolog A [Source:HGNC Symbol;Acc:HGNC:17217] | 0.7807 | 1.11E-09 |
| ENSG00000143554 | SLC27A3 | solute carrier family 27 member 3 [Source:HGNC Symbol;Acc:HGNC:10997] | 1.5900 | 1.12E-09 |
| ENSG00000133678 | TMEM254 | transmembrane protein 254 [Source:HGNC Symbol;Acc:HGNC:25804] | -0.7523 | 1.13E-09 |
| ENSG00000144747 | TMF1 | TATA element modulatory factor 1 [Source:HGNC Symbol;Acc:HGNC:11870] | 0.6579 | 1.14E-09 |
| ENSG00000004866 | ST7 | suppression of tumorigenicity 7 [Source:HGNC Symbol;Acc:HGNC:11351] | 0.6307 | 1.15E-09 |
| ENSG00000175538 | KCNE3 | potassium voltage-gated channel subfamily E regulatory subunit 3 [Source:HGNC Symbol;Acc:HGNC:6243] | -2.5544 | 1.17E-09 |
| ENSG00000189120 | SP6 | Sp6 transcription factor [Source:HGNC Symbol;Acc:HGNC:14530] | -1.3851 | 1.18E-09 |
| ENSG00000054654 | SYNE2 | spectrin repeat containing nuclear envelope protein 2 [Source:HGNC Symbol;Acc:HGNC:17084] | -0.6852 | 1.18E-09 |
| ENSG00000141401 | IMPA2 | inositol monophosphatase 2 [Source:HGNC Symbol;Acc:HGNC:6051] | -1.5348 | 1.19E-09 |
| ENSG00000110400 | NECTIN1 | nectin cell adhesion molecule 1 [Source:HGNC Symbol;Acc:HGNC:9706] | 2.2767 | 1.20E-09 |
| ENSG00000174028 | FAM3C2 | family with sequence similarity 3 member C2 (pseudogene) [Source:HGNC Symbol;Acc:HGNC:34501] | 0.8609 | 1.23E-09 |

| | | | | |
|---|---|---|---|---|
| ENSG00000256043 | CTSO | cathepsin O [Source:HGNC Symbol;Acc:HGNC:2542] | -1.0120 | 1.23E-09 |
| ENSG00000153310 | CYRIB | CYFIP related Rac1 interactor B [Source:HGNC Symbol;Acc:HGNC:25216] | 0.5235 | 1.27E-09 |
| ENSG00000158555 | GDPD5 | glycerophosphodiester phosphodiesterase domain containing 5 [Source:HGNC Symbol;Acc:HGNC:28804] | -1.1055 | 1.27E-09 |
| ENSG00000178919 | FOXE1 | forkhead box E1 [Source:HGNC Symbol;Acc:HGNC:3806] | -1.0133 | 1.29E-09 |
| ENSG00000162600 | OMA1 | OMA1 zinc metallopeptidase [Source:HGNC Symbol;Acc:HGNC:29661] | -1.3088 | 1.29E-09 |
| ENSG00000103064 | SLC7A6 | solute carrier family 7 member 6 [Source:HGNC Symbol;Acc:HGNC:11064] | 0.8809 | 1.30E-09 |
| ENSG00000005469 | CROT | carnitine O-octanoyltransferase [Source:HGNC Symbol;Acc:HGNC:2366] | -0.8857 | 1.31E-09 |
| ENSG00000162923 | WDR26 | WD repeat domain 26 [Source:HGNC Symbol;Acc:HGNC:21208] | 0.5266 | 1.32E-09 |
| ENSG00000136522 | MRPL47 | mitochondrial ribosomal protein L47 [Source:HGNC Symbol;Acc:HGNC:16652] | 0.5196 | 1.33E-09 |
| ENSG00000165775 | FUNDC2 | FUN14 domain containing 2 [Source:HGNC Symbol;Acc:HGNC:24925] | 0.5795 | 1.36E-09 |
| ENSG00000287769 | NA | novel transcript, antisense to SPIN1 | -1.0849 | 1.42E-09 |
| ENSG00000113391 | FAM172A | family with sequence similarity 172 member A [Source:HGNC Symbol;Acc:HGNC:25365] | -0.5890 | 1.43E-09 |
| ENSG00000117399 | CDC20 | cell division cycle 20 [Source:HGNC Symbol;Acc:HGNC:1723] | -0.6264 | 1.49E-09 |
| ENSG00000143891 | GALM | galactose mutarotase [Source:HGNC Symbol;Acc:HGNC:24063] | 0.6892 | 1.53E-09 |
| ENSG00000175029 | CTBP2 | C-terminal binding protein 2 [Source:HGNC Symbol;Acc:HGNC:2495] | 0.7593 | 1.53E-09 |
| ENSG00000141576 | RNF157 | ring finger protein 157 [Source:HGNC Symbol;Acc:HGNC:29402] | -0.9993 | 1.55E-09 |
| ENSG00000163704 | PRRT3 | proline rich transmembrane protein 3 [Source:HGNC Symbol;Acc:HGNC:26591] | -0.9970 | 1.59E-09 |
| ENSG00000119777 | TMEM214 | transmembrane protein 214 [Source:HGNC Symbol;Acc:HGNC:25983] | 0.5025 | 1.68E-09 |
| ENSG00000185561 | TLCD2 | TLC domain containing 2 [Source:HGNC Symbol;Acc:HGNC:33522] | 1.0931 | 1.71E-09 |
| ENSG00000029993 | HMGB3 | high mobility group box 3 [Source:HGNC Symbol;Acc:HGNC:5004] | -0.5784 | 1.78E-09 |
| ENSG00000166140 | ZFYVE19 | zinc finger FYVE-type containing 19 [Source:HGNC Symbol;Acc:HGNC:20758] | -0.6986 | 1.82E-09 |
| ENSG00000176842 | IRX5 | iroquois homeobox 5 [Source:HGNC Symbol;Acc:HGNC:14361] | -0.7568 | 1.84E-09 |
| ENSG00000133943 | DGLUCY | D-glutamate cyclase [Source:HGNC Symbol;Acc:HGNC:20498] | -0.6135 | 1.85E-09 |
| ENSG00000108306 | FBXL20 | F-box and leucine rich repeat protein 20 [Source:HGNC Symbol;Acc:HGNC:24679] | -1.1618 | 1.87E-09 |
| ENSG00000174010 | KLHL15 | kelch like family member 15 [Source:HGNC Symbol;Acc:HGNC:29347] | -0.6810 | 1.89E-09 |
| ENSG00000144136 | SLC20A1 | solute carrier family 20 member 1 [Source:HGNC Symbol;Acc:HGNC:10946] | 0.9926 | 1.94E-09 |
| ENSG00000172348 | RCAN2 | regulator of calcineurin 2 [Source:HGNC Symbol;Acc:HGNC:3041] | -1.2697 | 1.97E-09 |
| ENSG00000196456 | ZNF775 | zinc finger protein 775 [Source:HGNC Symbol;Acc:HGNC:28501] | -0.9149 | 1.99E-09 |
| ENSG00000156453 | PCDH1 | protocadherin 1 [Source:HGNC Symbol;Acc:HGNC:8655] | -1.1623 | 2.08E-09 |
| ENSG00000184007 | PTP4A2 | protein tyrosine phosphatase 4A2 [Source:HGNC Symbol;Acc:HGNC:9635] | 0.5536 | 2.12E-09 |
| ENSG00000140463 | BBS4 | Bardet-Biedl syndrome 4 [Source:HGNC Symbol;Acc:HGNC:969] | -0.5648 | 2.12E-09 |
| ENSG00000188739 | RBM34 | RNA binding motif protein 34 [Source:HGNC Symbol;Acc:HGNC:28965] | 0.8140 | 2.12E-09 |
| ENSG00000154309 | DISP1 | dispatched RND transporter family member 1 [Source:HGNC Symbol;Acc:HGNC:19711] | 1.2117 | 2.14E-09 |
| ENSG00000278934 | NA | TEC | -1.5673 | 2.15E-09 |
| ENSG00000172992 | DCAKD | dephospho-CoA kinase domain containing [Source:HGNC Symbol;Acc:HGNC:26238] | -0.6026 | 2.26E-09 |
| ENSG00000175426 | PCSK1 | proprotein convertase subtilisin/kexin type 1 [Source:HGNC Symbol;Acc:HGNC:8743] | 2.2903 | 2.29E-09 |
| ENSG00000127334 | DYRK2 | dual specificity tyrosine phosphorylation regulated kinase 2 [Source:HGNC Symbol;Acc:HGNC:3093] | 0.9991 | 2.33E-09 |
| ENSG00000135821 | GLUL | glutamate-ammonia ligase [Source:HGNC Symbol;Acc:HGNC:4341] | -0.8347 | 2.38E-09 |
| ENSG00000164023 | SGMS2 | sphingomyelin synthase 2 [Source:HGNC Symbol;Acc:HGNC:28395] | 1.5510 | 2.46E-09 |
| ENSG00000120068 | HOXB8 | homeobox B8 [Source:HGNC Symbol;Acc:HGNC:5119] | 1.0559 | 2.53E-09 |
| ENSG00000172071 | EIF2AK3 | eukaryotic translation initiation factor 2 alpha kinase 3 [Source:HGNC Symbol;Acc:HGNC:3255] | 0.8679 | 2.53E-09 |
| ENSG00000204560 | DHX16 | DEAH-box helicase 16 [Source:HGNC Symbol;Acc:HGNC:2739] | 0.5249 | 2.54E-09 |
| ENSG00000131652 | THOC6 | THO complex 6 [Source:HGNC Symbol;Acc:HGNC:28369] | -0.6569 | 2.59E-09 |

| | | | | |
|---|---|---|---|---|
| ENSG00000124479 | NDP | norrin cystine knot growth factor NDP [Source:HGNC Symbol;Acc:HGNC:7678] | -1.7979 | 2.71E-09 |
| ENSG00000167434 | CA4 | carbonic anhydrase 4 [Source:HGNC Symbol;Acc:HGNC:1375] | -1.3550 | 2.72E-09 |
| ENSG00000124225 | PMEPA1 | prostate transmembrane protein, androgen induced 1 [Source:HGNC Symbol;Acc:HGNC:14107] | 6.4905 | 2.81E-09 |
| ENSG00000165424 | ZCCHC24 | zinc finger CCHC-type containing 24 [Source:HGNC Symbol;Acc:HGNC:26911] | -0.9109 | 2.86E-09 |
| ENSG00000183166 | CALN1 | calneuron 1 [Source:HGNC Symbol;Acc:HGNC:13248] | -0.9015 | 2.92E-09 |
| ENSG00000272462 | NA | novel transcript | -0.7490 | 2.93E-09 |
| ENSG00000160216 | AGPAT3 | 1-acylglycerol-3-phosphate O-acyltransferase 3 [Source:HGNC Symbol;Acc:HGNC:326] | 0.5170 | 2.95E-09 |
| ENSG00000100271 | TTLL1 | tubulin tyrosine ligase like 1 [Source:HGNC Symbol;Acc:HGNC:1312] | -2.4727 | 2.99E-09 |
| ENSG00000264522 | OTUD7B | OTU deubiquitinase 7B [Source:HGNC Symbol;Acc:HGNC:16683] | 0.8184 | 3.06E-09 |
| ENSG00000100285 | NEFH | neurofilament heavy [Source:HGNC Symbol;Acc:HGNC:7737] | -1.8160 | 3.10E-09 |
| ENSG00000169299 | PGM2 | phosphoglucomutase 2 [Source:HGNC Symbol;Acc:HGNC:8906] | 1.0589 | 3.10E-09 |
| ENSG00000229953 | NA | novel transcript | 1.6960 | 3.10E-09 |
| ENSG00000114107 | CEP70 | centrosomal protein 70 [Source:HGNC Symbol;Acc:HGNC:29972] | 0.6635 | 3.14E-09 |
| ENSG00000280399 | NA | TEC | 1.5792 | 3.16E-09 |
| ENSG00000175221 | MED16 | mediator complex subunit 16 [Source:HGNC Symbol;Acc:HGNC:17556] | -0.6555 | 3.27E-09 |
| ENSG00000197444 | OGDHL | oxoglutarate dehydrogenase like [Source:HGNC Symbol;Acc:HGNC:25590] | -1.3243 | 3.32E-09 |
| ENSG00000122512 | PMS2 | PMS1 homolog 2, mismatch repair system component [Source:HGNC Symbol;Acc:HGNC:9122] | -0.7599 | 3.35E-09 |
| ENSG00000277586 | NEFL | neurofilament light [Source:HGNC Symbol;Acc:HGNC:7739] | -0.9601 | 3.39E-09 |
| ENSG00000106070 | GRB10 | growth factor receptor bound protein 10 [Source:HGNC Symbol;Acc:HGNC:4564] | 1.0428 | 3.39E-09 |
| ENSG00000198018 | ENTPD7 | ectonucleoside triphosphate diphosphohydrolase 7 [Source:HGNC Symbol;Acc:HGNC:19745] | 0.9087 | 3.39E-09 |
| ENSG00000067798 | NAV3 | neuron navigator 3 [Source:HGNC Symbol;Acc:HGNC:15998] | -0.6902 | 3.41E-09 |
| ENSG00000055483 | USP36 | ubiquitin specific peptidase 36 [Source:HGNC Symbol;Acc:HGNC:20062] | 0.5093 | 3.42E-09 |
| ENSG00000072840 | EVC | EvC ciliary complex subunit 1 [Source:HGNC Symbol;Acc:HGNC:3497] | -0.8605 | 3.42E-09 |
| ENSG00000162692 | VCAM1 | vascular cell adhesion molecule 1 [Source:HGNC Symbol;Acc:HGNC:12663] | 1.0447 | 3.42E-09 |
| ENSG00000185742 | C11orf87 | chromosome 11 open reading frame 87 [Source:HGNC Symbol;Acc:HGNC:33788] | 2.1544 | 3.43E-09 |
| ENSG00000157601 | MX1 | MX dynamin like GTPase 1 [Source:HGNC Symbol;Acc:HGNC:7532] | -1.3567 | 3.53E-09 |
| ENSG00000204025 | TRPC5OS | TRPC5 opposite strand [Source:HGNC Symbol;Acc:HGNC:40593] | 2.1958 | 3.63E-09 |
| ENSG00000205930 | C21orf62-AS1 | C21orf62 antisense RNA 1 [Source:HGNC Symbol;Acc:HGNC:1290] | -1.7357 | 3.69E-09 |
| ENSG00000107863 | ARHGAP21 | Rho GTPase activating protein 21 [Source:HGNC Symbol;Acc:HGNC:23725] | 0.7016 | 3.71E-09 |
| ENSG00000167553 | TUBA1C | tubulin alpha 1c [Source:HGNC Symbol;Acc:HGNC:20768] | 0.7312 | 3.74E-09 |
| ENSG00000145016 | RUBCN | rubicon autophagy regulator [Source:HGNC Symbol;Acc:HGNC:28991] | 0.7640 | 3.78E-09 |
| ENSG00000131015 | ULBP2 | UL16 binding protein 2 [Source:HGNC Symbol;Acc:HGNC:14894] | 1.5815 | 3.80E-09 |
| ENSG00000160325 | CACFD1 | calcium channel flower domain containing 1 [Source:HGNC Symbol;Acc:HGNC:1365] | -0.7990 | 3.82E-09 |
| ENSG00000151131 | C12orf45 | chromosome 12 open reading frame 45 [Source:HGNC Symbol;Acc:HGNC:28628] | 0.9705 | 3.83E-09 |
| ENSG00000156650 | KAT6B | lysine acetyltransferase 6B [Source:HGNC Symbol;Acc:HGNC:17582] | -0.5917 | 3.84E-09 |
| ENSG00000172458 | IL17D | interleukin 17D [Source:HGNC Symbol;Acc:HGNC:5984] | 0.8351 | 3.85E-09 |
| ENSG00000174307 | PHLDA3 | pleckstrin homology like domain family A member 3 [Source:HGNC Symbol;Acc:HGNC:8934] | 1.8546 | 3.85E-09 |
| ENSG00000174740 | PABPC5 | poly(A) binding protein cytoplasmic 5 [Source:HGNC Symbol;Acc:HGNC:13629] | -0.8750 | 3.87E-09 |
| ENSG00000149292 | TTC12 | tetratricopeptide repeat domain 12 [Source:HGNC Symbol;Acc:HGNC:23700] | -1.2954 | 3.93E-09 |
| ENSG00000184156 | KCNQ3 | potassium voltage-gated channel subfamily Q member 3 [Source:HGNC Symbol;Acc:HGNC:6297] | 5.0585 | 4.01E-09 |
| ENSG00000135540 | NHSL1 | NHS like 1 [Source:HGNC Symbol;Acc:HGNC:21021] | 1.5810 | 4.03E-09 |
| ENSG00000196549 | MME | membrane metalloendopeptidase [Source:HGNC Symbol;Acc:HGNC:7154] | -0.8943 | 4.18E-09 |
| ENSG00000275591 | XKR5 | XK related 5 [Source:HGNC Symbol;Acc:HGNC:20782] | -2.1488 | 4.24E-09 |

| | | | | |
|---|---|---|---|---|
| ENSG00000135441 | BLOC1S1 | biogenesis of lysosomal organelles complex 1 subunit 1 [Source:HGNC Symbol;Acc:HGNC:4200] | -0.6241 | 4.31E-09 |
| ENSG00000179364 | PACS2 | phosphofurin acidic cluster sorting protein 2 [Source:HGNC Symbol;Acc:HGNC:23794] | -0.7070 | 4.40E-09 |
| ENSG00000130706 | ADRM1 | adhesion regulating molecule 1 [Source:HGNC Symbol;Acc:HGNC:15759] | 0.5180 | 4.43E-09 |
| ENSG00000168061 | SAC3D1 | SAC3 domain containing 1 [Source:HGNC Symbol;Acc:HGNC:30179] | -0.9637 | 4.49E-09 |
| ENSG00000162873 | KLHDC8A | kelch domain containing 8A [Source:HGNC Symbol;Acc:HGNC:25573] | 0.9498 | 4.58E-09 |
| ENSG00000183873 | SCN5A | sodium voltage-gated channel alpha subunit 5 [Source:HGNC Symbol;Acc:HGNC:10593] | -1.6133 | 4.67E-09 |
| ENSG00000148225 | WDR31 | WD repeat domain 31 [Source:HGNC Symbol;Acc:HGNC:21421] | -1.3805 | 4.76E-09 |
| ENSG00000138738 | PRDM5 | PR/SET domain 5 [Source:HGNC Symbol;Acc:HGNC:9349] | -0.7192 | 4.81E-09 |
| ENSG00000073331 | ALPK1 | alpha kinase 1 [Source:HGNC Symbol;Acc:HGNC:20917] | -0.5394 | 4.87E-09 |
| ENSG00000177674 | AGTRAP | angiotensin II receptor associated protein [Source:HGNC Symbol;Acc:HGNC:13539] | -0.5006 | 4.89E-09 |
| ENSG00000175182 | FAM131A | family with sequence similarity 131 member A [Source:HGNC Symbol;Acc:HGNC:28308] | 0.8456 | 5.01E-09 |
| ENSG00000105419 | MEIS3 | Meis homeobox 3 [Source:HGNC Symbol;Acc:HGNC:29537] | 0.7805 | 5.02E-09 |
| ENSG00000139641 | ESYT1 | extended synaptotagmin 1 [Source:HGNC Symbol;Acc:HGNC:29534] | 0.5624 | 5.15E-09 |
| ENSG00000104154 | SLC30A4 | solute carrier family 30 member 4 [Source:HGNC Symbol;Acc:HGNC:11015] | -0.5501 | 5.24E-09 |
| ENSG00000157734 | SNX22 | sorting nexin 22 [Source:HGNC Symbol;Acc:HGNC:16315] | 0.8332 | 5.27E-09 |
| ENSG00000167549 | CORO6 | coronin 6 [Source:HGNC Symbol;Acc:HGNC:21356] | -1.5476 | 5.30E-09 |
| ENSG00000139428 | MMAB | metabolism of cobalamin associated B [Source:HGNC Symbol;Acc:HGNC:19331] | -0.8540 | 5.31E-09 |
| ENSG00000050405 | LIMA1 | LIM domain and actin binding 1 [Source:HGNC Symbol;Acc:HGNC:24636] | 0.9368 | 5.33E-09 |
| ENSG00000161082 | CELF5 | CUGBP Elav-like family member 5 [Source:HGNC Symbol;Acc:HGNC:14058] | -1.0342 | 5.33E-09 |
| ENSG00000184117 | NIPSNAP1 | nipsnap homolog 1 [Source:HGNC Symbol;Acc:HGNC:7827] | -0.6494 | 5.35E-09 |
| ENSG00000081026 | MAGI3 | membrane associated guanylate kinase, WW and PDZ domain containing 3 [Source:HGNC Symbol;Acc:HGNC:29647] | -0.7415 | 5.43E-09 |
| ENSG00000030110 | BAK1 | BCL2 antagonist/killer 1 [Source:HGNC Symbol;Acc:HGNC:949] | 0.6052 | 5.48E-09 |
| ENSG00000175600 | SUGCT | succinyl-CoA:glutarate-CoA transferase [Source:HGNC Symbol;Acc:HGNC:16001] | 2.5635 | 5.52E-09 |
| ENSG00000132640 | BTBD3 | BTB domain containing 3 [Source:HGNC Symbol;Acc:HGNC:15854] | 0.6223 | 5.52E-09 |
| ENSG00000136758 | YME1L1 | YME1 like 1 ATPase [Source:HGNC Symbol;Acc:HGNC:12843] | 0.5444 | 5.58E-09 |
| ENSG00000125844 | RRBP1 | ribosome binding protein 1 [Source:HGNC Symbol;Acc:HGNC:10448] | 0.7415 | 5.60E-09 |
| ENSG00000119711 | ALDH6A1 | aldehyde dehydrogenase 6 family member A1 [Source:HGNC Symbol;Acc:HGNC:7179] | -0.6545 | 5.67E-09 |
| ENSG00000165572 | KBTBD6 | kelch repeat and BTB domain containing 6 [Source:HGNC Symbol;Acc:HGNC:25340] | -0.6951 | 5.69E-09 |
| ENSG00000196660 | SLC30A10 | solute carrier family 30 member 10 [Source:HGNC Symbol;Acc:HGNC:25355] | -0.7220 | 5.78E-09 |
| ENSG00000138623 | SEMA7A | semaphorin 7A (John Milton Hagen blood group) [Source:HGNC Symbol;Acc:HGNC:10741] | 0.6143 | 5.78E-09 |
| ENSG00000205809 | KLRC2 | killer cell lectin like receptor C2 [Source:HGNC Symbol;Acc:HGNC:6375] | 4.6479 | 5.85E-09 |
| ENSG00000125740 | FOSB | FosB proto-oncogene, AP-1 transcription factor subunit [Source:HGNC Symbol;Acc:HGNC:3797] | 3.2228 | 5.87E-09 |
| ENSG00000180530 | NRIP1 | nuclear receptor interacting protein 1 [Source:HGNC Symbol;Acc:HGNC:8001] | 0.5625 | 5.91E-09 |
| ENSG00000280254 | NA | TEC | -1.2628 | 5.96E-09 |
| ENSG00000234465 | PINLYP | phospholipase A2 inhibitor and LY6/PLAUR domain containing [Source:HGNC Symbol;Acc:HGNC:44206] | 2.2844 | 6.03E-09 |
| ENSG00000170899 | GSTA4 | glutathione S-transferase alpha 4 [Source:HGNC Symbol;Acc:HGNC:4629] | -0.5749 | 6.05E-09 |
| ENSG00000137965 | IFI44 | interferon induced protein 44 [Source:HGNC Symbol;Acc:HGNC:16938] | -0.7205 | 6.20E-09 |
| ENSG00000177380 | PPFIA3 | PTPRF interacting protein alpha 3 [Source:HGNC Symbol;Acc:HGNC:9247] | -0.7241 | 6.20E-09 |
| ENSG00000184838 | PRR16 | proline rich 16 [Source:HGNC Symbol;Acc:HGNC:29654] | -0.9030 | 6.41E-09 |
| ENSG00000120805 | 1,00 ARL | ADP ribosylation factor like GTPase 1 [Source:HGNC Symbol;Acc:HGNC:692] | 0.6786 | 6.42E-09 |
| ENSG00000173917 | HOXB2 | homeobox B2 [Source:HGNC Symbol;Acc:HGNC:5113] | 1.4074 | 6.45E-09 |
| ENSG00000198964 | SGMS1 | sphingomyelin synthase 1 [Source:HGNC Symbol;Acc:HGNC:29799] | 0.9283 | 6.56E-09 |

| | | | | |
|---|---|---|---|---|
| ENSG00000167680 | SEMA6B | semaphorin 6B [Source:HGNC Symbol;Acc:HGNC:10739] | -1.0194 | 6.71E-09 |
| ENSG00000156381 | ANKRD9 | ankyrin repeat domain 9 [Source:HGNC Symbol;Acc:HGNC:20096] | -0.9577 | 6.72E-09 |
| ENSG00000145431 | PDGFC | platelet derived growth factor C [Source:HGNC Symbol;Acc:HGNC:8801] | -0.6972 | 6.88E-09 |
| ENSG00000167470 | MIDN | midnolin [Source:HGNC Symbol;Acc:HGNC:16298] | 0.8196 | 6.93E-09 |
| ENSG00000084112 | SSH1 | slingshot protein phosphatase 1 [Source:HGNC Symbol;Acc:HGNC:30579] | 0.6394 | 7.07E-09 |
| ENSG00000185483 | ROR1 | receptor tyrosine kinase like orphan receptor 1 [Source:HGNC Symbol;Acc:HGNC:10256] | -1.5017 | 7.10E-09 |
| ENSG00000136169 | SETDB2 | SET domain bifurcated histone lysine methyltransferase 2 [Source:HGNC Symbol;Acc:HGNC:20263] | -0.7685 | 7.15E-09 |
| ENSG00000171861 | MRM3 | mitochondrial rRNA methyltransferase 3 [Source:HGNC Symbol;Acc:HGNC:18485] | 0.7575 | 7.20E-09 |
| ENSG00000182575 | NXPH3 | neurexophilin 3 [Source:HGNC Symbol;Acc:HGNC:8077] | -1.2583 | 7.22E-09 |
| ENSG00000101901 | ALG13 | ALG13 UDP-N-acetylglucosaminyltransferase subunit [Source:HGNC Symbol;Acc:HGNC:30881] | 0.6246 | 7.27E-09 |
| ENSG00000130224 | LRCH2 | leucine rich repeats and calponin homology domain containing 2 [Source:HGNC Symbol;Acc:HGNC:29292] | -0.5667 | 7.38E-09 |
| ENSG00000144366 | GULP1 | GULP PTB domain containing engulfment adaptor 1 [Source:HGNC Symbol;Acc:HGNC:18649] | 0.6686 | 7.40E-09 |
| ENSG00000112763 | BTN2A1 | butyrophilin subfamily 2 member A1 [Source:HGNC Symbol;Acc:HGNC:1136] | 0.6956 | 7.49E-09 |
| ENSG00000174943 | KCTD13 | potassium channel tetramerization domain containing 13 [Source:HGNC Symbol;Acc:HGNC:22234] | 0.8761 | 7.53E-09 |
| ENSG00000205571 | SMN2 | survival of motor neuron 2, centromeric [Source:HGNC Symbol;Acc:HGNC:11118] | 0.5019 | 7.54E-09 |
| ENSG00000149599 | DUSP15 | dual specificity phosphatase 15 [Source:HGNC Symbol;Acc:HGNC:16236] | -1.1366 | 7.57E-09 |
| ENSG00000126368 | NR1D1 | nuclear receptor subfamily 1 group D member 1 [Source:HGNC Symbol;Acc:HGNC:7962] | -0.7456 | 7.59E-09 |
| ENSG00000103005 | USB1 | U6 snRNA biogenesis phosphodiesterase 1 [Source:HGNC Symbol;Acc:HGNC:25792] | 0.5407 | 7.65E-09 |
| ENSG00000237836 | PHKA2-AS1 | PHKA2 antisense RNA 1 [Source:HGNC Symbol;Acc:HGNC:44110] | 1.1412 | 7.69E-09 |
| ENSG00000136999 | CCN3 | cellular communication network factor 3 [Source:HGNC Symbol;Acc:HGNC:7885] | -0.8316 | 7.79E-09 |
| ENSG00000232434 | AJM1 | apical junction component 1 homolog [Source:HGNC Symbol;Acc:HGNC:37284] | 0.5899 | 8.06E-09 |
| ENSG00000167881 | SRP68 | signal recognition particle 68 [Source:HGNC Symbol;Acc:HGNC:11302] | 0.5295 | 8.17E-09 |
| ENSG00000188559 | RALGAPA2 | Ral GTPase activating protein catalytic subunit alpha 2 [Source:HGNC Symbol;Acc:HGNC:16207] | 0.6607 | 8.27E-09 |
| ENSG00000171161 | ZNF672 | zinc finger protein 672 [Source:HGNC Symbol;Acc:HGNC:26179] | 0.6154 | 8.39E-09 |
| ENSG00000142494 | SLC47A1 | solute carrier family 47 member 1 [Source:HGNC Symbol;Acc:HGNC:25588] | 0.8327 | 8.53E-09 |
| ENSG00000159479 | MED8 | mediator complex subunit 8 [Source:HGNC Symbol;Acc:HGNC:19971] | 0.5082 | 8.70E-09 |
| ENSG00000059758 | CDK17 | cyclin dependent kinase 17 [Source:HGNC Symbol;Acc:HGNC:8750] | 1.0248 | 8.88E-09 |
| ENSG00000128872 | TMOD2 | tropomodulin 2 [Source:HGNC Symbol;Acc:HGNC:11872] | -1.0282 | 8.95E-09 |
| ENSG00000111371 | SLC38A1 | solute carrier family 38 member 1 [Source:HGNC Symbol;Acc:HGNC:13447] | 0.7128 | 9.07E-09 |
| ENSG00000169398 | PTK2 | protein tyrosine kinase 2 [Source:HGNC Symbol;Acc:HGNC:9611] | 0.5658 | 9.17E-09 |
| ENSG00000109084 | TMEM97 | transmembrane protein 97 [Source:HGNC Symbol;Acc:HGNC:28106] | -0.5300 | 9.17E-09 |
| ENSG00000055732 | MCOLN3 | mucolipin 3 [Source:HGNC Symbol;Acc:HGNC:13358] | 0.7561 | 9.39E-09 |
| ENSG00000257108 | NHLRC4 | NHL repeat containing 4 [Source:HGNC Symbol;Acc:HGNC:26700] | -1.8049 | 9.79E-09 |
| ENSG00000232110 | NA | novel transcript, antisense to LIPA | 2.5996 | 9.84E-09 |
| ENSG00000137804 | NUSAP1 | nucleolar and spindle associated protein 1 [Source:HGNC Symbol;Acc:HGNC:18538] | -0.7737 | 9.88E-09 |
| ENSG00000115963 | RND3 | Rho family GTPase 3 [Source:HGNC Symbol;Acc:HGNC:671] | 2.1207 | 9.89E-09 |
| ENSG00000204851 | PNMA8B | PNMA family member 8B [Source:HGNC Symbol;Acc:HGNC:29206] | -0.8998 | 9.89E-09 |
| ENSG00000114999 | TTL | tubulin tyrosine ligase [Source:HGNC Symbol;Acc:HGNC:21586] | 0.6756 | 1.04E-08 |
| ENSG00000138613 | APH1B | aph-1 homolog B, gamma-secretase subunit [Source:HGNC Symbol;Acc:HGNC:24080] | -0.6456 | 1.05E-08 |
| ENSG00000158220 | ESYT3 | extended synaptotagmin 3 [Source:HGNC Symbol;Acc:HGNC:24295] | 1.6922 | 1.07E-08 |
| ENSG00000147996 | CBWD5 | COBW domain containing 5 [Source:HGNC Symbol;Acc:HGNC:24584] | 0.5718 | 1.09E-08 |
| ENSG00000186765 | FSCN2 | fascin actin-bundling protein 2, retinal [Source:HGNC Symbol;Acc:HGNC:3960] | 1.5757 | 1.11E-08 |

| | | | | |
|---|---|---|---|---|
| ENSG00000100319 | ZMAT5 | zinc finger matrin-type 5 [Source:HGNC Symbol;Acc:HGNC:28046] | -0.7078 | 1.12E-08 |
| ENSG00000231625 | SLC47A1P2 | SLC47A1 pseudogene 2 [Source:HGNC Symbol;Acc:HGNC:53866] | 1.9727 | 1.14E-08 |
| ENSG00000076826 | CAMSAP3 | calmodulin regulated spectrin associated protein family member 3 [Source:HGNC Symbol;Acc:HGNC:29307] | -0.9496 | 1.16E-08 |
| ENSG00000135766 | EGLN1 | egl-9 family hypoxia inducible factor 1 [Source:HGNC Symbol;Acc:HGNC:1232] | 0.7644 | 1.16E-08 |
| ENSG00000258315 | C17orf49 | chromosome 17 open reading frame 49 [Source:HGNC Symbol;Acc:HGNC:28737] | -0.5781 | 1.17E-08 |
| ENSG00000182463 | TSHZ2 | teashirt zinc finger homeobox 2 [Source:HGNC Symbol;Acc:HGNC:13010] | 1.2841 | 1.20E-08 |
| ENSG00000101349 | PAK5 | p21 (RAC1) activated kinase 5 [Source:HGNC Symbol;Acc:HGNC:15916] | -1.5810 | 1.27E-08 |
| ENSG00000141076 | UTP4 | UTP4 small subunit processome component [Source:HGNC Symbol;Acc:HGNC:1983] | 0.5058 | 1.27E-08 |
| ENSG00000144895 | EIF2A | eukaryotic translation initiation factor 2A [Source:HGNC Symbol;Acc:HGNC:3254] | 0.5004 | 1.29E-08 |
| ENSG00000183150 | GPR19 | G protein-coupled receptor 19 [Source:HGNC Symbol;Acc:HGNC:4473] | 0.6460 | 1.31E-08 |
| ENSG00000139926 | FRMD6 | FERM domain containing 6 [Source:HGNC Symbol;Acc:HGNC:19839] | -0.7504 | 1.32E-08 |
| ENSG00000102893 | PHKB | phosphorylase kinase regulatory subunit beta [Source:HGNC Symbol;Acc:HGNC:8927] | -0.5835 | 1.34E-08 |
| ENSG00000229320 | KRT8P12 | keratin 8 pseudogene 12 [Source:HGNC Symbol;Acc:HGNC:28057] | 0.9564 | 1.34E-08 |
| ENSG00000235049 | LINC00940 | long intergenic non-protein coding RNA 940 [Source:HGNC Symbol;Acc:HGNC:48634] | -2.4295 | 1.34E-08 |
| ENSG00000204103 | MAFB | MAF bZIP transcription factor B [Source:HGNC Symbol;Acc:HGNC:6408] | -1.3650 | 1.35E-08 |
| ENSG00000069974 | RAB27A | RAB27A, member RAS oncogene family [Source:HGNC Symbol;Acc:HGNC:9766] | 0.8669 | 1.36E-08 |
| ENSG00000260329 | NA | novel transcript, antisense to C12orf23 | -1.4224 | 1.38E-08 |
| ENSG00000168067 | MAP4K2 | mitogen-activated protein kinase kinase kinase kinase 2 [Source:HGNC Symbol;Acc:HGNC:6864] | -1.0718 | 1.38E-08 |
| ENSG00000138443 | ABI2 | abl interactor 2 [Source:HGNC Symbol;Acc:HGNC:24011] | 0.6604 | 1.38E-08 |
| ENSG00000184788 | SATL1 | spermidine/spermine N1-acetyl transferase like 1 [Source:HGNC Symbol;Acc:HGNC:27992] | -1.3351 | 1.39E-08 |
| ENSG00000197622 | CDC42SE1 | CDC42 small effector 1 [Source:HGNC Symbol;Acc:HGNC:17719] | 0.5389 | 1.46E-08 |
| ENSG00000122824 | NUDT10 | nudix hydrolase 10 [Source:HGNC Symbol;Acc:HGNC:17621] | -0.7453 | 1.46E-08 |
| ENSG00000176014 | TUBB6 | tubulin beta 6 class V [Source:HGNC Symbol;Acc:HGNC:20776] | 1.1548 | 1.47E-08 |
| ENSG00000129355 | CDKN2D | cyclin dependent kinase inhibitor 2D [Source:HGNC Symbol;Acc:HGNC:1790] | -0.9351 | 1.48E-08 |
| ENSG00000110906 | KCTD10 | potassium channel tetramerization domain containing 10 [Source:HGNC Symbol;Acc:HGNC:23236] | 0.5120 | 1.48E-08 |
| ENSG00000241399 | CD302 | CD302 molecule [Source:HGNC Symbol;Acc:HGNC:30843] | -0.7278 | 1.50E-08 |
| ENSG00000174136 | RGMB | repulsive guidance molecule BMP co-receptor b [Source:HGNC Symbol;Acc:HGNC:26896] | -0.8757 | 1.54E-08 |
| ENSG00000122068 | FYTTD1 | forty-two-three domain containing 1 [Source:HGNC Symbol;Acc:HGNC:25407] | 0.5836 | 1.54E-08 |
| ENSG00000247271 | ZBED5-AS1 | ZBED5 antisense RNA 1 [Source:HGNC Symbol;Acc:HGNC:48646] | -1.0048 | 1.56E-08 |
| ENSG00000165030 | NFIL3 | nuclear factor, interleukin 3 regulated [Source:HGNC Symbol;Acc:HGNC:7787] | 1.2227 | 1.59E-08 |
| ENSG00000168874 | ATOH8 | atonal bHLH transcription factor 8 [Source:HGNC Symbol;Acc:HGNC:24126] | -4.8346 | 1.59E-08 |
| ENSG00000245571 | FAM111A-DT | FAM111A divergent transcript [Source:HGNC Symbol;Acc:HGNC:53752] | -1.4090 | 1.61E-08 |
| ENSG00000128268 | MGAT3 | beta-1,4-mannosyl-glycoprotein 4-beta-N-acetylglucosaminyltransferase [Source:HGNC Symbol;Acc:HGNC:7046] | -1.3020 | 1.62E-08 |
| ENSG00000204020 | LIPN | lipase family member N [Source:HGNC Symbol;Acc:HGNC:23452] | 3.2966 | 1.62E-08 |
| ENSG00000112290 | WASF1 | WASP family member 1 [Source:HGNC Symbol;Acc:HGNC:12732] | -0.5241 | 1.64E-08 |
| ENSG00000162738 | VANGL2 | VANGL planar cell polarity protein 2 [Source:HGNC Symbol;Acc:HGNC:15511] | -0.7034 | 1.65E-08 |
| ENSG00000117394 | SLC2A1 | solute carrier family 2 member 1 [Source:HGNC Symbol;Acc:HGNC:11005] | 0.6495 | 1.66E-08 |
| ENSG00000140015 | KCNH5 | potassium voltage-gated channel subfamily H member 5 [Source:HGNC Symbol;Acc:HGNC:6254] | 2.4284 | 1.67E-08 |
| ENSG00000198915 | RASGEF1A | RasGEF domain family member 1A [Source:HGNC Symbol;Acc:HGNC:24246] | 2.0408 | 1.68E-08 |
| ENSG00000163050 | COQ8A | coenzyme Q8A [Source:HGNC Symbol;Acc:HGNC:16812] | -0.7801 | 1.68E-08 |
| ENSG00000198162 | MAN1A2 | mannosidase alpha class 1A member 2 [Source:HGNC Symbol;Acc:HGNC:6822] | 0.8011 | 1.71E-08 |
| ENSG00000122971 | ACADS | acyl-CoA dehydrogenase short chain [Source:HGNC Symbol;Acc:HGNC:90] | -0.8140 | 1.73E-08 |
| ENSG00000079691 | CARMIL1 | capping protein regulator and myosin 1 linker 1 [Source:HGNC Symbol;Acc:HGNC:21581] | 0.8788 | 1.81E-08 |

| | | | | |
|---|---|---|---|---|
| ENSG00000184828 | ZBTB7C | zinc finger and BTB domain containing 7C [Source:HGNC Symbol;Acc:HGNC:31700] | -0.6911 | 1.83E-08 |
| ENSG00000197121 | PGAP1 | post-GPI attachment to proteins inositol deacylase 1 [Source:HGNC Symbol;Acc:HGNC:25712] | 1.0860 | 1.83E-08 |
| ENSG00000158301 | GPRASP2 | G protein-coupled receptor associated sorting protein 2 [Source:HGNC Symbol;Acc:HGNC:25169] | -0.8375 | 1.83E-08 |
| ENSG00000157600 | TMEM164 | transmembrane protein 164 [Source:HGNC Symbol;Acc:HGNC:26217] | -0.7904 | 1.86E-08 |
| ENSG00000172349 | IL16 | interleukin 16 [Source:HGNC Symbol;Acc:HGNC:5980] | 4.8260 | 1.87E-08 |
| ENSG00000145555 | MYO10 | myosin X [Source:HGNC Symbol;Acc:HGNC:7593] | -0.7337 | 1.89E-08 |
| ENSG00000178301 | AQP11 | aquaporin 11 [Source:HGNC Symbol;Acc:HGNC:19940] | -1.5856 | 1.95E-08 |
| ENSG00000157617 | C2CD2 | C2 calcium dependent domain containing 2 [Source:HGNC Symbol;Acc:HGNC:1266] | -0.6823 | 2.00E-08 |
| ENSG00000137463 | MGARP | mitochondria localized glutamic acid rich protein [Source:HGNC Symbol;Acc:HGNC:29969] | -0.8772 | 2.00E-08 |
| ENSG00000257181 | NA | novel transcript | 0.9651 | 2.02E-08 |
| ENSG00000163659 | TIPARP | TCDD inducible poly(ADP-ribose) polymerase [Source:HGNC Symbol;Acc:HGNC:23696] | 1.1550 | 2.05E-08 |
| ENSG00000081052 | COL4A4 | collagen type IV alpha 4 chain [Source:HGNC Symbol;Acc:HGNC:2206] | -0.6244 | 2.05E-08 |
| ENSG00000097096 | SYDE2 | synapse defective Rho GTPase homolog 2 [Source:HGNC Symbol;Acc:HGNC:25841] | -0.7947 | 2.05E-08 |
| ENSG00000152804 | HHEX | hematopoietically expressed homeobox [Source:HGNC Symbol;Acc:HGNC:4901] | 0.9500 | 2.08E-08 |
| ENSG00000184160 | ADRA2C | adrenoceptor alpha 2C [Source:HGNC Symbol;Acc:HGNC:283] | -1.6120 | 2.08E-08 |
| ENSG00000144357 | UBR3 | ubiquitin protein ligase E3 component n-recognin 3 [Source:HGNC Symbol;Acc:HGNC:30467] | 0.5402 | 2.13E-08 |
| ENSG00000203943 | SAMD13 | sterile alpha motif domain containing 13 [Source:HGNC Symbol;Acc:HGNC:24582] | -1.9205 | 2.13E-08 |
| ENSG00000040487 | SLC66A1 | solute carrier family 66 member 1 [Source:HGNC Symbol;Acc:HGNC:26001] | 0.7399 | 2.14E-08 |
| ENSG00000104368 | PLAT | plasminogen activator, tissue type [Source:HGNC Symbol;Acc:HGNC:9051] | 1.0251 | 2.15E-08 |
| ENSG00000168546 | GFRA2 | GDNF family receptor alpha 2 [Source:HGNC Symbol;Acc:HGNC:4244] | 1.9247 | 2.15E-08 |
| ENSG00000140320 | BAHD1 | bromo adjacent homology domain containing 1 [Source:HGNC Symbol;Acc:HGNC:29153] | -0.6083 | 2.17E-08 |
| ENSG00000128283 | CDC42EP1 | CDC42 effector protein 1 [Source:HGNC Symbol;Acc:HGNC:17014] | -0.5361 | 2.18E-08 |
| ENSG00000179284 | DAND5 | DAN domain BMP antagonist family member 5 [Source:HGNC Symbol;Acc:HGNC:26780] | -2.6908 | 2.18E-08 |
| ENSG00000122642 | FKBP9 | FKBP prolyl isomerase 9 [Source:HGNC Symbol;Acc:HGNC:3725] | -0.6517 | 2.25E-08 |
| ENSG00000062716 | VMP1 | vacuole membrane protein 1 [Source:HGNC Symbol;Acc:HGNC:29559] | 0.5740 | 2.27E-08 |
| ENSG00000105281 | SLC1A5 | solute carrier family 1 member 5 [Source:HGNC Symbol;Acc:HGNC:10943] | 0.7866 | 2.30E-08 |
| ENSG00000107937 | GTPBP4 | GTP binding protein 4 [Source:HGNC Symbol;Acc:HGNC:21535] | 0.8670 | 2.45E-08 |
| ENSG00000177707 | NECTIN3 | nectin cell adhesion molecule 3 [Source:HGNC Symbol;Acc:HGNC:17664] | 0.8208 | 2.48E-08 |
| ENSG00000069667 | RORA | RAR related orphan receptor A [Source:HGNC Symbol;Acc:HGNC:10258] | 0.8431 | 2.53E-08 |
| ENSG00000250358 | LINC02200 | long intergenic non-protein coding RNA 2200 [Source:HGNC Symbol;Acc:HGNC:53066] | -2.5473 | 2.56E-08 |
| ENSG00000123560 | PLP1 | proteolipid protein 1 [Source:HGNC Symbol;Acc:HGNC:9086] | -1.3243 | 2.57E-08 |
| ENSG00000171988 | JMJD1C | jumonji domain containing 1C [Source:HGNC Symbol;Acc:HGNC:12313] | 0.6991 | 2.62E-08 |
| ENSG00000140876 | NUDT7 | nudix hydrolase 7 [Source:HGNC Symbol;Acc:HGNC:8054] | -1.2181 | 2.63E-08 |
| ENSG00000106415 | GLCCI1 | glucocorticoid induced 1 [Source:HGNC Symbol;Acc:HGNC:18713] | -0.9897 | 2.65E-08 |
| ENSG00000148408 | CACNA1B | calcium voltage-gated channel subunit alpha1 B [Source:HGNC Symbol;Acc:HGNC:1389] | -0.6283 | 2.67E-08 |
| ENSG00000129235 | TXNDC17 | thioredoxin domain containing 17 [Source:HGNC Symbol;Acc:HGNC:28218] | 0.5688 | 2.68E-08 |
| ENSG00000137106 | GRHPR | glyoxylate and hydroxypyruvate reductase [Source:HGNC Symbol;Acc:HGNC:4570] | -0.5004 | 2.71E-08 |
| ENSG00000172340 | SUCLG2 | succinate-CoA ligase GDP-forming subunit beta [Source:HGNC Symbol;Acc:HGNC:11450] | -0.5912 | 2.74E-08 |
| ENSG00000184557 | SOCS3 | suppressor of cytokine signaling 3 [Source:HGNC Symbol;Acc:HGNC:19391] | 2.2688 | 2.74E-08 |
| ENSG00000100522 | GNPNAT1 | glucosamine-phosphate N-acetyltransferase 1 [Source:HGNC Symbol;Acc:HGNC:19980] | 0.6795 | 2.77E-08 |
| ENSG00000123130 | ACOT9 | acyl-CoA thioesterase 9 [Source:HGNC Symbol;Acc:HGNC:17152] | 0.6226 | 2.82E-08 |
| ENSG00000144535 | DIS3L2 | DIS3 like 3'-5' exoribonuclease 2 [Source:HGNC Symbol;Acc:HGNC:28648] | 0.5260 | 2.83E-08 |
| ENSG00000135049 | AGTPBP1 | ATP/GTP binding protein 1 [Source:HGNC Symbol;Acc:HGNC:17258] | -0.8338 | 2.88E-08 |

| | | | | |
|---|---|---|---|---|
| ENSG00000128294 | TPST2 | tyrosylprotein sulfotransferase 2 [Source:HGNC Symbol;Acc:HGNC:12021] | -0.6484 | 2.91E-08 |
| ENSG00000130830 | MPP1 | membrane palmitoylated protein 1 [Source:HGNC Symbol;Acc:HGNC:7219] | -0.5397 | 2.93E-08 |
| ENSG00000154553 | PDLIM3 | PDZ and LIM domain 3 [Source:HGNC Symbol;Acc:HGNC:20767] | 2.3984 | 2.93E-08 |
| ENSG00000139970 | RTN1 | reticulon 1 [Source:HGNC Symbol;Acc:HGNC:10467] | -0.6527 | 2.99E-08 |
| ENSG00000114378 | HYAL1 | hyaluronidase 1 [Source:HGNC Symbol;Acc:HGNC:5320] | -0.8036 | 3.06E-08 |
| ENSG00000235961 | PNMA6A | PNMA family member 6A [Source:HGNC Symbol;Acc:HGNC:28248] | -0.8636 | 3.06E-08 |
| ENSG00000003989 | SLC7A2 | solute carrier family 7 member 2 [Source:HGNC Symbol;Acc:HGNC:11060] | -0.8326 | 3.19E-08 |
| ENSG00000121851 | POLR3GL | RNA polymerase III subunit G like [Source:HGNC Symbol;Acc:HGNC:28466] | -1.1321 | 3.29E-08 |
| ENSG00000286169 | PDCD6-AHRR | PDCD6-AHRR readthrough (NMD candidate) [Source:HGNC Symbol;Acc:HGNC:54724] | -0.6339 | 3.29E-08 |
| ENSG00000119718 | EIF2B2 | eukaryotic translation initiation factor 2B subunit beta [Source:HGNC Symbol;Acc:HGNC:3258] | 0.5656 | 3.32E-08 |
| ENSG00000126001 | CEP250 | centrosomal protein 250 [Source:HGNC Symbol;Acc:HGNC:1859] | -1.0755 | 3.34E-08 |
| ENSG00000135269 | TES | testin LIM domain protein [Source:HGNC Symbol;Acc:HGNC:14620] | 0.5961 | 3.39E-08 |
| ENSG00000239672 | NME1 | NME/NM23 nucleoside diphosphate kinase 1 [Source:HGNC Symbol;Acc:HGNC:7849] | 0.5580 | 3.41E-08 |
| ENSG00000125454 | SLC25A19 | solute carrier family 25 member 19 [Source:HGNC Symbol;Acc:HGNC:14409] | 0.9385 | 3.46E-08 |
| ENSG00000130304 | SLC27A1 | solute carrier family 27 member 1 [Source:HGNC Symbol;Acc:HGNC:10995] | -1.3688 | 3.46E-08 |
| ENSG00000198920 | KIAA0753 | KIAA0753 [Source:HGNC Symbol;Acc:HGNC:29110] | 0.6589 | 3.53E-08 |
| ENSG00000164938 | TP53INP1 | tumor protein p53 inducible nuclear protein 1 [Source:HGNC Symbol;Acc:HGNC:18022] | -0.8248 | 3.55E-08 |
| ENSG00000113657 | DPYSL3 | dihydropyrimidinase like 3 [Source:HGNC Symbol;Acc:HGNC:3015] | -0.8592 | 3.57E-08 |
| ENSG00000162616 | DNAJB4 | DnaJ heat shock protein family (Hsp40) member B4 [Source:HGNC Symbol;Acc:HGNC:14886] | 0.8320 | 3.57E-08 |
| ENSG00000168792 | ABHD15 | abhydrolase domain containing 15 [Source:HGNC Symbol;Acc:HGNC:26971] | -0.7866 | 3.63E-08 |
| ENSG00000036530 | CYP46A1 | cytochrome P450 family 46 subfamily A member 1 [Source:HGNC Symbol;Acc:HGNC:2641] | -1.0263 | 3.64E-08 |
| ENSG00000133121 | STARD13 | StAR related lipid transfer domain containing 13 [Source:HGNC Symbol;Acc:HGNC:19164] | 0.9553 | 3.64E-08 |
| ENSG00000171490 | RSL1D1 | ribosomal L1 domain containing 1 [Source:HGNC Symbol;Acc:HGNC:24534] | 0.6042 | 3.66E-08 |
| ENSG00000137942 | FNBP1L | formin binding protein 1 like [Source:HGNC Symbol;Acc:HGNC:20851] | -0.6804 | 3.70E-08 |
| ENSG00000139187 | KLRG1 | killer cell lectin like receptor G1 [Source:HGNC Symbol;Acc:HGNC:6380] | -2.3533 | 3.77E-08 |
| ENSG00000175832 | ETV4 | ETS variant transcription factor 4 [Source:HGNC Symbol;Acc:HGNC:3493] | 3.7654 | 3.84E-08 |
| ENSG00000168216 | LMBRD1 | LMBR1 domain containing 1 [Source:HGNC Symbol;Acc:HGNC:23038] | 0.5017 | 3.86E-08 |
| ENSG00000165548 | TMEM63C | transmembrane protein 63C [Source:HGNC Symbol;Acc:HGNC:23787] | -0.9331 | 3.91E-08 |
| ENSG00000143771 | CNIH4 | cornichon family AMPA receptor auxiliary protein 4 [Source:HGNC Symbol;Acc:HGNC:25013] | 0.5604 | 3.97E-08 |
| ENSG00000168268 | NT5DC2 | 5'-nucleotidase domain containing 2 [Source:HGNC Symbol;Acc:HGNC:25717] | -0.7989 | 4.05E-08 |
| ENSG00000115290 | GRB14 | growth factor receptor bound protein 14 [Source:HGNC Symbol;Acc:HGNC:4565] | -0.6869 | 4.08E-08 |
| ENSG00000253508 | NA | novel transcript | 1.8152 | 4.10E-08 |
| ENSG00000179041 | RRS1 | ribosome biogenesis regulator 1 homolog [Source:HGNC Symbol;Acc:HGNC:17083] | 0.6195 | 4.15E-08 |
| ENSG00000005882 | PDK2 | pyruvate dehydrogenase kinase 2 [Source:HGNC Symbol;Acc:HGNC:8810] | -0.8048 | 4.20E-08 |
| ENSG00000169826 | CSGALNACT2 | chondroitin sulfate N-acetylgalactosaminyltransferase 2 [Source:HGNC Symbol;Acc:HGNC:24292] | 0.5488 | 4.21E-08 |
| ENSG00000095585 | BLNK | B cell linker [Source:HGNC Symbol;Acc:HGNC:14211] | -1.3858 | 4.21E-08 |
| ENSG00000182165 | TP53TG1 | TP53 target 1 [Source:HGNC Symbol;Acc:HGNC:17026] | -1.0799 | 4.23E-08 |
| ENSG00000084070 | SMAP2 | small ArfGAP2 [Source:HGNC Symbol;Acc:HGNC:25082] | 0.6630 | 4.24E-08 |
| ENSG00000168542 | COL3A1 | collagen type III alpha 1 chain [Source:HGNC Symbol;Acc:HGNC:2201] | 2.6152 | 4.28E-08 |
| ENSG00000247626 | mars-02 | methionyl-tRNA synthetase 2, mitochondrial [Source:HGNC Symbol;Acc:HGNC:25133] | 0.8670 | 4.29E-08 |
| ENSG00000109881 | CCDC34 | coiled-coil domain containing 34 [Source:HGNC Symbol;Acc:HGNC:25079] | -0.8571 | 4.29E-08 |
| ENSG00000184619 | KRBA2 | KRAB-A domain containing 2 [Source:HGNC Symbol;Acc:HGNC:26989] | -1.2026 | 4.32E-08 |
| ENSG00000128510 | CPA4 | carboxypeptidase A4 [Source:HGNC Symbol;Acc:HGNC:15740] | 2.8501 | 4.33E-08 |

| | | | | |
|---|---|---|---|---|
| ENSG00000066926 | FECH | ferrochelatase [Source:HGNC Symbol;Acc:HGNC:3647] | -0.5801 | 4.41E-08 |
| ENSG00000153044 | CENPH | centromere protein H [Source:HGNC Symbol;Acc:HGNC:17268] | -0.6846 | 4.41E-08 |
| ENSG00000070495 | JMJD6 | jumonji domain containing 6, arginine demethylase and lysine hydroxylase [Source:HGNC Symbol;Acc:HGNC:19355] | 0.8727 | 4.45E-08 |
| ENSG00000119979 | DENND10 | DENN domain containing 10 [Source:HGNC Symbol;Acc:HGNC:31793] | -0.5528 | 4.62E-08 |
| ENSG00000135547 | HEY2 | hes related family bHLH transcription factor with YRPW motif 2 [Source:HGNC Symbol;Acc:HGNC:4881] | -1.2638 | 4.70E-08 |
| ENSG00000084731 | KIF3C | kinesin family member 3C [Source:HGNC Symbol;Acc:HGNC:6321] | -0.7893 | 4.71E-08 |
| ENSG00000141404 | GNAL | G protein subunit alpha L [Source:HGNC Symbol;Acc:HGNC:4388] | -1.1642 | 4.74E-08 |
| ENSG00000086189 | DIMT1 | DIMT1 rRNA methyltransferase and ribosome maturation factor [Source:HGNC Symbol;Acc:HGNC:30217] | 0.6045 | 4.83E-08 |
| ENSG00000273599 | NA | novel transcript, antisense to CTBP2 | 1.2000 | 4.95E-08 |
| ENSG00000287316 | NA | novel transcript | 1.9036 | 5.07E-08 |
| ENSG00000100297 | MCM5 | minichromosome maintenance complex component 5 [Source:HGNC Symbol;Acc:HGNC:6948] | -0.7586 | 5.12E-08 |
| ENSG00000124140 | SLC12A5 | solute carrier family 12 member 5 [Source:HGNC Symbol;Acc:HGNC:13818] | 0.6885 | 5.14E-08 |
| ENSG00000108557 | RAI1 | retinoic acid induced 1 [Source:HGNC Symbol;Acc:HGNC:9834] | 0.5798 | 5.19E-08 |
| ENSG00000136840 | ST6GALNAC4 | ST6 N-acetylgalactosaminide alpha-2,6-sialyltransferase 4 [Source:HGNC Symbol;Acc:HGNC:17846] | 0.6394 | 5.19E-08 |
| ENSG00000139410 | SDSL | serine dehydratase like [Source:HGNC Symbol;Acc:HGNC:30404] | 0.6668 | 5.22E-08 |
| ENSG00000149925 | ALDOA | aldolase, fructose-bisphosphate A [Source:HGNC Symbol;Acc:HGNC:414] | 0.7300 | 5.26E-08 |
| ENSG00000114209 | PDCD10 | programmed cell death 10 [Source:HGNC Symbol;Acc:HGNC:8761] | 0.5353 | 5.27E-08 |
| ENSG00000167767 | KRT80 | keratin 80 [Source:HGNC Symbol;Acc:HGNC:27056] | 1.8725 | 5.28E-08 |
| ENSG00000186205 | MTARC1 | mitochondrial amidoxime reducing component 1 [Source:HGNC Symbol;Acc:HGNC:26189] | -0.8972 | 5.29E-08 |
| ENSG00000204536 | CCHCR1 | coiled-coil alpha-helical rod protein 1 [Source:HGNC Symbol;Acc:HGNC:13930] | -0.7929 | 5.29E-08 |
| ENSG00000234390 | USP27X-AS1 | USP27X antisense RNA 1 (head to head) [Source:HGNC Symbol;Acc:HGNC:27249] | -1.1908 | 5.31E-08 |
| ENSG00000169641 | LUZP1 | leucine zipper protein 1 [Source:HGNC Symbol;Acc:HGNC:14985] | 0.6289 | 5.33E-08 |
| ENSG00000134215 | VAV3 | vav guanine nucleotide exchange factor 3 [Source:HGNC Symbol;Acc:HGNC:12659] | -0.7403 | 5.38E-08 |
| ENSG00000181788 | SIAH2 | siah E3 ubiquitin protein ligase 2 [Source:HGNC Symbol;Acc:HGNC:10858] | 0.5171 | 5.46E-08 |
| ENSG00000259884 | NR4A1AS | NR4A1 antisense RNA [Source:HGNC Symbol;Acc:HGNC:54409] | 1.8733 | 5.57E-08 |
| ENSG00000153162 | BMP6 | bone morphogenetic protein 6 [Source:HGNC Symbol;Acc:HGNC:1073] | 1.3897 | 5.58E-08 |
| ENSG00000187955 | COL14A1 | collagen type XIV alpha 1 chain [Source:HGNC Symbol;Acc:HGNC:2191] | -2.7202 | 5.60E-08 |
| ENSG00000173334 | TRIB1 | tribbles pseudokinase 1 [Source:HGNC Symbol;Acc:HGNC:16891] | 2.1216 | 5.60E-08 |
| ENSG00000035862 | TIMP2 | TIMP metallopeptidase inhibitor 2 [Source:HGNC Symbol;Acc:HGNC:11821] | -0.7462 | 5.61E-08 |
| ENSG00000163623 | NKX6-1 | NK6 homeobox 1 [Source:HGNC Symbol;Acc:HGNC:7839] | 1.3521 | 5.61E-08 |
| ENSG00000128513 | POT1 | protection of telomeres 1 [Source:HGNC Symbol;Acc:HGNC:17284] | -0.6429 | 5.63E-08 |
| ENSG00000109062 | SLC9A3R1 | SLC9A3 regulator 1 [Source:HGNC Symbol;Acc:HGNC:11075] | 1.2666 | 5.66E-08 |
| ENSG00000148123 | PLPPR1 | phospholipid phosphatase related 1 [Source:HGNC Symbol;Acc:HGNC:25993] | -0.7877 | 5.68E-08 |
| ENSG00000120913 | PDLIM2 | PDZ and LIM domain 2 [Source:HGNC Symbol;Acc:HGNC:13992] | -0.6837 | 5.71E-08 |
| ENSG00000143256 | PFDN2 | prefoldin subunit 2 [Source:HGNC Symbol;Acc:HGNC:8867] | 0.5379 | 5.79E-08 |
| ENSG00000185559 | DLK1 | delta like non-canonical Notch ligand 1 [Source:HGNC Symbol;Acc:HGNC:2907] | 0.5692 | 5.83E-08 |
| ENSG00000184304 | PRKD1 | protein kinase D1 [Source:HGNC Symbol;Acc:HGNC:9407] | -0.8423 | 5.88E-08 |
| ENSG00000248334 | WHAMMP2 | WAS protein homolog associated with actin, golgi membranes and microtubules pseudogene 2 [Source:HGNC Symbol;Acc:HGNC:32360] | -0.9826 | 5.91E-08 |
| ENSG00000157766 | ACAN | aggrecan [Source:HGNC Symbol;Acc:HGNC:319] | -1.3959 | 5.99E-08 |
| ENSG00000187555 | USP7 | ubiquitin specific peptidase 7 [Source:HGNC Symbol;Acc:HGNC:12630] | 0.5058 | 6.37E-08 |
| ENSG00000237880 | LINC01885 | long intergenic non-protein coding RNA 1885 [Source:HGNC Symbol;Acc:HGNC:52704] | -1.6943 | 6.37E-08 |

| | | | | |
|---|---|---|---|---|
| ENSG00000264207 | NA | novel transcript, antisense HIST2H4A | -1.1836 | 6.44E-08 |
| ENSG00000141551 | CSNK1D | casein kinase 1 delta [Source:HGNC Symbol;Acc:HGNC:2452] | 0.6703 | 6.61E-08 |
| ENSG00000116977 | LGALS8 | galectin 8 [Source:HGNC Symbol;Acc:HGNC:6569] | 0.9102 | 6.63E-08 |
| ENSG00000145687 | SSBP2 | single stranded DNA binding protein 2 [Source:HGNC Symbol;Acc:HGNC:15831] | -0.6278 | 6.63E-08 |
| ENSG00000184489 | PTP4A3 | protein tyrosine phosphatase 4A3 [Source:HGNC Symbol;Acc:HGNC:9636] | -1.8142 | 6.74E-08 |
| ENSG00000120324 | PCDHB10 | protocadherin beta 10 [Source:HGNC Symbol;Acc:HGNC:8681] | -0.8778 | 6.74E-08 |
| ENSG00000104883 | PEX11G | peroxisomal biogenesis factor 11 gamma [Source:HGNC Symbol;Acc:HGNC:20208] | -1.3732 | 6.76E-08 |
| ENSG00000140563 | MCTP2 | multiple C2 and transmembrane domain containing 2 [Source:HGNC Symbol;Acc:HGNC:25636] | 0.9371 | 6.79E-08 |
| ENSG00000185112 | FAM43A | family with sequence similarity 43 member A [Source:HGNC Symbol;Acc:HGNC:26888] | -0.7352 | 6.85E-08 |
| ENSG00000196072 | BLOC1S2 | biogenesis of lysosomal organelles complex 1 subunit 2 [Source:HGNC Symbol;Acc:HGNC:20984] | 0.7300 | 6.90E-08 |
| ENSG00000196497 | IPO4 | importin 4 [Source:HGNC Symbol;Acc:HGNC:19426] | 0.9321 | 6.91E-08 |
| ENSG00000099330 | OCEL1 | occludin/ELL domain containing 1 [Source:HGNC Symbol;Acc:HGNC:26221] | -0.9396 | 7.18E-08 |
| ENSG00000067445 | TRO | trophinin [Source:HGNC Symbol;Acc:HGNC:12326] | -0.5514 | 7.23E-08 |
| ENSG00000151229 | SLC2A13 | solute carrier family 2 member 13 [Source:HGNC Symbol;Acc:HGNC:15956] | 1.0330 | 7.32E-08 |
| ENSG00000204711 | C9orf135 | chromosome 9 open reading frame 135 [Source:HGNC Symbol;Acc:HGNC:31422] | 1.6269 | 7.36E-08 |
| ENSG00000115325 | DOK1 | docking protein 1 [Source:HGNC Symbol;Acc:HGNC:2990] | -0.8206 | 7.43E-08 |
| ENSG00000164877 | MICALL2 | MICAL like 2 [Source:HGNC Symbol;Acc:HGNC:29672] | 1.3406 | 7.64E-08 |
| ENSG00000176597 | B3GNT5 | UDP-GlcNAc:betaGal beta-1,3-N-acetylglucosaminyltransferase 5 [Source:HGNC Symbol;Acc:HGNC:15684] | 2.4719 | 7.77E-08 |
| ENSG00000254093 | PINX1 | PIN2 (TERF1) interacting telomerase inhibitor 1 [Source:HGNC Symbol;Acc:HGNC:30046] | 1.1755 | 7.77E-08 |
| ENSG00000179820 | MYADM | myeloid associated differentiation marker [Source:HGNC Symbol;Acc:HGNC:7544] | 0.8448 | 7.80E-08 |
| ENSG00000243244 | STON1 | stonin 1 [Source:HGNC Symbol;Acc:HGNC:17003] | -0.9600 | 7.93E-08 |
| ENSG00000114670 | NEK11 | NIMA related kinase 11 [Source:HGNC Symbol;Acc:HGNC:18593] | -1.0077 | 8.26E-08 |
| ENSG00000198843 | SELENOT | selenoprotein T [Source:HGNC Symbol;Acc:HGNC:18136] | 0.5558 | 8.39E-08 |
| ENSG00000198929 | NOS1AP | nitric oxide synthase 1 adaptor protein [Source:HGNC Symbol;Acc:HGNC:16859] | -1.1131 | 8.57E-08 |
| ENSG00000137959 | IFI44L | interferon induced protein 44 like [Source:HGNC Symbol;Acc:HGNC:17817] | -1.0678 | 8.60E-08 |
| ENSG00000221890 | NPTXR | neuronal pentraxin receptor [Source:HGNC Symbol;Acc:HGNC:7954] | -0.7358 | 8.65E-08 |
| ENSG00000099385 | BCL7C | BAF chromatin remodeling complex subunit BCL7C [Source:HGNC Symbol;Acc:HGNC:1006] | -0.6094 | 8.72E-08 |
| ENSG00000090971 | NAT14 | N-acetyltransferase 14 (putative) [Source:HGNC Symbol;Acc:HGNC:28918] | -0.6245 | 8.75E-08 |
| ENSG00000106080 | FKBP14 | FKBP prolyl isomerase 14 [Source:HGNC Symbol;Acc:HGNC:18625] | 0.6995 | 8.75E-08 |
| ENSG00000138759 | FRAS1 | Fraser extracellular matrix complex subunit 1 [Source:HGNC Symbol;Acc:HGNC:19185] | 0.9768 | 8.78E-08 |
| ENSG00000215808 | LINC01139 | long intergenic non-protein coding RNA 1139 [Source:HGNC Symbol;Acc:HGNC:27924] | -0.7296 | 8.80E-08 |
| ENSG00000071051 | NCK2 | NCK adaptor protein 2 [Source:HGNC Symbol;Acc:HGNC:7665] | -0.5626 | 8.81E-08 |
| ENSG00000127080 | IPPK | inositol-pentakisphosphate 2-kinase [Source:HGNC Symbol;Acc:HGNC:14645] | 0.5892 | 8.91E-08 |
| ENSG00000168078 | PBK | PDZ binding kinase [Source:HGNC Symbol;Acc:HGNC:18282] | -0.6526 | 8.95E-08 |
| ENSG00000134058 | CDK7 | cyclin dependent kinase 7 [Source:HGNC Symbol;Acc:HGNC:1778] | 0.6679 | 8.97E-08 |
| ENSG00000155189 | AGPAT5 | 1-acylglycerol-3-phosphate O-acyltransferase 5 [Source:HGNC Symbol;Acc:HGNC:20886] | 1.0149 | 8.97E-08 |
| ENSG00000149571 | KIRREL3 | kirre like nephrin family adhesion molecule 3 [Source:HGNC Symbol;Acc:HGNC:23204] | 1.3226 | 9.15E-08 |
| ENSG00000127252 | PLAAT1 | phospholipase A and acyltransferase 1 [Source:HGNC Symbol;Acc:HGNC:14922] | -1.0035 | 9.25E-08 |
| ENSG00000054967 | RELT | RELT TNF receptor [Source:HGNC Symbol;Acc:HGNC:13764] | 1.2157 | 9.39E-08 |
| ENSG00000125966 | MMP24 | matrix metallopeptidase 24 [Source:HGNC Symbol;Acc:HGNC:7172] | 1.1934 | 9.55E-08 |
| ENSG00000104147 | OIP5 | Opa interacting protein 5 [Source:HGNC Symbol;Acc:HGNC:20300] | -0.7942 | 9.86E-08 |
| ENSG00000132749 | TESMIN | testis expressed metallothionein like protein [Source:HGNC Symbol;Acc:HGNC:7446] | -1.0290 | 9.89E-08 |

| | | | | |
|---|---|---|---|---|
| ENSG00000113845 | TIMMDC1 | translocase of inner mitochondrial membrane domain containing 1 [Source:HGNC Symbol;Acc:HGNC:1321] | -0.6795 | 9.93E-08 |
| ENSG00000102781 | KATNAL1 | katanin catalytic subunit A1 like 1 [Source:HGNC Symbol;Acc:HGNC:28361] | -0.5890 | 9.97E-08 |
| ENSG00000165046 | LETM2 | leucine zipper and EF-hand containing transmembrane protein 2 [Source:HGNC Symbol;Acc:HGNC:14648] | 1.4089 | 1.01E-07 |
| ENSG00000174156 | GSTA3 | glutathione S-transferase alpha 3 [Source:HGNC Symbol;Acc:HGNC:4628] | -0.8028 | 1.02E-07 |
| ENSG00000124275 | MTRR | 5-methyltetrahydrofolate-homocysteine methyltransferase reductase [Source:HGNC Symbol;Acc:HGNC:7473] | 0.6371 | 1.03E-07 |
| ENSG00000035141 | FAM136A | family with sequence similarity 136 member A [Source:HGNC Symbol;Acc:HGNC:25911] | 0.5626 | 1.04E-07 |
| ENSG00000088854 | C20orf194 | chromosome 20 open reading frame 194 [Source:HGNC Symbol;Acc:HGNC:17721] | -0.7340 | 1.04E-07 |
| ENSG00000186462 | NAP1L2 | nucleosome assembly protein 1 like 2 [Source:HGNC Symbol;Acc:HGNC:7638] | -0.8006 | 1.09E-07 |
| ENSG00000139971 | ARMH4 | armadillo like helical domain containing 4 [Source:HGNC Symbol;Acc:HGNC:19846] | -0.6937 | 1.10E-07 |
| ENSG00000068308 | OTUD5 | OTU deubiquitinase 5 [Source:HGNC Symbol;Acc:HGNC:25402] | 0.5130 | 1.11E-07 |
| ENSG00000280071 | GATD3B | glutamine amidotransferase like class 1 domain containing 3B [Source:HGNC Symbol;Acc:HGNC:53816] | -0.5094 | 1.12E-07 |
| ENSG00000110619 | CARS1 | cysteinyl-tRNA synthetase 1 [Source:HGNC Symbol;Acc:HGNC:1493] | 0.6383 | 1.14E-07 |
| ENSG00000069849 | ATP1B3 | ATPase Na+/K+ transporting subunit beta 3 [Source:HGNC Symbol;Acc:HGNC:806] | 0.6396 | 1.15E-07 |
| ENSG00000167525 | PROCA1 | protein interacting with cyclin A1 [Source:HGNC Symbol;Acc:HGNC:28600] | -1.1662 | 1.15E-07 |
| ENSG00000227225 | MTND1P14 | MT-ND1 pseudogene 14 [Source:HGNC Symbol;Acc:HGNC:42063] | 2.2833 | 1.17E-07 |
| ENSG00000129003 | VPS13C | vacuolar protein sorting 13 homolog C [Source:HGNC Symbol;Acc:HGNC:23594] | 0.6367 | 1.19E-07 |
| ENSG00000196396 | PTPN1 | protein tyrosine phosphatase non-receptor type 1 [Source:HGNC Symbol;Acc:HGNC:9642] | 0.5074 | 1.20E-07 |
| ENSG00000139352 | ASCL1 | achaete-scute family bHLH transcription factor 1 [Source:HGNC Symbol;Acc:HGNC:738] | -1.7386 | 1.20E-07 |
| ENSG00000222009 | BTBD19 | BTB domain containing 19 [Source:HGNC Symbol;Acc:HGNC:27145] | 1.0646 | 1.20E-07 |
| ENSG00000287335 | NA | novel transcript | -0.9075 | 1.22E-07 |
| ENSG00000139899 | CBLN3 | cerebellin 3 precursor [Source:HGNC Symbol;Acc:HGNC:20146] | -1.0152 | 1.22E-07 |
| ENSG00000173276 | ZBTB21 | zinc finger and BTB domain containing 21 [Source:HGNC Symbol;Acc:HGNC:13083] | 0.8952 | 1.22E-07 |
| ENSG00000104998 | IL27RA | interleukin 27 receptor subunit alpha [Source:HGNC Symbol;Acc:HGNC:17290] | 1.5781 | 1.23E-07 |
| ENSG00000120594 | PLXDC2 | plexin domain containing 2 [Source:HGNC Symbol;Acc:HGNC:21013] | -0.6203 | 1.23E-07 |
| ENSG00000203791 | EEF1AKMT2 | EEF1A lysine methyltransferase 2 [Source:HGNC Symbol;Acc:HGNC:33787] | -0.6581 | 1.23E-07 |
| ENSG00000170836 | PPM1D | protein phosphatase, Mg2+/Mn2+ dependent 1D [Source:HGNC Symbol;Acc:HGNC:9277] | 0.6571 | 1.24E-07 |
| ENSG00000178467 | P4HTM | prolyl 4-hydroxylase, transmembrane [Source:HGNC Symbol;Acc:HGNC:28858] | 0.6180 | 1.24E-07 |
| ENSG00000070081 | NUCB2 | nucleobindin 2 [Source:HGNC Symbol;Acc:HGNC:8044] | 0.5584 | 1.26E-07 |
| ENSG00000088808 | PPP1R13B | protein phosphatase 1 regulatory subunit 13B [Source:HGNC Symbol;Acc:HGNC:14950] | -0.7748 | 1.27E-07 |
| ENSG00000100505 | TRIM9 | tripartite motif containing 9 [Source:HGNC Symbol;Acc:HGNC:16288] | 1.5040 | 1.28E-07 |
| ENSG00000153317 | ASAP1 | ArfGAP with SH3 domain, ankyrin repeat and PH domain 1 [Source:HGNC Symbol;Acc:HGNC:2720] | 0.5324 | 1.31E-07 |
| ENSG00000124406 | ATP8A1 | ATPase phospholipid transporting 8A1 [Source:HGNC Symbol;Acc:HGNC:13531] | -1.1465 | 1.33E-07 |
| ENSG00000187954 | CYHR1 | cysteine and histidine rich 1 [Source:HGNC Symbol;Acc:HGNC:17806] | -0.5741 | 1.33E-07 |
| ENSG00000132382 | MYBBP1A | MYB binding protein 1a [Source:HGNC Symbol;Acc:HGNC:7546] | 0.5436 | 1.33E-07 |
| ENSG00000126803 | HSPA2 | heat shock protein family A (Hsp70) member 2 [Source:HGNC Symbol;Acc:HGNC:5235] | -1.0840 | 1.36E-07 |
| ENSG00000143797 | MBOAT2 | membrane bound O-acyltransferase domain containing 2 [Source:HGNC Symbol;Acc:HGNC:25193] | -0.8909 | 1.38E-07 |
| ENSG00000157107 | FCHO2 | FCH and mu domain containing endocytic adaptor 2 [Source:HGNC Symbol;Acc:HGNC:25180] | -0.7541 | 1.38E-07 |
| ENSG00000146409 | SLC18B1 | solute carrier family 18 member B1 [Source:HGNC Symbol;Acc:HGNC:21573] | -0.5116 | 1.39E-07 |
| ENSG00000124784 | RIOK1 | RIO kinase 1 [Source:HGNC Symbol;Acc:HGNC:18656] | 0.7493 | 1.39E-07 |
| ENSG00000186470 | BTN3A2 | butyrophilin subfamily 3 member A2 [Source:HGNC Symbol;Acc:HGNC:1139] | -0.5412 | 1.40E-07 |
| ENSG00000138629 | UBL7 | ubiquitin like 7 [Source:HGNC Symbol;Acc:HGNC:28221] | -0.5809 | 1.42E-07 |
| ENSG00000161277 | THAP8 | THAP domain containing 8 [Source:HGNC Symbol;Acc:HGNC:23191] | -1.1522 | 1.42E-07 |

| | | | | |
|---|---|---|---|---|
| ENSG00000268560 | NA | novel transcript | -2.2067 | 1.43E-07 |
| ENSG00000163702 | IL17RC | interleukin 17 receptor C [Source:HGNC Symbol;Acc:HGNC:18358] | -0.8305 | 1.43E-07 |
| ENSG00000256977 | LIMS3 | LIM zinc finger domain containing 3 [Source:HGNC Symbol;Acc:HGNC:30047] | 1.5840 | 1.45E-07 |
| ENSG00000137502 | RAB30 | RAB30, member RAS oncogene family [Source:HGNC Symbol;Acc:HGNC:9770] | -0.8075 | 1.46E-07 |
| ENSG00000155495 | MAGEC1 | MAGE family member C1 [Source:HGNC Symbol;Acc:HGNC:6812] | 0.5345 | 1.47E-07 |
| ENSG00000120519 | SLC10A7 | solute carrier family 10 member 7 [Source:HGNC Symbol;Acc:HGNC:23088] | 1.4155 | 1.49E-07 |
| ENSG00000161551 | ZNF577 | zinc finger protein 577 [Source:HGNC Symbol;Acc:HGNC:28673] | -0.9243 | 1.49E-07 |
| ENSG00000119888 | EPCAM | epithelial cell adhesion molecule [Source:HGNC Symbol;Acc:HGNC:11529] | 1.3762 | 1.50E-07 |
| ENSG00000151224 | MAT1A | methionine adenosyltransferase 1A [Source:HGNC Symbol;Acc:HGNC:6903] | -2.8969 | 1.51E-07 |
| ENSG00000169689 | CENPX | centromere protein X [Source:HGNC Symbol;Acc:HGNC:11422] | -0.8693 | 1.51E-07 |
| ENSG00000127663 | KDM4B | lysine demethylase 4B [Source:HGNC Symbol;Acc:HGNC:29136] | 0.5167 | 1.55E-07 |
| ENSG00000107959 | PITRM1 | pitrilysin metallopeptidase 1 [Source:HGNC Symbol;Acc:HGNC:17663] | 0.5386 | 1.55E-07 |
| ENSG00000108219 | TSPAN14 | tetraspanin 14 [Source:HGNC Symbol;Acc:HGNC:23303] | -0.6586 | 1.57E-07 |
| ENSG00000259969 | NA | novel transcript, overlapping C14orf37 | -1.1378 | 1.58E-07 |
| ENSG00000272886 | DCP1A | decapping mRNA 1A [Source:HGNC Symbol;Acc:HGNC:18714] | 0.5339 | 1.58E-07 |
| ENSG00000165912 | PACSIN3 | protein kinase C and casein kinase substrate in neurons 3 [Source:HGNC Symbol;Acc:HGNC:8572] | -0.5016 | 1.60E-07 |
| ENSG00000203880 | PCMTD2 | protein-L-isoaspartate (D-aspartate) O-methyltransferase domain containing 2 [Source:HGNC Symbol;Acc:HGNC:15882] | -0.8004 | 1.63E-07 |
| ENSG00000165312 | OTUD1 | OTU deubiquitinase 1 [Source:HGNC Symbol;Acc:HGNC:27346] | -1.1381 | 1.63E-07 |
| ENSG00000165322 | ARHGAP12 | Rho GTPase activating protein 12 [Source:HGNC Symbol;Acc:HGNC:16348] | 0.5846 | 1.63E-07 |
| ENSG00000047634 | SCML1 | Scm polycomb group protein like 1 [Source:HGNC Symbol;Acc:HGNC:10580] | 0.5581 | 1.63E-07 |
| ENSG00000064763 | FAR2 | fatty acyl-CoA reductase 2 [Source:HGNC Symbol;Acc:HGNC:25531] | 0.9653 | 1.63E-07 |
| ENSG00000182158 | CREB3L2 | cAMP responsive element binding protein 3 like 2 [Source:HGNC Symbol;Acc:HGNC:23720] | 0.6560 | 1.66E-07 |
| ENSG00000067334 | DNTTIP2 | deoxynucleotidyltransferase terminal interacting protein 2 [Source:HGNC Symbol;Acc:HGNC:24013] | 0.5276 | 1.67E-07 |
| ENSG00000187624 | C17orf97 | chromosome 17 open reading frame 97 [Source:HGNC Symbol;Acc:HGNC:33800] | 1.1261 | 1.69E-07 |
| ENSG00000268001 | CARD8-AS1 | CARD8 antisense RNA 1 [Source:HGNC Symbol;Acc:HGNC:51408] | 1.0297 | 1.69E-07 |
| ENSG00000115652 | UXS1 | UDP-glucuronate decarboxylase 1 [Source:HGNC Symbol;Acc:HGNC:17729] | 0.6374 | 1.72E-07 |
| ENSG00000274253 | NA | novel transcript, sense overlapping NIPA1 | -0.8518 | 1.72E-07 |
| ENSG00000135336 | ORC3 | origin recognition complex subunit 3 [Source:HGNC Symbol;Acc:HGNC:8489] | -0.5873 | 1.75E-07 |
| ENSG00000213782 | DDX47 | DEAD-box helicase 47 [Source:HGNC Symbol;Acc:HGNC:18682] | 0.5204 | 1.75E-07 |
| ENSG00000164104 | HMGB2 | high mobility group box 2 [Source:HGNC Symbol;Acc:HGNC:5000] | -0.6014 | 1.81E-07 |
| ENSG00000159399 | HK2 | hexokinase 2 [Source:HGNC Symbol;Acc:HGNC:4923] | 2.2578 | 1.82E-07 |
| ENSG00000161010 | MRNIP | MRN complex interacting protein [Source:HGNC Symbol;Acc:HGNC:30817] | -0.6987 | 1.83E-07 |
| ENSG00000009780 | FAM76A | family with sequence similarity 76 member A [Source:HGNC Symbol;Acc:HGNC:28530] | -0.8533 | 1.84E-07 |
| ENSG00000100207 | TCF20 | transcription factor 20 [Source:HGNC Symbol;Acc:HGNC:11631] | -0.5978 | 1.84E-07 |
| ENSG00000273033 | LINC02035 | long intergenic non-protein coding RNA 2035 [Source:HGNC Symbol;Acc:HGNC:52875] | 0.8797 | 1.84E-07 |
| ENSG00000226137 | BAIAP2-DT | BAIAP2 divergent transcript [Source:HGNC Symbol;Acc:HGNC:44342] | -0.8489 | 1.85E-07 |
| ENSG00000099864 | PALM | paralemmin [Source:HGNC Symbol;Acc:HGNC:8594] | -0.9953 | 1.85E-07 |
| ENSG00000001461 | NIPAL3 | NIPA like domain containing 3 [Source:HGNC Symbol;Acc:HGNC:25233] | -0.5136 | 1.88E-07 |
| ENSG00000173988 | LRRC63 | leucine rich repeat containing 63 [Source:HGNC Symbol;Acc:HGNC:34296] | -0.8610 | 1.91E-07 |
| ENSG00000136152 | COG3 | component of oligomeric golgi complex 3 [Source:HGNC Symbol;Acc:HGNC:18619] | 0.6391 | 1.91E-07 |
| ENSG00000100345 | MYH9 | myosin heavy chain 9 [Source:HGNC Symbol;Acc:HGNC:7579] | 0.5275 | 1.92E-07 |
| ENSG00000136295 | TTYH3 | tweety family member 3 [Source:HGNC Symbol;Acc:HGNC:22222] | 0.6693 | 1.93E-07 |

| | | | | |
|---|---|---|---|---|
| ENSG00000171604 | CXXC5 | CXXC finger protein 5 [Source:HGNC Symbol;Acc:HGNC:26943] | -1.5859 | 1.95E-07 |
| ENSG00000116285 | ERRFI1 | ERBB receptor feedback inhibitor 1 [Source:HGNC Symbol;Acc:HGNC:18185] | 1.1308 | 1.98E-07 |
| ENSG00000078967 | UBE2D4 | ubiquitin conjugating enzyme E2 D4 (putative) [Source:HGNC Symbol;Acc:HGNC:21647] | -0.5115 | 2.00E-07 |
| ENSG00000130382 | MLLT1 | MLLT1 super elongation complex subunit [Source:HGNC Symbol;Acc:HGNC:7134] | -0.5221 | 2.00E-07 |
| ENSG00000233369 | GTF2IP4 | general transcription factor IIi pseudogene 4 [Source:HGNC Symbol;Acc:HGNC:51716] | -0.6501 | 2.04E-07 |
| ENSG00000156990 | RPUSD3 | RNA pseudouridine synthase D3 [Source:HGNC Symbol;Acc:HGNC:28437] | -0.6301 | 2.04E-07 |
| ENSG00000164308 | ERAP2 | endoplasmic reticulum aminopeptidase 2 [Source:HGNC Symbol;Acc:HGNC:29499] | -0.6114 | 2.08E-07 |
| ENSG00000163686 | ABHD6 | abhydrolase domain containing 6, acylglycerol lipase [Source:HGNC Symbol;Acc:HGNC:21398] | -0.6115 | 2.16E-07 |
| ENSG00000183960 | KCNH8 | potassium voltage-gated channel subfamily H member 8 [Source:HGNC Symbol;Acc:HGNC:18864] | 0.8286 | 2.17E-07 |
| ENSG00000140284 | SLC27A2 | solute carrier family 27 member 2 [Source:HGNC Symbol;Acc:HGNC:10996] | 2.8802 | 2.17E-07 |
| ENSG00000162552 | WNT4 | Wnt family member 4 [Source:HGNC Symbol;Acc:HGNC:12783] | -0.8429 | 2.18E-07 |
| ENSG00000163864 | NMNAT3 | nicotinamide nucleotide adenylyltransferase 3 [Source:HGNC Symbol;Acc:HGNC:20989] | -1.4239 | 2.18E-07 |
| ENSG00000138688 | KIAA1109 | KIAA1109 [Source:HGNC Symbol;Acc:HGNC:26953] | -0.5549 | 2.22E-07 |
| ENSG00000247121 | NA | novel transcript, antisense to LNPEP and ERAP2 | -0.8908 | 2.22E-07 |
| ENSG00000245648 | NA | novel transcript, antisense to KLRK1 | 1.5688 | 2.25E-07 |
| ENSG00000067606 | PRKCZ | protein kinase C zeta [Source:HGNC Symbol;Acc:HGNC:9412] | -0.6175 | 2.25E-07 |
| ENSG00000167965 | MLST8 | MTOR associated protein, LST8 homolog [Source:HGNC Symbol;Acc:HGNC:24825] | -0.5903 | 2.26E-07 |
| ENSG00000128829 | EIF2AK4 | eukaryotic translation initiation factor 2 alpha kinase 4 [Source:HGNC Symbol;Acc:HGNC:19687] | -0.5282 | 2.27E-07 |
| ENSG00000288586 | NA | novel transcript | 1.0148 | 2.27E-07 |
| ENSG00000064199 | SPA17 | sperm autoantigenic protein 17 [Source:HGNC Symbol;Acc:HGNC:11210] | -1.1130 | 2.30E-07 |
| ENSG00000113100 | CDH9 | cadherin 9 [Source:HGNC Symbol;Acc:HGNC:1768] | -1.2128 | 2.31E-07 |
| ENSG00000246465 | NA | novel transcript LOC100506705 | -1.0736 | 2.33E-07 |
| ENSG00000064652 | SNX24 | sorting nexin 24 [Source:HGNC Symbol;Acc:HGNC:21533] | -0.6752 | 2.36E-07 |
| ENSG00000138867 | GUCD1 | guanylyl cyclase domain containing 1 [Source:HGNC Symbol;Acc:HGNC:14237] | -0.5485 | 2.36E-07 |
| ENSG00000203859 | HSD3B2 | hydroxy-delta-5-steroid dehydrogenase, 3 beta- and steroid delta-isomerase 2 [Source:HGNC Symbol;Acc:HGNC:5218] | 2.6721 | 2.40E-07 |
| ENSG00000283633 | NA | novel transcript | 0.5002 | 2.47E-07 |
| ENSG00000104957 | CCDC130 | coiled-coil domain containing 130 [Source:HGNC Symbol;Acc:HGNC:28118] | 0.6130 | 2.47E-07 |
| ENSG00000110711 | AIP | aryl hydrocarbon receptor interacting protein [Source:HGNC Symbol;Acc:HGNC:358] | 0.5669 | 2.49E-07 |
| ENSG00000154839 | SKA1 | spindle and kinetochore associated complex subunit 1 [Source:HGNC Symbol;Acc:HGNC:28109] | -0.9816 | 2.53E-07 |
| ENSG00000136603 | SKIL | SKI like proto-oncogene [Source:HGNC Symbol;Acc:HGNC:10897] | 0.9495 | 2.53E-07 |
| ENSG00000230487 | PSMG3-AS1 | PSMG3 antisense RNA 1 (head to head) [Source:HGNC Symbol;Acc:HGNC:22230] | -1.1752 | 2.55E-07 |
| ENSG00000139163 | ETNK1 | ethanolamine kinase 1 [Source:HGNC Symbol;Acc:HGNC:24649] | 0.5194 | 2.58E-07 |
| ENSG00000186951 | PPARA | peroxisome proliferator activated receptor alpha [Source:HGNC Symbol;Acc:HGNC:9232] | -0.8258 | 2.61E-07 |
| ENSG00000141337 | ARSG | arylsulfatase G [Source:HGNC Symbol;Acc:HGNC:24102] | 1.0913 | 2.61E-07 |
| ENSG00000185436 | IFNLR1 | interferon lambda receptor 1 [Source:HGNC Symbol;Acc:HGNC:18584] | -0.5617 | 2.61E-07 |
| ENSG00000259235 | NA | novel transcript | -1.3646 | 2.62E-07 |
| ENSG00000244731 | C4A | complement C4A (Rodgers blood group) [Source:HGNC Symbol;Acc:HGNC:1323] | 1.1898 | 2.64E-07 |
| ENSG00000100234 | TIMP3 | TIMP metallopeptidase inhibitor 3 [Source:HGNC Symbol;Acc:HGNC:11822] | 0.6557 | 2.64E-07 |
| ENSG00000163738 | MTHFD2L | methylenetetrahydrofolate dehydrogenase (NADP+ dependent) 2 like [Source:HGNC Symbol;Acc:HGNC:31865] | 0.5683 | 2.65E-07 |
| ENSG00000247317 | LY6E-DT | LY6E divergent transcript [Source:HGNC Symbol;Acc:HGNC:53728] | 1.7979 | 2.73E-07 |
| ENSG00000144596 | GRIP2 | glutamate receptor interacting protein 2 [Source:HGNC Symbol;Acc:HGNC:23841] | -2.7665 | 2.79E-07 |
| ENSG00000280187 | NA | TEC | -0.7228 | 2.81E-07 |
| ENSG00000120925 | RNF170 | ring finger protein 170 [Source:HGNC Symbol;Acc:HGNC:25358] | -0.6549 | 2.83E-07 |

| | | | | |
|---|---|---|---|---|
| ENSG00000165409 | TSHR | thyroid stimulating hormone receptor [Source:HGNC Symbol;Acc:HGNC:12373] | 1.7482 | 2.84E-07 |
| ENSG00000150433 | TMEM218 | transmembrane protein 218 [Source:HGNC Symbol;Acc:HGNC:27344] | -1.1236 | 2.87E-07 |
| ENSG00000243725 | TTC4 | tetratricopeptide repeat domain 4 [Source:HGNC Symbol;Acc:HGNC:12394] | 0.5853 | 2.94E-07 |
| ENSG00000104267 | CA2 | carbonic anhydrase 2 [Source:HGNC Symbol;Acc:HGNC:1373] | 1.2841 | 2.95E-07 |
| ENSG00000173545 | ZNF622 | zinc finger protein 622 [Source:HGNC Symbol;Acc:HGNC:30958] | 0.5968 | 2.95E-07 |
| ENSG00000163171 | CDC42EP3 | CDC42 effector protein 3 [Source:HGNC Symbol;Acc:HGNC:16943] | 1.9274 | 2.96E-07 |
| ENSG00000006638 | TBXA2R | thromboxane A2 receptor [Source:HGNC Symbol;Acc:HGNC:11608] | -1.3137 | 3.02E-07 |
| ENSG00000099625 | CBARP | CACN subunit beta associated regulatory protein [Source:HGNC Symbol;Acc:HGNC:28617] | 1.7613 | 3.04E-07 |
| ENSG00000119333 | WDR34 | WD repeat domain 34 [Source:HGNC Symbol;Acc:HGNC:28296] | -0.7004 | 3.04E-07 |
| ENSG00000231721 | LINC-PINT | long intergenic non-protein coding RNA, p53 induced transcript [Source:HGNC Symbol;Acc:HGNC:26885] | 0.8036 | 3.05E-07 |
| ENSG00000255622 | PCDHB17P | protocadherin beta 17 pseudogene [Source:HGNC Symbol;Acc:HGNC:14547] | -0.8363 | 3.05E-07 |
| ENSG00000168685 | IL7R | interleukin 7 receptor [Source:HGNC Symbol;Acc:HGNC:6024] | 5.2958 | 3.07E-07 |
| ENSG00000100304 | TTLL12 | tubulin tyrosine ligase like 12 [Source:HGNC Symbol;Acc:HGNC:28974] | 0.5469 | 3.07E-07 |
| ENSG00000105605 | CACNG7 | calcium voltage-gated channel auxiliary subunit gamma 7 [Source:HGNC Symbol;Acc:HGNC:13626] | -0.8421 | 3.08E-07 |
| ENSG00000107882 | SUFU | SUFU negative regulator of hedgehog signaling [Source:HGNC Symbol;Acc:HGNC:16466] | -0.6715 | 3.10E-07 |
| ENSG00000279905 | NA | novel transcript | 2.5896 | 3.11E-07 |
| ENSG00000129566 | TEP1 | telomerase associated protein 1 [Source:HGNC Symbol;Acc:HGNC:11726] | 1.0510 | 3.26E-07 |
| ENSG00000140443 | IGF1R | insulin like growth factor 1 receptor [Source:HGNC Symbol;Acc:HGNC:5465] | 0.8142 | 3.29E-07 |
| ENSG00000123908 | AGO2 | argonaute RISC catalytic component 2 [Source:HGNC Symbol;Acc:HGNC:3263] | 0.7247 | 3.30E-07 |
| ENSG00000114646 | CSPG5 | chondroitin sulfate proteoglycan 5 [Source:HGNC Symbol;Acc:HGNC:2467] | -1.5429 | 3.42E-07 |
| ENSG00000119523 | ALG2 | ALG2 alpha-1,3/1,6-mannosyltransferase [Source:HGNC Symbol;Acc:HGNC:23159] | 0.5913 | 3.46E-07 |
| ENSG00000065029 | ZNF76 | zinc finger protein 76 [Source:HGNC Symbol;Acc:HGNC:13149] | -0.6048 | 3.60E-07 |
| ENSG00000257434 | NA | novel transcript, antisense to KCNC2 | -4.3272 | 3.60E-07 |
| ENSG00000049246 | PER3 | period circadian regulator 3 [Source:HGNC Symbol;Acc:HGNC:8847] | -1.3917 | 3.61E-07 |
| ENSG00000087842 | PIR | pirin [Source:HGNC Symbol;Acc:HGNC:30048] | -0.5389 | 3.63E-07 |
| ENSG00000101224 | CDC25B | cell division cycle 25B [Source:HGNC Symbol;Acc:HGNC:1726] | -0.5905 | 3.69E-07 |
| ENSG00000079950 | STX7 | syntaxin 7 [Source:HGNC Symbol;Acc:HGNC:11442] | -0.7652 | 3.69E-07 |
| ENSG00000101189 | MRGBP | MRG domain binding protein [Source:HGNC Symbol;Acc:HGNC:15866] | -0.7465 | 3.69E-07 |
| ENSG00000147647 | DPYS | dihydropyrimidinase [Source:HGNC Symbol;Acc:HGNC:3013] | 1.0598 | 3.73E-07 |
| ENSG00000149557 | FEZ1 | fasciculation and elongation protein zeta 1 [Source:HGNC Symbol;Acc:HGNC:3659] | -0.9645 | 3.78E-07 |
| ENSG00000187122 | SLIT1 | slit guidance ligand 1 [Source:HGNC Symbol;Acc:HGNC:11085] | -1.5006 | 3.80E-07 |
| ENSG00000131094 | C1QL1 | complement C1q like 1 [Source:HGNC Symbol;Acc:HGNC:24182] | -1.1233 | 3.80E-07 |
| ENSG00000136237 | RAPGEF5 | Rap guanine nucleotide exchange factor 5 [Source:HGNC Symbol;Acc:HGNC:16862] | 1.4418 | 3.87E-07 |
| ENSG00000115091 | ACTR3 | actin related protein 3 [Source:HGNC Symbol;Acc:HGNC:170] | 0.5204 | 3.88E-07 |
| ENSG00000196975 | ANXA4 | annexin A4 [Source:HGNC Symbol;Acc:HGNC:542] | -0.7220 | 3.88E-07 |
| ENSG00000177200 | CHD9 | chromodomain helicase DNA binding protein 9 [Source:HGNC Symbol;Acc:HGNC:25701] | 0.5464 | 3.88E-07 |
| ENSG00000175445 | LPL | lipoprotein lipase [Source:HGNC Symbol;Acc:HGNC:6677] | -1.2291 | 3.89E-07 |
| ENSG00000130939 | UBE4B | ubiquitination factor E4B [Source:HGNC Symbol;Acc:HGNC:12500] | -0.6117 | 3.90E-07 |
| ENSG00000101935 | AMMECR1 | AMMECR nuclear protein 1 [Source:HGNC Symbol;Acc:HGNC:467] | 0.6594 | 3.91E-07 |
| ENSG00000148541 | FAM13C | family with sequence similarity 13 member C [Source:HGNC Symbol;Acc:HGNC:19371] | -0.6919 | 3.94E-07 |
| ENSG00000140511 | HAPLN3 | hyaluronan and proteoglycan link protein 3 [Source:HGNC Symbol;Acc:HGNC:21446] | 1.2925 | 3.97E-07 |
| ENSG00000004777 | ARHGAP33 | Rho GTPase activating protein 33 [Source:HGNC Symbol;Acc:HGNC:23085] | -1.0171 | 3.99E-07 |
| ENSG00000165355 | FBXO33 | F-box protein 33 [Source:HGNC Symbol;Acc:HGNC:19833] | 0.5294 | 4.16E-07 |

| | | | | |
|---|---|---|---|---|
| ENSG00000177426 | TGIF1 | TGFB induced factor homeobox 1 [Source:HGNC Symbol;Acc:HGNC:11776] | 0.7712 | 4.18E-07 |
| ENSG00000100324 | TAB1 | TGF-beta activated kinase 1 (MAP3K7) binding protein 1 [Source:HGNC Symbol;Acc:HGNC:18157] | -0.6910 | 4.20E-07 |
| ENSG00000119900 | OGFRL1 | opioid growth factor receptor like 1 [Source:HGNC Symbol;Acc:HGNC:21378] | -0.9041 | 4.21E-07 |
| ENSG00000170345 | FOS | Fos proto-oncogene, AP-1 transcription factor subunit [Source:HGNC Symbol;Acc:HGNC:3796] | 4.3673 | 4.23E-07 |
| ENSG00000160505 | NLRP4 | NLR family pyrin domain containing 4 [Source:HGNC Symbol;Acc:HGNC:22943] | -1.6925 | 4.24E-07 |
| ENSG00000158164 | TMSB15A | thymosin beta 15a [Source:HGNC Symbol;Acc:HGNC:30744] | -0.5640 | 4.25E-07 |
| ENSG00000180815 | MAP3K15 | mitogen-activated protein kinase kinase kinase 15 [Source:HGNC Symbol;Acc:HGNC:31689] | -1.2250 | 4.25E-07 |
| ENSG00000100664 | EIF5 | eukaryotic translation initiation factor 5 [Source:HGNC Symbol;Acc:HGNC:3299] | 0.8702 | 4.32E-07 |
| ENSG00000127124 | HIVEP3 | HIVEP zinc finger 3 [Source:HGNC Symbol;Acc:HGNC:13561] | 1.4971 | 4.35E-07 |
| ENSG00000157911 | PEX10 | peroxisomal biogenesis factor 10 [Source:HGNC Symbol;Acc:HGNC:8851] | -0.7822 | 4.37E-07 |
| ENSG00000141376 | BCAS3 | BCAS3 microtubule associated cell migration factor [Source:HGNC Symbol;Acc:HGNC:14347] | -1.2376 | 4.44E-07 |
| ENSG00000110925 | CSRNP2 | cysteine and serine rich nuclear protein 2 [Source:HGNC Symbol;Acc:HGNC:16006] | 0.6076 | 4.49E-07 |
| ENSG00000128699 | ORMDL1 | ORMDL sphingolipid biosynthesis regulator 1 [Source:HGNC Symbol;Acc:HGNC:16036] | 0.5338 | 4.53E-07 |
| ENSG00000115257 | PCSK4 | proprotein convertase subtilisin/kexin type 4 [Source:HGNC Symbol;Acc:HGNC:8746] | -1.2114 | 4.57E-07 |
| ENSG00000196628 | TCF4 | transcription factor 4 [Source:HGNC Symbol;Acc:HGNC:11634] | 0.6083 | 4.58E-07 |
| ENSG00000163874 | ZC3H12A | zinc finger CCCH-type containing 12A [Source:HGNC Symbol;Acc:HGNC:26259] | 1.2832 | 4.59E-07 |
| ENSG00000100036 | SLC35E4 | solute carrier family 35 member E4 [Source:HGNC Symbol;Acc:HGNC:17058] | 1.0833 | 4.62E-07 |
| ENSG00000105639 | JAK3 | Janus kinase 3 [Source:HGNC Symbol;Acc:HGNC:6193] | -1.5738 | 4.70E-07 |
| ENSG00000232533 | NA | novel transcript | 1.6632 | 4.77E-07 |
| ENSG00000205213 | LGR4 | leucine rich repeat containing G protein-coupled receptor 4 [Source:HGNC Symbol;Acc:HGNC:13299] | -0.7279 | 4.79E-07 |
| ENSG00000130988 | RGN | regucalcin [Source:HGNC Symbol;Acc:HGNC:9989] | -0.7635 | 4.79E-07 |
| ENSG00000167207 | NOD2 | nucleotide binding oligomerization domain containing 2 [Source:HGNC Symbol;Acc:HGNC:5331] | -0.9432 | 4.80E-07 |
| ENSG00000137522 | RNF121 | ring finger protein 121 [Source:HGNC Symbol;Acc:HGNC:21070] | 0.5289 | 4.90E-07 |
| ENSG00000228725 | MTND2P12 | MT-ND2 pseudogene 12 [Source:HGNC Symbol;Acc:HGNC:42113] | 2.4331 | 5.00E-07 |
| ENSG00000273117 | NA | novel transcript | -1.3660 | 5.05E-07 |
| ENSG00000273038 | NA | novel transcript | 1.4750 | 5.06E-07 |
| ENSG00000117602 | RCAN3 | RCAN family member 3 [Source:HGNC Symbol;Acc:HGNC:3042] | -0.8727 | 5.17E-07 |
| ENSG00000260941 | LINC00622 | long intergenic non-protein coding RNA 622 [Source:HGNC Symbol;Acc:HGNC:44251] | 3.6827 | 5.18E-07 |
| ENSG00000228742 | LINC02577 | long intergenic non-protein coding RNA 2577 [Source:HGNC Symbol;Acc:HGNC:53749] | 2.3905 | 5.25E-07 |
| ENSG00000196850 | PPTC7 | protein phosphatase targeting COQ7 [Source:HGNC Symbol;Acc:HGNC:30695] | 0.5040 | 5.25E-07 |
| ENSG00000096872 | IFT74 | intraflagellar transport 74 [Source:HGNC Symbol;Acc:HGNC:21424] | -0.7260 | 5.26E-07 |
| ENSG00000104093 | DMXL2 | Dmx like 2 [Source:HGNC Symbol;Acc:HGNC:2938] | -0.6247 | 5.28E-07 |
| ENSG00000109911 | ELP4 | elongator acetyltransferase complex subunit 4 [Source:HGNC Symbol;Acc:HGNC:1171] | -0.8426 | 5.28E-07 |
| ENSG00000149289 | ZC3H12C | zinc finger CCCH-type containing 12C [Source:HGNC Symbol;Acc:HGNC:29362] | 0.5961 | 5.33E-07 |
| ENSG00000251369 | ZNF550 | zinc finger protein 550 [Source:HGNC Symbol;Acc:HGNC:28643] | -0.6075 | 5.33E-07 |
| ENSG00000139567 | ACVRL1 | activin A receptor like type 1 [Source:HGNC Symbol;Acc:HGNC:175] | -2.2951 | 5.38E-07 |
| ENSG00000186352 | ANKRD37 | ankyrin repeat domain 37 [Source:HGNC Symbol;Acc:HGNC:29593] | 2.1499 | 5.42E-07 |
| ENSG00000075651 | PLD1 | phospholipase D1 [Source:HGNC Symbol;Acc:HGNC:9067] | -0.8544 | 5.45E-07 |
| ENSG00000110693 | SOX6 | SRY-box transcription factor 6 [Source:HGNC Symbol;Acc:HGNC:16421] | -3.8250 | 5.50E-07 |
| ENSG00000196684 | HSH2D | hematopoietic SH2 domain containing [Source:HGNC Symbol;Acc:HGNC:24920] | 0.8236 | 5.57E-07 |
| ENSG00000185585 | OLFML2A | olfactomedin like 2A [Source:HGNC Symbol;Acc:HGNC:27270] | -1.5405 | 5.64E-07 |
| ENSG00000105063 | PPP6R1 | protein phosphatase 6 regulatory subunit 1 [Source:HGNC Symbol;Acc:HGNC:29195] | 0.5037 | 6.00E-07 |
| ENSG00000204371 | EHMT2 | euchromatic histone lysine methyltransferase 2 [Source:HGNC Symbol;Acc:HGNC:14129] | -0.7007 | 6.27E-07 |

| | | | | |
|---|---|---|---|---|
| ENSG00000161642 | ZNF385A | zinc finger protein 385A [Source:HGNC Symbol;Acc:HGNC:17521] | -0.6666 | 6.31E-07 |
| ENSG00000017260 | ATP2C1 | ATPase secretory pathway Ca2+ transporting 1 [Source:HGNC Symbol;Acc:HGNC:13211] | -0.5276 | 6.41E-07 |
| ENSG00000184675 | AMER1 | APC membrane recruitment protein 1 [Source:HGNC Symbol;Acc:HGNC:26837] | -0.6589 | 6.46E-07 |
| ENSG00000013375 | PGM3 | phosphoglucomutase 3 [Source:HGNC Symbol;Acc:HGNC:8907] | 0.9032 | 6.47E-07 |
| ENSG00000158321 | AUTS2 | activator of transcription and developmental regulator AUTS2 [Source:HGNC Symbol;Acc:HGNC:14262] | -0.9679 | 6.50E-07 |
| ENSG00000182093 | GET1 | guided entry of tail-anchored proteins factor 1 [Source:HGNC Symbol;Acc:HGNC:12790] | -0.5276 | 6.57E-07 |
| ENSG00000176909 | MAMSTR | MEF2 activating motif and SAP domain containing transcriptional regulator [Source:HGNC Symbol;Acc:HGNC:26689] | -0.6244 | 6.64E-07 |
| ENSG00000108924 | HLF | HLF transcription factor, PAR bZIP family member [Source:HGNC Symbol;Acc:HGNC:4977] | -0.9776 | 6.66E-07 |
| ENSG00000166900 | STX3 | syntaxin 3 [Source:HGNC Symbol;Acc:HGNC:11438] | 0.5621 | 6.73E-07 |
| ENSG00000125247 | TMTC4 | transmembrane O-mannosyltransferase targeting cadherins 4 [Source:HGNC Symbol;Acc:HGNC:25904] | -1.4121 | 6.81E-07 |
| ENSG00000123384 | LRP1 | LDL receptor related protein 1 [Source:HGNC Symbol;Acc:HGNC:6692] | -0.7906 | 6.91E-07 |
| ENSG00000279940 | NA | novel transcript | 3.3030 | 7.01E-07 |
| ENSG00000197479 | PCDHB11 | protocadherin beta 11 [Source:HGNC Symbol;Acc:HGNC:8682] | -1.0977 | 7.04E-07 |
| ENSG00000144711 | IQSEC1 | IQ motif and Sec7 domain ArfGEF 1 [Source:HGNC Symbol;Acc:HGNC:29112] | -0.5283 | 7.04E-07 |
| ENSG00000197363 | ZNF517 | zinc finger protein 517 [Source:HGNC Symbol;Acc:HGNC:27984] | -1.1800 | 7.04E-07 |
| ENSG00000240990 | HOXA11-AS | HOXA11 antisense RNA [Source:HGNC Symbol;Acc:HGNC:24957] | 0.5713 | 7.11E-07 |
| ENSG00000109079 | TNFAIP1 | TNF alpha induced protein 1 [Source:HGNC Symbol;Acc:HGNC:11894] | 0.6123 | 7.12E-07 |
| ENSG00000246627 | CACNA1C-AS1 | CACNA1C antisense RNA 1 [Source:HGNC Symbol;Acc:HGNC:40119] | 2.7553 | 7.12E-07 |
| ENSG00000242082 | SLC5A4-AS1 | SLC5A4 antisense RNA 1 [Source:HGNC Symbol;Acc:HGNC:53163] | -1.5431 | 7.16E-07 |
| ENSG00000149596 | JPH2 | junctophilin 2 [Source:HGNC Symbol;Acc:HGNC:14202] | -0.8044 | 7.23E-07 |
| ENSG00000157240 | FZD1 | frizzled class receptor 1 [Source:HGNC Symbol;Acc:HGNC:4038] | -1.4023 | 7.29E-07 |
| ENSG00000166974 | MAPRE2 | microtubule associated protein RP/EB family member 2 [Source:HGNC Symbol;Acc:HGNC:6891] | 0.6222 | 7.29E-07 |
| ENSG00000233435 | AGGF1P2 | angiogenic factor with G-patch and FHA domains 1 pseudogene 2 [Source:HGNC Symbol;Acc:HGNC:38052] | -0.9711 | 7.29E-07 |
| ENSG00000164338 | UTP15 | UTP15 small subunit processome component [Source:HGNC Symbol;Acc:HGNC:25758] | 0.8338 | 7.32E-07 |
| ENSG00000114450 | GNB4 | G protein subunit beta 4 [Source:HGNC Symbol;Acc:HGNC:20731] | -0.9787 | 7.33E-07 |
| ENSG00000196704 | AMZ2 | archaelysin family metallopeptidase 2 [Source:HGNC Symbol;Acc:HGNC:28041] | -0.5929 | 7.34E-07 |
| ENSG00000163214 | DHX57 | DExH-box helicase 57 [Source:HGNC Symbol;Acc:HGNC:20086] | 0.7941 | 7.35E-07 |
| ENSG00000089195 | TRMT6 | tRNA methyltransferase 6 [Source:HGNC Symbol;Acc:HGNC:20900] | 0.5877 | 7.38E-07 |
| ENSG00000105976 | MET | MET proto-oncogene, receptor tyrosine kinase [Source:HGNC Symbol;Acc:HGNC:7029] | -0.6263 | 7.59E-07 |
| ENSG00000173262 | SLC2A14 | solute carrier family 2 member 14 [Source:HGNC Symbol;Acc:HGNC:18301] | 0.6876 | 7.59E-07 |
| ENSG00000167272 | POP5 | POP5 homolog, ribonuclease P/MRP subunit [Source:HGNC Symbol;Acc:HGNC:17689] | -0.5931 | 7.70E-07 |
| ENSG00000166816 | LDHD | lactate dehydrogenase D [Source:HGNC Symbol;Acc:HGNC:19708] | -0.9700 | 7.72E-07 |
| ENSG00000139112 | GABARAPL1 | GABA type A receptor associated protein like 1 [Source:HGNC Symbol;Acc:HGNC:4068] | 0.7277 | 7.96E-07 |
| ENSG00000214367 | HAUS3 | HAUS augmin like complex subunit 3 [Source:HGNC Symbol;Acc:HGNC:28719] | -0.5020 | 8.03E-07 |
| ENSG00000113504 | SLC12A7 | solute carrier family 12 member 7 [Source:HGNC Symbol;Acc:HGNC:10915] | 0.6447 | 8.03E-07 |
| ENSG00000108018 | SORCS1 | sortilin related VPS10 domain containing receptor 1 [Source:HGNC Symbol;Acc:HGNC:16697] | 1.0908 | 8.20E-07 |
| ENSG00000083454 | P2RX5 | purinergic receptor P2X 5 [Source:HGNC Symbol;Acc:HGNC:8536] | 0.7311 | 8.44E-07 |
| ENSG00000180998 | GPR137C | G protein-coupled receptor 137C [Source:HGNC Symbol;Acc:HGNC:25445] | 1.4791 | 8.46E-07 |
| ENSG00000272163 | NA | novel transcript, antisense to NEFM | -1.7000 | 8.46E-07 |
| ENSG00000182944 | EWSR1 | EWS RNA binding protein 1 [Source:HGNC Symbol;Acc:HGNC:3508] | 0.5367 | 8.56E-07 |
| ENSG00000236501 | NA | novel transcript | 1.2665 | 8.63E-07 |
| ENSG00000185697 | MYBL1 | MYB proto-oncogene like 1 [Source:HGNC Symbol;Acc:HGNC:7547] | -0.7274 | 8.73E-07 |

| | | | | |
|---|---|---|---|---|
| ENSG00000174939 | ASPHD1 | aspartate beta-hydroxylase domain containing 1 [Source:HGNC Symbol;Acc:HGNC:27380] | 0.6938 | 8.76E-07 |
| ENSG00000164841 | TMEM74 | transmembrane protein 74 [Source:HGNC Symbol;Acc:HGNC:26409] | -1.1530 | 8.83E-07 |
| ENSG00000160606 | TLCD1 | TLC domain containing 1 [Source:HGNC Symbol;Acc:HGNC:25177] | -1.3860 | 8.84E-07 |
| ENSG00000158169 | FANCC | FA complementation group C [Source:HGNC Symbol;Acc:HGNC:3584] | -0.5782 | 8.87E-07 |
| ENSG00000153558 | FBXL2 | F-box and leucine rich repeat protein 2 [Source:HGNC Symbol;Acc:HGNC:13598] | -0.7882 | 8.89E-07 |
| ENSG00000152556 | PFKM | phosphofructokinase, muscle [Source:HGNC Symbol;Acc:HGNC:8877] | -0.5319 | 8.90E-07 |
| ENSG00000106100 | NOD1 | nucleotide binding oligomerization domain containing 1 [Source:HGNC Symbol;Acc:HGNC:16390] | -1.6470 | 8.99E-07 |
| ENSG00000117868 | ESYT2 | extended synaptotagmin 2 [Source:HGNC Symbol;Acc:HGNC:22211] | 0.6539 | 9.12E-07 |
| ENSG00000185127 | C6orf120 | chromosome 6 open reading frame 120 [Source:HGNC Symbol;Acc:HGNC:21247] | 0.5021 | 9.13E-07 |
| ENSG00000079277 | MKNK1 | MAPK interacting serine/threonine kinase 1 [Source:HGNC Symbol;Acc:HGNC:7110] | -0.6055 | 9.19E-07 |
| ENSG00000242247 | ARFGAP3 | ADP ribosylation factor GTPase activating protein 3 [Source:HGNC Symbol;Acc:HGNC:661] | 0.6382 | 9.24E-07 |
| ENSG00000108379 | WNT3 | Wnt family member 3 [Source:HGNC Symbol;Acc:HGNC:12782] | 0.6339 | 9.25E-07 |
| ENSG00000060642 | PIGV | phosphatidylinositol glycan anchor biosynthesis class V [Source:HGNC Symbol;Acc:HGNC:26031] | -0.8750 | 9.39E-07 |
| ENSG00000183837 | PNMA3 | PNMA family member 3 [Source:HGNC Symbol;Acc:HGNC:18742] | -1.6718 | 9.49E-07 |
| ENSG00000198715 | GLMP | glycosylated lysosomal membrane protein [Source:HGNC Symbol;Acc:HGNC:29436] | -0.5423 | 9.55E-07 |
| ENSG00000181830 | SLC35C1 | solute carrier family 35 member C1 [Source:HGNC Symbol;Acc:HGNC:20197] | 0.6564 | 9.57E-07 |
| ENSG00000120322 | PCDHB8 | protocadherin beta 8 [Source:HGNC Symbol;Acc:HGNC:8693] | -0.6896 | 9.58E-07 |
| ENSG00000163322 | ABRAXAS1 | abraxas 1, BRCA1 A complex subunit [Source:HGNC Symbol;Acc:HGNC:25829] | -0.7847 | 9.67E-07 |
| ENSG00000081320 | STK17B | serine/threonine kinase 17b [Source:HGNC Symbol;Acc:HGNC:11396] | -0.5796 | 9.71E-07 |
| ENSG00000067836 | ROGDI | rogdi atypical leucine zipper [Source:HGNC Symbol;Acc:HGNC:29478] | -0.6968 | 9.98E-07 |
| ENSG00000179630 | LACC1 | laccase domain containing 1 [Source:HGNC Symbol;Acc:HGNC:26789] | -1.6847 | 1.01E-06 |
| ENSG00000267221 | C17orf113 | chromosome 17 open reading frame 113 [Source:HGNC Symbol;Acc:HGNC:53437] | -1.7873 | 1.01E-06 |
| ENSG00000164934 | DCAF13 | DDB1 and CUL4 associated factor 13 [Source:HGNC Symbol;Acc:HGNC:24535] | 0.5529 | 1.02E-06 |
| ENSG00000065882 | TBC1D1 | TBC1 domain family member 1 [Source:HGNC Symbol;Acc:HGNC:11578] | -0.6524 | 1.04E-06 |
| ENSG00000154310 | TNIK | TRAF2 and NCK interacting kinase [Source:HGNC Symbol;Acc:HGNC:30765] | 1.0079 | 1.05E-06 |
| ENSG00000164532 | TBX20 | T-box transcription factor 20 [Source:HGNC Symbol;Acc:HGNC:11598] | 1.9703 | 1.05E-06 |
| ENSG00000135077 | HAVCR2 | hepatitis A virus cellular receptor 2 [Source:HGNC Symbol;Acc:HGNC:18437] | 1.6718 | 1.05E-06 |
| ENSG00000008735 | MAPK8IP2 | mitogen-activated protein kinase 8 interacting protein 2 [Source:HGNC Symbol;Acc:HGNC:6883] | -0.8130 | 1.06E-06 |
| ENSG00000223711 | NA | novel transcript | 1.7403 | 1.06E-06 |
| ENSG00000125652 | ALKBH7 | alkB homolog 7 [Source:HGNC Symbol;Acc:HGNC:21306] | -0.6250 | 1.07E-06 |
| ENSG00000132199 | ENOSF1 | enolase superfamily member 1 [Source:HGNC Symbol;Acc:HGNC:30365] | -1.0140 | 1.07E-06 |
| ENSG00000112234 | FBXL4 | F-box and leucine rich repeat protein 4 [Source:HGNC Symbol;Acc:HGNC:13601] | -0.6842 | 1.09E-06 |
| ENSG00000134285 | FKBP11 | FKBP prolyl isomerase 11 [Source:HGNC Symbol;Acc:HGNC:18624] | 0.9420 | 1.10E-06 |
| ENSG00000196586 | MYO6 | myosin VI [Source:HGNC Symbol;Acc:HGNC:7605] | 0.7061 | 1.10E-06 |
| ENSG00000163285 | GABRG1 | gamma-aminobutyric acid type A receptor subunit gamma1 [Source:HGNC Symbol;Acc:HGNC:4086] | -2.0830 | 1.11E-06 |
| ENSG00000273044 | NA | novel transcript | 2.0504 | 1.11E-06 |
| ENSG00000226306 | NPY6R | neuropeptide Y receptor Y6 (pseudogene) [Source:HGNC Symbol;Acc:HGNC:7959] | 0.5622 | 1.12E-06 |
| ENSG00000176834 | VSIG10 | V-set and immunoglobulin domain containing 10 [Source:HGNC Symbol;Acc:HGNC:26078] | -0.6470 | 1.14E-06 |
| ENSG00000198042 | MAK16 | MAK16 homolog [Source:HGNC Symbol;Acc:HGNC:13703] | 0.5825 | 1.14E-06 |
| ENSG00000144476 | ACKR3 | atypical chemokine receptor 3 [Source:HGNC Symbol;Acc:HGNC:23692] | 1.0862 | 1.14E-06 |
| ENSG00000272143 | FGF14-AS2 | FGF14 antisense RNA 2 [Source:HGNC Symbol;Acc:HGNC:44368] | -1.2679 | 1.14E-06 |
| ENSG00000183305 | MAGEA2B | MAGE family member A2B [Source:HGNC Symbol;Acc:HGNC:19340] | 0.5441 | 1.15E-06 |
| ENSG00000102897 | LYRM1 | LYR motif containing 1 [Source:HGNC Symbol;Acc:HGNC:25074] | 0.5310 | 1.17E-06 |

| | | | | |
|---|---|---|---|---|
| ENSG00000100307 | CBX7 | chromobox 7 [Source:HGNC Symbol;Acc:HGNC:1557] | -0.8976 | 1.18E-06 |
| ENSG00000175197 | DDIT3 | DNA damage inducible transcript 3 [Source:HGNC Symbol;Acc:HGNC:2726] | 0.7140 | 1.18E-06 |
| ENSG00000197208 | SLC22A4 | solute carrier family 22 member 4 [Source:HGNC Symbol;Acc:HGNC:10968] | 1.7355 | 1.18E-06 |
| ENSG00000108813 | DLX4 | distal-less homeobox 4 [Source:HGNC Symbol;Acc:HGNC:2917] | -0.9489 | 1.19E-06 |
| ENSG00000128694 | OSGEPL1 | O-sialoglycoprotein endopeptidase like 1 [Source:HGNC Symbol;Acc:HGNC:23075] | -0.8367 | 1.19E-06 |
| ENSG00000163694 | RBM47 | RNA binding motif protein 47 [Source:HGNC Symbol;Acc:HGNC:30358] | 0.5228 | 1.20E-06 |
| ENSG00000111665 | CDCA3 | cell division cycle associated 3 [Source:HGNC Symbol;Acc:HGNC:14624] | -0.8439 | 1.22E-06 |
| ENSG00000157212 | PAXIP1 | PAX interacting protein 1 [Source:HGNC Symbol;Acc:HGNC:8624] | -0.5830 | 1.22E-06 |
| ENSG00000258947 | TUBB3 | tubulin beta 3 class III [Source:HGNC Symbol;Acc:HGNC:20772] | 0.9223 | 1.23E-06 |
| ENSG00000102401 | ARMCX3 | armadillo repeat containing X-linked 3 [Source:HGNC Symbol;Acc:HGNC:24065] | 0.5620 | 1.25E-06 |
| ENSG00000251201 | TMED7-TICAM2 | TMED7-TICAM2 readthrough [Source:HGNC Symbol;Acc:HGNC:33945] | 0.9903 | 1.27E-06 |
| ENSG00000172301 | COPRS | coordinator of PRMT5 and differentiation stimulator [Source:HGNC Symbol;Acc:HGNC:28848] | 0.5414 | 1.28E-06 |
| ENSG00000057704 | TMCC3 | transmembrane and coiled-coil domain family 3 [Source:HGNC Symbol;Acc:HGNC:29199] | -1.6040 | 1.29E-06 |
| ENSG00000233452 | STXBP5-AS1 | STXBP5 antisense RNA 1 [Source:HGNC Symbol;Acc:HGNC:44183] | 1.8550 | 1.29E-06 |
| ENSG00000131080 | EDA2R | ectodysplasin A2 receptor [Source:HGNC Symbol;Acc:HGNC:17756] | -0.8306 | 1.30E-06 |
| ENSG00000185739 | SRL | sarcalumenin [Source:HGNC Symbol;Acc:HGNC:11295] | 3.8150 | 1.30E-06 |
| ENSG00000104660 | LEPROTL1 | leptin receptor overlapping transcript like 1 [Source:HGNC Symbol;Acc:HGNC:6555] | 0.6734 | 1.30E-06 |
| ENSG00000275325 | PDCD6IPP1 | PDCD6IP pseudogene 1 [Source:HGNC Symbol;Acc:HGNC:49872] | -0.7224 | 1.31E-06 |
| ENSG00000164221 | CCDC112 | coiled-coil domain containing 112 [Source:HGNC Symbol;Acc:HGNC:28599] | -0.7128 | 1.33E-06 |
| ENSG00000181444 | ZNF467 | zinc finger protein 467 [Source:HGNC Symbol;Acc:HGNC:23154] | -1.2351 | 1.33E-06 |
| ENSG00000274317 | LINC02334 | long intergenic non-protein coding RNA 2334 [Source:HGNC Symbol;Acc:HGNC:53254] | 2.2606 | 1.35E-06 |
| ENSG00000006652 | IFRD1 | interferon related developmental regulator 1 [Source:HGNC Symbol;Acc:HGNC:5456] | 0.7204 | 1.35E-06 |
| ENSG00000175161 | CADM2 | cell adhesion molecule 2 [Source:HGNC Symbol;Acc:HGNC:29849] | 0.9410 | 1.35E-06 |
| ENSG00000187244 | BCAM | basal cell adhesion molecule (Lutheran blood group) [Source:HGNC Symbol;Acc:HGNC:6722] | -0.6929 | 1.35E-06 |
| ENSG00000182489 | XKRX | XK related X-linked [Source:HGNC Symbol;Acc:HGNC:29845] | 1.3182 | 1.36E-06 |
| ENSG00000132600 | PRMT7 | protein arginine methyltransferase 7 [Source:HGNC Symbol;Acc:HGNC:25557] | -0.5675 | 1.38E-06 |
| ENSG00000198406 | BZW1P2 | basic leucine zipper and W2 domains 1 pseudogene 2 [Source:HGNC Symbol;Acc:HGNC:33954] | 0.8784 | 1.39E-06 |
| ENSG00000147082 | CCNB3 | cyclin B3 [Source:HGNC Symbol;Acc:HGNC:18709] | -0.6346 | 1.39E-06 |
| ENSG00000198026 | ZNF335 | zinc finger protein 335 [Source:HGNC Symbol;Acc:HGNC:15807] | 0.5568 | 1.43E-06 |
| ENSG00000108821 | COL1A1 | collagen type I alpha 1 chain [Source:HGNC Symbol;Acc:HGNC:2197] | -1.2986 | 1.43E-06 |
| ENSG00000197978 | GOLGA6L9 | golgin A6 family like 9 [Source:HGNC Symbol;Acc:HGNC:37229] | -0.9169 | 1.44E-06 |
| ENSG00000104856 | RELB | RELB proto-oncogene, NF-kB subunit [Source:HGNC Symbol;Acc:HGNC:9956] | 1.8837 | 1.44E-06 |
| ENSG00000171793 | CTPS1 | CTP synthase 1 [Source:HGNC Symbol;Acc:HGNC:2519] | 0.5009 | 1.45E-06 |
| ENSG00000152778 | IFIT5 | interferon induced protein with tetratricopeptide repeats 5 [Source:HGNC Symbol;Acc:HGNC:13328] | 0.6978 | 1.45E-06 |
| ENSG00000107443 | CCNJ | cyclin J [Source:HGNC Symbol;Acc:HGNC:23434] | 0.7234 | 1.47E-06 |
| ENSG00000197976 | AKAP17A | A-kinase anchoring protein 17A [Source:HGNC Symbol;Acc:HGNC:18783] | 0.5688 | 1.51E-06 |
| ENSG00000002079 | MYH16 | myosin heavy chain 16 pseudogene [Source:HGNC Symbol;Acc:HGNC:31038] | 5.1107 | 1.55E-06 |
| ENSG00000173905 | GOLIM4 | golgi integral membrane protein 4 [Source:HGNC Symbol;Acc:HGNC:15448] | 0.5367 | 1.56E-06 |
| ENSG00000221914 | PPP2R2A | protein phosphatase 2 regulatory subunit Balpha [Source:HGNC Symbol;Acc:HGNC:9304] | 0.5298 | 1.58E-06 |
| ENSG00000073111 | MCM2 | minichromosome maintenance complex component 2 [Source:HGNC Symbol;Acc:HGNC:6944] | -0.6971 | 1.58E-06 |
| ENSG00000081307 | UBA5 | ubiquitin like modifier activating enzyme 5 [Source:HGNC Symbol;Acc:HGNC:23230] | 0.5654 | 1.58E-06 |
| ENSG00000149927 | DOC2A | double C2 domain alpha [Source:HGNC Symbol;Acc:HGNC:2985] | -0.8760 | 1.59E-06 |
| ENSG00000141577 | CEP131 | centrosomal protein 131 [Source:HGNC Symbol;Acc:HGNC:29511] | -0.9378 | 1.59E-06 |

| | | | | |
|---|---|---|---|---|
| ENSG00000162729 | IGSF8 | immunoglobulin superfamily member 8 [Source:HGNC Symbol;Acc:HGNC:17813] | 0.5485 | 1.60E-06 |
| ENSG00000268941 | LINC01711 | long intergenic non-protein coding RNA 1711 [Source:HGNC Symbol;Acc:HGNC:28663] | 4.8749 | 1.60E-06 |
| ENSG00000110318 | CEP126 | centrosomal protein 126 [Source:HGNC Symbol;Acc:HGNC:29264] | -0.9655 | 1.61E-06 |
| ENSG00000164048 | ZNF589 | zinc finger protein 589 [Source:HGNC Symbol;Acc:HGNC:16747] | -0.9668 | 1.64E-06 |
| ENSG00000160753 | RUSC1 | RUN and SH3 domain containing 1 [Source:HGNC Symbol;Acc:HGNC:17153] | -0.5925 | 1.64E-06 |
| ENSG00000107404 | DVL1 | dishevelled segment polarity protein 1 [Source:HGNC Symbol;Acc:HGNC:3084] | -0.5429 | 1.65E-06 |
| ENSG00000176209 | SMIM19 | small integral membrane protein 19 [Source:HGNC Symbol;Acc:HGNC:25166] | -0.7049 | 1.68E-06 |
| ENSG00000072415 | MPP5 | membrane palmitoylated protein 5 [Source:HGNC Symbol;Acc:HGNC:18669] | -0.6250 | 1.69E-06 |
| ENSG00000109320 | NFKB1 | nuclear factor kappa B subunit 1 [Source:HGNC Symbol;Acc:HGNC:7794] | 0.5404 | 1.69E-06 |
| ENSG00000145283 | SLC10A6 | solute carrier family 10 member 6 [Source:HGNC Symbol;Acc:HGNC:30603] | 5.1925 | 1.71E-06 |
| ENSG00000089060 | SLC8B1 | solute carrier family 8 member B1 [Source:HGNC Symbol;Acc:HGNC:26175] | -0.5016 | 1.76E-06 |
| ENSG00000113594 | LIFR | LIF receptor subunit alpha [Source:HGNC Symbol;Acc:HGNC:6597] | -0.7527 | 1.77E-06 |
| ENSG00000144354 | CDCA7 | cell division cycle associated 7 [Source:HGNC Symbol;Acc:HGNC:14628] | -0.7635 | 1.79E-06 |
| ENSG00000236816 | ANKRD20A7P | ankyrin repeat domain 20 family member A7, pseudogene [Source:HGNC Symbol;Acc:HGNC:31980] | -0.5657 | 1.80E-06 |
| ENSG00000253868 | FER1L6-AS2 | FER1L6 antisense RNA 2 [Source:HGNC Symbol;Acc:HGNC:26534] | 0.7847 | 1.80E-06 |
| ENSG00000276122 | NA | novel transcript | 6.6992 | 1.80E-06 |
| ENSG00000157654 | PALM2AKAP2 | PALM2 and AKAP2 fusion [Source:HGNC Symbol;Acc:HGNC:33529] | 3.8897 | 1.81E-06 |
| ENSG00000183036 | PCP4 | Purkinje cell protein 4 [Source:HGNC Symbol;Acc:HGNC:8742] | 0.7944 | 1.81E-06 |
| ENSG00000196350 | ZNF729 | zinc finger protein 729 [Source:HGNC Symbol;Acc:HGNC:32464] | -1.2396 | 1.81E-06 |
| ENSG00000259673 | IQCH-AS1 | IQCH antisense RNA 1 [Source:HGNC Symbol;Acc:HGNC:44104] | -0.8002 | 1.81E-06 |
| ENSG00000187535 | IFT140 | intraflagellar transport 140 [Source:HGNC Symbol;Acc:HGNC:29077] | -0.5744 | 1.81E-06 |
| ENSG00000067113 | PLPP1 | phospholipid phosphatase 1 [Source:HGNC Symbol;Acc:HGNC:9228] | 0.6837 | 1.83E-06 |
| ENSG00000120262 | CCDC170 | coiled-coil domain containing 170 [Source:HGNC Symbol;Acc:HGNC:21177] | -2.1977 | 1.85E-06 |
| ENSG00000249740 | OSMR-AS1 | OSMR antisense RNA 1 (head to head) [Source:HGNC Symbol;Acc:HGNC:50296] | 1.5101 | 1.85E-06 |
| ENSG00000270069 | MIR222HG | miR222/221 cluster host gene [Source:HGNC Symbol;Acc:HGNC:49555] | 1.1297 | 1.87E-06 |
| ENSG00000119403 | PHF19 | PHD finger protein 19 [Source:HGNC Symbol;Acc:HGNC:24566] | 0.5701 | 1.89E-06 |
| ENSG00000131941 | RHPN2 | rhophilin Rho GTPase binding protein 2 [Source:HGNC Symbol;Acc:HGNC:19974] | 0.5997 | 1.89E-06 |
| ENSG00000183023 | SLC8A1 | solute carrier family 8 member A1 [Source:HGNC Symbol;Acc:HGNC:11068] | 1.5611 | 1.92E-06 |
| ENSG00000169991 | IFFO2 | intermediate filament family orphan 2 [Source:HGNC Symbol;Acc:HGNC:27006] | 0.5720 | 1.92E-06 |
| ENSG00000274349 | ZNF658 | zinc finger protein 658 [Source:HGNC Symbol;Acc:HGNC:25226] | -0.9079 | 1.97E-06 |
| ENSG00000134046 | MBD2 | methyl-CpG binding domain protein 2 [Source:HGNC Symbol;Acc:HGNC:6917] | 0.6536 | 1.99E-06 |
| ENSG00000160469 | BRSK1 | BR serine/threonine kinase 1 [Source:HGNC Symbol;Acc:HGNC:18994] | 0.8755 | 1.99E-06 |
| ENSG00000227051 | C14orf132 | chromosome 14 open reading frame 132 [Source:HGNC Symbol;Acc:HGNC:20346] | -0.7207 | 2.00E-06 |
| ENSG00000139835 | GRTP1 | growth hormone regulated TBC protein 1 [Source:HGNC Symbol;Acc:HGNC:20310] | -0.6688 | 2.02E-06 |
| ENSG00000250208 | FZD10-AS1 | FZD10 antisense divergent transcript [Source:HGNC Symbol;Acc:HGNC:48632] | 1.9130 | 2.05E-06 |
| ENSG00000135862 | LAMC1 | laminin subunit gamma 1 [Source:HGNC Symbol;Acc:HGNC:6492] | 0.6805 | 2.07E-06 |
| ENSG00000143878 | RHOB | ras homolog family member B [Source:HGNC Symbol;Acc:HGNC:668] | 1.4223 | 2.07E-06 |
| ENSG00000235431 | NA | novel transcript | 0.8560 | 2.07E-06 |
| ENSG00000174306 | ZHX3 | zinc fingers and homeoboxes 3 [Source:HGNC Symbol;Acc:HGNC:15935] | 0.5906 | 2.07E-06 |
| ENSG00000255112 | CHMP1B | charged multivesicular body protein 1B [Source:HGNC Symbol;Acc:HGNC:24287] | 0.5026 | 2.07E-06 |
| ENSG00000144152 | FBLN7 | fibulin 7 [Source:HGNC Symbol;Acc:HGNC:26740] | -1.7058 | 2.11E-06 |
| ENSG00000128185 | DGCR6L | DiGeorge syndrome critical region gene 6 like [Source:HGNC Symbol;Acc:HGNC:18551] | -0.5438 | 2.16E-06 |
| ENSG00000168778 | TCTN2 | tectonic family member 2 [Source:HGNC Symbol;Acc:HGNC:25774] | -0.6854 | 2.17E-06 |

| | | | | |
|---|---|---|---|---|
| ENSG00000130363 | RSPH3 | radial spoke head 3 [Source:HGNC Symbol;Acc:HGNC:21054] | 0.7088 | 2.20E-06 |
| ENSG00000124782 | RREB1 | ras responsive element binding protein 1 [Source:HGNC Symbol;Acc:HGNC:10449] | 0.6678 | 2.24E-06 |
| ENSG00000139624 | CERS5 | ceramide synthase 5 [Source:HGNC Symbol;Acc:HGNC:23749] | 0.5687 | 2.25E-06 |
| ENSG00000103449 | SALL1 | spalt like transcription factor 1 [Source:HGNC Symbol;Acc:HGNC:10524] | 1.9518 | 2.25E-06 |
| ENSG00000163378 | EOGT | EGF domain specific O-linked N-acetylglucosamine transferase [Source:HGNC Symbol;Acc:HGNC:28526] | 0.9084 | 2.26E-06 |
| ENSG00000167604 | NFKBID | NFKB inhibitor delta [Source:HGNC Symbol;Acc:HGNC:15671] | 1.0685 | 2.27E-06 |
| ENSG00000258708 | SLC25A21-AS1 | SLC25A21 antisense RNA 1 [Source:HGNC Symbol;Acc:HGNC:44298] | -1.5465 | 2.28E-06 |
| ENSG00000114026 | OGG1 | 8-oxoguanine DNA glycosylase [Source:HGNC Symbol;Acc:HGNC:8125] | -0.5935 | 2.28E-06 |
| ENSG00000105963 | ADAP1 | ArfGAP with dual PH domains 1 [Source:HGNC Symbol;Acc:HGNC:16486] | 2.1186 | 2.30E-06 |
| ENSG00000236213 | NA | novel transcript | 2.1612 | 2.31E-06 |
| ENSG00000184220 | CMSS1 | cms1 ribosomal small subunit homolog [Source:HGNC Symbol;Acc:HGNC:28666] | 0.5077 | 2.32E-06 |
| ENSG00000124104 | SNX21 | sorting nexin family member 21 [Source:HGNC Symbol;Acc:HGNC:16154] | -0.8057 | 2.39E-06 |
| ENSG00000237499 | WAKMAR2 | wound and keratinocyte migration associated lncRNA 2 [Source:HGNC Symbol;Acc:HGNC:53754] | 0.9332 | 2.39E-06 |
| ENSG00000116096 | SPR | sepiapterin reductase [Source:HGNC Symbol;Acc:HGNC:11257] | -0.6310 | 2.41E-06 |
| ENSG00000269609 | RPARP-AS1 | RPARP antisense RNA 1 [Source:HGNC Symbol;Acc:HGNC:45238] | -0.7937 | 2.41E-06 |
| ENSG00000204186 | ZDBF2 | zinc finger DBF-type containing 2 [Source:HGNC Symbol;Acc:HGNC:29313] | 0.7463 | 2.42E-06 |
| ENSG00000049192 | ADAMTS6 | ADAM metallopeptidase with thrombospondin type 1 motif 6 [Source:HGNC Symbol;Acc:HGNC:222] | 1.2805 | 2.43E-06 |
| ENSG00000277491 | NA | novel transcript | 2.9333 | 2.44E-06 |
| ENSG00000136542 | GALNT5 | polypeptide N-acetylgalactosaminyltransferase 5 [Source:HGNC Symbol;Acc:HGNC:4127] | 2.6643 | 2.44E-06 |
| ENSG00000263069 | RNF213-AS1 | RNF213 antisense RNA 1 [Source:HGNC Symbol;Acc:HGNC:54402] | 0.7547 | 2.46E-06 |
| ENSG00000150457 | LATS2 | large tumor suppressor kinase 2 [Source:HGNC Symbol;Acc:HGNC:6515] | 0.6582 | 2.46E-06 |
| ENSG00000177640 | CASC2 | cancer susceptibility 2 [Source:HGNC Symbol;Acc:HGNC:22933] | -1.1861 | 2.46E-06 |
| ENSG00000116205 | TCEANC2 | transcription elongation factor A N-terminal and central domain containing 2 [Source:HGNC Symbol;Acc:HGNC:26494] | -0.6336 | 2.48E-06 |
| ENSG00000120915 | EPHX2 | epoxide hydrolase 2 [Source:HGNC Symbol;Acc:HGNC:3402] | -1.1676 | 2.49E-06 |
| ENSG00000156206 | CFAP161 | cilia and flagella associated protein 161 [Source:HGNC Symbol;Acc:HGNC:26782] | 3.6437 | 2.49E-06 |
| ENSG00000203497 | PDCD4-AS1 | PDCD4 antisense RNA 1 [Source:HGNC Symbol;Acc:HGNC:27425] | -1.0802 | 2.53E-06 |
| ENSG00000248161 | NA | novel transcript | -1.9957 | 2.55E-06 |
| ENSG00000138780 | GSTCD | glutathione S-transferase C-terminal domain containing [Source:HGNC Symbol;Acc:HGNC:25806] | -0.5828 | 2.56E-06 |
| ENSG00000171503 | ETFDH | electron transfer flavoprotein dehydrogenase [Source:HGNC Symbol;Acc:HGNC:3483] | -0.5922 | 2.59E-06 |
| ENSG00000158710 | TAGLN2 | transgelin 2 [Source:HGNC Symbol;Acc:HGNC:11554] | 0.5542 | 2.62E-06 |
| ENSG00000180370 | PAK2 | p21 (RAC1) activated kinase 2 [Source:HGNC Symbol;Acc:HGNC:8591] | 0.6150 | 2.63E-06 |
| ENSG00000162494 | LRRC38 | leucine rich repeat containing 38 [Source:HGNC Symbol;Acc:HGNC:27005] | -1.1616 | 2.65E-06 |
| ENSG00000102007 | PLP2 | proteolipid protein 2 [Source:HGNC Symbol;Acc:HGNC:9087] | 0.5934 | 2.65E-06 |
| ENSG00000124596 | OARD1 | O-acyl-ADP-ribose deacylase 1 [Source:HGNC Symbol;Acc:HGNC:21257] | -0.5723 | 2.68E-06 |
| ENSG00000103184 | SEC14L5 | SEC14 like lipid binding 5 [Source:HGNC Symbol;Acc:HGNC:29032] | -2.1158 | 2.74E-06 |
| ENSG00000280594 | BTG3-AS1 | BTG3 antisense RNA 1 [Source:HGNC Symbol;Acc:HGNC:53145] | 1.3552 | 2.78E-06 |
| ENSG00000225595 | LINC01623 | long intergenic non-protein coding RNA 1623 [Source:HGNC Symbol;Acc:HGNC:52050] | -3.5257 | 2.81E-06 |
| ENSG00000198125 | MB | myoglobin [Source:HGNC Symbol;Acc:HGNC:6915] | -0.8821 | 2.83E-06 |
| ENSG00000154079 | SDHAF4 | succinate dehydrogenase complex assembly factor 4 [Source:HGNC Symbol;Acc:HGNC:20957] | -0.8627 | 2.84E-06 |
| ENSG00000285770 | NA | novel transcript, antisense to CLSTN3 | 4.6929 | 2.85E-06 |
| ENSG00000213949 | ITGA1 | integrin subunit alpha 1 [Source:HGNC Symbol;Acc:HGNC:6134] | 0.8012 | 2.86E-06 |
| ENSG00000148344 | PTGES | prostaglandin E synthase [Source:HGNC Symbol;Acc:HGNC:9599] | -1.7363 | 2.89E-06 |

| | | | | |
|---|---|---|---|---|
| ENSG00000117152 | RGS4 | regulator of G protein signaling 4 [Source:HGNC Symbol;Acc:HGNC:10000] | 3.5385 | 2.90E-06 |
| ENSG00000164970 | FAM219A | family with sequence similarity 219 member A [Source:HGNC Symbol;Acc:HGNC:19920] | 0.5797 | 2.91E-06 |
| ENSG00000184232 | OAF | out at first homolog [Source:HGNC Symbol;Acc:HGNC:28752] | 0.7039 | 2.91E-06 |
| ENSG00000198911 | SREBF2 | sterol regulatory element binding transcription factor 2 [Source:HGNC Symbol;Acc:HGNC:11290] | -0.5303 | 2.93E-06 |
| ENSG00000100802 | C14orf93 | chromosome 14 open reading frame 93 [Source:HGNC Symbol;Acc:HGNC:20162] | -0.9961 | 2.94E-06 |
| ENSG00000188396 | TCTEX1D4 | Tctex1 domain containing 4 [Source:HGNC Symbol;Acc:HGNC:32315] | 2.6066 | 2.98E-06 |
| ENSG00000109321 | AREG | amphiregulin [Source:HGNC Symbol;Acc:HGNC:651] | 5.2813 | 2.99E-06 |
| ENSG00000112796 | ENPP5 | ectonucleotide pyrophosphatase/phosphodiesterase family member 5 [Source:HGNC Symbol;Acc:HGNC:13717] | -0.5731 | 3.00E-06 |
| ENSG00000167191 | GPRC5B | G protein-coupled receptor class C group 5 member B [Source:HGNC Symbol;Acc:HGNC:13308] | -1.0595 | 3.00E-06 |
| ENSG00000188906 | LRRK2 | leucine rich repeat kinase 2 [Source:HGNC Symbol;Acc:HGNC:18618] | -1.3692 | 3.05E-06 |
| ENSG00000119125 | GDA | guanine deaminase [Source:HGNC Symbol;Acc:HGNC:4212] | 1.7247 | 3.12E-06 |
| ENSG00000182957 | SPATA13 | spermatogenesis associated 13 [Source:HGNC Symbol;Acc:HGNC:23222] | 0.5647 | 3.13E-06 |
| ENSG00000142235 | LMTK3 | lemur tyrosine kinase 3 [Source:HGNC Symbol;Acc:HGNC:19295] | -0.7366 | 3.15E-06 |
| ENSG00000145012 | LPP | LIM domain containing preferred translocation partner in lipoma [Source:HGNC Symbol;Acc:HGNC:6679] | 1.1734 | 3.18E-06 |
| ENSG00000198252 | STYX | serine/threonine/tyrosine interacting protein [Source:HGNC Symbol;Acc:HGNC:11447] | -0.5029 | 3.18E-06 |
| ENSG00000135074 | ADAM19 | ADAM metallopeptidase domain 19 [Source:HGNC Symbol;Acc:HGNC:197] | 1.6935 | 3.22E-06 |
| ENSG00000134333 | LDHA | lactate dehydrogenase A [Source:HGNC Symbol;Acc:HGNC:6535] | 0.7836 | 3.23E-06 |
| ENSG00000259129 | LINC00648 | long intergenic non-protein coding RNA 648 [Source:HGNC Symbol;Acc:HGNC:44302] | -1.3493 | 3.23E-06 |
| ENSG00000092020 | PPP2R3C | protein phosphatase 2 regulatory subunit B"gamma [Source:HGNC Symbol;Acc:HGNC:17485] | 0.6599 | 3.23E-06 |
| ENSG00000122877 | EGR2 | early growth response 2 [Source:HGNC Symbol;Acc:HGNC:3239] | 5.2759 | 3.26E-06 |
| ENSG00000119514 | GALNT12 | polypeptide N-acetylgalactosaminyltransferase 12 [Source:HGNC Symbol;Acc:HGNC:19877] | 1.3496 | 3.26E-06 |
| ENSG00000255641 | NA | novel protein | 4.5380 | 3.27E-06 |
| ENSG00000279821 | NA | TEC | 1.7345 | 3.29E-06 |
| ENSG00000187391 | MAGI2 | membrane associated guanylate kinase, WW and PDZ domain containing 2 [Source:HGNC Symbol;Acc:HGNC:18957] | -0.6691 | 3.32E-06 |
| ENSG00000188352 | FOCAD | focadhesin [Source:HGNC Symbol;Acc:HGNC:23377] | 0.6484 | 3.34E-06 |
| ENSG00000174460 | ZCCHC12 | zinc finger CCHC-type containing 12 [Source:HGNC Symbol;Acc:HGNC:27273] | 0.9164 | 3.37E-06 |
| ENSG00000178381 | ZFAND2A | zinc finger AN1-type containing 2A [Source:HGNC Symbol;Acc:HGNC:28073] | 0.7271 | 3.40E-06 |
| ENSG00000117000 | RLF | RLF zinc finger [Source:HGNC Symbol;Acc:HGNC:10025] | 0.5884 | 3.41E-06 |
| ENSG00000139880 | CDH24 | cadherin 24 [Source:HGNC Symbol;Acc:HGNC:14265] | 0.7658 | 3.45E-06 |
| ENSG00000178860 | MSC | musculin [Source:HGNC Symbol;Acc:HGNC:7321] | 1.5817 | 3.45E-06 |
| ENSG00000112972 | HMGCS1 | 3-hydroxy-3-methylglutaryl-CoA synthase 1 [Source:HGNC Symbol;Acc:HGNC:5007] | -0.6564 | 3.48E-06 |
| ENSG00000265972 | TXNIP | thioredoxin interacting protein [Source:HGNC Symbol;Acc:HGNC:16952] | -1.6163 | 3.62E-06 |
| ENSG00000116005 | PCYOX1 | prenylcysteine oxidase 1 [Source:HGNC Symbol;Acc:HGNC:20588] | -0.5093 | 3.64E-06 |
| ENSG00000244681 | MTHFD2P1 | methylenetetrahydrofolate dehydrogenase (NADP+ dependent) 2, methenyltetrahydrofolate cyclohydrolase pseudogene 1 [Source:HGNC Symbol;Acc:HGNC:48859] | 0.5549 | 3.66E-06 |
| ENSG00000033170 | FUT8 | fucosyltransferase 8 [Source:HGNC Symbol;Acc:HGNC:4019] | 0.6935 | 3.66E-06 |
| ENSG00000124222 | STX16 | syntaxin 16 [Source:HGNC Symbol;Acc:HGNC:11431] | 0.5425 | 3.69E-06 |
| ENSG00000136247 | ZDHHC4 | zinc finger DHHC-type palmitoyltransferase 4 [Source:HGNC Symbol;Acc:HGNC:18471] | -0.6317 | 3.69E-06 |
| ENSG00000101871 | MID1 | midline 1 [Source:HGNC Symbol;Acc:HGNC:7095] | -0.6828 | 3.69E-06 |
| ENSG00000163629 | PTPN13 | protein tyrosine phosphatase non-receptor type 13 [Source:HGNC Symbol;Acc:HGNC:9646] | 0.9271 | 3.73E-06 |
| ENSG00000266010 | GATA6-AS1 | GATA6 antisense RNA 1 (head to head) [Source:HGNC Symbol;Acc:HGNC:48840] | 1.2767 | 3.81E-06 |
| ENSG00000172175 | MALT1 | MALT1 paracaspase [Source:HGNC Symbol;Acc:HGNC:6819] | 0.5356 | 3.87E-06 |

| | | | | |
|---|---|---|---|---|
| ENSG00000140280 | LYSMD2 | LysM domain containing 2 [Source:HGNC Symbol;Acc:HGNC:28571] | 0.8886 | 3.90E-06 |
| ENSG00000144674 | GOLGA4 | golgin A4 [Source:HGNC Symbol;Acc:HGNC:4427] | 0.5101 | 3.96E-06 |
| ENSG00000118046 | STK11 | serine/threonine kinase 11 [Source:HGNC Symbol;Acc:HGNC:11389] | -0.5329 | 4.00E-06 |
| ENSG00000277715 | NA | novel transcript, antisense to CKAP4 | -1.6246 | 4.07E-06 |
| ENSG00000176049 | JAKMIP2 | janus kinase and microtubule interacting protein 2 [Source:HGNC Symbol;Acc:HGNC:29067] | -0.5854 | 4.07E-06 |
| ENSG00000259877 | NA | novel transcript, antisense to ZNF778 | -0.7404 | 4.08E-06 |
| ENSG00000135776 | ABCB10 | ATP binding cassette subfamily B member 10 [Source:HGNC Symbol;Acc:HGNC:41] | -0.5913 | 4.13E-06 |
| ENSG00000181634 | TNFSF15 | TNF superfamily member 15 [Source:HGNC Symbol;Acc:HGNC:11931] | 3.2808 | 4.18E-06 |
| ENSG00000134996 | OSTF1 | osteoclast stimulating factor 1 [Source:HGNC Symbol;Acc:HGNC:8510] | 0.6274 | 4.23E-06 |
| ENSG00000141441 | GAREM1 | GRB2 associated regulator of MAPK1 subtype 1 [Source:HGNC Symbol;Acc:HGNC:26136] | 1.9186 | 4.23E-06 |
| ENSG00000151413 | NUBPL | nucleotide binding protein like [Source:HGNC Symbol;Acc:HGNC:20278] | -0.6816 | 4.23E-06 |
| ENSG00000169189 | NSMCE1 | NSE1 homolog, SMC5-SMC6 complex component [Source:HGNC Symbol;Acc:HGNC:29897] | -0.5656 | 4.23E-06 |
| ENSG00000184381 | PLA2G6 | phospholipase A2 group VI [Source:HGNC Symbol;Acc:HGNC:9039] | -0.6253 | 4.27E-06 |
| ENSG00000074219 | TEAD2 | TEA domain transcription factor 2 [Source:HGNC Symbol;Acc:HGNC:11715] | -0.6708 | 4.29E-06 |
| ENSG00000272686 | WASL-DT | WASL divergent transcript [Source:HGNC Symbol;Acc:HGNC:54820] | -0.7130 | 4.30E-06 |
| ENSG00000135373 | EHF | ETS homologous factor [Source:HGNC Symbol;Acc:HGNC:3246] | -2.4701 | 4.31E-06 |
| ENSG00000151718 | WWC2 | WW and C2 domain containing 2 [Source:HGNC Symbol;Acc:HGNC:24148] | 0.7050 | 4.33E-06 |
| ENSG00000102967 | DHODH | dihydroorotate dehydrogenase (quinone) [Source:HGNC Symbol;Acc:HGNC:2867] | -0.7051 | 4.36E-06 |
| ENSG00000178537 | SLC25A20 | solute carrier family 25 member 20 [Source:HGNC Symbol;Acc:HGNC:1421] | -0.5775 | 4.38E-06 |
| ENSG00000154370 | TRIM11 | tripartite motif containing 11 [Source:HGNC Symbol;Acc:HGNC:16281] | 0.5923 | 4.39E-06 |
| ENSG00000115353 | TACR1 | tachykinin receptor 1 [Source:HGNC Symbol;Acc:HGNC:11526] | -1.6929 | 4.39E-06 |
| ENSG00000175395 | ZNF25 | zinc finger protein 25 [Source:HGNC Symbol;Acc:HGNC:13043] | -0.6179 | 4.43E-06 |
| ENSG00000182040 | USH1G | USH1 protein network component sans [Source:HGNC Symbol;Acc:HGNC:16356] | 0.8692 | 4.50E-06 |
| ENSG00000126070 | AGO3 | argonaute RISC catalytic component 3 [Source:HGNC Symbol;Acc:HGNC:18421] | 0.6485 | 4.61E-06 |
| ENSG00000279561 | NA | uncharacterized LOC100132249 [Source:NCBI gene (formerly Entrezgene);Acc:100132249] | -0.6260 | 4.61E-06 |
| ENSG00000213689 | TREX1 | three prime repair exonuclease 1 [Source:HGNC Symbol;Acc:HGNC:12269] | -0.5777 | 4.63E-06 |
| ENSG00000172819 | RARG | retinoic acid receptor gamma [Source:HGNC Symbol;Acc:HGNC:9866] | -0.5529 | 4.63E-06 |
| ENSG00000111331 | OAS3 | 2'-5'-oligoadenylate synthetase 3 [Source:HGNC Symbol;Acc:HGNC:8088] | -0.9019 | 4.71E-06 |
| ENSG00000105875 | WDR91 | WD repeat domain 91 [Source:HGNC Symbol;Acc:HGNC:24997] | 0.8897 | 4.75E-06 |
| ENSG00000059804 | SLC2A3 | solute carrier family 2 member 3 [Source:HGNC Symbol;Acc:HGNC:11007] | 0.8057 | 4.77E-06 |
| ENSG00000260992 | DOCK9-DT | DOCK9 divergent transcript [Source:HGNC Symbol;Acc:HGNC:43696] | -1.3715 | 4.77E-06 |
| ENSG00000178690 | DYNAP | dynactin associated protein [Source:HGNC Symbol;Acc:HGNC:26808] | -1.2727 | 4.82E-06 |
| ENSG00000180867 | PDIA3P1 | protein disulfide isomerase family A member 3 pseudogene 1 [Source:HGNC Symbol;Acc:HGNC:4607] | 0.6982 | 4.87E-06 |
| ENSG00000267280 | TBX2-AS1 | TBX2 antisense RNA 1 [Source:HGNC Symbol;Acc:HGNC:50355] | -1.4278 | 4.90E-06 |
| ENSG00000214425 | LRRC37A4P | leucine rich repeat containing 37 member A4, pseudogene [Source:HGNC Symbol;Acc:HGNC:25479] | -0.6043 | 5.05E-06 |
| ENSG00000164087 | POC1A | POC1 centriolar protein A [Source:HGNC Symbol;Acc:HGNC:24488] | -0.8143 | 5.06E-06 |
| ENSG00000142684 | ZNF593 | zinc finger protein 593 [Source:HGNC Symbol;Acc:HGNC:30943] | 0.6658 | 5.07E-06 |
| ENSG00000175806 | MSRA | methionine sulfoxide reductase A [Source:HGNC Symbol;Acc:HGNC:7377] | 0.6809 | 5.25E-06 |
| ENSG00000129315 | CCNT1 | cyclin T1 [Source:HGNC Symbol;Acc:HGNC:1599] | 0.5217 | 5.30E-06 |
| ENSG00000173193 | PARP14 | poly(ADP-ribose) polymerase family member 14 [Source:HGNC Symbol;Acc:HGNC:29232] | 0.6747 | 5.31E-06 |
| ENSG00000196562 | SULF2 | sulfatase 2 [Source:HGNC Symbol;Acc:HGNC:20392] | -1.3893 | 5.31E-06 |
| ENSG00000135643 | KCNMB4 | potassium calcium-activated channel subfamily M regulatory beta subunit 4 [Source:HGNC Symbol;Acc:HGNC:6289] | -0.5790 | 5.32E-06 |
| ENSG00000082512 | TRAF5 | TNF receptor associated factor 5 [Source:HGNC Symbol;Acc:HGNC:12035] | -1.1422 | 5.35E-06 |

| | | | | |
|---|---|---|---|---|
| ENSG00000239415 | NA | novel transcript, antisense to MCM3APAS | -1.3231 | 5.41E-06 |
| ENSG00000074660 | SCARF1 | scavenger receptor class F member 1 [Source:HGNC Symbol;Acc:HGNC:16820] | 2.3829 | 5.42E-06 |
| ENSG00000101098 | RIMS4 | regulating synaptic membrane exocytosis 4 [Source:HGNC Symbol;Acc:HGNC:16183] | -2.4771 | 5.44E-06 |
| ENSG00000257605 | MYG1-AS1 | MYG1 antisense RNA 1 [Source:HGNC Symbol;Acc:HGNC:54810] | 2.2783 | 5.57E-06 |
| ENSG00000189306 | RRP7A | ribosomal RNA processing 7 homolog A [Source:HGNC Symbol;Acc:HGNC:24286] | 0.5514 | 5.59E-06 |
| ENSG00000169087 | HSPBAP1 | HSPB1 associated protein 1 [Source:HGNC Symbol;Acc:HGNC:16389] | 0.5738 | 5.63E-06 |
| ENSG00000182264 | IZUMO1 | izumo sperm-egg fusion 1 [Source:HGNC Symbol;Acc:HGNC:28539] | -1.2601 | 5.63E-06 |
| ENSG00000099204 | ABLIM1 | actin binding LIM protein 1 [Source:HGNC Symbol;Acc:HGNC:78] | -1.6942 | 5.65E-06 |
| ENSG00000250519 | NA | novel transcript | -1.0051 | 5.67E-06 |
| ENSG00000127337 | YEATS4 | YEATS domain containing 4 [Source:HGNC Symbol;Acc:HGNC:24859] | -0.5025 | 5.68E-06 |
| ENSG00000181894 | ZNF329 | zinc finger protein 329 [Source:HGNC Symbol;Acc:HGNC:14209] | -0.5645 | 5.69E-06 |
| ENSG00000140691 | ARMC5 | armadillo repeat containing 5 [Source:HGNC Symbol;Acc:HGNC:25781] | 0.5244 | 5.72E-06 |
| ENSG00000163293 | NIPAL1 | NIPA like domain containing 1 [Source:HGNC Symbol;Acc:HGNC:27194] | 1.4514 | 5.73E-06 |
| ENSG00000198648 | STK39 | serine/threonine kinase 39 [Source:HGNC Symbol;Acc:HGNC:17717] | -0.5457 | 5.75E-06 |
| ENSG00000272604 | NA | novel transcript, antisense to EFCAB10 | 1.3064 | 5.76E-06 |
| ENSG00000198771 | RCSD1 | RCSD domain containing 1 [Source:HGNC Symbol;Acc:HGNC:28310] | -3.1103 | 5.87E-06 |
| ENSG00000149084 | HSD17B12 | hydroxysteroid 17-beta dehydrogenase 12 [Source:HGNC Symbol;Acc:HGNC:18646] | 0.6265 | 6.00E-06 |
| ENSG00000229891 | LINC01315 | long intergenic non-protein coding RNA 1315 [Source:HGNC Symbol;Acc:HGNC:50513] | -1.8343 | 6.09E-06 |
| ENSG00000233461 | NA | novel transcript | 1.3719 | 6.17E-06 |
| ENSG00000239440 | LINC02008 | long intergenic non-protein coding RNA 2008 [Source:HGNC Symbol;Acc:HGNC:52844] | 2.0127 | 6.21E-06 |
| ENSG00000187479 | C11orf96 | chromosome 11 open reading frame 96 [Source:HGNC Symbol;Acc:HGNC:38675] | 1.1167 | 6.22E-06 |
| ENSG00000280433 | NA | novel protein, similar to trafficking protein particle complex 10 TRAPPC10 | -0.6739 | 6.23E-06 |
| ENSG00000065621 | GSTO2 | glutathione S-transferase omega 2 [Source:HGNC Symbol;Acc:HGNC:23064] | -0.6222 | 6.23E-06 |
| ENSG00000166862 | CACNG2 | calcium voltage-gated channel auxiliary subunit gamma 2 [Source:HGNC Symbol;Acc:HGNC:1406] | -1.1976 | 6.26E-06 |
| ENSG00000176531 | PHLDB3 | pleckstrin homology like domain family B member 3 [Source:HGNC Symbol;Acc:HGNC:30499] | 1.0429 | 6.40E-06 |
| ENSG00000212993 | POU5F1B | POU class 5 homeobox 1B [Source:HGNC Symbol;Acc:HGNC:9223] | -1.8353 | 6.41E-06 |
| ENSG00000171453 | POLR1C | RNA polymerase I and III subunit C [Source:HGNC Symbol;Acc:HGNC:20194] | 0.5535 | 6.42E-06 |
| ENSG00000161921 | CXCL16 | C-X-C motif chemokine ligand 16 [Source:HGNC Symbol;Acc:HGNC:16642] | 1.6232 | 6.45E-06 |
| ENSG00000249899 | NA | novel transcript | 1.7336 | 6.46E-06 |
| ENSG00000083544 | TDRD3 | tudor domain containing 3 [Source:HGNC Symbol;Acc:HGNC:20612] | -0.7297 | 6.51E-06 |
| ENSG00000116117 | PARD3B | par-3 family cell polarity regulator beta [Source:HGNC Symbol;Acc:HGNC:14446] | -0.8150 | 6.55E-06 |
| ENSG00000155962 | CLIC2 | chloride intracellular channel 2 [Source:HGNC Symbol;Acc:HGNC:2063] | -1.0526 | 6.58E-06 |
| ENSG00000172572 | PDE3A | phosphodiesterase 3A [Source:HGNC Symbol;Acc:HGNC:8778] | 2.9668 | 6.63E-06 |
| ENSG00000171815 | PCDHB1 | protocadherin beta 1 [Source:HGNC Symbol;Acc:HGNC:8680] | -2.5634 | 6.71E-06 |
| ENSG00000264964 | NA | novel transcript | -2.5396 | 6.72E-06 |
| ENSG00000118508 | RAB32 | RAB32, member RAS oncogene family [Source:HGNC Symbol;Acc:HGNC:9772] | -0.6929 | 6.77E-06 |
| ENSG00000137877 | SPTBN5 | spectrin beta, non-erythrocytic 5 [Source:HGNC Symbol;Acc:HGNC:15680] | 1.9136 | 6.78E-06 |
| ENSG00000181215 | C4orf50 | chromosome 4 open reading frame 50 [Source:HGNC Symbol;Acc:HGNC:33766] | -3.5325 | 6.78E-06 |
| ENSG00000074621 | SLC24A1 | solute carrier family 24 member 1 [Source:HGNC Symbol;Acc:HGNC:10975] | -0.6400 | 6.80E-06 |
| ENSG00000013563 | DNASE1L1 | deoxyribonuclease 1 like 1 [Source:HGNC Symbol;Acc:HGNC:2957] | -0.6824 | 6.82E-06 |
| ENSG00000197948 | FCHSD1 | FCH and double SH3 domains 1 [Source:HGNC Symbol;Acc:HGNC:25463] | 0.5716 | 6.84E-06 |
| ENSG00000225313 | NA | novel transcript | 1.7787 | 6.85E-06 |
| ENSG00000247934 | NA | novel transcript, antisense to CCDC91 | -0.8601 | 6.90E-06 |

| | | | | |
|---|---|---|---|---|
| ENSG00000164638 | SLC29A4 | solute carrier family 29 member 4 [Source:HGNC Symbol;Acc:HGNC:23097] | -0.7228 | 6.92E-06 |
| ENSG00000059769 | DNAJC25 | DnaJ heat shock protein family (Hsp40) member C25 [Source:HGNC Symbol;Acc:HGNC:34187] | 0.5472 | 6.98E-06 |
| ENSG00000177910 | SPATA31C2 | SPATA31 subfamily C member 2 [Source:HGNC Symbol;Acc:HGNC:24508] | -0.8468 | 7.01E-06 |
| ENSG00000205476 | CCDC85C | coiled-coil domain containing 85C [Source:HGNC Symbol;Acc:HGNC:35459] | -0.8074 | 7.02E-06 |
| ENSG00000136205 | TNS3 | tensin 3 [Source:HGNC Symbol;Acc:HGNC:21616] | 1.6761 | 7.05E-06 |
| ENSG00000106003 | LFNG | LFNG O-fucosylpeptide 3-beta-N-acetylglucosaminyltransferase [Source:HGNC Symbol;Acc:HGNC:6560] | 1.4474 | 7.33E-06 |
| ENSG00000189376 | C8orf76 | chromosome 8 open reading frame 76 [Source:HGNC Symbol;Acc:HGNC:25924] | 0.5168 | 7.37E-06 |
| ENSG00000237596 | NA | novel transcript | 2.2598 | 7.37E-06 |
| ENSG00000211455 | STK38L | serine/threonine kinase 38 like [Source:HGNC Symbol;Acc:HGNC:17848] | 0.5419 | 7.38E-06 |
| ENSG00000129680 | MAP7D3 | MAP7 domain containing 3 [Source:HGNC Symbol;Acc:HGNC:25742] | 0.7225 | 7.58E-06 |
| ENSG00000140326 | CDAN1 | codanin 1 [Source:HGNC Symbol;Acc:HGNC:1713] | -0.7450 | 7.58E-06 |
| ENSG00000011566 | MAP4K3 | mitogen-activated protein kinase kinase kinase kinase 3 [Source:HGNC Symbol;Acc:HGNC:6865] | -0.5523 | 7.62E-06 |
| ENSG00000106078 | COBL | cordon-bleu WH2 repeat protein [Source:HGNC Symbol;Acc:HGNC:22199] | 4.1207 | 7.75E-06 |
| ENSG00000099785 | MARCHF2 | membrane associated ring-CH-type finger 2 [Source:HGNC Symbol;Acc:HGNC:28038] | -0.9968 | 7.84E-06 |
| ENSG00000254837 | NA | uncharacterized LOC100287896 [Source:NCBI gene (formerly Entrezgene);Acc:100287896] | -1.2098 | 7.95E-06 |
| ENSG00000114166 | KAT2B | lysine acetyltransferase 2B [Source:HGNC Symbol;Acc:HGNC:8638] | 0.7000 | 7.97E-06 |
| ENSG00000126243 | LRFN3 | leucine rich repeat and fibronectin type III domain containing 3 [Source:HGNC Symbol;Acc:HGNC:28370] | -0.5316 | 8.06E-06 |
| ENSG00000010295 | IFFO1 | intermediate filament family orphan 1 [Source:HGNC Symbol;Acc:HGNC:24970] | -0.6779 | 8.21E-06 |
| ENSG00000116641 | DOCK7 | dedicator of cytokinesis 7 [Source:HGNC Symbol;Acc:HGNC:19190] | -0.8955 | 8.25E-06 |
| ENSG00000124496 | TRERF1 | transcriptional regulating factor 1 [Source:HGNC Symbol;Acc:HGNC:18273] | -0.5687 | 8.27E-06 |
| ENSG00000153904 | DDAH1 | dimethylarginine dimethylaminohydrolase 1 [Source:HGNC Symbol;Acc:HGNC:2715] | 0.5193 | 8.30E-06 |
| ENSG00000174516 | PELI3 | pellino E3 ubiquitin protein ligase family member 3 [Source:HGNC Symbol;Acc:HGNC:30010] | -0.7428 | 8.54E-06 |
| ENSG00000169762 | TAPT1 | transmembrane anterior posterior transformation 1 [Source:HGNC Symbol;Acc:HGNC:26887] | 0.8506 | 8.56E-06 |
| ENSG00000272068 | NA | novel transcript | 0.9324 | 8.57E-06 |
| ENSG00000183826 | BTBD9 | BTB domain containing 9 [Source:HGNC Symbol;Acc:HGNC:21228] | 0.5906 | 8.64E-06 |
| ENSG00000228983 | SLC47A1P1 | SLC47A1 pseudogene 1 [Source:HGNC Symbol;Acc:HGNC:51849] | 1.7903 | 8.67E-06 |
| ENSG00000271913 | NA | novel transcript, antisense to TAGAP | -1.4375 | 8.68E-06 |
| ENSG00000135736 | CCDC102A | coiled-coil domain containing 102A [Source:HGNC Symbol;Acc:HGNC:28097] | -0.8382 | 8.77E-06 |
| ENSG00000152128 | TMEM163 | transmembrane protein 163 [Source:HGNC Symbol;Acc:HGNC:25380] | 1.3003 | 8.78E-06 |
| ENSG00000258232 | NA | novel transcript, antisense to TUBA1C | 0.9532 | 8.84E-06 |
| ENSG00000196155 | PLEKHG4 | pleckstrin homology and RhoGEF domain containing G4 [Source:HGNC Symbol;Acc:HGNC:24501] | 0.7014 | 8.87E-06 |
| ENSG00000133056 | PIK3C2B | phosphatidylinositol-4-phosphate 3-kinase catalytic subunit type 2 beta [Source:HGNC Symbol;Acc:HGNC:8972] | -1.3441 | 9.04E-06 |
| ENSG00000107551 | RASSF4 | Ras association domain family member 4 [Source:HGNC Symbol;Acc:HGNC:20793] | -1.2477 | 9.04E-06 |
| ENSG00000137135 | ARHGEF39 | Rho guanine nucleotide exchange factor 39 [Source:HGNC Symbol;Acc:HGNC:25909] | -0.6315 | 9.08E-06 |
| ENSG00000180354 | MTURN | maturin, neural progenitor differentiation regulator homolog [Source:HGNC Symbol;Acc:HGNC:25457] | -1.2653 | 9.11E-06 |
| ENSG00000074370 | ATP2A3 | ATPase sarcoplasmic/endoplasmic reticulum Ca2+ transporting 3 [Source:HGNC Symbol;Acc:HGNC:813] | -0.6872 | 9.15E-06 |
| ENSG00000162144 | CYB561A3 | cytochrome b561 family member A3 [Source:HGNC Symbol;Acc:HGNC:23014] | -0.5077 | 9.15E-06 |
| ENSG00000165449 | SLC16A9 | solute carrier family 16 member 9 [Source:HGNC Symbol;Acc:HGNC:23520] | -1.7342 | 9.15E-06 |
| ENSG00000239445 | ST3GAL6-AS1 | ST3GAL6 antisense RNA 1 [Source:HGNC Symbol;Acc:HGNC:40828] | -1.5396 | 9.17E-06 |
| ENSG00000175040 | CHST2 | carbohydrate sulfotransferase 2 [Source:HGNC Symbol;Acc:HGNC:1970] | 2.0898 | 9.18E-06 |
| ENSG00000160271 | RALGDS | ral guanine nucleotide dissociation stimulator [Source:HGNC Symbol;Acc:HGNC:9842] | -0.6077 | 9.26E-06 |
| ENSG00000231752 | EMBP1 | embigin pseudogene 1 [Source:HGNC Symbol;Acc:HGNC:38661] | -1.0677 | 9.26E-06 |
| ENSG00000163032 | VSNL1 | visinin like 1 [Source:HGNC Symbol;Acc:HGNC:12722] | -0.5590 | 9.26E-06 |

| | | | | |
|---|---|---|---|---|
| ENSG00000143674 | MAP3K21 | mitogen-activated protein kinase kinase kinase 21 [Source:HGNC Symbol;Acc:HGNC:29798] | -0.5592 | 9.28E-06 |
| ENSG00000174599 | TRAM1L1 | translocation associated membrane protein 1 like 1 [Source:HGNC Symbol;Acc:HGNC:28371] | -1.0832 | 9.33E-06 |
| ENSG00000074590 | NUAK1 | NUAK family kinase 1 [Source:HGNC Symbol;Acc:HGNC:14311] | 1.0181 | 9.41E-06 |
| ENSG00000270885 | RASL10B | RAS like family 10 member B [Source:HGNC Symbol;Acc:HGNC:30295] | 1.3659 | 9.68E-06 |
| ENSG00000090263 | MRPS33 | mitochondrial ribosomal protein S33 [Source:HGNC Symbol;Acc:HGNC:16634] | -0.5248 | 9.75E-06 |
| ENSG00000144847 | IGSF11 | immunoglobulin superfamily member 11 [Source:HGNC Symbol;Acc:HGNC:16669] | -1.0595 | 9.81E-06 |
| ENSG00000143322 | ABL2 | ABL proto-oncogene 2, non-receptor tyrosine kinase [Source:HGNC Symbol;Acc:HGNC:77] | 0.5166 | 9.82E-06 |
| ENSG00000214029 | ZNF891 | zinc finger protein 891 [Source:HGNC Symbol;Acc:HGNC:38709] | -0.9238 | 9.83E-06 |
| ENSG00000229776 | C4B-AS1 | C4B antisense RNA 1 [Source:HGNC Symbol;Acc:HGNC:39752] | 1.7734 | 1.01E-05 |
| ENSG00000233627 | C4A-AS1 | C4A antisense RNA 1 [Source:HGNC Symbol;Acc:HGNC:39753] | 1.7734 | 1.01E-05 |
| ENSG00000177335 | C8orf31 | chromosome 8 open reading frame 31 [Source:HGNC Symbol;Acc:HGNC:26731] | 0.7158 | 1.01E-05 |
| ENSG00000125534 | PPDPF | pancreatic progenitor cell differentiation and proliferation factor [Source:HGNC Symbol;Acc:HGNC:16142] | -0.6378 | 1.03E-05 |
| ENSG00000151148 | UBE3B | ubiquitin protein ligase E3B [Source:HGNC Symbol;Acc:HGNC:13478] | -0.5625 | 1.03E-05 |
| ENSG00000182050 | MGAT4C | MGAT4 family member C [Source:HGNC Symbol;Acc:HGNC:30871] | -1.2997 | 1.04E-05 |
| ENSG00000232160 | RAP2C-AS1 | RAP2C antisense RNA 1 [Source:HGNC Symbol;Acc:HGNC:40957] | -1.2002 | 1.05E-05 |
| ENSG00000172059 | KLF11 | Kruppel like factor 11 [Source:HGNC Symbol;Acc:HGNC:11811] | -0.5397 | 1.06E-05 |
| ENSG00000155636 | RBM45 | RNA binding motif protein 45 [Source:HGNC Symbol;Acc:HGNC:24468] | -0.5764 | 1.07E-05 |
| ENSG00000138463 | SLC49A4 | solute carrier family 49 member 4 [Source:HGNC Symbol;Acc:HGNC:16628] | 0.8343 | 1.08E-05 |
| ENSG00000197183 | NOL4L | nucleolar protein 4 like [Source:HGNC Symbol;Acc:HGNC:16106] | -0.7969 | 1.09E-05 |
| ENSG00000231566 | LINC02595 | long intergenic non-protein coding RNA 2595 [Source:HGNC Symbol;Acc:HGNC:53946] | 1.8993 | 1.10E-05 |
| ENSG00000101190 | TCFL5 | transcription factor like 5 [Source:HGNC Symbol;Acc:HGNC:11646] | -1.0443 | 1.11E-05 |
| ENSG00000171435 | KSR2 | kinase suppressor of ras 2 [Source:HGNC Symbol;Acc:HGNC:18610] | -0.8690 | 1.11E-05 |
| ENSG00000148814 | LRRC27 | leucine rich repeat containing 27 [Source:HGNC Symbol;Acc:HGNC:29346] | -1.0242 | 1.12E-05 |
| ENSG00000138002 | IFT172 | intraflagellar transport 172 [Source:HGNC Symbol;Acc:HGNC:30391] | -0.8816 | 1.12E-05 |
| ENSG00000100167 | SEPTIN3 | septin 3 [Source:HGNC Symbol;Acc:HGNC:10750] | -0.5493 | 1.13E-05 |
| ENSG00000112394 | SLC16A10 | solute carrier family 16 member 10 [Source:HGNC Symbol;Acc:HGNC:17027] | -1.3132 | 1.13E-05 |
| ENSG00000185024 | BRF1 | BRF1 RNA polymerase III transcription initiation factor subunit [Source:HGNC Symbol;Acc:HGNC:11551] | -0.8013 | 1.14E-05 |
| ENSG00000119912 | IDE | insulin degrading enzyme [Source:HGNC Symbol;Acc:HGNC:5381] | 0.5637 | 1.15E-05 |
| ENSG00000168913 | ENHO | energy homeostasis associated [Source:HGNC Symbol;Acc:HGNC:24838] | -1.9333 | 1.15E-05 |
| ENSG00000164199 | ADGRV1 | adhesion G protein-coupled receptor V1 [Source:HGNC Symbol;Acc:HGNC:17416] | 0.5010 | 1.15E-05 |
| ENSG00000138604 | GLCE | glucuronic acid epimerase [Source:HGNC Symbol;Acc:HGNC:17855] | -0.5390 | 1.16E-05 |
| ENSG00000130340 | SNX9 | sorting nexin 9 [Source:HGNC Symbol;Acc:HGNC:14973] | 0.5582 | 1.17E-05 |
| ENSG00000262877 | NA | novel transcript | -1.4510 | 1.17E-05 |
| ENSG00000213366 | GSTM2 | glutathione S-transferase mu 2 [Source:HGNC Symbol;Acc:HGNC:4634] | -0.6959 | 1.18E-05 |
| ENSG00000278175 | GLIDR | glioblastoma down-regulated RNA [Source:HGNC Symbol;Acc:HGNC:48823] | 0.7840 | 1.19E-05 |
| ENSG00000260958 | NA | novel transcript | -1.5775 | 1.20E-05 |
| ENSG00000161395 | PGAP3 | post-GPI attachment to proteins phospholipase 3 [Source:HGNC Symbol;Acc:HGNC:23719] | -0.5052 | 1.21E-05 |
| ENSG00000132639 | SNAP25 | synaptosome associated protein 25 [Source:HGNC Symbol;Acc:HGNC:11132] | -0.8046 | 1.21E-05 |
| ENSG00000074800 | ENO1 | enolase 1 [Source:HGNC Symbol;Acc:HGNC:3350] | 0.7884 | 1.22E-05 |
| ENSG00000213742 | ZNF337-AS1 | ZNF337 antisense RNA 1 [Source:HGNC Symbol;Acc:HGNC:40759] | -1.2445 | 1.22E-05 |
| ENSG00000286472 | NA | novel transcript | -1.6995 | 1.22E-05 |
| ENSG00000178338 | ZNF354B | zinc finger protein 354B [Source:HGNC Symbol;Acc:HGNC:17197] | -0.7589 | 1.24E-05 |
| ENSG00000260081 | NA | Morf4 family associated protein 1 (MRFAP1) pseudogene | -1.1355 | 1.24E-05 |

| | | | | |
|---|---|---|---|---|
| ENSG00000185650 | ZFP36L1 | ZFP36 ring finger protein like 1 [Source:HGNC Symbol;Acc:HGNC:1107] | 0.8854 | 1.25E-05 |
| ENSG00000280077 | NA | TEC | -0.9742 | 1.26E-05 |
| ENSG00000173653 | RCE1 | Ras converting CAAX endopeptidase 1 [Source:HGNC Symbol;Acc:HGNC:13721] | 0.5546 | 1.27E-05 |
| ENSG00000168495 | POLR3D | RNA polymerase III subunit D [Source:HGNC Symbol;Acc:HGNC:1080] | 0.5837 | 1.27E-05 |
| ENSG00000257453 | NA | novel transcript, antisense to PHLDA1 | 1.8452 | 1.27E-05 |
| ENSG00000188419 | CHM | CHM Rab escort protein [Source:HGNC Symbol;Acc:HGNC:1940] | 0.5096 | 1.27E-05 |
| ENSG00000226686 | LINC01535 | long intergenic non-protein coding RNA 1535 [Source:HGNC Symbol;Acc:HGNC:51282] | -1.1242 | 1.27E-05 |
| ENSG00000234327 | NA | novel transcript, antisense to ZNF232 | -1.0455 | 1.27E-05 |
| ENSG00000186326 | RGS9BP | regulator of G protein signaling 9 binding protein [Source:HGNC Symbol;Acc:HGNC:30304] | -1.1868 | 1.29E-05 |
| ENSG00000123892 | RAB38 | RAB38, member RAS oncogene family [Source:HGNC Symbol;Acc:HGNC:9776] | 0.8338 | 1.32E-05 |
| ENSG00000163132 | MSX1 | msh homeobox 1 [Source:HGNC Symbol;Acc:HGNC:7391] | -0.7430 | 1.32E-05 |
| ENSG00000159761 | C16orf86 | chromosome 16 open reading frame 86 [Source:HGNC Symbol;Acc:HGNC:33755] | -1.7688 | 1.33E-05 |
| ENSG00000176692 | FOXC2 | forkhead box C2 [Source:HGNC Symbol;Acc:HGNC:3801] | 3.4244 | 1.33E-05 |
| ENSG00000142621 | FHAD1 | forkhead associated phosphopeptide binding domain 1 [Source:HGNC Symbol;Acc:HGNC:29408] | -0.9156 | 1.34E-05 |
| ENSG00000061938 | TNK2 | tyrosine kinase non receptor 2 [Source:HGNC Symbol;Acc:HGNC:19297] | -0.6318 | 1.34E-05 |
| ENSG00000205643 | CDPF1 | cysteine rich DPF motif domain containing 1 [Source:HGNC Symbol;Acc:HGNC:33710] | -1.1027 | 1.34E-05 |
| ENSG00000166997 | CNPY4 | canopy FGF signaling regulator 4 [Source:HGNC Symbol;Acc:HGNC:28631] | -0.6218 | 1.35E-05 |
| ENSG00000223813 | NA | novel transcript, antisense to CHN2 | -1.4250 | 1.35E-05 |
| ENSG00000287104 | NA | novel transcript, antisense to TBC1D14 | -0.8854 | 1.35E-05 |
| ENSG00000083067 | TRPM3 | transient receptor potential cation channel subfamily M member 3 [Source:HGNC Symbol;Acc:HGNC:17992] | -2.0329 | 1.36E-05 |
| ENSG00000182389 | CACNB4 | calcium voltage-gated channel auxiliary subunit beta 4 [Source:HGNC Symbol;Acc:HGNC:1404] | 0.6501 | 1.36E-05 |
| ENSG00000176401 | EID2B | EP300 interacting inhibitor of differentiation 2B [Source:HGNC Symbol;Acc:HGNC:26796] | -0.6089 | 1.39E-05 |
| ENSG00000113231 | PDE8B | phosphodiesterase 8B [Source:HGNC Symbol;Acc:HGNC:8794] | 0.8876 | 1.41E-05 |
| ENSG00000144468 | RHBDD1 | rhomboid domain containing 1 [Source:HGNC Symbol;Acc:HGNC:23081] | 0.7125 | 1.41E-05 |
| ENSG00000007314 | SCN4A | sodium voltage-gated channel alpha subunit 4 [Source:HGNC Symbol;Acc:HGNC:10591] | -1.2264 | 1.43E-05 |
| ENSG00000264772 | NA | novel transcript | 0.6733 | 1.43E-05 |
| ENSG00000164649 | CDCA7L | cell division cycle associated 7 like [Source:HGNC Symbol;Acc:HGNC:30777] | -0.5291 | 1.44E-05 |
| ENSG00000258056 | NA | novel transcript, antisense to CD63 | -0.5564 | 1.44E-05 |
| ENSG00000198814 | GK | glycerol kinase [Source:HGNC Symbol;Acc:HGNC:4289] | 0.6256 | 1.44E-05 |
| ENSG00000112039 | FANCE | FA complementation group E [Source:HGNC Symbol;Acc:HGNC:3586] | -0.7105 | 1.44E-05 |
| ENSG00000267508 | ZNF285 | zinc finger protein 285 [Source:HGNC Symbol;Acc:HGNC:13079] | -1.3757 | 1.45E-05 |
| ENSG00000151014 | NOCT | nocturnin [Source:HGNC Symbol;Acc:HGNC:14254] | 1.1671 | 1.46E-05 |
| ENSG00000164440 | TXLNB | taxilin beta [Source:HGNC Symbol;Acc:HGNC:21617] | -1.2195 | 1.48E-05 |
| ENSG00000151240 | DIP2C | disco interacting protein 2 homolog C [Source:HGNC Symbol;Acc:HGNC:29150] | -0.5496 | 1.50E-05 |
| ENSG00000198517 | MAFK | MAF bZIP transcription factor K [Source:HGNC Symbol;Acc:HGNC:6782] | 0.9095 | 1.52E-05 |
| ENSG00000181467 | RAP2B | RAP2B, member of RAS oncogene family [Source:HGNC Symbol;Acc:HGNC:9862] | -0.5729 | 1.55E-05 |
| ENSG00000142173 | COL6A2 | collagen type VI alpha 2 chain [Source:HGNC Symbol;Acc:HGNC:2212] | 1.0100 | 1.56E-05 |
| ENSG00000197989 | SNHG12 | small nucleolar RNA host gene 12 [Source:HGNC Symbol;Acc:HGNC:30062] | 0.7322 | 1.56E-05 |
| ENSG00000108387 | SEPTIN4 | septin 4 [Source:HGNC Symbol;Acc:HGNC:9165] | -0.8112 | 1.58E-05 |
| ENSG00000131069 | ACSS2 | acyl-CoA synthetase short chain family member 2 [Source:HGNC Symbol;Acc:HGNC:15814] | -0.5460 | 1.58E-05 |
| ENSG00000196205 | EEF1A1P5 | eukaryotic translation elongation factor 1 alpha 1 pseudogene 5 [Source:HGNC Symbol;Acc:HGNC:3200] | 0.5647 | 1.61E-05 |
| ENSG00000232593 | KANTR | KDM5C adjacent transcript [Source:HGNC Symbol;Acc:HGNC:49510] | -0.8366 | 1.61E-05 |
| ENSG00000188603 | CLN3 | CLN3 lysosomal/endosomal transmembrane protein, battenin [Source:HGNC Symbol;Acc:HGNC:2074] | -0.6694 | 1.62E-05 |

| | | | | |
|---|---|---|---|---|
| ENSG00000197177 | ADGRA1 | adhesion G protein-coupled receptor A1 [Source:HGNC Symbol;Acc:HGNC:13838] | 4.7100 | 1.63E-05 |
| ENSG00000105227 | PRX | periaxin [Source:HGNC Symbol;Acc:HGNC:13797] | -0.7236 | 1.63E-05 |
| ENSG00000169740 | ZNF32 | zinc finger protein 32 [Source:HGNC Symbol;Acc:HGNC:13095] | -0.6318 | 1.71E-05 |
| ENSG00000204388 | HSPA1B | heat shock protein family A (Hsp70) member 1B [Source:HGNC Symbol;Acc:HGNC:5233] | -0.7427 | 1.71E-05 |
| ENSG00000224616 | RTCA-AS1 | RTCA antisense RNA 1 [Source:HGNC Symbol;Acc:HGNC:50573] | -1.4056 | 1.75E-05 |
| ENSG00000107954 | NEURL1 | neuralized E3 ubiquitin protein ligase 1 [Source:HGNC Symbol;Acc:HGNC:7761] | 5.0216 | 1.76E-05 |
| ENSG00000261456 | TUBB8 | tubulin beta 8 class VIII [Source:HGNC Symbol;Acc:HGNC:20773] | -2.0760 | 1.76E-05 |
| ENSG00000103319 | EEF2K | eukaryotic elongation factor 2 kinase [Source:HGNC Symbol;Acc:HGNC:24615] | -0.6176 | 1.83E-05 |
| ENSG00000115267 | IFIH1 | interferon induced with helicase C domain 1 [Source:HGNC Symbol;Acc:HGNC:18873] | -0.6378 | 1.83E-05 |
| ENSG00000136197 | C7orf25 | chromosome 7 open reading frame 25 [Source:HGNC Symbol;Acc:HGNC:21703] | -0.8370 | 1.83E-05 |
| ENSG00000270977 | NA | leucine rich repeat containing 37 member A3 (LRRC37A3) pseudogene | -2.0481 | 1.85E-05 |
| ENSG00000224888 | NA | novel transcript, antisense to FAM38A | 1.9505 | 1.89E-05 |
| ENSG00000128683 | GAD1 | glutamate decarboxylase 1 [Source:HGNC Symbol;Acc:HGNC:4092] | 0.6857 | 1.90E-05 |
| ENSG00000162408 | NOL9 | nucleolar protein 9 [Source:HGNC Symbol;Acc:HGNC:26265] | 0.6828 | 1.90E-05 |
| ENSG00000164946 | FREM1 | FRAS1 related extracellular matrix 1 [Source:HGNC Symbol;Acc:HGNC:23399] | -1.1538 | 1.90E-05 |
| ENSG00000253276 | CCDC71L | coiled-coil domain containing 71 like [Source:HGNC Symbol;Acc:HGNC:26685] | -0.5492 | 1.94E-05 |
| ENSG00000084628 | NKAIN1 | sodium/potassium transporting ATPase interacting 1 [Source:HGNC Symbol;Acc:HGNC:25743] | 2.6989 | 1.94E-05 |
| ENSG00000125731 | SH2D3A | SH2 domain containing 3A [Source:HGNC Symbol;Acc:HGNC:16885] | 0.7946 | 1.95E-05 |
| ENSG00000223653 | NA | novel transcript | 1.8263 | 1.97E-05 |
| ENSG00000156531 | PHF6 | PHD finger protein 6 [Source:HGNC Symbol;Acc:HGNC:18145] | 0.6687 | 1.98E-05 |
| ENSG00000115204 | MPV17 | mitochondrial inner membrane protein MPV17 [Source:HGNC Symbol;Acc:HGNC:7224] | 0.6019 | 1.98E-05 |
| ENSG00000221869 | CEBPD | CCAAT enhancer binding protein delta [Source:HGNC Symbol;Acc:HGNC:1835] | 1.5719 | 1.98E-05 |
| ENSG00000160191 | PDE9A | phosphodiesterase 9A [Source:HGNC Symbol;Acc:HGNC:8795] | -0.7830 | 1.98E-05 |
| ENSG00000197961 | ZNF121 | zinc finger protein 121 [Source:HGNC Symbol;Acc:HGNC:12904] | 0.5655 | 1.98E-05 |
| ENSG00000131480 | AOC2 | amine oxidase copper containing 2 [Source:HGNC Symbol;Acc:HGNC:549] | 1.1043 | 1.98E-05 |
| ENSG00000265787 | CYP4F35P | cytochrome P450 family 4 subfamily F member 35, pseudogene [Source:HGNC Symbol;Acc:HGNC:39954] | -2.1172 | 1.99E-05 |
| ENSG00000261488 | TBILA | TGF-beta induced lncRNA [Source:HGNC Symbol;Acc:HGNC:53943] | 0.9666 | 2.01E-05 |
| ENSG00000235173 | HGH1 | HGH1 homolog [Source:HGNC Symbol;Acc:HGNC:24161] | -0.5239 | 2.02E-05 |
| ENSG00000230551 | NA | novel transcript | 0.7116 | 2.02E-05 |
| ENSG00000144040 | SFXN5 | sideroflexin 5 [Source:HGNC Symbol;Acc:HGNC:16073] | 0.5765 | 2.03E-05 |
| ENSG00000140022 | STON2 | stonin 2 [Source:HGNC Symbol;Acc:HGNC:30652] | 0.6362 | 2.12E-05 |
| ENSG00000101974 | ATP11C | ATPase phospholipid transporting 11C [Source:HGNC Symbol;Acc:HGNC:13554] | 0.6829 | 2.14E-05 |
| ENSG00000159247 | TUBBP5 | tubulin beta pseudogene 5 [Source:HGNC Symbol;Acc:HGNC:23674] | -3.0072 | 2.15E-05 |
| ENSG00000184408 | KCND2 | potassium voltage-gated channel subfamily D member 2 [Source:HGNC Symbol;Acc:HGNC:6238] | 4.3422 | 2.16E-05 |
| ENSG00000185133 | INPP5J | inositol polyphosphate-5-phosphatase J [Source:HGNC Symbol;Acc:HGNC:8956] | -0.5469 | 2.20E-05 |
| ENSG00000137700 | SLC37A4 | solute carrier family 37 member 4 [Source:HGNC Symbol;Acc:HGNC:4061] | -0.5740 | 2.20E-05 |
| ENSG00000283050 | GTF2IP12 | general transcription factor IIi pseudogene 12 [Source:HGNC Symbol;Acc:HGNC:51723] | -0.8451 | 2.22E-05 |
| ENSG00000120885 | CLU | clusterin [Source:HGNC Symbol;Acc:HGNC:2095] | 0.5114 | 2.22E-05 |
| ENSG00000139914 | FITM1 | fat storage inducing transmembrane protein 1 [Source:HGNC Symbol;Acc:HGNC:33714] | 0.7904 | 2.23E-05 |
| ENSG00000153066 | TXNDC11 | thioredoxin domain containing 11 [Source:HGNC Symbol;Acc:HGNC:28030] | 0.5275 | 2.23E-05 |
| ENSG00000143793 | C1orf35 | chromosome 1 open reading frame 35 [Source:HGNC Symbol;Acc:HGNC:19032] | -0.5111 | 2.23E-05 |
| ENSG00000134871 | COL4A2 | collagen type IV alpha 2 chain [Source:HGNC Symbol;Acc:HGNC:2203] | 0.5599 | 2.25E-05 |

| | | | | |
|---|---|---|---|---|
| ENSG00000180537 | RNF182 | ring finger protein 182 [Source:HGNC Symbol;Acc:HGNC:28522] | -0.7656 | 2.27E-05 |
| ENSG00000213853 | EMP2 | epithelial membrane protein 2 [Source:HGNC Symbol;Acc:HGNC:3334] | -0.5995 | 2.27E-05 |
| ENSG00000138496 | PARP9 | poly(ADP-ribose) polymerase family member 9 [Source:HGNC Symbol;Acc:HGNC:24118] | -0.6366 | 2.29E-05 |
| ENSG00000224790 | NA | novel transcript | 0.8664 | 2.32E-05 |
| ENSG00000117245 | KIF17 | kinesin family member 17 [Source:HGNC Symbol;Acc:HGNC:19167] | 0.5315 | 2.33E-05 |
| ENSG00000152503 | TRIM36 | tripartite motif containing 36 [Source:HGNC Symbol;Acc:HGNC:16280] | -0.8230 | 2.33E-05 |
| ENSG00000141295 | SCRN2 | secernin 2 [Source:HGNC Symbol;Acc:HGNC:30381] | -0.5225 | 2.34E-05 |
| ENSG00000134532 | SOX5 | SRY-box transcription factor 5 [Source:HGNC Symbol;Acc:HGNC:11201] | 0.6883 | 2.34E-05 |
| ENSG00000129910 | CDH15 | cadherin 15 [Source:HGNC Symbol;Acc:HGNC:1754] | 3.3429 | 2.34E-05 |
| ENSG00000261340 | LINC01616 | long intergenic non-protein coding RNA 1616 [Source:HGNC Symbol;Acc:HGNC:51900] | 1.5436 | 2.37E-05 |
| ENSG00000225648 | SBDSP1 | SBDS pseudogene 1 [Source:HGNC Symbol;Acc:HGNC:21646] | 0.6577 | 2.37E-05 |
| ENSG00000133818 | RRAS2 | RAS related 2 [Source:HGNC Symbol;Acc:HGNC:17271] | -0.7366 | 2.37E-05 |
| ENSG00000070540 | WIPI1 | WD repeat domain, phosphoinositide interacting 1 [Source:HGNC Symbol;Acc:HGNC:25471] | 0.7414 | 2.38E-05 |
| ENSG00000130714 | POMT1 | protein O-mannosyltransferase 1 [Source:HGNC Symbol;Acc:HGNC:9202] | -0.5840 | 2.42E-05 |
| ENSG00000141447 | OSBPL1A | oxysterol binding protein like 1A [Source:HGNC Symbol;Acc:HGNC:16398] | -0.6405 | 2.42E-05 |
| ENSG00000163354 | DCST2 | DC-STAMP domain containing 2 [Source:HGNC Symbol;Acc:HGNC:26562] | -2.0267 | 2.43E-05 |
| ENSG00000143224 | PPOX | protoporphyrinogen oxidase [Source:HGNC Symbol;Acc:HGNC:9280] | -0.6266 | 2.46E-05 |
| ENSG00000258441 | LINC00641 | long intergenic non-protein coding RNA 641 [Source:HGNC Symbol;Acc:HGNC:27511] | 0.5973 | 2.46E-05 |
| ENSG00000272501 | NA | novel transcript, antisense to HCG27 | -1.0916 | 2.47E-05 |
| ENSG00000181873 | IBA57 | iron-sulfur cluster assembly factor IBA57 [Source:HGNC Symbol;Acc:HGNC:27302] | 0.5826 | 2.47E-05 |
| ENSG00000177932 | ZNF354C | zinc finger protein 354C [Source:HGNC Symbol;Acc:HGNC:16736] | -0.8713 | 2.48E-05 |
| ENSG00000163481 | RNF25 | ring finger protein 25 [Source:HGNC Symbol;Acc:HGNC:14662] | 0.5741 | 2.48E-05 |
| ENSG00000134602 | STK26 | serine/threonine kinase 26 [Source:HGNC Symbol;Acc:HGNC:18174] | 0.6816 | 2.50E-05 |
| ENSG00000185519 | FAM131C | family with sequence similarity 131 member C [Source:HGNC Symbol;Acc:HGNC:26717] | 0.7382 | 2.54E-05 |
| ENSG00000126215 | XRCC3 | X-ray repair cross complementing 3 [Source:HGNC Symbol;Acc:HGNC:12830] | -0.8101 | 2.56E-05 |
| ENSG00000138160 | KIF11 | kinesin family member 11 [Source:HGNC Symbol;Acc:HGNC:6388] | -0.5457 | 2.57E-05 |
| ENSG00000186395 | KRT10 | keratin 10 [Source:HGNC Symbol;Acc:HGNC:6413] | 0.5355 | 2.57E-05 |
| ENSG00000251580 | LINC02482 | long intergenic non-protein coding RNA 2482 [Source:HGNC Symbol;Acc:HGNC:53458] | -0.9168 | 2.62E-05 |
| ENSG00000103550 | KNOP1 | lysine rich nucleolar protein 1 [Source:HGNC Symbol;Acc:HGNC:34404] | -0.5424 | 2.62E-05 |
| ENSG00000116584 | ARHGEF2 | Rho/Rac guanine nucleotide exchange factor 2 [Source:HGNC Symbol;Acc:HGNC:682] | 0.6866 | 2.63E-05 |
| ENSG00000225760 | LINC00431 | long intergenic non-protein coding RNA 431 [Source:HGNC Symbol;Acc:HGNC:42766] | 5.8412 | 2.63E-05 |
| ENSG00000251562 | MALAT1 | metastasis associated lung adenocarcinoma transcript 1 [Source:HGNC Symbol;Acc:HGNC:29665] | 0.8215 | 2.65E-05 |
| ENSG00000133247 | KMT5C | lysine methyltransferase 5C [Source:HGNC Symbol;Acc:HGNC:28405] | -0.6331 | 2.68E-05 |
| ENSG00000156050 | FAM161B | FAM161 centrosomal protein B [Source:HGNC Symbol;Acc:HGNC:19854] | -1.4403 | 2.72E-05 |
| ENSG00000229589 | ACVR2B-AS1 | ACVR2B antisense RNA 1 [Source:HGNC Symbol;Acc:HGNC:44161] | -0.9606 | 2.73E-05 |
| ENSG00000120129 | DUSP1 | dual specificity phosphatase 1 [Source:HGNC Symbol;Acc:HGNC:3064] | 0.9925 | 2.74E-05 |
| ENSG00000102189 | EEA1 | early endosome antigen 1 [Source:HGNC Symbol;Acc:HGNC:3185] | 0.8726 | 2.75E-05 |
| ENSG00000114626 | ABTB1 | ankyrin repeat and BTB domain containing 1 [Source:HGNC Symbol;Acc:HGNC:18275] | -1.3833 | 2.75E-05 |
| ENSG00000177374 | HIC1 | HIC ZBTB transcriptional repressor 1 [Source:HGNC Symbol;Acc:HGNC:4909] | 0.8126 | 2.77E-05 |
| ENSG00000162594 | IL23R | interleukin 23 receptor [Source:HGNC Symbol;Acc:HGNC:19100] | 0.5699 | 2.78E-05 |
| ENSG00000138771 | SHROOM3 | shroom family member 3 [Source:HGNC Symbol;Acc:HGNC:30422] | -0.6075 | 2.80E-05 |
| ENSG00000233237 | LINC00472 | long intergenic non-protein coding RNA 472 [Source:HGNC Symbol;Acc:HGNC:21380] | -1.0944 | 2.82E-05 |
| ENSG00000118985 | ELL2 | elongation factor for RNA polymerase II 2 [Source:HGNC Symbol;Acc:HGNC:17064] | -0.6759 | 2.84E-05 |

| | | | | |
|---|---|---|---|---|
| ENSG00000177679 | SRRM3 | serine/arginine repetitive matrix 3 [Source:HGNC Symbol;Acc:HGNC:26729] | -0.5610 | 2.84E-05 |
| ENSG00000261325 | LINC02192 | long intergenic non-protein coding RNA 2192 [Source:HGNC Symbol;Acc:HGNC:53054] | -1.1338 | 2.85E-05 |
| ENSG00000116574 | RHOU | ras homolog family member U [Source:HGNC Symbol;Acc:HGNC:17794] | 0.8197 | 2.85E-05 |
| ENSG00000129682 | FGF13 | fibroblast growth factor 13 [Source:HGNC Symbol;Acc:HGNC:3670] | -0.5542 | 2.87E-05 |
| ENSG00000117305 | HMGCL | 3-hydroxy-3-methylglutaryl-CoA lyase [Source:HGNC Symbol;Acc:HGNC:5005] | -0.7445 | 2.88E-05 |
| ENSG00000228623 | ZNF883 | zinc finger protein 883 [Source:HGNC Symbol;Acc:HGNC:27271] | -0.9971 | 2.88E-05 |
| ENSG00000176974 | SHMT1 | serine hydroxymethyltransferase 1 [Source:HGNC Symbol;Acc:HGNC:10850] | -0.5366 | 2.95E-05 |
| ENSG00000143147 | GPR161 | G protein-coupled receptor 161 [Source:HGNC Symbol;Acc:HGNC:23694] | -0.7126 | 2.96E-05 |
| ENSG00000180044 | C3orf80 | chromosome 3 open reading frame 80 [Source:HGNC Symbol;Acc:HGNC:40048] | 1.3041 | 2.96E-05 |
| ENSG00000167646 | DNAAF3 | dynein axonemal assembly factor 3 [Source:HGNC Symbol;Acc:HGNC:30492] | -0.8052 | 2.98E-05 |
| ENSG00000147465 | STAR | steroidogenic acute regulatory protein [Source:HGNC Symbol;Acc:HGNC:11359] | 1.2116 | 3.01E-05 |
| ENSG00000204866 | IGFL2 | IGF like family member 2 [Source:HGNC Symbol;Acc:HGNC:32929] | 4.2202 | 3.02E-05 |
| ENSG00000118307 | CASC1 | cancer susceptibility 1 [Source:HGNC Symbol;Acc:HGNC:29599] | -2.0836 | 3.03E-05 |
| ENSG00000173801 | JUP | junction plakoglobin [Source:HGNC Symbol;Acc:HGNC:6207] | -0.7516 | 3.10E-05 |
| ENSG00000091844 | RGS17 | regulator of G protein signaling 17 [Source:HGNC Symbol;Acc:HGNC:14088] | -1.0179 | 3.11E-05 |
| ENSG00000070371 | CLTCL1 | clathrin heavy chain like 1 [Source:HGNC Symbol;Acc:HGNC:2093] | -0.7696 | 3.13E-05 |
| ENSG00000080608 | PUM3 | pumilio RNA binding family member 3 [Source:HGNC Symbol;Acc:HGNC:29676] | 0.5708 | 3.18E-05 |
| ENSG00000272341 | NA | novel transcript | -2.3952 | 3.20E-05 |
| ENSG00000215817 | ZC3H11B | zinc finger CCCH-type containing 11B [Source:HGNC Symbol;Acc:HGNC:25659] | 0.8990 | 3.21E-05 |
| ENSG00000243649 | CFB | complement factor B [Source:HGNC Symbol;Acc:HGNC:1037] | 1.7674 | 3.23E-05 |
| ENSG00000183691 | NOG | noggin [Source:HGNC Symbol;Acc:HGNC:7866] | -3.5413 | 3.23E-05 |
| ENSG00000185800 | DMWD | DM1 locus, WD repeat containing [Source:HGNC Symbol;Acc:HGNC:2936] | 0.5332 | 3.24E-05 |
| ENSG00000187867 | PALM3 | paralemmin 3 [Source:HGNC Symbol;Acc:HGNC:33274] | -1.2259 | 3.25E-05 |
| ENSG00000254319 | NA | novel transcript | -0.6830 | 3.25E-05 |
| ENSG00000284968 | NA | Novel transcript, antisense to AFF1 | -1.1423 | 3.26E-05 |
| ENSG00000184939 | ZFP90 | ZFP90 zinc finger protein [Source:HGNC Symbol;Acc:HGNC:23329] | -0.9130 | 3.27E-05 |
| ENSG00000266338 | NBPF15 | NBPF member 15 [Source:HGNC Symbol;Acc:HGNC:28791] | -0.5821 | 3.29E-05 |
| ENSG00000023572 | GLRX2 | glutaredoxin 2 [Source:HGNC Symbol;Acc:HGNC:16065] | 0.5634 | 3.31E-05 |
| ENSG00000183963 | SMTN | smoothelin [Source:HGNC Symbol;Acc:HGNC:11126] | -0.7782 | 3.35E-05 |
| ENSG00000286839 | NA | novel transcript | -1.7279 | 3.37E-05 |
| ENSG00000142675 | CNKSR1 | connector enhancer of kinase suppressor of Ras 1 [Source:HGNC Symbol;Acc:HGNC:19700] | 0.9029 | 3.38E-05 |
| ENSG00000117877 | CD3EAP | CD3e molecule associated protein [Source:HGNC Symbol;Acc:HGNC:24219] | 0.6440 | 3.42E-05 |
| ENSG00000230257 | NFE4 | nuclear factor, erythroid 4 [Source:HGNC Symbol;Acc:HGNC:29902] | 0.8592 | 3.47E-05 |
| ENSG00000125898 | FAM110A | family with sequence similarity 110 member A [Source:HGNC Symbol;Acc:HGNC:16188] | -0.7307 | 3.47E-05 |
| ENSG00000173838 | MARCHF10 | membrane associated ring-CH-type finger 10 [Source:HGNC Symbol;Acc:HGNC:26655] | 1.4075 | 3.51E-05 |
| ENSG00000113296 | THBS4 | thrombospondin 4 [Source:HGNC Symbol;Acc:HGNC:11788] | -0.9446 | 3.51E-05 |
| ENSG00000169131 | ZNF354A | zinc finger protein 354A [Source:HGNC Symbol;Acc:HGNC:11628] | -0.6215 | 3.54E-05 |
| ENSG00000180979 | LRRC57 | leucine rich repeat containing 57 [Source:HGNC Symbol;Acc:HGNC:26719] | -0.5257 | 3.57E-05 |
| ENSG00000058091 | CDK14 | cyclin dependent kinase 14 [Source:HGNC Symbol;Acc:HGNC:8883] | 1.2367 | 3.59E-05 |
| ENSG00000235026 | DPP10-AS1 | DPP10 antisense RNA 1 [Source:HGNC Symbol;Acc:HGNC:40941] | -0.6250 | 3.60E-05 |
| ENSG00000107130 | NCS1 | neuronal calcium sensor 1 [Source:HGNC Symbol;Acc:HGNC:3953] | 0.5874 | 3.65E-05 |
| ENSG00000162944 | RFTN2 | raftlin family member 2 [Source:HGNC Symbol;Acc:HGNC:26402] | 1.1486 | 3.66E-05 |
| ENSG00000165757 | JCAD | junctional cadherin 5 associated [Source:HGNC Symbol;Acc:HGNC:29283] | -1.2108 | 3.68E-05 |

| | | | | |
|---|---|---|---|---|
| ENSG00000272636 | DOC2B | double C2 domain beta [Source:HGNC Symbol;Acc:HGNC:2986] | 1.7757 | 3.69E-05 |
| ENSG00000171368 | TPPP | tubulin polymerization promoting protein [Source:HGNC Symbol;Acc:HGNC:24164] | -0.9548 | 3.71E-05 |
| ENSG00000069424 | KCNAB2 | potassium voltage-gated channel subfamily A regulatory beta subunit 2 [Source:HGNC Symbol;Acc:HGNC:6229] | -0.8126 | 3.75E-05 |
| ENSG00000152382 | TADA1 | transcriptional adaptor 1 [Source:HGNC Symbol;Acc:HGNC:30631] | -0.5471 | 3.76E-05 |
| ENSG00000154920 | EME1 | essential meiotic structure-specific endonuclease 1 [Source:HGNC Symbol;Acc:HGNC:24965] | -0.7372 | 3.81E-05 |
| ENSG00000287856 | NA | novel protein | -24.1629 | 3.81E-05 |
| ENSG00000215067 | ALOX12-AS1 | ALOX12 antisense RNA 1 [Source:HGNC Symbol;Acc:HGNC:51342] | -0.7350 | 3.85E-05 |
| ENSG00000001460 | STPG1 | sperm tail PG-rich repeat containing 1 [Source:HGNC Symbol;Acc:HGNC:28070] | -0.6015 | 3.86E-05 |
| ENSG00000170385 | SLC30A1 | solute carrier family 30 member 1 [Source:HGNC Symbol;Acc:HGNC:11012] | 0.5197 | 3.86E-05 |
| ENSG00000181908 | LINC02724 | long intergenic non-protein coding RNA 2724 [Source:HGNC Symbol;Acc:HGNC:54241] | -0.9860 | 3.87E-05 |
| ENSG00000196263 | ZNF471 | zinc finger protein 471 [Source:HGNC Symbol;Acc:HGNC:23226] | -0.7860 | 3.88E-05 |
| ENSG00000166839 | ANKDD1A | ankyrin repeat and death domain containing 1A [Source:HGNC Symbol;Acc:HGNC:28002] | -0.7006 | 3.90E-05 |
| ENSG00000100744 | GSKIP | GSK3B interacting protein [Source:HGNC Symbol;Acc:HGNC:20343] | 0.5941 | 3.94E-05 |
| ENSG00000246985 | SOCS2-AS1 | SOCS2 antisense RNA 1 [Source:HGNC Symbol;Acc:HGNC:27054] | 3.3307 | 3.95E-05 |
| ENSG00000175567 | UCP2 | uncoupling protein 2 [Source:HGNC Symbol;Acc:HGNC:12518] | -0.8872 | 3.97E-05 |
| ENSG00000141480 | ARRB2 | arrestin beta 2 [Source:HGNC Symbol;Acc:HGNC:712] | -0.5637 | 3.98E-05 |
| ENSG00000257704 | INAFM1 | InaF motif containing 1 [Source:HGNC Symbol;Acc:HGNC:27406] | 0.8778 | 3.98E-05 |
| ENSG00000265666 | RARA-AS1 | RARA antisense RNA 1 [Source:HGNC Symbol;Acc:HGNC:49577] | 1.3557 | 4.04E-05 |
| ENSG00000075891 | PAX2 | paired box 2 [Source:HGNC Symbol;Acc:HGNC:8616] | -1.1754 | 4.05E-05 |
| ENSG00000232712 | KIZ-AS1 | KIZ antisense RNA 1 [Source:HGNC Symbol;Acc:HGNC:51231] | 3.4934 | 4.08E-05 |
| ENSG00000108771 | DHX58 | DExH-box helicase 58 [Source:HGNC Symbol;Acc:HGNC:29517] | -0.9833 | 4.08E-05 |
| ENSG00000151665 | PIGF | phosphatidylinositol glycan anchor biosynthesis class F [Source:HGNC Symbol;Acc:HGNC:8962] | -0.5581 | 4.08E-05 |
| ENSG00000176896 | TCEANC | transcription elongation factor A N-terminal and central domain containing [Source:HGNC Symbol;Acc:HGNC:28277] | -1.6399 | 4.10E-05 |
| ENSG00000125895 | TMEM74B | transmembrane protein 74B [Source:HGNC Symbol;Acc:HGNC:15893] | -1.0569 | 4.11E-05 |
| ENSG00000164188 | RANBP3L | RAN binding protein 3 like [Source:HGNC Symbol;Acc:HGNC:26353] | -0.7109 | 4.13E-05 |
| ENSG00000259024 | TVP23C-CDRT4 | TVP23C-CDRT4 readthrough [Source:HGNC Symbol;Acc:HGNC:42961] | 0.6331 | 4.14E-05 |
| ENSG00000184634 | MED12 | mediator complex subunit 12 [Source:HGNC Symbol;Acc:HGNC:11957] | -0.5523 | 4.17E-05 |
| ENSG00000124429 | POF1B | POF1B actin binding protein [Source:HGNC Symbol;Acc:HGNC:13711] | -0.6268 | 4.18E-05 |
| ENSG00000135778 | NTPCR | nucleoside-triphosphatase, cancer-related [Source:HGNC Symbol;Acc:HGNC:28204] | -0.5228 | 4.18E-05 |
| ENSG00000171227 | TMEM37 | transmembrane protein 37 [Source:HGNC Symbol;Acc:HGNC:18216] | -1.2739 | 4.19E-05 |
| ENSG00000156508 | EEF1A1 | eukaryotic translation elongation factor 1 alpha 1 [Source:HGNC Symbol;Acc:HGNC:3189] | 0.5113 | 4.20E-05 |
| ENSG00000101298 | SNPH | syntaphilin [Source:HGNC Symbol;Acc:HGNC:15931] | -0.6785 | 4.20E-05 |
| ENSG00000133740 | E2F5 | E2F transcription factor 5 [Source:HGNC Symbol;Acc:HGNC:3119] | 0.6502 | 4.20E-05 |
| ENSG00000164010 | ERMAP | erythroblast membrane associated protein (Scianna blood group) [Source:HGNC Symbol;Acc:HGNC:15743] | -0.5983 | 4.23E-05 |
| ENSG00000160072 | ATAD3B | ATPase family AAA domain containing 3B [Source:HGNC Symbol;Acc:HGNC:24007] | 0.7310 | 4.30E-05 |
| ENSG00000217801 | NA | protein tyrosine phosphatase family pseudogene | 3.5875 | 4.31E-05 |
| ENSG00000123607 | TTC21B | tetratricopeptide repeat domain 21B [Source:HGNC Symbol;Acc:HGNC:25660] | -0.5957 | 4.32E-05 |
| ENSG00000180425 | C11orf71 | chromosome 11 open reading frame 71 [Source:HGNC Symbol;Acc:HGNC:25937] | -0.6788 | 4.34E-05 |
| ENSG00000183323 | CCDC125 | coiled-coil domain containing 125 [Source:HGNC Symbol;Acc:HGNC:28924] | -0.5025 | 4.34E-05 |
| ENSG00000197580 | BCO2 | beta-carotene oxygenase 2 [Source:HGNC Symbol;Acc:HGNC:18503] | 0.6826 | 4.35E-05 |
| ENSG00000226688 | ENTPD1-AS1 | ENTPD1 antisense RNA 1 [Source:HGNC Symbol;Acc:HGNC:45203] | 0.5159 | 4.43E-05 |
| ENSG00000172828 | CES3 | carboxylesterase 3 [Source:HGNC Symbol;Acc:HGNC:1865] | -0.9331 | 4.44E-05 |

| | | | | |
|---|---|---|---|---|
| ENSG00000138758 | SEPTIN11 | septin 11 [Source:HGNC Symbol;Acc:HGNC:25589] | 0.6042 | 4.53E-05 |
| ENSG00000100226 | GTPBP1 | GTP binding protein 1 [Source:HGNC Symbol;Acc:HGNC:4669] | 0.5801 | 4.54E-05 |
| ENSG00000240291 | NA | novel transcript, antisense to CACNB2 | 1.2679 | 4.56E-05 |
| ENSG00000151612 | ZNF827 | zinc finger protein 827 [Source:HGNC Symbol;Acc:HGNC:27193] | 0.5024 | 4.58E-05 |
| ENSG00000226200 | SGMS1-AS1 | SGMS1 antisense RNA 1 [Source:HGNC Symbol;Acc:HGNC:49683] | 1.0686 | 4.64E-05 |
| ENSG00000185551 | NR2F2 | nuclear receptor subfamily 2 group F member 2 [Source:HGNC Symbol;Acc:HGNC:7976] | -0.5984 | 4.74E-05 |
| ENSG00000178425 | NT5DC1 | 5'-nucleotidase domain containing 1 [Source:HGNC Symbol;Acc:HGNC:21556] | -0.5911 | 4.77E-05 |
| ENSG00000037897 | METTL1 | methyltransferase like 1 [Source:HGNC Symbol;Acc:HGNC:7030] | 0.6792 | 4.79E-05 |
| ENSG00000196693 | ZNF33B | zinc finger protein 33B [Source:HGNC Symbol;Acc:HGNC:13097] | -0.6134 | 4.79E-05 |
| ENSG00000089847 | ANKRD24 | ankyrin repeat domain 24 [Source:HGNC Symbol;Acc:HGNC:29424] | -1.2300 | 4.81E-05 |
| ENSG00000214106 | PAXIP1-AS2 | PAXIP1 antisense RNA 2 [Source:HGNC Symbol;Acc:HGNC:48958] | -0.7067 | 4.82E-05 |
| ENSG00000116191 | RALGPS2 | Ral GEF with PH domain and SH3 binding motif 2 [Source:HGNC Symbol;Acc:HGNC:30279] | -0.6605 | 4.84E-05 |
| ENSG00000180758 | GPR157 | G protein-coupled receptor 157 [Source:HGNC Symbol;Acc:HGNC:23687] | -1.0034 | 4.88E-05 |
| ENSG00000286699 | NA | novel transcript | -0.8572 | 4.89E-05 |
| ENSG00000162542 | TMCO4 | transmembrane and coiled-coil domains 4 [Source:HGNC Symbol;Acc:HGNC:27393] | -0.5570 | 4.91E-05 |
| ENSG00000284669 | NA | novel transcript, antisense CSRNP1 | 3.1455 | 4.91E-05 |
| ENSG00000196756 | SNHG17 | small nucleolar RNA host gene 17 [Source:HGNC Symbol;Acc:HGNC:48600] | 0.7044 | 4.97E-05 |
| ENSG00000196670 | ZFP62 | ZFP62 zinc finger protein [Source:HGNC Symbol;Acc:HGNC:23241] | -0.5344 | 4.99E-05 |
| ENSG00000130413 | STK33 | serine/threonine kinase 33 [Source:HGNC Symbol;Acc:HGNC:14568] | -0.7149 | 5.01E-05 |
| ENSG00000065989 | PDE4A | phosphodiesterase 4A [Source:HGNC Symbol;Acc:HGNC:8780] | 1.7930 | 5.02E-05 |
| ENSG00000054598 | FOXC1 | forkhead box C1 [Source:HGNC Symbol;Acc:HGNC:3800] | 0.9724 | 5.02E-05 |
| ENSG00000106804 | C5 | complement C5 [Source:HGNC Symbol;Acc:HGNC:1331] | -0.7572 | 5.05E-05 |
| ENSG00000236432 | MFF-DT | MFF divergent transcript [Source:HGNC Symbol;Acc:HGNC:41067] | -0.8283 | 5.07E-05 |
| ENSG00000088387 | DOCK9 | dedicator of cytokinesis 9 [Source:HGNC Symbol;Acc:HGNC:14132] | -0.6021 | 5.10E-05 |
| ENSG00000287279 | NA | novel transcript | -2.4199 | 5.16E-05 |
| ENSG00000160801 | PTH1R | parathyroid hormone 1 receptor [Source:HGNC Symbol;Acc:HGNC:9608] | -2.3117 | 5.19E-05 |
| ENSG00000167136 | ENDOG | endonuclease G [Source:HGNC Symbol;Acc:HGNC:3346] | -0.5683 | 5.20E-05 |
| ENSG00000141750 | STAC2 | SH3 and cysteine rich domain 2 [Source:HGNC Symbol;Acc:HGNC:23990] | -1.0386 | 5.26E-05 |
| ENSG00000285877 | NA | novel transcript, antisense to HELZ | 3.0833 | 5.29E-05 |
| ENSG00000232618 | NA | novel transcript | 3.9527 | 5.33E-05 |
| ENSG00000186889 | TMEM17 | transmembrane protein 17 [Source:HGNC Symbol;Acc:HGNC:26623] | -1.0382 | 5.36E-05 |
| ENSG00000223802 | CERS1 | ceramide synthase 1 [Source:HGNC Symbol;Acc:HGNC:14253] | -0.6029 | 5.36E-05 |
| ENSG00000235954 | TTC28-AS1 | TTC28 antisense RNA 1 [Source:HGNC Symbol;Acc:HGNC:29336] | -0.6420 | 5.45E-05 |
| ENSG00000183091 | NEB | nebulin [Source:HGNC Symbol;Acc:HGNC:7720] | 1.3800 | 5.51E-05 |
| ENSG00000265933 | LINC00668 | long intergenic non-protein coding RNA 668 [Source:HGNC Symbol;Acc:HGNC:44328] | -0.8905 | 5.52E-05 |
| ENSG00000139910 | NOVA1 | NOVA alternative splicing regulator 1 [Source:HGNC Symbol;Acc:HGNC:7886] | -0.6390 | 5.53E-05 |
| ENSG00000196220 | SRGAP3 | SLIT-ROBO Rho GTPase activating protein 3 [Source:HGNC Symbol;Acc:HGNC:19744] | -1.2331 | 5.59E-05 |
| ENSG00000116353 | MECR | mitochondrial trans-2-enoyl-CoA reductase [Source:HGNC Symbol;Acc:HGNC:19691] | -0.6779 | 5.60E-05 |
| ENSG00000267009 | NA | novel transcript | 1.3590 | 5.65E-05 |
| ENSG00000146592 | CREB5 | cAMP responsive element binding protein 5 [Source:HGNC Symbol;Acc:HGNC:16844] | 1.5500 | 5.66E-05 |
| ENSG00000167393 | PPP2R3B | protein phosphatase 2 regulatory subunit B''beta [Source:HGNC Symbol;Acc:HGNC:13417] | -1.1639 | 5.67E-05 |
| ENSG00000159259 | CHAF1B | chromatin assembly factor 1 subunit B [Source:HGNC Symbol;Acc:HGNC:1911] | -0.8421 | 5.70E-05 |
| ENSG00000168234 | TTC39C | tetratricopeptide repeat domain 39C [Source:HGNC Symbol;Acc:HGNC:26595] | 0.5908 | 5.77E-05 |

| | | | | |
|---|---|---|---|---|
| ENSG00000180340 | FZD2 | frizzled class receptor 2 [Source:HGNC Symbol;Acc:HGNC:4040] | -0.6705 | 5.78E-05 |
| ENSG00000172006 | ZNF554 | zinc finger protein 554 [Source:HGNC Symbol;Acc:HGNC:26629] | -1.0041 | 5.80E-05 |
| ENSG00000173218 | VANGL1 | VANGL planar cell polarity protein 1 [Source:HGNC Symbol;Acc:HGNC:15512] | -0.6963 | 5.80E-05 |
| ENSG00000085117 | CD82 | CD82 molecule [Source:HGNC Symbol;Acc:HGNC:6210] | 0.8480 | 5.81E-05 |
| ENSG00000162066 | AMDHD2 | amidohydrolase domain containing 2 [Source:HGNC Symbol;Acc:HGNC:24262] | -0.6155 | 5.81E-05 |
| ENSG00000259828 | NA | novel transcript | 1.0357 | 5.87E-05 |
| ENSG00000153936 | HS2ST1 | heparan sulfate 2-O-sulfotransferase 1 [Source:HGNC Symbol;Acc:HGNC:5193] | -0.5371 | 5.88E-05 |
| ENSG00000162769 | FLVCR1 | FLVCR heme transporter 1 [Source:HGNC Symbol;Acc:HGNC:24682] | 0.5008 | 5.91E-05 |
| ENSG00000180096 | SEPTIN1 | septin 1 [Source:HGNC Symbol;Acc:HGNC:2879] | -0.8604 | 5.93E-05 |
| ENSG00000282936 | NA | novel protein | 0.9270 | 5.93E-05 |
| ENSG00000167703 | SLC43A2 | solute carrier family 43 member 2 [Source:HGNC Symbol;Acc:HGNC:23087] | 1.2103 | 6.05E-05 |
| ENSG00000269378 | NA | integrin beta 1 pseudogene 1 | 0.8180 | 6.11E-05 |
| ENSG00000072864 | NDE1 | nudE neurodevelopment protein 1 [Source:HGNC Symbol;Acc:HGNC:17619] | -0.6148 | 6.14E-05 |
| ENSG00000287242 | NA | novel transcript, antisense to SCARB1 | 1.4508 | 6.17E-05 |
| ENSG00000227533 | SLC2A1-AS1 | SLC2A1 antisense RNA 1 [Source:HGNC Symbol;Acc:HGNC:44187] | -1.9173 | 6.17E-05 |
| ENSG00000166783 | MARF1 | meiosis regulator and mRNA stability factor 1 [Source:HGNC Symbol;Acc:HGNC:29562] | -0.5170 | 6.26E-05 |
| ENSG00000006459 | KDM7A | lysine demethylase 7A [Source:HGNC Symbol;Acc:HGNC:22224] | 1.5708 | 6.35E-05 |
| ENSG00000274828 | NA | novel transcript | -0.6434 | 6.41E-05 |
| ENSG00000113212 | PCDHB7 | protocadherin beta 7 [Source:HGNC Symbol;Acc:HGNC:8692] | -0.7032 | 6.41E-05 |
| ENSG00000271918 | NA | novel transcript, antisense to COMMD10 | -0.7930 | 6.42E-05 |
| ENSG00000151117 | TMEM86A | transmembrane protein 86A [Source:HGNC Symbol;Acc:HGNC:26890] | -1.2842 | 6.42E-05 |
| ENSG00000140386 | SCAPER | S-phase cyclin A associated protein in the ER [Source:HGNC Symbol;Acc:HGNC:13081] | -0.7032 | 6.42E-05 |
| ENSG00000261609 | GAN | gigaxonin [Source:HGNC Symbol;Acc:HGNC:4137] | 0.8656 | 6.50E-05 |
| ENSG00000156509 | FBXO43 | F-box protein 43 [Source:HGNC Symbol;Acc:HGNC:28521] | -1.0140 | 6.50E-05 |
| ENSG00000125459 | MSTO1 | misato mitochondrial distribution and morphology regulator 1 [Source:HGNC Symbol;Acc:HGNC:29678] | 0.6505 | 6.54E-05 |
| ENSG00000117533 | VAMP4 | vesicle associated membrane protein 4 [Source:HGNC Symbol;Acc:HGNC:12645] | 0.7119 | 6.56E-05 |
| ENSG00000119231 | SENP5 | SUMO specific peptidase 5 [Source:HGNC Symbol;Acc:HGNC:28407] | 0.5623 | 6.57E-05 |
| ENSG00000144026 | ZNF514 | zinc finger protein 514 [Source:HGNC Symbol;Acc:HGNC:25894] | -0.6602 | 6.63E-05 |
| ENSG00000255408 | PCDHA3 | protocadherin alpha 3 [Source:HGNC Symbol;Acc:HGNC:8669] | -0.9516 | 6.63E-05 |
| ENSG00000165886 | UBTD1 | ubiquitin domain containing 1 [Source:HGNC Symbol;Acc:HGNC:25683] | 0.9408 | 6.63E-05 |
| ENSG00000237560 | LINC01497 | long intergenic non-protein coding RNA 1497 [Source:HGNC Symbol;Acc:HGNC:51163] | 2.2986 | 6.66E-05 |
| ENSG00000075035 | WSCD2 | WSC domain containing 2 [Source:HGNC Symbol;Acc:HGNC:29117] | -2.2453 | 6.70E-05 |
| ENSG00000108799 | EZH1 | enhancer of zeste 1 polycomb repressive complex 2 subunit [Source:HGNC Symbol;Acc:HGNC:3526] | -0.5553 | 6.73E-05 |
| ENSG00000197226 | TBC1D9B | TBC1 domain family member 9B [Source:HGNC Symbol;Acc:HGNC:29097] | -0.5499 | 6.75E-05 |
| ENSG00000152348 | ATG10 | autophagy related 10 [Source:HGNC Symbol;Acc:HGNC:20315] | -0.5905 | 6.78E-05 |
| ENSG00000225151 | GOLGA2P7 | GOLGA2 pseudogene 7 [Source:HGNC Symbol;Acc:HGNC:49516] | -0.6736 | 6.85E-05 |
| ENSG00000166546 | BEAN1 | brain expressed associated with NEDD4 1 [Source:HGNC Symbol;Acc:HGNC:24160] | 1.1621 | 6.89E-05 |
| ENSG00000125703 | ATG4C | autophagy related 4C cysteine peptidase [Source:HGNC Symbol;Acc:HGNC:16040] | -0.6535 | 6.91E-05 |
| ENSG00000231806 | PCAT7 | prostate cancer associated transcript 7 [Source:HGNC Symbol;Acc:HGNC:48824] | -0.6222 | 6.94E-05 |
| ENSG00000115641 | FHL2 | four and a half LIM domains 2 [Source:HGNC Symbol;Acc:HGNC:3703] | -0.7810 | 7.02E-05 |
| ENSG00000137819 | PAQR5 | progestin and adipoQ receptor family member 5 [Source:HGNC Symbol;Acc:HGNC:29645] | -0.7074 | 7.07E-05 |
| ENSG00000266074 | BAHCC1 | BAH domain and coiled-coil containing 1 [Source:HGNC Symbol;Acc:HGNC:29279] | -2.3790 | 7.11E-05 |
| ENSG00000106012 | IQCE | IQ motif containing E [Source:HGNC Symbol;Acc:HGNC:29171] | 0.6573 | 7.15E-05 |

| | | | | |
|---|---|---|---|---|
| ENSG00000170522 | ELOVL6 | ELOVL fatty acid elongase 6 [Source:HGNC Symbol;Acc:HGNC:15829] | 0.8179 | 7.15E-05 |
| ENSG00000204991 | SPIRE2 | spire type actin nucleation factor 2 [Source:HGNC Symbol;Acc:HGNC:30623] | -0.6134 | 7.20E-05 |
| ENSG00000070961 | ATP2B1 | ATPase plasma membrane Ca2+ transporting 1 [Source:HGNC Symbol;Acc:HGNC:814] | 0.6297 | 7.27E-05 |
| ENSG00000196295 | GARS1-DT | GARS1 divergent transcript [Source:HGNC Symbol;Acc:HGNC:48951] | -0.6069 | 7.28E-05 |
| ENSG00000236603 | RANP1 | RAN pseudogene 1 [Source:HGNC Symbol;Acc:HGNC:21631] | 1.0578 | 7.33E-05 |
| ENSG00000079931 | MOXD1 | monooxygenase DBH like 1 [Source:HGNC Symbol;Acc:HGNC:21063] | -0.5356 | 7.34E-05 |
| ENSG00000228065 | LINC01515 | long intergenic non-protein coding RNA 1515 [Source:HGNC Symbol;Acc:HGNC:51210] | -0.5216 | 7.39E-05 |
| ENSG00000144369 | FAM171B | family with sequence similarity 171 member B [Source:HGNC Symbol;Acc:HGNC:29412] | -0.8182 | 7.45E-05 |
| ENSG00000095539 | SEMA4G | semaphorin 4G [Source:HGNC Symbol;Acc:HGNC:10735] | -0.8287 | 7.51E-05 |
| ENSG00000136943 | CTSV | cathepsin V [Source:HGNC Symbol;Acc:HGNC:2538] | 0.9698 | 7.53E-05 |
| ENSG00000150477 | KIAA1328 | KIAA1328 [Source:HGNC Symbol;Acc:HGNC:29248] | -0.7936 | 7.54E-05 |
| ENSG00000204116 | CHIC1 | cysteine rich hydrophobic domain 1 [Source:HGNC Symbol;Acc:HGNC:1934] | 0.5069 | 7.56E-05 |
| ENSG00000255020 | NA | novel transcript, antisense to TDH | -3.6588 | 7.62E-05 |
| ENSG00000181007 | ZFP82 | ZFP82 zinc finger protein [Source:HGNC Symbol;Acc:HGNC:28682] | -0.5719 | 7.67E-05 |
| ENSG00000108469 | RECQL5 | RecQ like helicase 5 [Source:HGNC Symbol;Acc:HGNC:9950] | -0.5977 | 7.68E-05 |
| ENSG00000125551 | PLGLB2 | plasminogen like B2 [Source:HGNC Symbol;Acc:HGNC:9073] | 0.9773 | 7.71E-05 |
| ENSG00000120729 | MYOT | myotilin [Source:HGNC Symbol;Acc:HGNC:12399] | 0.6197 | 7.75E-05 |
| ENSG00000085465 | OVGP1 | oviductal glycoprotein 1 [Source:HGNC Symbol;Acc:HGNC:8524] | 0.9791 | 7.78E-05 |
| ENSG00000068028 | RASSF1 | Ras association domain family member 1 [Source:HGNC Symbol;Acc:HGNC:9882] | 0.5901 | 7.80E-05 |
| ENSG00000103021 | CCDC113 | coiled-coil domain containing 113 [Source:HGNC Symbol;Acc:HGNC:25002] | -1.1353 | 7.83E-05 |
| ENSG00000162104 | ADCY9 | adenylate cyclase 9 [Source:HGNC Symbol;Acc:HGNC:240] | -0.6691 | 7.88E-05 |
| ENSG00000142949 | PTPRF | protein tyrosine phosphatase receptor type F [Source:HGNC Symbol;Acc:HGNC:9670] | -0.6318 | 7.91E-05 |
| ENSG00000153064 | BANK1 | B cell scaffold protein with ankyrin repeats 1 [Source:HGNC Symbol;Acc:HGNC:18233] | 1.8837 | 7.95E-05 |
| ENSG00000070669 | ASNS | asparagine synthetase (glutamine-hydrolyzing) [Source:HGNC Symbol;Acc:HGNC:753] | 0.6992 | 8.14E-05 |
| ENSG00000172687 | ZNF738 | zinc finger protein 738 [Source:HGNC Symbol;Acc:HGNC:32469] | -0.7850 | 8.16E-05 |
| ENSG00000087299 | L2HGDH | L-2-hydroxyglutarate dehydrogenase [Source:HGNC Symbol;Acc:HGNC:20499] | -0.5208 | 8.21E-05 |
| ENSG00000237649 | KIFC1 | kinesin family member C1 [Source:HGNC Symbol;Acc:HGNC:6389] | -0.6110 | 8.31E-05 |
| ENSG00000009413 | REV3L | REV3 like, DNA directed polymerase zeta catalytic subunit [Source:HGNC Symbol;Acc:HGNC:9968] | -0.6553 | 8.32E-05 |
| ENSG00000147894 | C9orf72 | C9orf72-SMCR8 complex subunit [Source:HGNC Symbol;Acc:HGNC:28337] | 0.5149 | 8.34E-05 |
| ENSG00000154721 | JAM2 | junctional adhesion molecule 2 [Source:HGNC Symbol;Acc:HGNC:14686] | -0.6279 | 8.41E-05 |
| ENSG00000089225 | TBX5 | T-box transcription factor 5 [Source:HGNC Symbol;Acc:HGNC:11604] | -2.2376 | 8.53E-05 |
| ENSG00000167508 | MVD | mevalonate diphosphate decarboxylase [Source:HGNC Symbol;Acc:HGNC:7529] | -0.5041 | 8.57E-05 |
| ENSG00000167280 | ENGASE | endo-beta-N-acetylglucosaminidase [Source:HGNC Symbol;Acc:HGNC:24622] | -0.9206 | 8.58E-05 |
| ENSG00000105643 | ARRDC2 | arrestin domain containing 2 [Source:HGNC Symbol;Acc:HGNC:25225] | 0.9619 | 8.59E-05 |
| ENSG00000267801 | NA | novel transcript, antisense to TRIM47 | 1.0858 | 8.66E-05 |
| ENSG00000275830 | NA | novel transcript | 1.8840 | 8.88E-05 |
| ENSG00000143344 | RGL1 | ral guanine nucleotide dissociation stimulator like 1 [Source:HGNC Symbol;Acc:HGNC:30281] | 0.6534 | 8.95E-05 |
| ENSG00000168661 | ZNF30 | zinc finger protein 30 [Source:HGNC Symbol;Acc:HGNC:13090] | -0.7568 | 8.96E-05 |
| ENSG00000176890 | TYMS | thymidylate synthetase [Source:HGNC Symbol;Acc:HGNC:12441] | -0.6464 | 9.02E-05 |
| ENSG00000136235 | GPNMB | glycoprotein nmb [Source:HGNC Symbol;Acc:HGNC:4462] | -0.5244 | 9.04E-05 |
| ENSG00000198003 | CCDC151 | coiled-coil domain containing 151 [Source:HGNC Symbol;Acc:HGNC:28303] | -0.9293 | 9.11E-05 |
| ENSG00000116668 | SWT1 | SWT1 RNA endoribonuclease homolog [Source:HGNC Symbol;Acc:HGNC:16785] | -0.9000 | 9.33E-05 |
| ENSG00000163898 | LIPH | lipase H [Source:HGNC Symbol;Acc:HGNC:18483] | -0.5444 | 9.41E-05 |

| | | | | |
|---|---|---|---|---|
| ENSG00000254211 | LINC01485 | long intergenic non-protein coding RNA 1485 [Source:HGNC Symbol;Acc:HGNC:51138] | 3.2178 | 9.43E-05 |
| ENSG00000198556 | ZNF789 | zinc finger protein 789 [Source:HGNC Symbol;Acc:HGNC:27801] | -0.7164 | 9.48E-05 |
| ENSG00000146151 | HMGCLL1 | 3-hydroxymethyl-3-methylglutaryl-CoA lyase like 1 [Source:HGNC Symbol;Acc:HGNC:21359] | -0.8715 | 9.71E-05 |
| ENSG00000070718 | AP3M2 | adaptor related protein complex 3 subunit mu 2 [Source:HGNC Symbol;Acc:HGNC:570] | -0.5802 | 9.72E-05 |
| ENSG00000166986 | mars-01 | methionyl-tRNA synthetase 1 [Source:HGNC Symbol;Acc:HGNC:6898] | 0.5169 | 9.75E-05 |
| ENSG00000183783 | KCTD8 | potassium channel tetramerization domain containing 8 [Source:HGNC Symbol;Acc:HGNC:22394] | -1.7408 | 9.75E-05 |
| ENSG00000259775 | NA | novel transcript, antisense to EIF5 | 1.2634 | 9.80E-05 |
| ENSG00000196787 | H2AC11 | H2A clustered histone 11 [Source:HGNC Symbol;Acc:HGNC:4737] | -0.5356 | 9.83E-05 |
| ENSG00000185168 | LINC00482 | long intergenic non-protein coding RNA 482 [Source:HGNC Symbol;Acc:HGNC:26816] | -1.9462 | 9.86E-05 |
| ENSG00000251692 | PTX4 | pentraxin 4 [Source:HGNC Symbol;Acc:HGNC:14171] | -1.5130 | 9.91E-05 |
| ENSG00000003249 | DBNDD1 | dysbindin domain containing 1 [Source:HGNC Symbol;Acc:HGNC:28455] | 0.5495 | 9.92E-05 |
| ENSG00000255240 | NA | novel transcript | -1.3617 | 0.00010 |
| ENSG00000180875 | GREM2 | gremlin 2, DAN family BMP antagonist [Source:HGNC Symbol;Acc:HGNC:17655] | 1.3348 | 0.00010 |
| ENSG00000273888 | FRMD6-AS1 | FRMD6 antisense RNA 1 [Source:HGNC Symbol;Acc:HGNC:20129] | -1.2370 | 0.00010 |
| ENSG00000124067 | SLC12A4 | solute carrier family 12 member 4 [Source:HGNC Symbol;Acc:HGNC:10913] | 0.5245 | 0.00010 |
| ENSG00000188385 | JAKMIP3 | Janus kinase and microtubule interacting protein 3 [Source:HGNC Symbol;Acc:HGNC:23523] | -1.1166 | 0.00010 |
| ENSG00000256870 | SLC5A8 | solute carrier family 5 member 8 [Source:HGNC Symbol;Acc:HGNC:19119] | -2.1836 | 0.00010 |
| ENSG00000111641 | NOP2 | NOP2 nucleolar protein [Source:HGNC Symbol;Acc:HGNC:7867] | 0.5100 | 0.00010 |
| ENSG00000099889 | ARVCF | ARVCF delta catenin family member [Source:HGNC Symbol;Acc:HGNC:728] | -0.5481 | 0.00011 |
| ENSG00000204389 | HSPA1A | heat shock protein family A (Hsp70) member 1A [Source:HGNC Symbol;Acc:HGNC:5232] | -0.8618 | 0.00011 |
| ENSG00000180938 | ZNF572 | zinc finger protein 572 [Source:HGNC Symbol;Acc:HGNC:26758] | -0.6657 | 0.00011 |
| ENSG00000183765 | CHEK2 | checkpoint kinase 2 [Source:HGNC Symbol;Acc:HGNC:16627] | -0.8557 | 0.00011 |
| ENSG00000114735 | HEMK1 | HemK methyltransferase family member 1 [Source:HGNC Symbol;Acc:HGNC:24923] | -0.5749 | 0.00011 |
| ENSG00000100065 | CARD10 | caspase recruitment domain family member 10 [Source:HGNC Symbol;Acc:HGNC:16422] | 0.7467 | 0.00011 |
| ENSG00000157214 | STEAP2 | STEAP2 metalloreductase [Source:HGNC Symbol;Acc:HGNC:17885] | -1.6182 | 0.00011 |
| ENSG00000164776 | PHKG1 | phosphorylase kinase catalytic subunit gamma 1 [Source:HGNC Symbol;Acc:HGNC:8930] | 1.1488 | 0.00011 |
| ENSG00000249859 | PVT1 | Pvt1 oncogene [Source:HGNC Symbol;Acc:HGNC:9709] | -1.1838 | 0.00011 |
| ENSG00000167220 | HDHD2 | haloacid dehalogenase like hydrolase domain containing 2 [Source:HGNC Symbol;Acc:HGNC:25364] | -0.6421 | 0.00011 |
| ENSG00000278921 | EPB41L4A-DT | EPB41L4A divergent transcript [Source:HGNC Symbol;Acc:HGNC:25643] | 0.7255 | 0.00011 |
| ENSG00000150995 | ITPR1 | inositol 1,4,5-trisphosphate receptor type 1 [Source:HGNC Symbol;Acc:HGNC:6180] | -0.5785 | 0.00011 |
| ENSG00000166971 | AKTIP | AKT interacting protein [Source:HGNC Symbol;Acc:HGNC:16710] | -0.6071 | 0.00011 |
| ENSG00000214530 | STARD10 | StAR related lipid transfer domain containing 10 [Source:HGNC Symbol;Acc:HGNC:10666] | -0.5520 | 0.00011 |
| ENSG00000154102 | C16orf74 | chromosome 16 open reading frame 74 [Source:HGNC Symbol;Acc:HGNC:23362] | -2.0529 | 0.00011 |
| ENSG00000118785 | SPP1 | secreted phosphoprotein 1 [Source:HGNC Symbol;Acc:HGNC:11255] | 0.9033 | 0.00011 |
| ENSG00000105877 | DNAH11 | dynein axonemal heavy chain 11 [Source:HGNC Symbol;Acc:HGNC:2942] | 0.8963 | 0.00012 |
| ENSG00000274895 | NA | novel transcript, antisense to PROX1 | 1.7425 | 0.00012 |
| ENSG00000140450 | ARRDC4 | arrestin domain containing 4 [Source:HGNC Symbol;Acc:HGNC:28087] | 0.8168 | 0.00012 |
| ENSG00000160691 | SHC1 | SHC adaptor protein 1 [Source:HGNC Symbol;Acc:HGNC:10840] | 0.5262 | 0.00012 |
| ENSG00000182511 | FES | FES proto-oncogene, tyrosine kinase [Source:HGNC Symbol;Acc:HGNC:3657] | 1.7337 | 0.00012 |
| ENSG00000263165 | NA | novel transcript, antisense to ANKFY1 | -6.5143 | 0.00012 |
| ENSG00000256340 | ABCC6P1 | ATP binding cassette subfamily C member 6 pseudogene 1 [Source:HGNC Symbol;Acc:HGNC:33352] | -1.7569 | 0.00012 |
| ENSG00000103227 | LMF1 | lipase maturation factor 1 [Source:HGNC Symbol;Acc:HGNC:14154] | -0.5476 | 0.00012 |
| ENSG00000196646 | ZNF136 | zinc finger protein 136 [Source:HGNC Symbol;Acc:HGNC:12920] | -0.6213 | 0.00012 |

| | | | | |
|---|---|---|---|---|
| ENSG00000205918 | PDPK2P | 3-phosphoinositide dependent protein kinase 2, pseudogene [Source:HGNC Symbol;Acc:HGNC:49897] | 0.9208 | 0.00012 |
| ENSG00000272556 | GTF2IP13 | general transcription factor IIi pseudogene 13 [Source:HGNC Symbol;Acc:HGNC:51725] | -0.5024 | 0.00012 |
| ENSG00000105771 | SMG9 | SMG9 nonsense mediated mRNA decay factor [Source:HGNC Symbol;Acc:HGNC:25763] | 0.5604 | 0.00012 |
| ENSG00000105612 | DNASE2 | deoxyribonuclease 2, lysosomal [Source:HGNC Symbol;Acc:HGNC:2960] | -0.5372 | 0.00012 |
| ENSG00000112182 | BACH2 | BTB domain and CNC homolog 2 [Source:HGNC Symbol;Acc:HGNC:14078] | 2.2503 | 0.00013 |
| ENSG00000088881 | EBF4 | EBF family member 4 [Source:HGNC Symbol;Acc:HGNC:29278] | 0.5666 | 0.00013 |
| ENSG00000204366 | ZBTB12 | zinc finger and BTB domain containing 12 [Source:HGNC Symbol;Acc:HGNC:19066] | -0.8235 | 0.00013 |
| ENSG00000025293 | PHF20 | PHD finger protein 20 [Source:HGNC Symbol;Acc:HGNC:16098] | -0.7157 | 0.00013 |
| ENSG00000179454 | KLHL28 | kelch like family member 28 [Source:HGNC Symbol;Acc:HGNC:19741] | 0.5014 | 0.00013 |
| ENSG00000159335 | PTMS | parathymosin [Source:HGNC Symbol;Acc:HGNC:9629] | 0.5330 | 0.00013 |
| ENSG00000187193 | MT1X | metallothionein 1X [Source:HGNC Symbol;Acc:HGNC:7405] | 0.8806 | 0.00013 |
| ENSG00000255583 | NA | T-box 20 (TBX20) pseudogene | 3.2648 | 0.00013 |
| ENSG00000160193 | WDR4 | WD repeat domain 4 [Source:HGNC Symbol;Acc:HGNC:12756] | 0.5325 | 0.00013 |
| ENSG00000197180 | NA | uncharacterized protein BC009467 [Source:NCBI gene (formerly Entrezgene);Acc:158960] | -0.7364 | 0.00013 |
| ENSG00000102984 | ZNF821 | zinc finger protein 821 [Source:HGNC Symbol;Acc:HGNC:28043] | 0.7439 | 0.00013 |
| ENSG00000165443 | PHYHIPL | phytanoyl-CoA 2-hydroxylase interacting protein like [Source:HGNC Symbol;Acc:HGNC:29378] | 0.6148 | 0.00013 |
| ENSG00000185361 | TNFAIP8L1 | TNF alpha induced protein 8 like 1 [Source:HGNC Symbol;Acc:HGNC:28279] | -0.6776 | 0.00014 |
| ENSG00000086717 | PPEF1 | protein phosphatase with EF-hand domain 1 [Source:HGNC Symbol;Acc:HGNC:9243] | 1.2678 | 0.00014 |
| ENSG00000181031 | RPH3AL | rabphilin 3A like (without C2 domains) [Source:HGNC Symbol;Acc:HGNC:10296] | 0.9353 | 0.00014 |
| ENSG00000206579 | XKR4 | XK related 4 [Source:HGNC Symbol;Acc:HGNC:29394] | 2.3260 | 0.00014 |
| ENSG00000186642 | PDE2A | phosphodiesterase 2A [Source:HGNC Symbol;Acc:HGNC:8777] | -0.6168 | 0.00014 |
| ENSG00000285830 | NA | novel transcript, antisense to TRAPPC6B | -1.5834 | 0.00014 |
| ENSG00000260948 | NA | novel transcript, sense overlapping WDR77 & OVGP1 | 1.1329 | 0.00014 |
| ENSG00000184979 | USP18 | ubiquitin specific peptidase 18 [Source:HGNC Symbol;Acc:HGNC:12616] | -1.1697 | 0.00014 |
| ENSG00000099251 | HSD17B7P2 | hydroxysteroid 17-beta dehydrogenase 7 pseudogene 2 [Source:HGNC Symbol;Acc:HGNC:28120] | -0.6875 | 0.00014 |
| ENSG00000172197 | MBOAT1 | membrane bound O-acyltransferase domain containing 1 [Source:HGNC Symbol;Acc:HGNC:21579] | -1.0160 | 0.00014 |
| ENSG00000174327 | SLC16A13 | solute carrier family 16 member 13 [Source:HGNC Symbol;Acc:HGNC:31037] | -0.7814 | 0.00014 |
| ENSG00000157764 | BRAF | B-Raf proto-oncogene, serine/threonine kinase [Source:HGNC Symbol;Acc:HGNC:1097] | 0.5546 | 0.00014 |
| ENSG00000180739 | S1PR5 | sphingosine-1-phosphate receptor 5 [Source:HGNC Symbol;Acc:HGNC:14299] | -0.9331 | 0.00014 |
| ENSG00000261135 | NA | novel transcript | -1.2658 | 0.00015 |
| ENSG00000171450 | CDK5R2 | cyclin dependent kinase 5 regulatory subunit 2 [Source:HGNC Symbol;Acc:HGNC:1776] | -1.0806 | 0.00015 |
| ENSG00000151338 | MIPOL1 | mirror-image polydactyly 1 [Source:HGNC Symbol;Acc:HGNC:21460] | -1.6461 | 0.00015 |
| ENSG00000115902 | SLC1A4 | solute carrier family 1 member 4 [Source:HGNC Symbol;Acc:HGNC:10942] | 0.5787 | 0.00015 |
| ENSG00000156639 | ZFAND3 | zinc finger AN1-type containing 3 [Source:HGNC Symbol;Acc:HGNC:18019] | 0.5037 | 0.00015 |
| ENSG00000174586 | ZNF497 | zinc finger protein 497 [Source:HGNC Symbol;Acc:HGNC:23714] | -0.7378 | 0.00015 |
| ENSG00000148516 | ZEB1 | zinc finger E-box binding homeobox 1 [Source:HGNC Symbol;Acc:HGNC:11642] | 0.9035 | 0.00015 |
| ENSG00000253696 | KBTBD11-OT1 | KBTBD11 overlapping transcript 1 [Source:HGNC Symbol;Acc:HGNC:49147] | 2.3906 | 0.00015 |
| ENSG00000143669 | LYST | lysosomal trafficking regulator [Source:HGNC Symbol;Acc:HGNC:1968] | 1.5678 | 0.00015 |
| ENSG00000221817 | PPP3CB-AS1 | PPP3CB antisense RNA 1 (head to head) [Source:HGNC Symbol;Acc:HGNC:50750] | -0.9839 | 0.00015 |
| ENSG00000255085 | NA | zinc finger protein 154 (ZNF154) pseudogene | -1.4477 | 0.00015 |
| ENSG00000249437 | NAIP | NLR family apoptosis inhibitory protein [Source:HGNC Symbol;Acc:HGNC:7634] | -0.5926 | 0.00015 |
| ENSG00000135127 | BICDL1 | BICD family like cargo adaptor 1 [Source:HGNC Symbol;Acc:HGNC:28095] | 0.9365 | 0.00015 |
| ENSG00000169548 | ZNF280A | zinc finger protein 280A [Source:HGNC Symbol;Acc:HGNC:18597] | 2.3035 | 0.00015 |

| | | | | |
|---|---|---|---|---|
| ENSG00000103642 | LACTB | lactamase beta [Source:HGNC Symbol;Acc:HGNC:16468] | -0.7317 | 0.00015 |
| ENSG00000175699 | CCDC197 | coiled-coil domain containing 197 [Source:HGNC Symbol;Acc:HGNC:19860] | 2.8995 | 0.00015 |
| ENSG00000166831 | RBPMS2 | RNA binding protein, mRNA processing factor 2 [Source:HGNC Symbol;Acc:HGNC:19098] | 0.5253 | 0.00016 |
| ENSG00000100478 | AP4S1 | adaptor related protein complex 4 subunit sigma 1 [Source:HGNC Symbol;Acc:HGNC:575] | 0.5865 | 0.00016 |
| ENSG00000042088 | TDP1 | tyrosyl-DNA phosphodiesterase 1 [Source:HGNC Symbol;Acc:HGNC:18884] | 0.5229 | 0.00016 |
| ENSG00000144451 | SPAG16 | sperm associated antigen 16 [Source:HGNC Symbol;Acc:HGNC:23225] | -0.7059 | 0.00016 |
| ENSG00000099260 | PALMD | palmdelphin [Source:HGNC Symbol;Acc:HGNC:15846] | -0.6413 | 0.00016 |
| ENSG00000260552 | NA | novel transcript, antisense to MOCOS | -1.2504 | 0.00016 |
| ENSG00000170837 | GPR27 | G protein-coupled receptor 27 [Source:HGNC Symbol;Acc:HGNC:4482] | -1.2939 | 0.00016 |
| ENSG00000276141 | WHAMMP3 | WAS protein homolog associated with actin, golgi membranes and microtubules pseudogene 3 [Source:HGNC Symbol;Acc:HGNC:27892] | -0.8501 | 0.00016 |
| ENSG00000260077 | NA | novel transcript | -1.4047 | 0.00016 |
| ENSG00000214274 | ANG | angiogenin [Source:HGNC Symbol;Acc:HGNC:483] | -1.1252 | 0.00016 |
| ENSG00000288611 | NPBWR1 | neuropeptides B and W receptor 1 [Source:HGNC Symbol;Acc:HGNC:4522] | 1.4551 | 0.00016 |
| ENSG00000225791 | TRAM2-AS1 | TRAM2 antisense RNA 1 (head to head) [Source:HGNC Symbol;Acc:HGNC:48663] | -0.5531 | 0.00016 |
| ENSG00000063761 | ADCK1 | aarF domain containing kinase 1 [Source:HGNC Symbol;Acc:HGNC:19038] | -0.8504 | 0.00016 |
| ENSG00000243646 | IL10RB | interleukin 10 receptor subunit beta [Source:HGNC Symbol;Acc:HGNC:5965] | -0.6738 | 0.00016 |
| ENSG00000141040 | ZNF287 | zinc finger protein 287 [Source:HGNC Symbol;Acc:HGNC:13502] | -0.8477 | 0.00016 |
| ENSG00000111012 | CYP27B1 | cytochrome P450 family 27 subfamily B member 1 [Source:HGNC Symbol;Acc:HGNC:2606] | 0.9999 | 0.00016 |
| ENSG00000132326 | PER2 | period circadian regulator 2 [Source:HGNC Symbol;Acc:HGNC:8846] | 1.3478 | 0.00016 |
| ENSG00000182255 | KCNA4 | potassium voltage-gated channel subfamily A member 4 [Source:HGNC Symbol;Acc:HGNC:6222] | 0.9009 | 0.00016 |
| ENSG00000181722 | ZBTB20 | zinc finger and BTB domain containing 20 [Source:HGNC Symbol;Acc:HGNC:13503] | 0.6584 | 0.00016 |
| ENSG00000227755 | NA | MAD1 mitotic arrest deficient-like 1 (yeast)(MAD1L1) pseudogene | -3.1385 | 0.00016 |
| ENSG00000204839 | MROH6 | maestro heat like repeat family member 6 [Source:HGNC Symbol;Acc:HGNC:27814] | 0.9667 | 0.00016 |
| ENSG00000153827 | TRIP12 | thyroid hormone receptor interactor 12 [Source:HGNC Symbol;Acc:HGNC:12306] | 0.5463 | 0.00016 |
| ENSG00000100418 | DESI1 | desumoylating isopeptidase 1 [Source:HGNC Symbol;Acc:HGNC:24577] | 0.6419 | 0.00016 |
| ENSG00000171116 | HSFX1 | heat shock transcription factor family, X-linked 1 [Source:HGNC Symbol;Acc:HGNC:29603] | -1.2641 | 0.00016 |
| ENSG00000119866 | BCL11A | BAF chromatin remodeling complex subunit BCL11A [Source:HGNC Symbol;Acc:HGNC:13221] | -0.6090 | 0.00017 |
| ENSG00000150054 | MPP7 | membrane palmitoylated protein 7 [Source:HGNC Symbol;Acc:HGNC:26542] | -1.1828 | 0.00017 |
| ENSG00000154556 | SORBS2 | sorbin and SH3 domain containing 2 [Source:HGNC Symbol;Acc:HGNC:24098] | -0.5519 | 0.00017 |
| ENSG00000101928 | MOSPD1 | motile sperm domain containing 1 [Source:HGNC Symbol;Acc:HGNC:25235] | 0.5277 | 0.00017 |
| ENSG00000182718 | ANXA2 | annexin A2 [Source:HGNC Symbol;Acc:HGNC:537] | 2.6906 | 0.00017 |
| ENSG00000170011 | MYRIP | myosin VIIA and Rab interacting protein [Source:HGNC Symbol;Acc:HGNC:19156] | -0.8232 | 0.00017 |
| ENSG00000118242 | MREG | melanoregulin [Source:HGNC Symbol;Acc:HGNC:25478] | -0.9577 | 0.00017 |
| ENSG00000155792 | DEPTOR | DEP domain containing MTOR interacting protein [Source:HGNC Symbol;Acc:HGNC:22953] | -0.7101 | 0.00017 |
| ENSG00000165821 | SALL2 | spalt like transcription factor 2 [Source:HGNC Symbol;Acc:HGNC:10526] | -0.5419 | 0.00018 |
| ENSG00000273702 | NA | novel transcript | -0.8599 | 0.00018 |
| ENSG00000261773 | NA | novel transcript, overlapping to FAM3A | 2.2248 | 0.00018 |
| ENSG00000102547 | CAB39L | calcium binding protein 39 like [Source:HGNC Symbol;Acc:HGNC:20290] | -1.0456 | 0.00018 |
| ENSG00000146205 | ANO7 | anoctamin 7 [Source:HGNC Symbol;Acc:HGNC:31677] | 1.1617 | 0.00018 |
| ENSG00000233058 | LINC00884 | long intergenic non-protein coding RNA 884 [Source:HGNC Symbol;Acc:HGNC:48570] | 0.9505 | 0.00018 |
| ENSG00000145335 | SNCA | synuclein alpha [Source:HGNC Symbol;Acc:HGNC:11138] | 2.0018 | 0.00018 |
| ENSG00000124613 | ZNF391 | zinc finger protein 391 [Source:HGNC Symbol;Acc:HGNC:18779] | -1.0009 | 0.00018 |

| | | | | |
|---|---|---|---|---|
| ENSG00000163689 | CFAP20DC | CFAP20 domain containing [Source:HGNC Symbol;Acc:HGNC:24763] | -0.6487 | 0.00018 |
| ENSG00000230082 | PRRT3-AS1 | PRRT3 antisense RNA 1 [Source:HGNC Symbol;Acc:HGNC:41151] | 2.6903 | 0.00018 |
| ENSG00000159433 | STARD9 | StAR related lipid transfer domain containing 9 [Source:HGNC Symbol;Acc:HGNC:19162] | -0.7359 | 0.00018 |
| ENSG00000102393 | GLA | galactosidase alpha [Source:HGNC Symbol;Acc:HGNC:4296] | 0.5396 | 0.00018 |
| ENSG00000069998 | HDHD5 | haloacid dehalogenase like hydrolase domain containing 5 [Source:HGNC Symbol;Acc:HGNC:1843] | -0.5769 | 0.00018 |
| ENSG00000146350 | TBC1D32 | TBC1 domain family member 32 [Source:HGNC Symbol;Acc:HGNC:21485] | -0.7446 | 0.00018 |
| ENSG00000184792 | OSBP2 | oxysterol binding protein 2 [Source:HGNC Symbol;Acc:HGNC:8504] | 0.6893 | 0.00018 |
| ENSG00000155011 | 2,00 DKK | dickkopf WNT signaling pathway inhibitor 2 [Source:HGNC Symbol;Acc:HGNC:2892] | 1.5831 | 0.00018 |
| ENSG00000149499 | EML3 | EMAP like 3 [Source:HGNC Symbol;Acc:HGNC:26666] | 0.6578 | 0.00018 |
| ENSG00000102871 | TRADD | TNFRSF1A associated via death domain [Source:HGNC Symbol;Acc:HGNC:12030] | -0.7527 | 0.00018 |
| ENSG00000225742 | LINC02036 | long intergenic non-protein coding RNA 2036 [Source:HGNC Symbol;Acc:HGNC:52876] | -1.8773 | 0.00018 |
| ENSG00000162543 | UBXN10 | UBX domain protein 10 [Source:HGNC Symbol;Acc:HGNC:26354] | 5.3672 | 0.00018 |
| ENSG00000111602 | TIMELESS | timeless circadian regulator [Source:HGNC Symbol;Acc:HGNC:11813] | -0.6771 | 0.00019 |
| ENSG00000110871 | COQ5 | coenzyme Q5, methyltransferase [Source:HGNC Symbol;Acc:HGNC:28722] | -0.5943 | 0.00019 |
| ENSG00000212123 | PRR22 | proline rich 22 [Source:HGNC Symbol;Acc:HGNC:28354] | 0.6125 | 0.00019 |
| ENSG00000258743 | LINC02301 | long intergenic non-protein coding RNA 2301 [Source:HGNC Symbol;Acc:HGNC:53220] | -1.8248 | 0.00019 |
| ENSG00000112149 | CD83 | CD83 molecule [Source:HGNC Symbol;Acc:HGNC:1703] | 0.5039 | 0.00019 |
| ENSG00000154447 | SH3RF1 | SH3 domain containing ring finger 1 [Source:HGNC Symbol;Acc:HGNC:17650] | 0.5825 | 0.00019 |
| ENSG00000197566 | ZNF624 | zinc finger protein 624 [Source:HGNC Symbol;Acc:HGNC:29254] | -0.7942 | 0.00019 |
| ENSG00000013810 | TACC3 | transforming acidic coiled-coil containing protein 3 [Source:HGNC Symbol;Acc:HGNC:11524] | -0.5578 | 0.00019 |
| ENSG00000080644 | CHRNA3 | cholinergic receptor nicotinic alpha 3 subunit [Source:HGNC Symbol;Acc:HGNC:1957] | -1.1323 | 0.00019 |
| ENSG00000196368 | NUDT11 | nudix hydrolase 11 [Source:HGNC Symbol;Acc:HGNC:18011] | -0.5264 | 0.00019 |
| ENSG00000187688 | TRPV2 | transient receptor potential cation channel subfamily V member 2 [Source:HGNC Symbol;Acc:HGNC:18082] | 0.7105 | 0.00019 |
| ENSG00000174928 | C3orf33 | chromosome 3 open reading frame 33 [Source:HGNC Symbol;Acc:HGNC:26434] | -0.6530 | 0.00019 |
| ENSG00000153832 | FBXO36 | F-box protein 36 [Source:HGNC Symbol;Acc:HGNC:27020] | -1.1512 | 0.00019 |
| ENSG00000163273 | NPPC | natriuretic peptide C [Source:HGNC Symbol;Acc:HGNC:7941] | 2.3031 | 0.00019 |
| ENSG00000283663 | NA | novel transcript | -1.6288 | 0.00020 |
| ENSG00000197763 | TXNRD3 | thioredoxin reductase 3 [Source:HGNC Symbol;Acc:HGNC:20667] | -1.0554 | 0.00020 |
| ENSG00000128218 | VPREB3 | V-set pre-B cell surrogate light chain 3 [Source:HGNC Symbol;Acc:HGNC:12710] | -0.5948 | 0.00020 |
| ENSG00000005189 | REXO5 | RNA exonuclease 5 [Source:HGNC Symbol;Acc:HGNC:24661] | -0.8503 | 0.00020 |
| ENSG00000138642 | HERC6 | HECT and RLD domain containing E3 ubiquitin protein ligase family member 6 [Source:HGNC Symbol;Acc:HGNC:26072] | -1.3320 | 0.00020 |
| ENSG00000230513 | THAP7-AS1 | THAP7 antisense RNA 1 [Source:HGNC Symbol;Acc:HGNC:41013] | -0.5486 | 0.00020 |
| ENSG00000164070 | HSPA4L | heat shock protein family A (Hsp70) member 4 like [Source:HGNC Symbol;Acc:HGNC:17041] | 0.5859 | 0.00020 |
| ENSG00000279443 | NA | TEC | 0.9702 | 0.00020 |
| ENSG00000204228 | HSD17B8 | hydroxysteroid 17-beta dehydrogenase 8 [Source:HGNC Symbol;Acc:HGNC:3554] | -0.7908 | 0.00020 |
| ENSG00000223553 | SMPD4P1 | sphingomyelin phosphodiesterase 4 pseudogene 1 [Source:HGNC Symbol;Acc:HGNC:39673] | -2.1571 | 0.00020 |
| ENSG00000108825 | PTGES3L-AARSD1 | PTGES3L-AARSD1 readthrough [Source:HGNC Symbol;Acc:HGNC:43946] | 1.7146 | 0.00020 |
| ENSG00000168065 | SLC22A11 | solute carrier family 22 member 11 [Source:HGNC Symbol;Acc:HGNC:18120] | -0.7855 | 0.00021 |
| ENSG00000135631 | RAB11FIP5 | RAB11 family interacting protein 5 [Source:HGNC Symbol;Acc:HGNC:24845] | -0.5846 | 0.00021 |
| ENSG00000102362 | SYTL4 | synaptotagmin like 4 [Source:HGNC Symbol;Acc:HGNC:15588] | -0.8888 | 0.00021 |
| ENSG00000159733 | ZFYVE28 | zinc finger FYVE-type containing 28 [Source:HGNC Symbol;Acc:HGNC:29334] | -0.6908 | 0.00021 |

| | | | | |
|---|---|---|---|---|
| ENSG00000163558 | PRKCI | protein kinase C iota [Source:HGNC Symbol;Acc:HGNC:9404] | 0.6121 | 0.00021 |
| ENSG00000103222 | ABCC1 | ATP binding cassette subfamily C member 1 [Source:HGNC Symbol;Acc:HGNC:51] | 0.5101 | 0.00021 |
| ENSG00000112769 | LAMA4 | laminin subunit alpha 4 [Source:HGNC Symbol;Acc:HGNC:6484] | -0.7990 | 0.00021 |
| ENSG00000244165 | P2RY11 | purinergic receptor P2Y11 [Source:HGNC Symbol;Acc:HGNC:8540] | 0.5436 | 0.00021 |
| ENSG00000231233 | CFAP58-DT | CFAP58 divergent transcript [Source:HGNC Symbol;Acc:HGNC:45243] | 1.0107 | 0.00021 |
| ENSG00000153914 | SREK1 | splicing regulatory glutamic acid and lysine rich protein 1 [Source:HGNC Symbol;Acc:HGNC:17882] | 0.6567 | 0.00021 |
| ENSG00000205464 | ATP6AP1L | ATPase H+ transporting accessory protein 1 like [Source:HGNC Symbol;Acc:HGNC:28091] | -0.6622 | 0.00021 |
| ENSG00000142512 | SIGLEC10 | sialic acid binding Ig like lectin 10 [Source:HGNC Symbol;Acc:HGNC:15620] | -2.0267 | 0.00021 |
| ENSG00000197619 | ZNF615 | zinc finger protein 615 [Source:HGNC Symbol;Acc:HGNC:24740] | -0.5114 | 0.00021 |
| ENSG00000236901 | MIR600HG | MIR600 host gene [Source:HGNC Symbol;Acc:HGNC:23642] | -0.7136 | 0.00021 |
| ENSG00000182890 | GLUD2 | glutamate dehydrogenase 2 [Source:HGNC Symbol;Acc:HGNC:4336] | 0.5564 | 0.00022 |
| ENSG00000246273 | SBF2-AS1 | SBF2 antisense RNA 1 [Source:HGNC Symbol;Acc:HGNC:27438] | -0.8370 | 0.00022 |
| ENSG00000267892 | NA | novel transcript, antisense to CAPN12 | 1.7668 | 0.00022 |
| ENSG00000091262 | ABCC6 | ATP binding cassette subfamily C member 6 [Source:HGNC Symbol;Acc:HGNC:57] | -1.3608 | 0.00022 |
| ENSG00000099999 | RNF215 | ring finger protein 215 [Source:HGNC Symbol;Acc:HGNC:33434] | -0.5842 | 0.00022 |
| ENSG00000066629 | EML1 | EMAP like 1 [Source:HGNC Symbol;Acc:HGNC:3330] | -0.7926 | 0.00022 |
| ENSG00000273162 | NA | novel transcript | -1.0723 | 0.00022 |
| ENSG00000183092 | BEGAIN | brain enriched guanylate kinase associated [Source:HGNC Symbol;Acc:HGNC:24163] | 1.3409 | 0.00023 |
| ENSG00000116852 | KIF21B | kinesin family member 21B [Source:HGNC Symbol;Acc:HGNC:29442] | -2.7794 | 0.00023 |
| ENSG00000171310 | CHST11 | carbohydrate sulfotransferase 11 [Source:HGNC Symbol;Acc:HGNC:17422] | 1.3054 | 0.00023 |
| ENSG00000279970 | NA | TEC | 1.1726 | 0.00023 |
| ENSG00000140939 | NOL3 | nucleolar protein 3 [Source:HGNC Symbol;Acc:HGNC:7869] | -0.6236 | 0.00023 |
| ENSG00000173376 | NDNF | neuron derived neurotrophic factor [Source:HGNC Symbol;Acc:HGNC:26256] | -0.6306 | 0.00023 |
| ENSG00000141068 | KSR1 | kinase suppressor of ras 1 [Source:HGNC Symbol;Acc:HGNC:6465] | -0.5058 | 0.00023 |
| ENSG00000155729 | KCTD18 | potassium channel tetramerization domain containing 18 [Source:HGNC Symbol;Acc:HGNC:26446] | -0.7202 | 0.00023 |
| ENSG00000188266 | HYKK | hydroxylysine kinase [Source:HGNC Symbol;Acc:HGNC:34403] | -1.3169 | 0.00023 |
| ENSG00000161682 | FAM171A2 | family with sequence similarity 171 member A2 [Source:HGNC Symbol;Acc:HGNC:30480] | -0.8838 | 0.00024 |
| ENSG00000109906 | ZBTB16 | zinc finger and BTB domain containing 16 [Source:HGNC Symbol;Acc:HGNC:12930] | -1.4563 | 0.00024 |
| ENSG00000164989 | CCDC171 | coiled-coil domain containing 171 [Source:HGNC Symbol;Acc:HGNC:29828] | -1.3993 | 0.00024 |
| ENSG00000147324 | MFHAS1 | malignant fibrous histiocytoma amplified sequence 1 [Source:HGNC Symbol;Acc:HGNC:16982] | -0.8739 | 0.00024 |
| ENSG00000176490 | DIRAS1 | DIRAS family GTPase 1 [Source:HGNC Symbol;Acc:HGNC:19127] | -0.6695 | 0.00024 |
| ENSG00000107105 | ELAVL2 | ELAV like RNA binding protein 2 [Source:HGNC Symbol;Acc:HGNC:3313] | 0.5960 | 0.00025 |
| ENSG00000105641 | SLC5A5 | solute carrier family 5 member 5 [Source:HGNC Symbol;Acc:HGNC:11040] | -1.9062 | 0.00025 |
| ENSG00000147041 | SYTL5 | synaptotagmin like 5 [Source:HGNC Symbol;Acc:HGNC:15589] | 2.7612 | 0.00025 |
| ENSG00000188613 | NANOS1 | nanos C2HC-type zinc finger 1 [Source:HGNC Symbol;Acc:HGNC:23044] | -1.4874 | 0.00025 |
| ENSG00000250510 | GPR162 | G protein-coupled receptor 162 [Source:HGNC Symbol;Acc:HGNC:16693] | -0.6953 | 0.00025 |
| ENSG00000205060 | SLC35B4 | solute carrier family 35 member B4 [Source:HGNC Symbol;Acc:HGNC:20584] | -0.5789 | 0.00025 |
| ENSG00000269834 | ZNF528-AS1 | ZNF528 antisense RNA 1 [Source:HGNC Symbol;Acc:HGNC:51305] | -0.6790 | 0.00025 |
| ENSG00000105664 | COMP | cartilage oligomeric matrix protein [Source:HGNC Symbol;Acc:HGNC:2227] | -2.0534 | 0.00026 |
| ENSG00000188878 | FBF1 | Fas binding factor 1 [Source:HGNC Symbol;Acc:HGNC:24674] | -0.6760 | 0.00026 |
| ENSG00000268996 | MAN1B1-DT | MAN1B1 divergent transcript [Source:HGNC Symbol;Acc:HGNC:48715] | -0.6286 | 0.00026 |
| ENSG00000183751 | TBL3 | transducin beta like 3 [Source:HGNC Symbol;Acc:HGNC:11587] | -0.5050 | 0.00026 |
| ENSG00000187730 | GABRD | gamma-aminobutyric acid type A receptor subunit delta [Source:HGNC Symbol;Acc:HGNC:4084] | 2.6636 | 0.00026 |

| | | | | |
|---|---|---|---|---|
| ENSG00000155629 | PIK3AP1 | phosphoinositide-3-kinase adaptor protein 1 [Source:HGNC Symbol;Acc:HGNC:30034] | -1.1243 | 0.00026 |
| ENSG00000276846 | NA | novel transcript, antisense to HKR1 | -1.5132 | 0.00026 |
| ENSG00000259363 | NA | novel transcript | -0.8522 | 0.00027 |
| ENSG00000198056 | PRIM1 | DNA primase subunit 1 [Source:HGNC Symbol;Acc:HGNC:9369] | -0.5456 | 0.00027 |
| ENSG00000054277 | OPN3 | opsin 3 [Source:HGNC Symbol;Acc:HGNC:14007] | -0.5509 | 0.00027 |
| ENSG00000258818 | RNASE4 | ribonuclease A family member 4 [Source:HGNC Symbol;Acc:HGNC:10047] | -1.4782 | 0.00027 |
| ENSG00000267052 | NA | novel transcript | 2.6518 | 0.00027 |
| ENSG00000165300 | SLITRK5 | SLIT and NTRK like family member 5 [Source:HGNC Symbol;Acc:HGNC:20295] | -0.8049 | 0.00027 |
| ENSG00000115425 | PECR | peroxisomal trans-2-enoyl-CoA reductase [Source:HGNC Symbol;Acc:HGNC:18281] | -0.6303 | 0.00027 |
| ENSG00000128045 | RASL11B | RAS like family 11 member B [Source:HGNC Symbol;Acc:HGNC:23804] | -1.0312 | 0.00027 |
| ENSG00000129595 | EPB41L4A | erythrocyte membrane protein band 4.1 like 4A [Source:HGNC Symbol;Acc:HGNC:13278] | 0.5866 | 0.00027 |
| ENSG00000285190 | NA | novel transcript, antisense to CCNH | 1.6528 | 0.00027 |
| ENSG00000242615 | NA | ribosomal protein L17 (RPL17) pseudogene | -1.0167 | 0.00027 |
| ENSG00000170820 | FSHR | follicle stimulating hormone receptor [Source:HGNC Symbol;Acc:HGNC:3969] | -1.1366 | 0.00028 |
| ENSG00000253227 | NA | novel transcript | 5.9421 | 0.00028 |
| ENSG00000164465 | DCBLD1 | discoidin, CUB and LCCL domain containing 1 [Source:HGNC Symbol;Acc:HGNC:21479] | -0.5907 | 0.00028 |
| ENSG00000167074 | TEF | TEF transcription factor, PAR bZIP family member [Source:HGNC Symbol;Acc:HGNC:11722] | -0.9886 | 0.00028 |
| ENSG00000172671 | ZFAND4 | zinc finger AN1-type containing 4 [Source:HGNC Symbol;Acc:HGNC:23504] | -0.6879 | 0.00028 |
| ENSG00000183054 | RGPD6 | RANBP2 like and GRIP domain containing 6 [Source:HGNC Symbol;Acc:HGNC:32419] | 0.7619 | 0.00028 |
| ENSG00000176136 | MC5R | melanocortin 5 receptor [Source:HGNC Symbol;Acc:HGNC:6933] | 2.4613 | 0.00028 |
| ENSG00000166326 | TRIM44 | tripartite motif containing 44 [Source:HGNC Symbol;Acc:HGNC:19016] | 0.6094 | 0.00028 |
| ENSG00000198934 | MAGEE1 | MAGE family member E1 [Source:HGNC Symbol;Acc:HGNC:24934] | -0.5704 | 0.00028 |
| ENSG00000165806 | CASP7 | caspase 7 [Source:HGNC Symbol;Acc:HGNC:1508] | 0.5578 | 0.00028 |
| ENSG00000272870 | SAP30-DT | SAP30 divergent transcript [Source:HGNC Symbol;Acc:HGNC:54424] | -1.4033 | 0.00029 |
| ENSG00000128596 | CCDC136 | coiled-coil domain containing 136 [Source:HGNC Symbol;Acc:HGNC:22225] | 0.6189 | 0.00029 |
| ENSG00000105499 | PLA2G4C | phospholipase A2 group IVC [Source:HGNC Symbol;Acc:HGNC:9037] | 1.0050 | 0.00029 |
| ENSG00000100385 | IL2RB | interleukin 2 receptor subunit beta [Source:HGNC Symbol;Acc:HGNC:6009] | 4.2629 | 0.00029 |
| ENSG00000101210 | EEF1A2 | eukaryotic translation elongation factor 1 alpha 2 [Source:HGNC Symbol;Acc:HGNC:3192] | 0.6148 | 0.00029 |
| ENSG00000142871 | CCN1 | cellular communication network factor 1 [Source:HGNC Symbol;Acc:HGNC:2654] | 1.7487 | 0.00029 |
| ENSG00000111300 | NAA25 | N-alpha-acetyltransferase 25, NatB auxiliary subunit [Source:HGNC Symbol;Acc:HGNC:25783] | 0.5179 | 0.00029 |
| ENSG00000143156 | NME7 | NME/NM23 family member 7 [Source:HGNC Symbol;Acc:HGNC:20461] | 0.5751 | 0.00029 |
| ENSG00000131781 | FMO5 | flavin containing dimethylaniline monoxygenase 5 [Source:HGNC Symbol;Acc:HGNC:3773] | -1.1087 | 0.00030 |
| ENSG00000167700 | MFSD3 | major facilitator superfamily domain containing 3 [Source:HGNC Symbol;Acc:HGNC:25157] | -0.7371 | 0.00030 |
| ENSG00000126010 | GRPR | gastrin releasing peptide receptor [Source:HGNC Symbol;Acc:HGNC:4609] | -1.4637 | 0.00030 |
| ENSG00000166415 | WDR72 | WD repeat domain 72 [Source:HGNC Symbol;Acc:HGNC:26790] | -0.6745 | 0.00030 |
| ENSG00000285467 | NA | dehydrogenase/reductase 4 like 1 [Source:NCBI gene (formerly Entrezgene);Acc:728635] | -1.8956 | 0.00030 |
| ENSG00000227487 | NCAM1-AS1 | NCAM1 antisense RNA1 [Source:HGNC Symbol;Acc:HGNC:48675] | 0.8413 | 0.00030 |
| ENSG00000234383 | CTBP2P8 | CTBP2 pseudogene 8 [Source:HGNC Symbol;Acc:HGNC:45200] | 2.2006 | 0.00031 |
| ENSG00000218226 | TATDN2P2 | TatD DNase domain containing 2 pseudogene 2 [Source:HGNC Symbol;Acc:HGNC:39255] | 1.0009 | 0.00031 |
| ENSG00000103154 | NECAB2 | N-terminal EF-hand calcium binding protein 2 [Source:HGNC Symbol;Acc:HGNC:23746] | 1.3965 | 0.00031 |
| ENSG00000161149 | TUBA3FP | tubulin alpha 3f pseudogene [Source:HGNC Symbol;Acc:HGNC:24067] | -2.9039 | 0.00031 |
| ENSG00000186567 | CEACAM19 | CEA cell adhesion molecule 19 [Source:HGNC Symbol;Acc:HGNC:31951] | 1.2041 | 0.00031 |
| ENSG00000137274 | BPHL | biphenyl hydrolase like [Source:HGNC Symbol;Acc:HGNC:1094] | -0.5574 | 0.00031 |

| | | | | |
|---|---|---|---|---|
| ENSG00000091128 | LAMB4 | laminin subunit beta 4 [Source:HGNC Symbol;Acc:HGNC:6491] | -1.0745 | 0.00032 |
| ENSG00000245849 | RAD51-AS1 | RAD51 antisense RNA 1 [Source:HGNC Symbol;Acc:HGNC:48621] | -0.6680 | 0.00032 |
| ENSG00000232300 | FAM215B | family with sequence similarity 215 member B [Source:HGNC Symbol;Acc:HGNC:43639] | 0.8159 | 0.00032 |
| ENSG00000103174 | NAGPA | N-acetylglucosamine-1-phosphodiester alpha-N-acetylglucosaminidase [Source:HGNC Symbol;Acc:HGNC:17378] | -0.6953 | 0.00032 |
| ENSG00000251081 | NA | novel transcript | -0.9775 | 0.00033 |
| ENSG00000238207 | NA | novel transcript | -1.0734 | 0.00033 |
| ENSG00000228801 | NA | novel transcript | -0.7802 | 0.00033 |
| ENSG00000235636 | NUS1P1 | NUS1 pseudogene 1 [Source:HGNC Symbol;Acc:HGNC:38472] | 0.9348 | 0.00033 |
| ENSG00000131153 | GINS2 | GINS complex subunit 2 [Source:HGNC Symbol;Acc:HGNC:24575] | -0.6068 | 0.00034 |
| ENSG00000256417 | NA | novel transcript | -1.7646 | 0.00034 |
| ENSG00000275549 | STPG3-AS1 | STPG3 antisense RNA 1 [Source:HGNC Symbol;Acc:HGNC:51176] | -1.2839 | 0.00034 |
| ENSG00000196366 | C9orf163 | chromosome 9 open reading frame 163 [Source:HGNC Symbol;Acc:HGNC:26718] | -1.0214 | 0.00034 |
| ENSG00000138311 | ZNF365 | zinc finger protein 365 [Source:HGNC Symbol;Acc:HGNC:18194] | 0.9180 | 0.00034 |
| ENSG00000079102 | RUNX1T1 | RUNX1 partner transcriptional co-repressor 1 [Source:HGNC Symbol;Acc:HGNC:1535] | -1.4197 | 0.00035 |
| ENSG00000242600 | MBL1P | mannose binding lectin 1, pseudogene [Source:HGNC Symbol;Acc:HGNC:6921] | 1.9390 | 0.00035 |
| ENSG00000231584 | FAHD2CP | fumarylacetoacetate hydrolase domain containing 2C, pseudogene [Source:HGNC Symbol;Acc:HGNC:44135] | -1.5471 | 0.00035 |
| ENSG00000055955 | ITIH4 | inter-alpha-trypsin inhibitor heavy chain 4 [Source:HGNC Symbol;Acc:HGNC:6169] | -1.0067 | 0.00035 |
| ENSG00000235162 | C12orf75 | chromosome 12 open reading frame 75 [Source:HGNC Symbol;Acc:HGNC:35164] | 0.5137 | 0.00036 |
| ENSG00000143375 | CGN | cingulin [Source:HGNC Symbol;Acc:HGNC:17429] | -0.9924 | 0.00036 |
| ENSG00000125510 | OPRL1 | opioid related nociceptin receptor 1 [Source:HGNC Symbol;Acc:HGNC:8155] | -2.0498 | 0.00036 |
| ENSG00000089250 | NOS1 | nitric oxide synthase 1 [Source:HGNC Symbol;Acc:HGNC:7872] | 4.4483 | 0.00036 |
| ENSG00000227543 | SPAG5-AS1 | SPAG5 antisense RNA 1 [Source:HGNC Symbol;Acc:HGNC:41140] | 0.7135 | 0.00036 |
| ENSG00000166704 | ZNF606 | zinc finger protein 606 [Source:HGNC Symbol;Acc:HGNC:25879] | -0.5517 | 0.00036 |
| ENSG00000262528 | NA | novel transcript, antisense to WDR90 | -0.7803 | 0.00036 |
| ENSG00000261211 | NA | novel transcript | -0.8421 | 0.00036 |
| ENSG00000250303 | LINC02762 | long intergenic non-protein coding RNA 2762 [Source:HGNC Symbol;Acc:HGNC:27443] | -0.6226 | 0.00036 |
| ENSG00000106125 | MINDY4 | MINDY lysine 48 deubiquitinase 4 [Source:HGNC Symbol;Acc:HGNC:21916] | -0.5198 | 0.00036 |
| ENSG00000227398 | KIF9-AS1 | KIF9 antisense RNA 1 [Source:HGNC Symbol;Acc:HGNC:26822] | -0.6432 | 0.00037 |
| ENSG00000152969 | JAKMIP1 | janus kinase and microtubule interacting protein 1 [Source:HGNC Symbol;Acc:HGNC:26460] | -1.6086 | 0.00037 |
| ENSG00000126970 | ZC4H2 | zinc finger C4H2-type containing [Source:HGNC Symbol;Acc:HGNC:24931] | -0.5687 | 0.00037 |
| ENSG00000149548 | CCDC15 | coiled-coil domain containing 15 [Source:HGNC Symbol;Acc:HGNC:25798] | -0.9679 | 0.00037 |
| ENSG00000167642 | SPINT2 | serine peptidase inhibitor, Kunitz type 2 [Source:HGNC Symbol;Acc:HGNC:11247] | -0.5404 | 0.00037 |
| ENSG00000119917 | IFIT3 | interferon induced protein with tetratricopeptide repeats 3 [Source:HGNC Symbol;Acc:HGNC:5411] | 0.6999 | 0.00038 |
| ENSG00000271590 | NA | novel transcript | 1.2078 | 0.00038 |
| ENSG00000168461 | RAB31 | RAB31, member RAS oncogene family [Source:HGNC Symbol;Acc:HGNC:9771] | 0.7394 | 0.00038 |
| ENSG00000251247 | ZNF345 | zinc finger protein 345 [Source:HGNC Symbol;Acc:HGNC:16367] | -0.6315 | 0.00038 |
| ENSG00000255983 | NA | novel transcript | -1.4188 | 0.00038 |
| ENSG00000164303 | ENPP6 | ectonucleotide pyrophosphatase/phosphodiesterase 6 [Source:HGNC Symbol;Acc:HGNC:23409] | -0.9677 | 0.00038 |
| ENSG00000165813 | CCDC186 | coiled-coil domain containing 186 [Source:HGNC Symbol;Acc:HGNC:24349] | 0.6627 | 0.00039 |
| ENSG00000168014 | C2CD3 | C2 domain containing 3 centriole elongation regulator [Source:HGNC Symbol;Acc:HGNC:24564] | 0.5977 | 0.00039 |
| ENSG00000169598 | DFFB | DNA fragmentation factor subunit beta [Source:HGNC Symbol;Acc:HGNC:2773] | -1.0952 | 0.00039 |
| ENSG00000134253 | TRIM45 | tripartite motif containing 45 [Source:HGNC Symbol;Acc:HGNC:19018] | -0.9237 | 0.00039 |
| ENSG00000176912 | TYMSOS | TYMS opposite strand [Source:HGNC Symbol;Acc:HGNC:29553] | -1.4950 | 0.00039 |

| | | | | |
|---|---|---|---|---|
| ENSG00000176533 | GNG7 | G protein subunit gamma 7 [Source:HGNC Symbol;Acc:HGNC:4410] | -1.9319 | 0.00039 |
| ENSG00000148985 | PGAP2 | post-GPI attachment to proteins 2 [Source:HGNC Symbol;Acc:HGNC:17893] | -0.9461 | 0.00040 |
| ENSG00000271013 | LRRC37A9P | leucine rich repeat containing 37 member A9, pseudogene [Source:HGNC Symbol;Acc:HGNC:43813] | -1.4910 | 0.00040 |
| ENSG00000141560 | FN3KRP | fructosamine 3 kinase related protein [Source:HGNC Symbol;Acc:HGNC:25700] | -0.5199 | 0.00040 |
| ENSG00000261373 | VPS9D1-AS1 | VPS9D1 antisense RNA 1 [Source:HGNC Symbol;Acc:HGNC:48915] | 0.6871 | 0.00040 |
| ENSG00000100116 | GCAT | glycine C-acetyltransferase [Source:HGNC Symbol;Acc:HGNC:4188] | 0.5301 | 0.00041 |
| ENSG00000166165 | CKB | creatine kinase B [Source:HGNC Symbol;Acc:HGNC:1991] | 0.5981 | 0.00041 |
| ENSG00000196876 | SCN8A | sodium voltage-gated channel alpha subunit 8 [Source:HGNC Symbol;Acc:HGNC:10596] | -0.6614 | 0.00041 |
| ENSG00000119042 | SATB2 | SATB homeobox 2 [Source:HGNC Symbol;Acc:HGNC:21637] | 0.6721 | 0.00041 |
| ENSG00000197063 | MAFG | MAF bZIP transcription factor G [Source:HGNC Symbol;Acc:HGNC:6781] | 0.5743 | 0.00042 |
| ENSG00000117010 | ZNF684 | zinc finger protein 684 [Source:HGNC Symbol;Acc:HGNC:28418] | 0.6164 | 0.00042 |
| ENSG00000247134 | NA | novel transcript | -0.6195 | 0.00042 |
| ENSG00000203392 | NA | novel transcript, antisense to CSPG4 | 3.4652 | 0.00042 |
| ENSG00000246859 | STARD4-AS1 | STARD4 antisense RNA 1 [Source:HGNC Symbol;Acc:HGNC:44117] | 0.5851 | 0.00043 |
| ENSG00000250685 | NA | novel transcript, antisense to ADAD2 | -4.1415 | 0.00043 |
| ENSG00000147852 | VLDLR | very low density lipoprotein receptor [Source:HGNC Symbol;Acc:HGNC:12698] | -0.8999 | 0.00043 |
| ENSG00000116544 | DLGAP3 | DLG associated protein 3 [Source:HGNC Symbol;Acc:HGNC:30368] | -0.7410 | 0.00043 |
| ENSG00000160401 | CFAP157 | cilia and flagella associated protein 157 [Source:HGNC Symbol;Acc:HGNC:27843] | -0.8275 | 0.00044 |
| ENSG00000174718 | RESF1 | retroelement silencing factor 1 [Source:HGNC Symbol;Acc:HGNC:25559] | -0.5491 | 0.00044 |
| ENSG00000266469 | NA | novel transcript, antisense to MED1 & FBXL20 | -5.8815 | 0.00044 |
| ENSG00000121743 | GJA3 | gap junction protein alpha 3 [Source:HGNC Symbol;Acc:HGNC:4277] | 1.8917 | 0.00044 |
| ENSG00000227354 | RBM26-AS1 | RBM26 antisense RNA 1 [Source:HGNC Symbol;Acc:HGNC:39805] | 0.5888 | 0.00044 |
| ENSG00000106733 | NMRK1 | nicotinamide riboside kinase 1 [Source:HGNC Symbol;Acc:HGNC:26057] | -1.0914 | 0.00044 |
| ENSG00000124151 | NCOA3 | nuclear receptor coactivator 3 [Source:HGNC Symbol;Acc:HGNC:7670] | 0.5757 | 0.00044 |
| ENSG00000168538 | TRAPPC11 | trafficking protein particle complex 11 [Source:HGNC Symbol;Acc:HGNC:25751] | -0.6774 | 0.00044 |
| ENSG00000132481 | TRIM47 | tripartite motif containing 47 [Source:HGNC Symbol;Acc:HGNC:19020] | 0.5441 | 0.00045 |
| ENSG00000128039 | SRD5A3 | steroid 5 alpha-reductase 3 [Source:HGNC Symbol;Acc:HGNC:25812] | -0.7614 | 0.00045 |
| ENSG00000181035 | SLC25A42 | solute carrier family 25 member 42 [Source:HGNC Symbol;Acc:HGNC:28380] | -0.5451 | 0.00045 |
| ENSG00000133874 | RNF122 | ring finger protein 122 [Source:HGNC Symbol;Acc:HGNC:21147] | -0.6437 | 0.00045 |
| ENSG00000166106 | ADAMTS15 | ADAM metallopeptidase with thrombospondin type 1 motif 15 [Source:HGNC Symbol;Acc:HGNC:16305] | -1.5109 | 0.00045 |
| ENSG00000205744 | DENND1C | DENN domain containing 1C [Source:HGNC Symbol;Acc:HGNC:26225] | -0.9762 | 0.00045 |
| ENSG00000135824 | RGS8 | regulator of G protein signaling 8 [Source:HGNC Symbol;Acc:HGNC:16810] | -1.9259 | 0.00045 |
| ENSG00000132837 | DMGDH | dimethylglycine dehydrogenase [Source:HGNC Symbol;Acc:HGNC:24475] | -1.7286 | 0.00046 |
| ENSG00000286329 | NA | novel transcript | -0.9229 | 0.00046 |
| ENSG00000204740 | MALRD1 | MAM and LDL receptor class A domain containing 1 [Source:HGNC Symbol;Acc:HGNC:24331] | -0.9878 | 0.00046 |
| ENSG00000185875 | THNSL1 | threonine synthase like 1 [Source:HGNC Symbol;Acc:HGNC:26160] | -0.5315 | 0.00046 |
| ENSG00000059122 | FLYWCH1 | FLYWCH-type zinc finger 1 [Source:HGNC Symbol;Acc:HGNC:25404] | -0.7240 | 0.00046 |
| ENSG00000109103 | UNC119 | unc-119 lipid binding chaperone [Source:HGNC Symbol;Acc:HGNC:12565] | -0.6060 | 0.00046 |
| ENSG00000187123 | LYPD6 | LY6/PLAUR domain containing 6 [Source:HGNC Symbol;Acc:HGNC:28751] | -1.8847 | 0.00046 |
| ENSG00000225872 | LINC01529 | long intergenic non-protein coding RNA 1529 [Source:HGNC Symbol;Acc:HGNC:51268] | -1.0025 | 0.00047 |
| ENSG00000188620 | HMX3 | H6 family homeobox 3 [Source:HGNC Symbol;Acc:HGNC:5019] | 1.7389 | 0.00047 |
| ENSG00000271605 | MILR1 | mast cell immunoglobulin like receptor 1 [Source:HGNC Symbol;Acc:HGNC:27570] | 3.1237 | 0.00047 |
| ENSG00000029559 | IBSP | integrin binding sialoprotein [Source:HGNC Symbol;Acc:HGNC:5341] | -2.0420 | 0.00047 |

| | | | | |
|---|---|---|---|---|
| ENSG00000178460 | MCMDC2 | minichromosome maintenance domain containing 2 [Source:HGNC Symbol;Acc:HGNC:26368] | -1.5513 | 0.00047 |
| ENSG00000196167 | COLCA1 | colorectal cancer associated 1 [Source:HGNC Symbol;Acc:HGNC:33789] | -1.7614 | 0.00047 |
| ENSG00000099219 | ERMP1 | endoplasmic reticulum metallopeptidase 1 [Source:HGNC Symbol;Acc:HGNC:23703] | -0.5256 | 0.00047 |
| ENSG00000102760 | RGCC | regulator of cell cycle [Source:HGNC Symbol;Acc:HGNC:20369] | 3.3157 | 0.00047 |
| ENSG00000287134 | NA | novel transcript, antisense to DLC1 | 1.7537 | 0.00048 |
| ENSG00000140104 | CLBA1 | clathrin binding box of aftiphilin containing 1 [Source:HGNC Symbol;Acc:HGNC:20126] | 0.6733 | 0.00048 |
| ENSG00000089094 | KDM2B | lysine demethylase 2B [Source:HGNC Symbol;Acc:HGNC:13610] | -0.5489 | 0.00048 |
| ENSG00000279667 | NA | TEC | 2.5228 | 0.00048 |
| ENSG00000082146 | STRADB | STE20 related adaptor beta [Source:HGNC Symbol;Acc:HGNC:13205] | -0.6867 | 0.00048 |
| ENSG00000166352 | IFTAP | intraflagellar transport associated protein [Source:HGNC Symbol;Acc:HGNC:25142] | -0.7089 | 0.00048 |
| ENSG00000170161 | NA | uncharacterized LOC554249 [Source:NCBI gene (formerly Entrezgene);Acc:554249] | 0.7901 | 0.00049 |
| ENSG00000248383 | PCDHAC1 | protocadherin alpha subfamily C, 1 [Source:HGNC Symbol;Acc:HGNC:8676] | -0.6540 | 0.00049 |
| ENSG00000102595 | UGGT2 | UDP-glucose glycoprotein glucosyltransferase 2 [Source:HGNC Symbol;Acc:HGNC:15664] | 0.5415 | 0.00049 |
| ENSG00000089916 | GPATCH2L | G-patch domain containing 2 like [Source:HGNC Symbol;Acc:HGNC:20210] | 0.6796 | 0.00050 |
| ENSG00000109738 | GLRB | glycine receptor beta [Source:HGNC Symbol;Acc:HGNC:4329] | -0.7503 | 0.00050 |
| ENSG00000287151 | NA | chromosome 2 open reading frame 27A [Source:NCBI gene (formerly Entrezgene);Acc:29798] | 0.7280 | 0.00050 |
| ENSG00000114547 | ROPN1B | rhophilin associated tail protein 1B [Source:HGNC Symbol;Acc:HGNC:31927] | -1.2904 | 0.00050 |
| ENSG00000219626 | FAM228B | family with sequence similarity 228 member B [Source:HGNC Symbol;Acc:HGNC:24736] | -0.9269 | 0.00050 |
| ENSG00000261308 | FIGNL2 | fidgetin like 2 [Source:HGNC Symbol;Acc:HGNC:13287] | 0.8188 | 0.00050 |
| ENSG00000176907 | TCIM | transcriptional and immune response regulator [Source:HGNC Symbol;Acc:HGNC:1357] | 1.4637 | 0.00050 |
| ENSG00000110881 | ASIC1 | acid sensing ion channel subunit 1 [Source:HGNC Symbol;Acc:HGNC:100] | -1.1722 | 0.00050 |
| ENSG00000161888 | SPC24 | SPC24 component of NDC80 kinetochore complex [Source:HGNC Symbol;Acc:HGNC:26913] | -0.8278 | 0.00050 |
| ENSG00000102879 | CORO1A | coronin 1A [Source:HGNC Symbol;Acc:HGNC:2252] | 1.3547 | 0.00051 |
| ENSG00000287126 | NA | novel transcript | -0.8252 | 0.00051 |
| ENSG00000248127 | NA | novel transcript | 1.0996 | 0.00051 |
| ENSG00000165996 | HACD1 | 3-hydroxyacyl-CoA dehydratase 1 [Source:HGNC Symbol;Acc:HGNC:9639] | 0.6311 | 0.00052 |
| ENSG00000158486 | DNAH3 | dynein axonemal heavy chain 3 [Source:HGNC Symbol;Acc:HGNC:2949] | -2.0469 | 0.00052 |
| ENSG00000113946 | CLDN16 | claudin 16 [Source:HGNC Symbol;Acc:HGNC:2037] | -1.8423 | 0.00052 |
| ENSG00000229848 | NA | novel transcript, antisense to ACTG1 | 1.6289 | 0.00052 |
| ENSG00000100628 | ASB2 | ankyrin repeat and SOCS box containing 2 [Source:HGNC Symbol;Acc:HGNC:16012] | 3.7095 | 0.00052 |
| ENSG00000106633 | GCK | glucokinase [Source:HGNC Symbol;Acc:HGNC:4195] | -1.0799 | 0.00052 |
| ENSG00000145349 | CAMK2D | calcium/calmodulin dependent protein kinase II delta [Source:HGNC Symbol;Acc:HGNC:1462] | -0.5207 | 0.00052 |
| ENSG00000235897 | TM4SF19-AS1 | TM4SF19 antisense RNA 1 [Source:HGNC Symbol;Acc:HGNC:41085] | -3.0542 | 0.00052 |
| ENSG00000272668 | NA | uncharacterized LOC107985216 [Source:NCBI gene (formerly Entrezgene);Acc:107985216] | -1.4848 | 0.00053 |
| ENSG00000108187 | PBLD | phenazine biosynthesis like protein domain containing [Source:HGNC Symbol;Acc:HGNC:23301] | -0.8198 | 0.00053 |
| ENSG00000130475 | FCHO1 | FCH and mu domain containing endocytic adaptor 1 [Source:HGNC Symbol;Acc:HGNC:29002] | -0.9369 | 0.00053 |
| ENSG00000135346 | CGA | glycoprotein hormones, alpha polypeptide [Source:HGNC Symbol;Acc:HGNC:1885] | 3.4026 | 0.00053 |
| ENSG00000152518 | ZFP36L2 | ZFP36 ring finger protein like 2 [Source:HGNC Symbol;Acc:HGNC:1108] | 0.5934 | 0.00053 |
| ENSG00000185839 | NA | adenylate kinase 2 (AK2) pseudogene | 0.9501 | 0.00054 |
| ENSG00000173404 | INSM1 | INSM transcriptional repressor 1 [Source:HGNC Symbol;Acc:HGNC:6090] | -2.7520 | 0.00054 |
| ENSG00000137573 | SULF1 | sulfatase 1 [Source:HGNC Symbol;Acc:HGNC:20391] | -0.9379 | 0.00054 |
| ENSG00000273899 | NOL12 | nucleolar protein 12 [Source:HGNC Symbol;Acc:HGNC:28585] | 0.5342 | 0.00054 |
| ENSG00000237651 | C2orf74 | chromosome 2 open reading frame 74 [Source:HGNC Symbol;Acc:HGNC:34439] | -0.7680 | 0.00054 |

| | | | | |
|---|---|---|---|---|
| ENSG00000165097 | KDM1B | lysine demethylase 1B [Source:HGNC Symbol;Acc:HGNC:21577] | -0.5188 | 0.00054 |
| ENSG00000104356 | POP1 | POP1 homolog, ribonuclease P/MRP subunit [Source:HGNC Symbol;Acc:HGNC:30129] | 0.6247 | 0.00054 |
| ENSG00000228775 | WEE2-AS1 | WEE2 antisense RNA 1 [Source:HGNC Symbol;Acc:HGNC:48669] | -1.0511 | 0.00054 |
| ENSG00000189409 | MMP23B | matrix metallopeptidase 23B [Source:HGNC Symbol;Acc:HGNC:7171] | 1.5234 | 0.00054 |
| ENSG00000187024 | PTRH1 | peptidyl-tRNA hydrolase 1 homolog [Source:HGNC Symbol;Acc:HGNC:27039] | -0.6012 | 0.00054 |
| ENSG00000128591 | FLNC | filamin C [Source:HGNC Symbol;Acc:HGNC:3756] | 1.8775 | 0.00055 |
| ENSG00000198416 | ZNF658B | zinc finger protein 658B (pseudogene) [Source:HGNC Symbol;Acc:HGNC:32033] | -0.8371 | 0.00055 |
| ENSG00000225377 | NRSN2-AS1 | NRSN2 antisense RNA 1 [Source:HGNC Symbol;Acc:HGNC:51222] | -0.6249 | 0.00055 |
| ENSG00000131788 | PIAS3 | protein inhibitor of activated STAT 3 [Source:HGNC Symbol;Acc:HGNC:16861] | -0.7860 | 0.00055 |
| ENSG00000088992 | TESC | tescalcin [Source:HGNC Symbol;Acc:HGNC:26065] | -0.5287 | 0.00055 |
| ENSG00000237988 | OR2I1P | olfactory receptor family 2 subfamily I member 1 pseudogene [Source:HGNC Symbol;Acc:HGNC:8258] | -5.2701 | 0.00055 |
| ENSG00000134802 | SLC43A3 | solute carrier family 43 member 3 [Source:HGNC Symbol;Acc:HGNC:17466] | 0.7504 | 0.00055 |
| ENSG00000132718 | SYT11 | synaptotagmin 11 [Source:HGNC Symbol;Acc:HGNC:19239] | -0.8760 | 0.00055 |
| ENSG00000165521 | EML5 | EMAP like 5 [Source:HGNC Symbol;Acc:HGNC:18197] | -0.5105 | 0.00055 |
| ENSG00000140323 | DISP2 | dispatched RND transporter family member 2 [Source:HGNC Symbol;Acc:HGNC:19712] | -1.1116 | 0.00055 |
| ENSG00000229132 | EIF4A1P10 | eukaryotic translation initiation factor 4A1 pseudogene 10 [Source:HGNC Symbol;Acc:HGNC:37930] | 0.7199 | 0.00056 |
| ENSG00000257732 | NA | novel transcript, antisense to TXNRD1 | 4.3963 | 0.00056 |
| ENSG00000227155 | NA | novel transcript | 2.3306 | 0.00056 |
| ENSG00000166452 | AKIP1 | A-kinase interacting protein 1 [Source:HGNC Symbol;Acc:HGNC:1170] | -0.5709 | 0.00056 |
| ENSG00000167720 | SRR | serine racemase [Source:HGNC Symbol;Acc:HGNC:14398] | -0.7023 | 0.00056 |
| ENSG00000129932 | DOHH | deoxyhypusine hydroxylase [Source:HGNC Symbol;Acc:HGNC:28662] | 0.5015 | 0.00056 |
| ENSG00000224945 | NA | novel transcript | 1.4661 | 0.00056 |
| ENSG00000115590 | IL1R2 | interleukin 1 receptor type 2 [Source:HGNC Symbol;Acc:HGNC:5994] | -1.1065 | 0.00056 |
| ENSG00000259030 | FPGT-TNNI3K | FPGT-TNNI3K readthrough [Source:HGNC Symbol;Acc:HGNC:42952] | -1.9038 | 0.00056 |
| ENSG00000146243 | IRAK1BP1 | interleukin 1 receptor associated kinase 1 binding protein 1 [Source:HGNC Symbol;Acc:HGNC:17368] | -0.6660 | 0.00058 |
| ENSG00000168135 | KCNJ4 | potassium inwardly rectifying channel subfamily J member 4 [Source:HGNC Symbol;Acc:HGNC:6265] | -0.8238 | 0.00058 |
| ENSG00000268592 | NA | RAET1E antisense RNA 1 [Source:NCBI gene (formerly Entrezgene);Acc:100652739] | -1.8537 | 0.00058 |
| ENSG00000284024 | NA | heat shock protein family A (Hsp70) member 14 [Source:NCBI gene (formerly Entrezgene);Acc:51182] | 0.5069 | 0.00058 |
| ENSG00000127564 | PKMYT1 | protein kinase, membrane associated tyrosine/threonine 1 [Source:HGNC Symbol;Acc:HGNC:29650] | -0.7128 | 0.00059 |
| ENSG00000279041 | NA | TEC | 0.8824 | 0.00059 |
| ENSG00000230699 | NA | novel transcript | -1.4712 | 0.00059 |
| ENSG00000158163 | DZIP1L | DAZ interacting zinc finger protein 1 like [Source:HGNC Symbol;Acc:HGNC:26551] | -0.5173 | 0.00059 |
| ENSG00000117480 | FAAH | fatty acid amide hydrolase [Source:HGNC Symbol;Acc:HGNC:3553] | -0.5164 | 0.00060 |
| ENSG00000183979 | NPB | neuropeptide B [Source:HGNC Symbol;Acc:HGNC:30099] | -2.1854 | 0.00060 |
| ENSG00000163870 | TPRA1 | transmembrane protein adipocyte associated 1 [Source:HGNC Symbol;Acc:HGNC:30413] | -0.6000 | 0.00060 |
| ENSG00000244128 | LINC01322 | long intergenic non-protein coding RNA 1322 [Source:HGNC Symbol;Acc:HGNC:50528] | -0.8998 | 0.00060 |
| ENSG00000188549 | CCDC9B | coiled-coil domain containing 9B [Source:HGNC Symbol;Acc:HGNC:33488] | -1.3056 | 0.00060 |
| ENSG00000163517 | HDAC11 | histone deacetylase 11 [Source:HGNC Symbol;Acc:HGNC:19086] | -0.8588 | 0.00061 |
| ENSG00000261879 | NA | novel transcript, antisense to C17orf87 | -1.1848 | 0.00061 |
| ENSG00000235944 | ZNF815P | zinc finger protein 815, pseudogene [Source:HGNC Symbol;Acc:HGNC:22029] | -1.3274 | 0.00061 |
| ENSG00000128891 | CCDC32 | coiled-coil domain containing 32 [Source:HGNC Symbol;Acc:HGNC:28295] | 0.7199 | 0.00061 |
| ENSG00000237149 | ZNF503-AS2 | ZNF503 antisense RNA 2 [Source:HGNC Symbol;Acc:HGNC:23525] | 0.6181 | 0.00062 |
| ENSG00000112379 | ARFGEF3 | ARFGEF family member 3 [Source:HGNC Symbol;Acc:HGNC:21213] | 2.3095 | 0.00062 |

| | | | | |
|---|---|---|---|---|
| ENSG00000223956 | LINC01767 | long intergenic non-protein coding RNA 1767 [Source:HGNC Symbol;Acc:HGNC:52557] | 3.7040 | 0.00062 |
| ENSG00000224281 | SLC25A5-AS1 | SLC25A5 antisense RNA 1 [Source:HGNC Symbol;Acc:HGNC:43438] | -0.8645 | 0.00062 |
| ENSG00000203999 | LINC01270 | long intergenic non-protein coding RNA 1270 [Source:HGNC Symbol;Acc:HGNC:27658] | -1.3808 | 0.00063 |
| ENSG00000081803 | CADPS2 | calcium dependent secretion activator 2 [Source:HGNC Symbol;Acc:HGNC:16018] | -0.7041 | 0.00063 |
| ENSG00000213780 | GTF2H4 | general transcription factor IIH subunit 4 [Source:HGNC Symbol;Acc:HGNC:4658] | 0.5583 | 0.00064 |
| ENSG00000231312 | MAP4K3-DT | MAP4K3 divergent transcript [Source:HGNC Symbol;Acc:HGNC:54056] | -0.6151 | 0.00064 |
| ENSG00000250565 | ATP6V1E2 | ATPase H+ transporting V1 subunit E2 [Source:HGNC Symbol;Acc:HGNC:18125] | -0.8896 | 0.00064 |
| ENSG00000110042 | DTX4 | deltex E3 ubiquitin ligase 4 [Source:HGNC Symbol;Acc:HGNC:29151] | -1.4274 | 0.00065 |
| ENSG00000114812 | VIPR1 | vasoactive intestinal peptide receptor 1 [Source:HGNC Symbol;Acc:HGNC:12694] | -0.8161 | 0.00065 |
| ENSG00000272688 | NA | novel transcript | 1.3001 | 0.00065 |
| ENSG00000203907 | OOEP | oocyte expressed protein [Source:HGNC Symbol;Acc:HGNC:21382] | -0.6980 | 0.00066 |
| ENSG00000198225 | FKBP1C | FKBP prolyl isomerase 1C [Source:HGNC Symbol;Acc:HGNC:21376] | 0.6300 | 0.00066 |
| ENSG00000279965 | NA | TEC | 2.0547 | 0.00066 |
| ENSG00000053918 | KCNQ1 | potassium voltage-gated channel subfamily Q member 1 [Source:HGNC Symbol;Acc:HGNC:6294] | 1.2088 | 0.00066 |
| ENSG00000187650 | VMAC | vimentin type intermediate filament associated coiled-coil protein [Source:HGNC Symbol;Acc:HGNC:33803] | -0.5679 | 0.00067 |
| ENSG00000173320 | STOX2 | storkhead box 2 [Source:HGNC Symbol;Acc:HGNC:25450] | 0.8480 | 0.00067 |
| ENSG00000196381 | ZNF781 | zinc finger protein 781 [Source:HGNC Symbol;Acc:HGNC:26745] | -0.8433 | 0.00067 |
| ENSG00000213963 | NA | novel transcript | -0.7122 | 0.00067 |
| ENSG00000176142 | TMEM39A | transmembrane protein 39A [Source:HGNC Symbol;Acc:HGNC:25600] | 0.5490 | 0.00068 |
| ENSG00000141540 | TTYH2 | tweety family member 2 [Source:HGNC Symbol;Acc:HGNC:13877] | -2.1712 | 0.00069 |
| ENSG00000286190 | NA | uncharacterized LOC728392 [Source:NCBI gene (formerly Entrezgene);Acc:728392] | 0.6855 | 0.00069 |
| ENSG00000136367 | ZFHX2 | zinc finger homeobox 2 [Source:HGNC Symbol;Acc:HGNC:20152] | 0.7416 | 0.00069 |
| ENSG00000160796 | NBEAL2 | neurobeachin like 2 [Source:HGNC Symbol;Acc:HGNC:31928] | -0.6386 | 0.00070 |
| ENSG00000187952 | HS6ST1P1 | heparan sulfate 6-O-sulfotransferase 1 pseudogene 1 [Source:HGNC Symbol;Acc:HGNC:31835] | -1.7411 | 0.00070 |
| ENSG00000256955 | NA | novel transcript | 3.7403 | 0.00070 |
| ENSG00000165474 | GJB2 | gap junction protein beta 2 [Source:HGNC Symbol;Acc:HGNC:4284] | 1.1059 | 0.00070 |
| ENSG00000169515 | CCDC8 | coiled-coil domain containing 8 [Source:HGNC Symbol;Acc:HGNC:25367] | -1.8160 | 0.00070 |
| ENSG00000225582 | NA | novel transcript | 3.4410 | 0.00070 |
| ENSG00000138380 | CARF | calcium responsive transcription factor [Source:HGNC Symbol;Acc:HGNC:14435] | -0.8827 | 0.00070 |
| ENSG00000081059 | TCF7 | transcription factor 7 [Source:HGNC Symbol;Acc:HGNC:11639] | 0.5381 | 0.00070 |
| ENSG00000169242 | EFNA1 | ephrin A1 [Source:HGNC Symbol;Acc:HGNC:3221] | -0.5945 | 0.00071 |
| ENSG00000234420 | ZNF37BP | zinc finger protein 37B, pseudogene [Source:HGNC Symbol;Acc:HGNC:13103] | -0.6117 | 0.00071 |
| ENSG00000267291 | NA | novel transcript | -1.3604 | 0.00071 |
| ENSG00000179588 | ZFPM1 | zinc finger protein, FOG family member 1 [Source:HGNC Symbol;Acc:HGNC:19762] | 0.9442 | 0.00071 |
| ENSG00000163840 | DTX3L | deltex E3 ubiquitin ligase 3L [Source:HGNC Symbol;Acc:HGNC:30323] | -0.6238 | 0.00072 |
| ENSG00000181274 | FRAT2 | FRAT regulator of WNT signaling pathway 2 [Source:HGNC Symbol;Acc:HGNC:16048] | -0.5979 | 0.00072 |
| ENSG00000266970 | NA | novel transcript | 2.7135 | 0.00072 |
| ENSG00000102384 | CENPI | centromere protein I [Source:HGNC Symbol;Acc:HGNC:3968] | -0.5157 | 0.00072 |
| ENSG00000229124 | VIM-AS1 | VIM antisense RNA 1 [Source:HGNC Symbol;Acc:HGNC:44879] | -1.1879 | 0.00072 |
| ENSG00000179299 | NSUN7 | NOP2/Sun RNA methyltransferase family member 7 [Source:HGNC Symbol;Acc:HGNC:25857] | -0.9017 | 0.00073 |
| ENSG00000270988 | NA | novel transcript | 2.5689 | 0.00073 |
| ENSG00000228589 | SPCS2P4 | signal peptidase complex subunit 2 pseudogene 4 [Source:HGNC Symbol;Acc:HGNC:45237] | 0.6466 | 0.00073 |

| | | | | |
|---|---|---|---|---|
| ENSG00000282851 | BISPR | BST2 interferon stimulated positive regulator [Source:HGNC Symbol;Acc:HGNC:51290] | -0.9867 | 0.00073 |
| ENSG00000260400 | NA | novel transcript, overlapping to TET1 | -0.8933 | 0.00073 |
| ENSG00000027869 | SH2D2A | SH2 domain containing 2A [Source:HGNC Symbol;Acc:HGNC:10821] | 3.8776 | 0.00074 |
| ENSG00000126337 | KRT36 | keratin 36 [Source:HGNC Symbol;Acc:HGNC:6454] | 2.8417 | 0.00074 |
| ENSG00000113209 | PCDHB5 | protocadherin beta 5 [Source:HGNC Symbol;Acc:HGNC:8690] | -0.5803 | 0.00075 |
| ENSG00000205002 | AARD | alanine and arginine rich domain containing protein [Source:HGNC Symbol;Acc:HGNC:33842] | -0.7045 | 0.00075 |
| ENSG00000224958 | PGM5-AS1 | PGM5 antisense RNA 1 [Source:HGNC Symbol;Acc:HGNC:44181] | -2.4364 | 0.00075 |
| ENSG00000183199 | HSP90AB3P | heat shock protein 90 alpha family class B member 3, pseudogene [Source:HGNC Symbol;Acc:HGNC:5259] | 1.0273 | 0.00075 |
| ENSG00000104321 | TRPA1 | transient receptor potential cation channel subfamily A member 1 [Source:HGNC Symbol;Acc:HGNC:497] | 4.3178 | 0.00076 |
| ENSG00000266256 | LINC00683 | long intergenic non-protein coding RNA 683 [Source:HGNC Symbol;Acc:HGNC:44467] | -1.4736 | 0.00076 |
| ENSG00000198670 | LPA | lipoprotein(a) [Source:HGNC Symbol;Acc:HGNC:6667] | -0.6998 | 0.00076 |
| ENSG00000159713 | TPPP3 | tubulin polymerization promoting protein family member 3 [Source:HGNC Symbol;Acc:HGNC:24162] | -0.8235 | 0.00076 |
| ENSG00000058335 | RASGRF1 | Ras protein specific guanine nucleotide releasing factor 1 [Source:HGNC Symbol;Acc:HGNC:9875] | 0.5199 | 0.00076 |
| ENSG00000171860 | C3AR1 | complement C3a receptor 1 [Source:HGNC Symbol;Acc:HGNC:1319] | 3.8984 | 0.00077 |
| ENSG00000274220 | NA | novel transcript | -0.8267 | 0.00077 |
| ENSG00000180616 | SSTR2 | somatostatin receptor 2 [Source:HGNC Symbol;Acc:HGNC:11331] | 0.7842 | 0.00077 |
| ENSG00000101977 | 2,00 MCF | MCF.2 cell line derived transforming sequence [Source:HGNC Symbol;Acc:HGNC:6940] | -0.7248 | 0.00078 |
| ENSG00000126091 | ST3GAL3 | ST3 beta-galactoside alpha-2,3-sialyltransferase 3 [Source:HGNC Symbol;Acc:HGNC:10866] | -0.5087 | 0.00079 |
| ENSG00000269952 | NA | novel transcript, sense intronic to CREM | 1.4731 | 0.00079 |
| ENSG00000160949 | TONSL | tonsoku like, DNA repair protein [Source:HGNC Symbol;Acc:HGNC:7801] | -0.7723 | 0.00079 |
| ENSG00000231528 | FAM225A | family with sequence similarity 225 member A [Source:HGNC Symbol;Acc:HGNC:27855] | -0.7018 | 0.00079 |
| ENSG00000187775 | DNAH17 | dynein axonemal heavy chain 17 [Source:HGNC Symbol;Acc:HGNC:2946] | 1.5395 | 0.00079 |
| ENSG00000234602 | MCIDAS | multiciliate differentiation and DNA synthesis associated cell cycle protein [Source:HGNC Symbol;Acc:HGNC:40050] | 1.0539 | 0.00080 |
| ENSG00000110436 | SLC1A2 | solute carrier family 1 member 2 [Source:HGNC Symbol;Acc:HGNC:10940] | 1.1000 | 0.00080 |
| ENSG00000173535 | TNFRSF10C | TNF receptor superfamily member 10c [Source:HGNC Symbol;Acc:HGNC:11906] | -1.0289 | 0.00080 |
| ENSG00000141956 | PRDM15 | PR/SET domain 15 [Source:HGNC Symbol;Acc:HGNC:13999] | 0.9082 | 0.00080 |
| ENSG00000112759 | SLC29A1 | solute carrier family 29 member 1 (Augustine blood group) [Source:HGNC Symbol;Acc:HGNC:11003] | -0.5392 | 0.00080 |
| ENSG00000286416 | NA | novel transcript | -1.2893 | 0.00080 |
| ENSG00000242732 | RTL5 | retrotransposon Gag like 5 [Source:HGNC Symbol;Acc:HGNC:29430] | -0.6695 | 0.00080 |
| ENSG00000280739 | EIF1B-AS1 | EIF1B antisense RNA 1 [Source:HGNC Symbol;Acc:HGNC:44555] | -0.8672 | 0.00080 |
| ENSG00000109674 | NEIL3 | nei like DNA glycosylase 3 [Source:HGNC Symbol;Acc:HGNC:24573] | -0.6816 | 0.00081 |
| ENSG00000287248 | NA | novel transcript | -0.6593 | 0.00081 |
| ENSG00000123600 | METTL8 | methyltransferase like 8 [Source:HGNC Symbol;Acc:HGNC:25856] | -0.6403 | 0.00082 |
| ENSG00000185189 | NRBP2 | nuclear receptor binding protein 2 [Source:HGNC Symbol;Acc:HGNC:19339] | -0.6538 | 0.00082 |
| ENSG00000161509 | GRIN2C | glutamate ionotropic receptor NMDA type subunit 2C [Source:HGNC Symbol;Acc:HGNC:4587] | 1.2384 | 0.00082 |
| ENSG00000130164 | LDLR | low density lipoprotein receptor [Source:HGNC Symbol;Acc:HGNC:6547] | 0.5537 | 0.00084 |
| ENSG00000279358 | NA | TEC | 3.2957 | 0.00084 |
| ENSG00000273374 | NA | novel transcript | -1.4489 | 0.00085 |
| ENSG00000235831 | BHLHE40-AS1 | BHLHE40 antisense RNA 1 [Source:HGNC Symbol;Acc:HGNC:44471] | 2.7778 | 0.00086 |
| ENSG00000157782 | CABP1 | calcium binding protein 1 [Source:HGNC Symbol;Acc:HGNC:1384] | 0.5281 | 0.00086 |
| ENSG00000288552 | NA | novel transcript | -0.9523 | 0.00086 |
| ENSG00000144959 | NCEH1 | neutral cholesterol ester hydrolase 1 [Source:HGNC Symbol;Acc:HGNC:29260] | 0.5597 | 0.00086 |
| ENSG00000249693 | THEGL | theg spermatid protein like [Source:HGNC Symbol;Acc:HGNC:43771] | -1.7804 | 0.00087 |

| | | | | |
|---|---|---|---|---|
| ENSG00000122786 | CALD1 | caldesmon 1 [Source:HGNC Symbol;Acc:HGNC:1441] | 0.7321 | 0.00087 |
| ENSG00000184922 | FMNL1 | formin like 1 [Source:HGNC Symbol;Acc:HGNC:1212] | 1.0323 | 0.00088 |
| ENSG00000103351 | CLUAP1 | clusterin associated protein 1 [Source:HGNC Symbol;Acc:HGNC:19009] | -0.6445 | 0.00088 |
| ENSG00000128578 | STRIP2 | striatin interacting protein 2 [Source:HGNC Symbol;Acc:HGNC:22209] | -0.8170 | 0.00090 |
| ENSG00000278974 | NA | tec | 0.6832 | 0.00091 |
| ENSG00000215424 | MCM3AP-AS1 | MCM3AP antisense RNA 1 [Source:HGNC Symbol;Acc:HGNC:16417] | -0.6470 | 0.00091 |
| ENSG00000136883 | KIF12 | kinesin family member 12 [Source:HGNC Symbol;Acc:HGNC:21495] | -1.4598 | 0.00091 |
| ENSG00000187097 | ENTPD5 | ectonucleoside triphosphate diphosphohydrolase 5 (inactive) [Source:HGNC Symbol;Acc:HGNC:3367] | -0.7362 | 0.00092 |
| ENSG00000120693 | SMAD9 | SMAD family member 9 [Source:HGNC Symbol;Acc:HGNC:6774] | 0.6499 | 0.00093 |
| ENSG00000197696 | NMB | neuromedin B [Source:HGNC Symbol;Acc:HGNC:7842] | 0.8727 | 0.00093 |
| ENSG00000250571 | GLI4 | GLI family zinc finger 4 [Source:HGNC Symbol;Acc:HGNC:4320] | -0.6527 | 0.00093 |
| ENSG00000258311 | NA | novel protein | -0.8111 | 0.00093 |
| ENSG00000126246 | IGFLR1 | IGF like family receptor 1 [Source:HGNC Symbol;Acc:HGNC:23620] | -0.9555 | 0.00093 |
| ENSG00000253882 | NA | family with sequence similarity 115, member C (FAM115C) pseudogene | -0.9699 | 0.00093 |
| ENSG00000188825 | LINC00910 | long intergenic non-protein coding RNA 910 [Source:HGNC Symbol;Acc:HGNC:44361] | -1.2897 | 0.00094 |
| ENSG00000102786 | INTS6 | integrator complex subunit 6 [Source:HGNC Symbol;Acc:HGNC:14879] | 0.5891 | 0.00095 |
| ENSG00000247809 | NR2F2-AS1 | NR2F2 antisense RNA 1 [Source:HGNC Symbol;Acc:HGNC:44222] | -2.2265 | 0.00095 |
| ENSG00000249087 | ZNF436-AS1 | ZNF436 antisense RNA 1 [Source:HGNC Symbol;Acc:HGNC:25122] | -0.5740 | 0.00096 |
| ENSG00000162702 | ZNF281 | zinc finger protein 281 [Source:HGNC Symbol;Acc:HGNC:13075] | 0.5052 | 0.00096 |
| ENSG00000248837 | NA | novel transcript | 1.0280 | 0.00096 |
| ENSG00000230499 | NA | novel transcript | 2.2567 | 0.00097 |
| ENSG00000105011 | ASF1B | anti-silencing function 1B histone chaperone [Source:HGNC Symbol;Acc:HGNC:20996] | -0.5543 | 0.00098 |
| ENSG00000234842 | MTCO2P16 | MT-CO2 pseudogene 16 [Source:HGNC Symbol;Acc:HGNC:52032] | -1.9264 | 0.00098 |
| ENSG00000130584 | ZBTB46 | zinc finger and BTB domain containing 46 [Source:HGNC Symbol;Acc:HGNC:16094] | -0.7449 | 0.00099 |
| ENSG00000158716 | DUSP23 | dual specificity phosphatase 23 [Source:HGNC Symbol;Acc:HGNC:21480] | 0.5938 | 0.00099 |
| ENSG00000144583 | MARCHF4 | membrane associated ring-CH-type finger 4 [Source:HGNC Symbol;Acc:HGNC:29269] | 0.8396 | 0.00100 |
| ENSG00000118965 | WDR35 | WD repeat domain 35 [Source:HGNC Symbol;Acc:HGNC:29250] | 0.6587 | 0.00100 |
| ENSG00000154889 | MPPE1 | metallophosphoesterase 1 [Source:HGNC Symbol;Acc:HGNC:15988] | -0.5409 | 0.00102 |
| ENSG00000264735 | NA | novel transcript | -2.4142 | 0.00102 |
| ENSG00000236810 | ELOA-AS1 | ELOA antisense RNA 1 [Source:HGNC Symbol;Acc:HGNC:50582] | -0.7938 | 0.00102 |
| ENSG00000105559 | PLEKHA4 | pleckstrin homology domain containing A4 [Source:HGNC Symbol;Acc:HGNC:14339] | 0.6135 | 0.00104 |
| ENSG00000287031 | NA | novel transcript | 2.3882 | 0.00104 |
| ENSG00000241684 | ADAMTS9-AS2 | ADAMTS9 antisense RNA 2 [Source:HGNC Symbol;Acc:HGNC:42435] | 1.2918 | 0.00104 |
| ENSG00000235706 | DICER1-AS1 | DICER1 antisense RNA 1 [Source:HGNC Symbol;Acc:HGNC:43017] | -0.5817 | 0.00105 |
| ENSG00000245750 | DRAIC | downregulated RNA in cancer, inhibitor of cell invasion and migration [Source:HGNC Symbol;Acc:HGNC:27082] | -1.0292 | 0.00105 |
| ENSG00000128165 | ADM2 | adrenomedullin 2 [Source:HGNC Symbol;Acc:HGNC:28898] | 0.8937 | 0.00105 |
| ENSG00000251026 | LINC02163 | long intergenic non-protein coding RNA 2163 [Source:HGNC Symbol;Acc:HGNC:53024] | -0.7110 | 0.00105 |
| ENSG00000236045 | NA | novel transcript | -1.7866 | 0.00107 |
| ENSG00000273172 | LINC02091 | long intergenic non-protein coding RNA 2091 [Source:HGNC Symbol;Acc:HGNC:52942] | 1.2172 | 0.00107 |
| ENSG00000180458 | NA | novel transcript, antisense to ZNF793 | -0.7861 | 0.00108 |
| ENSG00000226964 | RHEBP2 | RHEB pseudogene 2 [Source:HGNC Symbol;Acc:HGNC:44991] | -3.6416 | 0.00108 |
| ENSG00000162971 | TYW5 | tRNA-yW synthesizing protein 5 [Source:HGNC Symbol;Acc:HGNC:26754] | 0.5007 | 0.00109 |
| ENSG00000259425 | NA | Golgin subfamily A member 6 pseudogene | -0.9201 | 0.00109 |

| | | | | |
|---|---|---|---|---|
| ENSG00000267952 | NA | novel transcript | -5.2898 | 0.00109 |
| ENSG00000130222 | GADD45G | growth arrest and DNA damage inducible gamma [Source:HGNC Symbol;Acc:HGNC:4097] | 0.5509 | 0.00109 |
| ENSG00000122547 | EEPD1 | endonuclease/exonuclease/phosphatase family domain containing 1 [Source:HGNC Symbol;Acc:HGNC:22223] | -0.7080 | 0.00109 |
| ENSG00000163666 | HESX1 | HESX homeobox 1 [Source:HGNC Symbol;Acc:HGNC:4877] | -0.9346 | 0.00111 |
| ENSG00000274333 | NA | uncharacterized LOC102724219 [Source:NCBI gene (formerly Entrezgene);Acc:102724219] | -1.0001 | 0.00111 |
| ENSG00000254706 | NA | novel protein | -1.6350 | 0.00111 |
| ENSG00000119782 | FKBP1B | FKBP prolyl isomerase 1B [Source:HGNC Symbol;Acc:HGNC:3712] | -0.7752 | 0.00112 |
| ENSG00000229676 | ZNF492 | zinc finger protein 492 [Source:HGNC Symbol;Acc:HGNC:23707] | -0.5667 | 0.00113 |
| ENSG00000237424 | FOXD2-AS1 | FOXD2 adjacent opposite strand RNA 1 [Source:HGNC Symbol;Acc:HGNC:44256] | -0.7529 | 0.00113 |
| ENSG00000167094 | TTC16 | tetratricopeptide repeat domain 16 [Source:HGNC Symbol;Acc:HGNC:26536] | -1.2565 | 0.00114 |
| ENSG00000157557 | ETS2 | ETS proto-oncogene 2, transcription factor [Source:HGNC Symbol;Acc:HGNC:3489] | 1.6680 | 0.00114 |
| ENSG00000113494 | PRLR | prolactin receptor [Source:HGNC Symbol;Acc:HGNC:9446] | 1.5962 | 0.00114 |
| ENSG00000196208 | GREB1 | growth regulating estrogen receptor binding 1 [Source:HGNC Symbol;Acc:HGNC:24885] | -1.1667 | 0.00115 |
| ENSG00000132669 | RIN2 | Ras and Rab interactor 2 [Source:HGNC Symbol;Acc:HGNC:18750] | -0.7083 | 0.00115 |
| ENSG00000189212 | DPY19L2P1 | DPY19L2 pseudogene 1 [Source:HGNC Symbol;Acc:HGNC:22305] | 0.5172 | 0.00116 |
| ENSG00000219891 | ZSCAN12P1 | zinc finger and SCAN domain containing 12 pseudogene 1 [Source:HGNC Symbol;Acc:HGNC:13850] | 0.8376 | 0.00116 |
| ENSG00000138678 | GPAT3 | glycerol-3-phosphate acyltransferase 3 [Source:HGNC Symbol;Acc:HGNC:28157] | 0.6213 | 0.00116 |
| ENSG00000085552 | IGSF9 | immunoglobulin superfamily member 9 [Source:HGNC Symbol;Acc:HGNC:18132] | -0.9054 | 0.00116 |
| ENSG00000134508 | CABLES1 | Cdk5 and Abl enzyme substrate 1 [Source:HGNC Symbol;Acc:HGNC:25097] | 0.8293 | 0.00117 |
| ENSG00000180011 | ZADH2 | zinc binding alcohol dehydrogenase domain containing 2 [Source:HGNC Symbol;Acc:HGNC:28697] | -0.6315 | 0.00117 |
| ENSG00000164171 | ITGA2 | integrin subunit alpha 2 [Source:HGNC Symbol;Acc:HGNC:6137] | 0.8010 | 0.00118 |
| ENSG00000148082 | SHC3 | SHC adaptor protein 3 [Source:HGNC Symbol;Acc:HGNC:18181] | 0.9479 | 0.00119 |
| ENSG00000276116 | FUT8-AS1 | FUT8 antisense RNA 1 [Source:HGNC Symbol;Acc:HGNC:44294] | 2.2770 | 0.00119 |
| ENSG00000165752 | STK32C | serine/threonine kinase 32C [Source:HGNC Symbol;Acc:HGNC:21332] | -0.5424 | 0.00119 |
| ENSG00000170917 | NUDT6 | nudix hydrolase 6 [Source:HGNC Symbol;Acc:HGNC:8053] | -0.6499 | 0.00119 |
| ENSG00000101463 | SYNDIG1 | synapse differentiation inducing 1 [Source:HGNC Symbol;Acc:HGNC:15885] | 1.3284 | 0.00120 |
| ENSG00000181778 | TMEM252 | transmembrane protein 252 [Source:HGNC Symbol;Acc:HGNC:28537] | -1.9564 | 0.00120 |
| ENSG00000147144 | CCDC120 | coiled-coil domain containing 120 [Source:HGNC Symbol;Acc:HGNC:28910] | -0.5906 | 0.00120 |
| ENSG00000161040 | FBXL13 | F-box and leucine rich repeat protein 13 [Source:HGNC Symbol;Acc:HGNC:21658] | 0.8132 | 0.00120 |
| ENSG00000175785 | PRIMA1 | proline rich membrane anchor 1 [Source:HGNC Symbol;Acc:HGNC:18319] | -0.7768 | 0.00121 |
| ENSG00000234927 | HMGN1P18 | high mobility group nucleosome binding domain 1 pseudogene 18 [Source:HGNC Symbol;Acc:HGNC:39362] | 2.5426 | 0.00122 |
| ENSG00000245468 | LINC02447 | long intergenic non-protein coding RNA 2447 [Source:HGNC Symbol;Acc:HGNC:53379] | -1.5813 | 0.00122 |
| ENSG00000250392 | LINC02502 | long intergenic non-protein coding RNA 2502 [Source:HGNC Symbol;Acc:HGNC:53491] | -2.4227 | 0.00123 |
| ENSG00000111850 | SMIM8 | small integral membrane protein 8 [Source:HGNC Symbol;Acc:HGNC:21401] | -0.6258 | 0.00124 |
| ENSG00000010361 | FUZ | fuzzy planar cell polarity protein [Source:HGNC Symbol;Acc:HGNC:26219] | -0.5639 | 0.00124 |
| ENSG00000115137 | DNAJC27 | DnaJ heat shock protein family (Hsp40) member C27 [Source:HGNC Symbol;Acc:HGNC:30290] | -0.6160 | 0.00124 |
| ENSG00000152932 | RAB3C | RAB3C, member RAS oncogene family [Source:HGNC Symbol;Acc:HGNC:30269] | 0.6056 | 0.00125 |
| ENSG00000028839 | TBPL1 | TATA-box binding protein like 1 [Source:HGNC Symbol;Acc:HGNC:11589] | 0.5018 | 0.00125 |
| ENSG00000160223 | ICOSLG | inducible T cell costimulator ligand [Source:HGNC Symbol;Acc:HGNC:17087] | 2.0994 | 0.00125 |
| ENSG00000188315 | C3orf62 | chromosome 3 open reading frame 62 [Source:HGNC Symbol;Acc:HGNC:24771] | 0.6084 | 0.00126 |
| ENSG00000186493 | C5orf38 | chromosome 5 open reading frame 38 [Source:HGNC Symbol;Acc:HGNC:24226] | -1.2903 | 0.00126 |
| ENSG00000175879 | HOXD8 | homeobox D8 [Source:HGNC Symbol;Acc:HGNC:5139] | -1.1704 | 0.00126 |
| ENSG00000148357 | HMCN2 | hemicentin 2 [Source:HGNC Symbol;Acc:HGNC:21293] | 2.9479 | 0.00127 |

| | | | | |
|---|---|---|---|---|
| ENSG00000104361 | NIPAL2 | NIPA like domain containing 2 [Source:HGNC Symbol;Acc:HGNC:25854] | 0.7688 | 0.00128 |
| ENSG00000166821 | PEX11A | peroxisomal biogenesis factor 11 alpha [Source:HGNC Symbol;Acc:HGNC:8852] | -1.1074 | 0.00128 |
| ENSG00000248508 | SRP14-AS1 | SRP14 antisense RNA1 (head to head) [Source:HGNC Symbol;Acc:HGNC:48619] | -0.7581 | 0.00128 |
| ENSG00000177108 | ZDHHC22 | zinc finger DHHC-type palmitoyltransferase 22 [Source:HGNC Symbol;Acc:HGNC:20106] | -1.0971 | 0.00128 |
| ENSG00000189410 | SH2D5 | SH2 domain containing 5 [Source:HGNC Symbol;Acc:HGNC:28819] | 1.0632 | 0.00128 |
| ENSG00000178662 | CSRNP3 | cysteine and serine rich nuclear protein 3 [Source:HGNC Symbol;Acc:HGNC:30729] | -0.6067 | 0.00129 |
| ENSG00000189056 | RELN | reelin [Source:HGNC Symbol;Acc:HGNC:9957] | 0.6151 | 0.00130 |
| ENSG00000008056 | SYN1 | synapsin I [Source:HGNC Symbol;Acc:HGNC:11494] | 1.1291 | 0.00131 |
| ENSG00000156970 | BUB1B | BUB1 mitotic checkpoint serine/threonine kinase B [Source:HGNC Symbol;Acc:HGNC:1149] | -0.5002 | 0.00131 |
| ENSG00000174948 | GPR149 | G protein-coupled receptor 149 [Source:HGNC Symbol;Acc:HGNC:23627] | 2.8032 | 0.00131 |
| ENSG00000198959 | TGM2 | transglutaminase 2 [Source:HGNC Symbol;Acc:HGNC:11778] | -0.5854 | 0.00131 |
| ENSG00000130513 | GDF15 | growth differentiation factor 15 [Source:HGNC Symbol;Acc:HGNC:30142] | 2.3784 | 0.00132 |
| ENSG00000173540 | GMPPB | GDP-mannose pyrophosphorylase B [Source:HGNC Symbol;Acc:HGNC:22932] | 0.5410 | 0.00132 |
| ENSG00000205084 | TMEM231 | transmembrane protein 231 [Source:HGNC Symbol;Acc:HGNC:37234] | -0.5902 | 0.00133 |
| ENSG00000121964 | GTDC1 | glycosyltransferase like domain containing 1 [Source:HGNC Symbol;Acc:HGNC:20887] | -0.6026 | 0.00134 |
| ENSG00000134343 | ANO3 | anoctamin 3 [Source:HGNC Symbol;Acc:HGNC:14004] | 0.6100 | 0.00134 |
| ENSG00000181433 | SAGE1 | sarcoma antigen 1 [Source:HGNC Symbol;Acc:HGNC:30369] | -0.8649 | 0.00135 |
| ENSG00000163191 | S100A11 | S100 calcium binding protein A11 [Source:HGNC Symbol;Acc:HGNC:10488] | 0.6470 | 0.00135 |
| ENSG00000110328 | GALNT18 | polypeptide N-acetylgalactosaminyltransferase 18 [Source:HGNC Symbol;Acc:HGNC:30488] | -1.1206 | 0.00135 |
| ENSG00000179348 | GATA2 | GATA binding protein 2 [Source:HGNC Symbol;Acc:HGNC:4171] | -0.9601 | 0.00136 |
| ENSG00000164690 | SHH | sonic hedgehog signaling molecule [Source:HGNC Symbol;Acc:HGNC:10848] | 0.6332 | 0.00136 |
| ENSG00000173930 | SLCO4C1 | solute carrier organic anion transporter family member 4C1 [Source:HGNC Symbol;Acc:HGNC:23612] | -1.4130 | 0.00136 |
| ENSG00000283228 | NA | novel transcript | 0.9388 | 0.00138 |
| ENSG00000272690 | LINC02018 | long intergenic non-protein coding RNA 2018 [Source:HGNC Symbol;Acc:HGNC:52853] | -1.5361 | 0.00138 |
| ENSG00000154760 | SLFN13 | schlafen family member 13 [Source:HGNC Symbol;Acc:HGNC:26481] | 1.0761 | 0.00140 |
| ENSG00000178409 | BEND3 | BEN domain containing 3 [Source:HGNC Symbol;Acc:HGNC:23040] | 0.5878 | 0.00142 |
| ENSG00000109089 | CDR2L | cerebellar degeneration related protein 2 like [Source:HGNC Symbol;Acc:HGNC:29999] | -0.6599 | 0.00142 |
| ENSG00000158805 | ZNF276 | zinc finger protein 276 [Source:HGNC Symbol;Acc:HGNC:23330] | 0.6362 | 0.00142 |
| ENSG00000162772 | ATF3 | activating transcription factor 3 [Source:HGNC Symbol;Acc:HGNC:785] | 1.3228 | 0.00142 |
| ENSG00000278266 | NA | novel transcript | -0.5914 | 0.00144 |
| ENSG00000168309 | FAM107A | family with sequence similarity 107 member A [Source:HGNC Symbol;Acc:HGNC:30827] | -1.7343 | 0.00145 |
| ENSG00000122862 | SRGN | serglycin [Source:HGNC Symbol;Acc:HGNC:9361] | 0.8147 | 0.00145 |
| ENSG00000156687 | UNC5D | unc-5 netrin receptor D [Source:HGNC Symbol;Acc:HGNC:18634] | 0.6383 | 0.00146 |
| ENSG00000214652 | ZNF727 | zinc finger protein 727 [Source:HGNC Symbol;Acc:HGNC:22785] | -0.9515 | 0.00148 |
| ENSG00000204758 | NA | novel transcript, antisense to RPL26L1 | 1.1548 | 0.00149 |
| ENSG00000169607 | CKAP2L | cytoskeleton associated protein 2 like [Source:HGNC Symbol;Acc:HGNC:26877] | -0.5071 | 0.00150 |
| ENSG00000165695 | AK8 | adenylate kinase 8 [Source:HGNC Symbol;Acc:HGNC:26526] | -0.8584 | 0.00151 |
| ENSG00000167523 | SPATA33 | spermatogenesis associated 33 [Source:HGNC Symbol;Acc:HGNC:26463] | -0.5757 | 0.00152 |
| ENSG00000274422 | NA | novel transcript | -0.8054 | 0.00152 |
| ENSG00000162997 | PRORSD1P | prolyl-tRNA synthetase associated domain containing 1, pseudogene [Source:HGNC Symbol;Acc:HGNC:34379] | -1.0742 | 0.00152 |
| ENSG00000272221 | NA | novel transcript | -1.1914 | 0.00153 |
| ENSG00000138794 | CASP6 | caspase 6 [Source:HGNC Symbol;Acc:HGNC:1507] | -0.7781 | 0.00153 |
| ENSG00000099875 | MKNK2 | MAPK interacting serine/threonine kinase 2 [Source:HGNC Symbol;Acc:HGNC:7111] | -0.5028 | 0.00153 |

| | | | | |
|---|---|---|---|---|
| ENSG00000272121 | NA | novel transcript | -0.6488 | 0.00153 |
| ENSG00000280119 | NA | TEC | -1.7256 | 0.00154 |
| ENSG00000161558 | TMEM143 | transmembrane protein 143 [Source:HGNC Symbol;Acc:HGNC:25603] | -0.8596 | 0.00155 |
| ENSG00000156711 | MAPK13 | mitogen-activated protein kinase 13 [Source:HGNC Symbol;Acc:HGNC:6875] | -0.7613 | 0.00155 |
| ENSG00000171466 | ZNF562 | zinc finger protein 562 [Source:HGNC Symbol;Acc:HGNC:25950] | 0.5411 | 0.00155 |
| ENSG00000286638 | NA | novel transcript | -0.6753 | 0.00155 |
| ENSG00000165810 | BTNL9 | butyrophilin like 9 [Source:HGNC Symbol;Acc:HGNC:24176] | -0.8637 | 0.00156 |
| ENSG00000174130 | TLR6 | toll like receptor 6 [Source:HGNC Symbol;Acc:HGNC:16711] | 0.6651 | 0.00156 |
| ENSG00000183018 | SPNS2 | sphingolipid transporter 2 [Source:HGNC Symbol;Acc:HGNC:26992] | 1.9422 | 0.00156 |
| ENSG00000243083 | LINC00870 | long intergenic non-protein coding RNA 870 [Source:HGNC Symbol;Acc:HGNC:27319] | -2.5655 | 0.00156 |
| ENSG00000176641 | RNF152 | ring finger protein 152 [Source:HGNC Symbol;Acc:HGNC:26811] | -0.6752 | 0.00158 |
| ENSG00000183569 | SERHL2 | serine hydrolase like 2 [Source:HGNC Symbol;Acc:HGNC:29446] | 1.0272 | 0.00159 |
| ENSG00000277619 | NA | novel transcript, antisense to PITX1 | -2.0737 | 0.00159 |
| ENSG00000187372 | PCDHB13 | protocadherin beta 13 [Source:HGNC Symbol;Acc:HGNC:8684] | -0.5517 | 0.00159 |
| ENSG00000260653 | NA | novel transcript | -0.9251 | 0.00160 |
| ENSG00000248898 | NA | novel transcript | 1.3282 | 0.00161 |
| ENSG00000287067 | NA | zinc finger BED-type containing 9 (ZBED9) pseudogene | 1.0292 | 0.00161 |
| ENSG00000172927 | MYEOV | myeloma overexpressed [Source:HGNC Symbol;Acc:HGNC:7563] | -2.5935 | 0.00162 |
| ENSG00000104936 | DMPK | DM1 protein kinase [Source:HGNC Symbol;Acc:HGNC:2933] | 0.5889 | 0.00162 |
| ENSG00000186481 | ANKRD20A5P | ankyrin repeat domain 20 family member A5, pseudogene [Source:HGNC Symbol;Acc:HGNC:33833] | -1.0513 | 0.00163 |
| ENSG00000169862 | CTNND2 | catenin delta 2 [Source:HGNC Symbol;Acc:HGNC:2516] | 2.0878 | 0.00164 |
| ENSG00000149403 | GRIK4 | glutamate ionotropic receptor kainate type subunit 4 [Source:HGNC Symbol;Acc:HGNC:4582] | -1.2666 | 0.00165 |
| ENSG00000280099 | NA | TEC | -1.4061 | 0.00165 |
| ENSG00000213397 | HAUS7 | HAUS augmin like complex subunit 7 [Source:HGNC Symbol;Acc:HGNC:32979] | -0.5119 | 0.00165 |
| ENSG00000124444 | ZNF576 | zinc finger protein 576 [Source:HGNC Symbol;Acc:HGNC:28357] | 0.5049 | 0.00166 |
| ENSG00000128016 | ZFP36 | ZFP36 ring finger protein [Source:HGNC Symbol;Acc:HGNC:12862] | 0.6256 | 0.00166 |
| ENSG00000046647 | GEMIN8 | gem nuclear organelle associated protein 8 [Source:HGNC Symbol;Acc:HGNC:26044] | -0.7391 | 0.00166 |
| ENSG00000137267 | TUBB2A | tubulin beta 2A class IIa [Source:HGNC Symbol;Acc:HGNC:12412] | 0.5617 | 0.00167 |
| ENSG00000287844 | NA | novel transcript | -1.9740 | 0.00167 |
| ENSG00000136535 | TBR1 | T-box brain transcription factor 1 [Source:HGNC Symbol;Acc:HGNC:11590] | 1.6148 | 0.00167 |
| ENSG00000188042 | ARL4C | ADP ribosylation factor like GTPase 4C [Source:HGNC Symbol;Acc:HGNC:698] | -0.9203 | 0.00167 |
| ENSG00000198870 | STKLD1 | serine/threonine kinase like domain containing 1 [Source:HGNC Symbol;Acc:HGNC:28669] | -0.9961 | 0.00168 |
| ENSG00000276390 | NA | novel transcript, sense intronic to RPAP3 | 1.2807 | 0.00168 |
| ENSG00000130827 | PLXNA3 | plexin A3 [Source:HGNC Symbol;Acc:HGNC:9101] | 0.5958 | 0.00168 |
| ENSG00000196230 | TUBB | tubulin beta class I [Source:HGNC Symbol;Acc:HGNC:20778] | -0.5899 | 0.00169 |
| ENSG00000158850 | B4GALT3 | beta-1,4-galactosyltransferase 3 [Source:HGNC Symbol;Acc:HGNC:926] | 0.5210 | 0.00170 |
| ENSG00000112782 | CLIC5 | chloride intracellular channel 5 [Source:HGNC Symbol;Acc:HGNC:13517] | 1.6002 | 0.00170 |
| ENSG00000132570 | PCBD2 | pterin-4 alpha-carbinolamine dehydratase 2 [Source:HGNC Symbol;Acc:HGNC:24474] | -0.5961 | 0.00171 |
| ENSG00000243696 | NA | novel MUSTN1-ITIH4 readthrough | -0.8565 | 0.00171 |
| ENSG00000287409 | NA | novel transcript | 0.9974 | 0.00171 |
| ENSG00000284948 | NA | novel transcript | -0.5245 | 0.00171 |
| ENSG00000129667 | RHBDF2 | rhomboid 5 homolog 2 [Source:HGNC Symbol;Acc:HGNC:20788] | -0.7778 | 0.00172 |
| ENSG00000168916 | ZNF608 | zinc finger protein 608 [Source:HGNC Symbol;Acc:HGNC:29238] | 0.6669 | 0.00173 |

| | | | | |
|---|---|---|---|---|
| ENSG00000130813 | SHFL | shiftless antiviral inhibitor of ribosomal frameshifting [Source:HGNC Symbol;Acc:HGNC:25649] | -1.0006 | 0.00173 |
| ENSG00000151623 | NR3C2 | nuclear receptor subfamily 3 group C member 2 [Source:HGNC Symbol;Acc:HGNC:7979] | -1.5800 | 0.00174 |
| ENSG00000184898 | RBM43 | RNA binding motif protein 43 [Source:HGNC Symbol;Acc:HGNC:24790] | -0.6072 | 0.00175 |
| ENSG00000212978 | NA | novel transcript | -0.6452 | 0.00176 |
| ENSG00000174137 | FAM53A | family with sequence similarity 53 member A [Source:HGNC Symbol;Acc:HGNC:31860] | -1.8165 | 0.00176 |
| ENSG00000005187 | ACSM3 | acyl-CoA synthetase medium chain family member 3 [Source:HGNC Symbol;Acc:HGNC:10522] | -0.8107 | 0.00176 |
| ENSG00000005156 | LIG3 | DNA ligase 3 [Source:HGNC Symbol;Acc:HGNC:6600] | 0.5451 | 0.00177 |
| ENSG00000142227 | EMP3 | epithelial membrane protein 3 [Source:HGNC Symbol;Acc:HGNC:3335] | -0.8059 | 0.00180 |
| ENSG00000258964 | NA | novel transcript | 1.4218 | 0.00181 |
| ENSG00000285533 | RELA-DT | RELA divergent transcript [Source:HGNC Symbol;Acc:HGNC:54185] | -1.0748 | 0.00181 |
| ENSG00000115266 | APC2 | APC regulator of WNT signaling pathway 2 [Source:HGNC Symbol;Acc:HGNC:24036] | -0.6051 | 0.00182 |
| ENSG00000272077 | NA | novel transcript | 0.7291 | 0.00183 |
| ENSG00000140470 | ADAMTS17 | ADAM metallopeptidase with thrombospondin type 1 motif 17 [Source:HGNC Symbol;Acc:HGNC:17109] | 3.2354 | 0.00183 |
| ENSG00000163703 | CRELD1 | cysteine rich with EGF like domains 1 [Source:HGNC Symbol;Acc:HGNC:14630] | 0.6302 | 0.00183 |
| ENSG00000074803 | SLC12A1 | solute carrier family 12 member 1 [Source:HGNC Symbol;Acc:HGNC:10910] | -1.0257 | 0.00184 |
| ENSG00000117600 | PLPPR4 | phospholipid phosphatase related 4 [Source:HGNC Symbol;Acc:HGNC:23496] | -3.0716 | 0.00185 |
| ENSG00000254838 | GVINP1 | GTPase, very large interferon inducible pseudogene 1 [Source:HGNC Symbol;Acc:HGNC:25813] | 2.1619 | 0.00185 |
| ENSG00000214193 | SH3D21 | SH3 domain containing 21 [Source:HGNC Symbol;Acc:HGNC:26236] | 0.8539 | 0.00185 |
| ENSG00000150676 | CCDC83 | coiled-coil domain containing 83 [Source:HGNC Symbol;Acc:HGNC:28535] | 3.4785 | 0.00187 |
| ENSG00000271579 | NA | novel transcript | 0.7055 | 0.00187 |
| ENSG00000146411 | SLC2A12 | solute carrier family 2 member 12 [Source:HGNC Symbol;Acc:HGNC:18067] | -0.7716 | 0.00188 |
| ENSG00000196275 | GTF2IRD2 | GTF2I repeat domain containing 2 [Source:HGNC Symbol;Acc:HGNC:30775] | -0.7093 | 0.00188 |
| ENSG00000185052 | SLC24A3 | solute carrier family 24 member 3 [Source:HGNC Symbol;Acc:HGNC:10977] | 1.5737 | 0.00188 |
| ENSG00000149654 | CDH22 | cadherin 22 [Source:HGNC Symbol;Acc:HGNC:13251] | -3.9994 | 0.00189 |
| ENSG00000272669 | NA | novel transcript, antisense to SUN2 | 0.9930 | 0.00189 |
| ENSG00000285447 | NA | zinc finger protein 883 [Source:NCBI gene (formerly Entrezgene);Acc:169834] | 20.7723 | 0.00190 |
| ENSG00000287408 | NA | novel transcript, antisense to TMED3 | 1.8977 | 0.00190 |
| ENSG00000134569 | LRP4 | LDL receptor related protein 4 [Source:HGNC Symbol;Acc:HGNC:6696] | -1.1344 | 0.00190 |
| ENSG00000205485 | NA | pseudogene similar to part of postmeiotic segregation increased 2-like 3 PMS2L3 | 0.7628 | 0.00191 |
| ENSG00000187609 | EXD3 | exonuclease 3'-5' domain containing 3 [Source:HGNC Symbol;Acc:HGNC:26023] | -0.5407 | 0.00191 |
| ENSG00000197774 | EME2 | essential meiotic structure-specific endonuclease subunit 2 [Source:HGNC Symbol;Acc:HGNC:27289] | 0.6089 | 0.00192 |
| ENSG00000196972 | SMIM10L2B | small integral membrane protein 10 like 2B [Source:HGNC Symbol;Acc:HGNC:34500] | -0.9581 | 0.00192 |
| ENSG00000187416 | LHFPL3 | LHFPL tetraspan subfamily member 3 [Source:HGNC Symbol;Acc:HGNC:6589] | -1.2048 | 0.00192 |
| ENSG00000132003 | ZSWIM4 | zinc finger SWIM-type containing 4 [Source:HGNC Symbol;Acc:HGNC:25704] | 0.6681 | 0.00194 |
| ENSG00000108932 | SLC16A6 | solute carrier family 16 member 6 [Source:HGNC Symbol;Acc:HGNC:10927] | 1.1725 | 0.00194 |
| ENSG00000163646 | CLRN1 | clarin 1 [Source:HGNC Symbol;Acc:HGNC:12605] | 0.5993 | 0.00195 |
| ENSG00000280200 | NA | novel transcript | 3.4101 | 0.00197 |
| ENSG00000248971 | KRT8P46 | keratin 8 pseudogene 46 [Source:HGNC Symbol;Acc:HGNC:39880] | 1.2211 | 0.00197 |
| ENSG00000121716 | PILRB | paired immunoglobin like type 2 receptor beta [Source:HGNC Symbol;Acc:HGNC:18297] | 0.7533 | 0.00199 |
| ENSG00000170464 | DNAJC18 | DnaJ heat shock protein family (Hsp40) member C18 [Source:HGNC Symbol;Acc:HGNC:28429] | 0.6005 | 0.00200 |
| ENSG00000196812 | ZSCAN16 | zinc finger and SCAN domain containing 16 [Source:HGNC Symbol;Acc:HGNC:20813] | 0.8058 | 0.00200 |
| ENSG00000134698 | AGO4 | argonaute RISC component 4 [Source:HGNC Symbol;Acc:HGNC:18424] | -0.9503 | 0.00201 |
| ENSG00000260220 | CCDC187 | coiled-coil domain containing 187 [Source:HGNC Symbol;Acc:HGNC:30942] | 3.3151 | 0.00201 |

| | | | | |
|---|---|---|---|---|
| ENSG00000230701 | FBXW4P1 | F-box and WD repeat domain containing 4 pseudogene 1 [Source:HGNC Symbol;Acc:HGNC:13609] | 0.6800 | 0.00201 |
| ENSG00000142252 | GEMIN7 | gem nuclear organelle associated protein 7 [Source:HGNC Symbol;Acc:HGNC:20045] | 0.6367 | 0.00202 |
| ENSG00000173114 | LRRN3 | leucine rich repeat neuronal 3 [Source:HGNC Symbol;Acc:HGNC:17200] | -2.0849 | 0.00203 |
| ENSG00000242779 | ZNF702P | zinc finger protein 702, pseudogene [Source:HGNC Symbol;Acc:HGNC:25775] | -1.1975 | 0.00203 |
| ENSG00000135454 | B4GALNT1 | beta-1,4-N-acetyl-galactosaminyltransferase 1 [Source:HGNC Symbol;Acc:HGNC:4117] | 0.5850 | 0.00205 |
| ENSG00000278768 | BACE1-AS | BACE1 antisense RNA [Source:HGNC Symbol;Acc:HGNC:37125] | -0.5558 | 0.00205 |
| ENSG00000182631 | RXFP3 | relaxin family peptide receptor 3 [Source:HGNC Symbol;Acc:HGNC:24883] | 2.8920 | 0.00206 |
| ENSG00000227627 | NA | novel transcript | -1.4070 | 0.00206 |
| ENSG00000108106 | UBE2S | ubiquitin conjugating enzyme E2 S [Source:HGNC Symbol;Acc:HGNC:17895] | 0.5329 | 0.00207 |
| ENSG00000196358 | NTNG2 | netrin G2 [Source:HGNC Symbol;Acc:HGNC:14288] | -0.5297 | 0.00208 |
| ENSG00000264448 | MIR378D2HG | MIR378D2 host gene [Source:HGNC Symbol;Acc:HGNC:54166] | 3.1461 | 0.00209 |
| ENSG00000133466 | C1QTNF6 | C1q and TNF related 6 [Source:HGNC Symbol;Acc:HGNC:14343] | -0.7726 | 0.00210 |
| ENSG00000104967 | NOVA2 | NOVA alternative splicing regulator 2 [Source:HGNC Symbol;Acc:HGNC:7887] | -0.5269 | 0.00212 |
| ENSG00000251664 | PCDHA12 | protocadherin alpha 12 [Source:HGNC Symbol;Acc:HGNC:8666] | -0.7401 | 0.00213 |
| ENSG00000131398 | KCNC3 | potassium voltage-gated channel subfamily C member 3 [Source:HGNC Symbol;Acc:HGNC:6235] | 0.7938 | 0.00214 |
| ENSG00000254231 | NA | novel transcript | -1.4623 | 0.00214 |
| ENSG00000100592 | DAAM1 | dishevelled associated activator of morphogenesis 1 [Source:HGNC Symbol;Acc:HGNC:18142] | -0.6313 | 0.00215 |
| ENSG00000226413 | OR8T1P | olfactory receptor family 8 subfamily T member 1 pseudogene [Source:HGNC Symbol;Acc:HGNC:19630] | 2.5213 | 0.00216 |
| ENSG00000204429 | NA | potassium channel tetramerisation domain containing 1 (KCTD1) pseudogene | -1.2353 | 0.00217 |
| ENSG00000143382 | ADAMTSL4 | ADAMTS like 4 [Source:HGNC Symbol;Acc:HGNC:19706] | 1.6565 | 0.00219 |
| ENSG00000163485 | ADORA1 | adenosine A1 receptor [Source:HGNC Symbol;Acc:HGNC:262] | -0.7823 | 0.00219 |
| ENSG00000136014 | USP44 | ubiquitin specific peptidase 44 [Source:HGNC Symbol;Acc:HGNC:20064] | 1.3197 | 0.00220 |
| ENSG00000103150 | MLYCD | malonyl-CoA decarboxylase [Source:HGNC Symbol;Acc:HGNC:7150] | 0.6889 | 0.00220 |
| ENSG00000135924 | DNAJB2 | DnaJ heat shock protein family (Hsp40) member B2 [Source:HGNC Symbol;Acc:HGNC:5228] | 0.5002 | 0.00222 |
| ENSG00000030419 | IKZF2 | IKAROS family zinc finger 2 [Source:HGNC Symbol;Acc:HGNC:13177] | -0.8486 | 0.00222 |
| ENSG00000138375 | SMARCAL1 | SWI/SNF related, matrix associated, actin dependent regulator of chromatin, subfamily a like 1 [Source:HGNC Symbol;Acc:HGNC:11102] | -0.5882 | 0.00222 |
| ENSG00000146094 | DOK3 | docking protein 3 [Source:HGNC Symbol;Acc:HGNC:24583] | 0.6798 | 0.00223 |
| ENSG00000232000 | CLCN3P1 | chloride voltage-gated channel 3 pseudogene 1 [Source:HGNC Symbol;Acc:HGNC:49775] | -0.6667 | 0.00225 |
| ENSG00000286381 | NA | novel transcript | -0.9679 | 0.00225 |
| ENSG00000245498 | NA | novel transcript, antisense to C11orf61 | -0.6128 | 0.00227 |
| ENSG00000120328 | PCDHB12 | protocadherin beta 12 [Source:HGNC Symbol;Acc:HGNC:8683] | -0.7574 | 0.00227 |
| ENSG00000148803 | FUOM | fucose mutarotase [Source:HGNC Symbol;Acc:HGNC:24733] | 0.7233 | 0.00227 |
| ENSG00000220008 | LINGO3 | leucine rich repeat and Ig domain containing 3 [Source:HGNC Symbol;Acc:HGNC:21206] | -4.6204 | 0.00227 |
| ENSG00000243955 | GSTA1 | glutathione S-transferase alpha 1 [Source:HGNC Symbol;Acc:HGNC:4626] | -0.5539 | 0.00227 |
| ENSG00000125730 | C3 | complement C3 [Source:HGNC Symbol;Acc:HGNC:1318] | 1.8421 | 0.00228 |
| ENSG00000243660 | ZNF487 | zinc finger protein 487 [Source:HGNC Symbol;Acc:HGNC:23488] | -1.3681 | 0.00229 |
| ENSG00000170854 | RIOX2 | ribosomal oxygenase 2 [Source:HGNC Symbol;Acc:HGNC:19441] | -0.5303 | 0.00229 |
| ENSG00000285555 | NA | novel transcript, antisense to CACNA1C | 2.0060 | 0.00230 |
| ENSG00000168152 | THAP9 | THAP domain containing 9 [Source:HGNC Symbol;Acc:HGNC:23192] | -0.5158 | 0.00231 |
| ENSG00000107872 | FBXL15 | F-box and leucine rich repeat protein 15 [Source:HGNC Symbol;Acc:HGNC:28155] | -0.5894 | 0.00232 |
| ENSG00000138336 | TET1 | tet methylcytosine dioxygenase 1 [Source:HGNC Symbol;Acc:HGNC:29484] | -0.5561 | 0.00232 |
| ENSG00000234460 | NA | cathepsin L (CTSL) pseudogene | -1.2945 | 0.00232 |

| | | | | |
|---|---|---|---|---|
| ENSG00000141736 | ERBB2 | erb-b2 receptor tyrosine kinase 2 [Source:HGNC Symbol;Acc:HGNC:3430] | -0.7326 | 0.00233 |
| ENSG00000159905 | ZNF221 | zinc finger protein 221 [Source:HGNC Symbol;Acc:HGNC:13014] | -0.8214 | 0.00236 |
| ENSG00000258839 | MC1R | melanocortin 1 receptor [Source:HGNC Symbol;Acc:HGNC:6929] | 0.5369 | 0.00236 |
| ENSG00000177839 | PCDHB9 | protocadherin beta 9 [Source:HGNC Symbol;Acc:HGNC:8694] | -0.9602 | 0.00236 |
| ENSG00000100314 | CABP7 | calcium binding protein 7 [Source:HGNC Symbol;Acc:HGNC:20834] | -0.8747 | 0.00237 |
| ENSG00000197608 | ZNF841 | zinc finger protein 841 [Source:HGNC Symbol;Acc:HGNC:27611] | 0.6008 | 0.00238 |
| ENSG00000272732 | NA | novel transcript | -2.0354 | 0.00239 |
| ENSG00000173083 | HPSE | heparanase [Source:HGNC Symbol;Acc:HGNC:5164] | 0.5825 | 0.00240 |
| ENSG00000169629 | RGPD8 | RANBP2 like and GRIP domain containing 8 [Source:HGNC Symbol;Acc:HGNC:9849] | 0.8462 | 0.00240 |
| ENSG00000278817 | NA | protein DGCR6 [Source:NCBI gene (formerly Entrezgene);Acc:102724770] | -0.9937 | 0.00240 |
| ENSG00000204516 | MICB | MHC class I polypeptide-related sequence B [Source:HGNC Symbol;Acc:HGNC:7091] | 0.6288 | 0.00241 |
| ENSG00000123213 | NLN | neurolysin [Source:HGNC Symbol;Acc:HGNC:16058] | -0.6065 | 0.00242 |
| ENSG00000226629 | LINC00974 | long intergenic non-protein coding RNA 974 [Source:HGNC Symbol;Acc:HGNC:27105] | 3.2676 | 0.00242 |
| ENSG00000166262 | FAM227B | family with sequence similarity 227 member B [Source:HGNC Symbol;Acc:HGNC:26543] | -0.5865 | 0.00243 |
| ENSG00000127578 | WFIKKN1 | WAP, follistatin/kazal, immunoglobulin, kunitz and netrin domain containing 1 [Source:HGNC Symbol;Acc:HGNC:30912] | -0.7337 | 0.00243 |
| ENSG00000106714 | CNTNAP3 | contactin associated protein family member 3 [Source:HGNC Symbol;Acc:HGNC:13834] | -1.5818 | 0.00243 |
| ENSG00000203697 | CAPN8 | calpain 8 [Source:HGNC Symbol;Acc:HGNC:1485] | 3.2673 | 0.00243 |
| ENSG00000253554 | LINC01414 | long intergenic non-protein coding RNA 1414 [Source:HGNC Symbol;Acc:HGNC:50707] | -1.7096 | 0.00247 |
| ENSG00000230077 | MTAPP2 | methylthioadenosine phosphorylase pseudogene 2 [Source:HGNC Symbol;Acc:HGNC:42443] | 3.4041 | 0.00247 |
| ENSG00000087076 | HSD17B14 | hydroxysteroid 17-beta dehydrogenase 14 [Source:HGNC Symbol;Acc:HGNC:23238] | -0.6643 | 0.00248 |
| ENSG00000257283 | NA | novel transcript, antisense to RRP12 | 2.4796 | 0.00248 |
| ENSG00000140332 | TLE3 | TLE family member 3, transcriptional corepressor [Source:HGNC Symbol;Acc:HGNC:11839] | 1.2793 | 0.00248 |
| ENSG00000132329 | RAMP1 | receptor activity modifying protein 1 [Source:HGNC Symbol;Acc:HGNC:9843] | 2.3112 | 0.00248 |
| ENSG00000221883 | ARIH2OS | ARIH2 opposite strand lncRNA [Source:HGNC Symbol;Acc:HGNC:34425] | -0.7847 | 0.00249 |
| ENSG00000260528 | FAM157C | family with sequence similarity 157 member C [Source:HGNC Symbol;Acc:HGNC:34081] | -0.6305 | 0.00249 |
| ENSG00000171940 | ZNF217 | zinc finger protein 217 [Source:HGNC Symbol;Acc:HGNC:13009] | 0.5248 | 0.00249 |
| ENSG00000173612 | GPRC6A | G protein-coupled receptor class C group 6 member A [Source:HGNC Symbol;Acc:HGNC:18510] | 2.4557 | 0.00250 |
| ENSG00000132801 | ZSWIM3 | zinc finger SWIM-type containing 3 [Source:HGNC Symbol;Acc:HGNC:16157] | 0.6780 | 0.00251 |
| ENSG00000089692 | LAG3 | lymphocyte activating 3 [Source:HGNC Symbol;Acc:HGNC:6476] | 0.8618 | 0.00253 |
| ENSG00000253301 | LINC01606 | long intergenic non-protein coding RNA 1606 [Source:HGNC Symbol;Acc:HGNC:51656] | -0.5600 | 0.00253 |
| ENSG00000135363 | LMO2 | LIM domain only 2 [Source:HGNC Symbol;Acc:HGNC:6642] | -0.6957 | 0.00255 |
| ENSG00000107485 | GATA3 | GATA binding protein 3 [Source:HGNC Symbol;Acc:HGNC:4172] | -0.7635 | 0.00256 |
| ENSG00000250033 | SLC7A11-AS1 | SLC7A11 antisense RNA 1 [Source:HGNC Symbol;Acc:HGNC:44064] | 2.2059 | 0.00256 |
| ENSG00000135749 | PCNX2 | pecanex 2 [Source:HGNC Symbol;Acc:HGNC:8736] | 0.5338 | 0.00256 |
| ENSG00000163947 | ARHGEF3 | Rho guanine nucleotide exchange factor 3 [Source:HGNC Symbol;Acc:HGNC:683] | -0.5632 | 0.00257 |
| ENSG00000228109 | MELTF-AS1 | MELTF antisense RNA 1 [Source:HGNC Symbol;Acc:HGNC:40373] | -1.5315 | 0.00257 |
| ENSG00000230736 | NA | novel transcript | 1.5940 | 0.00257 |
| ENSG00000272047 | GTF2H5 | general transcription factor IIH subunit 5 [Source:HGNC Symbol;Acc:HGNC:21157] | -0.5858 | 0.00257 |
| ENSG00000154059 | IMPACT | impact RWD domain protein [Source:HGNC Symbol;Acc:HGNC:20387] | -0.5073 | 0.00258 |
| ENSG00000186446 | ZNF501 | zinc finger protein 501 [Source:HGNC Symbol;Acc:HGNC:23717] | -0.6001 | 0.00261 |
| ENSG00000261490 | NA | novel transcript, overlapping WDR1 | 1.0523 | 0.00261 |
| ENSG00000221829 | FANCG | FA complementation group G [Source:HGNC Symbol;Acc:HGNC:3588] | -0.5841 | 0.00263 |

| | | | | |
|---|---|---|---|---|
| ENSG00000225950 | NTF4 | neurotrophin 4 [Source:HGNC Symbol;Acc:HGNC:8024] | -1.2464 | 0.00263 |
| ENSG00000160179 | ABCG1 | ATP binding cassette subfamily G member 1 [Source:HGNC Symbol;Acc:HGNC:73] | -1.8883 | 0.00263 |
| ENSG00000135604 | STX11 | syntaxin 11 [Source:HGNC Symbol;Acc:HGNC:11429] | 1.5452 | 0.00263 |
| ENSG00000150471 | ADGRL3 | adhesion G protein-coupled receptor L3 [Source:HGNC Symbol;Acc:HGNC:20974] | -1.0345 | 0.00264 |
| ENSG00000197550 | NA | zinc finger pseudogene | -1.3599 | 0.00264 |
| ENSG00000157978 | LDLRAP1 | low density lipoprotein receptor adaptor protein 1 [Source:HGNC Symbol;Acc:HGNC:18640] | -0.5293 | 0.00264 |
| ENSG00000156042 | CFAP70 | cilia and flagella associated protein 70 [Source:HGNC Symbol;Acc:HGNC:30726] | -0.6473 | 0.00264 |
| ENSG00000260369 | NA | novel transcript | -1.3047 | 0.00264 |
| ENSG00000102048 | ASB9 | ankyrin repeat and SOCS box containing 9 [Source:HGNC Symbol;Acc:HGNC:17184] | -2.0046 | 0.00264 |
| ENSG00000227855 | DPY19L2P3 | DPY19L2 pseudogene 3 [Source:HGNC Symbol;Acc:HGNC:22367] | -0.7427 | 0.00265 |
| ENSG00000169715 | MT1E | metallothionein 1E [Source:HGNC Symbol;Acc:HGNC:7397] | 0.6160 | 0.00267 |
| ENSG00000179044 | EXOC3L1 | exocyst complex component 3 like 1 [Source:HGNC Symbol;Acc:HGNC:27540] | -1.0219 | 0.00267 |
| ENSG00000268364 | SMC5-AS1 | SMC5 antisense RNA 1 (head to head) [Source:HGNC Symbol;Acc:HGNC:48718] | 1.0228 | 0.00268 |
| ENSG00000112699 | GMDS | GDP-mannose 4,6-dehydratase [Source:HGNC Symbol;Acc:HGNC:4369] | 0.6441 | 0.00268 |
| ENSG00000236790 | LINC00299 | long intergenic non-protein coding RNA 299 [Source:HGNC Symbol;Acc:HGNC:27940] | -0.7909 | 0.00268 |
| ENSG00000223797 | ENTPD3-AS1 | ENTPD3 antisense RNA 1 [Source:HGNC Symbol;Acc:HGNC:26710] | -0.9430 | 0.00269 |
| ENSG00000286248 | NA | novel transcript, antisense to P2RX7 | -1.7569 | 0.00269 |
| ENSG00000247982 | LINC00926 | long intergenic non-protein coding RNA 926 [Source:HGNC Symbol;Acc:HGNC:27514] | 1.0546 | 0.00269 |
| ENSG00000272463 | NA | novel transcript | -1.5397 | 0.00269 |
| ENSG00000095383 | TBC1D2 | TBC1 domain family member 2 [Source:HGNC Symbol;Acc:HGNC:18026] | -0.5307 | 0.00270 |
| ENSG00000077044 | DGKD | diacylglycerol kinase delta [Source:HGNC Symbol;Acc:HGNC:2851] | 0.6153 | 0.00271 |
| ENSG00000019186 | CYP24A1 | cytochrome P450 family 24 subfamily A member 1 [Source:HGNC Symbol;Acc:HGNC:2602] | 2.6564 | 0.00271 |
| ENSG00000075073 | TACR2 | tachykinin receptor 2 [Source:HGNC Symbol;Acc:HGNC:11527] | 2.0621 | 0.00271 |
| ENSG00000234076 | TPRG1-AS1 | TPRG1 antisense RNA 1 [Source:HGNC Symbol;Acc:HGNC:42391] | -1.4117 | 0.00271 |
| ENSG00000178665 | ZNF713 | zinc finger protein 713 [Source:HGNC Symbol;Acc:HGNC:22043] | -0.6516 | 0.00272 |
| ENSG00000214510 | SPINK13 | serine peptidase inhibitor Kazal type 13 [Source:HGNC Symbol;Acc:HGNC:27200] | 0.6090 | 0.00272 |
| ENSG00000039523 | RIPOR1 | RHO family interacting cell polarization regulator 1 [Source:HGNC Symbol;Acc:HGNC:25836] | -0.5935 | 0.00273 |
| ENSG00000138641 | HERC3 | HECT and RLD domain containing E3 ubiquitin protein ligase 3 [Source:HGNC Symbol;Acc:HGNC:4876] | -0.5198 | 0.00273 |
| ENSG00000150627 | WDR17 | WD repeat domain 17 [Source:HGNC Symbol;Acc:HGNC:16661] | -1.1828 | 0.00273 |
| ENSG00000243970 | PPIEL | peptidylprolyl isomerase E like pseudogene [Source:HGNC Symbol;Acc:HGNC:33195] | 1.2729 | 0.00273 |
| ENSG00000111110 | PPM1H | protein phosphatase, Mg2+/Mn2+ dependent 1H [Source:HGNC Symbol;Acc:HGNC:18583] | -1.4352 | 0.00274 |
| ENSG00000172216 | CEBPB | CCAAT enhancer binding protein beta [Source:HGNC Symbol;Acc:HGNC:1834] | -0.5952 | 0.00275 |
| ENSG00000196335 | STK31 | serine/threonine kinase 31 [Source:HGNC Symbol;Acc:HGNC:11407] | -2.0570 | 0.00277 |
| ENSG00000223745 | CCDC18-AS1 | CCDC18 antisense RNA 1 [Source:HGNC Symbol;Acc:HGNC:52262] | -0.7326 | 0.00277 |
| ENSG00000245614 | DDX11-AS1 | DDX11 antisense RNA 1 [Source:HGNC Symbol;Acc:HGNC:44176] | -0.8549 | 0.00277 |
| ENSG00000075461 | CACNG4 | calcium voltage-gated channel auxiliary subunit gamma 4 [Source:HGNC Symbol;Acc:HGNC:1408] | -0.9616 | 0.00277 |
| ENSG00000141505 | ASGR1 | asialoglycoprotein receptor 1 [Source:HGNC Symbol;Acc:HGNC:742] | -0.5818 | 0.00278 |
| ENSG00000160953 | PWWP3A | PWWP domain containing 3A, DNA repair factor [Source:HGNC Symbol;Acc:HGNC:29641] | -0.5725 | 0.00282 |
| ENSG00000254416 | LINC02732 | long intergenic non-protein coding RNA 2732 [Source:HGNC Symbol;Acc:HGNC:54249] | 2.5059 | 0.00284 |
| ENSG00000234511 | C5orf58 | chromosome 5 open reading frame 58 [Source:HGNC Symbol;Acc:HGNC:37272] | -0.9949 | 0.00285 |
| ENSG00000095587 | TLL2 | tolloid like 2 [Source:HGNC Symbol;Acc:HGNC:11844] | -1.0995 | 0.00287 |
| ENSG00000183386 | FHL3 | four and a half LIM domains 3 [Source:HGNC Symbol;Acc:HGNC:3704] | 0.5460 | 0.00288 |
| ENSG00000263823 | NA | novel transcript, antisense to KIAA1012 | 1.5821 | 0.00289 |

| | | | | |
|---|---|---|---|---|
| ENSG00000227124 | ZNF717 | zinc finger protein 717 [Source:HGNC Symbol;Acc:HGNC:29448] | -0.5355 | 0.00290 |
| ENSG00000270276 | H4C15 | H4 clustered histone 15 [Source:HGNC Symbol;Acc:HGNC:29607] | -0.6909 | 0.00290 |
| ENSG00000137310 | TCF19 | transcription factor 19 [Source:HGNC Symbol;Acc:HGNC:11629] | -0.5584 | 0.00291 |
| ENSG00000196081 | ZNF724 | zinc finger protein 724 [Source:HGNC Symbol;Acc:HGNC:32460] | -0.5256 | 0.00291 |
| ENSG00000275481 | NA | novel transcript | -1.7359 | 0.00291 |
| ENSG00000272316 | NA | novel transcript | -0.7047 | 0.00297 |
| ENSG00000099984 | GSTT2 | glutathione S-transferase theta 2 (gene/pseudogene) [Source:HGNC Symbol;Acc:HGNC:4642] | -1.1336 | 0.00297 |
| ENSG00000149260 | CAPN5 | calpain 5 [Source:HGNC Symbol;Acc:HGNC:1482] | -0.6185 | 0.00298 |
| ENSG00000145248 | SLC10A4 | solute carrier family 10 member 4 [Source:HGNC Symbol;Acc:HGNC:22980] | -1.6057 | 0.00298 |
| ENSG00000188916 | INSYN2A | inhibitory synaptic factor 2A [Source:HGNC Symbol;Acc:HGNC:33859] | 2.5692 | 0.00298 |
| ENSG00000283431 | NA | novel zinc finger protein pseudogene | -1.0951 | 0.00299 |
| ENSG00000185306 | C12orf56 | chromosome 12 open reading frame 56 [Source:HGNC Symbol;Acc:HGNC:26967] | -4.7276 | 0.00299 |
| ENSG00000224254 | MTCO1P39 | MT-CO1 pseudogene 39 [Source:HGNC Symbol;Acc:HGNC:52104] | 1.9751 | 0.00300 |
| ENSG00000205810 | KLRC3 | killer cell lectin like receptor C3 [Source:HGNC Symbol;Acc:HGNC:6376] | 4.4635 | 0.00300 |
| ENSG00000143882 | ATP6V1C2 | ATPase H+ transporting V1 subunit C2 [Source:HGNC Symbol;Acc:HGNC:18264] | 1.1802 | 0.00301 |
| ENSG00000260611 | NA | novel transcript | -0.7710 | 0.00302 |
| ENSG00000104889 | RNASEH2A | ribonuclease H2 subunit A [Source:HGNC Symbol;Acc:HGNC:18518] | -0.5598 | 0.00304 |
| ENSG00000234072 | NA | novel transcript, antisense to GTF3C2 and EIF2B4 | 0.7281 | 0.00304 |
| ENSG00000272583 | NA | novel transcript | -0.8097 | 0.00305 |
| ENSG00000204745 | NA | anaphase promoting complex subunit 1 (ANAPC1) pseudogene | 0.5389 | 0.00305 |
| ENSG00000006047 | YBX2 | Y-box binding protein 2 [Source:HGNC Symbol;Acc:HGNC:17948] | 1.4036 | 0.00309 |
| ENSG00000009950 | MLXIPL | MLX interacting protein like [Source:HGNC Symbol;Acc:HGNC:12744] | -0.7983 | 0.00309 |
| ENSG00000242759 | LINC00882 | long intergenic non-protein coding RNA 882 [Source:HGNC Symbol;Acc:HGNC:48568] | -0.8080 | 0.00309 |
| ENSG00000257524 | NA | novel protein | 0.7529 | 0.00309 |
| ENSG00000198909 | MAP3K3 | mitogen-activated protein kinase kinase kinase 3 [Source:HGNC Symbol;Acc:HGNC:6855] | -0.7100 | 0.00310 |
| ENSG00000227486 | NA | novel transcript | -1.0009 | 0.00310 |
| ENSG00000087510 | TFAP2C | transcription factor AP-2 gamma [Source:HGNC Symbol;Acc:HGNC:11744] | 0.7461 | 0.00310 |
| ENSG00000178685 | PARP10 | poly(ADP-ribose) polymerase family member 10 [Source:HGNC Symbol;Acc:HGNC:25895] | -0.5204 | 0.00313 |
| ENSG00000266903 | NA | novel transcript, antisense to CEACAM16, CEACAM19, PVR & IGSF23 | -0.7047 | 0.00314 |
| ENSG00000139737 | SLAIN1 | SLAIN motif family member 1 [Source:HGNC Symbol;Acc:HGNC:26387] | -0.5381 | 0.00315 |
| ENSG00000180902 | D2HGDH | D-2-hydroxyglutarate dehydrogenase [Source:HGNC Symbol;Acc:HGNC:28358] | -0.6998 | 0.00316 |
| ENSG00000260000 | NA | novel transcript, antisense to ASCC3 | -0.6515 | 0.00316 |
| ENSG00000135517 | MIP | major intrinsic protein of lens fiber [Source:HGNC Symbol;Acc:HGNC:7103] | 2.3124 | 0.00319 |
| ENSG00000286443 | NA | novel transcript, antisense to PLXDC1 | -3.9692 | 0.00319 |
| ENSG00000104765 | BNIP3L | BCL2 interacting protein 3 like [Source:HGNC Symbol;Acc:HGNC:1085] | -0.5231 | 0.00321 |
| ENSG00000136867 | SLC31A2 | solute carrier family 31 member 2 [Source:HGNC Symbol;Acc:HGNC:11017] | -1.0739 | 0.00323 |
| ENSG00000162267 | ITIH3 | inter-alpha-trypsin inhibitor heavy chain 3 [Source:HGNC Symbol;Acc:HGNC:6168] | -0.8859 | 0.00326 |
| ENSG00000184486 | POU3F2 | POU class 3 homeobox 2 [Source:HGNC Symbol;Acc:HGNC:9215] | -0.6739 | 0.00329 |
| ENSG00000259234 | ANKRD34C-AS1 | ANKRD34C antisense RNA 1 [Source:HGNC Symbol;Acc:HGNC:48618] | 3.2923 | 0.00331 |
| ENSG00000084734 | GCKR | glucokinase regulator [Source:HGNC Symbol;Acc:HGNC:4196] | 3.4769 | 0.00332 |
| ENSG00000109220 | CHIC2 | cysteine rich hydrophobic domain 2 [Source:HGNC Symbol;Acc:HGNC:1935] | 0.5074 | 0.00333 |
| ENSG00000176428 | VPS37D | VPS37D subunit of ESCRT-I [Source:HGNC Symbol;Acc:HGNC:18287] | -1.1506 | 0.00333 |
| ENSG00000025423 | HSD17B6 | hydroxysteroid 17-beta dehydrogenase 6 [Source:HGNC Symbol;Acc:HGNC:23316] | -0.8571 | 0.00334 |

| | | | | |
|---|---|---|---|---|
| ENSG00000196839 | ADA | adenosine deaminase [Source:HGNC Symbol;Acc:HGNC:186] | -1.1674 | 0.00335 |
| ENSG00000274020 | LINC01138 | long intergenic non-protein coding RNA 1138 [Source:HGNC Symbol;Acc:HGNC:49454] | 0.9191 | 0.00335 |
| ENSG00000151445 | VIPAS39 | VPS33B interacting protein, apical-basolateral polarity regulator, spe-39 homolog [Source:HGNC Symbol;Acc:HGNC:20347] | -0.5137 | 0.00337 |
| ENSG00000120903 | CHRNA2 | cholinergic receptor nicotinic alpha 2 subunit [Source:HGNC Symbol;Acc:HGNC:1956] | -3.2201 | 0.00337 |
| ENSG00000276900 | NA | novel transcript, antisense to AMN1 | -0.5076 | 0.00337 |
| ENSG00000271647 | KRT8P47 | keratin 8 pseudogene 47 [Source:HGNC Symbol;Acc:HGNC:44461] | 0.8613 | 0.00338 |
| ENSG00000279368 | NA | TEC | -0.7759 | 0.00338 |
| ENSG00000132622 | HSPA12B | heat shock protein family A (Hsp70) member 12B [Source:HGNC Symbol;Acc:HGNC:16193] | -0.9414 | 0.00339 |
| ENSG00000231824 | AKAIN1 | A-kinase anchor inhibitor 1 [Source:HGNC Symbol;Acc:HGNC:28285] | -1.8118 | 0.00341 |
| ENSG00000266094 | RASSF5 | Ras association domain family member 5 [Source:HGNC Symbol;Acc:HGNC:17609] | -0.6512 | 0.00344 |
| ENSG00000102302 | FGD1 | FYVE, RhoGEF and PH domain containing 1 [Source:HGNC Symbol;Acc:HGNC:3663] | -0.5057 | 0.00344 |
| ENSG00000142544 | CTU1 | cytosolic thiouridylase subunit 1 [Source:HGNC Symbol;Acc:HGNC:29590] | 0.5379 | 0.00344 |
| ENSG00000110721 | CHKA | choline kinase alpha [Source:HGNC Symbol;Acc:HGNC:1937] | -0.5069 | 0.00345 |
| ENSG00000169926 | KLF13 | Kruppel like factor 13 [Source:HGNC Symbol;Acc:HGNC:13672] | -0.5939 | 0.00346 |
| ENSG00000185002 | RFX6 | regulatory factor X6 [Source:HGNC Symbol;Acc:HGNC:21478] | 2.5891 | 0.00346 |
| ENSG00000131323 | TRAF3 | TNF receptor associated factor 3 [Source:HGNC Symbol;Acc:HGNC:12033] | -0.5900 | 0.00347 |
| ENSG00000214942 | NA | novel transcript | 2.1634 | 0.00347 |
| ENSG00000025434 | NR1H3 | nuclear receptor subfamily 1 group H member 3 [Source:HGNC Symbol;Acc:HGNC:7966] | -0.7021 | 0.00347 |
| ENSG00000103034 | NDRG4 | NDRG family member 4 [Source:HGNC Symbol;Acc:HGNC:14466] | 0.8915 | 0.00348 |
| ENSG00000091490 | SEL1L3 | SEL1L family member 3 [Source:HGNC Symbol;Acc:HGNC:29108] | 0.8303 | 0.00348 |
| ENSG00000147251 | DOCK11 | dedicator of cytokinesis 11 [Source:HGNC Symbol;Acc:HGNC:23483] | -0.6401 | 0.00351 |
| ENSG00000237232 | ZNF295-AS1 | ZNF295 antisense RNA 1 [Source:HGNC Symbol;Acc:HGNC:23130] | 1.6456 | 0.00353 |
| ENSG00000285407 | NA | novel transcript | -1.1585 | 0.00353 |
| ENSG00000159423 | ALDH4A1 | aldehyde dehydrogenase 4 family member A1 [Source:HGNC Symbol;Acc:HGNC:406] | -0.5792 | 0.00354 |
| ENSG00000224043 | CCNT2-AS1 | CCNT2 antisense RNA 1 [Source:HGNC Symbol;Acc:HGNC:40130] | 0.8965 | 0.00357 |
| ENSG00000186409 | CCDC30 | coiled-coil domain containing 30 [Source:HGNC Symbol;Acc:HGNC:26103] | -0.5714 | 0.00358 |
| ENSG00000169744 | LDB2 | LIM domain binding 2 [Source:HGNC Symbol;Acc:HGNC:6533] | -0.6010 | 0.00358 |
| ENSG00000244486 | SCARF2 | scavenger receptor class F member 2 [Source:HGNC Symbol;Acc:HGNC:19869] | -0.6728 | 0.00359 |
| ENSG00000118514 | ALDH8A1 | aldehyde dehydrogenase 8 family member A1 [Source:HGNC Symbol;Acc:HGNC:15471] | 0.6437 | 0.00359 |
| ENSG00000253327 | RAD21-AS1 | RAD21 antisense RNA 1 [Source:HGNC Symbol;Acc:HGNC:32158] | -0.8692 | 0.00360 |
| ENSG00000241489 | NA | novel protein | 1.0547 | 0.00363 |
| ENSG00000158458 | NRG2 | neuregulin 2 [Source:HGNC Symbol;Acc:HGNC:7998] | -1.2109 | 0.00363 |
| ENSG00000241860 | NA | novel transcript | -0.7377 | 0.00363 |
| ENSG00000280649 | NA | TEC | 0.6649 | 0.00365 |
| ENSG00000103202 | NME4 | NME/NM23 nucleoside diphosphate kinase 4 [Source:HGNC Symbol;Acc:HGNC:7852] | -0.9755 | 0.00368 |
| ENSG00000173267 | SNCG | synuclein gamma [Source:HGNC Symbol;Acc:HGNC:11141] | -0.5828 | 0.00369 |
| ENSG00000203797 | DDO | D-aspartate oxidase [Source:HGNC Symbol;Acc:HGNC:2727] | -0.9730 | 0.00371 |
| ENSG00000161791 | FMNL3 | formin like 3 [Source:HGNC Symbol;Acc:HGNC:23698] | 0.6179 | 0.00373 |
| ENSG00000129167 | TPH1 | tryptophan hydroxylase 1 [Source:HGNC Symbol;Acc:HGNC:12008] | 1.3481 | 0.00373 |
| ENSG00000278592 | NA | NA | 1.7816 | 0.00373 |
| ENSG00000287738 | NA | novel transcript | 2.7632 | 0.00373 |
| ENSG00000115507 | OTX1 | orthodenticle homeobox 1 [Source:HGNC Symbol;Acc:HGNC:8521] | -0.8473 | 0.00377 |

| | | | | |
|---|---|---|---|---|
| ENSG00000247199 | NA | novel transcript, antisense to SPINK9, SPINK7 and SPINK13 | 0.5474 | 0.00377 |
| ENSG00000250264 | NA | novel protein, TAP2-HLA-DOB readthrough | -2.2263 | 0.00380 |
| ENSG00000110060 | PUS3 | pseudouridine synthase 3 [Source:HGNC Symbol;Acc:HGNC:25461] | 0.5326 | 0.00380 |
| ENSG00000175920 | DOK7 | docking protein 7 [Source:HGNC Symbol;Acc:HGNC:26594] | 1.7244 | 0.00381 |
| ENSG00000177990 | DPY19L2 | dpy-19 like 2 [Source:HGNC Symbol;Acc:HGNC:19414] | -0.5589 | 0.00382 |
| ENSG00000258986 | TMEM179 | transmembrane protein 179 [Source:HGNC Symbol;Acc:HGNC:20137] | -0.5967 | 0.00384 |
| ENSG00000084674 | APOB | apolipoprotein B [Source:HGNC Symbol;Acc:HGNC:603] | 0.5398 | 0.00384 |
| ENSG00000178031 | ADAMTSL1 | ADAMTS like 1 [Source:HGNC Symbol;Acc:HGNC:14632] | 0.5320 | 0.00390 |
| ENSG00000140279 | DUOX2 | dual oxidase 2 [Source:HGNC Symbol;Acc:HGNC:13273] | 1.0037 | 0.00390 |
| ENSG00000280623 | PCAT14 | prostate cancer associated transcript 14 [Source:HGNC Symbol;Acc:HGNC:48977] | -1.5226 | 0.00393 |
| ENSG00000224950 | NA | novel transcript | 2.6617 | 0.00394 |
| ENSG00000119929 | CUTC | cutC copper transporter [Source:HGNC Symbol;Acc:HGNC:24271] | -0.5308 | 0.00397 |
| ENSG00000250790 | NA | novel transcript | -1.6889 | 0.00399 |
| ENSG00000206337 | HCP5 | HLA complex P5 [Source:HGNC Symbol;Acc:HGNC:21659] | -0.6966 | 0.00401 |
| ENSG00000165171 | METTL27 | methyltransferase like 27 [Source:HGNC Symbol;Acc:HGNC:19068] | -0.8429 | 0.00402 |
| ENSG00000265417 | NA | ATPase family, AAA domain containing (ATD1) pseudogene | -2.0438 | 0.00402 |
| ENSG00000169169 | CPT1C | carnitine palmitoyltransferase 1C [Source:HGNC Symbol;Acc:HGNC:18540] | -0.5028 | 0.00403 |
| ENSG00000248429 | FAM198B-AS1 | FAM198B antisense RNA 1 [Source:HGNC Symbol;Acc:HGNC:53132] | -1.1789 | 0.00409 |
| ENSG00000026508 | CD44 | CD44 molecule (Indian blood group) [Source:HGNC Symbol;Acc:HGNC:1681] | 2.0829 | 0.00409 |
| ENSG00000165071 | TMEM71 | transmembrane protein 71 [Source:HGNC Symbol;Acc:HGNC:26572] | 1.2493 | 0.00410 |
| ENSG00000125931 | CITED1 | Cbp/p300 interacting transactivator with Glu/Asp rich carboxy-terminal domain 1 [Source:HGNC Symbol;Acc:HGNC:1986] | 1.2082 | 0.00411 |
| ENSG00000142082 | SIRT3 | sirtuin 3 [Source:HGNC Symbol;Acc:HGNC:14931] | -0.5011 | 0.00413 |
| ENSG00000166432 | ZMAT1 | zinc finger matrin-type 1 [Source:HGNC Symbol;Acc:HGNC:29377] | -0.7273 | 0.00414 |
| ENSG00000167037 | SGSM1 | small G protein signaling modulator 1 [Source:HGNC Symbol;Acc:HGNC:29410] | -1.5575 | 0.00414 |
| ENSG00000138764 | CCNG2 | cyclin G2 [Source:HGNC Symbol;Acc:HGNC:1593] | -0.7257 | 0.00416 |
| ENSG00000272589 | ZSWIM8-AS1 | ZSWIM8 antisense RNA 1 [Source:HGNC Symbol;Acc:HGNC:45103] | -1.2254 | 0.00416 |
| ENSG00000081377 | CDC14B | cell division cycle 14B [Source:HGNC Symbol;Acc:HGNC:1719] | -0.5444 | 0.00417 |
| ENSG00000275527 | NA | novel transcript | -2.3925 | 0.00419 |
| ENSG00000274718 | NA | novel transcript | -0.8704 | 0.00419 |
| ENSG00000236844 | NA | ribosomal protein L24 (RPL24) pseudogene | 1.6600 | 0.00419 |
| ENSG00000155265 | GOLGA7B | golgin A7 family member B [Source:HGNC Symbol;Acc:HGNC:31668] | 0.5356 | 0.00420 |
| ENSG00000105088 | OLFM2 | olfactomedin 2 [Source:HGNC Symbol;Acc:HGNC:17189] | -0.5398 | 0.00420 |
| ENSG00000224939 | LINC00184 | long intergenic non-protein coding RNA 184 [Source:HGNC Symbol;Acc:HGNC:37192] | -3.1590 | 0.00420 |
| ENSG00000166228 | PCBD1 | pterin-4 alpha-carbinolamine dehydratase 1 [Source:HGNC Symbol;Acc:HGNC:8646] | -0.7884 | 0.00421 |
| ENSG00000167723 | TRPV3 | transient receptor potential cation channel subfamily V member 3 [Source:HGNC Symbol;Acc:HGNC:18084] | 3.0517 | 0.00424 |
| ENSG00000223685 | LINC00571 | long intergenic non-protein coding RNA 571 [Source:HGNC Symbol;Acc:HGNC:43721] | 1.3591 | 0.00425 |
| ENSG00000136274 | NACAD | NAC alpha domain containing [Source:HGNC Symbol;Acc:HGNC:22196] | 2.3726 | 0.00426 |
| ENSG00000269896 | NA | small nuclear ribonucleoprotein N (SNRPN) pseudogene | -1.6817 | 0.00429 |
| ENSG00000173805 | HAP1 | huntingtin associated protein 1 [Source:HGNC Symbol;Acc:HGNC:4812] | 0.6442 | 0.00433 |
| ENSG00000120899 | PTK2B | protein tyrosine kinase 2 beta [Source:HGNC Symbol;Acc:HGNC:9612] | -0.7755 | 0.00434 |
| ENSG00000135953 | MFSD9 | major facilitator superfamily domain containing 9 [Source:HGNC Symbol;Acc:HGNC:28158] | 0.5098 | 0.00434 |

| | | | | |
|---|---|---|---|---|
| ENSG00000008517 | IL32 | interleukin 32 [Source:HGNC Symbol;Acc:HGNC:16830] | 1.4886 | 0.00434 |
| ENSG00000260772 | NA | novel transcript | 0.6577 | 0.00434 |
| ENSG00000179104 | TMTC2 | transmembrane O-mannosyltransferase targeting cadherins 2 [Source:HGNC Symbol;Acc:HGNC:25440] | -1.4811 | 0.00435 |
| ENSG00000283662 | NA | novel transcript | -1.3189 | 0.00436 |
| ENSG00000187134 | AKR1C1 | aldo-keto reductase family 1 member C1 [Source:HGNC Symbol;Acc:HGNC:384] | -0.5478 | 0.00437 |
| ENSG00000279865 | NA | TEC | -1.1847 | 0.00437 |
| ENSG00000206127 | GOLGA8O | golgin A8 family member O [Source:HGNC Symbol;Acc:HGNC:44406] | -0.7213 | 0.00438 |
| ENSG00000215475 | SIAH3 | siah E3 ubiquitin protein ligase family member 3 [Source:HGNC Symbol;Acc:HGNC:30553] | -0.8096 | 0.00439 |
| ENSG00000255248 | MIR100HG | mir-100-let-7a-2-mir-125b-1 cluster host gene [Source:HGNC Symbol;Acc:HGNC:39522] | -0.7340 | 0.00439 |
| ENSG00000183682 | BMP8A | bone morphogenetic protein 8a [Source:HGNC Symbol;Acc:HGNC:21650] | 0.9108 | 0.00441 |
| ENSG00000136048 | DRAM1 | DNA damage regulated autophagy modulator 1 [Source:HGNC Symbol;Acc:HGNC:25645] | -2.2275 | 0.00444 |
| ENSG00000170044 | ZPLD1 | zona pellucida like domain containing 1 [Source:HGNC Symbol;Acc:HGNC:27022] | -0.5568 | 0.00447 |
| ENSG00000256894 | NA | novel transcript, antisense to SSPN | 1.5818 | 0.00449 |
| ENSG00000130699 | TAF4 | TATA-box binding protein associated factor 4 [Source:HGNC Symbol;Acc:HGNC:11537] | -0.5819 | 0.00449 |
| ENSG00000264578 | NA | novel transcript, antisense to AGPAT6 | 1.3648 | 0.00450 |
| ENSG00000250420 | AACSP1 | acetoacetyl-CoA synthetase pseudogene 1 [Source:HGNC Symbol;Acc:HGNC:18226] | -0.6314 | 0.00453 |
| ENSG00000261485 | PAN3-AS1 | PAN3 antisense RNA 1 [Source:HGNC Symbol;Acc:HGNC:39932] | 0.6340 | 0.00458 |
| ENSG00000176092 | CRYBG2 | crystallin beta-gamma domain containing 2 [Source:HGNC Symbol;Acc:HGNC:17295] | -1.3632 | 0.00458 |
| ENSG00000176566 | DCAF4L2 | DDB1 and CUL4 associated factor 4 like 2 [Source:HGNC Symbol;Acc:HGNC:26657] | -0.7331 | 0.00458 |
| ENSG00000271584 | LINC02550 | long intergenic non-protein coding RNA 2550 [Source:HGNC Symbol;Acc:HGNC:53585] | -1.0911 | 0.00459 |
| ENSG00000272505 | NA | novel transcript | 1.0567 | 0.00459 |
| ENSG00000266718 | NA | novel transcript, antisense MYO1D | 1.9187 | 0.00460 |
| ENSG00000261572 | NA | novel transcript, intronic to CMTM8 | -0.6425 | 0.00460 |
| ENSG00000152582 | SPEF2 | sperm flagellar 2 [Source:HGNC Symbol;Acc:HGNC:26293] | -0.6699 | 0.00461 |
| ENSG00000258943 | NA | novel transcript | 1.9878 | 0.00461 |
| ENSG00000166833 | NAV2 | neuron navigator 2 [Source:HGNC Symbol;Acc:HGNC:15997] | 1.3422 | 0.00462 |
| ENSG00000186479 | RGS7BP | regulator of G protein signaling 7 binding protein [Source:HGNC Symbol;Acc:HGNC:23271] | 2.2269 | 0.00462 |
| ENSG00000271380 | NA | uncharacterized LOC101928120 [Source:NCBI gene (formerly Entrezgene);Acc:101928120] | -0.8263 | 0.00462 |
| ENSG00000182472 | CAPN12 | calpain 12 [Source:HGNC Symbol;Acc:HGNC:13249] | 1.5062 | 0.00463 |
| ENSG00000148459 | PDSS1 | decaprenyl diphosphate synthase subunit 1 [Source:HGNC Symbol;Acc:HGNC:17759] | 0.5417 | 0.00468 |
| ENSG00000168772 | CXXC4 | CXXC finger protein 4 [Source:HGNC Symbol;Acc:HGNC:24593] | -1.5879 | 0.00471 |
| ENSG00000250874 | NA | novel transcript | -0.6956 | 0.00472 |
| ENSG00000234928 | LINC01659 | long intergenic non-protein coding RNA 1659 [Source:HGNC Symbol;Acc:HGNC:52447] | -2.4017 | 0.00474 |
| ENSG00000269054 | NA | novel transcript, antisense to ZNF497 | -1.2379 | 0.00475 |
| ENSG00000253731 | PCDHGA6 | protocadherin gamma subfamily A, 6 [Source:HGNC Symbol;Acc:HGNC:8704] | -0.6641 | 0.00475 |
| ENSG00000100625 | SIX4 | SIX homeobox 4 [Source:HGNC Symbol;Acc:HGNC:10890] | 0.6951 | 0.00478 |
| ENSG00000196458 | ZNF605 | zinc finger protein 605 [Source:HGNC Symbol;Acc:HGNC:28068] | -0.5158 | 0.00481 |
| ENSG00000127311 | HELB | DNA helicase B [Source:HGNC Symbol;Acc:HGNC:17196] | 0.7079 | 0.00481 |
| ENSG00000230844 | ZNF674-AS1 | ZNF674 antisense RNA 1 (head to head) [Source:HGNC Symbol;Acc:HGNC:44266] | 0.5531 | 0.00481 |
| ENSG00000286161 | NA | novel transcript | 1.3158 | 0.00481 |
| ENSG00000253669 | GASAL1 | growth arrest associated lncRNA 1 [Source:HGNC Symbol;Acc:HGNC:53461] | -1.1033 | 0.00483 |
| ENSG00000102390 | PBDC1 | polysaccharide biosynthesis domain containing 1 [Source:HGNC Symbol;Acc:HGNC:28790] | 0.5040 | 0.00484 |
| ENSG00000139445 | FOXN4 | forkhead box N4 [Source:HGNC Symbol;Acc:HGNC:21399] | -0.9980 | 0.00484 |

| | | | | |
|---|---|---|---|---|
| ENSG00000240583 | AQP1 | aquaporin 1 (Colton blood group) [Source:HGNC Symbol;Acc:HGNC:633] | -0.5728 | 0.00485 |
| ENSG00000140682 | TGFB1I1 | transforming growth factor beta 1 induced transcript 1 [Source:HGNC Symbol;Acc:HGNC:11767] | 0.7507 | 0.00488 |
| ENSG00000115318 | LOXL3 | lysyl oxidase like 3 [Source:HGNC Symbol;Acc:HGNC:13869] | -0.5892 | 0.00490 |
| ENSG00000277957 | SENP3-EIF4A1 | SENP3-EIF4A1 readthrough (NMD candidate) [Source:HGNC Symbol;Acc:HGNC:49182] | 0.8548 | 0.00495 |
| ENSG00000196605 | ZNF846 | zinc finger protein 846 [Source:HGNC Symbol;Acc:HGNC:27260] | -0.6176 | 0.00495 |
| ENSG00000186088 | GSAP | gamma-secretase activating protein [Source:HGNC Symbol;Acc:HGNC:28042] | -0.6863 | 0.00496 |
| ENSG00000150938 | CRIM1 | cysteine rich transmembrane BMP regulator 1 [Source:HGNC Symbol;Acc:HGNC:2359] | 1.5577 | 0.00501 |
| ENSG00000012504 | NR1H4 | nuclear receptor subfamily 1 group H member 4 [Source:HGNC Symbol;Acc:HGNC:7967] | -0.5671 | 0.00502 |
| ENSG00000164124 | TMEM144 | transmembrane protein 144 [Source:HGNC Symbol;Acc:HGNC:25633] | -0.7148 | 0.00506 |
| ENSG00000234197 | ETV5-AS1 | ETV5 antisense RNA 1 [Source:HGNC Symbol;Acc:HGNC:40222] | 2.9161 | 0.00508 |
| ENSG00000214999 | NA | novel transcript, antisense to ALOX12B | -2.2864 | 0.00508 |
| ENSG00000227467 | LINC01537 | long intergenic non-protein coding RNA 1537 [Source:HGNC Symbol;Acc:HGNC:51301] | -1.0010 | 0.00508 |
| ENSG00000254852 | NPIPA2 | nuclear pore complex interacting protein family member A2 [Source:HGNC Symbol;Acc:HGNC:41979] | -2.1384 | 0.00511 |
| ENSG00000223547 | ZNF844 | zinc finger protein 844 [Source:HGNC Symbol;Acc:HGNC:25932] | -0.8230 | 0.00512 |
| ENSG00000279833 | NA | novel transcript, antisense to CBX7 | -1.4472 | 0.00513 |
| ENSG00000164398 | ACSL6 | acyl-CoA synthetase long chain family member 6 [Source:HGNC Symbol;Acc:HGNC:16496] | -0.9164 | 0.00513 |
| ENSG00000163449 | TMEM169 | transmembrane protein 169 [Source:HGNC Symbol;Acc:HGNC:25130] | -0.6308 | 0.00516 |
| ENSG00000274818 | NA | novel transcript | 1.1126 | 0.00517 |
| ENSG00000213967 | ZNF726 | zinc finger protein 726 [Source:HGNC Symbol;Acc:HGNC:32462] | -0.6531 | 0.00518 |
| ENSG00000171320 | ESCO2 | establishment of sister chromatid cohesion N-acetyltransferase 2 [Source:HGNC Symbol;Acc:HGNC:27230] | -0.5677 | 0.00519 |
| ENSG00000188177 | ZC3H6 | zinc finger CCCH-type containing 6 [Source:HGNC Symbol;Acc:HGNC:24762] | -0.7144 | 0.00521 |
| ENSG00000182870 | GALNT9 | polypeptide N-acetylgalactosaminyltransferase 9 [Source:HGNC Symbol;Acc:HGNC:4131] | 2.2055 | 0.00521 |
| ENSG00000171763 | SPATA5L1 | spermatogenesis associated 5 like 1 [Source:HGNC Symbol;Acc:HGNC:28762] | 0.5106 | 0.00522 |
| ENSG00000227060 | LINC00629 | long intergenic non-protein coding RNA 629 [Source:HGNC Symbol;Acc:HGNC:44262] | -1.3216 | 0.00524 |
| ENSG00000275765 | NA | novel transcript | -1.3200 | 0.00524 |
| ENSG00000167065 | DUSP18 | dual specificity phosphatase 18 [Source:HGNC Symbol;Acc:HGNC:18484] | 0.5931 | 0.00524 |
| ENSG00000228146 | CASP16P | caspase 16, pseudogene [Source:HGNC Symbol;Acc:HGNC:27290] | -1.0139 | 0.00526 |
| ENSG00000140043 | PTGR2 | prostaglandin reductase 2 [Source:HGNC Symbol;Acc:HGNC:20149] | -0.5743 | 0.00526 |
| ENSG00000169696 | ASPSCR1 | ASPSCR1 tether for SLC2A4, UBX domain containing [Source:HGNC Symbol;Acc:HGNC:13825] | -0.5153 | 0.00529 |
| ENSG00000026559 | KCNG1 | potassium voltage-gated channel modifier subfamily G member 1 [Source:HGNC Symbol;Acc:HGNC:6248] | -2.3893 | 0.00531 |
| ENSG00000286311 | NA | novel transcript, antisense to NOL4L | -2.6154 | 0.00531 |
| ENSG00000227992 | NA | pseudogene similar to part of ubiquitin carboxyl-terminal hydrolase L5 UCHL5 | 1.6090 | 0.00532 |
| ENSG00000125798 | FOXA2 | forkhead box A2 [Source:HGNC Symbol;Acc:HGNC:5022] | -1.1168 | 0.00533 |
| ENSG00000004848 | ARX | aristaless related homeobox [Source:HGNC Symbol;Acc:HGNC:18060] | 1.5095 | 0.00533 |
| ENSG00000078328 | RBFOX1 | RNA binding fox-1 homolog 1 [Source:HGNC Symbol;Acc:HGNC:18222] | -2.7605 | 0.00536 |
| ENSG00000220785 | MTMR9LP | myotubularin related protein 9 like, pseudogene [Source:HGNC Symbol;Acc:HGNC:27920] | -0.8630 | 0.00537 |
| ENSG00000157470 | FAM81A | family with sequence similarity 81 member A [Source:HGNC Symbol;Acc:HGNC:28379] | -0.9901 | 0.00540 |
| ENSG00000163359 | COL6A3 | collagen type VI alpha 3 chain [Source:HGNC Symbol;Acc:HGNC:2213] | -1.9225 | 0.00540 |
| ENSG00000120254 | MTHFD1L | methylenetetrahydrofolate dehydrogenase (NADP+ dependent) 1 like [Source:HGNC Symbol;Acc:HGNC:21055] | 0.6594 | 0.00541 |
| ENSG00000234571 | H2BP2 | H2B histone pseudogene 2 [Source:HGNC Symbol;Acc:HGNC:20654] | -0.8777 | 0.00544 |
| ENSG00000180447 | GAS1 | growth arrest specific 1 [Source:HGNC Symbol;Acc:HGNC:4165] | 0.8057 | 0.00545 |

| | | | | |
|---|---|---|---|---|
| ENSG00000267072 | NAGPA-AS1 | NAGPA antisense RNA 1 [Source:HGNC Symbol;Acc:HGNC:44184] | -1.8437 | 0.00547 |
| ENSG00000153885 | KCTD15 | potassium channel tetramerization domain containing 15 [Source:HGNC Symbol;Acc:HGNC:23297] | 0.9213 | 0.00548 |
| ENSG00000267454 | ZNF582-AS1 | ZNF582 antisense RNA 1 (head to head) [Source:HGNC Symbol;Acc:HGNC:25213] | -1.4261 | 0.00550 |
| ENSG00000285314 | NA | novel transcript | -1.8863 | 0.00550 |
| ENSG00000283122 | HYMAI | hydatidiform mole associated and imprinted [Source:HGNC Symbol;Acc:HGNC:5326] | -0.6876 | 0.00551 |
| ENSG00000287737 | NA | novel transcript | 0.5143 | 0.00553 |
| ENSG00000155158 | TTC39B | tetratricopeptide repeat domain 39B [Source:HGNC Symbol;Acc:HGNC:23704] | -0.5022 | 0.00557 |
| ENSG00000283312 | NA | GTF2I repeat domain containing 2(GTF2IRD2) pseudogene | -1.0525 | 0.00557 |
| ENSG00000170365 | SMAD1 | SMAD family member 1 [Source:HGNC Symbol;Acc:HGNC:6767] | -0.6234 | 0.00560 |
| ENSG00000079482 | OPHN1 | oligophrenin 1 [Source:HGNC Symbol;Acc:HGNC:8148] | -0.6489 | 0.00561 |
| ENSG00000182272 | B4GALNT4 | beta-1,4-N-acetyl-galactosaminyltransferase 4 [Source:HGNC Symbol;Acc:HGNC:26315] | -0.7125 | 0.00567 |
| ENSG00000279489 | NA | tec | 1.4346 | 0.00568 |
| ENSG00000285804 | NA | novel transcript | -0.7563 | 0.00568 |
| ENSG00000230091 | TMEM254-AS1 | TMEM254 antisense RNA 1 [Source:HGNC Symbol;Acc:HGNC:27340] | -1.0796 | 0.00569 |
| ENSG00000272831 | NA | novel transcript, antisense to RABGEF1 | -0.8456 | 0.00569 |
| ENSG00000139200 | PIANP | PILR alpha associated neural protein [Source:HGNC Symbol;Acc:HGNC:25338] | -0.5692 | 0.00570 |
| ENSG00000223478 | NA | novel transcript | -0.6044 | 0.00571 |
| ENSG00000197951 | ZNF71 | zinc finger protein 71 [Source:HGNC Symbol;Acc:HGNC:13141] | -0.5144 | 0.00572 |
| ENSG00000213139 | CRYGS | crystallin gamma S [Source:HGNC Symbol;Acc:HGNC:2417] | 1.0663 | 0.00572 |
| ENSG00000117525 | 3,00 F | coagulation factor III, tissue factor [Source:HGNC Symbol;Acc:HGNC:3541] | 0.8246 | 0.00572 |
| ENSG00000268203 | NA | novel transcript, antisense to TUBB4A | -1.5848 | 0.00579 |
| ENSG00000276644 | DACH1 | dachshund family transcription factor 1 [Source:HGNC Symbol;Acc:HGNC:2663] | -0.5274 | 0.00579 |
| ENSG00000176714 | CCDC121 | coiled-coil domain containing 121 [Source:HGNC Symbol;Acc:HGNC:25833] | -0.9688 | 0.00580 |
| ENSG00000169418 | 1,00 NPR | natriuretic peptide receptor 1 [Source:HGNC Symbol;Acc:HGNC:7943] | 1.4512 | 0.00580 |
| ENSG00000048405 | ZNF800 | zinc finger protein 800 [Source:HGNC Symbol;Acc:HGNC:27267] | 0.5126 | 0.00580 |
| ENSG00000149636 | DSN1 | DSN1 component of MIS12 kinetochore complex [Source:HGNC Symbol;Acc:HGNC:16165] | -0.5335 | 0.00581 |
| ENSG00000261613 | NA | novel transcript, antisense to PDPK1 | 0.6055 | 0.00584 |
| ENSG00000106536 | POU6F2 | POU class 6 homeobox 2 [Source:HGNC Symbol;Acc:HGNC:21694] | 0.5840 | 0.00586 |
| ENSG00000185158 | LRRC37B | leucine rich repeat containing 37B [Source:HGNC Symbol;Acc:HGNC:29070] | -0.5280 | 0.00587 |
| ENSG00000278616 | BEND3P3 | BEN domain containing 3 pseudogene 3 [Source:HGNC Symbol;Acc:HGNC:45016] | 0.7297 | 0.00587 |
| ENSG00000285580 | NA | Novel transcript | 1.8964 | 0.00587 |
| ENSG00000242396 | NA | novel transcript | 1.4876 | 0.00590 |
| ENSG00000242612 | DECR2 | 2,4-dienoyl-CoA reductase 2 [Source:HGNC Symbol;Acc:HGNC:2754] | -0.5945 | 0.00591 |
| ENSG00000137098 | SPAG8 | sperm associated antigen 8 [Source:HGNC Symbol;Acc:HGNC:14105] | -1.1237 | 0.00592 |
| ENSG00000008226 | DLEC1 | DLEC1 cilia and flagella associated protein [Source:HGNC Symbol;Acc:HGNC:2899] | 1.3853 | 0.00593 |
| ENSG00000164088 | PPM1M | protein phosphatase, Mg2+/Mn2+ dependent 1M [Source:HGNC Symbol;Acc:HGNC:26506] | -0.5352 | 0.00596 |
| ENSG00000235033 | DAAM2-AS1 | DAAM2 antisense RNA 1 [Source:HGNC Symbol;Acc:HGNC:40830] | 1.0625 | 0.00597 |
| ENSG00000172817 | CYP7B1 | cytochrome P450 family 7 subfamily B member 1 [Source:HGNC Symbol;Acc:HGNC:2652] | 0.8529 | 0.00598 |
| ENSG00000177045 | SIX5 | SIX homeobox 5 [Source:HGNC Symbol;Acc:HGNC:10891] | -0.6222 | 0.00600 |
| ENSG00000227036 | LINC00511 | long intergenic non-protein coding RNA 511 [Source:HGNC Symbol;Acc:HGNC:43564] | 0.8450 | 0.00603 |
| ENSG00000105376 | ICAM5 | intercellular adhesion molecule 5 [Source:HGNC Symbol;Acc:HGNC:5348] | 0.7227 | 0.00605 |
| ENSG00000205791 | LOH12CR2 | loss of heterozygosity on chromosome 12, region 2 [Source:HGNC Symbol;Acc:HGNC:26524] | -0.9510 | 0.00606 |
| ENSG00000225361 | PPP1R26-AS1 | PPP1R26 antisense RNA 1 [Source:HGNC Symbol;Acc:HGNC:48717] | -1.2809 | 0.00614 |

| | | | | |
|---|---|---|---|---|
| ENSG00000229932 | YWHAZP3 | tyrosine 3-monooxygenase/tryptophan 5-monooxygenase activation protein zeta pseudogene 3 [Source:HGNC Symbol;Acc:HGNC:31101] | 1.3923 | 0.00614 |
| ENSG00000125744 | RTN2 | reticulon 2 [Source:HGNC Symbol;Acc:HGNC:10468] | 0.6016 | 0.00616 |
| ENSG00000160588 | MPZL3 | myelin protein zero like 3 [Source:HGNC Symbol;Acc:HGNC:27279] | 0.9503 | 0.00620 |
| ENSG00000178814 | OPLAH | 5-oxoprolinase, ATP-hydrolysing [Source:HGNC Symbol;Acc:HGNC:8149] | -0.7083 | 0.00622 |
| ENSG00000254893 | NA | RAP1B, member of RAS oncogene family pseudogene | 0.5141 | 0.00622 |
| ENSG00000256433 | NA | novel transcript | 0.8681 | 0.00625 |
| ENSG00000250546 | NA | novel transcript | -0.7021 | 0.00626 |
| ENSG00000092758 | COL9A3 | collagen type IX alpha 3 chain [Source:HGNC Symbol;Acc:HGNC:2219] | -4.6027 | 0.00627 |
| ENSG00000271858 | NA | cytochrome b561 family member D2 [Source:NCBI gene (formerly Entrezgene);Acc:11068] | -1.0771 | 0.00629 |
| ENSG00000232656 | IDI2-AS1 | IDI2 antisense RNA 1 [Source:HGNC Symbol;Acc:HGNC:30885] | -0.7151 | 0.00631 |
| ENSG00000206140 | TMEM191C | transmembrane protein 191C [Source:HGNC Symbol;Acc:HGNC:33601] | -0.8158 | 0.00632 |
| ENSG00000221886 | ZBED8 | zinc finger BED-type containing 8 [Source:HGNC Symbol;Acc:HGNC:30804] | -0.6087 | 0.00632 |
| ENSG00000124208 | TMEM189-UBE2V1 | TMEM189-UBE2V1 readthrough [Source:HGNC Symbol;Acc:HGNC:33521] | 7.1689 | 0.00633 |
| ENSG00000107020 | PLGRKT | plasminogen receptor with a C-terminal lysine [Source:HGNC Symbol;Acc:HGNC:23633] | -0.6821 | 0.00633 |
| ENSG00000131061 | ZNF341 | zinc finger protein 341 [Source:HGNC Symbol;Acc:HGNC:15992] | -0.6477 | 0.00637 |
| ENSG00000287537 | NA | novel transcript | -2.2329 | 0.00637 |
| ENSG00000237054 | PRMT5-AS1 | PRMT5 antisense RNA 1 [Source:HGNC Symbol;Acc:HGNC:40533] | 1.0650 | 0.00639 |
| ENSG00000267296 | CEBPA-DT | CEBPA divergent transcript [Source:HGNC Symbol;Acc:HGNC:25710] | -1.7444 | 0.00639 |
| ENSG00000273419 | NA | novel transcript, antisense to ZNF862 | 0.6655 | 0.00642 |
| ENSG00000188910 | GJB3 | gap junction protein beta 3 [Source:HGNC Symbol;Acc:HGNC:4285] | 1.9993 | 0.00643 |
| ENSG00000175463 | TBC1D10C | TBC1 domain family member 10C [Source:HGNC Symbol;Acc:HGNC:24702] | 1.2277 | 0.00645 |
| ENSG00000141485 | SLC13A5 | solute carrier family 13 member 5 [Source:HGNC Symbol;Acc:HGNC:23089] | -2.0806 | 0.00650 |
| ENSG00000260742 | NA | novel transcript, antisense to SSFA2 | 0.9899 | 0.00650 |
| ENSG00000143869 | GDF7 | growth differentiation factor 7 [Source:HGNC Symbol;Acc:HGNC:4222] | 1.5963 | 0.00650 |
| ENSG00000270332 | SMC2-AS1 | SMC2 antisense RNA 1 (head to head) [Source:HGNC Symbol;Acc:HGNC:50827] | -1.7138 | 0.00652 |
| ENSG00000180479 | ZNF571 | zinc finger protein 571 [Source:HGNC Symbol;Acc:HGNC:25000] | -0.5198 | 0.00656 |
| ENSG00000272419 | LINC01145 | long intergenic non-protein coding RNA 1145 [Source:HGNC Symbol;Acc:HGNC:49462] | 0.7655 | 0.00657 |
| ENSG00000285727 | NA | novel transcript | -3.1477 | 0.00657 |
| ENSG00000165072 | MAMDC2 | MAM domain containing 2 [Source:HGNC Symbol;Acc:HGNC:23673] | 1.1473 | 0.00657 |
| ENSG00000279619 | NA | novel transcript | -0.6948 | 0.00658 |
| ENSG00000230021 | NA | septin 14 (SEPT14) pseudogene | -3.5098 | 0.00659 |
| ENSG00000111271 | ACAD10 | acyl-CoA dehydrogenase family member 10 [Source:HGNC Symbol;Acc:HGNC:21597] | -0.5778 | 0.00660 |
| ENSG00000197483 | ZNF628 | zinc finger protein 628 [Source:HGNC Symbol;Acc:HGNC:28054] | 0.5844 | 0.00661 |
| ENSG00000077092 | RARB | retinoic acid receptor beta [Source:HGNC Symbol;Acc:HGNC:9865] | -0.7298 | 0.00663 |
| ENSG00000081041 | CXCL2 | C-X-C motif chemokine ligand 2 [Source:HGNC Symbol;Acc:HGNC:4603] | 1.8547 | 0.00663 |
| ENSG00000108878 | CACNG1 | calcium voltage-gated channel auxiliary subunit gamma 1 [Source:HGNC Symbol;Acc:HGNC:1405] | -2.6031 | 0.00665 |
| ENSG00000277449 | CEBPB-AS1 | CEBPB antisense RNA 1 [Source:HGNC Symbol;Acc:HGNC:51226] | -0.7816 | 0.00665 |
| ENSG00000167555 | ZNF528 | zinc finger protein 528 [Source:HGNC Symbol;Acc:HGNC:29384] | -0.7051 | 0.00667 |
| ENSG00000250423 | KIAA1210 | KIAA1210 [Source:HGNC Symbol;Acc:HGNC:29218] | -2.6871 | 0.00669 |
| ENSG00000129173 | E2F8 | E2F transcription factor 8 [Source:HGNC Symbol;Acc:HGNC:24727] | 0.5436 | 0.00673 |
| ENSG00000254990 | NA | novel transcript | 2.0769 | 0.00675 |
| ENSG00000166105 | GLB1L3 | galactosidase beta 1 like 3 [Source:HGNC Symbol;Acc:HGNC:25147] | 0.5711 | 0.00675 |

| | | | | |
|---|---|---|---|---|
| ENSG00000260793 | NA | novel transcript, antisense to UBTF | 0.6513 | 0.00678 |
| ENSG00000132510 | KDM6B | lysine demethylase 6B [Source:HGNC Symbol;Acc:HGNC:29012] | 0.6150 | 0.00679 |
| ENSG00000250012 | NA | novel transcript | -0.9235 | 0.00682 |
| ENSG00000179071 | CCDC89 | coiled-coil domain containing 89 [Source:HGNC Symbol;Acc:HGNC:26762] | -0.6214 | 0.00686 |
| ENSG00000204956 | PCDHGA1 | protocadherin gamma subfamily A, 1 [Source:HGNC Symbol;Acc:HGNC:8696] | -0.5019 | 0.00689 |
| ENSG00000166002 | SMCO4 | single-pass membrane protein with coiled-coil domains 4 [Source:HGNC Symbol;Acc:HGNC:24810] | 1.3888 | 0.00698 |
| ENSG00000214273 | AGGF1P1 | angiogenic factor with G-patch and FHA domains 1 pseudogene 1 [Source:HGNC Symbol;Acc:HGNC:38051] | -1.4315 | 0.00699 |
| ENSG00000101134 | DOK5 | docking protein 5 [Source:HGNC Symbol;Acc:HGNC:16173] | 2.0840 | 0.00700 |
| ENSG00000147434 | CHRNA6 | cholinergic receptor nicotinic alpha 6 subunit [Source:HGNC Symbol;Acc:HGNC:15963] | 2.5415 | 0.00701 |
| ENSG00000254294 | IMPDH1P6 | inosine monophosphate dehydrogenase 1 pseudogene 6 [Source:HGNC Symbol;Acc:HGNC:33961] | -1.4336 | 0.00703 |
| ENSG00000274605 | PCCA-DT | PCCA divergent transcript [Source:HGNC Symbol;Acc:HGNC:53266] | 0.6225 | 0.00712 |
| ENSG00000197279 | ZNF165 | zinc finger protein 165 [Source:HGNC Symbol;Acc:HGNC:12953] | 0.6594 | 0.00714 |
| ENSG00000258365 | NA | novel transcript, antisense to PLXNC1 | 1.4508 | 0.00715 |
| ENSG00000277969 | NA | novel transcript | -0.5077 | 0.00717 |
| ENSG00000267080 | ASB16-AS1 | ASB16 antisense RNA 1 [Source:HGNC Symbol;Acc:HGNC:25442] | -0.5531 | 0.00719 |
| ENSG00000122483 | CCDC18 | coiled-coil domain containing 18 [Source:HGNC Symbol;Acc:HGNC:30370] | -0.6213 | 0.00721 |
| ENSG00000281912 | LINC01144 | long intergenic non-protein coding RNA 1144 [Source:HGNC Symbol;Acc:HGNC:49460] | -0.7182 | 0.00721 |
| ENSG00000261147 | NA | novel protein | 2.0905 | 0.00723 |
| ENSG00000132016 | C19orf57 | chromosome 19 open reading frame 57 [Source:HGNC Symbol;Acc:HGNC:28153] | -0.7136 | 0.00726 |
| ENSG00000088002 | SULT2B1 | sulfotransferase family 2B member 1 [Source:HGNC Symbol;Acc:HGNC:11459] | -1.1686 | 0.00727 |
| ENSG00000168071 | CCDC88B | coiled-coil domain containing 88B [Source:HGNC Symbol;Acc:HGNC:26757] | -0.6222 | 0.00727 |
| ENSG00000283294 | NA | novel transcript | -1.3944 | 0.00728 |
| ENSG00000092529 | CAPN3 | calpain 3 [Source:HGNC Symbol;Acc:HGNC:1480] | -0.6509 | 0.00729 |
| ENSG00000162643 | WDR63 | WD repeat domain 63 [Source:HGNC Symbol;Acc:HGNC:30711] | -1.3059 | 0.00732 |
| ENSG00000212864 | RNF208 | ring finger protein 208 [Source:HGNC Symbol;Acc:HGNC:25420] | -0.5984 | 0.00739 |
| ENSG00000188013 | MEIS3P2 | Meis homeobox 3 pseudogene 2 [Source:HGNC Symbol;Acc:HGNC:17638] | 0.9638 | 0.00741 |
| ENSG00000121797 | CCRL2 | C-C motif chemokine receptor like 2 [Source:HGNC Symbol;Acc:HGNC:1612] | 2.0884 | 0.00741 |
| ENSG00000160602 | NEK8 | NIMA related kinase 8 [Source:HGNC Symbol;Acc:HGNC:13387] | -0.8047 | 0.00741 |
| ENSG00000166689 | PLEKHA7 | pleckstrin homology domain containing A7 [Source:HGNC Symbol;Acc:HGNC:27049] | -0.9465 | 0.00743 |
| ENSG00000279875 | NA | TEC | 1.2973 | 0.00743 |
| ENSG00000142197 | DOP1B | DOP1 leucine zipper like protein B [Source:HGNC Symbol;Acc:HGNC:1291] | 0.7209 | 0.00745 |
| ENSG00000102678 | FGF9 | fibroblast growth factor 9 [Source:HGNC Symbol;Acc:HGNC:3687] | -0.6517 | 0.00746 |
| ENSG00000160471 | COX6B2 | cytochrome c oxidase subunit 6B2 [Source:HGNC Symbol;Acc:HGNC:24380] | -1.4948 | 0.00746 |
| ENSG00000224116 | INHBA-AS1 | INHBA antisense RNA 1 [Source:HGNC Symbol;Acc:HGNC:40303] | 2.9282 | 0.00746 |
| ENSG00000268486 | NA | ankyrin repeat domain 30B (ANKRD30B) pseudogene | -1.5036 | 0.00747 |
| ENSG00000141469 | SLC14A1 | solute carrier family 14 member 1 (Kidd blood group) [Source:HGNC Symbol;Acc:HGNC:10918] | 2.3634 | 0.00747 |
| ENSG00000244119 | PDCL3P4 | PDCL3 pseudogene 4 [Source:HGNC Symbol;Acc:HGNC:44505] | -0.8323 | 0.00747 |
| ENSG00000168676 | KCTD19 | potassium channel tetramerization domain containing 19 [Source:HGNC Symbol;Acc:HGNC:24753] | -0.9203 | 0.00747 |
| ENSG00000235910 | APOA1-AS | APOA1 antisense RNA [Source:HGNC Symbol;Acc:HGNC:40079] | -1.5237 | 0.00747 |
| ENSG00000121897 | LIAS | lipoic acid synthetase [Source:HGNC Symbol;Acc:HGNC:16429] | -0.5630 | 0.00749 |
| ENSG00000227906 | SNAP25-AS1 | SNAP25 antisense RNA 1 [Source:HGNC Symbol;Acc:HGNC:44312] | -1.0111 | 0.00752 |
| ENSG00000142149 | HUNK | hormonally up-regulated Neu-associated kinase [Source:HGNC Symbol;Acc:HGNC:13326] | 1.5576 | 0.00753 |
| ENSG00000224429 | LINC00539 | long intergenic non-protein coding RNA 539 [Source:HGNC Symbol;Acc:HGNC:43672] | -0.9606 | 0.00757 |

| | | | | |
|---|---|---|---|---|
| ENSG00000278376 | NA | novel transcript | -0.5975 | 0.00766 |
| ENSG00000173706 | HEG1 | heart development protein with EGF like domains 1 [Source:HGNC Symbol;Acc:HGNC:29227] | -0.6896 | 0.00767 |
| ENSG00000111834 | RSPH4A | radial spoke head component 4A [Source:HGNC Symbol;Acc:HGNC:21558] | 0.8012 | 0.00769 |
| ENSG00000164404 | GDF9 | growth differentiation factor 9 [Source:HGNC Symbol;Acc:HGNC:4224] | -0.9516 | 0.00771 |
| ENSG00000235381 | NA | novel transcript | 0.6434 | 0.00773 |
| ENSG00000260190 | NA | novel transcript, overlapping FBXW5 | 1.4321 | 0.00774 |
| ENSG00000163116 | STPG2 | sperm tail PG-rich repeat containing 2 [Source:HGNC Symbol;Acc:HGNC:28712] | -0.9913 | 0.00777 |
| ENSG00000272256 | NA | novel transcript, antisense to TMEM66 | 1.1042 | 0.00777 |
| ENSG00000146001 | PCDHB18P | protocadherin beta 18 pseudogene [Source:HGNC Symbol;Acc:HGNC:14548] | -0.7884 | 0.00781 |
| ENSG00000198865 | CCDC152 | coiled-coil domain containing 152 [Source:HGNC Symbol;Acc:HGNC:34438] | -0.6023 | 0.00788 |
| ENSG00000279673 | NA | TEC | -0.9634 | 0.00788 |
| ENSG00000159263 | SIM2 | SIM bHLH transcription factor 2 [Source:HGNC Symbol;Acc:HGNC:10883] | -1.7675 | 0.00788 |
| ENSG00000242104 | LINC02618 | long intergenic non-protein coding RNA 2618 [Source:HGNC Symbol;Acc:HGNC:54091] | -2.0528 | 0.00794 |
| ENSG00000174236 | REP15 | RAB15 effector protein [Source:HGNC Symbol;Acc:HGNC:33748] | 1.6308 | 0.00794 |
| ENSG00000197302 | ZNF720 | zinc finger protein 720 [Source:HGNC Symbol;Acc:HGNC:26987] | 0.5325 | 0.00795 |
| ENSG00000105855 | ITGB8 | integrin subunit beta 8 [Source:HGNC Symbol;Acc:HGNC:6163] | 1.4077 | 0.00796 |
| ENSG00000278662 | GOLGA6L10 | golgin A6 family like 10 [Source:HGNC Symbol;Acc:HGNC:37228] | -1.3984 | 0.00798 |
| ENSG00000216906 | NA | multimerin 2 (MMRN2) pseudogene | -1.4930 | 0.00801 |
| ENSG00000173809 | TDRD12 | tudor domain containing 12 [Source:HGNC Symbol;Acc:HGNC:25044] | -1.3357 | 0.00802 |
| ENSG00000164093 | PITX2 | paired like homeodomain 2 [Source:HGNC Symbol;Acc:HGNC:9005] | 1.1823 | 0.00804 |
| ENSG00000110719 | TCIRG1 | T cell immune regulator 1, ATPase H+ transporting V0 subunit a3 [Source:HGNC Symbol;Acc:HGNC:11647] | 0.8794 | 0.00812 |
| ENSG00000119227 | PIGZ | phosphatidylinositol glycan anchor biosynthesis class Z [Source:HGNC Symbol;Acc:HGNC:30596] | -0.9860 | 0.00813 |
| ENSG00000251537 | NA | novel tripartite motif-containing 16 (TRIM16) and CMT1A duplicated region transcript 1 (CDRT1) protein | 0.9119 | 0.00815 |
| ENSG00000159164 | SV2A | synaptic vesicle glycoprotein 2A [Source:HGNC Symbol;Acc:HGNC:20566] | 1.0087 | 0.00816 |
| ENSG00000280164 | NA | novel transcript | -1.3772 | 0.00817 |
| ENSG00000253686 | LINC01484 | long intergenic non-protein coding RNA 1484 [Source:HGNC Symbol;Acc:HGNC:51136] | 2.6505 | 0.00826 |
| ENSG00000152457 | DCLRE1C | DNA cross-link repair 1C [Source:HGNC Symbol;Acc:HGNC:17642] | 0.5685 | 0.00827 |
| ENSG00000260404 | NA | general transcription factor IIi (GTF2I) pseudogene | 0.5329 | 0.00828 |
| ENSG00000286231 | NA | novel protein | -1.4860 | 0.00832 |
| ENSG00000186451 | SPATA12 | spermatogenesis associated 12 [Source:HGNC Symbol;Acc:HGNC:23221] | 2.6273 | 0.00834 |
| ENSG00000221963 | APOL6 | apolipoprotein L6 [Source:HGNC Symbol;Acc:HGNC:14870] | -0.7376 | 0.00841 |
| ENSG00000158201 | ABHD3 | abhydrolase domain containing 3, phospholipase [Source:HGNC Symbol;Acc:HGNC:18718] | 0.5217 | 0.00841 |
| ENSG00000260806 | NA | novel transcript, antisense to PAPOLA | 0.8508 | 0.00851 |
| ENSG00000286883 | NA | novel transcript, antisense to CCDC142and MRPL53and MOGS | 0.7257 | 0.00853 |
| ENSG00000035115 | SH3YL1 | SH3 and SYLF domain containing 1 [Source:HGNC Symbol;Acc:HGNC:29546] | -0.6527 | 0.00855 |
| ENSG00000125246 | CLYBL | citramalyl-CoA lyase [Source:HGNC Symbol;Acc:HGNC:18355] | -0.5400 | 0.00855 |
| ENSG00000138101 | DTNB | dystrobrevin beta [Source:HGNC Symbol;Acc:HGNC:3058] | -0.6134 | 0.00856 |
| ENSG00000159917 | ZNF235 | zinc finger protein 235 [Source:HGNC Symbol;Acc:HGNC:12866] | -0.7255 | 0.00859 |
| ENSG00000127528 | KLF2 | Kruppel like factor 2 [Source:HGNC Symbol;Acc:HGNC:6347] | 0.7449 | 0.00860 |
| ENSG00000227517 | LINC01483 | long intergenic non-protein coding RNA 1483 [Source:HGNC Symbol;Acc:HGNC:51130] | 2.4549 | 0.00866 |
| ENSG00000243694 | LINC02027 | long intergenic non-protein coding RNA 2027 [Source:HGNC Symbol;Acc:HGNC:52862] | -1.5748 | 0.00866 |
| ENSG00000253730 | NA | novel transcript | -2.4333 | 0.00868 |
| ENSG00000259803 | SLC22A31 | solute carrier family 22 member 31 [Source:HGNC Symbol;Acc:HGNC:27091] | 1.9221 | 0.00869 |

| | | | | |
|---|---|---|---|---|
| ENSG00000272711 | NA | novel transcript | 1.1889 | 0.00869 |
| ENSG00000234776 | C11orf94 | chromosome 11 open reading frame 94 [Source:HGNC Symbol;Acc:HGNC:37213] | 0.8266 | 0.00871 |
| ENSG00000249274 | PDLIM1P4 | PDZ and LIM domain 1 pseudogene 4 [Source:HGNC Symbol;Acc:HGNC:48947] | -1.9715 | 0.00874 |
| ENSG00000178828 | RNF186 | ring finger protein 186 [Source:HGNC Symbol;Acc:HGNC:25978] | -0.8301 | 0.00875 |
| ENSG00000262209 | PCDHGB3 | protocadherin gamma subfamily B, 3 [Source:HGNC Symbol;Acc:HGNC:8710] | -0.7140 | 0.00875 |
| ENSG00000145476 | CYP4V2 | cytochrome P450 family 4 subfamily V member 2 [Source:HGNC Symbol;Acc:HGNC:23198] | -0.5671 | 0.00876 |
| ENSG00000245322 | H2AZ1-DT | H2AZ1 divergent transcript [Source:HGNC Symbol;Acc:HGNC:40271] | -0.9456 | 0.00877 |
| ENSG00000183250 | LINC01547 | long intergenic non-protein coding RNA 1547 [Source:HGNC Symbol;Acc:HGNC:15707] | -0.5246 | 0.00877 |
| ENSG00000138650 | PCDH10 | protocadherin 10 [Source:HGNC Symbol;Acc:HGNC:13404] | -1.6647 | 0.00881 |
| ENSG00000223701 | RAET1E-AS1 | RAET1E antisense RNA 1 [Source:HGNC Symbol;Acc:HGNC:48994] | -2.5023 | 0.00882 |
| ENSG00000018236 | CNTN1 | contactin 1 [Source:HGNC Symbol;Acc:HGNC:2171] | 0.6703 | 0.00883 |
| ENSG00000177455 | CD19 | CD19 molecule [Source:HGNC Symbol;Acc:HGNC:1633] | -0.7562 | 0.00884 |
| ENSG00000283761 | NA | novel protein | 1.4680 | 0.00886 |
| ENSG00000178403 | NEUROG2 | neurogenin 2 [Source:HGNC Symbol;Acc:HGNC:13805] | 0.9954 | 0.00890 |
| ENSG00000268297 | CLEC4GP1 | C-type lectin domain family 4 member G pseudogene 1 [Source:HGNC Symbol;Acc:HGNC:33502] | -4.2453 | 0.00890 |
| ENSG00000176826 | FKBP9P1 | FKBP prolyl isomerase 9 pseudogene 1 [Source:HGNC Symbol;Acc:HGNC:23568] | -1.4568 | 0.00890 |
| ENSG00000184343 | SRPK3 | SRSF protein kinase 3 [Source:HGNC Symbol;Acc:HGNC:11402] | 1.1384 | 0.00891 |
| ENSG00000261963 | CCDC92B | coiled-coil domain containing 92B [Source:HGNC Symbol;Acc:HGNC:52279] | -1.8845 | 0.00891 |
| ENSG00000286010 | NA | novel transcript | 0.5070 | 0.00892 |
| ENSG00000256250 | NA | novel transcript | 2.5298 | 0.00893 |
| ENSG00000226057 | PHF2P2 | PHD finger protein 2 pseudogene 2 [Source:HGNC Symbol;Acc:HGNC:38808] | -0.6893 | 0.00893 |
| ENSG00000250451 | HOXC-AS1 | HOXC cluster antisense RNA 1 [Source:HGNC Symbol;Acc:HGNC:43749] | -2.2614 | 0.00895 |
| ENSG00000139263 | LRIG3 | leucine rich repeats and immunoglobulin like domains 3 [Source:HGNC Symbol;Acc:HGNC:30991] | -0.6373 | 0.00903 |
| ENSG00000257842 | LINC02588 | long intergenic non-protein coding RNA 2588 [Source:HGNC Symbol;Acc:HGNC:19827] | -0.9554 | 0.00907 |
| ENSG00000106366 | SERPINE1 | serpin family E member 1 [Source:HGNC Symbol;Acc:HGNC:8583] | 1.9685 | 0.00916 |
| ENSG00000187017 | ESPN | espin [Source:HGNC Symbol;Acc:HGNC:13281] | -4.4480 | 0.00916 |
| ENSG00000287574 | NA | novel transcript | -0.5526 | 0.00921 |
| ENSG00000235859 | NA | proteasome (prosome, macropain) 26S subunit, ATPase, 5 (PSMC5) pseudogene | -0.9502 | 0.00922 |
| ENSG00000070601 | FRMPD1 | FERM and PDZ domain containing 1 [Source:HGNC Symbol;Acc:HGNC:29159] | -2.1608 | 0.00923 |
| ENSG00000287816 | NA | novel transcript | -1.5757 | 0.00923 |
| ENSG00000061337 | LZTS1 | leucine zipper tumor suppressor 1 [Source:HGNC Symbol;Acc:HGNC:13861] | -0.7803 | 0.00927 |
| ENSG00000250049 | NA | novel transcript | -1.1394 | 0.00928 |
| ENSG00000135185 | TMEM243 | transmembrane protein 243 [Source:HGNC Symbol;Acc:HGNC:21707] | 0.5517 | 0.00931 |
| ENSG00000188070 | C11orf95 | chromosome 11 open reading frame 95 [Source:HGNC Symbol;Acc:HGNC:28449] | -0.5697 | 0.00934 |
| ENSG00000166664 | CHRFAM7A | CHRNA7 (exons 5-10) and FAM7A (exons A-E) fusion [Source:HGNC Symbol;Acc:HGNC:15781] | 2.2366 | 0.00938 |
| ENSG00000288095 | NA | novel transcript | -1.8335 | 0.00940 |
| ENSG00000163923 | RPL39L | ribosomal protein L39 like [Source:HGNC Symbol;Acc:HGNC:17094] | -0.6287 | 0.00940 |
| ENSG00000196593 | ANKRD20A19P | ankyrin repeat domain 20 family member A19, pseudogene [Source:HGNC Symbol;Acc:HGNC:42737] | -3.0948 | 0.00944 |
| ENSG00000272008 | NA | novel transcript, antisense to ZNF292 | -1.0170 | 0.00944 |
| ENSG00000273840 | NA | novel transcript | 0.6542 | 0.00944 |
| ENSG00000161692 | DBF4B | DBF4 zinc finger B [Source:HGNC Symbol;Acc:HGNC:17883] | 0.5487 | 0.00946 |
| ENSG00000225265 | TAF1A-AS1 | TAF1A antisense RNA 1 [Source:HGNC Symbol;Acc:HGNC:40573] | -0.7791 | 0.00947 |
| ENSG00000158122 | PRXL2C | peroxiredoxin like 2C [Source:HGNC Symbol;Acc:HGNC:16881] | 0.5811 | 0.00950 |

| | | | | |
|---|---|---|---|---|
| ENSG00000174501 | ANKRD36C | ankyrin repeat domain 36C [Source:HGNC Symbol;Acc:HGNC:32946] | -0.7355 | 0.00950 |
| ENSG00000254402 | LRRC24 | leucine rich repeat containing 24 [Source:HGNC Symbol;Acc:HGNC:28947] | -0.5668 | 0.00950 |
| ENSG00000237310 | NA | uncharacterized LOC100289098 [Source:NCBI gene (formerly Entrezgene);Acc:100289098] | -0.7065 | 0.00951 |
| ENSG00000184068 | SREBF2-AS1 | SREBF2 antisense RNA 1 [Source:HGNC Symbol;Acc:HGNC:53953] | -0.5405 | 0.00951 |
| ENSG00000160298 | C21orf58 | chromosome 21 open reading frame 58 [Source:HGNC Symbol;Acc:HGNC:1300] | -0.5905 | 0.00952 |
| ENSG00000196951 | SCOC-AS1 | SCOC antisense RNA 1 [Source:HGNC Symbol;Acc:HGNC:50601] | -0.7623 | 0.00957 |
| ENSG00000105613 | MAST1 | microtubule associated serine/threonine kinase 1 [Source:HGNC Symbol;Acc:HGNC:19034] | -0.5299 | 0.00959 |
| ENSG00000276166 | NA | novel transcript, antisense to KIFC3 | 1.6200 | 0.00962 |
| ENSG00000152689 | RASGRP3 | RAS guanyl releasing protein 3 [Source:HGNC Symbol;Acc:HGNC:14545] | 1.7422 | 0.00963 |
| ENSG00000142733 | MAP3K6 | mitogen-activated protein kinase kinase kinase 6 [Source:HGNC Symbol;Acc:HGNC:6858] | 0.6779 | 0.00968 |
| ENSG00000275457 | NA | novel transcript, antisense to XRN2 | 0.7271 | 0.00970 |
| ENSG00000261634 | NA | novel transcript, antisense to PAQR5 | -0.8442 | 0.00971 |
| ENSG00000160999 | SH2B2 | SH2B adaptor protein 2 [Source:HGNC Symbol;Acc:HGNC:17381] | 1.0321 | 0.00971 |
| ENSG00000113600 | C9 | complement C9 [Source:HGNC Symbol;Acc:HGNC:1358] | 0.9829 | 0.00973 |
| ENSG00000154240 | CEP112 | centrosomal protein 112 [Source:HGNC Symbol;Acc:HGNC:28514] | -0.5187 | 0.00973 |
| ENSG00000196337 | CGB7 | chorionic gonadotropin subunit beta 7 [Source:HGNC Symbol;Acc:HGNC:16451] | -0.9752 | 0.00974 |
| ENSG00000238276 | NA | uncharacterized LOC101927760 [Source:NCBI gene (formerly Entrezgene);Acc:101927760] | -0.9596 | 0.00982 |
| ENSG00000261604 | NA | novel transcript, antisense to HMGCS1 | -1.1902 | 0.00985 |
| ENSG00000066923 | STAG3 | stromal antigen 3 [Source:HGNC Symbol;Acc:HGNC:11356] | -0.5621 | 0.00990 |
| ENSG00000197415 | VEPH1 | ventricular zone expressed PH domain containing 1 [Source:HGNC Symbol;Acc:HGNC:25735] | -1.7119 | 0.00991 |
| ENSG00000244097 | RPS4XP17 | ribosomal protein S4X pseudogene 17 [Source:HGNC Symbol;Acc:HGNC:36789] | 2.1918 | 0.00994 |
| ENSG00000078900 | TP73 | tumor protein p73 [Source:HGNC Symbol;Acc:HGNC:12003] | -0.5287 | 0.00997 |
| ENSG00000058453 | CROCC | ciliary rootlet coiled-coil, rootletin [Source:HGNC Symbol;Acc:HGNC:21299] | -0.5033 | 0.00998 |
| ENSG00000236829 | NA | ribosomal protein L23a (RPL23A) pseudogene | -1.5174 | 0.01001 |
| ENSG00000123154 | WDR83 | WD repeat domain 83 [Source:HGNC Symbol;Acc:HGNC:32672] | 0.5416 | 0.01007 |
| ENSG00000160447 | PKN3 | protein kinase N3 [Source:HGNC Symbol;Acc:HGNC:17999] | -0.5622 | 0.01008 |
| ENSG00000249709 | ZNF564 | zinc finger protein 564 [Source:HGNC Symbol;Acc:HGNC:31106] | -0.6186 | 0.01010 |
| ENSG00000168961 | LGALS9 | galectin 9 [Source:HGNC Symbol;Acc:HGNC:6570] | -0.8058 | 0.01012 |
| ENSG00000149575 | SCN2B | sodium voltage-gated channel beta subunit 2 [Source:HGNC Symbol;Acc:HGNC:10589] | -2.7526 | 0.01014 |
| ENSG00000280341 | NA | novel transcript | -0.8151 | 0.01016 |
| ENSG00000116016 | EPAS1 | endothelial PAS domain protein 1 [Source:HGNC Symbol;Acc:HGNC:3374] | 0.8804 | 0.01016 |
| ENSG00000198673 | TAFA2 | TAFA chemokine like family member 2 [Source:HGNC Symbol;Acc:HGNC:21589] | 0.9555 | 0.01020 |
| ENSG00000267493 | CIRBP-AS1 | CIRBP antisense RNA 1 [Source:HGNC Symbol;Acc:HGNC:28588] | 0.9357 | 0.01021 |
| ENSG00000221874 | ZNF816-ZNF321P | ZNF816-ZNF321P readthrough [Source:HGNC Symbol;Acc:HGNC:38879] | 5.5960 | 0.01033 |
| ENSG00000255150 | EID3 | EP300 interacting inhibitor of differentiation 3 [Source:HGNC Symbol;Acc:HGNC:32961] | 1.4487 | 0.01035 |
| ENSG00000101412 | E2F1 | E2F transcription factor 1 [Source:HGNC Symbol;Acc:HGNC:3113] | -0.5488 | 0.01039 |
| ENSG00000104881 | PPP1R13L | protein phosphatase 1 regulatory subunit 13 like [Source:HGNC Symbol;Acc:HGNC:18838] | 0.5318 | 0.01041 |
| ENSG00000165828 | PRAP1 | proline rich acidic protein 1 [Source:HGNC Symbol;Acc:HGNC:23304] | 2.0858 | 0.01041 |
| ENSG00000271551 | NA | novel transcript, antisense to ARID1B | -0.8971 | 0.01050 |
| ENSG00000005513 | SOX8 | SRY-box transcription factor 8 [Source:HGNC Symbol;Acc:HGNC:11203] | 0.5084 | 0.01054 |
| ENSG00000266885 | NA | chromosome 17 open reading frame 51 (C17orf51) pseudogene | -1.2079 | 0.01056 |
| ENSG00000148204 | CRB2 | crumbs cell polarity complex component 2 [Source:HGNC Symbol;Acc:HGNC:18688] | 0.6635 | 0.01062 |
| ENSG00000271826 | PLS3-AS1 | PLS3 antisense RNA 1 [Source:HGNC Symbol;Acc:HGNC:50343] | -0.8538 | 0.01062 |

| | | | | |
|---|---|---|---|---|
| ENSG00000121207 | LRAT | lecithin retinol acyltransferase [Source:HGNC Symbol;Acc:HGNC:6685] | -1.5802 | 0.01065 |
| ENSG00000162572 | SCNN1D | sodium channel epithelial 1 subunit delta [Source:HGNC Symbol;Acc:HGNC:10601] | -0.8975 | 0.01067 |
| ENSG00000261675 | NA | novel transcript | -1.0287 | 0.01075 |
| ENSG00000286136 | NA | novel transcript | 1.6047 | 0.01077 |
| ENSG00000165323 | FAT3 | FAT atypical cadherin 3 [Source:HGNC Symbol;Acc:HGNC:23112] | -1.7148 | 0.01077 |
| ENSG00000286257 | NA | novel transcript | 0.5728 | 0.01083 |
| ENSG00000167748 | KLK1 | kallikrein 1 [Source:HGNC Symbol;Acc:HGNC:6357] | -1.0659 | 0.01083 |
| ENSG00000228232 | GAPDHP1 | glyceraldehyde-3-phosphate dehydrogenase pseudogene 1 [Source:HGNC Symbol;Acc:HGNC:4159] | 0.5334 | 0.01100 |
| ENSG00000228624 | HDAC2-AS2 | HDAC2 and HS3ST5 antisense RNA 2 [Source:HGNC Symbol;Acc:HGNC:43590] | -0.9598 | 0.01101 |
| ENSG00000133985 | TTC9 | tetratricopeptide repeat domain 9 [Source:HGNC Symbol;Acc:HGNC:20267] | -0.8859 | 0.01103 |
| ENSG00000171812 | COL8A2 | collagen type VIII alpha 2 chain [Source:HGNC Symbol;Acc:HGNC:2216] | 0.9309 | 0.01104 |
| ENSG00000233672 | RNASEH2B-AS1 | RNASEH2B antisense RNA 1 [Source:HGNC Symbol;Acc:HGNC:39967] | -1.5507 | 0.01105 |
| ENSG00000284954 | NA | novel transcript | -1.0195 | 0.01106 |
| ENSG00000182771 | GRID1 | glutamate ionotropic receptor delta type subunit 1 [Source:HGNC Symbol;Acc:HGNC:4575] | 1.0603 | 0.01109 |
| ENSG00000188747 | NOXA1 | NADPH oxidase activator 1 [Source:HGNC Symbol;Acc:HGNC:10668] | -0.8984 | 0.01117 |
| ENSG00000231703 | NA | novel transcript, antisense to TSHZ2 | 2.4108 | 0.01117 |
| ENSG00000162433 | AK4 | adenylate kinase 4 [Source:HGNC Symbol;Acc:HGNC:363] | 1.1768 | 0.01126 |
| ENSG00000043039 | BARX2 | BARX homeobox 2 [Source:HGNC Symbol;Acc:HGNC:956] | -2.3875 | 0.01127 |
| ENSG00000081019 | RSBN1 | round spermatid basic protein 1 [Source:HGNC Symbol;Acc:HGNC:25642] | 0.5065 | 0.01127 |
| ENSG00000100346 | CACNA1I | calcium voltage-gated channel subunit alpha1 I [Source:HGNC Symbol;Acc:HGNC:1396] | -0.9758 | 0.01131 |
| ENSG00000283757 | NA | novel transcript | -1.2675 | 0.01138 |
| ENSG00000152154 | TMEM178A | transmembrane protein 178A [Source:HGNC Symbol;Acc:HGNC:28517] | -1.2873 | 0.01141 |
| ENSG00000205978 | NYNRIN | NYN domain and retroviral integrase containing [Source:HGNC Symbol;Acc:HGNC:20165] | -0.5864 | 0.01144 |
| ENSG00000280177 | NA | TEC | -2.3025 | 0.01145 |
| ENSG00000273137 | NA | novel transcript, antisense to SELO | 1.5311 | 0.01149 |
| ENSG00000164379 | FOXQ1 | forkhead box Q1 [Source:HGNC Symbol;Acc:HGNC:20951] | 1.5223 | 0.01149 |
| ENSG00000091831 | ESR1 | estrogen receptor 1 [Source:HGNC Symbol;Acc:HGNC:3467] | 0.6044 | 0.01152 |
| ENSG00000260265 | LINC02562 | long intergenic non-protein coding RNA 2562 [Source:HGNC Symbol;Acc:HGNC:53602] | -2.7727 | 0.01152 |
| ENSG00000251161 | NA | novel transcript | 1.1455 | 0.01154 |
| ENSG00000226445 | NA | novel transcript | 0.5787 | 0.01158 |
| ENSG00000249962 | NA | pantothenate kinase 1 (PANK1) pseudogene | -0.9925 | 0.01159 |
| ENSG00000187527 | ATP13A5 | ATPase 13A5 [Source:HGNC Symbol;Acc:HGNC:31789] | -1.5861 | 0.01160 |
| ENSG00000104341 | LAPTM4B | lysosomal protein transmembrane 4 beta [Source:HGNC Symbol;Acc:HGNC:13646] | 0.7319 | 0.01162 |
| ENSG00000249948 | GBA3 | glucosylceramidase beta 3 (gene/pseudogene) [Source:HGNC Symbol;Acc:HGNC:19069] | -1.2661 | 0.01162 |
| ENSG00000132938 | MTUS2 | microtubule associated scaffold protein 2 [Source:HGNC Symbol;Acc:HGNC:20595] | 0.8296 | 0.01166 |
| ENSG00000244067 | GSTA2 | glutathione S-transferase alpha 2 [Source:HGNC Symbol;Acc:HGNC:4627] | -1.0991 | 0.01169 |
| ENSG00000213445 | SIPA1 | signal-induced proliferation-associated 1 [Source:HGNC Symbol;Acc:HGNC:10885] | -0.5058 | 0.01172 |
| ENSG00000242294 | STAG3L5P | stromal antigen 3-like 5 pseudogene [Source:HGNC Symbol;Acc:HGNC:48896] | 0.6632 | 0.01174 |
| ENSG00000082196 | C1QTNF3 | C1q and TNF related 3 [Source:HGNC Symbol;Acc:HGNC:14326] | -0.6938 | 0.01176 |
| ENSG00000069399 | BCL3 | BCL3 transcription coactivator [Source:HGNC Symbol;Acc:HGNC:998] | 0.6651 | 0.01177 |
| ENSG00000128606 | LRRC17 | leucine rich repeat containing 17 [Source:HGNC Symbol;Acc:HGNC:16895] | -0.7947 | 0.01182 |
| ENSG00000186026 | ZNF284 | zinc finger protein 284 [Source:HGNC Symbol;Acc:HGNC:13078] | -0.5857 | 0.01187 |
| ENSG00000111799 | COL12A1 | collagen type XII alpha 1 chain [Source:HGNC Symbol;Acc:HGNC:2188] | 1.6063 | 0.01200 |

| | | | | |
|---|---|---|---|---|
| ENSG00000197536 | IRF1-AS1 | IRF1 antisense RNA 1 [Source:HGNC Symbol;Acc:HGNC:33838] | -0.5552 | 0.01202 |
| ENSG00000255874 | PRECSIT | p53 regulated carcinoma associated Stat3 activating long intergenic non-protein coding transcript [Source:HGNC Symbol;Acc:HGNC:27492] | 1.4684 | 0.01202 |
| ENSG00000287811 | NA | novel transcript | 1.2599 | 0.01202 |
| ENSG00000234660 | LINC00440 | long intergenic non-protein coding RNA 440 [Source:HGNC Symbol;Acc:HGNC:42777] | -0.6967 | 0.01207 |
| ENSG00000273045 | C2orf15 | chromosome 2 open reading frame 15 [Source:HGNC Symbol;Acc:HGNC:28436] | 0.5152 | 0.01208 |
| ENSG00000242574 | HLA-DMB | major histocompatibility complex, class II, DM beta [Source:HGNC Symbol;Acc:HGNC:4935] | -1.6902 | 0.01208 |
| ENSG00000160712 | IL6R | interleukin 6 receptor [Source:HGNC Symbol;Acc:HGNC:6019] | 2.1230 | 0.01212 |
| ENSG00000111186 | WNT5B | Wnt family member 5B [Source:HGNC Symbol;Acc:HGNC:16265] | 1.1061 | 0.01215 |
| ENSG00000073711 | PPP2R3A | protein phosphatase 2 regulatory subunit B''alpha [Source:HGNC Symbol;Acc:HGNC:9307] | -0.5287 | 0.01216 |
| ENSG00000260588 | NA | novel transcript | -1.9839 | 0.01216 |
| ENSG00000186148 | NA | two-pore calcium channel 3 (TPCN3) pseudogene | 1.5127 | 0.01217 |
| ENSG00000189143 | CLDN4 | claudin 4 [Source:HGNC Symbol;Acc:HGNC:2046] | -0.6727 | 0.01218 |
| ENSG00000287861 | NA | novel transcript | -0.5791 | 0.01225 |
| ENSG00000286996 | NA | novel transcript | 1.3572 | 0.01226 |
| ENSG00000250563 | KNOP1P5 | lysine rich nucleolar protein 1 pseudogene 5 [Source:HGNC Symbol;Acc:HGNC:48923] | -0.9045 | 0.01228 |
| ENSG00000236425 | NA | ribosomal protein L31 (RPL31) pseudogene | 1.8979 | 0.01229 |
| ENSG00000224057 | EGFR-AS1 | EGFR antisense RNA 1 [Source:HGNC Symbol;Acc:HGNC:40207] | 6.3316 | 0.01230 |
| ENSG00000260763 | NA | novel transcript | -0.9411 | 0.01232 |
| ENSG00000232320 | MXRA7P1 | MXRA7 pseudogene 1 [Source:HGNC Symbol;Acc:HGNC:54643] | 1.6997 | 0.01237 |
| ENSG00000189157 | FAM47E | family with sequence similarity 47 member E [Source:HGNC Symbol;Acc:HGNC:34343] | -1.3255 | 0.01243 |
| ENSG00000111325 | OGFOD2 | 2-oxoglutarate and iron dependent oxygenase domain containing 2 [Source:HGNC Symbol;Acc:HGNC:25823] | -0.5224 | 0.01243 |
| ENSG00000140025 | EFCAB11 | EF-hand calcium binding domain 11 [Source:HGNC Symbol;Acc:HGNC:20357] | -0.6758 | 0.01244 |
| ENSG00000133794 | ARNTL | aryl hydrocarbon receptor nuclear translocator like [Source:HGNC Symbol;Acc:HGNC:701] | 0.5472 | 0.01250 |
| ENSG00000285999 | NA | novel transcipt | 1.0702 | 0.01250 |
| ENSG00000258317 | NA | novel transcript, antisense to ZC3H10 and ESYT1 | 1.1074 | 0.01251 |
| ENSG00000079156 | OSBPL6 | oxysterol binding protein like 6 [Source:HGNC Symbol;Acc:HGNC:16388] | -0.6052 | 0.01251 |
| ENSG00000124523 | SIRT5 | sirtuin 5 [Source:HGNC Symbol;Acc:HGNC:14933] | -0.6927 | 0.01261 |
| ENSG00000250222 | NA | novel transcript | -1.1020 | 0.01261 |
| ENSG00000101442 | ACTR5 | actin related protein 5 [Source:HGNC Symbol;Acc:HGNC:14671] | -0.5829 | 0.01262 |
| ENSG00000272434 | NA | novel transcript, antisense to IMPDH2 and QRICH1 | 0.9521 | 0.01263 |
| ENSG00000260293 | NA | novel transcript, intronic to TBC1D24 | -1.0238 | 0.01272 |
| ENSG00000163958 | ZDHHC19 | zinc finger DHHC-type palmitoyltransferase 19 [Source:HGNC Symbol;Acc:HGNC:20713] | -0.5395 | 0.01275 |
| ENSG00000259291 | ZNF710-AS1 | ZNF710 antisense RNA 1 [Source:HGNC Symbol;Acc:HGNC:53141] | -0.9391 | 0.01285 |
| ENSG00000164089 | ETNPPL | ethanolamine-phosphate phospho-lyase [Source:HGNC Symbol;Acc:HGNC:14404] | 0.6096 | 0.01286 |
| ENSG00000258754 | LINC01579 | long intergenic non-protein coding RNA 1579 [Source:HGNC Symbol;Acc:HGNC:27519] | 1.1963 | 0.01290 |
| ENSG00000233175 | NA | novel transcript, antisense to FMNL1 | 0.7799 | 0.01290 |
| ENSG00000279655 | NA | TEC | 2.0453 | 0.01290 |
| ENSG00000229404 | LINC00858 | long intergenic non-protein coding RNA 858 [Source:HGNC Symbol;Acc:HGNC:27276] | -0.7878 | 0.01293 |
| ENSG00000049247 | UTS2 | urotensin 2 [Source:HGNC Symbol;Acc:HGNC:12636] | 2.9486 | 0.01293 |
| ENSG00000135476 | ESPL1 | extra spindle pole bodies like 1, separase [Source:HGNC Symbol;Acc:HGNC:16856] | -0.5612 | 0.01295 |
| ENSG00000213781 | PSMC1P2 | proteasome 26S subunit, ATPase 1 pseudogene 2 [Source:HGNC Symbol;Acc:HGNC:39777] | 2.1218 | 0.01299 |
| ENSG00000164270 | HTR4 | 5-hydroxytryptamine receptor 4 [Source:HGNC Symbol;Acc:HGNC:5299] | 2.8636 | 0.01303 |

| | | | | |
|---|---|---|---|---|
| ENSG00000215912 | TTC34 | tetratricopeptide repeat domain 34 [Source:HGNC Symbol;Acc:HGNC:34297] | -0.9638 | 0.01305 |
| ENSG00000105327 | BBC3 | BCL2 binding component 3 [Source:HGNC Symbol;Acc:HGNC:17868] | -1.2956 | 0.01314 |
| ENSG00000188599 | NPIPP1 | nuclear pore complex interacting protein pseudogene 1 [Source:HGNC Symbol;Acc:HGNC:35407] | -0.5710 | 0.01322 |
| ENSG00000075223 | SEMA3C | semaphorin 3C [Source:HGNC Symbol;Acc:HGNC:10725] | 1.8622 | 0.01327 |
| ENSG00000100218 | RSPH14 | radial spoke head 14 homolog [Source:HGNC Symbol;Acc:HGNC:13437] | -0.6771 | 0.01336 |
| ENSG00000228175 | GEMIN8P4 | gem nuclear organelle associated protein 8 pseudogene 4 [Source:HGNC Symbol;Acc:HGNC:37979] | -1.0179 | 0.01338 |
| ENSG00000244541 | LINC01213 | long intergenic non-protein coding RNA 1213 [Source:HGNC Symbol;Acc:HGNC:49648] | -1.6154 | 0.01339 |
| ENSG00000249631 | NA | novel transcript | 0.8837 | 0.01341 |
| ENSG00000137218 | FRS3 | fibroblast growth factor receptor substrate 3 [Source:HGNC Symbol;Acc:HGNC:16970] | -0.5660 | 0.01346 |
| ENSG00000075429 | CACNG5 | calcium voltage-gated channel auxiliary subunit gamma 5 [Source:HGNC Symbol;Acc:HGNC:1409] | -2.6784 | 0.01350 |
| ENSG00000115084 | SLC35F5 | solute carrier family 35 member F5 [Source:HGNC Symbol;Acc:HGNC:23617] | 0.5094 | 0.01352 |
| ENSG00000286449 | NA | novel transcript | -0.9303 | 0.01356 |
| ENSG00000286757 | NA | novel transcript, antisense to CLYBL | -1.7719 | 0.01358 |
| ENSG00000267546 | NA | novel transcript, antisense to PRCD | -1.7961 | 0.01363 |
| ENSG00000101000 | PROCR | protein C receptor [Source:HGNC Symbol;Acc:HGNC:9452] | 0.5330 | 0.01365 |
| ENSG00000203877 | RIPPLY2 | ripply transcriptional repressor 2 [Source:HGNC Symbol;Acc:HGNC:21390] | -3.1544 | 0.01368 |
| ENSG00000256463 | SALL3 | spalt like transcription factor 3 [Source:HGNC Symbol;Acc:HGNC:10527] | 2.4412 | 0.01371 |
| ENSG00000265590 | CFAP298-TCP10L | CFAP298-TCP10L readthrough [Source:HGNC Symbol;Acc:HGNC:54636] | 0.8699 | 0.01373 |
| ENSG00000177409 | SAMD9L | sterile alpha motif domain containing 9 like [Source:HGNC Symbol;Acc:HGNC:1349] | -1.1009 | 0.01376 |
| ENSG00000237984 | PTENP1 | phosphatase and tensin homolog pseudogene 1 [Source:HGNC Symbol;Acc:HGNC:9589] | -0.6498 | 0.01377 |
| ENSG00000197329 | PELI1 | pellino E3 ubiquitin protein ligase 1 [Source:HGNC Symbol;Acc:HGNC:8827] | 0.7090 | 0.01378 |
| ENSG00000198300 | PEG3 | paternally expressed 3 [Source:HGNC Symbol;Acc:HGNC:8826] | -1.3333 | 0.01378 |
| ENSG00000260630 | SNAI3-AS1 | SNAI3 antisense RNA 1 [Source:HGNC Symbol;Acc:HGNC:28327] | -0.6129 | 0.01384 |
| ENSG00000107984 | 1,00 DKK | dickkopf WNT signaling pathway inhibitor 1 [Source:HGNC Symbol;Acc:HGNC:2891] | 0.9583 | 0.01392 |
| ENSG00000261046 | C2orf69P1 | chromosome 2 open reading frame 69 pseudogene 1 [Source:HGNC Symbol;Acc:HGNC:51800] | -1.0448 | 0.01394 |
| ENSG00000066248 | NGEF | neuronal guanine nucleotide exchange factor [Source:HGNC Symbol;Acc:HGNC:7807] | 1.0879 | 0.01395 |
| ENSG00000085831 | TTC39A | tetratricopeptide repeat domain 39A [Source:HGNC Symbol;Acc:HGNC:18657] | 0.7961 | 0.01395 |
| ENSG00000176387 | HSD11B2 | hydroxysteroid 11-beta dehydrogenase 2 [Source:HGNC Symbol;Acc:HGNC:5209] | -1.3321 | 0.01396 |
| ENSG00000099250 | NRP1 | neuropilin 1 [Source:HGNC Symbol;Acc:HGNC:8004] | 0.5730 | 0.01403 |
| ENSG00000158865 | SLC5A11 | solute carrier family 5 member 11 [Source:HGNC Symbol;Acc:HGNC:23091] | -1.1169 | 0.01403 |
| ENSG00000163794 | UCN | urocortin [Source:HGNC Symbol;Acc:HGNC:12516] | 0.8339 | 0.01404 |
| ENSG00000262096 | PCDHB19P | protocadherin beta 19 pseudogene [Source:HGNC Symbol;Acc:HGNC:14549] | -1.0594 | 0.01404 |
| ENSG00000182687 | GALR2 | galanin receptor 2 [Source:HGNC Symbol;Acc:HGNC:4133] | 1.8945 | 0.01416 |
| ENSG00000069943 | PIGB | phosphatidylinositol glycan anchor biosynthesis class B [Source:HGNC Symbol;Acc:HGNC:8959] | 0.7449 | 0.01426 |
| ENSG00000081479 | LRP2 | LDL receptor related protein 2 [Source:HGNC Symbol;Acc:HGNC:6694] | -4.2621 | 0.01437 |
| ENSG00000260691 | ANKRD20A1 | ankyrin repeat domain 20 family member A1 [Source:HGNC Symbol;Acc:HGNC:23665] | -0.7253 | 0.01444 |
| ENSG00000249286 | AMD1P3 | adenosylmethionine decarboxylase 1 pseudogene 3 [Source:HGNC Symbol;Acc:HGNC:51608] | -1.3388 | 0.01444 |
| ENSG00000166793 | YPEL4 | yippee like 4 [Source:HGNC Symbol;Acc:HGNC:18328] | 0.6769 | 0.01446 |
| ENSG00000259985 | NA | novel transcript, antisense to B4GALT6 | -0.6957 | 0.01457 |
| ENSG00000162849 | KIF26B | kinesin family member 26B [Source:HGNC Symbol;Acc:HGNC:25484] | -0.8690 | 0.01459 |
| ENSG00000166128 | RAB8B | RAB8B, member RAS oncogene family [Source:HGNC Symbol;Acc:HGNC:30273] | -0.5055 | 0.01464 |
| ENSG00000267248 | NA | novel transcript | -0.8624 | 0.01467 |
| ENSG00000141449 | GREB1L | GREB1 like retinoic acid receptor coactivator [Source:HGNC Symbol;Acc:HGNC:31042] | -0.7291 | 0.01469 |

| | | | | |
|---|---|---|---|---|
| ENSG00000172020 | GAP43 | growth associated protein 43 [Source:HGNC Symbol;Acc:HGNC:4140] | 0.5921 | 0.01469 |
| ENSG00000180178 | FAR2P1 | fatty acyl-CoA reductase 2 pseudogene 1 [Source:HGNC Symbol;Acc:HGNC:49284] | -0.5051 | 0.01469 |
| ENSG00000204338 | CYP21A1P | cytochrome P450 family 21 subfamily A member 1, pseudogene [Source:HGNC Symbol;Acc:HGNC:2599] | 1.1613 | 0.01470 |
| ENSG00000226363 | HAGLROS | HAGLR opposite strand lncRNA [Source:HGNC Symbol;Acc:HGNC:50646] | -1.5279 | 0.01475 |
| ENSG00000273253 | NA | novel transcript, antisense TRABD | -1.8961 | 0.01485 |
| ENSG00000188897 | NA | novel lipoprotein amino terminal region containing protein | -1.8325 | 0.01486 |
| ENSG00000268869 | ESPNP | espin pseudogene [Source:HGNC Symbol;Acc:HGNC:23285] | -1.5769 | 0.01486 |
| ENSG00000239282 | CASTOR1 | cytosolic arginine sensor for mTORC1 subunit 1 [Source:HGNC Symbol;Acc:HGNC:34423] | -0.9252 | 0.01490 |
| ENSG00000276535 | NA | arginine/serine-rich coiled-coil 1 (RSRC1) pseudogene | -1.1753 | 0.01494 |
| ENSG00000121210 | TMEM131L | transmembrane 131 like [Source:HGNC Symbol;Acc:HGNC:29146] | 0.6446 | 0.01504 |
| ENSG00000244055 | NA | novel transcript, antisense to ERVW-1 and PEX1 | 0.7345 | 0.01525 |
| ENSG00000140832 | MARVELD3 | MARVEL domain containing 3 [Source:HGNC Symbol;Acc:HGNC:30525] | -0.8931 | 0.01530 |
| ENSG00000225968 | ELFN1 | extracellular leucine rich repeat and fibronectin type III domain containing 1 [Source:HGNC Symbol;Acc:HGNC:33154] | -0.9921 | 0.01533 |
| ENSG00000162814 | SPATA17 | spermatogenesis associated 17 [Source:HGNC Symbol;Acc:HGNC:25184] | -0.9208 | 0.01537 |
| ENSG00000234996 | NA | actin, gamma 1 (ACTG1) pseudogene | -1.2349 | 0.01548 |
| ENSG00000281357 | ARRDC3-AS1 | ARRDC3 antisense RNA 1 [Source:HGNC Symbol;Acc:HGNC:44145] | -0.6790 | 0.01569 |
| ENSG00000224635 | NA | novel transcript | 0.9852 | 0.01572 |
| ENSG00000070759 | TESK2 | testis associated actin remodelling kinase 2 [Source:HGNC Symbol;Acc:HGNC:11732] | -0.5613 | 0.01575 |
| ENSG00000031081 | ARHGAP31 | Rho GTPase activating protein 31 [Source:HGNC Symbol;Acc:HGNC:29216] | 0.6236 | 0.01580 |
| ENSG00000273420 | NA | novel transcript | -0.6778 | 0.01589 |
| ENSG00000125775 | SDCBP2 | syndecan binding protein 2 [Source:HGNC Symbol;Acc:HGNC:15756] | 1.0103 | 0.01590 |
| ENSG00000196890 | H2BU1 | H2B.U histone 1 [Source:HGNC Symbol;Acc:HGNC:20514] | -0.6722 | 0.01590 |
| ENSG00000283189 | NA | novel protein | -1.6930 | 0.01591 |
| ENSG00000134256 | CD101 | CD101 molecule [Source:HGNC Symbol;Acc:HGNC:5949] | 0.7902 | 0.01593 |
| ENSG00000139144 | PIK3C2G | phosphatidylinositol-4-phosphate 3-kinase catalytic subunit type 2 gamma [Source:HGNC Symbol;Acc:HGNC:8973] | -0.6964 | 0.01598 |
| ENSG00000236754 | NA | novel transcript | 1.9003 | 0.01602 |
| ENSG00000110203 | FOLR3 | folate receptor gamma [Source:HGNC Symbol;Acc:HGNC:3795] | -1.8902 | 0.01619 |
| ENSG00000165181 | SHOC1 | shortage in chiasmata 1 [Source:HGNC Symbol;Acc:HGNC:26535] | 0.5180 | 0.01622 |
| ENSG00000134042 | MRO | maestro [Source:HGNC Symbol;Acc:HGNC:24121] | -1.4233 | 0.01626 |
| ENSG00000203858 | HSD3BP2 | hydroxy-delta-5-steroid dehydrogenase, 3 beta, pseudogene 2 [Source:HGNC Symbol;Acc:HGNC:5220] | 2.1469 | 0.01628 |
| ENSG00000260403 | NA | novel transcript, antisense to CACNA1H | -2.6034 | 0.01629 |
| ENSG00000183154 | NA | novel transcript, antisense to ERLIN2 | 1.1212 | 0.01632 |
| ENSG00000112238 | PRDM13 | PR/SET domain 13 [Source:HGNC Symbol;Acc:HGNC:13998] | -0.7615 | 0.01632 |
| ENSG00000215158 | NA | Putative beta-glucuronidase-like protein FLJ75287 pseudogene | -0.6938 | 0.01633 |
| ENSG00000174576 | NPAS4 | neuronal PAS domain protein 4 [Source:HGNC Symbol;Acc:HGNC:18983] | 2.9711 | 0.01637 |
| ENSG00000074317 | SNCB | synuclein beta [Source:HGNC Symbol;Acc:HGNC:11140] | -2.7643 | 0.01637 |
| ENSG00000232046 | LINC01798 | long intergenic non-protein coding RNA 1798 [Source:HGNC Symbol;Acc:HGNC:52588] | -0.9673 | 0.01637 |
| ENSG00000102886 | GDPD3 | glycerophosphodiester phosphodiesterase domain containing 3 [Source:HGNC Symbol;Acc:HGNC:28638] | -0.8404 | 0.01641 |
| ENSG00000102890 | ELMO3 | engulfment and cell motility 3 [Source:HGNC Symbol;Acc:HGNC:17289] | 0.8061 | 0.01648 |
| ENSG00000111181 | SLC6A12 | solute carrier family 6 member 12 [Source:HGNC Symbol;Acc:HGNC:11045] | 1.9284 | 0.01648 |
| ENSG00000100626 | GALNT16 | polypeptide N-acetylgalactosaminyltransferase 16 [Source:HGNC Symbol;Acc:HGNC:23233] | 0.5786 | 0.01654 |
| ENSG00000162722 | TRIM58 | tripartite motif containing 58 [Source:HGNC Symbol;Acc:HGNC:24150] | -1.1170 | 0.01656 |

| | | | | |
|---|---|---|---|---|
| ENSG00000276417 | NA | novel transcript | -2.2713 | 0.01658 |
| ENSG00000283403 | NA | novel transcript | -0.6961 | 0.01659 |
| ENSG00000205293 | LINC01602 | long intergenic non-protein coding RNA 1602 [Source:HGNC Symbol;Acc:HGNC:51634] | 1.7180 | 0.01661 |
| ENSG00000215045 | GRID2IP | Grid2 interacting protein [Source:HGNC Symbol;Acc:HGNC:18464] | -1.4639 | 0.01663 |
| ENSG00000159648 | TEPP | testis, prostate and placenta expressed [Source:HGNC Symbol;Acc:HGNC:33745] | -1.7281 | 0.01665 |
| ENSG00000188868 | ZNF563 | zinc finger protein 563 [Source:HGNC Symbol;Acc:HGNC:30498] | -0.5760 | 0.01685 |
| ENSG00000119950 | MXI1 | MAX interactor 1, dimerization protein [Source:HGNC Symbol;Acc:HGNC:7534] | -0.5463 | 0.01693 |
| ENSG00000160683 | CXCR5 | C-X-C motif chemokine receptor 5 [Source:HGNC Symbol;Acc:HGNC:1060] | -5.4573 | 0.01694 |
| ENSG00000156384 | SFR1 | SWI5 dependent homologous recombination repair protein 1 [Source:HGNC Symbol;Acc:HGNC:29574] | -0.5307 | 0.01701 |
| ENSG00000188833 | ENTPD8 | ectonucleoside triphosphate diphosphohydrolase 8 [Source:HGNC Symbol;Acc:HGNC:24860] | 0.6225 | 0.01703 |
| ENSG00000151846 | PABPC3 | poly(A) binding protein cytoplasmic 3 [Source:HGNC Symbol;Acc:HGNC:8556] | 1.0665 | 0.01705 |
| ENSG00000137494 | ANKRD42 | ankyrin repeat domain 42 [Source:HGNC Symbol;Acc:HGNC:26752] | 0.5507 | 0.01714 |
| ENSG00000163075 | CFAP221 | cilia and flagella associated protein 221 [Source:HGNC Symbol;Acc:HGNC:33720] | -0.5989 | 0.01721 |
| ENSG00000258674 | NA | novel protein | 0.7545 | 0.01722 |
| ENSG00000263280 | NA | novel transcript | -1.2886 | 0.01722 |
| ENSG00000235972 | NA | novel transcript | -3.2047 | 0.01727 |
| ENSG00000135931 | ARMC9 | armadillo repeat containing 9 [Source:HGNC Symbol;Acc:HGNC:20730] | 0.5883 | 0.01727 |
| ENSG00000288066 | NA | novel transcript, antisense to ARL5A | 0.5164 | 0.01728 |
| ENSG00000256001 | NA | novel transcript | -1.9796 | 0.01729 |
| ENSG00000233967 | NA | novel transcript | -0.8611 | 0.01731 |
| ENSG00000143476 | DTL | denticleless E3 ubiquitin protein ligase homolog [Source:HGNC Symbol;Acc:HGNC:30288] | -0.5775 | 0.01732 |
| ENSG00000279976 | NA | novel transcript | 3.6065 | 0.01733 |
| ENSG00000258301 | VASH1-AS1 | VASH1 antisense RNA 1 [Source:HGNC Symbol;Acc:HGNC:53215] | -0.8732 | 0.01742 |
| ENSG00000079385 | CEACAM1 | CEA cell adhesion molecule 1 [Source:HGNC Symbol;Acc:HGNC:1814] | 1.4111 | 0.01745 |
| ENSG00000137962 | ARHGAP29 | Rho GTPase activating protein 29 [Source:HGNC Symbol;Acc:HGNC:30207] | 1.0430 | 0.01746 |
| ENSG00000213091 | PHBP1 | prohibitin pseudogene 1 [Source:HGNC Symbol;Acc:HGNC:8913] | -2.3053 | 0.01758 |
| ENSG00000265399 | NA | novel transcript, antisense to MYOM1 | 0.6278 | 0.01760 |
| ENSG00000197768 | STPG3 | sperm-tail PG-rich repeat containing 3 [Source:HGNC Symbol;Acc:HGNC:37285] | -1.0572 | 0.01760 |
| ENSG00000272836 | NA | novel transcript, antisense to SELO | 1.5606 | 0.01761 |
| ENSG00000213638 | ADAT3 | adenosine deaminase tRNA specific 3 [Source:HGNC Symbol;Acc:HGNC:25151] | 0.5016 | 0.01763 |
| ENSG00000182901 | RGS7 | regulator of G protein signaling 7 [Source:HGNC Symbol;Acc:HGNC:10003] | 0.5894 | 0.01763 |
| ENSG00000104518 | GSDMD | gasdermin D [Source:HGNC Symbol;Acc:HGNC:25697] | -0.7486 | 0.01764 |
| ENSG00000281398 | SNHG4 | small nucleolar RNA host gene 4 [Source:HGNC Symbol;Acc:HGNC:32964] | 0.6200 | 0.01766 |
| ENSG00000267136 | NA | novel transcript, antisense C18orf1 | 2.2989 | 0.01767 |
| ENSG00000236618 | PITPNA-AS1 | PITPNA antisense RNA 1 [Source:HGNC Symbol;Acc:HGNC:44116] | -0.6376 | 0.01768 |
| ENSG00000175066 | GK5 | glycerol kinase 5 [Source:HGNC Symbol;Acc:HGNC:28635] | -0.5723 | 0.01771 |
| ENSG00000175707 | KDF1 | keratinocyte differentiation factor 1 [Source:HGNC Symbol;Acc:HGNC:26624] | -0.8054 | 0.01771 |
| ENSG00000286989 | NA | novel transcript | -1.5801 | 0.01775 |
| ENSG00000079335 | CDC14A | cell division cycle 14A [Source:HGNC Symbol;Acc:HGNC:1718] | 0.5963 | 0.01780 |
| ENSG00000177464 | GPR4 | G protein-coupled receptor 4 [Source:HGNC Symbol;Acc:HGNC:4497] | 0.7918 | 0.01794 |
| ENSG00000130643 | CALY | calcyon neuron specific vesicular protein [Source:HGNC Symbol;Acc:HGNC:17938] | 2.0537 | 0.01796 |
| ENSG00000125657 | TNFSF9 | TNF superfamily member 9 [Source:HGNC Symbol;Acc:HGNC:11939] | 0.7570 | 0.01803 |
| ENSG00000157315 | TMED6 | transmembrane p24 trafficking protein 6 [Source:HGNC Symbol;Acc:HGNC:28331] | 1.3555 | 0.01806 |

| | | | | |
|---|---|---|---|---|
| ENSG00000166407 | LMO1 | LIM domain only 1 [Source:HGNC Symbol;Acc:HGNC:6641] | -0.8389 | 0.01814 |
| ENSG00000260997 | NA | novel transcript, overlapping MYO1G | 1.5941 | 0.01816 |
| ENSG00000118473 | SGIP1 | SH3GL interacting endocytic adaptor 1 [Source:HGNC Symbol;Acc:HGNC:25412] | 1.5849 | 0.01817 |
| ENSG00000224825 | RORB-AS1 | RORB antisense RNA 1 [Source:HGNC Symbol;Acc:HGNC:49803] | -0.5085 | 0.01817 |
| ENSG00000279110 | NA | novel transcript | 1.4871 | 0.01818 |
| ENSG00000261542 | NA | novel transcript | 1.3729 | 0.01818 |
| ENSG00000260276 | NA | novel transcript, antisense to CARHSP1 | 0.8232 | 0.01821 |
| ENSG00000287542 | NA | HECT and RLD domain containing E3 ubiquitin protein ligase 3 [Source:NCBI gene (formerly Entrezgene);Acc:8916] | -0.9694 | 0.01821 |
| ENSG00000128886 | ELL3 | elongation factor for RNA polymerase II 3 [Source:HGNC Symbol;Acc:HGNC:23113] | -1.2024 | 0.01823 |
| ENSG00000275713 | H2BC9 | H2B clustered histone 9 [Source:HGNC Symbol;Acc:HGNC:4755] | -0.5922 | 0.01823 |
| ENSG00000102349 | KLF8 | Kruppel like factor 8 [Source:HGNC Symbol;Acc:HGNC:6351] | -0.6638 | 0.01823 |
| ENSG00000145911 | N4BP3 | NEDD4 binding protein 3 [Source:HGNC Symbol;Acc:HGNC:29852] | -0.6048 | 0.01823 |
| ENSG00000113555 | PCDH12 | protocadherin 12 [Source:HGNC Symbol;Acc:HGNC:8657] | 0.7492 | 0.01824 |
| ENSG00000056487 | PHF21B | PHD finger protein 21B [Source:HGNC Symbol;Acc:HGNC:25161] | 0.9168 | 0.01827 |
| ENSG00000106013 | ANKRD7 | ankyrin repeat domain 7 [Source:HGNC Symbol;Acc:HGNC:18588] | -1.2061 | 0.01828 |
| ENSG00000276007 | NA | novel transcript, antisense WWOX | -1.4348 | 0.01840 |
| ENSG00000285591 | NA | novel transcript | -1.0914 | 0.01844 |
| ENSG00000173442 | EHBP1L1 | EH domain binding protein 1 like 1 [Source:HGNC Symbol;Acc:HGNC:30682] | 0.5760 | 0.01846 |
| ENSG00000223466 | LINC01825 | long intergenic non-protein coding RNA 1825 [Source:HGNC Symbol;Acc:HGNC:52631] | -1.8356 | 0.01856 |
| ENSG00000225386 | NA | novel transcript | -0.5175 | 0.01861 |
| ENSG00000248830 | ZNF807P | zinc finger protein 807, pseudogene [Source:HGNC Symbol;Acc:HGNC:33229] | -1.5108 | 0.01869 |
| ENSG00000173597 | SULT1B1 | sulfotransferase family 1B member 1 [Source:HGNC Symbol;Acc:HGNC:17845] | -1.0008 | 0.01870 |
| ENSG00000215374 | FAM66B | family with sequence similarity 66 member B [Source:HGNC Symbol;Acc:HGNC:28890] | -1.7599 | 0.01870 |
| ENSG00000286116 | NA | FAS antisense RNA 1 | 1.0437 | 0.01874 |
| ENSG00000217236 | SP9 | Sp9 transcription factor [Source:HGNC Symbol;Acc:HGNC:30690] | -0.6182 | 0.01880 |
| ENSG00000262772 | LINC01977 | long intergenic non-protein coding RNA 1977 [Source:HGNC Symbol;Acc:HGNC:52804] | 0.8983 | 0.01880 |
| ENSG00000067601 | PMS2P4 | PMS1 homolog 2, mismatch repair system component pseudogene 4 [Source:HGNC Symbol;Acc:HGNC:9129] | -0.8575 | 0.01883 |
| ENSG00000204396 | VWA7 | von Willebrand factor A domain containing 7 [Source:HGNC Symbol;Acc:HGNC:13939] | 1.7115 | 0.01883 |
| ENSG00000278860 | NA | TEC | 2.0726 | 0.01894 |
| ENSG00000144642 | RBMS3 | RNA binding motif single stranded interacting protein 3 [Source:HGNC Symbol;Acc:HGNC:13427] | -0.6861 | 0.01897 |
| ENSG00000196867 | ZFP28 | ZFP28 zinc finger protein [Source:HGNC Symbol;Acc:HGNC:17801] | -0.6011 | 0.01897 |
| ENSG00000274964 | NA | novel transcript, sense intronic to BICD1 | 0.9155 | 0.01898 |
| ENSG00000261247 | GOLGA8T | golgin A8 family member T [Source:HGNC Symbol;Acc:HGNC:44410] | -1.4780 | 0.01904 |
| ENSG00000130045 | NXNL2 | nucleoredoxin like 2 [Source:HGNC Symbol;Acc:HGNC:30482] | -1.9838 | 0.01911 |
| ENSG00000077009 | NMRK2 | nicotinamide riboside kinase 2 [Source:HGNC Symbol;Acc:HGNC:17871] | 0.7599 | 0.01918 |
| ENSG00000100092 | SH3BP1 | SH3 domain binding protein 1 [Source:HGNC Symbol;Acc:HGNC:10824] | 0.6487 | 0.01925 |
| ENSG00000230707 | NA | novel pseudogene | -2.3977 | 0.01926 |
| ENSG00000176732 | PFN4 | profilin family member 4 [Source:HGNC Symbol;Acc:HGNC:31103] | -1.1798 | 0.01928 |
| ENSG00000230183 | CNOT6LP1 | CNOT6L pseudogene 1 [Source:HGNC Symbol;Acc:HGNC:32355] | 1.6327 | 0.01932 |
| ENSG00000124490 | CRISP2 | cysteine rich secretory protein 2 [Source:HGNC Symbol;Acc:HGNC:12024] | 1.8599 | 0.01935 |
| ENSG00000185324 | CDK10 | cyclin dependent kinase 10 [Source:HGNC Symbol;Acc:HGNC:1770] | -0.5026 | 0.01944 |
| ENSG00000140519 | RHCG | Rh family C glycoprotein [Source:HGNC Symbol;Acc:HGNC:18140] | -0.5738 | 0.01945 |
| ENSG00000132275 | RRP8 | ribosomal RNA processing 8 [Source:HGNC Symbol;Acc:HGNC:29030] | 0.5943 | 0.01945 |

| | | | | |
|---|---|---|---|---|
| ENSG00000260249 | NA | novel transcript, antisense to SNX20 | -0.6963 | 0.01945 |
| ENSG00000269937 | NA | novel transcript, antisense to PDPK1 | 0.5525 | 0.01946 |
| ENSG00000272829 | NA | novel transcript | -0.9997 | 0.01946 |
| ENSG00000177791 | MYOZ1 | myozenin 1 [Source:HGNC Symbol;Acc:HGNC:13752] | -0.8810 | 0.01949 |
| ENSG00000256671 | LIMS4 | LIM zinc finger domain containing 4 [Source:HGNC Symbol;Acc:HGNC:39941] | 1.2903 | 0.01967 |
| ENSG00000232692 | NA | novel transcript | 0.8783 | 0.01976 |
| ENSG00000196878 | LAMB3 | laminin subunit beta 3 [Source:HGNC Symbol;Acc:HGNC:6490] | 1.9468 | 0.01979 |
| ENSG00000125384 | PTGER2 | prostaglandin E receptor 2 [Source:HGNC Symbol;Acc:HGNC:9594] | -1.1768 | 0.01983 |
| ENSG00000171680 | PLEKHG5 | pleckstrin homology and RhoGEF domain containing G5 [Source:HGNC Symbol;Acc:HGNC:29105] | 0.9316 | 0.01984 |
| ENSG00000288550 | NA | novel protein | -5.2824 | 0.01984 |
| ENSG00000176946 | THAP4 | THAP domain containing 4 [Source:HGNC Symbol;Acc:HGNC:23187] | 0.6126 | 0.01985 |
| ENSG00000228395 | NA | novel transcript | -0.9626 | 0.01987 |
| ENSG00000229950 | TFAP2A-AS1 | TFAP2A antisense RNA 1 [Source:HGNC Symbol;Acc:HGNC:40579] | -2.1907 | 0.01988 |
| ENSG00000251595 | ABCA11P | ATP binding cassette subfamily A member 11, pseudogene [Source:HGNC Symbol;Acc:HGNC:31] | -1.2441 | 0.01989 |
| ENSG00000132330 | SCLY | selenocysteine lyase [Source:HGNC Symbol;Acc:HGNC:18161] | -0.5054 | 0.01990 |
| ENSG00000285935 | NA | novel transcript, antisense to WASF3 | -2.1691 | 0.01990 |
| ENSG00000198237 | NA | glucuronidase, beta (GUSB) pseudogene | -0.6295 | 0.01992 |
| ENSG00000204261 | PSMB8-AS1 | PSMB8 antisense RNA 1 (head to head) [Source:HGNC Symbol;Acc:HGNC:39758] | -0.8744 | 0.01995 |
| ENSG00000130529 | TRPM4 | transient receptor potential cation channel subfamily M member 4 [Source:HGNC Symbol;Acc:HGNC:17993] | -0.6991 | 0.01997 |
| ENSG00000221995 | TIAF1 | TGFB1-induced anti-apoptotic factor 1 [Source:HGNC Symbol;Acc:HGNC:11803] | -0.6280 | 0.02003 |
| ENSG00000272791 | NA | novel transcript | -1.1181 | 0.02009 |
| ENSG00000154342 | WNT3A | Wnt family member 3A [Source:HGNC Symbol;Acc:HGNC:15983] | 2.1892 | 0.02037 |
| ENSG00000285420 | PDGFRL2P | platelet derived growth factor receptor like 2, pseudogene [Source:HGNC Symbol;Acc:HGNC:54778] | -1.3622 | 0.02041 |
| ENSG00000149922 | TBX6 | T-box transcription factor 6 [Source:HGNC Symbol;Acc:HGNC:11605] | -0.7208 | 0.02046 |
| ENSG00000225173 | NA | novel transcript | -1.2243 | 0.02054 |
| ENSG00000118777 | ABCG2 | ATP binding cassette subfamily G member 2 (Junior blood group) [Source:HGNC Symbol;Acc:HGNC:74] | 1.8890 | 0.02054 |
| ENSG00000182747 | SLC35D3 | solute carrier family 35 member D3 [Source:HGNC Symbol;Acc:HGNC:15621] | 1.1091 | 0.02059 |
| ENSG00000178977 | LINC00324 | long intergenic non-protein coding RNA 324 [Source:HGNC Symbol;Acc:HGNC:26628] | -0.5714 | 0.02072 |
| ENSG00000198590 | C3orf35 | chromosome 3 open reading frame 35 [Source:HGNC Symbol;Acc:HGNC:24082] | 1.0923 | 0.02074 |
| ENSG00000118922 | KLF12 | Kruppel like factor 12 [Source:HGNC Symbol;Acc:HGNC:6346] | 1.6838 | 0.02076 |
| ENSG00000273820 | USP27X | ubiquitin specific peptidase 27 X-linked [Source:HGNC Symbol;Acc:HGNC:13486] | -0.6489 | 0.02081 |
| ENSG00000272572 | NA | novel transcript, antisense to FAM178A | -0.9057 | 0.02082 |
| ENSG00000153237 | CCDC148 | coiled-coil domain containing 148 [Source:HGNC Symbol;Acc:HGNC:25191] | -1.6949 | 0.02089 |
| ENSG00000112280 | COL9A1 | collagen type IX alpha 1 chain [Source:HGNC Symbol;Acc:HGNC:2217] | -0.8054 | 0.02092 |
| ENSG00000248890 | HHIP-AS1 | HHIP antisense RNA 1 [Source:HGNC Symbol;Acc:HGNC:44182] | -0.6179 | 0.02093 |
| ENSG00000196639 | HRH1 | histamine receptor H1 [Source:HGNC Symbol;Acc:HGNC:5182] | 0.6078 | 0.02100 |
| ENSG00000273706 | LHX1 | LIM homeobox 1 [Source:HGNC Symbol;Acc:HGNC:6593] | -2.9511 | 0.02102 |
| ENSG00000198353 | HOXC4 | homeobox C4 [Source:HGNC Symbol;Acc:HGNC:5126] | -0.6445 | 0.02105 |
| ENSG00000137101 | CD72 | CD72 molecule [Source:HGNC Symbol;Acc:HGNC:1696] | -0.5744 | 0.02108 |
| ENSG00000198133 | TMEM229B | transmembrane protein 229B [Source:HGNC Symbol;Acc:HGNC:20130] | -0.6615 | 0.02109 |
| ENSG00000137960 | GIPC2 | GIPC PDZ domain containing family member 2 [Source:HGNC Symbol;Acc:HGNC:18177] | 1.7074 | 0.02111 |
| ENSG00000266066 | POLRMTP1 | RNA polymerase mitochondrial pseudogene 1 [Source:HGNC Symbol;Acc:HGNC:28865] | 0.6188 | 0.02115 |
| ENSG00000129194 | SOX15 | SRY-box transcription factor 15 [Source:HGNC Symbol;Acc:HGNC:11196] | 0.6418 | 0.02117 |

| | | | | |
|---|---|---|---|---|
| ENSG00000286064 | NA | aldo-keto reductase family pseudogene | 1.2199 | 0.02118 |
| ENSG00000285542 | NA | novel protein, PPM1B-SLC3A1 readthrough | -2.0369 | 0.02128 |
| ENSG00000160323 | ADAMTS13 | ADAM metallopeptidase with thrombospondin type 1 motif 13 [Source:HGNC Symbol;Acc:HGNC:1366] | -0.9394 | 0.02129 |
| ENSG00000268883 | PNMA6B | PNMA family member 6B (pseudogene) [Source:HGNC Symbol;Acc:HGNC:26681] | -0.7319 | 0.02158 |
| ENSG00000157653 | C9orf43 | chromosome 9 open reading frame 43 [Source:HGNC Symbol;Acc:HGNC:23570] | -0.6865 | 0.02158 |
| ENSG00000244115 | DNAJC25-GNG10 | DNAJC25-GNG10 readthrough [Source:HGNC Symbol;Acc:HGNC:37501] | 0.5558 | 0.02158 |
| ENSG00000287725 | NA | novel protein, TPCN2 - SMIM38 readthrough | -1.9316 | 0.02159 |
| ENSG00000156500 | FAM122C | family with sequence similarity 122C [Source:HGNC Symbol;Acc:HGNC:25202] | -0.6187 | 0.02170 |
| ENSG00000006611 | USH1C | USH1 protein network component harmonin [Source:HGNC Symbol;Acc:HGNC:12597] | -1.3035 | 0.02171 |
| ENSG00000234043 | NUDT9P1 | nudix hydrolase 9 pseudogene 1 [Source:HGNC Symbol;Acc:HGNC:28523] | -1.4856 | 0.02183 |
| ENSG00000279442 | NA | novel transcript | -1.1495 | 0.02199 |
| ENSG00000183421 | RIPK4 | receptor interacting serine/threonine kinase 4 [Source:HGNC Symbol;Acc:HGNC:496] | -0.8414 | 0.02210 |
| ENSG00000161653 | NAGS | N-acetylglutamate synthase [Source:HGNC Symbol;Acc:HGNC:17996] | 0.6861 | 0.02218 |
| ENSG00000115850 | LCT | lactase [Source:HGNC Symbol;Acc:HGNC:6530] | 0.7339 | 0.02224 |
| ENSG00000196227 | FAM217B | family with sequence similarity 217 member B [Source:HGNC Symbol;Acc:HGNC:16170] | -0.5334 | 0.02232 |
| ENSG00000287721 | NA | novel transcript | 0.6733 | 0.02235 |
| ENSG00000232931 | LINC00342 | long intergenic non-protein coding RNA 342 [Source:HGNC Symbol;Acc:HGNC:42470] | -0.5852 | 0.02236 |
| ENSG00000286485 | NA | novel transcript, antisense to ANKRD54and IFT27 | -0.5253 | 0.02240 |
| ENSG00000275395 | FCGBP | Fc fragment of IgG binding protein [Source:HGNC Symbol;Acc:HGNC:13572] | -0.8202 | 0.02244 |
| ENSG00000227218 | NA | novel transcript | 0.8533 | 0.02244 |
| ENSG00000225975 | LINC01534 | long intergenic non-protein coding RNA 1534 [Source:HGNC Symbol;Acc:HGNC:51281] | -0.6464 | 0.02252 |
| ENSG00000263818 | RDM1P5 | RDM1 pseudogene 5 [Source:HGNC Symbol;Acc:HGNC:53921] | -2.0785 | 0.02258 |
| ENSG00000287575 | NA | novel transcript | 1.0817 | 0.02258 |
| ENSG00000183476 | SH2D7 | SH2 domain containing 7 [Source:HGNC Symbol;Acc:HGNC:34549] | -1.0683 | 0.02259 |
| ENSG00000101331 | CCM2L | CCM2 like scaffold protein [Source:HGNC Symbol;Acc:HGNC:16153] | -1.2788 | 0.02266 |
| ENSG00000135976 | ANKRD36 | ankyrin repeat domain 36 [Source:HGNC Symbol;Acc:HGNC:24079] | -0.6652 | 0.02282 |
| ENSG00000181333 | HEPHL1 | hephaestin like 1 [Source:HGNC Symbol;Acc:HGNC:30477] | -1.8575 | 0.02283 |
| ENSG00000122507 | BBS9 | Bardet-Biedl syndrome 9 [Source:HGNC Symbol;Acc:HGNC:30000] | -0.5461 | 0.02291 |
| ENSG00000198732 | SMOC1 | SPARC related modular calcium binding 1 [Source:HGNC Symbol;Acc:HGNC:20318] | 4.1071 | 0.02292 |
| ENSG00000215883 | CYB5RL | cytochrome b5 reductase like [Source:HGNC Symbol;Acc:HGNC:32220] | -0.6066 | 0.02292 |
| ENSG00000280310 | NA | tec | -1.5286 | 0.02292 |
| ENSG00000171126 | KCNG3 | potassium voltage-gated channel modifier subfamily G member 3 [Source:HGNC Symbol;Acc:HGNC:18306] | 0.7255 | 0.02305 |
| ENSG00000011638 | TMEM159 | transmembrane protein 159 [Source:HGNC Symbol;Acc:HGNC:30136] | -0.9871 | 0.02307 |
| ENSG00000261115 | TMEM178B | transmembrane protein 178B [Source:HGNC Symbol;Acc:HGNC:44112] | -1.1118 | 0.02308 |
| ENSG00000186862 | PDZD7 | PDZ domain containing 7 [Source:HGNC Symbol;Acc:HGNC:26257] | -0.7652 | 0.02309 |
| ENSG00000234807 | LINC01135 | long intergenic non-protein coding RNA 1135 [Source:HGNC Symbol;Acc:HGNC:49450] | -1.1908 | 0.02313 |
| ENSG00000174600 | CMKLR1 | chemerin chemokine-like receptor 1 [Source:HGNC Symbol;Acc:HGNC:2121] | -3.4707 | 0.02320 |
| ENSG00000234290 | NA | novel transcript | -0.6832 | 0.02320 |
| ENSG00000233527 | ZNF529-AS1 | ZNF529 antisense RNA 1 [Source:HGNC Symbol;Acc:HGNC:51275] | -0.5060 | 0.02321 |
| ENSG00000099365 | STX1B | syntaxin 1B [Source:HGNC Symbol;Acc:HGNC:18539] | 0.6332 | 0.02323 |
| ENSG00000118004 | COLEC11 | collectin subfamily member 11 [Source:HGNC Symbol;Acc:HGNC:17213] | -1.0271 | 0.02323 |
| ENSG00000286379 | NA | novel transcript | 0.8533 | 0.02324 |
| ENSG00000147573 | TRIM55 | tripartite motif containing 55 [Source:HGNC Symbol;Acc:HGNC:14215] | 1.4470 | 0.02329 |

| | | | | |
|---|---|---|---|---|
| ENSG00000170122 | FOXD4 | forkhead box D4 [Source:HGNC Symbol;Acc:HGNC:3805] | 0.7571 | 0.02337 |
| ENSG00000213906 | LTB4R2 | leukotriene B4 receptor 2 [Source:HGNC Symbol;Acc:HGNC:19260] | 0.5094 | 0.02341 |
| ENSG00000271780 | NA | novel transcript | -0.9020 | 0.02343 |
| ENSG00000267216 | NA | ZNF8-ERVK3-1 readthrough [Source:NCBI gene (formerly Entrezgene);Acc:108903150] | 0.6504 | 0.02344 |
| ENSG00000254332 | NA | tropomyosin 4 (TPM4) pseudogene | 1.2788 | 0.02346 |
| ENSG00000258667 | HIF1A-AS3 | HIF1A antisense RNA 3 [Source:HGNC Symbol;Acc:HGNC:54284] | 1.7799 | 0.02356 |
| ENSG00000214770 | NA | novel transcript | -0.8772 | 0.02358 |
| ENSG00000187800 | PEAR1 | platelet endothelial aggregation receptor 1 [Source:HGNC Symbol;Acc:HGNC:33631] | 2.4175 | 0.02358 |
| ENSG00000111728 | ST8SIA1 | ST8 alpha-N-acetyl-neuraminide alpha-2,8-sialyltransferase 1 [Source:HGNC Symbol;Acc:HGNC:10869] | -1.0627 | 0.02359 |
| ENSG00000075275 | CELSR1 | cadherin EGF LAG seven-pass G-type receptor 1 [Source:HGNC Symbol;Acc:HGNC:1850] | 1.2466 | 0.02367 |
| ENSG00000204428 | LY6G5C | lymphocyte antigen 6 family member G5C [Source:HGNC Symbol;Acc:HGNC:13932] | 1.3309 | 0.02375 |
| ENSG00000228274 | NA | novel transcript, antisense to CBY1 | -0.6148 | 0.02381 |
| ENSG00000110811 | P3H3 | prolyl 3-hydroxylase 3 [Source:HGNC Symbol;Acc:HGNC:19318] | -0.6648 | 0.02392 |
| ENSG00000234509 | NA | novel transcript | -2.2261 | 0.02413 |
| ENSG00000175820 | CCDC168 | coiled-coil domain containing 168 [Source:HGNC Symbol;Acc:HGNC:26851] | 1.0982 | 0.02414 |
| ENSG00000214973 | CHCHD3P3 | coiled-coil-helix-coiled-coil-helix domain containing 3 pseudogene 3 [Source:HGNC Symbol;Acc:HGNC:44699] | 1.4773 | 0.02415 |
| ENSG00000234264 | DEPDC1-AS1 | DEPDC1 antisense RNA 1 [Source:HGNC Symbol;Acc:HGNC:50592] | 0.9721 | 0.02416 |
| ENSG00000287923 | NA | novel transcript | -1.0850 | 0.02421 |
| ENSG00000188176 | SMTNL2 | smoothelin like 2 [Source:HGNC Symbol;Acc:HGNC:24764] | -1.9522 | 0.02425 |
| ENSG00000253877 | LINC01608 | long intergenic non-protein coding RNA 1608 [Source:HGNC Symbol;Acc:HGNC:51662] | -0.5816 | 0.02425 |
| ENSG00000227368 | CDK8P2 | cyclin dependent kinase 8 pseudogene 2 [Source:HGNC Symbol;Acc:HGNC:52296] | -1.8461 | 0.02431 |
| ENSG00000146674 | IGFBP3 | insulin like growth factor binding protein 3 [Source:HGNC Symbol;Acc:HGNC:5472] | 0.9106 | 0.02433 |
| ENSG00000143494 | VASH2 | vasohibin 2 [Source:HGNC Symbol;Acc:HGNC:25723] | -0.6813 | 0.02437 |
| ENSG00000259920 | NA | novel transcript | 1.4238 | 0.02443 |
| ENSG00000123329 | ARHGAP9 | Rho GTPase activating protein 9 [Source:HGNC Symbol;Acc:HGNC:14130] | 2.1892 | 0.02449 |
| ENSG00000227896 | NA | novel transcript | -0.9201 | 0.02450 |
| ENSG00000258405 | ZNF578 | zinc finger protein 578 [Source:HGNC Symbol;Acc:HGNC:26449] | -0.6151 | 0.02452 |
| ENSG00000109794 | FAM149A | family with sequence similarity 149 member A [Source:HGNC Symbol;Acc:HGNC:24527] | -1.3375 | 0.02455 |
| ENSG00000249413 | NA | novel transcript | -1.0386 | 0.02465 |
| ENSG00000167550 | RHEBL1 | RHEB like 1 [Source:HGNC Symbol;Acc:HGNC:21166] | 0.5555 | 0.02467 |
| ENSG00000197332 | NA | novel transcript | -0.7042 | 0.02468 |
| ENSG00000104728 | ARHGEF10 | Rho guanine nucleotide exchange factor 10 [Source:HGNC Symbol;Acc:HGNC:14103] | 0.5953 | 0.02475 |
| ENSG00000170153 | RNF150 | ring finger protein 150 [Source:HGNC Symbol;Acc:HGNC:23138] | 0.5941 | 0.02491 |
| ENSG00000129295 | LRRC6 | leucine rich repeat containing 6 [Source:HGNC Symbol;Acc:HGNC:16725] | 0.6502 | 0.02498 |
| ENSG00000092929 | UNC13D | unc-13 homolog D [Source:HGNC Symbol;Acc:HGNC:23147] | 1.1526 | 0.02523 |
| ENSG00000176681 | LRRC37A | leucine rich repeat containing 37A [Source:HGNC Symbol;Acc:HGNC:29069] | 0.5099 | 0.02523 |
| ENSG00000163545 | NUAK2 | NUAK family kinase 2 [Source:HGNC Symbol;Acc:HGNC:29558] | -0.8038 | 0.02533 |
| ENSG00000196218 | RYR1 | ryanodine receptor 1 [Source:HGNC Symbol;Acc:HGNC:10483] | -1.5683 | 0.02533 |
| ENSG00000164746 | C7orf57 | chromosome 7 open reading frame 57 [Source:HGNC Symbol;Acc:HGNC:22247] | -1.7954 | 0.02543 |
| ENSG00000170509 | HSD17B13 | hydroxysteroid 17-beta dehydrogenase 13 [Source:HGNC Symbol;Acc:HGNC:18685] | -0.5632 | 0.02547 |
| ENSG00000168356 | SCN11A | sodium voltage-gated channel alpha subunit 11 [Source:HGNC Symbol;Acc:HGNC:10583] | -1.5989 | 0.02549 |
| ENSG00000261644 | NA | novel transcript, antisense to CYLD and NOD2 | 1.9431 | 0.02550 |
| ENSG00000267327 | NA | novel transcript | -0.5090 | 0.02551 |

| | | | | |
|---|---|---|---|---|
| ENSG00000203761 | MSTO2P | misato family member 2, pseudogene [Source:HGNC Symbol;Acc:HGNC:26329] | 0.6467 | 0.02552 |
| ENSG00000262155 | LINC02175 | long intergenic non-protein coding RNA 2175 [Source:HGNC Symbol;Acc:HGNC:27550] | -0.5028 | 0.02554 |
| ENSG00000197587 | DMBX1 | diencephalon/mesencephalon homeobox 1 [Source:HGNC Symbol;Acc:HGNC:19026] | -1.0508 | 0.02566 |
| ENSG00000267886 | NA | novel transcript | -0.8738 | 0.02577 |
| ENSG00000270605 | NA | novel transcript | -1.8403 | 0.02585 |
| ENSG00000183066 | WBP2NL | WBP2 N-terminal like [Source:HGNC Symbol;Acc:HGNC:28389] | -1.4507 | 0.02587 |
| ENSG00000268055 | NA | novel transcript, antisense to RYR1 | -0.9733 | 0.02588 |
| ENSG00000236255 | NA | novel transcript | 0.6543 | 0.02594 |
| ENSG00000280047 | NA | TEC | -0.7783 | 0.02595 |
| ENSG00000286945 | NA | novel transcript | -2.1061 | 0.02603 |
| ENSG00000103490 | PYCARD | PYD and CARD domain containing [Source:HGNC Symbol;Acc:HGNC:16608] | 5.1732 | 0.02611 |
| ENSG00000242866 | STRC | stereocilin [Source:HGNC Symbol;Acc:HGNC:16035] | 0.9012 | 0.02612 |
| ENSG00000287038 | NA | novel transcript | -2.0337 | 0.02638 |
| ENSG00000066382 | MPPED2 | metallophosphoesterase domain containing 2 [Source:HGNC Symbol;Acc:HGNC:1180] | -1.1144 | 0.02639 |
| ENSG00000267727 | NA | novel transcript | 5.0665 | 0.02650 |
| ENSG00000231083 | NA | novel transcript | -1.0314 | 0.02650 |
| ENSG00000214439 | FAM185BP | family with sequence similarity 185 member B, pseudogene [Source:HGNC Symbol;Acc:HGNC:21813] | -0.6944 | 0.02650 |
| ENSG00000268119 | NA | novel transcript | -1.5977 | 0.02651 |
| ENSG00000269069 | NA | protein associated with topoisomerase II homolog 1 (yeast) (PATL1) pseudogene | 0.8436 | 0.02652 |
| ENSG00000127325 | BEST3 | bestrophin 3 [Source:HGNC Symbol;Acc:HGNC:17105] | 1.7524 | 0.02656 |
| ENSG00000285822 | NA | novel transcript | -0.7503 | 0.02663 |
| ENSG00000232079 | LINC01697 | long intergenic non-protein coding RNA 1697 [Source:HGNC Symbol;Acc:HGNC:52485] | 1.4100 | 0.02664 |
| ENSG00000272275 | NA | novel transcript | 0.7564 | 0.02664 |
| ENSG00000254166 | CASC19 | cancer susceptibility 19 [Source:HGNC Symbol;Acc:HGNC:49476] | -1.6118 | 0.02669 |
| ENSG00000263020 | NA | novel protein | 1.1471 | 0.02672 |
| ENSG00000253910 | PCDHGB2 | protocadherin gamma subfamily B, 2 [Source:HGNC Symbol;Acc:HGNC:8709] | -0.6172 | 0.02676 |
| ENSG00000287431 | NA | novel transcript | -0.8651 | 0.02678 |
| ENSG00000188818 | ZDHHC11 | zinc finger DHHC-type containing 11 [Source:HGNC Symbol;Acc:HGNC:19158] | 0.6773 | 0.02683 |
| ENSG00000140092 | FBLN5 | fibulin 5 [Source:HGNC Symbol;Acc:HGNC:3602] | -1.0670 | 0.02685 |
| ENSG00000171004 | HS6ST2 | heparan sulfate 6-O-sulfotransferase 2 [Source:HGNC Symbol;Acc:HGNC:19133] | -0.6876 | 0.02689 |
| ENSG00000169508 | GPR183 | G protein-coupled receptor 183 [Source:HGNC Symbol;Acc:HGNC:3128] | -0.6010 | 0.02691 |
| ENSG00000231007 | CDC20P1 | cell division cycle 20 pseudogene 1 [Source:HGNC Symbol;Acc:HGNC:29487] | -0.8227 | 0.02703 |
| ENSG00000127507 | ADGRE2 | adhesion G protein-coupled receptor E2 [Source:HGNC Symbol;Acc:HGNC:3337] | 2.2699 | 0.02717 |
| ENSG00000215869 | NA | THAP domain containing, apoptosis associated protein 3 (THAP3) pseudogene | -2.5281 | 0.02737 |
| ENSG00000256061 | DNAAF4 | dynein axonemal assembly factor 4 [Source:HGNC Symbol;Acc:HGNC:21493] | -0.8230 | 0.02745 |
| ENSG00000126822 | PLEKHG3 | pleckstrin homology and RhoGEF domain containing G3 [Source:HGNC Symbol;Acc:HGNC:20364] | -0.6987 | 0.02748 |
| ENSG00000271553 | NA | novel transcript | -0.9001 | 0.02757 |
| ENSG00000093217 | XYLB | xylulokinase [Source:HGNC Symbol;Acc:HGNC:12839] | -0.6129 | 0.02767 |
| ENSG00000229212 | NA | golgin subfamily A pseudogene | -3.6934 | 0.02775 |
| ENSG00000279170 | TSTD3 | thiosulfate sulfurtransferase like domain containing 3 [Source:HGNC Symbol;Acc:HGNC:40910] | -0.5944 | 0.02779 |
| ENSG00000229043 | NA | novel transcript | -0.8027 | 0.02785 |
| ENSG00000235531 | MSC-AS1 | MSC antisense RNA 1 [Source:HGNC Symbol;Acc:HGNC:48724] | 0.9983 | 0.02787 |
| ENSG00000238129 | NA | novel transcript | 0.5439 | 0.02788 |

| | | | | |
|---|---|---|---|---|
| ENSG00000240069 | GPAA1P1 | glycosylphosphatidylinositol anchor attachment 1 pseudogene 1 [Source:HGNC Symbol;Acc:HGNC:4447] | 2.6877 | 0.02791 |
| ENSG00000172830 | SSH3 | slingshot protein phosphatase 3 [Source:HGNC Symbol;Acc:HGNC:30581] | 0.9835 | 0.02792 |
| ENSG00000169962 | TAS1R3 | taste 1 receptor member 3 [Source:HGNC Symbol;Acc:HGNC:15661] | 0.6029 | 0.02792 |
| ENSG00000213981 | NA | novel transcript, antisense to MYO3B | -1.2119 | 0.02807 |
| ENSG00000225746 | MEG8 | maternally expressed 8, small nucleolar RNA host gene [Source:HGNC Symbol;Acc:HGNC:14574] | 0.5278 | 0.02814 |
| ENSG00000283154 | IQCJ-SCHIP1 | IQCJ-SCHIP1 readthrough [Source:HGNC Symbol;Acc:HGNC:38842] | 1.3712 | 0.02817 |
| ENSG00000276115 | NA | novel transcript, sense intronic to BICD1 | 1.1650 | 0.02820 |
| ENSG00000053438 | NNAT | neuronatin [Source:HGNC Symbol;Acc:HGNC:7860] | 0.9925 | 0.02821 |
| ENSG00000125571 | IL37 | interleukin 37 [Source:HGNC Symbol;Acc:HGNC:15563] | -2.3157 | 0.02830 |
| ENSG00000185437 | SH3BGR | SH3 domain binding glutamate rich protein [Source:HGNC Symbol;Acc:HGNC:10822] | -0.6842 | 0.02831 |
| ENSG00000158859 | ADAMTS4 | ADAM metallopeptidase with thrombospondin type 1 motif 4 [Source:HGNC Symbol;Acc:HGNC:220] | -1.1427 | 0.02832 |
| ENSG00000236508 | ATP13A5-AS1 | ATP13A5 antisense RNA 1 [Source:HGNC Symbol;Acc:HGNC:41281] | -1.4223 | 0.02834 |
| ENSG00000249464 | LINC01091 | long intergenic non-protein coding RNA 1091 [Source:HGNC Symbol;Acc:HGNC:27721] | -1.3761 | 0.02840 |
| ENSG00000259692 | NA | novel transcript | -1.0053 | 0.02850 |
| ENSG00000185386 | MAPK11 | mitogen-activated protein kinase 11 [Source:HGNC Symbol;Acc:HGNC:6873] | -1.3193 | 0.02853 |
| ENSG00000244021 | NA | ribosomal protein L21 (RPL21) pseudogene | -4.0951 | 0.02854 |
| ENSG00000185291 | IL3RA | interleukin 3 receptor subunit alpha [Source:HGNC Symbol;Acc:HGNC:6012] | 1.1991 | 0.02864 |
| ENSG00000226031 | FGF13-AS1 | FGF13 antisense RNA 1 [Source:HGNC Symbol;Acc:HGNC:44264] | -1.4903 | 0.02864 |
| ENSG00000109576 | AADAT | aminoadipate aminotransferase [Source:HGNC Symbol;Acc:HGNC:17929] | -0.6887 | 0.02870 |
| ENSG00000172250 | SERHL | serine hydrolase like (pseudogene) [Source:HGNC Symbol;Acc:HGNC:14408] | 1.6152 | 0.02871 |
| ENSG00000269437 | NXF2B | nuclear RNA export factor 2B [Source:HGNC Symbol;Acc:HGNC:23984] | 0.9451 | 0.02872 |
| ENSG00000141664 | ZCCHC2 | zinc finger CCHC-type containing 2 [Source:HGNC Symbol;Acc:HGNC:22916] | -0.5007 | 0.02872 |
| ENSG00000189186 | DCAF8L2 | DDB1 and CUL4 associated factor 8 like 2 [Source:HGNC Symbol;Acc:HGNC:31811] | -0.5135 | 0.02874 |
| ENSG00000089820 | ARHGAP4 | Rho GTPase activating protein 4 [Source:HGNC Symbol;Acc:HGNC:674] | -0.5167 | 0.02879 |
| ENSG00000125355 | TMEM255A | transmembrane protein 255A [Source:HGNC Symbol;Acc:HGNC:26086] | -0.6866 | 0.02888 |
| ENSG00000268182 | SMIM17 | small integral membrane protein 17 [Source:HGNC Symbol;Acc:HGNC:27114] | -1.3822 | 0.02888 |
| ENSG00000105750 | ZNF85 | zinc finger protein 85 [Source:HGNC Symbol;Acc:HGNC:13160] | -0.6123 | 0.02890 |
| ENSG00000259248 | USP3-AS1 | USP3 antisense RNA 1 [Source:HGNC Symbol;Acc:HGNC:44140] | -0.7420 | 0.02893 |
| ENSG00000074527 | NTN4 | netrin 4 [Source:HGNC Symbol;Acc:HGNC:13658] | 0.5237 | 0.02898 |
| ENSG00000178162 | FAR2P2 | fatty acyl-CoA reductase 2 pseudogene 2 [Source:HGNC Symbol;Acc:HGNC:49279] | -0.8400 | 0.02898 |
| ENSG00000225180 | PVALEF | parvalbumin like EF-hand containing [Source:HGNC Symbol;Acc:HGNC:40053] | -2.5445 | 0.02898 |
| ENSG00000227210 | NA | novel transcript | -1.1549 | 0.02898 |
| ENSG00000287539 | NA | novel transcript | -0.8164 | 0.02901 |
| ENSG00000187699 | C2orf88 | chromosome 2 open reading frame 88 [Source:HGNC Symbol;Acc:HGNC:28191] | 1.6265 | 0.02908 |
| ENSG00000233006 | MIR3936HG | MIR3936 host gene [Source:HGNC Symbol;Acc:HGNC:40538] | -0.5457 | 0.02909 |
| ENSG00000165507 | DEPP1 | DEPP1 autophagy regulator [Source:HGNC Symbol;Acc:HGNC:23355] | -1.0472 | 0.02911 |
| ENSG00000162669 | HFM1 | helicase for meiosis 1 [Source:HGNC Symbol;Acc:HGNC:20193] | -1.5277 | 0.02924 |
| ENSG00000105655 | ISYNA1 | inositol-3-phosphate synthase 1 [Source:HGNC Symbol;Acc:HGNC:29821] | 0.9184 | 0.02930 |
| ENSG00000283464 | NA | novel pseudogene identical to IGHVII-44-2D | -0.8476 | 0.02935 |
| ENSG00000033867 | SLC4A7 | solute carrier family 4 member 7 [Source:HGNC Symbol;Acc:HGNC:11033] | 0.5819 | 0.02938 |
| ENSG00000005108 | THSD7A | thrombospondin type 1 domain containing 7A [Source:HGNC Symbol;Acc:HGNC:22207] | -0.7194 | 0.02944 |
| ENSG00000267978 | MAGEA9B | MAGE family member A9B [Source:HGNC Symbol;Acc:HGNC:31909] | -0.5747 | 0.02944 |
| ENSG00000223760 | MED15P9 | mediator complex subunit 15 pseudogene 9 [Source:HGNC Symbol;Acc:HGNC:44130] | -0.7143 | 0.02953 |

| | | | | |
|---|---|---|---|---|
| ENSG00000230658 | KLHL7-DT | KLHL7 divergent transcript [Source:HGNC Symbol;Acc:HGNC:43431] | -0.8700 | 0.02953 |
| ENSG00000128482 | RNF112 | ring finger protein 112 [Source:HGNC Symbol;Acc:HGNC:12968] | -0.8007 | 0.02956 |
| ENSG00000175279 | CENPS | centromere protein S [Source:HGNC Symbol;Acc:HGNC:23163] | -0.5184 | 0.02956 |
| ENSG00000186231 | KLHL32 | kelch like family member 32 [Source:HGNC Symbol;Acc:HGNC:21221] | -1.3050 | 0.02961 |
| ENSG00000222764 | RN7SKP106 | RN7SK pseudogene 106 [Source:HGNC Symbol;Acc:HGNC:45830] | 1.0553 | 0.02972 |
| ENSG00000272701 | MESTIT1 | MEST intronic transcript 1, antisense RNA [Source:HGNC Symbol;Acc:HGNC:17991] | 1.0688 | 0.02972 |
| ENSG00000273151 | NA | novel transcript, antisense to GET4 | 0.5623 | 0.02972 |
| ENSG00000244151 | NA | novel transcript | 0.5915 | 0.02984 |
| ENSG00000256069 | A2MP1 | alpha-2-macroglobulin pseudogene 1 [Source:HGNC Symbol;Acc:HGNC:8] | 0.9169 | 0.02986 |
| ENSG00000226007 | NA | novel transcript | 0.9809 | 0.02987 |
| ENSG00000275493 | NA | NA | 1.3603 | 0.02987 |
| ENSG00000197044 | ZNF441 | zinc finger protein 441 [Source:HGNC Symbol;Acc:HGNC:20875] | -0.6307 | 0.02993 |
| ENSG00000258655 | ARHGAP5-AS1 | ARHGAP5 antisense RNA 1 (head to head) [Source:HGNC Symbol;Acc:HGNC:20279] | -0.5607 | 0.02997 |
| ENSG00000137273 | FOXF2 | forkhead box F2 [Source:HGNC Symbol;Acc:HGNC:3810] | -0.6796 | 0.03009 |
| ENSG00000281571 | NA | TEC | 0.6432 | 0.03015 |
| ENSG00000137843 | PAK6 | p21 (RAC1) activated kinase 6 [Source:HGNC Symbol;Acc:HGNC:16061] | 1.0528 | 0.03020 |
| ENSG00000198576 | ARC | activity regulated cytoskeleton associated protein [Source:HGNC Symbol;Acc:HGNC:648] | 2.4022 | 0.03025 |
| ENSG00000237914 | SIRPG-AS1 | SIRPG antisense RNA 1 [Source:HGNC Symbol;Acc:HGNC:51229] | -2.0841 | 0.03025 |
| ENSG00000113083 | LOX | lysyl oxidase [Source:HGNC Symbol;Acc:HGNC:6664] | 0.6790 | 0.03030 |
| ENSG00000257207 | NA | novel transcript | 2.8483 | 0.03030 |
| ENSG00000186280 | KDM4D | lysine demethylase 4D [Source:HGNC Symbol;Acc:HGNC:25498] | -0.6480 | 0.03067 |
| ENSG00000100276 | RASL10A | RAS like family 10 member A [Source:HGNC Symbol;Acc:HGNC:16954] | 1.1372 | 0.03068 |
| ENSG00000233858 | LINC02599 | long intergenic non-protein coding RNA 2599 [Source:HGNC Symbol;Acc:HGNC:53931] | -0.8148 | 0.03075 |
| ENSG00000165115 | KIF27 | kinesin family member 27 [Source:HGNC Symbol;Acc:HGNC:18632] | -0.5359 | 0.03077 |
| ENSG00000157404 | KIT | KIT proto-oncogene, receptor tyrosine kinase [Source:HGNC Symbol;Acc:HGNC:6342] | -2.7217 | 0.03084 |
| ENSG00000285427 | SOD2-OT1 | SOD2 overlapping transcript 1 [Source:HGNC Symbol;Acc:HGNC:53445] | -0.6140 | 0.03100 |
| ENSG00000163406 | SLC15A2 | solute carrier family 15 member 2 [Source:HGNC Symbol;Acc:HGNC:10921] | -1.5394 | 0.03112 |
| ENSG00000225492 | GBP1P1 | guanylate binding protein 1 pseudogene 1 [Source:HGNC Symbol;Acc:HGNC:39561] | -3.4577 | 0.03114 |
| ENSG00000227098 | NA | novel transcript | 1.3352 | 0.03125 |
| ENSG00000260805 | NA | novel transcript | -0.8114 | 0.03137 |
| ENSG00000233288 | NA | novel transcript | -1.0021 | 0.03167 |
| ENSG00000286545 | NA | novel transcript, antisense to ZC3H8 | -0.5650 | 0.03175 |
| ENSG00000117971 | CHRNB4 | cholinergic receptor nicotinic beta 4 subunit [Source:HGNC Symbol;Acc:HGNC:1964] | -1.9163 | 0.03179 |
| ENSG00000085514 | PILRA | paired immunoglobin like type 2 receptor alpha [Source:HGNC Symbol;Acc:HGNC:20396] | 0.6254 | 0.03188 |
| ENSG00000264569 | DCXR-DT | DCXR divergent transcript [Source:HGNC Symbol;Acc:HGNC:54083] | -1.7649 | 0.03189 |
| ENSG00000288612 | NA | novel transcript | 1.6482 | 0.03190 |
| ENSG00000273327 | OR6L2P | olfactory receptor family 6 subfamily L member 2 pseudogene [Source:HGNC Symbol;Acc:HGNC:15125] | 1.0393 | 0.03195 |
| ENSG00000151572 | ANO4 | anoctamin 4 [Source:HGNC Symbol;Acc:HGNC:23837] | -1.0641 | 0.03196 |
| ENSG00000170703 | TTLL6 | tubulin tyrosine ligase like 6 [Source:HGNC Symbol;Acc:HGNC:26664] | -2.1066 | 0.03208 |
| ENSG00000288473 | NA | novel protein | 0.9408 | 0.03209 |
| ENSG00000175505 | CLCF1 | cardiotrophin like cytokine factor 1 [Source:HGNC Symbol;Acc:HGNC:17412] | 1.9682 | 0.03211 |
| ENSG00000287966 | NA | novel transcript | -0.6205 | 0.03240 |
| ENSG00000270673 | YTHDF3-AS1 | YTHDF3 antisense RNA 1 (head to head) [Source:HGNC Symbol;Acc:HGNC:48728] | -0.6405 | 0.03245 |

| | | | | |
|---|---|---|---|---|
| ENSG00000275494 | NA | novel transcript | -0.8644 | 0.03246 |
| ENSG00000105675 | ATP4A | ATPase H+/K+ transporting subunit alpha [Source:HGNC Symbol;Acc:HGNC:819] | -1.0941 | 0.03264 |
| ENSG00000213612 | FAM220CP | family with sequence similarity 220 member C, pseudogene [Source:HGNC Symbol;Acc:HGNC:43640] | -2.0036 | 0.03269 |
| ENSG00000101276 | SLC52A3 | solute carrier family 52 member 3 [Source:HGNC Symbol;Acc:HGNC:16187] | -1.7111 | 0.03273 |
| ENSG00000188095 | MESP2 | mesoderm posterior bHLH transcription factor 2 [Source:HGNC Symbol;Acc:HGNC:29659] | -3.0855 | 0.03274 |
| ENSG00000254933 | NA | novel transcript | -1.5308 | 0.03275 |
| ENSG00000170425 | ADORA2B | adenosine A2b receptor [Source:HGNC Symbol;Acc:HGNC:264] | 1.3194 | 0.03285 |
| ENSG00000229083 | PSMA6P2 | proteasome subunit alpha 6 pseudogene 2 [Source:HGNC Symbol;Acc:HGNC:39607] | 0.9936 | 0.03288 |
| ENSG00000230991 | NA | novel transcript | -1.0262 | 0.03291 |
| ENSG00000160219 | GAB3 | GRB2 associated binding protein 3 [Source:HGNC Symbol;Acc:HGNC:17515] | -0.7410 | 0.03292 |
| ENSG00000128512 | DOCK4 | dedicator of cytokinesis 4 [Source:HGNC Symbol;Acc:HGNC:19192] | -0.7671 | 0.03299 |
| ENSG00000286305 | NA | novel transcript, antisense to TRRAP | 1.1613 | 0.03311 |
| ENSG00000261420 | NA | novel transcript, antisense to BRP44L | -1.0007 | 0.03311 |
| ENSG00000279278 | NA | novel transcript | -0.6791 | 0.03313 |
| ENSG00000174500 | GCSAM | germinal center associated signaling and motility [Source:HGNC Symbol;Acc:HGNC:20253] | 0.8963 | 0.03332 |
| ENSG00000269951 | NA | novel transcript, sense intronic to TSPAN3 | 0.6854 | 0.03338 |
| ENSG00000159899 | 2,00 NPR | natriuretic peptide receptor 2 [Source:HGNC Symbol;Acc:HGNC:7944] | -0.5729 | 0.03338 |
| ENSG00000092051 | JPH4 | junctophilin 4 [Source:HGNC Symbol;Acc:HGNC:20156] | -0.6832 | 0.03348 |
| ENSG00000245164 | LINC00861 | long intergenic non-protein coding RNA 861 [Source:HGNC Symbol;Acc:HGNC:45133] | -0.8778 | 0.03351 |
| ENSG00000251034 | NA | novel transcript | 1.7393 | 0.03361 |
| ENSG00000250075 | NA | novel transcript | -1.9986 | 0.03362 |
| ENSG00000137841 | PLCB2 | phospholipase C beta 2 [Source:HGNC Symbol;Acc:HGNC:9055] | -3.3912 | 0.03366 |
| ENSG00000080224 | EPHA6 | EPH receptor A6 [Source:HGNC Symbol;Acc:HGNC:19296] | -0.9029 | 0.03372 |
| ENSG00000103335 | PIEZO1 | piezo type mechanosensitive ion channel component 1 [Source:HGNC Symbol;Acc:HGNC:28993] | 0.8970 | 0.03372 |
| ENSG00000224383 | PRR29 | proline rich 29 [Source:HGNC Symbol;Acc:HGNC:25673] | -1.1651 | 0.03376 |
| ENSG00000146677 | NA | ribosomal protein L32 (RPL32) pseudogene | 0.5398 | 0.03378 |
| ENSG00000272606 | NA | novel transcript | -0.9127 | 0.03398 |
| ENSG00000279196 | NA | novel transcript | 0.7041 | 0.03398 |
| ENSG00000133116 | KL | klotho [Source:HGNC Symbol;Acc:HGNC:6344] | 2.4353 | 0.03398 |
| ENSG00000163814 | CDCP1 | CUB domain containing protein 1 [Source:HGNC Symbol;Acc:HGNC:24357] | 1.6348 | 0.03403 |
| ENSG00000152217 | SETBP1 | SET binding protein 1 [Source:HGNC Symbol;Acc:HGNC:15573] | 2.7308 | 0.03413 |
| ENSG00000011422 | PLAUR | plasminogen activator, urokinase receptor [Source:HGNC Symbol;Acc:HGNC:9053] | -0.6703 | 0.03416 |
| ENSG00000118292 | C1orf54 | chromosome 1 open reading frame 54 [Source:HGNC Symbol;Acc:HGNC:26258] | 0.6964 | 0.03426 |
| ENSG00000228440 | MTND5P6 | MT-ND5 pseudogene 6 [Source:HGNC Symbol;Acc:HGNC:42268] | -1.6235 | 0.03431 |
| ENSG00000283095 | NA | novel transcript | -1.0184 | 0.03437 |
| ENSG00000169621 | APLF | aprataxin and PNKP like factor [Source:HGNC Symbol;Acc:HGNC:28724] | -0.8908 | 0.03441 |
| ENSG00000279154 | NA | tec | -0.8871 | 0.03455 |
| ENSG00000253320 | AZIN1-AS1 | AZIN1 antisense RNA 1 [Source:HGNC Symbol;Acc:HGNC:51558] | -0.9993 | 0.03457 |
| ENSG00000235946 | H2BW3P | H2B.W histone 3, pseudogene [Source:HGNC Symbol;Acc:HGNC:44390] | -1.1512 | 0.03474 |
| ENSG00000211689 | TRGC1 | T cell receptor gamma constant 1 [Source:HGNC Symbol;Acc:HGNC:12275] | -2.7972 | 0.03476 |
| ENSG00000237638 | LINC02245 | long intergenic non-protein coding RNA 2245 [Source:HGNC Symbol;Acc:HGNC:53134] | -1.1752 | 0.03488 |
| ENSG00000188523 | CFAP77 | cilia and flagella associated protein 77 [Source:HGNC Symbol;Acc:HGNC:33776] | -0.8244 | 0.03488 |
| ENSG00000197291 | RAMP2-AS1 | RAMP2 antisense RNA 1 [Source:HGNC Symbol;Acc:HGNC:44358] | -1.2995 | 0.03501 |

| | | | | |
|---|---|---|---|---|
| ENSG00000163219 | ARHGAP25 | Rho GTPase activating protein 25 [Source:HGNC Symbol;Acc:HGNC:28951] | 1.6509 | 0.03501 |
| ENSG00000165194 | PCDH19 | protocadherin 19 [Source:HGNC Symbol;Acc:HGNC:14270] | 0.8962 | 0.03501 |
| ENSG00000109771 | LRP2BP | LRP2 binding protein [Source:HGNC Symbol;Acc:HGNC:25434] | 1.0624 | 0.03508 |
| ENSG00000263400 | TMEM220-AS1 | TMEM220 antisense RNA 1 [Source:HGNC Symbol;Acc:HGNC:44357] | -1.3700 | 0.03515 |
| ENSG00000247240 | UBL7-AS1 | UBL7 antisense RNA 1 (head to head) [Source:HGNC Symbol;Acc:HGNC:48614] | -0.5700 | 0.03517 |
| ENSG00000286098 | NA | zinc finger protein 844 [Source:NCBI gene (formerly Entrezgene);Acc:284391] | -1.1533 | 0.03517 |
| ENSG00000169918 | OTUD7A | OTU deubiquitinase 7A [Source:HGNC Symbol;Acc:HGNC:20718] | 2.9793 | 0.03518 |
| ENSG00000225766 | DHRS4L1 | dehydrogenase/reductase 4 like 1 [Source:HGNC Symbol;Acc:HGNC:19732] | -1.9073 | 0.03529 |
| ENSG00000269958 | NA | novel transcript, sense intronic to KLC1 | 0.5581 | 0.03535 |
| ENSG00000280138 | NA | novel transcript | 0.6412 | 0.03536 |
| ENSG00000170421 | KRT8 | keratin 8 [Source:HGNC Symbol;Acc:HGNC:6446] | 0.5328 | 0.03566 |
| ENSG00000260857 | NA | novel transcript | -1.1796 | 0.03588 |
| ENSG00000287261 | NA | novel transcript, antisense to HPS1 | 1.6481 | 0.03594 |
| ENSG00000145087 | STXBP5L | syntaxin binding protein 5 like [Source:HGNC Symbol;Acc:HGNC:30757] | -0.5705 | 0.03601 |
| ENSG00000286340 | NA | novel transcript, antisense to PHIP | -0.7343 | 0.03608 |
| ENSG00000108702 | CCL1 | C-C motif chemokine ligand 1 [Source:HGNC Symbol;Acc:HGNC:10609] | 2.3137 | 0.03612 |
| ENSG00000248213 | CICP16 | capicua transcriptional repressor pseudogene 16 [Source:HGNC Symbol;Acc:HGNC:38550] | 0.5751 | 0.03623 |
| ENSG00000105866 | SP4 | Sp4 transcription factor [Source:HGNC Symbol;Acc:HGNC:11209] | 0.5582 | 0.03625 |
| ENSG00000232653 | GOLGA8N | golgin A8 family member N [Source:HGNC Symbol;Acc:HGNC:44405] | -0.5439 | 0.03632 |
| ENSG00000277050 | NA | novel transcript, antisense to FRMD6 | -1.8567 | 0.03636 |
| ENSG00000186301 | MST1P2 | macrophage stimulating 1 pseudogene 2 [Source:HGNC Symbol;Acc:HGNC:7383] | -0.7381 | 0.03637 |
| ENSG00000279762 | NA | novel transcript, antisense to PSMD11 | 1.9491 | 0.03646 |
| ENSG00000218418 | NA | Rho GTPase activating protein 21 (ARHGAP21) pseudogene | 1.1205 | 0.03650 |
| ENSG00000163633 | C4orf36 | chromosome 4 open reading frame 36 [Source:HGNC Symbol;Acc:HGNC:28386] | 1.4071 | 0.03658 |
| ENSG00000226763 | SRRM5 | serine/arginine repetitive matrix 5 [Source:HGNC Symbol;Acc:HGNC:37248] | 1.0419 | 0.03674 |
| ENSG00000197405 | C5AR1 | complement C5a receptor 1 [Source:HGNC Symbol;Acc:HGNC:1338] | 1.1483 | 0.03675 |
| ENSG00000288536 | NA | novel transcript | 1.6725 | 0.03684 |
| ENSG00000284292 | NA | novel protein, ARPC1A and ARPC1B readthrough | 1.0923 | 0.03692 |
| ENSG00000261924 | NA | novel transcript, antisense to RPTOR | 0.5807 | 0.03698 |
| ENSG00000154274 | C4orf19 | chromosome 4 open reading frame 19 [Source:HGNC Symbol;Acc:HGNC:25618] | -0.7236 | 0.03706 |
| ENSG00000254461 | NA | novel transcript | 2.1598 | 0.03707 |
| ENSG00000266079 | SNORA59B | small nucleolar RNA, H/ACA box 59B [Source:HGNC Symbol;Acc:HGNC:32658] | 0.8487 | 0.03715 |
| ENSG00000278952 | NA | TEC | 1.2006 | 0.03721 |
| ENSG00000197813 | NA | novel transcript | -0.8307 | 0.03721 |
| ENSG00000088826 | SMOX | spermine oxidase [Source:HGNC Symbol;Acc:HGNC:15862] | 0.7492 | 0.03740 |
| ENSG00000213866 | YBX1P10 | Y-box binding protein 1 pseudogene 10 [Source:HGNC Symbol;Acc:HGNC:42432] | 0.5435 | 0.03753 |
| ENSG00000278238 | NA | novel transcript | -0.7049 | 0.03791 |
| ENSG00000261008 | LINC01572 | long intergenic non-protein coding RNA 1572 [Source:HGNC Symbol;Acc:HGNC:51385] | -0.9303 | 0.03794 |
| ENSG00000183773 | AIFM3 | apoptosis inducing factor mitochondria associated 3 [Source:HGNC Symbol;Acc:HGNC:26398] | -0.8251 | 0.03796 |
| ENSG00000178146 | NA | glycerol kinase (GK) pseudogene | 0.9000 | 0.03797 |
| ENSG00000279407 | NA | TEC | 0.8517 | 0.03812 |
| ENSG00000106034 | CPED1 | cadherin like and PC-esterase domain containing 1 [Source:HGNC Symbol;Acc:HGNC:26159] | 0.7593 | 0.03818 |
| ENSG00000263006 | ROCK1P1 | Rho associated coiled-coil containing protein kinase 1 pseudogene 1 [Source:HGNC Symbol;Acc:HGNC:37832] | -0.8789 | 0.03829 |

| | | | | |
|---|---|---|---|---|
| ENSG00000138744 | NAAA | N-acylethanolamine acid amidase [Source:HGNC Symbol;Acc:HGNC:736] | -0.7960 | 0.03830 |
| ENSG00000164406 | LEAP2 | liver enriched antimicrobial peptide 2 [Source:HGNC Symbol;Acc:HGNC:29571] | 0.6182 | 0.03841 |
| ENSG00000144644 | GADL1 | glutamate decarboxylase like 1 [Source:HGNC Symbol;Acc:HGNC:27949] | 1.4318 | 0.03843 |
| ENSG00000242193 | CRYZL2P | crystallin zeta like 2, pseudogene [Source:HGNC Symbol;Acc:HGNC:52164] | -0.5653 | 0.03848 |
| ENSG00000183346 | CABCOCO1 | ciliary associated calcium binding coiled-coil 1 [Source:HGNC Symbol;Acc:HGNC:28678] | -1.1541 | 0.03861 |
| ENSG00000230876 | LINC00486 | long intergenic non-protein coding RNA 486 [Source:HGNC Symbol;Acc:HGNC:42946] | -1.1249 | 0.03861 |
| ENSG00000172476 | RAB40A | RAB40A, member RAS oncogene family [Source:HGNC Symbol;Acc:HGNC:18283] | 0.5780 | 0.03881 |
| ENSG00000174628 | IQCK | IQ motif containing K [Source:HGNC Symbol;Acc:HGNC:28556] | -0.6669 | 0.03881 |
| ENSG00000215375 | MYL5 | myosin light chain 5 [Source:HGNC Symbol;Acc:HGNC:7586] | -0.6794 | 0.03884 |
| ENSG00000165879 | FRAT1 | FRAT regulator of WNT signaling pathway 1 [Source:HGNC Symbol;Acc:HGNC:3944] | -0.5666 | 0.03885 |
| ENSG00000204802 | NA | novel transcript | 0.7390 | 0.03885 |
| ENSG00000181826 | RELL1 | RELT like 1 [Source:HGNC Symbol;Acc:HGNC:27379] | -0.7279 | 0.03887 |
| ENSG00000105479 | CCDC114 | coiled-coil domain containing 114 [Source:HGNC Symbol;Acc:HGNC:26560] | -0.9466 | 0.03896 |
| ENSG00000099994 | SUSD2 | sushi domain containing 2 [Source:HGNC Symbol;Acc:HGNC:30667] | 1.0839 | 0.03900 |
| ENSG00000235545 | NA | novel transcript | -2.8457 | 0.03901 |
| ENSG00000238121 | LINC00426 | long intergenic non-protein coding RNA 426 [Source:HGNC Symbol;Acc:HGNC:42761] | -2.5100 | 0.03929 |
| ENSG00000185332 | NA | TMEM105 long non-coding RNA [Source:NCBI gene (formerly Entrezgene);Acc:284186] | -2.6868 | 0.03941 |
| ENSG00000143127 | ITGA10 | integrin subunit alpha 10 [Source:HGNC Symbol;Acc:HGNC:6135] | 1.0821 | 0.03947 |
| ENSG00000279530 | NA | TEC | 0.6393 | 0.03950 |
| ENSG00000236297 | NA | eukaryotic translation elongation factor 1 alpha 1 (EEF1A1) pseudogene | 1.3566 | 0.03951 |
| ENSG00000185261 | KIAA0825 | KIAA0825 [Source:HGNC Symbol;Acc:HGNC:28532] | -0.6330 | 0.03975 |
| ENSG00000215595 | C20orf202 | chromosome 20 open reading frame 202 [Source:HGNC Symbol;Acc:HGNC:37254] | -1.3675 | 0.03976 |
| ENSG00000104894 | CD37 | CD37 molecule [Source:HGNC Symbol;Acc:HGNC:1666] | -0.6036 | 0.03976 |
| ENSG00000185347 | TEDC1 | tubulin epsilon and delta complex 1 [Source:HGNC Symbol;Acc:HGNC:20127] | -0.5515 | 0.03992 |
| ENSG00000196436 | NPIPB15 | nuclear pore complex interacting protein family member B15 [Source:HGNC Symbol;Acc:HGNC:34409] | 0.5628 | 0.03994 |
| ENSG00000244094 | SPRR2F | small proline rich protein 2F [Source:HGNC Symbol;Acc:HGNC:11266] | -1.7383 | 0.03997 |
| ENSG00000279232 | NA | novel transcript, antisense to RGMB | -0.9326 | 0.03997 |
| ENSG00000144406 | UNC80 | unc-80 homolog, NALCN channel complex subunit [Source:HGNC Symbol;Acc:HGNC:26582] | -1.1881 | 0.04004 |
| ENSG00000253540 | FAM86HP | family with sequence similarity 86 member H, pseudogene [Source:HGNC Symbol;Acc:HGNC:42359] | 1.0014 | 0.04012 |
| ENSG00000132334 | PTPRE | protein tyrosine phosphatase receptor type E [Source:HGNC Symbol;Acc:HGNC:9669] | 0.8852 | 0.04019 |
| ENSG00000168959 | GRM5 | glutamate metabotropic receptor 5 [Source:HGNC Symbol;Acc:HGNC:4597] | -1.2994 | 0.04027 |
| ENSG00000271576 | NA | novel transcript | -0.6539 | 0.04028 |
| ENSG00000150637 | CD226 | CD226 molecule [Source:HGNC Symbol;Acc:HGNC:16961] | 1.1601 | 0.04033 |
| ENSG00000255502 | LINC02713 | long intergenic non-protein coding RNA 2713 [Source:HGNC Symbol;Acc:HGNC:54230] | -1.9621 | 0.04050 |
| ENSG00000155465 | SLC7A7 | solute carrier family 7 member 7 [Source:HGNC Symbol;Acc:HGNC:11065] | -1.1754 | 0.04052 |
| ENSG00000170827 | CELP | carboxyl ester lipase pseudogene [Source:HGNC Symbol;Acc:HGNC:1849] | 2.3636 | 0.04052 |
| ENSG00000250038 | NA | novel transcript | 0.5055 | 0.04053 |
| ENSG00000255441 | SIGLEC10-AS1 | SIGLEC10 antisense RNA 1 [Source:HGNC Symbol;Acc:HGNC:40719] | -1.2217 | 0.04062 |
| ENSG00000260019 | LINC01992 | long intergenic non-protein coding RNA 1992 [Source:HGNC Symbol;Acc:HGNC:52824] | 0.6385 | 0.04066 |
| ENSG00000279518 | NA | TEC | 0.9048 | 0.04068 |
| ENSG00000229980 | TOB1-AS1 | TOB1 antisense RNA 1 [Source:HGNC Symbol;Acc:HGNC:44340] | -0.5633 | 0.04068 |
| ENSG00000261462 | NA | novel transcript | -0.9356 | 0.04069 |
| ENSG00000149781 | FERMT3 | fermitin family member 3 [Source:HGNC Symbol;Acc:HGNC:23151] | 0.9035 | 0.04076 |

| | | | | |
|---|---|---|---|---|
| ENSG00000180152 | XIAPP3 | X-linked inhibitor of apoptosis pseudogene 3 [Source:HGNC Symbol;Acc:HGNC:52377] | -1.3506 | 0.04079 |
| ENSG00000174827 | PDZK1 | PDZ domain containing 1 [Source:HGNC Symbol;Acc:HGNC:8821] | -0.8223 | 0.04081 |
| ENSG00000198756 | COLGALT2 | collagen beta(1-O)galactosyltransferase 2 [Source:HGNC Symbol;Acc:HGNC:16790] | -0.5963 | 0.04084 |
| ENSG00000135835 | KIAA1614 | KIAA1614 [Source:HGNC Symbol;Acc:HGNC:29327] | 1.1904 | 0.04087 |
| ENSG00000120669 | SOHLH2 | spermatogenesis and oogenesis specific basic helix-loop-helix 2 [Source:HGNC Symbol;Acc:HGNC:26026] | -0.8517 | 0.04099 |
| ENSG00000184524 | CEND1 | cell cycle exit and neuronal differentiation 1 [Source:HGNC Symbol;Acc:HGNC:24153] | -0.5219 | 0.04108 |
| ENSG00000169247 | SH3TC2 | SH3 domain and tetratricopeptide repeats 2 [Source:HGNC Symbol;Acc:HGNC:29427] | 2.0738 | 0.04109 |
| ENSG00000232472 | EEF1B2P3 | eukaryotic translation elongation factor 1 beta 2 pseudogene 3 [Source:HGNC Symbol;Acc:HGNC:3210] | 0.6485 | 0.04109 |
| ENSG00000072682 | P4HA2 | prolyl 4-hydroxylase subunit alpha 2 [Source:HGNC Symbol;Acc:HGNC:8547] | 0.5424 | 0.04116 |
| ENSG00000260585 | NA | novel transcript | -1.2059 | 0.04126 |
| ENSG00000204581 | ACOXL-AS1 | ACOXL antisense RNA 1 [Source:HGNC Symbol;Acc:HGNC:41112] | -1.7450 | 0.04127 |
| ENSG00000260735 | NA | phospholipase A2 group X (PLA2G10) pseudogene | -1.5878 | 0.04133 |
| ENSG00000267707 | NA | novel transcript, antisense to CELF4 | -1.9779 | 0.04134 |
| ENSG00000242781 | LINC02050 | long intergenic non-protein coding RNA 2050 [Source:HGNC Symbol;Acc:HGNC:52890] | -0.9545 | 0.04146 |
| ENSG00000100024 | UPB1 | beta-ureidopropionase 1 [Source:HGNC Symbol;Acc:HGNC:16297] | -1.3064 | 0.04148 |
| ENSG00000171044 | XKR6 | XK related 6 [Source:HGNC Symbol;Acc:HGNC:27806] | 0.8500 | 0.04152 |
| ENSG00000162676 | GFI1 | growth factor independent 1 transcriptional repressor [Source:HGNC Symbol;Acc:HGNC:4237] | 1.2516 | 0.04157 |
| ENSG00000170775 | GPR37 | G protein-coupled receptor 37 [Source:HGNC Symbol;Acc:HGNC:4494] | -1.2760 | 0.04162 |
| ENSG00000214837 | LINC01347 | long intergenic non-protein coding RNA 1347 [Source:HGNC Symbol;Acc:HGNC:50566] | -0.6874 | 0.04177 |
| ENSG00000175130 | MARCKSL1 | MARCKS like 1 [Source:HGNC Symbol;Acc:HGNC:7142] | 0.6180 | 0.04180 |
| ENSG00000111886 | GABRR2 | gamma-aminobutyric acid type A receptor subunit rho2 [Source:HGNC Symbol;Acc:HGNC:4091] | 0.8839 | 0.04187 |
| ENSG00000286535 | NA | novel transcript, antisense to ASAP1 | -0.8486 | 0.04188 |
| ENSG00000251566 | HMGB1P35 | high mobility group box 1 pseudogene 35 [Source:HGNC Symbol;Acc:HGNC:39175] | 4.2283 | 0.04191 |
| ENSG00000255052 | FAM66D | family with sequence similarity 66 member D [Source:HGNC Symbol;Acc:HGNC:24159] | -0.9514 | 0.04196 |
| ENSG00000273064 | NA | novel transcript | -1.9694 | 0.04197 |
| ENSG00000197565 | COL4A6 | collagen type IV alpha 6 chain [Source:HGNC Symbol;Acc:HGNC:2208] | 2.6079 | 0.04216 |
| ENSG00000271086 | NAMA | non-protein coding RNA, associated with MAP kinase pathway and growth arrest [Source:HGNC Symbol;Acc:HGNC:42408] | 1.5806 | 0.04244 |
| ENSG00000248727 | LINC01948 | long intergenic non-protein coding RNA 1948 [Source:HGNC Symbol;Acc:HGNC:52772] | -1.7656 | 0.04250 |
| ENSG00000278917 | NA | TEC | -0.7585 | 0.04259 |
| ENSG00000168209 | DDIT4 | DNA damage inducible transcript 4 [Source:HGNC Symbol;Acc:HGNC:24944] | -0.6498 | 0.04274 |
| ENSG00000121769 | FABP3 | fatty acid binding protein 3 [Source:HGNC Symbol;Acc:HGNC:3557] | 0.7400 | 0.04289 |
| ENSG00000114805 | PLCH1 | phospholipase C eta 1 [Source:HGNC Symbol;Acc:HGNC:29185] | -1.7565 | 0.04293 |
| ENSG00000156096 | UGT2B4 | UDP glucuronosyltransferase family 2 member B4 [Source:HGNC Symbol;Acc:HGNC:12553] | -0.6250 | 0.04301 |
| ENSG00000286373 | NA | novel transcript | 1.7727 | 0.04312 |
| ENSG00000119121 | TRPM6 | transient receptor potential cation channel subfamily M member 6 [Source:HGNC Symbol;Acc:HGNC:17995] | 0.9409 | 0.04316 |
| ENSG00000168333 | PPDPFL | pancreatic progenitor cell differentiation and proliferation factor like [Source:HGNC Symbol;Acc:HGNC:31745] | -0.7030 | 0.04325 |
| ENSG00000272973 | NA | novel transcript, antisense to SLC2A11 | -0.8040 | 0.04332 |
| ENSG00000253842 | NA | novel transcript | 2.2691 | 0.04337 |
| ENSG00000116661 | FBXO2 | F-box protein 2 [Source:HGNC Symbol;Acc:HGNC:13581] | -0.5235 | 0.04344 |
| ENSG00000185681 | MORN5 | MORN repeat containing 5 [Source:HGNC Symbol;Acc:HGNC:17841] | -1.9514 | 0.04348 |
| ENSG00000168229 | PTGDR | prostaglandin D2 receptor [Source:HGNC Symbol;Acc:HGNC:9591] | -3.5457 | 0.04349 |

| | | | | |
|---|---|---|---|---|
| ENSG00000133246 | PRAM1 | PML-RARA regulated adaptor molecule 1 [Source:HGNC Symbol;Acc:HGNC:30091] | 1.8287 | 0.04354 |
| ENSG00000166924 | NYAP1 | neuronal tyrosine phosphorylated phosphoinositide-3-kinase adaptor 1 [Source:HGNC Symbol;Acc:HGNC:22009] | -0.5173 | 0.04362 |
| ENSG00000278514 | NA | novel transcript | -0.6974 | 0.04363 |
| ENSG00000077942 | FBLN1 | fibulin 1 [Source:HGNC Symbol;Acc:HGNC:3600] | -0.5502 | 0.04365 |
| ENSG00000224184 | MIR3681HG | MIR3681 host gene [Source:HGNC Symbol;Acc:HGNC:52001] | 1.4196 | 0.04366 |
| ENSG00000151491 | EPS8 | epidermal growth factor receptor pathway substrate 8 [Source:HGNC Symbol;Acc:HGNC:3420] | -0.5366 | 0.04408 |
| ENSG00000285679 | NA | novel transcript | -1.1567 | 0.04421 |
| ENSG00000118523 | CCN2 | cellular communication network factor 2 [Source:HGNC Symbol;Acc:HGNC:2500] | 2.4506 | 0.04421 |
| ENSG00000233932 | CTXN2 | cortexin 2 [Source:HGNC Symbol;Acc:HGNC:31109] | -0.8173 | 0.04421 |
| ENSG00000255725 | TDGP1 | thymine-DNA glycosylase pseudogene 1 [Source:HGNC Symbol;Acc:HGNC:49891] | 1.4083 | 0.04432 |
| ENSG00000178802 | MPI | mannose phosphate isomerase [Source:HGNC Symbol;Acc:HGNC:7216] | -0.5139 | 0.04438 |
| ENSG00000231829 | NA | novel transcript | -0.6008 | 0.04446 |
| ENSG00000037757 | MRI1 | methylthioribose-1-phosphate isomerase 1 [Source:HGNC Symbol;Acc:HGNC:28469] | 0.6002 | 0.04448 |
| ENSG00000068078 | FGFR3 | fibroblast growth factor receptor 3 [Source:HGNC Symbol;Acc:HGNC:3690] | -1.2468 | 0.04456 |
| ENSG00000183784 | DOCK8-AS1 | DOCK8 antisense RNA 1 [Source:HGNC Symbol;Acc:HGNC:26436] | 1.4978 | 0.04471 |
| ENSG00000215018 | COL28A1 | collagen type XXVIII alpha 1 chain [Source:HGNC Symbol;Acc:HGNC:22442] | -0.9621 | 0.04478 |
| ENSG00000267702 | NA | novel transcript, sense intronic C18orf1 | 1.7568 | 0.04522 |
| ENSG00000282907 | NA | novel transcript | -2.1307 | 0.04535 |
| ENSG00000214548 | MEG3 | maternally expressed 3 [Source:HGNC Symbol;Acc:HGNC:14575] | 0.5780 | 0.04573 |
| ENSG00000236532 | LINC01695 | long intergenic non-protein coding RNA 1695 [Source:HGNC Symbol;Acc:HGNC:52483] | 0.8214 | 0.04592 |
| ENSG00000260551 | PWRN2 | Prader-Willi region non-protein coding RNA 2 [Source:HGNC Symbol;Acc:HGNC:33236] | -2.7456 | 0.04608 |
| ENSG00000245017 | LINC02453 | long intergenic non-protein coding RNA 2453 [Source:HGNC Symbol;Acc:HGNC:53392] | -1.2696 | 0.04619 |
| ENSG00000257027 | NA | novel transcript | -0.6748 | 0.04642 |
| ENSG00000255292 | NA | novel transcript | 2.9353 | 0.04667 |
| ENSG00000258017 | NA | novel transcript, antisense to TUBA1B | -0.5222 | 0.04667 |
| ENSG00000214888 | NA | nuclear pore associated protein 1 (NPAP1) pseudogene | -1.0334 | 0.04672 |
| ENSG00000166960 | CCDC178 | coiled-coil domain containing 178 [Source:HGNC Symbol;Acc:HGNC:29588] | -1.0727 | 0.04678 |
| ENSG00000278341 | NA | novel transcript, antisense to CTU2 | 1.7164 | 0.04684 |
| ENSG00000157890 | MEGF11 | multiple EGF like domains 11 [Source:HGNC Symbol;Acc:HGNC:29635] | 0.6418 | 0.04692 |
| ENSG00000153157 | SYCP2L | synaptonemal complex protein 2 like [Source:HGNC Symbol;Acc:HGNC:21537] | -0.7411 | 0.04695 |
| ENSG00000241549 | GUSBP2 | GUSB pseudogene 2 [Source:HGNC Symbol;Acc:HGNC:18792] | -0.8295 | 0.04700 |
| ENSG00000288187 | NA | novel transcript | -0.9187 | 0.04700 |
| ENSG00000183929 | DUSP5P1 | dual specificity phosphatase 5 pseudogene 1 [Source:HGNC Symbol;Acc:HGNC:32020] | -1.2477 | 0.04706 |
| ENSG00000272146 | ARF4-AS1 | ARF4 antisense RNA 1 [Source:HGNC Symbol;Acc:HGNC:51593] | -1.1668 | 0.04711 |
| ENSG00000253438 | PCAT1 | prostate cancer associated transcript 1 [Source:HGNC Symbol;Acc:HGNC:43022] | -0.9727 | 0.04720 |
| ENSG00000196968 | FUT11 | fucosyltransferase 11 [Source:HGNC Symbol;Acc:HGNC:19233] | 0.6463 | 0.04741 |
| ENSG00000261684 | NA | novel transcript, antisense to SNAP23 | 1.3900 | 0.04741 |
| ENSG00000243055 | GK-AS1 | GK antisense RNA 1 [Source:HGNC Symbol;Acc:HGNC:40255] | 2.1048 | 0.04744 |
| ENSG00000254827 | SLC22A18AS | solute carrier family 22 member 18 antisense [Source:HGNC Symbol;Acc:HGNC:10965] | -1.1138 | 0.04748 |
| ENSG00000159212 | CLIC6 | chloride intracellular channel 6 [Source:HGNC Symbol;Acc:HGNC:2065] | -0.8534 | 0.04750 |
| ENSG00000159958 | TNFRSF13C | TNF receptor superfamily member 13C [Source:HGNC Symbol;Acc:HGNC:17755] | -0.8015 | 0.04763 |
| ENSG00000220891 | NA | uncharacterized LOC648691 [Source:NCBI gene (formerly Entrezgene);Acc:648691] | -2.2308 | 0.04770 |
| ENSG00000173451 | THAP2 | THAP domain containing 2 [Source:HGNC Symbol;Acc:HGNC:20854] | 0.5842 | 0.04773 |

| | | | | |
|---|---|---|---|---|
| ENSG00000254198 | NA | Family with sequence similarity 9, member C (FAM9C) pseudogene | 3.1289 | 0.04802 |
| ENSG00000268658 | LINC00664 | long intergenic non-protein coding RNA 664 [Source:HGNC Symbol;Acc:HGNC:44319] | -1.4927 | 0.04806 |
| ENSG00000102271 | KLHL4 | kelch like family member 4 [Source:HGNC Symbol;Acc:HGNC:6355] | -0.8561 | 0.04807 |
| ENSG00000259657 | PIGHP1 | phosphatidylinositol glycan anchor biosynthesis class H pseudogene 1 [Source:HGNC Symbol;Acc:HGNC:45152] | 1.5221 | 0.04810 |
| ENSG00000163071 | SPATA18 | spermatogenesis associated 18 [Source:HGNC Symbol;Acc:HGNC:29579] | -1.2276 | 0.04812 |
| ENSG00000115414 | FN1 | fibronectin 1 [Source:HGNC Symbol;Acc:HGNC:3778] | 0.6321 | 0.04815 |
| ENSG00000267058 | NA | novel transcript | -0.8287 | 0.04815 |
| ENSG00000257869 | NA | novel transcript | -1.0422 | 0.04823 |
| ENSG00000138172 | CALHM2 | calcium homeostasis modulator family member 2 [Source:HGNC Symbol;Acc:HGNC:23493] | 0.5760 | 0.04823 |
| ENSG00000270179 | NA | novel transcript | -1.5446 | 0.04834 |
| ENSG00000285863 | NA | novel transcript, antisense to VPS13C | 1.0928 | 0.04852 |
| ENSG00000086730 | LAT2 | linker for activation of T cells family member 2 [Source:HGNC Symbol;Acc:HGNC:12749] | 0.8863 | 0.04852 |
| ENSG00000132832 | NA | novel transcript | -2.1848 | 0.04862 |
| ENSG00000225544 | NA | pseudogene similar to part of metallophosphoesterase 1 (MPPE1) | -1.7388 | 0.04862 |
| ENSG00000283913 | NA | novel transcript | 1.3501 | 0.04864 |
| ENSG00000288253 | NA | novel transcript | -1.2559 | 0.04864 |
| ENSG00000142661 | MYOM3 | myomesin 3 [Source:HGNC Symbol;Acc:HGNC:26679] | -1.5287 | 0.04869 |
| ENSG00000009724 | MASP2 | mannan binding lectin serine peptidase 2 [Source:HGNC Symbol;Acc:HGNC:6902] | 0.8479 | 0.04874 |
| ENSG00000250995 | NA | novel pseudogene | 1.3780 | 0.04885 |
| ENSG00000149527 | PLCH2 | phospholipase C eta 2 [Source:HGNC Symbol;Acc:HGNC:29037] | 1.6965 | 0.04896 |
| ENSG00000159184 | HOXB13 | homeobox B13 [Source:HGNC Symbol;Acc:HGNC:5112] | -0.5955 | 0.04896 |
| ENSG00000105695 | MAG | myelin associated glycoprotein [Source:HGNC Symbol;Acc:HGNC:6783] | -3.0411 | 0.04902 |
| ENSG00000272021 | NA | novel transcript | -1.6514 | 0.04916 |
| ENSG00000135083 | CCNJL | cyclin J like [Source:HGNC Symbol;Acc:HGNC:25876] | -0.5959 | 0.04968 |
| ENSG00000156265 | MAP3K7CL | MAP3K7 C-terminal like [Source:HGNC Symbol;Acc:HGNC:16457] | 0.7280 | 0.04972 |
| ENSG00000232093 | DCST1-AS1 | DCST1 antisense RNA 1 [Source:HGNC Symbol;Acc:HGNC:41147] | -0.8169 | 0.04977 |
| ENSG00000161860 | SYCE2 | synaptonemal complex central element protein 2 [Source:HGNC Symbol;Acc:HGNC:27411] | -0.7471 | 0.04980 |
| ENSG00000239626 | RPSAP41 | ribosomal protein SA pseudogene 41 [Source:HGNC Symbol;Acc:HGNC:36663] | -0.9212 | 0.04981 |
| ENSG00000242220 | TCP10L | t-complex 10 like [Source:HGNC Symbol;Acc:HGNC:11657] | 1.3449 | 0.04990 |
