## Supplemental Table 5 for "Modulation of calcium signaling on demand to decipher the molecular mechanisms of primary aldosteronism"

**Supplementary Table 5. List of differentially expressed genes in H295R_S2 cells expressing the α7-5HT3 receptor in response to 8h treatment with 12mM K+**

| Ensembl Gene ID | Gene Symbol | Gene Name | Log2 Fold Change | pAdj |
|---|---|---|---|---|
| ENSG00000155827 | RNF20 | ring finger protein 20 [Source:HGNC Symbol;Acc:HGNC:10062] | 1.4926 | 1.86E-36 |
| ENSG00000116514 | RNF19B | ring finger protein 19B [Source:HGNC Symbol;Acc:HGNC:26886] | 1.8598 | 1.60E-31 |
| ENSG00000101230 | ISM1 | isthmin 1 [Source:HGNC Symbol;Acc:HGNC:16213] | -1.6114 | 7.76E-29 |
| ENSG00000101605 | MYOM1 | myomesin 1 [Source:HGNC Symbol;Acc:HGNC:7613] | 2.1157 | 2.93E-27 |
| ENSG00000196782 | MAML3 | mastermind like transcriptional coactivator 3 [Source:HGNC Symbol;Acc:HGNC:16272] | 1.0236 | 7.04E-27 |
| ENSG00000151835 | SACS | sacsin molecular chaperone [Source:HGNC Symbol;Acc:HGNC:10519] | 1.1059 | 2.92E-26 |
| ENSG00000196352 | CD55 | CD55 molecule (Cromer blood group) [Source:HGNC Symbol;Acc:HGNC:2665] | 1.3377 | 1.37E-25 |
| ENSG00000076716 | GPC4 | glypican 4 [Source:HGNC Symbol;Acc:HGNC:4452] | 1.4109 | 8.19E-24 |
| ENSG00000006468 | ETV1 | ETS variant transcription factor 1 [Source:HGNC Symbol;Acc:HGNC:3490] | 1.1826 | 4.58E-23 |
| ENSG00000081181 | ARG2 | arginase 2 [Source:HGNC Symbol;Acc:HGNC:664] | 1.4894 | 5.23E-22 |
| ENSG00000165152 | PGAP4 | post-GPI attachment to proteins GalNAc transferase 4 [Source:HGNC Symbol;Acc:HGNC:28180] | 0.8478 | 1.85E-21 |
| ENSG00000143153 | ATP1B1 | ATPase Na+/K+ transporting subunit beta 1 [Source:HGNC Symbol;Acc:HGNC:804] | 1.1310 | 3.01E-21 |
| ENSG00000138759 | FRAS1 | Fraser extracellular matrix complex subunit 1 [Source:HGNC Symbol;Acc:HGNC:19185] | 1.7230 | 4.17E-21 |
| ENSG00000150893 | FREM2 | FRAS1 related extracellular matrix 2 [Source:HGNC Symbol;Acc:HGNC:25396] | 1.4529 | 5.69E-21 |
| ENSG00000111424 | VDR | vitamin D receptor [Source:HGNC Symbol;Acc:HGNC:12679] | 1.4224 | 8.01E-21 |
| ENSG00000122012 | SV2C | synaptic vesicle glycoprotein 2C [Source:HGNC Symbol;Acc:HGNC:30670] | 1.5320 | 1.56E-18 |
| ENSG00000109689 | STIM2 | stromal interaction molecule 2 [Source:HGNC Symbol;Acc:HGNC:19205] | 1.3655 | 2.06E-17 |
| ENSG00000228536 | LYPLAL1-AS1 | LYPLAL1 antisense RNA 1 [Source:HGNC Symbol;Acc:HGNC:54054] | 1.6180 | 3.45E-17 |
| ENSG00000137575 | SDCBP | syndecan binding protein [Source:HGNC Symbol;Acc:HGNC:10662] | 1.0099 | 5.20E-17 |
| ENSG00000176697 | BDNF | brain derived neurotrophic factor [Source:HGNC Symbol;Acc:HGNC:1033] | 1.9516 | 5.20E-17 |
| ENSG00000108946 | PRKAR1A | protein kinase cAMP-dependent type I regulatory subunit alpha [Source:HGNC Symbol;Acc:HGNC:9388] | 0.5418 | 9.49E-17 |
| ENSG00000119698 | PPP4R4 | protein phosphatase 4 regulatory subunit 4 [Source:HGNC Symbol;Acc:HGNC:23788] | 1.6406 | 1.38E-16 |
| ENSG00000101445 | PPP1R16B | protein phosphatase 1 regulatory subunit 16B [Source:HGNC Symbol;Acc:HGNC:15850] | 0.9690 | 1.67E-16 |
| ENSG00000151276 | MAGI1 | membrane associated guanylate kinase, WW and PDZ domain containing 1 [Source:HGNC Symbol;Acc:HGNC:946] | 0.6187 | 2.28E-16 |
| ENSG00000184349 | EFNA5 | ephrin A5 [Source:HGNC Symbol;Acc:HGNC:3225] | 1.9795 | 5.65E-16 |
| ENSG00000068878 | PSME4 | proteasome activator subunit 4 [Source:HGNC Symbol;Acc:HGNC:20635] | 0.6697 | 8.66E-16 |
| ENSG00000177875 | CCDC184 | coiled-coil domain containing 184 [Source:HGNC Symbol;Acc:HGNC:33749] | 1.3884 | 3.43E-15 |
| ENSG00000125520 | SLC2A4RG | SLC2A4 regulator [Source:HGNC Symbol;Acc:HGNC:15930] | -0.9230 | 9.75E-15 |
| ENSG00000073969 | NSF | N-ethylmaleimide sensitive factor, vesicle fusing ATPase [Source:HGNC Symbol;Acc:HGNC:8016] | 0.9535 | 1.03E-14 |
| ENSG00000010818 | HIVEP2 | HIVEP zinc finger 2 [Source:HGNC Symbol;Acc:HGNC:4921] | 1.0762 | 1.86E-14 |
| ENSG00000139154 | AEBP2 | AE binding protein 2 [Source:HGNC Symbol;Acc:HGNC:24051] | 0.6487 | 3.17E-14 |
| ENSG00000162745 | OLFML2B | olfactomedin like 2B [Source:HGNC Symbol;Acc:HGNC:24558] | -2.0276 | 1.41E-13 |
| ENSG00000196586 | MYO6 | myosin VI [Source:HGNC Symbol;Acc:HGNC:7605] | 1.0729 | 1.90E-13 |
| ENSG00000108176 | DNAJC12 | DnaJ heat shock protein family (Hsp40) member C12 [Source:HGNC Symbol;Acc:HGNC:28908] | 1.4285 | 2.62E-13 |
| ENSG00000135605 | TEC | tec protein tyrosine kinase [Source:HGNC Symbol;Acc:HGNC:11719] | 0.6312 | 2.62E-13 |
| ENSG00000188517 | COL25A1 | collagen type XXV alpha 1 chain [Source:HGNC Symbol;Acc:HGNC:18603] | 0.9155 | 1.12E-12 |
| ENSG00000104043 | ATP8B4 | ATPase phospholipid transporting 8B4 (putative) [Source:HGNC Symbol;Acc:HGNC:13536] | 1.8232 | 1.21E-12 |
| ENSG00000113231 | PDE8B | phosphodiesterase 8B [Source:HGNC Symbol;Acc:HGNC:8794] | 1.4642 | 1.61E-12 |
| ENSG00000197635 | DPP4 | dipeptidyl peptidase 4 [Source:HGNC Symbol;Acc:HGNC:3009] | 0.9502 | 1.83E-12 |
| ENSG00000198597 | ZNF536 | zinc finger protein 536 [Source:HGNC Symbol;Acc:HGNC:29025] | 0.9491 | 1.41E-11 |
| ENSG00000108861 | DUSP3 | dual specificity phosphatase 3 [Source:HGNC Symbol;Acc:HGNC:3069] | 0.5075 | 1.42E-11 |
| ENSG00000167528 | ZNF641 | zinc finger protein 641 [Source:HGNC Symbol;Acc:HGNC:31834] | 1.1764 | 1.85E-11 |
| ENSG00000058866 | DGKG | diacylglycerol kinase gamma [Source:HGNC Symbol;Acc:HGNC:2853] | 0.7290 | 2.46E-11 |
| ENSG00000235431 | NA | novel transcript | 1.2414 | 2.84E-11 |
| ENSG00000008405 | CRY1 | cryptochrome circadian regulator 1 [Source:HGNC Symbol;Acc:HGNC:2384] | 0.8020 | 3.18E-11 |
| ENSG00000091428 | RAPGEF4 | Rap guanine nucleotide exchange factor 4 [Source:HGNC Symbol;Acc:HGNC:16626] | 1.9233 | 3.60E-11 |
| ENSG00000152049 | KCNE4 | potassium voltage-gated channel subfamily E regulatory subunit 4 [Source:HGNC Symbol;Acc:HGNC:6244] | 1.3644 | 3.62E-11 |
| ENSG00000101974 | ATP11C | ATPase phospholipid transporting 11C [Source:HGNC Symbol;Acc:HGNC:13554] | 1.0795 | 5.70E-11 |
| ENSG00000065183 | WDR3 | WD repeat domain 3 [Source:HGNC Symbol;Acc:HGNC:12755] | -0.6249 | 7.60E-11 |
| ENSG00000070061 | ELP1 | elongator complex protein 1 [Source:HGNC Symbol;Acc:HGNC:5959] | 0.9378 | 7.97E-11 |
| ENSG00000060656 | PTPRU | protein tyrosine phosphatase receptor type U [Source:HGNC Symbol;Acc:HGNC:9683] | -0.7799 | 7.99E-11 |

| | | | | |
|---|---|---|---|---|
| ENSG00000163431 | LMOD1 | leiomodin 1 [Source:HGNC Symbol;Acc:HGNC:6647] | -1.3127 | 8.72E-11 |
| ENSG00000092068 | SLC7A8 | solute carrier family 7 member 8 [Source:HGNC Symbol;Acc:HGNC:11066] | -0.8163 | 1.15E-10 |
| ENSG00000138767 | CNOT6L | CCR4-NOT transcription complex subunit 6 like [Source:HGNC Symbol;Acc:HGNC:18042] | 0.6950 | 1.18E-10 |
| ENSG00000185231 | MC2R | melanocortin 2 receptor [Source:HGNC Symbol;Acc:HGNC:6930] | 2.5064 | 1.25E-10 |
| ENSG00000244405 | ETV5 | ETS variant transcription factor 5 [Source:HGNC Symbol;Acc:HGNC:3494] | 1.8231 | 1.48E-10 |
| ENSG00000254615 | NA | novel transcript | 2.2497 | 2.29E-10 |
| ENSG00000128283 | CDC42EP1 | CDC42 effector protein 1 [Source:HGNC Symbol;Acc:HGNC:17014] | -0.6410 | 2.45E-10 |
| ENSG00000165434 | PGM2L1 | phosphoglucomutase 2 like 1 [Source:HGNC Symbol;Acc:HGNC:20898] | 1.3519 | 3.42E-10 |
| ENSG00000270757 | HSPE1-MOB4 | HSPE1-MOB4 readthrough [Source:HGNC Symbol;Acc:HGNC:49184] | 0.9324 | 3.51E-10 |
| ENSG00000090530 | P3H2 | prolyl 3-hydroxylase 2 [Source:HGNC Symbol;Acc:HGNC:19317] | 0.8498 | 3.91E-10 |
| ENSG00000072110 | ACTN1 | actinin alpha 1 [Source:HGNC Symbol;Acc:HGNC:163] | 0.8648 | 4.26E-10 |
| ENSG00000090776 | EFNB1 | ephrin B1 [Source:HGNC Symbol;Acc:HGNC:3226] | -0.7746 | 5.26E-10 |
| ENSG00000150594 | ADRA2A | adrenoceptor alpha 2A [Source:HGNC Symbol;Acc:HGNC:281] | -1.5015 | 6.97E-10 |
| ENSG00000183654 | MARCHF11 | membrane associated ring-CH-type finger 11 [Source:HGNC Symbol;Acc:HGNC:33609] | 1.0730 | 7.17E-10 |
| ENSG00000119927 | GPAM | glycerol-3-phosphate acyltransferase, mitochondrial [Source:HGNC Symbol;Acc:HGNC:24865] | -0.7818 | 8.66E-10 |
| ENSG00000157514 | TSC22D3 | TSC22 domain family member 3 [Source:HGNC Symbol;Acc:HGNC:3051] | 0.7115 | 9.95E-10 |
| ENSG00000092445 | TYRO3 | TYRO3 protein tyrosine kinase [Source:HGNC Symbol;Acc:HGNC:12446] | 0.6333 | 1.06E-09 |
| ENSG00000126368 | NR1D1 | nuclear receptor subfamily 1 group D member 1 [Source:HGNC Symbol;Acc:HGNC:7962] | -0.8166 | 3.60E-09 |
| ENSG00000134602 | STK26 | serine/threonine kinase 26 [Source:HGNC Symbol;Acc:HGNC:18174] | 0.9882 | 3.60E-09 |
| ENSG00000109323 | MANBA | mannosidase beta [Source:HGNC Symbol;Acc:HGNC:6831] | 0.5989 | 3.75E-09 |
| ENSG00000158966 | CACHD1 | cache domain containing 1 [Source:HGNC Symbol;Acc:HGNC:29314] | 0.5903 | 4.38E-09 |
| ENSG00000165702 | GFI1B | growth factor independent 1B transcriptional repressor [Source:HGNC Symbol;Acc:HGNC:4238] | -1.8961 | 6.38E-09 |
| ENSG00000134294 | SLC38A2 | solute carrier family 38 member 2 [Source:HGNC Symbol;Acc:HGNC:13448] | 0.5564 | 6.42E-09 |
| ENSG00000072422 | RHOBTB1 | Rho related BTB domain containing 1 [Source:HGNC Symbol;Acc:HGNC:18738] | -0.7411 | 7.55E-09 |
| ENSG00000255036 | STRA6LP | STRA6 like, pseudogene [Source:HGNC Symbol;Acc:HGNC:53830] | 0.8612 | 8.97E-09 |
| ENSG00000187231 | SESTD1 | SEC14 and spectrin domain containing 1 [Source:HGNC Symbol;Acc:HGNC:18379] | 0.6090 | 9.43E-09 |
| ENSG00000165240 | ATP7A | ATPase copper transporting alpha [Source:HGNC Symbol;Acc:HGNC:869] | 0.6242 | 1.55E-08 |
| ENSG00000119915 | ELOVL3 | ELOVL fatty acid elongase 3 [Source:HGNC Symbol;Acc:HGNC:18047] | -1.3594 | 1.99E-08 |
| ENSG00000106025 | TSPAN12 | tetraspanin 12 [Source:HGNC Symbol;Acc:HGNC:21641] | 0.7912 | 2.01E-08 |
| ENSG00000005249 | PRKAR2B | protein kinase cAMP-dependent type II regulatory subunit beta [Source:HGNC Symbol;Acc:HGNC:9392] | 0.8571 | 2.24E-08 |
| ENSG00000187479 | C11orf96 | chromosome 11 open reading frame 96 [Source:HGNC Symbol;Acc:HGNC:38675] | 1.4453 | 2.26E-08 |
| ENSG00000188766 | SPRED3 | sprouty related EVH1 domain containing 3 [Source:HGNC Symbol;Acc:HGNC:31041] | 1.4788 | 2.29E-08 |
| ENSG00000160882 | CYP11B1 | cytochrome P450 family 11 subfamily B member 1 [Source:HGNC Symbol;Acc:HGNC:2591] | 1.9336 | 2.32E-08 |
| ENSG00000085741 | WNT11 | Wnt family member 11 [Source:HGNC Symbol;Acc:HGNC:12776] | -0.6632 | 2.50E-08 |
| ENSG00000147894 | C9orf72 | C9orf72-SMCR8 complex subunit [Source:HGNC Symbol;Acc:HGNC:28337] | 0.7591 | 2.52E-08 |
| ENSG00000102780 | DGKH | diacylglycerol kinase eta [Source:HGNC Symbol;Acc:HGNC:2854] | 0.8980 | 2.75E-08 |
| ENSG00000121361 | KCNJ8 | potassium inwardly rectifying channel subfamily J member 8 [Source:HGNC Symbol;Acc:HGNC:6269] | 1.1554 | 2.80E-08 |
| ENSG00000092969 | TGFB2 | transforming growth factor beta 2 [Source:HGNC Symbol;Acc:HGNC:11768] | 1.3502 | 3.56E-08 |
| ENSG00000163833 | FBXO40 | F-box protein 40 [Source:HGNC Symbol;Acc:HGNC:29816] | -1.2326 | 3.56E-08 |
| ENSG00000140859 | KIFC3 | kinesin family member C3 [Source:HGNC Symbol;Acc:HGNC:6326] | 0.8981 | 3.87E-08 |
| ENSG00000250986 | LINC02600 | long intergenic non-protein coding RNA 2600 [Source:HGNC Symbol;Acc:HGNC:53177] | -1.0753 | 3.87E-08 |
| ENSG00000112541 | PDE10A | phosphodiesterase 10A [Source:HGNC Symbol;Acc:HGNC:8772] | 1.2541 | 3.97E-08 |
| ENSG00000137714 | FDX1 | ferredoxin 1 [Source:HGNC Symbol;Acc:HGNC:3638] | 0.9324 | 4.43E-08 |
| ENSG00000146122 | DAAM2 | dishevelled associated activator of morphogenesis 2 [Source:HGNC Symbol;Acc:HGNC:18143] | 0.6691 | 4.43E-08 |
| ENSG00000147027 | TMEM47 | transmembrane protein 47 [Source:HGNC Symbol;Acc:HGNC:18515] | 0.8741 | 4.43E-08 |
| ENSG00000167580 | AQP2 | aquaporin 2 [Source:HGNC Symbol;Acc:HGNC:634] | 1.2313 | 5.52E-08 |
| ENSG00000078053 | AMPH | amphiphysin [Source:HGNC Symbol;Acc:HGNC:471] | 0.5156 | 6.01E-08 |
| ENSG00000185070 | FLRT2 | fibronectin leucine rich transmembrane protein 2 [Source:HGNC Symbol;Acc:HGNC:3761] | 0.8556 | 6.18E-08 |
| ENSG00000084710 | EFR3B | EFR3 homolog B [Source:HGNC Symbol;Acc:HGNC:29155] | 1.1272 | 6.31E-08 |
| ENSG00000176406 | RIMS2 | regulating synaptic membrane exocytosis 2 [Source:HGNC Symbol;Acc:HGNC:17283] | 0.5508 | 6.67E-08 |
| ENSG00000198846 | TOX | thymocyte selection associated high mobility group box [Source:HGNC Symbol;Acc:HGNC:18988] | 1.4668 | 6.67E-08 |
| ENSG00000131791 | PRKAB2 | protein kinase AMP-activated non-catalytic subunit beta 2 [Source:HGNC Symbol;Acc:HGNC:9379] | -0.5806 | 8.45E-08 |
| ENSG00000151067 | CACNA1C | calcium voltage-gated channel subunit alpha1 C [Source:HGNC Symbol;Acc:HGNC:1390] | 1.0072 | 9.41E-08 |
| ENSG00000130758 | MAP3K10 | mitogen-activated protein kinase kinase kinase 10 [Source:HGNC Symbol;Acc:HGNC:6849] | 0.6122 | 9.64E-08 |

| | | | | |
|---|---|---|---|---|
| ENSG00000145779 | TNFAIP8 | TNF alpha induced protein 8 [Source:HGNC Symbol;Acc:HGNC:17260] | 1.1355 | 1.04E-07 |
| ENSG00000151914 | DST | dystonin [Source:HGNC Symbol;Acc:HGNC:1090] | 0.6309 | 1.05E-07 |
| ENSG00000118432 | CNR1 | cannabinoid receptor 1 [Source:HGNC Symbol;Acc:HGNC:2159] | -0.7344 | 1.15E-07 |
| ENSG00000060709 | RIMBP2 | RIMS binding protein 2 [Source:HGNC Symbol;Acc:HGNC:30339] | 1.4438 | 1.26E-07 |
| ENSG00000127824 | TUBA4A | tubulin alpha 4a [Source:HGNC Symbol;Acc:HGNC:12407] | 1.2518 | 1.31E-07 |
| ENSG00000167971 | CASKIN1 | CASK interacting protein 1 [Source:HGNC Symbol;Acc:HGNC:20879] | 1.5269 | 1.43E-07 |
| ENSG00000095397 | WHRN | whirlin [Source:HGNC Symbol;Acc:HGNC:16361] | 0.7797 | 1.52E-07 |
| ENSG00000149212 | SESN3 | sestrin 3 [Source:HGNC Symbol;Acc:HGNC:23060] | 0.8770 | 1.52E-07 |
| ENSG00000170485 | NPAS2 | neuronal PAS domain protein 2 [Source:HGNC Symbol;Acc:HGNC:7895] | 1.0865 | 1.60E-07 |
| ENSG00000186642 | PDE2A | phosphodiesterase 2A [Source:HGNC Symbol;Acc:HGNC:8777] | -0.8923 | 1.61E-07 |
| ENSG00000146250 | PRSS35 | serine protease 35 [Source:HGNC Symbol;Acc:HGNC:21387] | 1.2006 | 1.64E-07 |
| ENSG00000160305 | DIP2A | disco interacting protein 2 homolog A [Source:HGNC Symbol;Acc:HGNC:17217] | 0.7244 | 2.30E-07 |
| ENSG00000109163 | GNRHR | gonadotropin releasing hormone receptor [Source:HGNC Symbol;Acc:HGNC:4421] | 0.8726 | 2.35E-07 |
| ENSG00000148841 | ITPRIP | inositol 1,4,5-trisphosphate receptor interacting protein [Source:HGNC Symbol;Acc:HGNC:29370] | 0.8934 | 3.00E-07 |
| ENSG00000145555 | MYO10 | myosin X [Source:HGNC Symbol;Acc:HGNC:7593] | -0.7262 | 3.28E-07 |
| ENSG00000225376 | TMEM246-AS1 | TMEM246 antisense RNA 1 [Source:HGNC Symbol;Acc:HGNC:51191] | 2.1190 | 3.33E-07 |
| ENSG00000148842 | CNNM2 | cyclin and CBS domain divalent metal cation transport mediator 2 [Source:HGNC Symbol;Acc:HGNC:103] | 0.6559 | 3.54E-07 |
| ENSG00000004660 | CAMKK1 | calcium/calmodulin dependent protein kinase kinase 1 [Source:HGNC Symbol;Acc:HGNC:1469] | -0.9719 | 4.24E-07 |
| ENSG00000143641 | GALNT2 | polypeptide N-acetylgalactosaminyltransferase 2 [Source:HGNC Symbol;Acc:HGNC:4124] | 0.6241 | 4.52E-07 |
| ENSG00000152661 | GJA1 | gap junction protein alpha 1 [Source:HGNC Symbol;Acc:HGNC:4274] | 0.9339 | 4.87E-07 |
| ENSG00000107731 | UNC5B | unc-5 netrin receptor B [Source:HGNC Symbol;Acc:HGNC:12568] | -1.1539 | 5.00E-07 |
| ENSG00000116741 | RGS2 | regulator of G protein signaling 2 [Source:HGNC Symbol;Acc:HGNC:9998] | 0.8088 | 5.03E-07 |
| ENSG00000213694 | S1PR3 | sphingosine-1-phosphate receptor 3 [Source:HGNC Symbol;Acc:HGNC:3167] | 0.6782 | 5.05E-07 |
| ENSG00000139988 | RDH12 | retinol dehydrogenase 12 [Source:HGNC Symbol;Acc:HGNC:19977] | 1.2145 | 5.47E-07 |
| ENSG00000006283 | CACNA1G | calcium voltage-gated channel subunit alpha1 G [Source:HGNC Symbol;Acc:HGNC:1394] | -0.8295 | 5.83E-07 |
| ENSG00000140563 | MCTP2 | multiple C2 and transmembrane domain containing 2 [Source:HGNC Symbol;Acc:HGNC:25636] | 0.9403 | 6.60E-07 |
| ENSG00000134508 | CABLES1 | Cdk5 and Abl enzyme substrate 1 [Source:HGNC Symbol;Acc:HGNC:25097] | 1.3080 | 8.18E-07 |
| ENSG00000069849 | ATP1B3 | ATPase Na+/K+ transporting subunit beta 3 [Source:HGNC Symbol;Acc:HGNC:806] | 0.6484 | 8.43E-07 |
| ENSG00000108244 | KRT23 | keratin 23 [Source:HGNC Symbol;Acc:HGNC:6438] | 2.0726 | 9.44E-07 |
| ENSG00000122824 | NUDT10 | nudix hydrolase 10 [Source:HGNC Symbol;Acc:HGNC:17621] | 0.6778 | 9.44E-07 |
| ENSG00000132938 | MTUS2 | microtubule associated scaffold protein 2 [Source:HGNC Symbol;Acc:HGNC:20595] | 1.6340 | 9.51E-07 |
| ENSG00000139973 | SYT16 | synaptotagmin 16 [Source:HGNC Symbol;Acc:HGNC:23142] | 1.8052 | 9.75E-07 |
| ENSG00000108984 | MAP2K6 | mitogen-activated protein kinase kinase 6 [Source:HGNC Symbol;Acc:HGNC:6846] | -1.0388 | 1.01E-06 |
| ENSG00000124374 | PAIP2B | poly(A) binding protein interacting protein 2B [Source:HGNC Symbol;Acc:HGNC:29200] | -0.6936 | 1.01E-06 |
| ENSG00000119408 | NEK6 | NIMA related kinase 6 [Source:HGNC Symbol;Acc:HGNC:7749] | 1.0967 | 1.09E-06 |
| ENSG00000171843 | MLLT3 | MLLT3 super elongation complex subunit [Source:HGNC Symbol;Acc:HGNC:7136] | 0.5555 | 1.11E-06 |
| ENSG00000057252 | SOAT1 | sterol O-acyltransferase 1 [Source:HGNC Symbol;Acc:HGNC:11177] | 0.5674 | 1.18E-06 |
| ENSG00000117586 | TNFSF4 | TNF superfamily member 4 [Source:HGNC Symbol;Acc:HGNC:11934] | 0.8525 | 1.18E-06 |
| ENSG00000146242 | TPBG | trophoblast glycoprotein [Source:HGNC Symbol;Acc:HGNC:12004] | 0.5775 | 1.18E-06 |
| ENSG00000185338 | SOCS1 | suppressor of cytokine signaling 1 [Source:HGNC Symbol;Acc:HGNC:19383] | 0.9984 | 1.18E-06 |
| ENSG00000115540 | MOB4 | MOB family member 4, phocein [Source:HGNC Symbol;Acc:HGNC:17261] | 0.8354 | 1.21E-06 |
| ENSG00000182796 | TMEM198B | transmembrane protein 198B (pseudogene) [Source:HGNC Symbol;Acc:HGNC:43629] | -0.5763 | 1.23E-06 |
| ENSG00000154639 | CXADR | CXADR Ig-like cell adhesion molecule [Source:HGNC Symbol;Acc:HGNC:2559] | 0.8533 | 1.24E-06 |
| ENSG00000153930 | ANKFN1 | ankyrin repeat and fibronectin type III domain containing 1 [Source:HGNC Symbol;Acc:HGNC:26766] | -1.0605 | 1.38E-06 |
| ENSG00000084674 | APOB | apolipoprotein B [Source:HGNC Symbol;Acc:HGNC:603] | 0.9172 | 1.53E-06 |
| ENSG00000119771 | KLHL29 | kelch like family member 29 [Source:HGNC Symbol;Acc:HGNC:29404] | 0.8090 | 1.55E-06 |
| ENSG00000095794 | CREM | cAMP responsive element modulator [Source:HGNC Symbol;Acc:HGNC:2352] | 0.6807 | 1.58E-06 |
| ENSG00000170500 | LONRF2 | LON peptidase N-terminal domain and ring finger 2 [Source:HGNC Symbol;Acc:HGNC:24788] | 0.7596 | 1.64E-06 |
| ENSG00000120306 | CYSTM1 | cysteine rich transmembrane module containing 1 [Source:HGNC Symbol;Acc:HGNC:30239] | 0.6326 | 1.70E-06 |
| ENSG00000131773 | KHDRBS3 | KH RNA binding domain containing, signal transduction associated 3 [Source:HGNC Symbol;Acc:HGNC:18117] | -0.6241 | 1.70E-06 |
| ENSG00000086300 | SNX10 | sorting nexin 10 [Source:HGNC Symbol;Acc:HGNC:14974] | 0.5448 | 1.71E-06 |
| ENSG00000180628 | PCGF5 | polycomb group ring finger 5 [Source:HGNC Symbol;Acc:HGNC:28264] | 0.5253 | 1.84E-06 |
| ENSG00000129195 | PIMREG | PICALM interacting mitotic regulator [Source:HGNC Symbol;Acc:HGNC:25483] | -0.5036 | 2.01E-06 |
| ENSG00000168675 | LDLRAD4 | low density lipoprotein receptor class A domain containing 4 [Source:HGNC Symbol;Acc:HGNC:1224] | 1.1681 | 2.01E-06 |

| | | | | |
|---|---|---|---|---|
| ENSG00000146733 | PSPH | phosphoserine phosphatase [Source:HGNC Symbol;Acc:HGNC:9577] | -0.5303 | 2.12E-06 |
| ENSG00000156531 | PHF6 | PHD finger protein 6 [Source:HGNC Symbol;Acc:HGNC:18145] | 0.7985 | 2.29E-06 |
| ENSG00000145147 | SLIT2 | slit guidance ligand 2 [Source:HGNC Symbol;Acc:HGNC:11086] | 0.9099 | 2.34E-06 |
| ENSG00000165698 | SPACA9 | sperm acrosome associated 9 [Source:HGNC Symbol;Acc:HGNC:1367] | -0.8128 | 2.44E-06 |
| ENSG00000064651 | SLC12A2 | solute carrier family 12 member 2 [Source:HGNC Symbol;Acc:HGNC:10911] | 0.7313 | 2.47E-06 |
| ENSG00000179142 | CYP11B2 | cytochrome P450 family 11 subfamily B member 2 [Source:HGNC Symbol;Acc:HGNC:2592] | 2.5510 | 2.47E-06 |
| ENSG00000145632 | PLK2 | polo like kinase 2 [Source:HGNC Symbol;Acc:HGNC:19699] | 1.8516 | 2.85E-06 |
| ENSG00000161681 | SHANK1 | SH3 and multiple ankyrin repeat domains 1 [Source:HGNC Symbol;Acc:HGNC:15474] | -0.5459 | 2.92E-06 |
| ENSG00000162552 | WNT4 | Wnt family member 4 [Source:HGNC Symbol;Acc:HGNC:12783] | -0.8323 | 3.20E-06 |
| ENSG00000151718 | WWC2 | WW and C2 domain containing 2 [Source:HGNC Symbol;Acc:HGNC:24148] | 0.7719 | 3.77E-06 |
| ENSG00000133612 | AGAP3 | ArfGAP with GTPase domain, ankyrin repeat and PH domain 3 [Source:HGNC Symbol;Acc:HGNC:16923] | 0.5578 | 3.81E-06 |
| ENSG00000106031 | HOXA13 | homeobox A13 [Source:HGNC Symbol;Acc:HGNC:5102] | 0.8093 | 3.83E-06 |
| ENSG00000286257 | NA | novel transcript | 1.0396 | 3.85E-06 |
| ENSG00000173334 | TRIB1 | tribbles pseudokinase 1 [Source:HGNC Symbol;Acc:HGNC:16891] | 1.9923 | 3.90E-06 |
| ENSG00000106546 | AHR | aryl hydrocarbon receptor [Source:HGNC Symbol;Acc:HGNC:348] | 0.9966 | 4.21E-06 |
| ENSG00000230918 | DPP4-DT | DPP4 divergent transcript [Source:HGNC Symbol;Acc:HGNC:40191] | 1.6442 | 4.34E-06 |
| ENSG00000188386 | PPP3R2 | protein phosphatase 3 regulatory subunit B, beta [Source:HGNC Symbol;Acc:HGNC:9318] | 2.0360 | 4.59E-06 |
| ENSG00000175264 | CHST1 | carbohydrate sulfotransferase 1 [Source:HGNC Symbol;Acc:HGNC:1969] | 0.5087 | 4.78E-06 |
| ENSG00000196517 | SLC6A9 | solute carrier family 6 member 9 [Source:HGNC Symbol;Acc:HGNC:11056] | 0.6372 | 4.78E-06 |
| ENSG00000267432 | DNAH17-AS1 | DNAH17 antisense RNA 1 [Source:HGNC Symbol;Acc:HGNC:48594] | 1.4255 | 4.86E-06 |
| ENSG00000226950 | DANCR | differentiation antagonizing non-protein coding RNA [Source:HGNC Symbol;Acc:HGNC:28964] | 0.5236 | 5.30E-06 |
| ENSG00000068383 | INPP5A | inositol polyphosphate-5-phosphatase A [Source:HGNC Symbol;Acc:HGNC:6076] | 0.7233 | 5.62E-06 |
| ENSG00000261340 | LINC01616 | long intergenic non-protein coding RNA 1616 [Source:HGNC Symbol;Acc:HGNC:51900] | 1.7842 | 5.68E-06 |
| ENSG00000166917 | MIR202HG | MIR202 host gene [Source:HGNC Symbol;Acc:HGNC:49402] | -1.5857 | 6.35E-06 |
| ENSG00000055118 | KCNH2 | potassium voltage-gated channel subfamily H member 2 [Source:HGNC Symbol;Acc:HGNC:6251] | -0.7312 | 6.49E-06 |
| ENSG00000167766 | ZNF83 | zinc finger protein 83 [Source:HGNC Symbol;Acc:HGNC:13158] | -0.5491 | 6.58E-06 |
| ENSG00000167693 | NXN | nucleoredoxin [Source:HGNC Symbol;Acc:HGNC:18008] | 1.1343 | 6.65E-06 |
| ENSG00000181773 | GPR3 | G protein-coupled receptor 3 [Source:HGNC Symbol;Acc:HGNC:4484] | 2.0246 | 7.02E-06 |
| ENSG00000280061 | NA | TEC | 0.7844 | 7.55E-06 |
| ENSG00000170145 | SIK2 | salt inducible kinase 2 [Source:HGNC Symbol;Acc:HGNC:21680] | 0.6171 | 8.26E-06 |
| ENSG00000179314 | WSCD1 | WSC domain containing 1 [Source:HGNC Symbol;Acc:HGNC:29060] | 1.5135 | 8.71E-06 |
| ENSG00000197580 | BCO2 | beta-carotene oxygenase 2 [Source:HGNC Symbol;Acc:HGNC:18503] | 0.8081 | 8.85E-06 |
| ENSG00000145246 | ATP10D | ATPase phospholipid transporting 10D (putative) [Source:HGNC Symbol;Acc:HGNC:13549] | 0.8653 | 9.28E-06 |
| ENSG00000118257 | NRP2 | neuropilin 2 [Source:HGNC Symbol;Acc:HGNC:8005] | 0.8830 | 1.04E-05 |
| ENSG00000150593 | PDCD4 | programmed cell death 4 [Source:HGNC Symbol;Acc:HGNC:8763] | -0.5606 | 1.19E-05 |
| ENSG00000204682 | MIR1915HG | MIR1915 host gene [Source:HGNC Symbol;Acc:HGNC:31448] | 0.7422 | 1.23E-05 |
| ENSG00000272398 | CD24 | CD24 molecule [Source:HGNC Symbol;Acc:HGNC:1645] | 1.5894 | 1.25E-05 |
| ENSG00000019144 | PHLDB1 | pleckstrin homology like domain family B member 1 [Source:HGNC Symbol;Acc:HGNC:23697] | -0.5037 | 1.27E-05 |
| ENSG00000006210 | CX3CL1 | C-X3-C motif chemokine ligand 1 [Source:HGNC Symbol;Acc:HGNC:10647] | 0.5747 | 1.31E-05 |
| ENSG00000118513 | MYB | MYB proto-oncogene, transcription factor [Source:HGNC Symbol;Acc:HGNC:7545] | 1.1113 | 1.33E-05 |
| ENSG00000115738 | ID2 | inhibitor of DNA binding 2 [Source:HGNC Symbol;Acc:HGNC:5361] | -0.5913 | 1.37E-05 |
| ENSG00000183508 | TENT5C | terminal nucleotidyltransferase 5C [Source:HGNC Symbol;Acc:HGNC:24712] | 1.2573 | 1.39E-05 |
| ENSG00000139793 | MBNL2 | muscleblind like splicing regulator 2 [Source:HGNC Symbol;Acc:HGNC:16746] | 0.5305 | 1.52E-05 |
| ENSG00000136928 | GABBR2 | gamma-aminobutyric acid type B receptor subunit 2 [Source:HGNC Symbol;Acc:HGNC:4507] | -0.5637 | 1.53E-05 |
| ENSG00000167434 | CA4 | carbonic anhydrase 4 [Source:HGNC Symbol;Acc:HGNC:1375] | -1.0954 | 1.60E-05 |
| ENSG00000171408 | PDE7B | phosphodiesterase 7B [Source:HGNC Symbol;Acc:HGNC:8792] | 0.6076 | 1.62E-05 |
| ENSG00000236008 | LINC01814 | long intergenic non-protein coding RNA 1814 [Source:HGNC Symbol;Acc:HGNC:52618] | -0.7002 | 1.62E-05 |
| ENSG00000257365 | FNTB | farnesyltransferase, CAAX box, beta [Source:HGNC Symbol;Acc:HGNC:3785] | 0.5091 | 1.64E-05 |
| ENSG00000139998 | RAB15 | RAB15, member RAS oncogene family [Source:HGNC Symbol;Acc:HGNC:20150] | 0.6391 | 1.69E-05 |
| ENSG00000179630 | LACC1 | laccase domain containing 1 [Source:HGNC Symbol;Acc:HGNC:26789] | -1.6106 | 1.79E-05 |
| ENSG00000261325 | LINC02192 | long intergenic non-protein coding RNA 2192 [Source:HGNC Symbol;Acc:HGNC:53054] | 1.1041 | 1.84E-05 |
| ENSG00000143061 | IGSF3 | immunoglobulin superfamily member 3 [Source:HGNC Symbol;Acc:HGNC:5950] | -0.6089 | 1.91E-05 |
| ENSG00000151729 | SLC25A4 | solute carrier family 25 member 4 [Source:HGNC Symbol;Acc:HGNC:10990] | 0.6297 | 1.93E-05 |
| ENSG00000106852 | LHX6 | LIM homeobox 6 [Source:HGNC Symbol;Acc:HGNC:21735] | -1.0947 | 2.03E-05 |

| | | | | |
|---|---|---|---|---|
| ENSG00000136044 | APPL2 | adaptor protein, phosphotyrosine interacting with PH domain and leucine zipper 2 [Source:HGNC Symbol;Acc:HGNC:182 | 0.5209 | 2.35E-05 |
| ENSG00000069020 | MAST4 | microtubule associated serine/threonine kinase family member 4 [Source:HGNC Symbol;Acc:HGNC:19037] | 0.6540 | 2.43E-05 |
| ENSG00000134780 | DAGLA | diacylglycerol lipase alpha [Source:HGNC Symbol;Acc:HGNC:1165] | -0.7741 | 2.47E-05 |
| ENSG00000116717 | GADD45A | growth arrest and DNA damage inducible alpha [Source:HGNC Symbol;Acc:HGNC:4095] | 0.8349 | 2.47E-05 |
| ENSG00000065809 | FAM107B | family with sequence similarity 107 member B [Source:HGNC Symbol;Acc:HGNC:23726] | 0.6864 | 2.55E-05 |
| ENSG00000118407 | FILIP1 | filamin A interacting protein 1 [Source:HGNC Symbol;Acc:HGNC:21015] | -0.8426 | 2.55E-05 |
| ENSG00000162545 | CAMK2N1 | calcium/calmodulin dependent protein kinase II inhibitor 1 [Source:HGNC Symbol;Acc:HGNC:24190] | 1.0486 | 2.55E-05 |
| ENSG00000118503 | TNFAIP3 | TNF alpha induced protein 3 [Source:HGNC Symbol;Acc:HGNC:11896] | 0.7322 | 2.59E-05 |
| ENSG00000101298 | SNPH | syntaphilin [Source:HGNC Symbol;Acc:HGNC:15931] | 0.7306 | 2.60E-05 |
| ENSG00000151229 | SLC2A13 | solute carrier family 2 member 13 [Source:HGNC Symbol;Acc:HGNC:15956] | 0.9052 | 2.65E-05 |
| ENSG00000169302 | STK32A | serine/threonine kinase 32A [Source:HGNC Symbol;Acc:HGNC:28317] | -0.8241 | 2.65E-05 |
| ENSG00000259479 | SORD2P | sorbitol dehydrogenase 2, pseudogene [Source:HGNC Symbol;Acc:HGNC:49919] | 0.5290 | 2.85E-05 |
| ENSG00000184160 | ADRA2C | adrenoceptor alpha 2C [Source:HGNC Symbol;Acc:HGNC:283] | -1.3335 | 3.08E-05 |
| ENSG00000153162 | BMP6 | bone morphogenetic protein 6 [Source:HGNC Symbol;Acc:HGNC:1073] | 1.1955 | 3.40E-05 |
| ENSG00000072210 | ALDH3A2 | aldehyde dehydrogenase 3 family member A2 [Source:HGNC Symbol;Acc:HGNC:403] | 0.6032 | 3.52E-05 |
| ENSG00000152932 | RAB3C | RAB3C, member RAS oncogene family [Source:HGNC Symbol;Acc:HGNC:30269] | 0.8270 | 4.38E-05 |
| ENSG00000105642 | KCNN1 | potassium calcium-activated channel subfamily N member 1 [Source:HGNC Symbol;Acc:HGNC:6290] | -0.9237 | 4.45E-05 |
| ENSG00000140263 | SORD | sorbitol dehydrogenase [Source:HGNC Symbol;Acc:HGNC:11184] | 0.5341 | 4.67E-05 |
| ENSG00000104976 | SNAPC2 | small nuclear RNA activating complex polypeptide 2 [Source:HGNC Symbol;Acc:HGNC:11135] | -0.5421 | 5.10E-05 |
| ENSG00000137959 | IFI44L | interferon induced protein 44 like [Source:HGNC Symbol;Acc:HGNC:17817] | -0.9026 | 5.13E-05 |
| ENSG00000153234 | NR4A2 | nuclear receptor subfamily 4 group A member 2 [Source:HGNC Symbol;Acc:HGNC:7981] | 2.5281 | 5.13E-05 |
| ENSG00000245864 | MEF2C-AS2 | MEF2C antisense RNA 2 [Source:HGNC Symbol;Acc:HGNC:53115] | 0.5617 | 5.58E-05 |
| ENSG00000273419 | NA | novel transcript, antisense to ZNF862 | 1.0284 | 5.65E-05 |
| ENSG00000120068 | HOXB8 | homeobox B8 [Source:HGNC Symbol;Acc:HGNC:5119] | 0.8175 | 5.67E-05 |
| ENSG00000182255 | KCNA4 | potassium voltage-gated channel subfamily A member 4 [Source:HGNC Symbol;Acc:HGNC:6222] | 1.0544 | 5.67E-05 |
| ENSG00000177910 | SPATA31C2 | SPATA31 subfamily C member 2 [Source:HGNC Symbol;Acc:HGNC:24508] | -0.8453 | 5.68E-05 |
| ENSG00000125148 | MT2A | metallothionein 2A [Source:HGNC Symbol;Acc:HGNC:7406] | 0.7217 | 5.91E-05 |
| ENSG00000079308 | TNS1 | tensin 1 [Source:HGNC Symbol;Acc:HGNC:11973] | -0.6120 | 5.91E-05 |
| ENSG00000164690 | SHH | sonic hedgehog signaling molecule [Source:HGNC Symbol;Acc:HGNC:10848] | 0.8555 | 5.91E-05 |
| ENSG00000185269 | NOTUM | notum, palmitoleoyl-protein carboxylesterase [Source:HGNC Symbol;Acc:HGNC:27106] | -0.9700 | 6.89E-05 |
| ENSG00000049883 | PTCD2 | pentatricopeptide repeat domain 2 [Source:HGNC Symbol;Acc:HGNC:25734] | -0.5315 | 7.14E-05 |
| ENSG00000266401 | NA | novel transcript, antisense to DLGAP1 | 1.3519 | 7.16E-05 |
| ENSG00000143590 | EFNA3 | ephrin A3 [Source:HGNC Symbol;Acc:HGNC:3223] | -0.7592 | 7.35E-05 |
| ENSG00000173548 | SNX33 | sorting nexin 33 [Source:HGNC Symbol;Acc:HGNC:28468] | 0.9394 | 7.41E-05 |
| ENSG00000154640 | BTG3 | BTG anti-proliferation factor 3 [Source:HGNC Symbol;Acc:HGNC:1132] | 0.7348 | 7.43E-05 |
| ENSG00000166292 | TMEM100 | transmembrane protein 100 [Source:HGNC Symbol;Acc:HGNC:25607] | 0.6280 | 8.00E-05 |
| ENSG00000185630 | PBX1 | PBX homeobox 1 [Source:HGNC Symbol;Acc:HGNC:8632] | 0.5274 | 8.00E-05 |
| ENSG00000102996 | MMP15 | matrix metallopeptidase 15 [Source:HGNC Symbol;Acc:HGNC:7161] | -0.6998 | 8.22E-05 |
| ENSG00000165194 | PCDH19 | protocadherin 19 [Source:HGNC Symbol;Acc:HGNC:14270] | 1.7038 | 8.59E-05 |
| ENSG00000133800 | LYVE1 | lymphatic vessel endothelial hyaluronan receptor 1 [Source:HGNC Symbol;Acc:HGNC:14687] | 1.0671 | 8.76E-05 |
| ENSG00000204291 | COL15A1 | collagen type XV alpha 1 chain [Source:HGNC Symbol;Acc:HGNC:2192] | 0.7591 | 9.19E-05 |
| ENSG00000165617 | DACT1 | dishevelled binding antagonist of beta catenin 1 [Source:HGNC Symbol;Acc:HGNC:17748] | 0.8900 | 9.37E-05 |
| ENSG00000119508 | NR4A3 | nuclear receptor subfamily 4 group A member 3 [Source:HGNC Symbol;Acc:HGNC:7982] | 1.2982 | 0.00011 |
| ENSG00000119599 | DCAF4 | DDB1 and CUL4 associated factor 4 [Source:HGNC Symbol;Acc:HGNC:20229] | -0.8024 | 0.00011 |
| ENSG00000115963 | RND3 | Rho family GTPase 3 [Source:HGNC Symbol;Acc:HGNC:671] | 1.6411 | 0.00011 |
| ENSG00000070961 | ATP2B1 | ATPase plasma membrane Ca2+ transporting 1 [Source:HGNC Symbol;Acc:HGNC:814] | 0.6803 | 0.00012 |
| ENSG00000204060 | FOXO6 | forkhead box O6 [Source:HGNC Symbol;Acc:HGNC:24814] | -0.9952 | 0.00012 |
| ENSG00000078804 | TP53INP2 | tumor protein p53 inducible nuclear protein 2 [Source:HGNC Symbol;Acc:HGNC:16104] | 0.6233 | 0.00013 |
| ENSG00000169783 | LINGO1 | leucine rich repeat and Ig domain containing 1 [Source:HGNC Symbol;Acc:HGNC:21205] | -0.9359 | 0.00013 |
| ENSG00000105849 | TWISTNB | TWIST neighbor [Source:HGNC Symbol;Acc:HGNC:18027] | 0.7728 | 0.00014 |
| ENSG00000078814 | MYH7B | myosin heavy chain 7B [Source:HGNC Symbol;Acc:HGNC:15906] | -0.8195 | 0.00014 |
| ENSG00000164089 | ETNPPL | ethanolamine-phosphate phospho-lyase [Source:HGNC Symbol;Acc:HGNC:14404] | 0.9843 | 0.00014 |
| ENSG00000158186 | MRAS | muscle RAS oncogene homolog [Source:HGNC Symbol;Acc:HGNC:7227] | 0.5315 | 0.00014 |
| ENSG00000281832 | LINC00602 | long intergenic non-protein coding RNA 602 [Source:HGNC Symbol;Acc:HGNC:43917] | 1.6457 | 0.00015 |

| | | | | |
|---|---|---|---|---|
| ENSG00000105875 | WDR91 | WD repeat domain 91 [Source:HGNC Symbol;Acc:HGNC:24997] | 0.8345 | 0.00015 |
| ENSG00000164442 | CITED2 | Cbp/p300 interacting transactivator with Glu/Asp rich carboxy-terminal domain 2 [Source:HGNC Symbol;Acc:HGNC:198' | 0.7349 | 0.00015 |
| ENSG00000172551 | MUCL1 | mucin like 1 [Source:HGNC Symbol;Acc:HGNC:30588] | -0.6318 | 0.00015 |
| ENSG00000244968 | LIFR-AS1 | LIFR antisense RNA 1 [Source:HGNC Symbol;Acc:HGNC:43600] | -0.7206 | 0.00016 |
| ENSG00000111432 | FZD10 | frizzled class receptor 10 [Source:HGNC Symbol;Acc:HGNC:4039] | 1.7799 | 0.00016 |
| ENSG00000162769 | FLVCR1 | FLVCR heme transporter 1 [Source:HGNC Symbol;Acc:HGNC:24682] | 0.5254 | 0.00016 |
| ENSG00000184675 | AMER1 | APC membrane recruitment protein 1 [Source:HGNC Symbol;Acc:HGNC:26837] | -0.5679 | 0.00016 |
| ENSG00000204516 | MICB | MHC class I polypeptide-related sequence B [Source:HGNC Symbol;Acc:HGNC:7091] | 0.8408 | 0.00017 |
| ENSG00000197852 | INKA2 | inka box actin regulator 2 [Source:HGNC Symbol;Acc:HGNC:28045] | 0.5758 | 0.00017 |
| ENSG00000039560 | RAI14 | retinoic acid induced 14 [Source:HGNC Symbol;Acc:HGNC:14873] | 0.5415 | 0.00018 |
| ENSG00000180263 | FGD6 | FYVE, RhoGEF and PH domain containing 6 [Source:HGNC Symbol;Acc:HGNC:21740] | 0.7747 | 0.00018 |
| ENSG00000064666 | CNN2 | calponin 2 [Source:HGNC Symbol;Acc:HGNC:2156] | 0.8109 | 0.00019 |
| ENSG00000173269 | MMRN2 | multimerin 2 [Source:HGNC Symbol;Acc:HGNC:19888] | -1.2220 | 0.00019 |
| ENSG00000253276 | CCDC71L | coiled-coil domain containing 71 like [Source:HGNC Symbol;Acc:HGNC:26685] | -0.5394 | 0.00020 |
| ENSG00000183826 | BTBD9 | BTB domain containing 9 [Source:HGNC Symbol;Acc:HGNC:21228] | 0.5612 | 0.00021 |
| ENSG00000186594 | MIR22HG | MIR22 host gene [Source:HGNC Symbol;Acc:HGNC:28219] | 0.7762 | 0.00021 |
| ENSG00000198598 | MMP17 | matrix metallopeptidase 17 [Source:HGNC Symbol;Acc:HGNC:7163] | 1.6202 | 0.00021 |
| ENSG00000144290 | SLC4A10 | solute carrier family 4 member 10 [Source:HGNC Symbol;Acc:HGNC:13811] | 0.9755 | 0.00023 |
| ENSG00000142677 | IL22RA1 | interleukin 22 receptor subunit alpha 1 [Source:HGNC Symbol;Acc:HGNC:13700] | -0.9563 | 0.00023 |
| ENSG00000120875 | DUSP4 | dual specificity phosphatase 4 [Source:HGNC Symbol;Acc:HGNC:3070] | 0.9181 | 0.00024 |
| ENSG00000130066 | SAT1 | spermidine/spermine N1-acetyltransferase 1 [Source:HGNC Symbol;Acc:HGNC:10540] | 0.8358 | 0.00026 |
| ENSG00000151224 | MAT1A | methionine adenosyltransferase 1A [Source:HGNC Symbol;Acc:HGNC:6903] | -1.9508 | 0.00027 |
| ENSG00000106689 | LHX2 | LIM homeobox 2 [Source:HGNC Symbol;Acc:HGNC:6594] | 1.0401 | 0.00028 |
| ENSG00000188582 | PAQR9 | progestin and adipoQ receptor family member 9 [Source:HGNC Symbol;Acc:HGNC:30131] | 1.2803 | 0.00030 |
| ENSG00000180447 | GAS1 | growth arrest specific 1 [Source:HGNC Symbol;Acc:HGNC:4165] | 1.1361 | 0.00032 |
| ENSG00000180758 | GPR157 | G protein-coupled receptor 157 [Source:HGNC Symbol;Acc:HGNC:23687] | -1.0093 | 0.00032 |
| ENSG00000141448 | GATA6 | GATA binding protein 6 [Source:HGNC Symbol;Acc:HGNC:4174] | 0.7890 | 0.00032 |
| ENSG00000161082 | CELF5 | CUGBP Elav-like family member 5 [Source:HGNC Symbol;Acc:HGNC:14058] | -0.7179 | 0.00032 |
| ENSG00000165757 | JCAD | junctional cadherin 5 associated [Source:HGNC Symbol;Acc:HGNC:29283] | -1.1814 | 0.00033 |
| ENSG00000135541 | AHI1 | Abelson helper integration site 1 [Source:HGNC Symbol;Acc:HGNC:21575] | 0.5278 | 0.00033 |
| ENSG00000032444 | PNPLA6 | patatin like phospholipase domain containing 6 [Source:HGNC Symbol;Acc:HGNC:16268] | 0.6334 | 0.00034 |
| ENSG00000154217 | PITPNC1 | phosphatidylinositol transfer protein cytoplasmic 1 [Source:HGNC Symbol;Acc:HGNC:21045] | 0.7703 | 0.00034 |
| ENSG00000026103 | FAS | Fas cell surface death receptor [Source:HGNC Symbol;Acc:HGNC:11920] | 0.6599 | 0.00035 |
| ENSG00000267221 | C17orf113 | chromosome 17 open reading frame 113 [Source:HGNC Symbol;Acc:HGNC:53437] | -1.4026 | 0.00035 |
| ENSG00000105825 | TFPI2 | tissue factor pathway inhibitor 2 [Source:HGNC Symbol;Acc:HGNC:11761] | 1.2353 | 0.00036 |
| ENSG00000067992 | PDK3 | pyruvate dehydrogenase kinase 3 [Source:HGNC Symbol;Acc:HGNC:8811] | 0.6249 | 0.00036 |
| ENSG00000137713 | PPP2R1B | protein phosphatase 2 scaffold subunit Abeta [Source:HGNC Symbol;Acc:HGNC:9303] | 0.6754 | 0.00036 |
| ENSG00000158125 | XDH | xanthine dehydrogenase [Source:HGNC Symbol;Acc:HGNC:12805] | -2.4457 | 0.00036 |
| ENSG00000111371 | SLC38A1 | solute carrier family 38 member 1 [Source:HGNC Symbol;Acc:HGNC:13447] | 0.5164 | 0.00037 |
| ENSG00000112893 | MAN2A1 | mannosidase alpha class 2A member 1 [Source:HGNC Symbol;Acc:HGNC:6824] | 0.5701 | 0.00037 |
| ENSG00000266094 | RASSF5 | Ras association domain family member 5 [Source:HGNC Symbol;Acc:HGNC:17609] | -0.8810 | 0.00037 |
| ENSG00000185532 | PRKG1 | protein kinase cGMP-dependent 1 [Source:HGNC Symbol;Acc:HGNC:9414] | 0.6376 | 0.00038 |
| ENSG00000124588 | NQO2 | N-ribosyldihydronicotinamide:quinone reductase 2 [Source:HGNC Symbol;Acc:HGNC:7856] | 0.6047 | 0.00039 |
| ENSG00000067113 | PLPP1 | phospholipid phosphatase 1 [Source:HGNC Symbol;Acc:HGNC:9228] | 0.5868 | 0.00040 |
| ENSG00000064199 | SPA17 | sperm autoantigenic protein 17 [Source:HGNC Symbol;Acc:HGNC:11210] | -0.8727 | 0.00041 |
| ENSG00000133710 | SPINK5 | serine peptidase inhibitor Kazal type 5 [Source:HGNC Symbol;Acc:HGNC:15464] | 0.7395 | 0.00041 |
| ENSG00000164741 | DLC1 | DLC1 Rho GTPase activating protein [Source:HGNC Symbol;Acc:HGNC:2897] | 0.5053 | 0.00043 |
| ENSG00000186806 | VSIG10L | V-set and immunoglobulin domain containing 10 like [Source:HGNC Symbol;Acc:HGNC:27111] | -0.5587 | 0.00044 |
| ENSG00000197121 | PGAP1 | post-GPI attachment to proteins inositol deacylase 1 [Source:HGNC Symbol;Acc:HGNC:25712] | 0.7984 | 0.00044 |
| ENSG00000111452 | ADGRD1 | adhesion G protein-coupled receptor D1 [Source:HGNC Symbol;Acc:HGNC:19893] | 1.4982 | 0.00044 |
| ENSG00000240086 | NA | novel transcript, antisense to EPHB1 | -0.8370 | 0.00044 |
| ENSG00000074621 | SLC24A1 | solute carrier family 24 member 1 [Source:HGNC Symbol;Acc:HGNC:10975] | -0.5737 | 0.00045 |
| ENSG00000157110 | RBPMS | RNA binding protein, mRNA processing factor [Source:HGNC Symbol;Acc:HGNC:19097] | 0.6365 | 0.00045 |
| ENSG00000165905 | LARGE2 | LARGE xylosyl- and glucuronyltransferase 2 [Source:HGNC Symbol;Acc:HGNC:16522] | -0.6738 | 0.00045 |

| | | | | |
|---|---|---|---|---|
| ENSG00000102962 | CCL22 | C-C motif chemokine ligand 22 [Source:HGNC Symbol;Acc:HGNC:10621] | 0.6138 | 0.00047 |
| ENSG00000287409 | NA | novel transcript | 1.2229 | 0.00048 |
| ENSG00000140284 | SLC27A2 | solute carrier family 27 member 2 [Source:HGNC Symbol;Acc:HGNC:10996] | 2.2888 | 0.00048 |
| ENSG00000163285 | GABRG1 | gamma-aminobutyric acid type A receptor subunit gamma1 [Source:HGNC Symbol;Acc:HGNC:4086] | -1.6023 | 0.00050 |
| ENSG00000233581 | NA | novel transcript | 1.3375 | 0.00051 |
| ENSG00000106789 | CORO2A | coronin 2A [Source:HGNC Symbol;Acc:HGNC:2255] | -0.5789 | 0.00051 |
| ENSG00000182118 | FAM89A | family with sequence similarity 89 member A [Source:HGNC Symbol;Acc:HGNC:25057] | 0.7219 | 0.00051 |
| ENSG00000127920 | GNG11 | G protein subunit gamma 11 [Source:HGNC Symbol;Acc:HGNC:4403] | 0.7467 | 0.00053 |
| ENSG00000144893 | MED12L | mediator complex subunit 12L [Source:HGNC Symbol;Acc:HGNC:16050] | 0.5428 | 0.00054 |
| ENSG00000272556 | GTF2IP13 | general transcription factor IIi pseudogene 13 [Source:HGNC Symbol;Acc:HGNC:51725] | -0.5136 | 0.00054 |
| ENSG00000144802 | NFKBIZ | NFKB inhibitor zeta [Source:HGNC Symbol;Acc:HGNC:29805] | 0.8577 | 0.00054 |
| ENSG00000173221 | GLRX | glutaredoxin [Source:HGNC Symbol;Acc:HGNC:4330] | 0.8565 | 0.00054 |
| ENSG00000050030 | NEXMIF | neurite extension and migration factor [Source:HGNC Symbol;Acc:HGNC:29433] | -0.9780 | 0.00059 |
| ENSG00000135766 | EGLN1 | egl-9 family hypoxia inducible factor 1 [Source:HGNC Symbol;Acc:HGNC:1232] | 0.5419 | 0.00059 |
| ENSG00000121966 | CXCR4 | C-X-C motif chemokine receptor 4 [Source:HGNC Symbol;Acc:HGNC:2561] | 0.5473 | 0.00060 |
| ENSG00000155090 | KLF10 | Kruppel like factor 10 [Source:HGNC Symbol;Acc:HGNC:11810] | -0.8969 | 0.00060 |
| ENSG00000196562 | SULF2 | sulfatase 2 [Source:HGNC Symbol;Acc:HGNC:20392] | -1.1703 | 0.00063 |
| ENSG00000259070 | LINC00639 | long intergenic non-protein coding RNA 639 [Source:HGNC Symbol;Acc:HGNC:27502] | 0.9830 | 0.00064 |
| ENSG00000100302 | RASD2 | RASD family member 2 [Source:HGNC Symbol;Acc:HGNC:18229] | 0.6040 | 0.00064 |
| ENSG00000170011 | MYRIP | myosin VIIA and Rab interacting protein [Source:HGNC Symbol;Acc:HGNC:19156] | 0.8372 | 0.00064 |
| ENSG00000265972 | TXNIP | thioredoxin interacting protein [Source:HGNC Symbol;Acc:HGNC:16952] | -1.3787 | 0.00064 |
| ENSG00000198369 | SPRED2 | sprouty related EVH1 domain containing 2 [Source:HGNC Symbol;Acc:HGNC:17722] | 0.8552 | 0.00069 |
| ENSG00000272688 | NA | novel transcript | 1.4524 | 0.00071 |
| ENSG00000100246 | DNAL4 | dynein axonemal light chain 4 [Source:HGNC Symbol;Acc:HGNC:2955] | -0.5187 | 0.00072 |
| ENSG00000136295 | TTYH3 | tweety family member 3 [Source:HGNC Symbol;Acc:HGNC:22222] | 0.5095 | 0.00074 |
| ENSG00000140807 | NKD1 | NKD inhibitor of WNT signaling pathway 1 [Source:HGNC Symbol;Acc:HGNC:17045] | -0.5584 | 0.00074 |
| ENSG00000167094 | TTC16 | tetratricopeptide repeat domain 16 [Source:HGNC Symbol;Acc:HGNC:26536] | -1.4999 | 0.00078 |
| ENSG00000175928 | LRRN1 | leucine rich repeat neuronal 1 [Source:HGNC Symbol;Acc:HGNC:20980] | 1.1686 | 0.00079 |
| ENSG00000163625 | WDFY3 | WD repeat and FYVE domain containing 3 [Source:HGNC Symbol;Acc:HGNC:20751] | 0.5516 | 0.00080 |
| ENSG00000178764 | ZHX2 | zinc fingers and homeoboxes 2 [Source:HGNC Symbol;Acc:HGNC:18513] | 0.5615 | 0.00084 |
| ENSG00000110880 | CORO1C | coronin 1C [Source:HGNC Symbol;Acc:HGNC:2254] | 0.5104 | 0.00085 |
| ENSG00000189046 | ALKBH2 | alkB homolog 2, alpha-ketoglutarate dependent dioxygenase [Source:HGNC Symbol;Acc:HGNC:32487] | -0.6104 | 0.00085 |
| ENSG00000215817 | ZC3H11B | zinc finger CCCH-type containing 11B [Source:HGNC Symbol;Acc:HGNC:25659] | 0.8345 | 0.00085 |
| ENSG00000236432 | MFF-DT | MFF divergent transcript [Source:HGNC Symbol;Acc:HGNC:41067] | -0.7728 | 0.00086 |
| ENSG00000139318 | DUSP6 | dual specificity phosphatase 6 [Source:HGNC Symbol;Acc:HGNC:3072] | 1.0627 | 0.00092 |
| ENSG00000142494 | SLC47A1 | solute carrier family 47 member 1 [Source:HGNC Symbol;Acc:HGNC:25588] | 0.5688 | 0.00092 |
| ENSG00000076555 | ACACB | acetyl-CoA carboxylase beta [Source:HGNC Symbol;Acc:HGNC:85] | 0.6069 | 0.00093 |
| ENSG00000230797 | YY2 | YY2 transcription factor [Source:HGNC Symbol;Acc:HGNC:31684] | -0.6879 | 0.00095 |
| ENSG00000108960 | MMD | monocyte to macrophage differentiation associated [Source:HGNC Symbol;Acc:HGNC:7153] | 0.6819 | 0.00097 |
| ENSG00000160867 | FGFR4 | fibroblast growth factor receptor 4 [Source:HGNC Symbol;Acc:HGNC:3691] | -0.6154 | 0.00097 |
| ENSG00000136859 | ANGPTL2 | angiopoietin like 2 [Source:HGNC Symbol;Acc:HGNC:490] | -0.7530 | 0.00106 |
| ENSG00000070540 | WIPI1 | WD repeat domain, phosphoinositide interacting 1 [Source:HGNC Symbol;Acc:HGNC:25471] | 0.6679 | 0.00106 |
| ENSG00000185742 | C11orf87 | chromosome 11 open reading frame 87 [Source:HGNC Symbol;Acc:HGNC:33788] | 1.4516 | 0.00107 |
| ENSG00000164237 | CMBL | carboxymethylenebutenolidase homolog [Source:HGNC Symbol;Acc:HGNC:25090] | -0.5180 | 0.00107 |
| ENSG00000173281 | PPP1R3B | protein phosphatase 1 regulatory subunit 3B [Source:HGNC Symbol;Acc:HGNC:14942] | -0.9179 | 0.00110 |
| ENSG00000278530 | CHMP1B2P | charged multivesicular body protein 1B2, pseudogene [Source:HGNC Symbol;Acc:HGNC:49380] | 0.7384 | 0.00112 |
| ENSG00000119900 | OGFRL1 | opioid growth factor receptor like 1 [Source:HGNC Symbol;Acc:HGNC:21378] | 0.6643 | 0.00114 |
| ENSG00000285830 | NA | novel transcript, antisense to TRAPPC6B | -1.5456 | 0.00115 |
| ENSG00000122884 | P4HA1 | prolyl 4-hydroxylase subunit alpha 1 [Source:HGNC Symbol;Acc:HGNC:8546] | 0.6447 | 0.00117 |
| ENSG00000136997 | MYC | MYC proto-oncogene, bHLH transcription factor [Source:HGNC Symbol;Acc:HGNC:7553] | -0.5674 | 0.00117 |
| ENSG00000125744 | RTN2 | reticulon 2 [Source:HGNC Symbol;Acc:HGNC:10468] | 0.8080 | 0.00119 |
| ENSG00000141376 | BCAS3 | BCAS3 microtubule associated cell migration factor [Source:HGNC Symbol;Acc:HGNC:14347] | -0.9252 | 0.00119 |
| ENSG00000175093 | SPSB4 | splA/ryanodine receptor domain and SOCS box containing 4 [Source:HGNC Symbol;Acc:HGNC:30630] | -0.7024 | 0.00120 |
| ENSG00000164070 | HSPA4L | heat shock protein family A (Hsp70) member 4 like [Source:HGNC Symbol;Acc:HGNC:17041] | 0.5850 | 0.00123 |

| | | | | |
|---|---|---|---|---|
| ENSG00000133687 | TMTC1 | transmembrane O-mannosyltransferase targeting cadherins 1 [Source:HGNC Symbol;Acc:HGNC:24099] | 0.5896 | 0.00133 |
| ENSG00000203326 | ZNF525 | zinc finger protein 525 [Source:HGNC Symbol;Acc:HGNC:29423] | -0.5819 | 0.00133 |
| ENSG00000231721 | LINC-PINT | long intergenic non-protein coding RNA, p53 induced transcript [Source:HGNC Symbol;Acc:HGNC:26885] | 0.5975 | 0.00141 |
| ENSG00000111450 | STX2 | syntaxin 2 [Source:HGNC Symbol;Acc:HGNC:3403] | 0.5265 | 0.00142 |
| ENSG00000069667 | RORA | RAR related orphan receptor A [Source:HGNC Symbol;Acc:HGNC:10258] | 0.5765 | 0.00146 |
| ENSG00000187720 | THSD4 | thrombospondin type 1 domain containing 4 [Source:HGNC Symbol;Acc:HGNC:25835] | 0.5781 | 0.00148 |
| ENSG00000272323 | NA | novel transcript, antisense to TTC23L | 0.7002 | 0.00151 |
| ENSG00000171951 | SCG2 | secretogranin II [Source:HGNC Symbol;Acc:HGNC:10575] | 1.4429 | 0.00156 |
| ENSG00000100311 | PDGFB | platelet derived growth factor subunit B [Source:HGNC Symbol;Acc:HGNC:8800] | -1.2451 | 0.00166 |
| ENSG00000150977 | RILPL2 | Rab interacting lysosomal protein like 2 [Source:HGNC Symbol;Acc:HGNC:28787] | 0.5196 | 0.00169 |
| ENSG00000163958 | ZDHHC19 | zinc finger DHHC-type palmitoyltransferase 19 [Source:HGNC Symbol;Acc:HGNC:20713] | -0.7634 | 0.00170 |
| ENSG00000137872 | SEMA6D | semaphorin 6D [Source:HGNC Symbol;Acc:HGNC:16770] | 0.8086 | 0.00170 |
| ENSG00000183023 | SLC8A1 | solute carrier family 8 member A1 [Source:HGNC Symbol;Acc:HGNC:11068] | 1.2193 | 0.00175 |
| ENSG00000172164 | SNTB1 | syntrophin beta 1 [Source:HGNC Symbol;Acc:HGNC:11168] | 0.6291 | 0.00177 |
| ENSG00000138134 | STAMBPL1 | STAM binding protein like 1 [Source:HGNC Symbol;Acc:HGNC:24105] | 0.6064 | 0.00177 |
| ENSG00000198915 | RASGEF1A | RasGEF domain family member 1A [Source:HGNC Symbol;Acc:HGNC:24246] | 1.3897 | 0.00180 |
| ENSG00000260329 | NA | novel transcript, antisense to C12orf23 | -0.8690 | 0.00186 |
| ENSG00000280187 | NA | TEC | -0.5135 | 0.00186 |
| ENSG00000100889 | PCK2 | phosphoenolpyruvate carboxykinase 2, mitochondrial [Source:HGNC Symbol;Acc:HGNC:8725] | -0.5015 | 0.00193 |
| ENSG00000135953 | MFSD9 | major facilitator superfamily domain containing 9 [Source:HGNC Symbol;Acc:HGNC:28158] | 0.6249 | 0.00196 |
| ENSG00000248008 | NRAV | negative regulator of antiviral response [Source:HGNC Symbol;Acc:HGNC:48588] | -0.5287 | 0.00196 |
| ENSG00000156097 | GPR61 | G protein-coupled receptor 61 [Source:HGNC Symbol;Acc:HGNC:13300] | -0.8604 | 0.00202 |
| ENSG00000139344 | AMDHD1 | amidohydrolase domain containing 1 [Source:HGNC Symbol;Acc:HGNC:28577] | 0.6652 | 0.00202 |
| ENSG00000119737 | GPR75 | G protein-coupled receptor 75 [Source:HGNC Symbol;Acc:HGNC:4526] | 0.7189 | 0.00205 |
| ENSG00000136531 | SCN2A | sodium voltage-gated channel alpha subunit 2 [Source:HGNC Symbol;Acc:HGNC:10588] | 0.6103 | 0.00211 |
| ENSG00000173846 | PLK3 | polo like kinase 3 [Source:HGNC Symbol;Acc:HGNC:2154] | 0.9220 | 0.00211 |
| ENSG00000273702 | NA | novel transcript | -0.8142 | 0.00214 |
| ENSG00000258430 | NA | novel transcript, antisense to AKT1 | 0.9924 | 0.00217 |
| ENSG00000166415 | WDR72 | WD repeat domain 72 [Source:HGNC Symbol;Acc:HGNC:26790] | -0.6564 | 0.00218 |
| ENSG00000262001 | DLGAP1-AS2 | DLGAP1 antisense RNA 2 [Source:HGNC Symbol;Acc:HGNC:28146] | 0.5951 | 0.00221 |
| ENSG00000148082 | SHC3 | SHC adaptor protein 3 [Source:HGNC Symbol;Acc:HGNC:18181] | 1.0259 | 0.00226 |
| ENSG00000232300 | FAM215B | family with sequence similarity 215 member B [Source:HGNC Symbol;Acc:HGNC:43639] | 0.8046 | 0.00226 |
| ENSG00000171867 | PRNP | prion protein [Source:HGNC Symbol;Acc:HGNC:9449] | 0.5209 | 0.00237 |
| ENSG00000111674 | ENO2 | enolase 2 [Source:HGNC Symbol;Acc:HGNC:3353] | 0.5177 | 0.00240 |
| ENSG00000109762 | SNX25 | sorting nexin 25 [Source:HGNC Symbol;Acc:HGNC:21883] | 0.9018 | 0.00248 |
| ENSG00000155974 | GRIP1 | glutamate receptor interacting protein 1 [Source:HGNC Symbol;Acc:HGNC:18708] | 0.7572 | 0.00248 |
| ENSG00000158258 | CLSTN2 | calsyntenin 2 [Source:HGNC Symbol;Acc:HGNC:17448] | 1.1659 | 0.00256 |
| ENSG00000166068 | SPRED1 | sprouty related EVH1 domain containing 1 [Source:HGNC Symbol;Acc:HGNC:20249] | 0.7456 | 0.00258 |
| ENSG00000183044 | ABAT | 4-aminobutyrate aminotransferase [Source:HGNC Symbol;Acc:HGNC:23] | -0.5942 | 0.00259 |
| ENSG00000183688 | RFLNB | refilin B [Source:HGNC Symbol;Acc:HGNC:28705] | 1.0190 | 0.00272 |
| ENSG00000180875 | GREM2 | gremlin 2, DAN family BMP antagonist [Source:HGNC Symbol;Acc:HGNC:17655] | 1.2110 | 0.00276 |
| ENSG00000141639 | MAPK4 | mitogen-activated protein kinase 4 [Source:HGNC Symbol;Acc:HGNC:6878] | 0.6812 | 0.00281 |
| ENSG00000104332 | SFRP1 | secreted frizzled related protein 1 [Source:HGNC Symbol;Acc:HGNC:10776] | 1.7872 | 0.00282 |
| ENSG00000148339 | SLC25A25 | solute carrier family 25 member 25 [Source:HGNC Symbol;Acc:HGNC:20663] | 0.5453 | 0.00282 |
| ENSG00000279875 | NA | TEC | 1.6129 | 0.00283 |
| ENSG00000279041 | NA | TEC | 0.8894 | 0.00289 |
| ENSG00000107485 | GATA3 | GATA binding protein 3 [Source:HGNC Symbol;Acc:HGNC:4172] | -0.8424 | 0.00296 |
| ENSG00000141258 | SGSM2 | small G protein signaling modulator 2 [Source:HGNC Symbol;Acc:HGNC:29026] | 0.5090 | 0.00298 |
| ENSG00000135373 | EHF | ETS homologous factor [Source:HGNC Symbol;Acc:HGNC:3246] | -1.8159 | 0.00300 |
| ENSG00000171877 | FRMD5 | FERM domain containing 5 [Source:HGNC Symbol;Acc:HGNC:28214] | 1.1902 | 0.00304 |
| ENSG00000139428 | MMAB | metabolism of cobalamin associated B [Source:HGNC Symbol;Acc:HGNC:19331] | -0.5197 | 0.00313 |
| ENSG00000168243 | GNG4 | G protein subunit gamma 4 [Source:HGNC Symbol;Acc:HGNC:4407] | 0.8699 | 0.00320 |
| ENSG00000181409 | AATK | apoptosis associated tyrosine kinase [Source:HGNC Symbol;Acc:HGNC:21] | -0.6210 | 0.00327 |
| ENSG00000120833 | SOCS2 | suppressor of cytokine signaling 2 [Source:HGNC Symbol;Acc:HGNC:19382] | 1.3918 | 0.00327 |

| | | | | |
|---|---|---|---|---|
| ENSG00000109906 | ZBTB16 | zinc finger and BTB domain containing 16 [Source:HGNC Symbol;Acc:HGNC:12930] | -1.3419 | 0.00337 |
| ENSG00000124766 | SOX4 | SRY-box transcription factor 4 [Source:HGNC Symbol;Acc:HGNC:11200] | 0.7669 | 0.00345 |
| ENSG00000067141 | NEO1 | neogenin 1 [Source:HGNC Symbol;Acc:HGNC:7754] | 0.7672 | 0.00348 |
| ENSG00000231528 | FAM225A | family with sequence similarity 225 member A [Source:HGNC Symbol;Acc:HGNC:27855] | -0.7124 | 0.00349 |
| ENSG00000165030 | NFIL3 | nuclear factor, interleukin 3 regulated [Source:HGNC Symbol;Acc:HGNC:7787] | 0.7717 | 0.00353 |
| ENSG00000165475 | CRYL1 | crystallin lambda 1 [Source:HGNC Symbol;Acc:HGNC:18246] | -0.5343 | 0.00353 |
| ENSG00000215218 | UBE2QL1 | ubiquitin conjugating enzyme E2 Q family like 1 [Source:HGNC Symbol;Acc:HGNC:37269] | 0.5769 | 0.00353 |
| ENSG00000243364 | EFNA4 | ephrin A4 [Source:HGNC Symbol;Acc:HGNC:3224] | -0.5251 | 0.00353 |
| ENSG00000273044 | NA | novel transcript | 1.5232 | 0.00353 |
| ENSG00000134343 | ANO3 | anoctamin 3 [Source:HGNC Symbol;Acc:HGNC:14004] | 0.6431 | 0.00354 |
| ENSG00000079257 | LXN | latexin [Source:HGNC Symbol;Acc:HGNC:13347] | 0.6598 | 0.00362 |
| ENSG00000100346 | CACNA1I | calcium voltage-gated channel subunit alpha1 I [Source:HGNC Symbol;Acc:HGNC:1396] | -1.2877 | 0.00367 |
| ENSG00000143156 | NME7 | NME/NM23 family member 7 [Source:HGNC Symbol;Acc:HGNC:20461] | 0.5441 | 0.00368 |
| ENSG00000144647 | POMGNT2 | protein O-linked mannose N-acetylglucosaminyltransferase 2 (beta 1,4-) [Source:HGNC Symbol;Acc:HGNC:25902] | 0.5218 | 0.00370 |
| ENSG00000137502 | RAB30 | RAB30, member RAS oncogene family [Source:HGNC Symbol;Acc:HGNC:9770] | -0.5377 | 0.00373 |
| ENSG00000149571 | KIRREL3 | kirre like nephrin family adhesion molecule 3 [Source:HGNC Symbol;Acc:HGNC:23204] | 0.8754 | 0.00376 |
| ENSG00000120756 | PLS1 | plastin 1 [Source:HGNC Symbol;Acc:HGNC:9090] | 0.6642 | 0.00379 |
| ENSG00000258708 | SLC25A21-AS1 | SLC25A21 antisense RNA 1 [Source:HGNC Symbol;Acc:HGNC:44298] | -1.0644 | 0.00401 |
| ENSG00000168389 | MFSD2A | major facilitator superfamily domain containing 2A [Source:HGNC Symbol;Acc:HGNC:25897] | 1.2382 | 0.00414 |
| ENSG00000104267 | CA2 | carbonic anhydrase 2 [Source:HGNC Symbol;Acc:HGNC:1373] | 0.8758 | 0.00416 |
| ENSG00000257524 | NA | novel protein | 0.8446 | 0.00417 |
| ENSG00000228624 | HDAC2-AS2 | HDAC2 and HS3ST5 antisense RNA 2 [Source:HGNC Symbol;Acc:HGNC:43590] | -1.2825 | 0.00417 |
| ENSG00000184408 | KCND2 | potassium voltage-gated channel subfamily D member 2 [Source:HGNC Symbol;Acc:HGNC:6238] | 3.5146 | 0.00419 |
| ENSG00000105516 | DBP | D-box binding PAR bZIP transcription factor [Source:HGNC Symbol;Acc:HGNC:2697] | -0.6541 | 0.00426 |
| ENSG00000138193 | PLCE1 | phospholipase C epsilon 1 [Source:HGNC Symbol;Acc:HGNC:17175] | -0.7208 | 0.00426 |
| ENSG00000092421 | SEMA6A | semaphorin 6A [Source:HGNC Symbol;Acc:HGNC:10738] | -0.7581 | 0.00442 |
| ENSG00000115850 | LCT | lactase [Source:HGNC Symbol;Acc:HGNC:6530] | 1.0182 | 0.00450 |
| ENSG00000124208 | TMEM189-UBE2V1 | TMEM189-UBE2V1 readthrough [Source:HGNC Symbol;Acc:HGNC:33521] | 8.5414 | 0.00460 |
| ENSG00000189056 | RELN | reelin [Source:HGNC Symbol;Acc:HGNC:9957] | 0.6330 | 0.00471 |
| ENSG00000221923 | ZNF880 | zinc finger protein 880 [Source:HGNC Symbol;Acc:HGNC:37249] | -0.6827 | 0.00474 |
| ENSG00000121207 | LRAT | lecithin retinol acyltransferase [Source:HGNC Symbol;Acc:HGNC:6685] | -1.9265 | 0.00486 |
| ENSG00000124479 | NDP | norrin cystine knot growth factor NDP [Source:HGNC Symbol;Acc:HGNC:7678] | -0.9495 | 0.00490 |
| ENSG00000136367 | ZFHX2 | zinc finger homeobox 2 [Source:HGNC Symbol;Acc:HGNC:20152] | 0.7253 | 0.00499 |
| ENSG00000077943 | ITGA8 | integrin subunit alpha 8 [Source:HGNC Symbol;Acc:HGNC:6144] | 0.6277 | 0.00499 |
| ENSG00000127955 | GNAI1 | G protein subunit alpha i1 [Source:HGNC Symbol;Acc:HGNC:4384] | 0.5109 | 0.00511 |
| ENSG00000145730 | PAM | peptidylglycine alpha-amidating monooxygenase [Source:HGNC Symbol;Acc:HGNC:8596] | 0.6211 | 0.00521 |
| ENSG00000146233 | CYP39A1 | cytochrome P450 family 39 subfamily A member 1 [Source:HGNC Symbol;Acc:HGNC:17449] | 0.6408 | 0.00532 |
| ENSG00000176654 | NANOGP1 | Nanog homeobox pseudogene 1 [Source:HGNC Symbol;Acc:HGNC:23099] | -0.8601 | 0.00552 |
| ENSG00000183837 | PNMA3 | PNMA family member 3 [Source:HGNC Symbol;Acc:HGNC:18742] | -1.0620 | 0.00552 |
| ENSG00000131650 | KREMEN2 | kringle containing transmembrane protein 2 [Source:HGNC Symbol;Acc:HGNC:18797] | -0.5758 | 0.00557 |
| ENSG00000106100 | NOD1 | nucleotide binding oligomerization domain containing 1 [Source:HGNC Symbol;Acc:HGNC:16390] | -1.0225 | 0.00568 |
| ENSG00000186446 | ZNF501 | zinc finger protein 501 [Source:HGNC Symbol;Acc:HGNC:23717] | -0.6325 | 0.00572 |
| ENSG00000247708 | STX18-AS1 | STX18 antisense RNA 1 (head to head) [Source:HGNC Symbol;Acc:HGNC:48877] | -0.5189 | 0.00597 |
| ENSG00000166979 | EVA1C | eva-1 homolog C [Source:HGNC Symbol;Acc:HGNC:13239] | 0.9172 | 0.00599 |
| ENSG00000175600 | SUGCT | succinyl-CoA:glutarate-CoA transferase [Source:HGNC Symbol;Acc:HGNC:16001] | 1.5430 | 0.00612 |
| ENSG00000253554 | LINC01414 | long intergenic non-protein coding RNA 1414 [Source:HGNC Symbol;Acc:HGNC:50707] | -1.6385 | 0.00625 |
| ENSG00000270112 | NA | novel transcript, antisense to ST8SIA5 | -0.5583 | 0.00627 |
| ENSG00000175832 | ETV4 | ETS variant transcription factor 4 [Source:HGNC Symbol;Acc:HGNC:3493] | 2.3178 | 0.00627 |
| ENSG00000140015 | KCNH5 | potassium voltage-gated channel subfamily H member 5 [Source:HGNC Symbol;Acc:HGNC:6254] | 1.4716 | 0.00629 |
| ENSG00000072864 | NDE1 | nudE neurodevelopment protein 1 [Source:HGNC Symbol;Acc:HGNC:17619] | -0.5014 | 0.00640 |
| ENSG00000110328 | GALNT18 | polypeptide N-acetylgalactosaminyltransferase 18 [Source:HGNC Symbol;Acc:HGNC:30488] | -1.1232 | 0.00643 |
| ENSG00000196273 | LINC00523 | long intergenic non-protein coding RNA 523 [Source:HGNC Symbol;Acc:HGNC:20117] | -0.5820 | 0.00669 |
| ENSG00000117595 | IRF6 | interferon regulatory factor 6 [Source:HGNC Symbol;Acc:HGNC:6121] | -0.6454 | 0.00678 |
| ENSG00000081803 | CADPS2 | calcium dependent secretion activator 2 [Source:HGNC Symbol;Acc:HGNC:16018] | 0.6233 | 0.00707 |

| | | | | |
|---|---|---|---|---|
| ENSG00000173210 | ABLIM3 | actin binding LIM protein family member 3 [Source:HGNC Symbol;Acc:HGNC:29132] | -0.6300 | 0.00717 |
| ENSG00000034152 | MAP2K3 | mitogen-activated protein kinase kinase 3 [Source:HGNC Symbol;Acc:HGNC:6843] | 0.5988 | 0.00719 |
| ENSG00000136297 | MMD2 | monocyte to macrophage differentiation associated 2 [Source:HGNC Symbol;Acc:HGNC:30133] | -0.7010 | 0.00738 |
| ENSG00000153064 | BANK1 | B cell scaffold protein with ankyrin repeats 1 [Source:HGNC Symbol;Acc:HGNC:18233] | 1.5683 | 0.00738 |
| ENSG00000269416 | LINC01224 | long intergenic non-protein coding RNA 1224 [Source:HGNC Symbol;Acc:HGNC:49676] | -0.6360 | 0.00758 |
| ENSG00000122786 | CALD1 | caldesmon 1 [Source:HGNC Symbol;Acc:HGNC:1441] | 0.7007 | 0.00758 |
| ENSG00000204934 | ATP6V0E2-AS1 | ATP6V0E2 antisense RNA 1 [Source:HGNC Symbol;Acc:HGNC:44180] | 0.5290 | 0.00758 |
| ENSG00000283050 | GTF2IP12 | general transcription factor IIi pseudogene 12 [Source:HGNC Symbol;Acc:HGNC:51723] | -0.6414 | 0.00762 |
| ENSG00000047230 | CTPS2 | CTP synthase 2 [Source:HGNC Symbol;Acc:HGNC:2520] | -0.5478 | 0.00786 |
| ENSG00000166963 | MAP1A | microtubule associated protein 1A [Source:HGNC Symbol;Acc:HGNC:6835] | -0.5639 | 0.00792 |
| ENSG00000178297 | TMPRSS9 | transmembrane serine protease 9 [Source:HGNC Symbol;Acc:HGNC:30079] | -0.7115 | 0.00799 |
| ENSG00000129355 | CDKN2D | cyclin dependent kinase inhibitor 2D [Source:HGNC Symbol;Acc:HGNC:1790] | -0.5362 | 0.00805 |
| ENSG00000225465 | RFPL1S | RFPL1 antisense RNA 1 [Source:HGNC Symbol;Acc:HGNC:9978] | -0.9992 | 0.00827 |
| ENSG00000258405 | ZNF578 | zinc finger protein 578 [Source:HGNC Symbol;Acc:HGNC:26449] | -0.8352 | 0.00839 |
| ENSG00000129595 | EPB41L4A | erythrocyte membrane protein band 4.1 like 4A [Source:HGNC Symbol;Acc:HGNC:13278] | 0.5130 | 0.00847 |
| ENSG00000146674 | IGFBP3 | insulin like growth factor binding protein 3 [Source:HGNC Symbol;Acc:HGNC:5472] | 1.2154 | 0.00853 |
| ENSG00000117707 | PROX1 | prospero homeobox 1 [Source:HGNC Symbol;Acc:HGNC:9459] | 0.6230 | 0.00857 |
| ENSG00000180616 | SSTR2 | somatostatin receptor 2 [Source:HGNC Symbol;Acc:HGNC:11331] | 0.7342 | 0.00876 |
| ENSG00000196335 | STK31 | serine/threonine kinase 31 [Source:HGNC Symbol;Acc:HGNC:11407] | -2.0148 | 0.00878 |
| ENSG00000150627 | WDR17 | WD repeat domain 17 [Source:HGNC Symbol;Acc:HGNC:16661] | -1.2028 | 0.00898 |
| ENSG00000145794 | MEGF10 | multiple EGF like domains 10 [Source:HGNC Symbol;Acc:HGNC:29634] | 1.5031 | 0.00929 |
| ENSG00000274718 | NA | novel transcript | 0.8730 | 0.00933 |
| ENSG00000131831 | RAI2 | retinoic acid induced 2 [Source:HGNC Symbol;Acc:HGNC:9835] | -0.5248 | 0.00936 |
| ENSG00000105639 | JAK3 | Janus kinase 3 [Source:HGNC Symbol;Acc:HGNC:6193] | -0.9407 | 0.00941 |
| ENSG00000049246 | PER3 | period circadian regulator 3 [Source:HGNC Symbol;Acc:HGNC:8847] | -0.8664 | 0.00945 |
| ENSG00000224790 | NA | novel transcript | 0.6629 | 0.00946 |
| ENSG00000164761 | TNFRSF11B | TNF receptor superfamily member 11b [Source:HGNC Symbol;Acc:HGNC:11909] | 0.5777 | 0.00982 |
| ENSG00000197444 | OGDHL | oxoglutarate dehydrogenase like [Source:HGNC Symbol;Acc:HGNC:25590] | -0.6895 | 0.00984 |
| ENSG00000099860 | GADD45B | growth arrest and DNA damage inducible beta [Source:HGNC Symbol;Acc:HGNC:4096] | -0.5446 | 0.00988 |
| ENSG00000197321 | SVIL | supervillin [Source:HGNC Symbol;Acc:HGNC:11480] | -1.1028 | 0.00988 |
| ENSG00000013392 | RWDD2A | RWD domain containing 2A [Source:HGNC Symbol;Acc:HGNC:21385] | 0.5272 | 0.00992 |
| ENSG00000144847 | IGSF11 | immunoglobulin superfamily member 11 [Source:HGNC Symbol;Acc:HGNC:16669] | -0.7389 | 0.00995 |
| ENSG00000246627 | CACNA1C-AS1 | CACNA1C antisense RNA 1 [Source:HGNC Symbol;Acc:HGNC:40119] | 1.8323 | 0.00998 |
| ENSG00000095303 | PTGS1 | prostaglandin-endoperoxide synthase 1 [Source:HGNC Symbol;Acc:HGNC:9604] | 0.8853 | 0.00998 |
| ENSG00000265148 | TSPOAP1-AS1 | TSPOAP1, SUPT4H1 and RNF43 antisense RNA 1 [Source:HGNC Symbol;Acc:HGNC:44148] | -0.7535 | 0.01030 |
| ENSG00000126215 | XRCC3 | X-ray repair cross complementing 3 [Source:HGNC Symbol;Acc:HGNC:12830] | -0.6061 | 0.01035 |
| ENSG00000132003 | ZSWIM4 | zinc finger SWIM-type containing 4 [Source:HGNC Symbol;Acc:HGNC:25704] | 0.6610 | 0.01037 |
| ENSG00000185022 | MAFF | MAF bZIP transcription factor F [Source:HGNC Symbol;Acc:HGNC:6780] | 1.3723 | 0.01047 |
| ENSG00000189120 | SP6 | Sp6 transcription factor [Source:HGNC Symbol;Acc:HGNC:14530] | -0.7184 | 0.01049 |
| ENSG00000279865 | NA | TEC | -1.2699 | 0.01054 |
| ENSG00000223553 | SMPD4P1 | sphingomyelin phosphodiesterase 4 pseudogene 1 [Source:HGNC Symbol;Acc:HGNC:39673] | -1.7044 | 0.01072 |
| ENSG00000250358 | LINC02200 | long intergenic non-protein coding RNA 2200 [Source:HGNC Symbol;Acc:HGNC:53066] | -1.3517 | 0.01072 |
| ENSG00000287861 | NA | novel transcript | -0.6974 | 0.01074 |
| ENSG00000250420 | AACSP1 | acetoacetyl-CoA synthetase pseudogene 1 [Source:HGNC Symbol;Acc:HGNC:18226] | -0.6742 | 0.01090 |
| ENSG00000184557 | SOCS3 | suppressor of cytokine signaling 3 [Source:HGNC Symbol;Acc:HGNC:19391] | 1.3226 | 0.01095 |
| ENSG00000237651 | C2orf74 | chromosome 2 open reading frame 74 [Source:HGNC Symbol;Acc:HGNC:34439] | -0.6746 | 0.01099 |
| ENSG00000178233 | TMEM151B | transmembrane protein 151B [Source:HGNC Symbol;Acc:HGNC:21315] | 0.7754 | 0.01105 |
| ENSG00000088970 | KIZ | kizuna centrosomal protein [Source:HGNC Symbol;Acc:HGNC:15865] | 0.5341 | 0.01123 |
| ENSG00000169193 | CCDC126 | coiled-coil domain containing 126 [Source:HGNC Symbol;Acc:HGNC:22398] | 0.8955 | 0.01134 |
| ENSG00000227467 | LINC01537 | long intergenic non-protein coding RNA 1537 [Source:HGNC Symbol;Acc:HGNC:51301] | -1.0785 | 0.01134 |
| ENSG00000132204 | LINC00470 | long intergenic non-protein coding RNA 470 [Source:HGNC Symbol;Acc:HGNC:1225] | -0.8128 | 0.01148 |
| ENSG00000196653 | ZNF502 | zinc finger protein 502 [Source:HGNC Symbol;Acc:HGNC:23718] | -0.5105 | 0.01159 |
| ENSG00000187098 | MITF | melanocyte inducing transcription factor [Source:HGNC Symbol;Acc:HGNC:7105] | 0.9402 | 0.01159 |
| ENSG00000160401 | CFAP157 | cilia and flagella associated protein 157 [Source:HGNC Symbol;Acc:HGNC:27843] | -0.7163 | 0.01162 |

| | | | | |
|---|---|---|---|---|
| ENSG00000256870 | SLC5A8 | solute carrier family 5 member 8 [Source:HGNC Symbol;Acc:HGNC:19119] | -1.5904 | 0.01166 |
| ENSG00000279720 | SOWAHCP2 | SOWAHC pseudogene 2 [Source:HGNC Symbol;Acc:HGNC:54506] | -0.7934 | 0.01167 |
| ENSG00000187140 | FOXD3 | forkhead box D3 [Source:HGNC Symbol;Acc:HGNC:3804] | 0.9116 | 0.01215 |
| ENSG00000162944 | RFTN2 | raftlin family member 2 [Source:HGNC Symbol;Acc:HGNC:26402] | 0.8645 | 0.01225 |
| ENSG00000074966 | TXK | TXK tyrosine kinase [Source:HGNC Symbol;Acc:HGNC:12434] | 0.6355 | 0.01226 |
| ENSG00000227906 | SNAP25-AS1 | SNAP25 antisense RNA 1 [Source:HGNC Symbol;Acc:HGNC:44312] | 0.9500 | 0.01249 |
| ENSG00000148120 | AOPEP | aminopeptidase O (putative) [Source:HGNC Symbol;Acc:HGNC:1361] | -0.5367 | 0.01252 |
| ENSG00000223749 | MIR503HG | MIR503 host gene [Source:HGNC Symbol;Acc:HGNC:28258] | 0.5483 | 0.01252 |
| ENSG00000103647 | CORO2B | coronin 2B [Source:HGNC Symbol;Acc:HGNC:2256] | -0.6512 | 0.01257 |
| ENSG00000137501 | SYTL2 | synaptotagmin like 2 [Source:HGNC Symbol;Acc:HGNC:15585] | -0.5609 | 0.01257 |
| ENSG00000175785 | PRIMA1 | proline rich membrane anchor 1 [Source:HGNC Symbol;Acc:HGNC:18319] | -0.7185 | 0.01262 |
| ENSG00000204740 | MALRD1 | MAM and LDL receptor class A domain containing 1 [Source:HGNC Symbol;Acc:HGNC:24331] | -0.8331 | 0.01289 |
| ENSG00000132837 | DMGDH | dimethylglycine dehydrogenase [Source:HGNC Symbol;Acc:HGNC:24475] | -1.3137 | 0.01291 |
| ENSG00000269486 | ERVK9-11 | endogenous retrovirus group K9, member 11 [Source:HGNC Symbol;Acc:HGNC:51160] | -0.5031 | 0.01306 |
| ENSG00000204442 | FAM155A | family with sequence similarity 155 member A [Source:HGNC Symbol;Acc:HGNC:33877] | 0.5454 | 0.01322 |
| ENSG00000136155 | SCEL | sciellin [Source:HGNC Symbol;Acc:HGNC:10573] | -1.1382 | 0.01323 |
| ENSG00000181634 | TNFSF15 | TNF superfamily member 15 [Source:HGNC Symbol;Acc:HGNC:11931] | 2.2720 | 0.01327 |
| ENSG00000132669 | RIN2 | Ras and Rab interactor 2 [Source:HGNC Symbol;Acc:HGNC:18750] | -0.6490 | 0.01360 |
| ENSG00000285907 | NA | novel transcript | -0.8562 | 0.01391 |
| ENSG00000027075 | PRKCH | protein kinase C eta [Source:HGNC Symbol;Acc:HGNC:9403] | 0.8474 | 0.01402 |
| ENSG00000110427 | KIAA1549L | KIAA1549 like [Source:HGNC Symbol;Acc:HGNC:24836] | 0.7590 | 0.01406 |
| ENSG00000139211 | AMIGO2 | adhesion molecule with Ig like domain 2 [Source:HGNC Symbol;Acc:HGNC:24073] | 0.7718 | 0.01406 |
| ENSG00000167550 | RHEBL1 | RHEB like 1 [Source:HGNC Symbol;Acc:HGNC:21166] | 0.6958 | 0.01406 |
| ENSG00000018236 | CNTN1 | contactin 1 [Source:HGNC Symbol;Acc:HGNC:2171] | 0.7423 | 0.01445 |
| ENSG00000089041 | P2RX7 | purinergic receptor P2X 7 [Source:HGNC Symbol;Acc:HGNC:8537] | -0.6338 | 0.01476 |
| ENSG00000187678 | SPRY4 | sprouty RTK signaling antagonist 4 [Source:HGNC Symbol;Acc:HGNC:15533] | 2.2038 | 0.01514 |
| ENSG00000130413 | STK33 | serine/threonine kinase 33 [Source:HGNC Symbol;Acc:HGNC:14568] | -0.5291 | 0.01525 |
| ENSG00000188010 | MORN2 | MORN repeat containing 2 [Source:HGNC Symbol;Acc:HGNC:30166] | 0.5386 | 0.01526 |
| ENSG00000187736 | NHEJ1 | non-homologous end joining factor 1 [Source:HGNC Symbol;Acc:HGNC:25737] | 0.5134 | 0.01542 |
| ENSG00000255529 | POLR2M | RNA polymerase II subunit M [Source:HGNC Symbol;Acc:HGNC:14862] | 0.7017 | 0.01580 |
| ENSG00000163697 | APBB2 | amyloid beta precursor protein binding family B member 2 [Source:HGNC Symbol;Acc:HGNC:582] | 0.5300 | 0.01600 |
| ENSG00000251562 | MALAT1 | metastasis associated lung adenocarcinoma transcript 1 [Source:HGNC Symbol;Acc:HGNC:29665] | 0.5907 | 0.01604 |
| ENSG00000146243 | IRAK1BP1 | interleukin 1 receptor associated kinase 1 binding protein 1 [Source:HGNC Symbol;Acc:HGNC:17368] | -0.5632 | 0.01615 |
| ENSG00000205464 | ATP6AP1L | ATPase H+ transporting accessory protein 1 like [Source:HGNC Symbol;Acc:HGNC:28091] | -0.5269 | 0.01620 |
| ENSG00000171105 | INSR | insulin receptor [Source:HGNC Symbol;Acc:HGNC:6091] | 0.5632 | 0.01651 |
| ENSG00000163072 | NOSTRIN | nitric oxide synthase trafficking [Source:HGNC Symbol;Acc:HGNC:20203] | -0.5911 | 0.01655 |
| ENSG00000162620 | LRRIQ3 | leucine rich repeats and IQ motif containing 3 [Source:HGNC Symbol;Acc:HGNC:28318] | -0.5718 | 0.01662 |
| ENSG00000203667 | COX20 | cytochrome c oxidase assembly factor COX20 [Source:HGNC Symbol;Acc:HGNC:26970] | 0.5128 | 0.01666 |
| ENSG00000088899 | LZTS3 | leucine zipper tumor suppressor family member 3 [Source:HGNC Symbol;Acc:HGNC:30139] | 0.5269 | 0.01687 |
| ENSG00000163840 | DTX3L | deltex E3 ubiquitin ligase 3L [Source:HGNC Symbol;Acc:HGNC:30323] | -0.5407 | 0.01688 |
| ENSG00000128039 | SRD5A3 | steroid 5 alpha-reductase 3 [Source:HGNC Symbol;Acc:HGNC:25812] | -0.6046 | 0.01704 |
| ENSG00000128849 | CGNL1 | cingulin like 1 [Source:HGNC Symbol;Acc:HGNC:25931] | 0.6110 | 0.01731 |
| ENSG00000197122 | SRC | SRC proto-oncogene, non-receptor tyrosine kinase [Source:HGNC Symbol;Acc:HGNC:11283] | -1.0995 | 0.01753 |
| ENSG00000163623 | NKX6-1 | NK6 homeobox 1 [Source:HGNC Symbol;Acc:HGNC:7839] | 0.7674 | 0.01758 |
| ENSG00000144868 | TMEM108 | transmembrane protein 108 [Source:HGNC Symbol;Acc:HGNC:28451] | 0.5423 | 0.01836 |
| ENSG00000039523 | RIPOR1 | RHO family interacting cell polarization regulator 1 [Source:HGNC Symbol;Acc:HGNC:25836] | -0.5506 | 0.01836 |
| ENSG00000005187 | ACSM3 | acyl-CoA synthetase medium chain family member 3 [Source:HGNC Symbol;Acc:HGNC:10522] | -0.7302 | 0.01849 |
| ENSG00000106003 | LFNG | LFNG O-fucosylpeptide 3-beta-N-acetylglucosaminyltransferase [Source:HGNC Symbol;Acc:HGNC:6560] | 0.9740 | 0.01852 |
| ENSG00000188483 | IER5L | immediate early response 5 like [Source:HGNC Symbol;Acc:HGNC:23679] | -0.6192 | 0.01889 |
| ENSG00000100285 | NEFH | neurofilament heavy [Source:HGNC Symbol;Acc:HGNC:7737] | -0.8668 | 0.01909 |
| ENSG00000170345 | FOS | Fos proto-oncogene, AP-1 transcription factor subunit [Source:HGNC Symbol;Acc:HGNC:3796] | 2.6072 | 0.01911 |
| ENSG00000228725 | MTND2P12 | MT-ND2 pseudogene 12 [Source:HGNC Symbol;Acc:HGNC:42113] | 1.4835 | 0.01925 |
| ENSG00000142621 | FHAD1 | forkhead associated phosphopeptide binding domain 1 [Source:HGNC Symbol;Acc:HGNC:29408] | -0.6154 | 0.01927 |
| ENSG00000163646 | CLRN1 | clarin 1 [Source:HGNC Symbol;Acc:HGNC:12605] | 0.5533 | 0.01963 |

| | | | | |
|---|---|---|---|---|
| ENSG00000113296 | THBS4 | thrombospondin 4 [Source:HGNC Symbol;Acc:HGNC:11788] | -0.6424 | 0.01977 |
| ENSG00000204052 | LRRC73 | leucine rich repeat containing 73 [Source:HGNC Symbol;Acc:HGNC:21375] | 0.5222 | 0.01977 |
| ENSG00000227124 | ZNF717 | zinc finger protein 717 [Source:HGNC Symbol;Acc:HGNC:29448] | -0.5067 | 0.02000 |
| ENSG00000260793 | NA | novel transcript, antisense to UBTF | 0.6806 | 0.02009 |
| ENSG00000117152 | RGS4 | regulator of G protein signaling 4 [Source:HGNC Symbol;Acc:HGNC:10000] | 2.2755 | 0.02025 |
| ENSG00000123364 | HOXC13 | homeobox C13 [Source:HGNC Symbol;Acc:HGNC:5125] | 0.5887 | 0.02065 |
| ENSG00000125657 | TNFSF9 | TNF superfamily member 9 [Source:HGNC Symbol;Acc:HGNC:11939] | 0.8792 | 0.02065 |
| ENSG00000113389 | 3 | natriuretic peptide receptor 3 [Source:HGNC Symbol;Acc:HGNC:7945] | 0.5692 | 0.02079 |
| ENSG00000160172 | FAM86C2P | family with sequence similarity 86 member C2, pseudogene [Source:HGNC Symbol;Acc:HGNC:42392] | -1.0856 | 0.02081 |
| ENSG00000147174 | GCNA | germ cell nuclear acidic peptidase [Source:HGNC Symbol;Acc:HGNC:15805] | 1.3074 | 0.02104 |
| ENSG00000168016 | TRANK1 | tetratricopeptide repeat and ankyrin repeat containing 1 [Source:HGNC Symbol;Acc:HGNC:29011] | -0.9999 | 0.02104 |
| ENSG00000185585 | OLFML2A | olfactomedin like 2A [Source:HGNC Symbol;Acc:HGNC:27270] | -0.8812 | 0.02104 |
| ENSG00000162755 | KLHDC9 | kelch domain containing 9 [Source:HGNC Symbol;Acc:HGNC:28489] | -0.7552 | 0.02106 |
| ENSG00000087085 | ACHE | acetylcholinesterase (Cartwright blood group) [Source:HGNC Symbol;Acc:HGNC:108] | 0.7899 | 0.02106 |
| ENSG00000163617 | CCDC191 | coiled-coil domain containing 191 [Source:HGNC Symbol;Acc:HGNC:29272] | -0.6098 | 0.02131 |
| ENSG00000064309 | CDON | cell adhesion associated, oncogene regulated [Source:HGNC Symbol;Acc:HGNC:17104] | -0.6258 | 0.02134 |
| ENSG00000126733 | DACH2 | dachshund family transcription factor 2 [Source:HGNC Symbol;Acc:HGNC:16814] | 0.6904 | 0.02134 |
| ENSG00000163644 | PPM1K | protein phosphatase, Mg2+/Mn2+ dependent 1K [Source:HGNC Symbol;Acc:HGNC:25415] | 0.5049 | 0.02136 |
| ENSG00000235903 | CPB2-AS1 | CPB2 antisense RNA 1 [Source:HGNC Symbol;Acc:HGNC:39898] | 0.7974 | 0.02216 |
| ENSG00000135919 | SERPINE2 | serpin family E member 2 [Source:HGNC Symbol;Acc:HGNC:8951] | 0.6049 | 0.02271 |
| ENSG00000093217 | XYLB | xylulokinase [Source:HGNC Symbol;Acc:HGNC:12839] | -0.7282 | 0.02281 |
| ENSG00000177508 | IRX3 | iroquois homeobox 3 [Source:HGNC Symbol;Acc:HGNC:14360] | 0.5076 | 0.02281 |
| ENSG00000227225 | MTND1P14 | MT-ND1 pseudogene 14 [Source:HGNC Symbol;Acc:HGNC:42063] | 1.3173 | 0.02281 |
| ENSG00000141542 | RAB40B | RAB40B, member RAS oncogene family [Source:HGNC Symbol;Acc:HGNC:18284] | -0.5912 | 0.02294 |
| ENSG00000141449 | GREB1L | GREB1 like retinoic acid receptor coactivator [Source:HGNC Symbol;Acc:HGNC:31042] | -0.8177 | 0.02301 |
| ENSG00000150967 | ABCB9 | ATP binding cassette subfamily B member 9 [Source:HGNC Symbol;Acc:HGNC:50] | -0.6529 | 0.02321 |
| ENSG00000167207 | NOD2 | nucleotide binding oligomerization domain containing 2 [Source:HGNC Symbol;Acc:HGNC:5331] | -0.5387 | 0.02324 |
| ENSG00000159712 | ANKRD18CP | ankyrin repeat domain 18C, pseudogene [Source:HGNC Symbol;Acc:HGNC:43601] | 1.3509 | 0.02339 |
| ENSG00000067057 | PFKP | phosphofructokinase, platelet [Source:HGNC Symbol;Acc:HGNC:8878] | 0.5629 | 0.02350 |
| ENSG00000056972 | TRAF3IP2 | TRAF3 interacting protein 2 [Source:HGNC Symbol;Acc:HGNC:1343] | -0.6282 | 0.02353 |
| ENSG00000100314 | CABP7 | calcium binding protein 7 [Source:HGNC Symbol;Acc:HGNC:20834] | -0.8025 | 0.02353 |
| ENSG00000168671 | UGT3A2 | UDP glycosyltransferase family 3 member A2 [Source:HGNC Symbol;Acc:HGNC:27266] | 0.5561 | 0.02381 |
| ENSG00000141506 | PIK3R5 | phosphoinositide-3-kinase regulatory subunit 5 [Source:HGNC Symbol;Acc:HGNC:30035] | -1.3582 | 0.02403 |
| ENSG00000231625 | SLC47A1P2 | SLC47A1 pseudogene 2 [Source:HGNC Symbol;Acc:HGNC:53866] | 1.0306 | 0.02422 |
| ENSG00000185168 | LINC00482 | long intergenic non-protein coding RNA 482 [Source:HGNC Symbol;Acc:HGNC:26816] | -1.3356 | 0.02426 |
| ENSG00000007866 | TEAD3 | TEA domain transcription factor 3 [Source:HGNC Symbol;Acc:HGNC:11716] | 0.7938 | 0.02435 |
| ENSG00000118242 | MREG | melanoregulin [Source:HGNC Symbol;Acc:HGNC:25478] | -0.7089 | 0.02448 |
| ENSG00000168539 | CHRM1 | cholinergic receptor muscarinic 1 [Source:HGNC Symbol;Acc:HGNC:1950] | 1.6488 | 0.02508 |
| ENSG00000251692 | PTX4 | pentraxin 4 [Source:HGNC Symbol;Acc:HGNC:14171] | -1.0493 | 0.02513 |
| ENSG00000261037 | NA | novel transcript | 0.7826 | 0.02542 |
| ENSG00000247809 | NR2F2-AS1 | NR2F2 antisense RNA 1 [Source:HGNC Symbol;Acc:HGNC:44222] | -1.7374 | 0.02547 |
| ENSG00000148219 | ASTN2 | astrotactin 2 [Source:HGNC Symbol;Acc:HGNC:17021] | -0.5885 | 0.02609 |
| ENSG00000122641 | INHBA | inhibin subunit beta A [Source:HGNC Symbol;Acc:HGNC:6066] | 1.6596 | 0.02706 |
| ENSG00000204020 | LIPN | lipase family member N [Source:HGNC Symbol;Acc:HGNC:23452] | 1.7648 | 0.02706 |
| ENSG00000233485 | FHAD1-AS1 | FHAD1 antisense RNA 1 [Source:HGNC Symbol;Acc:HGNC:41241] | -0.6422 | 0.02719 |
| ENSG00000151689 | INPP1 | inositol polyphosphate-1-phosphatase [Source:HGNC Symbol;Acc:HGNC:6071] | 0.5527 | 0.02779 |
| ENSG00000154545 | MAGED4 | MAGE family member D4 [Source:HGNC Symbol;Acc:HGNC:23793] | 0.5659 | 0.02910 |
| ENSG00000111886 | GABRR2 | gamma-aminobutyric acid type A receptor subunit rho2 [Source:HGNC Symbol;Acc:HGNC:4091] | 1.1221 | 0.02972 |
| ENSG00000253882 | NA | family with sequence similarity 115, member C (FAM115C) pseudogene | -0.7715 | 0.03043 |
| ENSG00000131242 | RAB11FIP4 | RAB11 family interacting protein 4 [Source:HGNC Symbol;Acc:HGNC:30267] | -0.8430 | 0.03067 |
| ENSG00000277715 | NA | novel transcript, antisense to CKAP4 | -0.8870 | 0.03091 |
| ENSG00000076258 | FMO4 | flavin containing dimethylaniline monoxygenase 4 [Source:HGNC Symbol;Acc:HGNC:3772] | -1.2728 | 0.03095 |
| ENSG00000144285 | SCN1A | sodium voltage-gated channel alpha subunit 1 [Source:HGNC Symbol;Acc:HGNC:10585] | -0.7358 | 0.03100 |
| ENSG00000169184 | MN1 | MN1 proto-oncogene, transcriptional regulator [Source:HGNC Symbol;Acc:HGNC:7180] | 1.6996 | 0.03118 |

| | | | | |
|---|---|---|---|---|
| ENSG00000196696 | NA | nuclear pore complex-interacting protein [Source:NCBI gene (formerly Entrezgene);Acc:283970] | 0.5007 | 0.03169 |
| ENSG00000172828 | CES3 | carboxylesterase 3 [Source:HGNC Symbol;Acc:HGNC:1865] | -0.6144 | 0.03178 |
| ENSG00000108821 | COL1A1 | collagen type I alpha 1 chain [Source:HGNC Symbol;Acc:HGNC:2197] | -0.7422 | 0.03185 |
| ENSG00000166432 | ZMAT1 | zinc finger matrin-type 1 [Source:HGNC Symbol;Acc:HGNC:29377] | -0.6724 | 0.03231 |
| ENSG00000167705 | RILP | Rab interacting lysosomal protein [Source:HGNC Symbol;Acc:HGNC:30266] | 1.0855 | 0.03249 |
| ENSG00000159247 | TUBBP5 | tubulin beta pseudogene 5 [Source:HGNC Symbol;Acc:HGNC:23674] | -1.3313 | 0.03255 |
| ENSG00000115226 | FNDC4 | fibronectin type III domain containing 4 [Source:HGNC Symbol;Acc:HGNC:20239] | 0.5517 | 0.03322 |
| ENSG00000168542 | COL3A1 | collagen type III alpha 1 chain [Source:HGNC Symbol;Acc:HGNC:2201] | 1.3882 | 0.03356 |
| ENSG00000117318 | ID3 | inhibitor of DNA binding 3, HLH protein [Source:HGNC Symbol;Acc:HGNC:5362] | -1.1115 | 0.03384 |
| ENSG00000286282 | NA | novel transcript, antisense to NOV | -0.7056 | 0.03385 |
| ENSG00000177181 | RIMKLA | ribosomal modification protein rimK like family member A [Source:HGNC Symbol;Acc:HGNC:28725] | 0.6535 | 0.03419 |
| ENSG00000152377 | SPOCK1 | SPARC (osteonectin), cwcv and kazal like domains proteoglycan 1 [Source:HGNC Symbol;Acc:HGNC:11251] | -0.5910 | 0.03449 |
| ENSG00000239445 | ST3GAL6-AS1 | ST3GAL6 antisense RNA 1 [Source:HGNC Symbol;Acc:HGNC:40828] | -0.9014 | 0.03484 |
| ENSG00000163093 | BBS5 | Bardet-Biedl syndrome 5 [Source:HGNC Symbol;Acc:HGNC:970] | -0.5983 | 0.03489 |
| ENSG00000232160 | RAP2C-AS1 | RAP2C antisense RNA 1 [Source:HGNC Symbol;Acc:HGNC:40957] | -0.6834 | 0.03498 |
| ENSG00000272341 | NA | novel transcript | -1.3311 | 0.03498 |
| ENSG00000102981 | PARD6A | par-6 family cell polarity regulator alpha [Source:HGNC Symbol;Acc:HGNC:15943] | 0.9109 | 0.03514 |
| ENSG00000168071 | CCDC88B | coiled-coil domain containing 88B [Source:HGNC Symbol;Acc:HGNC:26757] | -0.5881 | 0.03518 |
| ENSG00000180287 | PLD5 | phospholipase D family member 5 [Source:HGNC Symbol;Acc:HGNC:26879] | 0.5087 | 0.03539 |
| ENSG00000189367 | KIAA0408 | KIAA0408 [Source:HGNC Symbol;Acc:HGNC:21636] | -0.6731 | 0.03550 |
| ENSG00000174137 | FAM53A | family with sequence similarity 53 member A [Source:HGNC Symbol;Acc:HGNC:31860] | -1.4199 | 0.03569 |
| ENSG00000106948 | AKNA | AT-hook transcription factor [Source:HGNC Symbol;Acc:HGNC:24108] | 0.7033 | 0.03584 |
| ENSG00000160282 | FTCD | formimidoyltransferase cyclodeaminase [Source:HGNC Symbol;Acc:HGNC:3974] | 0.9293 | 0.03589 |
| ENSG00000116661 | FBXO2 | F-box protein 2 [Source:HGNC Symbol;Acc:HGNC:13581] | 0.5911 | 0.03619 |
| ENSG00000185561 | TLCD2 | TLC domain containing 2 [Source:HGNC Symbol;Acc:HGNC:33522] | 0.5150 | 0.03627 |
| ENSG00000183873 | SCN5A | sodium voltage-gated channel alpha subunit 5 [Source:HGNC Symbol;Acc:HGNC:10593] | -0.7255 | 0.03639 |
| ENSG00000198929 | NOS1AP | nitric oxide synthase 1 adaptor protein [Source:HGNC Symbol;Acc:HGNC:16859] | -0.5401 | 0.03666 |
| ENSG00000165124 | SVEP1 | sushi, von Willebrand factor type A, EGF and pentraxin domain containing 1 [Source:HGNC Symbol;Acc:HGNC:15985] | 0.6250 | 0.03667 |
| ENSG00000286478 | NA | novel transcript, antisense to HIBADH | 0.8135 | 0.03680 |
| ENSG00000166828 | SCNN1G | sodium channel epithelial 1 subunit gamma [Source:HGNC Symbol;Acc:HGNC:10602] | -1.8444 | 0.03728 |
| ENSG00000163545 | NUAK2 | NUAK family kinase 2 [Source:HGNC Symbol;Acc:HGNC:29558] | -0.9229 | 0.03749 |
| ENSG00000168546 | GFRA2 | GDNF family receptor alpha 2 [Source:HGNC Symbol;Acc:HGNC:4244] | 0.9827 | 0.03780 |
| ENSG00000272143 | FGF14-AS2 | FGF14 antisense RNA 2 [Source:HGNC Symbol;Acc:HGNC:44368] | -0.6774 | 0.03780 |
| ENSG00000261572 | NA | novel transcript, intronic to CMTM8 | -0.5873 | 0.03809 |
| ENSG00000126010 | GRPR | gastrin releasing peptide receptor [Source:HGNC Symbol;Acc:HGNC:4609] | -1.0087 | 0.03846 |
| ENSG00000279673 | NA | TEC | -0.9301 | 0.03884 |
| ENSG00000197816 | CCDC180 | coiled-coil domain containing 180 [Source:HGNC Symbol;Acc:HGNC:29303] | 0.8239 | 0.03925 |
| ENSG00000119125 | GDA | guanine deaminase [Source:HGNC Symbol;Acc:HGNC:4212] | 1.0326 | 0.03973 |
| ENSG00000177788 | NA | novel transcript | -0.5499 | 0.03985 |
| ENSG00000167912 | NA | novel transcript | 0.9720 | 0.03993 |
| ENSG00000265666 | RARA-AS1 | RARA antisense RNA 1 [Source:HGNC Symbol;Acc:HGNC:49577] | 0.9080 | 0.04032 |
| ENSG00000180011 | ZADH2 | zinc binding alcohol dehydrogenase domain containing 2 [Source:HGNC Symbol;Acc:HGNC:28697] | -0.5079 | 0.04053 |
| ENSG00000167074 | TEF | TEF transcription factor, PAR bZIP family member [Source:HGNC Symbol;Acc:HGNC:11722] | -0.7128 | 0.04072 |
| ENSG00000234719 | NPIPB2 | nuclear pore complex interacting protein family member B2 [Source:HGNC Symbol;Acc:HGNC:37451] | -0.7919 | 0.04082 |
| ENSG00000105641 | SLC5A5 | solute carrier family 5 member 5 [Source:HGNC Symbol;Acc:HGNC:11040] | -1.3118 | 0.04084 |
| ENSG00000117643 | MAN1C1 | mannosidase alpha class 1C member 1 [Source:HGNC Symbol;Acc:HGNC:19080] | -0.6683 | 0.04118 |
| ENSG00000215032 | GNL3LP1 | G protein nucleolar 3 like pseudogene 1 [Source:HGNC Symbol;Acc:HGNC:25733] | -0.9397 | 0.04120 |
| ENSG00000136158 | SPRY2 | sprouty RTK signaling antagonist 2 [Source:HGNC Symbol;Acc:HGNC:11270] | 0.6495 | 0.04223 |
| ENSG00000259248 | USP3-AS1 | USP3 antisense RNA 1 [Source:HGNC Symbol;Acc:HGNC:44140] | -0.8340 | 0.04226 |
| ENSG00000149823 | VPS51 | VPS51 subunit of GARP complex [Source:HGNC Symbol;Acc:HGNC:1172] | 0.5024 | 0.04254 |
| ENSG00000079335 | CDC14A | cell division cycle 14A [Source:HGNC Symbol;Acc:HGNC:1718] | 0.6292 | 0.04261 |
| ENSG00000150630 | VEGFC | vascular endothelial growth factor C [Source:HGNC Symbol;Acc:HGNC:12682] | 0.6359 | 0.04268 |
| ENSG00000137462 | TLR2 | toll like receptor 2 [Source:HGNC Symbol;Acc:HGNC:11848] | 0.7112 | 0.04268 |
| ENSG00000170122 | FOXD4 | forkhead box D4 [Source:HGNC Symbol;Acc:HGNC:3805] | 0.8303 | 0.04268 |

| | | | | |
|---|---|---|---|---|
| ENSG00000110436 | SLC1A2 | solute carrier family 1 member 2 [Source:HGNC Symbol;Acc:HGNC:10940] | 0.8553 | 0.04309 |
| ENSG00000179431 | FJX1 | four-jointed box kinase 1 [Source:HGNC Symbol;Acc:HGNC:17166] | 0.5779 | 0.04309 |
| ENSG00000276740 | NA | novel transcript, sense intronic to FAM155A | 1.0701 | 0.04309 |
| ENSG00000278934 | NA | TEC | -0.6560 | 0.04309 |
| ENSG00000224596 | ZMIZ1-AS1 | ZMIZ1 antisense RNA 1 [Source:HGNC Symbol;Acc:HGNC:27433] | -1.3321 | 0.04312 |
| ENSG00000182901 | RGS7 | regulator of G protein signaling 7 [Source:HGNC Symbol;Acc:HGNC:10003] | 0.6219 | 0.04329 |
| ENSG00000130294 | KIF1A | kinesin family member 1A [Source:HGNC Symbol;Acc:HGNC:888] | 0.6166 | 0.04339 |
| ENSG00000101098 | RIMS4 | regulating synaptic membrane exocytosis 4 [Source:HGNC Symbol;Acc:HGNC:16183] | -1.2801 | 0.04363 |
| ENSG00000105650 | PDE4C | phosphodiesterase 4C [Source:HGNC Symbol;Acc:HGNC:8782] | 1.7010 | 0.04475 |
| ENSG00000100592 | DAAM1 | dishevelled associated activator of morphogenesis 1 [Source:HGNC Symbol;Acc:HGNC:18142] | -0.5284 | 0.04484 |
| ENSG00000180458 | NA | novel transcript, antisense to ZNF793 | -0.6114 | 0.04493 |
| ENSG00000129295 | LRRC6 | leucine rich repeat containing 6 [Source:HGNC Symbol;Acc:HGNC:16725] | 0.7175 | 0.04493 |
| ENSG00000170837 | GPR27 | G protein-coupled receptor 27 [Source:HGNC Symbol;Acc:HGNC:4482] | -0.8466 | 0.04542 |
| ENSG00000150433 | TMEM218 | transmembrane protein 218 [Source:HGNC Symbol;Acc:HGNC:27344] | -0.5726 | 0.04634 |
| ENSG00000057294 | PKP2 | plakophilin 2 [Source:HGNC Symbol;Acc:HGNC:9024] | 0.5135 | 0.04667 |
| ENSG00000109927 | TECTA | tectorin alpha [Source:HGNC Symbol;Acc:HGNC:11720] | -0.5265 | 0.04680 |
| ENSG00000134253 | TRIM45 | tripartite motif containing 45 [Source:HGNC Symbol;Acc:HGNC:19018] | -0.6703 | 0.04696 |
| ENSG00000214827 | MTCP1 | mature T cell proliferation 1 [Source:HGNC Symbol;Acc:HGNC:7423] | -0.5298 | 0.04719 |
| ENSG00000214193 | SH3D21 | SH3 domain containing 21 [Source:HGNC Symbol;Acc:HGNC:26236] | 0.7053 | 0.04740 |
| ENSG00000166407 | LMO1 | LIM domain only 1 [Source:HGNC Symbol;Acc:HGNC:6641] | -0.8776 | 0.04772 |
| ENSG00000140092 | FBLN5 | fibulin 5 [Source:HGNC Symbol;Acc:HGNC:3602] | -1.1591 | 0.04806 |
| ENSG00000049192 | ADAMTS6 | ADAM metallopeptidase with thrombospondin type 1 motif 6 [Source:HGNC Symbol;Acc:HGNC:222] | 0.7264 | 0.04807 |
| ENSG00000153885 | KCTD15 | potassium channel tetramerization domain containing 15 [Source:HGNC Symbol;Acc:HGNC:23297] | 0.8342 | 0.04844 |
| ENSG00000188906 | LRRK2 | leucine rich repeat kinase 2 [Source:HGNC Symbol;Acc:HGNC:18618] | -0.7669 | 0.04844 |
| ENSG00000168952 | STXBP6 | syntaxin binding protein 6 [Source:HGNC Symbol;Acc:HGNC:19666] | -0.5744 | 0.04873 |
| ENSG00000250208 | FZD10-AS1 | FZD10 antisense divergent transcript [Source:HGNC Symbol;Acc:HGNC:48632] | 1.0907 | 0.04873 |
| ENSG00000120915 | EPHX2 | epoxide hydrolase 2 [Source:HGNC Symbol;Acc:HGNC:3402] | -0.6138 | 0.04905 |
| ENSG00000053747 | LAMA3 | laminin subunit alpha 3 [Source:HGNC Symbol;Acc:HGNC:6483] | 0.6628 | 0.04929 |
| ENSG00000161653 | NAGS | N-acetylglutamate synthase [Source:HGNC Symbol;Acc:HGNC:17996] | 0.7326 | 0.04929 |
