## Supplemental Table 6 for "Modulation of calcium signaling on demand to decipher the molecular mechanisms of primary aldosteronism"

**Supplementary Table 6. List of significantly enriched pathways in H295R_S2 cells expressing the _7-5HT3 receptor in response to 8h treatment with 10-8M AngII**

| Pathway | Fold Enrichment | PValue |
|---|---|---|
| hsa04141:Protein processing in endoplasmic reticulum | 2.68 | 7.14E-14 |
| GO:0034976~response to endoplasmic reticulum stress | 2.95 | 4.04E-08 |
| hsa04934:Cushing syndrome | 2.24 | 9.22E-08 |
| GO:0045893~positive regulation of transcription, DNA-templated | 1.52 | 3.55E-07 |
| hsa04925:Aldosterone synthesis and secretion | 2.49 | 8.24E-07 |
| hsa04927:Cortisol synthesis and secretion | 2.85 | 1.67E-06 |
| hsa04928:Parathyroid hormone synthesis, secretion and action | 2.37 | 1.87E-06 |
| hsa00513:Various types of N-glycan biosynthesis | 3.35 | 2.79E-06 |
| GO:0030308~negative regulation of cell growth | 2.28 | 3.24E-06 |
| hsa04010:MAPK signaling pathway | 1.72 | 4.70E-06 |
| GO:0045944~positive regulation of transcription from RNA polymerase II promoter | 1.35 | 4.73E-06 |
| GO:0032922~circadian regulation of gene expression | 2.70 | 1.00E-05 |
| GO:0006986~response to unfolded protein | 2.89 | 1.32E-05 |
| GO:0034599~cellular response to oxidative stress | 2.40 | 1.74E-05 |
| hsa05130:Pathogenic Escherichia coli infection | 1.84 | 2.33E-05 |
| hsa00510:N-Glycan biosynthesis | 2.79 | 3.25E-05 |
| GO:0007399~nervous system development | 1.57 | 3.64E-05 |
| GO:0007165~signal transduction | 1.30 | 3.76E-05 |
| hsa04068:FoxO signaling pathway | 2.03 | 3.93E-05 |
| GO:0009410~response to xenobiotic stimulus | 1.74 | 5.21E-05 |
| GO:0043065~positive regulation of apoptotic process | 1.64 | 5.21E-05 |
| GO:0019082~viral protein processing | 3.49 | 6.14E-05 |
| hsa04926:Relaxin signaling pathway | 2.01 | 6.80E-05 |
| GO:0001649~osteoblast differentiation | 2.08 | 6.98E-05 |
| GO:0034975~protein folding in endoplasmic reticulum | 5.81 | 9.94E-05 |
| hsa04151:PI3K-Akt signaling pathway | 1.55 | 9.96E-05 |
| GO:0007507~heart development | 1.77 | 1.20E-04 |
| GO:0042542~response to hydrogen peroxide | 2.97 | 1.35E-04 |
| GO:0034097~response to cytokine | 2.83 | 1.50E-04 |
| GO:0006468~protein phosphorylation | 1.45 | 1.68E-04 |
| GO:0006886~intracellular protein transport | 1.59 | 1.70E-04 |
| GO:0007264~small GTPase mediated signal transduction | 2.03 | 1.85E-04 |
| hsa04210:Apoptosis | 1.90 | 2.08E-04 |
| hsa05132:Salmonella infection | 1.63 | 2.16E-04 |
| GO:0060070~canonical Wnt signaling pathway | 2.11 | 2.64E-04 |
| hsa04261:Adrenergic signaling in cardiomyocytes | 1.83 | 2.64E-04 |
| GO:0000079~regulation of cyclin-dependent protein serine/threonine kinase activity | 2.61 | 2.66E-04 |
| GO:0016477~cell migration | 1.63 | 2.68E-04 |
| GO:0001843~neural tube closure | 2.27 | 2.92E-04 |
| GO:0015031~protein transport | 1.48 | 3.19E-04 |
| GO:0035556~intracellular signal transduction | 1.46 | 3.31E-04 |
| hsa04931:Insulin resistance | 1.99 | 3.90E-04 |
| GO:0006749~glutathione metabolic process | 2.66 | 5.28E-04 |
| GO:0043547~positive regulation of GTPase activity | 1.73 | 5.48E-04 |
| GO:0006915~apoptotic process | 1.37 | 5.60E-04 |
| hsa04728:Dopaminergic synapse | 1.85 | 5.68E-04 |
| hsa05131:Shigellosis | 1.59 | 5.93E-04 |
| GO:0007411~axon guidance | 1.72 | 6.12E-04 |
| GO:0006487~protein N-linked glycosylation | 2.51 | 6.75E-04 |
| GO:0030433~ubiquitin-dependent ERAD pathway | 2.17 | 7.79E-04 |
| GO:0008585~female gonad development | 3.63 | 7.97E-04 |
| GO:0008360~regulation of cell shape | 1.81 | 7.99E-04 |
| hsa04510:Focal adhesion | 1.64 | 8.15E-04 |
| GO:0042981~regulation of apoptotic process | 1.61 | 8.43E-04 |
| hsa05215:Prostate cancer | 1.98 | 8.60E-04 |

| | | |
|---|---|---|
| GO:0030522~intracellular receptor signaling pathway | 3.09 | 8.86E-04 |
| GO:0007265~Ras protein signal transduction | 2.06 | 9.14E-04 |
| GO:0043066~negative regulation of apoptotic process | 1.39 | 9.36E-04 |
| hsa04668:TNF signaling pathway | 1.88 | 9.86E-04 |
| hsa05205:Proteoglycans in cancer | 1.62 | 0.0010 |
| h_p53hypoxiaPathway:Hypoxia and p53 in the Cardiovascular system | 2.70 | 0.0010 |
| hsa04360:Axon guidance | 1.67 | 0.0010 |
| GO:0030036~actin cytoskeleton organization | 1.69 | 0.0011 |
| hsa04935:Growth hormone synthesis, secretion and action | 1.85 | 0.0011 |
| GO:0000902~cell morphogenesis | 2.04 | 0.0011 |
| hsa04710:Circadian rhythm | 2.83 | 0.0011 |
| GO:0071560~cellular response to transforming growth factor beta stimulus | 2.26 | 0.0012 |
| GO:0045047~protein targeting to ER | 4.26 | 0.0012 |
| GO:1904382~mannose trimming involved in glycoprotein ERAD pathway | 5.99 | 0.0012 |
| GO:0071474~cellular hyperosmotic response | 5.99 | 0.0012 |
| GO:0007015~actin filament organization | 1.76 | 0.0013 |
| GO:0070507~regulation of microtubule cytoskeleton organization | 2.81 | 0.0013 |
| GO:0060412~ventricular septum morphogenesis | 2.81 | 0.0013 |
| GO:0007179~transforming growth factor beta receptor signaling pathway | 1.98 | 0.0013 |
| hsa05169:Epstein-Barr virus infection | 1.61 | 0.0014 |
| hsa05418:Fluid shear stress and atherosclerosis | 1.76 | 0.0014 |
| hsa05225:Hepatocellular carcinoma | 1.67 | 0.0015 |
| GO:0000122~negative regulation of transcription from RNA polymerase II promoter | 1.26 | 0.0015 |
| GO:0000226~microtubule cytoskeleton organization | 1.79 | 0.0016 |
| hsa04015:Rap1 signaling pathway | 1.59 | 0.0017 |
| GO:0006983~ER overload response | 4.66 | 0.0018 |
| hsa05202:Transcriptional misregulation in cancer | 1.61 | 0.0018 |
| hsa04725:Cholinergic synapse | 1.83 | 0.0019 |
| GO:0030501~positive regulation of bone mineralization | 2.54 | 0.0022 |
| hsa04146:Peroxisome | 1.99 | 0.0024 |
| GO:0001525~angiogenesis | 1.53 | 0.0024 |
| GO:0060316~positive regulation of ryanodine-sensitive calcium-release channel | 5.32 | 0.0025 |
| GO:1902570~protein localization to nucleolus | 5.32 | 0.0025 |
| hsa04915:Estrogen signaling pathway | 1.72 | 0.0026 |
| hsa04919:Thyroid hormone signaling pathway | 1.77 | 0.0026 |
| GO:0001822~kidney development | 1.83 | 0.0027 |
| hsa05200:Pathways in cancer | 1.32 | 0.0027 |
| GO:2001046~positive regulation of integrin-mediated signaling pathway | 4.30 | 0.0030 |
| GO:0030838~positive regulation of actin filament polymerization | 2.35 | 0.0032 |
| hsa03250:Viral life cycle - HIV-1 | 2.11 | 0.0033 |
| GO:0050772~positive regulation of axonogenesis | 2.83 | 0.0034 |
| GO:0001933~negative regulation of protein phosphorylation | 2.02 | 0.0034 |
| GO:0001701~in utero embryonic development | 1.57 | 0.0034 |
| hsa04014:Ras signaling pathway | 1.50 | 0.0035 |
| GO:0030010~establishment of cell polarity | 2.66 | 0.0035 |
| GO:0034260~negative regulation of GTPase activity | 2.66 | 0.0035 |
| GO:0070059~intrinsic apoptotic signaling pathway in response to endoplasmic reticulum stress | 2.66 | 0.0035 |
| GO:0001657~ureteric bud development | 2.66 | 0.0035 |
| GO:0000278~mitotic cell cycle | 1.69 | 0.0035 |
| GO:0006470~protein dephosphorylation | 1.69 | 0.0035 |
| GO:0009749~response to glucose | 2.11 | 0.0036 |
| GO:0007030~Golgi organization | 1.74 | 0.0037 |
| GO:0008285~negative regulation of cell proliferation | 1.36 | 0.0039 |
| GO:0016192~vesicle-mediated transport | 1.53 | 0.0039 |
| GO:0001570~vasculogenesis | 2.15 | 0.0039 |
| GO:0007005~mitochondrion organization | 1.95 | 0.0041 |

| | | |
|---|---|---|
| GO:0030335~positive regulation of cell migration | 1.49 | 0.0042 |
| GO:0042149~cellular response to glucose starvation | 2.20 | 0.0043 |
| GO:0017157~regulation of exocytosis | 2.59 | 0.0044 |
| GO:0009267~cellular response to starvation | 2.02 | 0.0044 |
| GO:0042752~regulation of circadian rhythm | 2.12 | 0.0046 |
| hsa05417:Lipid and atherosclerosis | 1.51 | 0.0047 |
| hsa04066:HIF-1 signaling pathway | 1.77 | 0.0048 |
| GO:0035024~negative regulation of Rho protein signal transduction | 3.12 | 0.0051 |
| hsa04520:Adherens junction | 1.83 | 0.0054 |
| GO:0006486~protein glycosylation | 1.76 | 0.0054 |
| GO:0072583~clathrin-dependent endocytosis | 2.85 | 0.0055 |
| GO:0098609~cell-cell adhesion | 1.56 | 0.0062 |
| GO:0036499~PERK-mediated unfolded protein response | 5.70 | 0.0063 |
| GO:0019852~L-ascorbic acid metabolic process | 5.70 | 0.0063 |
| GO:0060021~palate development | 2.06 | 0.0064 |
| hsa04933:AGE-RAGE signaling pathway in diabetic complications | 1.78 | 0.0064 |
| hsa04140:Autophagy - animal | 1.63 | 0.0068 |
| GO:1903826~arginine transmembrane transport | 3.73 | 0.0070 |
| GO:0035924~cellular response to vascular endothelial growth factor stimulus | 2.58 | 0.0071 |
| GO:0035910~ascending aorta morphogenesis | 7.99 | 0.0071 |
| GO:0032057~negative regulation of translational initiation in response to stress | 7.99 | 0.0071 |
| GO:0045892~negative regulation of transcription, DNA-templated | 1.29 | 0.0073 |
| GO:0019896~axonal transport of mitochondrion | 4.36 | 0.0074 |
| hsa04922:Glucagon signaling pathway | 1.73 | 0.0076 |
| GO:0002062~chondrocyte differentiation | 2.14 | 0.0078 |
| GO:0030154~cell differentiation | 1.26 | 0.0080 |
| hsa03060:Protein export | 2.90 | 0.0081 |
| GO:0006366~transcription from RNA polymerase II promoter | 1.48 | 0.0082 |
| GO:0001764~neuron migration | 1.70 | 0.0084 |
| hsa04022:cGMP-PKG signaling pathway | 1.55 | 0.0084 |
| GO:0001837~epithelial to mesenchymal transition | 2.19 | 0.0086 |
| GO:0048010~vascular endothelial growth factor receptor signaling pathway | 2.66 | 0.0091 |
| GO:0031397~negative regulation of protein ubiquitination | 2.10 | 0.0092 |
| hsa05161:Hepatitis B | 1.55 | 0.0093 |
| GO:0001666~response to hypoxia | 1.59 | 0.0093 |
| GO:0010976~positive regulation of neuron projection development | 1.71 | 0.0096 |
| GO:0086010~membrane depolarization during action potential | 3.49 | 0.0100 |
| GO:0030970~retrograde protein transport, ER to cytosol | 3.49 | 0.0100 |
| GO:0030177~positive regulation of Wnt signaling pathway | 2.34 | 0.0102 |
| GO:0016055~Wnt signaling pathway | 1.52 | 0.0105 |
| GO:0048675~axon extension | 2.44 | 0.0109 |
| GO:0070588~calcium ion transmembrane transport | 1.71 | 0.0109 |
| hsa05212:Pancreatic cancer | 1.85 | 0.0110 |
| hsa05030:Cocaine addiction | 2.11 | 0.0111 |
| GO:0030837~negative regulation of actin filament polymerization | 3.04 | 0.0113 |
| GO:0015804~neutral amino acid transport | 3.04 | 0.0113 |
| GO:0046620~regulation of organ growth | 4.99 | 0.0113 |
| GO:1903401~L-lysine transmembrane transport | 4.99 | 0.0113 |
| GO:0071493~cellular response to UV-B | 3.99 | 0.0113 |
| GO:0031098~stress-activated protein kinase signaling cascade | 3.99 | 0.0113 |
| GO:0043117~positive regulation of vascular permeability | 3.99 | 0.0113 |
| GO:0046697~decidualization | 2.76 | 0.0116 |
| GO:0007155~cell adhesion | 1.28 | 0.0117 |
| GO:0051056~regulation of small GTPase mediated signal transduction | 1.68 | 0.0118 |
| GO:1902895~positive regulation of pri-miRNA transcription from RNA polymerase II promoter | 2.11 | 0.0120 |
| GO:0006874~cellular calcium ion homeostasis | 1.70 | 0.0121 |
| GO:0006865~amino acid transport | 2.28 | 0.0122 |

| | | |
|---|---|---|
| GO:0043406~positive regulation of MAP kinase activity | 1.87 | 0.0132 |
| hsa05145:Toxoplasmosis | 1.65 | 0.0133 |
| GO:0018105~peptidyl-serine phosphorylation | 1.51 | 0.0135 |
| hsa04211:Longevity regulating pathway | 1.75 | 0.0137 |
| hsa04727:GABAergic synapse | 1.75 | 0.0137 |
| GO:0008654~phospholipid biosynthetic process | 2.50 | 0.0141 |
| GO:0048008~platelet-derived growth factor receptor signaling pathway | 2.50 | 0.0141 |
| GO:0046777~protein autophosphorylation | 1.52 | 0.0141 |
| GO:0009409~response to cold | 2.23 | 0.0146 |
| GO:0043409~negative regulation of MAPK cascade | 2.23 | 0.0146 |
| GO:0045666~positive regulation of neuron differentiation | 1.77 | 0.0147 |
| hsa04340:Hedgehog signaling pathway | 1.98 | 0.0147 |
| GO:0006479~protein methylation | 2.66 | 0.0147 |
| GO:0072332~intrinsic apoptotic signaling pathway by p53 class mediator | 2.66 | 0.0147 |
| GO:0050679~positive regulation of epithelial cell proliferation | 1.89 | 0.0150 |
| hsa04910:Insulin signaling pathway | 1.57 | 0.0150 |
| GO:0045578~negative regulation of B cell differentiation | 6.39 | 0.0161 |
| GO:1903912~negative regulation of endoplasmic reticulum stress-induced eIF2 alpha phosphorylation | 6.39 | 0.0161 |
| GO:0031670~cellular response to nutrient | 6.39 | 0.0161 |
| GO:0048167~regulation of synaptic plasticity | 2.03 | 0.0163 |
| GO:0090398~cellular senescence | 2.03 | 0.0163 |
| GO:0032355~response to estradiol | 1.76 | 0.0164 |
| hsa04930:Type II diabetes mellitus | 2.09 | 0.0165 |
| GO:2000279~negative regulation of DNA biosynthetic process | 3.69 | 0.0165 |
| GO:0040008~regulation of growth | 1.96 | 0.0167 |
| GO:0032570~response to progesterone | 2.42 | 0.0173 |
| GO:0003281~ventricular septum development | 2.42 | 0.0173 |
| GO:0035633~maintenance of permeability of blood-brain barrier | 2.42 | 0.0173 |
| hsa04390:Hippo signaling pathway | 1.51 | 0.0176 |
| hsa04371:Apelin signaling pathway | 1.54 | 0.0181 |
| GO:0034614~cellular response to reactive oxygen species | 2.08 | 0.0183 |
| GO:0008354~germ cell migration | 4.44 | 0.0183 |
| GO:0071243~cellular response to arsenic-containing substance | 4.44 | 0.0183 |
| GO:0060231~mesenchymal to epithelial transition | 4.44 | 0.0183 |
| GO:0003143~embryonic heart tube morphogenesis | 4.44 | 0.0183 |
| GO:0031204~posttranslational protein targeting to membrane, translocation | 4.44 | 0.0183 |
| GO:1902949~positive regulation of tau-protein kinase activity | 4.44 | 0.0183 |
| GO:0006198~cAMP catabolic process | 4.44 | 0.0183 |
| GO:0046755~viral budding | 4.44 | 0.0183 |
| GO:0030307~positive regulation of cell growth | 1.74 | 0.0184 |
| GO:0006259~DNA metabolic process | 2.57 | 0.0184 |
| hsa00561:Glycerolipid metabolism | 1.88 | 0.0184 |
| GO:0055088~lipid homeostasis | 2.00 | 0.0189 |
| GO:0060307~regulation of ventricular cardiac muscle cell membrane repolarization | 2.78 | 0.0190 |
| GO:2000378~negative regulation of reactive oxygen species metabolic process | 2.78 | 0.0190 |
| GO:0002244~hematopoietic progenitor cell differentiation | 1.80 | 0.0190 |
| hsa04911:Insulin secretion | 1.72 | 0.0190 |
| GO:0050919~negative chemotaxis | 2.25 | 0.0192 |
| GO:0009306~protein secretion | 1.93 | 0.0192 |
| hsa05165:Human papillomavirus infection | 1.32 | 0.0204 |
| GO:0045747~positive regulation of Notch signaling pathway | 2.13 | 0.0205 |
| GO:0045648~positive regulation of erythrocyte differentiation | 2.35 | 0.0210 |
| GO:0030968~endoplasmic reticulum unfolded protein response | 2.04 | 0.0213 |
| GO:0007173~epidermal growth factor receptor signaling pathway | 2.04 | 0.0213 |
| GO:0042552~myelination | 2.04 | 0.0213 |

| | | |
|---|---|---|
| GO:0000462~maturation of SSU-rRNA from tricistronic rRNA transcript (SSU-rRNA, 5.8S rRNA, LSU-rRNA) | 2.48 | 0.0227 |
| GO:0008625~extrinsic apoptotic signaling pathway via death domain receptors | 2.20 | 0.0228 |
| GO:0035094~response to nicotine | 2.20 | 0.0228 |
| GO:0050767~regulation of neurogenesis | 2.20 | 0.0228 |
| GO:0001974~blood vessel remodeling | 2.20 | 0.0228 |
| GO:0060996~dendritic spine development | 3.42 | 0.0231 |
| GO:0043278~response to morphine | 2.66 | 0.0240 |
| GO:0051571~positive regulation of histone H3-K4 methylation | 2.66 | 0.0240 |
| GO:0018279~protein N-linked glycosylation via asparagine | 2.66 | 0.0240 |
| GO:0006418~tRNA aminoacylation for protein translation | 2.66 | 0.0240 |
| GO:0007420~brain development | 1.38 | 0.0240 |
| GO:0071260~cellular response to mechanical stimulus | 1.79 | 0.0244 |
| GO:1902236~negative regulation of endoplasmic reticulum stress-induced intrinsic apoptotic signaling pathway | 2.94 | 0.0244 |
| GO:0006414~translational elongation | 2.94 | 0.0244 |
| GO:0042326~negative regulation of phosphorylation | 2.94 | 0.0244 |
| GO:0006491~N-glycan processing | 2.94 | 0.0244 |
| GO:0097191~extrinsic apoptotic signaling pathway | 2.00 | 0.0246 |
| GO:0048666~neuron development | 1.93 | 0.0249 |
| hsa05170:Human immunodeficiency virus 1 infection | 1.40 | 0.0257 |
| hsa05100:Bacterial invasion of epithelial cells | 1.73 | 0.0258 |
| GO:0071456~cellular response to hypoxia | 1.55 | 0.0260 |
| hsa05163:Human cytomegalovirus infection | 1.38 | 0.0261 |
| GO:0019216~regulation of lipid metabolic process | 2.14 | 0.0269 |
| GO:0021987~cerebral cortex development | 1.76 | 0.0273 |
| GO:0015820~leucine transport | 3.99 | 0.0276 |
| GO:0060840~artery development | 3.99 | 0.0276 |
| GO:0010907~positive regulation of glucose metabolic process | 3.99 | 0.0276 |
| GO:0060371~regulation of atrial cardiac muscle cell membrane depolarization | 3.99 | 0.0276 |
| GO:0032836~glomerular basement membrane development | 3.99 | 0.0276 |
| GO:0043085~positive regulation of catalytic activity | 2.40 | 0.0276 |
| GO:0060271~cilium assembly | 1.39 | 0.0277 |
| GO:0030509~BMP signaling pathway | 1.66 | 0.0278 |
| GO:0048538~thymus development | 2.04 | 0.0278 |
| GO:1901224~positive regulation of NIK/NF-kappaB signaling | 1.80 | 0.0279 |
| hsa05223:Non-small cell lung cancer | 1.75 | 0.0284 |
| GO:0002931~response to ischemia | 1.89 | 0.0284 |
| h_stressPathway:TNF/Stress Related Signaling | 2.16 | 0.0285 |
| GO:1903575~cornified envelope assembly | 5.32 | 0.0292 |
| GO:0097398~cellular response to interleukin-17 | 5.32 | 0.0292 |
| GO:0046985~positive regulation of hemoglobin biosynthetic process | 5.32 | 0.0292 |
| GO:0048713~regulation of oligodendrocyte differentiation | 5.32 | 0.0292 |
| GO:1902747~negative regulation of lens fiber cell differentiation | 5.32 | 0.0292 |
| GO:0014031~mesenchymal cell development | 5.32 | 0.0292 |
| GO:0098885~modification of postsynaptic actin cytoskeleton | 5.32 | 0.0292 |
| GO:0051891~positive regulation of cardioblast differentiation | 5.32 | 0.0292 |
| GO:0040020~regulation of meiotic nuclear division | 5.32 | 0.0292 |
| GO:1903898~negative regulation of PERK-mediated unfolded protein response | 5.32 | 0.0292 |
| GO:0072659~protein localization to plasma membrane | 1.48 | 0.0293 |
| GO:0060038~cardiac muscle cell proliferation | 2.56 | 0.0298 |
| GO:0045216~cell-cell junction organization | 2.56 | 0.0298 |
| hsa05162:Measles | 1.49 | 0.0309 |
| hsa05110:Vibrio cholerae infection | 1.92 | 0.0311 |
| GO:0051301~cell division | 1.29 | 0.0312 |
| GO:0000470~maturation of LSU-rRNA | 3.19 | 0.0312 |
| GO:0030220~platelet formation | 2.80 | 0.0313 |
| GO:0060122~inner ear receptor stereocilium organization | 2.80 | 0.0313 |

| | | |
|---|---|---|
| GO:0086091~regulation of heart rate by cardiac conduction | 2.09 | 0.0314 |
| GO:0032870~cellular response to hormone stimulus | 2.09 | 0.0314 |
| GO:0043087~regulation of GTPase activity | 1.67 | 0.0318 |
| hsa01524:Platinum drug resistance | 1.72 | 0.0319 |
| GO:0001568~blood vessel development | 2.00 | 0.0322 |
| hsa04152:AMPK signaling pathway | 1.53 | 0.0322 |
| GO:2000045~regulation of G1/S transition of mitotic cell cycle | 1.92 | 0.0324 |
| hsa04012:ErbB signaling pathway | 1.65 | 0.0328 |
| hsa05032:Morphine addiction | 1.63 | 0.0330 |
| hsa04713:Circadian entrainment | 1.60 | 0.0330 |
| GO:0000165~MAPK cascade | 1.48 | 0.0332 |
| GO:0006457~protein folding | 1.43 | 0.0332 |
| GO:0007026~negative regulation of microtubule depolymerization | 2.32 | 0.0333 |
| GO:0070536~protein K63-linked deubiquitination | 2.32 | 0.0333 |
| h_telPathway:Telomeres, Telomerase, Cellular Aging, and Immortality | 2.40 | 0.0341 |
| GO:0051603~proteolysis involved in cellular protein catabolic process | 2.16 | 0.0354 |
| GO:0051017~actin filament bundle assembly | 2.16 | 0.0354 |
| GO:0006654~phosphatidic acid biosynthetic process | 2.16 | 0.0354 |
| GO:0048536~spleen development | 2.16 | 0.0354 |
| GO:0071466~cellular response to xenobiotic stimulus | 1.79 | 0.0358 |
| GO:0043123~positive regulation of I-kappaB kinase/NF-kappaB signaling | 1.40 | 0.0362 |
| hsa05210:Colorectal cancer | 1.63 | 0.0365 |
| GO:0090103~cochlea morphogenesis | 2.46 | 0.0365 |
| GO:0042098~T cell proliferation | 2.04 | 0.0365 |
| GO:0042307~positive regulation of protein import into nucleus | 2.04 | 0.0365 |
| GO:0032728~positive regulation of interferon-beta production | 2.04 | 0.0365 |
| GO:0007568~aging | 1.49 | 0.0368 |
| GO:0001934~positive regulation of protein phosphorylation | 1.39 | 0.0368 |
| GO:0006094~gluconeogenesis | 1.96 | 0.0370 |
| GO:0050821~protein stabilization | 1.37 | 0.0372 |
| GO:0010628~positive regulation of gene expression | 1.23 | 0.0378 |
| hsa05217:Basal cell carcinoma | 1.76 | 0.0387 |
| GO:0006613~cotranslational protein targeting to membrane | 3.63 | 0.0391 |
| GO:0006002~fructose 6-phosphate metabolic process | 3.63 | 0.0391 |
| GO:0045048~protein insertion into ER membrane | 3.63 | 0.0391 |
| GO:2001020~regulation of response to DNA damage stimulus | 3.63 | 0.0391 |
| GO:0006646~phosphatidylethanolamine biosynthetic process | 3.63 | 0.0391 |
| GO:0070231~T cell apoptotic process | 3.63 | 0.0391 |
| GO:0016081~synaptic vesicle docking | 3.63 | 0.0391 |
| GO:0033554~cellular response to stress | 2.66 | 0.0393 |
| GO:0006977~DNA damage response, signal transduction by p53 class mediator resulting in cell cycle arrest | 2.66 | 0.0393 |
| GO:0045722~positive regulation of gluconeogenesis | 2.66 | 0.0393 |
| hsa04024:cAMP signaling pathway | 1.35 | 0.0393 |
| hsa04920:Adipocytokine signaling pathway | 1.72 | 0.0395 |
| GO:0060325~face morphogenesis | 2.25 | 0.0396 |
| hsa04918:Thyroid hormone synthesis | 1.68 | 0.0401 |
| hsa00564:Glycerophospholipid metabolism | 1.57 | 0.0401 |
| hsa04912:GnRH signaling pathway | 1.59 | 0.0403 |
| GO:0007517~muscle organ development | 1.60 | 0.0405 |
| GO:0090263~positive regulation of canonical Wnt signaling pathway | 1.50 | 0.0407 |
| GO:0002021~response to dietary excess | 2.99 | 0.0408 |
| GO:1990869~cellular response to chemokine | 2.99 | 0.0408 |
| GO:0050848~regulation of calcium-mediated signaling | 2.99 | 0.0408 |
| GO:0032516~positive regulation of phosphoprotein phosphatase activity | 2.99 | 0.0408 |
| GO:0010634~positive regulation of epithelial cell migration | 2.10 | 0.0414 |
| GO:0098703~calcium ion import across plasma membrane | 2.10 | 0.0414 |
| GO:0060071~Wnt signaling pathway, planar cell polarity pathway | 1.85 | 0.0418 |

| | | |
|---|---|---|
| GO:0045600~positive regulation of fat cell differentiation | 1.85 | 0.0418 |
| GO:0048863~stem cell differentiation | 2.00 | 0.0421 |
| GO:0071364~cellular response to epidermal growth factor stimulus | 2.00 | 0.0421 |
| GO:0050714~positive regulation of protein secretion | 1.92 | 0.0422 |
| GO:0036508~protein alpha-1,2-demannosylation | 7.99 | 0.0431 |
| GO:0032342~aldosterone biosynthetic process | 7.99 | 0.0431 |
| GO:0033693~neurofilament bundle assembly | 7.99 | 0.0431 |
| GO:0010638~positive regulation of organelle organization | 7.99 | 0.0431 |
| GO:1901534~positive regulation of hematopoietic progenitor cell differentiation | 7.99 | 0.0431 |
| GO:1904381~Golgi apparatus mannose trimming | 7.99 | 0.0431 |
| GO:0072102~glomerulus morphogenesis | 7.99 | 0.0431 |
| GO:0034651~cortisol biosynthetic process | 7.99 | 0.0431 |
| GO:0070859~positive regulation of bile acid biosynthetic process | 7.99 | 0.0431 |
| GO:0050893~sensory processing | 7.99 | 0.0431 |
| GO:1990737~response to manganese-induced endoplasmic reticulum stress | 7.99 | 0.0431 |
| h_PparaPathway:Mechanism of Gene Regulation by Peroxisome Proliferators via PPARa(alpha) | 1.66 | 0.0431 |
| GO:0032868~response to insulin | 1.70 | 0.0433 |
| GO:0001938~positive regulation of endothelial cell proliferation | 1.70 | 0.0433 |
| hsa05230:Central carbon metabolism in cancer | 1.69 | 0.0443 |
| GO:0032869~cellular response to insulin stimulus | 1.58 | 0.0443 |
| GO:0021766~hippocampus development | 1.74 | 0.0447 |
| GO:0006469~negative regulation of protein kinase activity | 1.74 | 0.0447 |
| hsa00514:Other types of O-glycan biosynthesis | 1.89 | 0.0450 |
| GO:0051216~cartilage development | 1.77 | 0.0460 |
| GO:0060613~fat pad development | 4.56 | 0.0464 |
| GO:0036089~cleavage furrow formation | 4.56 | 0.0464 |
| GO:1900076~regulation of cellular response to insulin stimulus | 4.56 | 0.0464 |
| GO:0032364~oxygen homeostasis | 4.56 | 0.0464 |
| GO:0051239~regulation of multicellular organismal process | 4.56 | 0.0464 |
| GO:0003406~retinal pigment epithelium development | 4.56 | 0.0464 |
| GO:0061669~spontaneous neurotransmitter secretion | 4.56 | 0.0464 |
| GO:0046321~positive regulation of fatty acid oxidation | 4.56 | 0.0464 |
| GO:0007435~salivary gland morphogenesis | 4.56 | 0.0464 |
| GO:0035356~cellular triglyceride homeostasis | 4.56 | 0.0464 |
| GO:2000210~positive regulation of anoikis | 4.56 | 0.0464 |
| GO:0006914~autophagy | 1.44 | 0.0464 |
| hsa04961:Endocrine and other factor-regulated calcium reabsorption | 1.81 | 0.0468 |
| GO:2000300~regulation of synaptic vesicle exocytosis | 1.82 | 0.0472 |
| GO:0009612~response to mechanical stimulus | 1.82 | 0.0472 |
| GO:0006888~ER to Golgi vesicle-mediated transport | 1.48 | 0.0473 |
| 99.NF-kB_activation | 2.42 | 0.0476 |
| GO:0030199~collagen fibril organization | 1.68 | 0.0479 |
| GO:0009636~response to toxic substance | 1.68 | 0.0479 |
| GO:0003151~outflow tract morphogenesis | 1.88 | 0.0480 |
| GO:0060173~limb development | 2.05 | 0.0480 |
| GO:0033077~T cell differentiation in thymus | 2.05 | 0.0480 |
| hsa04916:Melanogenesis | 1.54 | 0.0482 |
| GO:0055075~potassium ion homeostasis | 2.54 | 0.0485 |
| GO:1901214~regulation of neuron death | 2.54 | 0.0485 |
| GO:0030517~negative regulation of axon extension | 2.54 | 0.0485 |
| GO:0051726~regulation of cell cycle | 1.32 | 0.0490 |
| GO:0008284~positive regulation of cell proliferation | 1.21 | 0.0491 |
| hsa04512:ECM-receptor interaction | 1.58 | 0.0492 |
| hsa05412:Arrhythmogenic right ventricular cardiomyopathy | 1.63 | 0.0496 |
