## Supplemental Table 7 for "Modulation of calcium signaling on demand to decipher the molecular mechanisms of primary aldosteronism"

**Supplementary Table 7. List of significantly enriched pathways in H295R_S2 cells expressing the _7-5HT3 receptor in response to 8h treatment with 12mM K+**

| Pathway | Fold Enrichment | PValue |
|---|---|---|
| GO:0009410~response to xenobiotic stimulus | 3,63 | 3,15E-08 |
| GO:0035556~intracellular signal transduction | 2,52 | 2,24E-06 |
| GO:0007411~axon guidance | 3,41 | 1,29E-05 |
| GO:0001764~neuron migration | 4,19 | 1,38E-05 |
| GO:0001822~kidney development | 4,04 | 4,29E-05 |
| GO:0086010~membrane depolarization during action potential | 12,77 | 7,10E-05 |
| GO:0001933~negative regulation of protein phosphorylation | 4,74 | 1,01E-04 |
| GO:0098703~calcium ion import across plasma membrane | 7,17 | 1,03E-04 |
| GO:0045944~positive regulation of transcription from RNA polymerase II promoter | 1,67 | 1,11E-04 |
| GO:0051770~positive regulation of nitric-oxide synthase biosynthetic process | 11,35 | 1,33E-04 |
| GO:0007507~heart development | 2,89 | 1,79E-04 |
| hsa04710:Circadian rhythm | 6,28 | 2,13E-04 |
| GO:0006814~sodium ion transport | 4,21 | 2,75E-04 |
| GO:0050731~positive regulation of peptidyl-tyrosine phosphorylation | 4,12 | 3,30E-04 |
| hsa04010:MAPK signaling pathway | 2,21 | 3,75E-04 |
| GO:0030336~negative regulation of cell migration | 2.95 | 3,76E-04 |
| GO:0045893~positive regulation of transcription, DNA-templated | 1.83 | 4,08E-04 |
| GO:0043406~positive regulation of MAP kinase activity | 4.42 | 4,10E-04 |
| GO:0086064~cell communication by electrical coupling involved in cardiac conduction | 13.10 | 4,23E-04 |
| hsa04727:GABAergic synapse | 3.60 | 4,46E-04 |
| GO:1902747~negative regulation of lens fiber cell differentiation | 22.71 | 4,69E-04 |
| GO:0035725~sodium ion transmembrane transport | 3.62 | 4,87E-04 |
| GO:1904322~cellular response to forskolin | 12.16 | 5,78E-04 |
| GO:0071456~cellular response to hypoxia | 3.30 | 5,87E-04 |
| GO:0060412~ventricular septum morphogenesis | 6.44 | 6,62E-04 |
| GO:0030168~platelet activation | 4.51 | 8,16E-04 |
| GO:0050919~negative chemotaxis | 6.11 | 8,84E-04 |
| GO:0086002~cardiac muscle cell action potential involved in contraction | 10.64 | 0.0010 |
| GO:0010719~negative regulation of epithelial to mesenchymal transition | 5.96 | 0.0010 |
| GO:0009611~response to wounding | 4.32 | 0.0011 |
| GO:0007165~signal transduction | 1.54 | 0.0011 |
| GO:0030154~cell differentiation | 1.78 | 0.0012 |
| hsa04360:Axon guidance | 2.49 | 0.0012 |
| GO:0007155~cell adhesion | 1.88 | 0.0013 |
| GO:0032870~cellular response to hormone stimulus | 5.68 | 0.0013 |
| GO:0060048~cardiac muscle contraction | 5.68 | 0.0013 |
| hsa04022:cGMP-PKG signaling pathway | 2.56 | 0.0014 |
| GO:0043409~negative regulation of MAPK cascade | 5.54 | 0.0015 |
| GO:0006198~cAMP catabolic process | 15.14 | 0.0018 |
| GO:0045664~regulation of neuron differentiation | 6.59 | 0.0019 |
| hsa05032:Morphine addiction | 3.23 | 0.0020 |
| hsa04015:Rap1 signaling pathway | 2.29 | 0.0022 |
| GO:0006468~protein phosphorylation | 1.84 | 0.0024 |
| GO:1990646~cellular response to prolactin | 34.06 | 0.0025 |
| GO:0070373~negative regulation of ERK1 and ERK2 cascade | 3.69 | 0.0030 |
| GO:1990830~cellular response to leukemia inhibitory factor | 3.31 | 0.0033 |
| GO:0050714~positive regulation of protein secretion | 4.77 | 0.0033 |
| hsa04151:PI3K-Akt signaling pathway | 1.89 | 0.0033 |
| GO:0019228~neuronal action potential | 5.84 | 0.0033 |
| hsa04925:Aldosterone synthesis and secretion | 3.00 | 0.0035 |
| GO:0042632~cholesterol homeostasis | 3.27 | 0.0035 |
| GO:0038083~peptidyl-tyrosine autophosphorylation | 7.74 | 0.0035 |
| GO:0042476~odontogenesis | 5.68 | 0.0038 |
| GO:0030036~actin cytoskeleton organization | 2.52 | 0.0039 |
| GO:0060307~regulation of ventricular cardiac muscle cell membrane repolarization | 7.40 | 0.0041 |

| | | |
|---|---|---|
| GO:0007417~central nervous system development | 2.76 | 0.0043 |
| GO:0018108~peptidyl-tyrosine phosphorylation | 3.89 | 0.0044 |
| GO:0001701~in utero embryonic development | 2.39 | 0.0044 |
| GO:0048251~elastic fiber assembly | 11.35 | 0.0045 |
| GO:0032922~circadian regulation of gene expression | 3.84 | 0.0047 |
| GO:0006883~cellular sodium ion homeostasis | 7.10 | 0.0049 |
| GO:0043949~regulation of cAMP-mediated signaling | 25.54 | 0.0049 |
| GO:0016055~Wnt signaling pathway | 2.45 | 0.0051 |
| GO:0014912~negative regulation of smooth muscle cell migration | 10.48 | 0.0058 |
| GO:0032024~positive regulation of insulin secretion | 4.26 | 0.0058 |
| GO:0034765~regulation of ion transmembrane transport | 2.80 | 0.0061 |
| GO:0060348~bone development | 4.18 | 0.0063 |
| GO:0009612~response to mechanical stimulus | 4.18 | 0.0063 |
| GO:0043433~negative regulation of sequence-specific DNA binding transcription facto | 3.26 | 0.0064 |
| GO:0009267~cellular response to starvation | 3.63 | 0.0064 |
| GO:0001938~positive regulation of endothelial cell proliferation | 3.63 | 0.0064 |
| GO:0000122~negative regulation of transcription from RNA polymerase II promoter | 1.52 | 0.0064 |
| GO:0030177~positive regulation of Wnt signaling pathway | 4.98 | 0.0067 |
| GO:0071260~cellular response to mechanical stimulus | 3.59 | 0.0068 |
| GO:0045165~cell fate commitment | 4.11 | 0.0069 |
| GO:0071356~cellular response to tumor necrosis factor | 2.73 | 0.0072 |
| GO:0010955~negative regulation of protein processing | 9.73 | 0.0072 |
| GO:0086091~regulation of heart rate by cardiac conduction | 4.87 | 0.0074 |
| GO:0001666~response to hypoxia | 2.54 | 0.0080 |
| GO:0086045~membrane depolarization during AV node cell action potential | 20.44 | 0.0081 |
| GO:0045578~negative regulation of B cell differentiation | 20.44 | 0.0081 |
| GO:0031999~negative regulation of fatty acid beta-oxidation | 20.44 | 0.0081 |
| GO:0014856~skeletal muscle cell proliferation | 20.44 | 0.0081 |
| GO:0019640~glucuronate catabolic process to xylulose 5-phosphate | 20.44 | 0.0081 |
| GO:0051965~positive regulation of synapse assembly | 3.97 | 0.0081 |
| GO:0008285~negative regulation of cell proliferation | 1.79 | 0.0087 |
| GO:0030198~extracellular matrix organization | 2.38 | 0.0089 |
| GO:0030501~positive regulation of bone mineralization | 4.64 | 0.0090 |
| GO:0010628~positive regulation of gene expression | 1.73 | 0.0093 |
| GO:0045860~positive regulation of protein kinase activity | 3.85 | 0.0095 |
| hsa04964:Proximal tubule bicarbonate reclamation | 5.81 | 0.0096 |
| GO:0001656~metanephros development | 5.87 | 0.0097 |
| GO:0007275~multicellular organism development | 2.60 | 0.0100 |
| GO:0043547~positive regulation of GTPase activity | 2.34 | 0.0100 |
| hsa04713:Circadian entrainment | 2.75 | 0.0101 |
| hsa04927:Cortisol synthesis and secretion | 3.29 | 0.0103 |
| GO:0030522~intracellular receptor signaling pathway | 5.49 | 0.0123 |
| GO:0016525~negative regulation of angiogenesis | 2.68 | 0.0125 |
| GO:0071375~cellular response to peptide hormone stimulus | 8.01 | 0.0125 |
| GO:0051895~negative regulation of focal adhesion assembly | 8.01 | 0.0125 |
| GO:0001779~natural killer cell differentiation | 8.01 | 0.0125 |
| GO:0022409~positive regulation of cell-cell adhesion | 8.01 | 0.0125 |
| GO:0086005~ventricular cardiac muscle cell action potential | 8.01 | 0.0125 |
| GO:0098609~cell-cell adhesion | 2.27 | 0.0126 |
| hsa04934:Cushing syndrome | 2.24 | 0.0131 |
| hsa05202:Transcriptional misregulation in cancer | 2.08 | 0.0132 |
| hsa04910:Insulin signaling pathway | 2.34 | 0.0133 |
| GO:0048706~embryonic skeletal system development | 5.32 | 0.0137 |
| GO:0071773~cellular response to BMP stimulus | 5.32 | 0.0137 |
| GO:0032148~activation of protein kinase B activity | 5.32 | 0.0137 |
| GO:0007613~memory | 3.13 | 0.0139 |
| GO:0031667~response to nutrient levels | 4.17 | 0.0140 |
| hsa04935:Growth hormone synthesis, secretion and action | 2.45 | 0.0141 |

| | | |
|---|---|---|
| hsa04974:Protein digestion and absorption | 2.59 | 0.0146 |
| GO:2000573~positive regulation of DNA biosynthetic process | 7.57 | 0.0147 |
| GO:0060065~uterus development | 7.57 | 0.0147 |
| GO:0010812~negative regulation of cell-substrate adhesion | 7.57 | 0.0147 |
| GO:0008584~male gonad development | 2.81 | 0.0149 |
| GO:0006954~inflammatory response | 1.75 | 0.0153 |
| GO:0007015~actin filament organization | 2.43 | 0.0155 |
| GO:0001837~epithelial to mesenchymal transition | 4.01 | 0.0165 |
| GO:0045666~positive regulation of neuron differentiation | 3.03 | 0.0165 |
| GO:0090263~positive regulation of canonical Wnt signaling pathway | 2.56 | 0.0165 |
| GO:0043065~positive regulation of apoptotic process | 1.87 | 0.0167 |
| GO:0031668~cellular response to extracellular stimulus | 7.17 | 0.0171 |
| GO:0032355~response to estradiol | 2.99 | 0.0175 |
| h_dreamPathway:Repression of Pain Sensation by the Transcriptional Regulator DREAI | 6.72 | 0.0184 |
| GO:0050679~positive regulation of epithelial cell proliferation | 3.31 | 0.0189 |
| GO:1902895~positive regulation of pri-miRNA transcription from RNA polymerase II pi | 3.86 | 0.0192 |
| hsa04350:TGF-beta signaling pathway | 2.47 | 0.0193 |
| GO:2000179~positive regulation of neural precursor cell proliferation | 6.81 | 0.0197 |
| GO:0030539~male genitalia development | 6.81 | 0.0197 |
| GO:0032956~regulation of actin cytoskeleton organization | 2.90 | 0.0205 |
| h_il2rbPathway:IL-2 Receptor Beta Chain in T cell Activation | 3.62 | 0.0212 |
| GO:0043410~positive regulation of MAPK cascade | 2.31 | 0.0213 |
| GO:0021516~dorsal spinal cord development | 12.77 | 0.0214 |
| GO:0048167~regulation of synaptic plasticity | 3.72 | 0.0222 |
| GO:0007420~brain development | 1.96 | 0.0222 |
| GO:0007616~long-term memory | 4.60 | 0.0224 |
| GO:0048536~spleen development | 4.60 | 0.0224 |
| GO:0017157~regulation of exocytosis | 4.60 | 0.0224 |
| GO:0006366~transcription from RNA polymerase II promoter | 2.00 | 0.0237 |
| hsa04978:Mineral absorption | 3.12 | 0.0238 |
| GO:0042593~glucose homeostasis | 2.58 | 0.0239 |
| GO:0032147~activation of protein kinase activity | 4.48 | 0.0245 |
| GO:0007193~adenylate cyclase-inhibiting G-protein coupled receptor signaling pathwa | 3.59 | 0.0255 |
| GO:0014911~positive regulation of smooth muscle cell migration | 6.19 | 0.0255 |
| GO:0040037~negative regulation of fibroblast growth factor receptor signaling pathwa | 6.19 | 0.0255 |
| h_gata3Pathway:GATA3 participate in activating the Th2 cytokine genes expression | 5.88 | 0.0266 |
| GO:0060042~retina morphogenesis in camera-type eye | 11.35 | 0.0269 |
| GO:0034109~homotypic cell-cell adhesion | 11.35 | 0.0269 |
| GO:0051057~positive regulation of small GTPase mediated signal transduction | 11.35 | 0.0269 |
| GO:0050728~negative regulation of inflammatory response | 2.35 | 0.0272 |
| hsa04930:Type II diabetes mellitus | 3.48 | 0.0275 |
| GO:0042391~regulation of membrane potential | 2.72 | 0.0277 |
| hsa04724:Glutamatergic synapse | 2.32 | 0.0278 |
| GO:0007179~transforming growth factor beta receptor signaling pathway | 2.70 | 0.0290 |
| GO:0043627~response to estrogen | 3.46 | 0.0291 |
| GO:0060070~canonical Wnt signaling pathway | 2.67 | 0.0304 |
| GO:0007405~neuroblast proliferation | 4.15 | 0.0314 |
| GO:0007595~lactation | 4.15 | 0.0314 |
| hsa04218:Cellular senescence | 2.05 | 0.0315 |
| GO:0001889~liver development | 2.94 | 0.0316 |
| GO:0010906~regulation of glucose metabolic process | 5.68 | 0.0321 |
| GO:0030308~negative regulation of cell growth | 2.43 | 0.0322 |
| GO:0060371~regulation of atrial cardiac muscle cell membrane depolarization | 10.22 | 0.0330 |
| h_no1Pathway:Actions of Nitric Oxide in the Heart | 3.92 | 0.0339 |
| GO:0043588~skin development | 4.05 | 0.0340 |
| GO:0036120~cellular response to platelet-derived growth factor stimulus | 5.45 | 0.0357 |
| GO:0070509~calcium ion import | 5.45 | 0.0357 |
| GO:0019222~regulation of metabolic process | 5.45 | 0.0357 |

| | | |
|---|---|---|
| GO:0060038~cardiac muscle cell proliferation | 5.45 | 0.0357 |
| GO:0090316~positive regulation of intracellular protein transport | 5.45 | 0.0357 |
| GO:0030217~T cell differentiation | 3.96 | 0.0366 |
| GO:0045471~response to ethanol | 2.55 | 0.0380 |
| GO:0046777~protein autophosphorylation | 2.09 | 0.0385 |
| GO:0007267~cell-cell signaling | 1.93 | 0.0391 |
| GO:0001658~branching involved in ureteric bud morphogenesis | 3.87 | 0.0394 |
| GO:0042752~regulation of circadian rhythm | 3.19 | 0.0394 |
| GO:0030325~adrenal gland development | 5.24 | 0.0395 |
| GO:0048714~positive regulation of oligodendrocyte differentiation | 5.24 | 0.0395 |
| GO:1990535~neuron projection maintenance | 9.29 | 0.0396 |
| GO:0086009~membrane repolarization | 9.29 | 0.0396 |
| GO:0015816~glycine transport | 9.29 | 0.0396 |
| GO:0002237~response to molecule of bacterial origin | 9.29 | 0.0396 |
| GO:0021953~central nervous system neuron differentiation | 9.29 | 0.0396 |
| GO:0007010~cytoskeleton organization | 2.18 | 0.0406 |
| GO:0000165~MAPK cascade | 2.17 | 0.0420 |
| hsa04024:cAMP signaling pathway | 1.78 | 0.0421 |
| GO:0007189~adenylate cyclase-activating G-protein coupled receptor signaling pathwa | 2.30 | 0.0422 |
| GO:1990573~potassium ion import across plasma membrane | 3.78 | 0.0422 |
| GO:0002088~lens development in camera-type eye | 3.78 | 0.0422 |
| GO:0006836~neurotransmitter transport | 3.78 | 0.0422 |
| GO:0007205~protein kinase C-activating G-protein coupled receptor signaling pathway | 5.05 | 0.0435 |
| GO:0032689~negative regulation of interferon-gamma production | 3.70 | 0.0452 |
| GO:0048711~positive regulation of astrocyte differentiation | 8.51 | 0.0466 |
| GO:2001223~negative regulation of neuron migration | 8.51 | 0.0466 |
| GO:0048668~collateral sprouting | 8.51 | 0.0466 |
| GO:0090331~negative regulation of platelet aggregation | 8.51 | 0.0466 |
| GO:0086012~membrane depolarization during cardiac muscle cell action potential | 8.51 | 0.0466 |
| GO:0022407~regulation of cell-cell adhesion | 8.51 | 0.0466 |
| GO:0031098~stress-activated protein kinase signaling cascade | 8.51 | 0.0466 |
| GO:0034379~very-low-density lipoprotein particle assembly | 8.51 | 0.0466 |
| GO:0070588~calcium ion transmembrane transport | 2.43 | 0.0467 |
| GO:0032703~negative regulation of interleukin-2 production | 4.87 | 0.0477 |
| GO:0010842~retina layer formation | 4.87 | 0.0477 |
| GO:0007214~gamma-aminobutyric acid signaling pathway | 4.87 | 0.0477 |
| GO:0071280~cellular response to copper ion | 4.87 | 0.0477 |
| hsa04512:ECM-receptor interaction | 2.40 | 0.0481 |
| GO:0014823~response to activity | 3.62 | 0.0484 |
| GO:0001755~neural crest cell migration | 3.62 | 0.0484 |
