## Supplemental Table 8 for "Modulation of calcium signaling on demand to decipher the molecular mechanisms of primary aldosteronism"

### Supplementary Table 8. List of differentially expressed genes in H295R_S2 cells expressing the α7-5HT3 receptor in response to 24h treatment with 10-8M AngII

| Ensembl Gene ID | Gene Symbol | Gene Name | log2FoldChange | padj |
|---|---|---|---|---|
| ENSG00000128564 | VGF | VGF nerve growth factor inducible [Source:HGNC Symbol;Acc:HGNC:12684] | 2.19 | 7.15E-49 |
| ENSG00000126368 | NR1D1 | nuclear receptor subfamily 1 group D member 1 [Source:HGNC Symbol;Acc:HGNC:7962] | 1.25 | 2.19E-27 |
| ENSG00000049130 | KITLG | KIT ligand [Source:HGNC Symbol;Acc:HGNC:6343] | 1.96 | 7.90E-26 |
| ENSG00000198795 | ZNF521 | zinc finger protein 521 [Source:HGNC Symbol;Acc:HGNC:24605] | -0.71 | 1.66E-21 |
| ENSG00000095303 | PTGS1 | prostaglandin-endoperoxide synthase 1 [Source:HGNC Symbol;Acc:HGNC:9604] | 2.26 | 2.17E-16 |
| ENSG00000135919 | SERPINE2 | serpin family E member 2 [Source:HGNC Symbol;Acc:HGNC:8951] | 1.73 | 3.05E-16 |
| ENSG00000128050 | PAICS | phosphoribosylaminoimidazole carboxylase and phosphoribosylaminoimidazolesuccinocarboxamide synthase [Source:HGNC Symbol;Acc:HGNC:8587] | -0.63 | 7.67E-16 |
| ENSG00000057252 | SOAT1 | sterol O-acyltransferase 1 [Source:HGNC Symbol;Acc:HGNC:11177] | -0.88 | 3.38E-15 |
| ENSG00000137713 | PPP2R1B | protein phosphatase 2 scaffold subunit Abeta [Source:HGNC Symbol;Acc:HGNC:9303] | 1.33 | 4.57E-15 |
| ENSG00000166557 | TMED3 | transmembrane p24 trafficking protein 3 [Source:HGNC Symbol;Acc:HGNC:28889] | 1.02 | 7.09E-15 |
| ENSG00000145604 | SKP2 | S-phase kinase associated protein 2 [Source:HGNC Symbol;Acc:HGNC:10901] | -0.60 | 8.88E-14 |
| ENSG00000150893 | FREM2 | FRAS1 related extracellular matrix 2 [Source:HGNC Symbol;Acc:HGNC:25396] | 1.19 | 2.27E-13 |
| ENSG00000008405 | CRY1 | cryptochrome circadian regulator 1 [Source:HGNC Symbol;Acc:HGNC:2384] | -0.94 | 2.32E-13 |
| ENSG00000182512 | GLRX5 | glutaredoxin 5 [Source:HGNC Symbol;Acc:HGNC:20134] | -0.76 | 1.58E-12 |
| ENSG00000196611 | MMP1 | matrix metallopeptidase 1 [Source:HGNC Symbol;Acc:HGNC:7155] | 5.22 | 1.58E-12 |
| ENSG00000133687 | TMTC1 | transmembrane O-mannosyltransferase targeting cadherins 1 [Source:HGNC Symbol;Acc:HGNC:24099] | -1.16 | 1.72E-12 |
| ENSG00000243955 | GSTA1 | glutathione S-transferase alpha 1 [Source:HGNC Symbol;Acc:HGNC:4626] | -1.25 | 5.15E-12 |
| ENSG00000151790 | TDO2 | tryptophan 2,3-dioxygenase [Source:HGNC Symbol;Acc:HGNC:11708] | 1.44 | 5.80E-12 |
| ENSG00000081803 | CADPS2 | calcium dependent secretion activator 2 [Source:HGNC Symbol;Acc:HGNC:16018] | -1.41 | 7.80E-12 |
| ENSG00000160161 | CILP2 | cartilage intermediate layer protein 2 [Source:HGNC Symbol;Acc:HGNC:24213] | 1.40 | 7.80E-12 |
| ENSG00000101986 | ABCD1 | ATP binding cassette subfamily D member 1 [Source:HGNC Symbol;Acc:HGNC:61] | 0.72 | 7.80E-12 |
| ENSG00000180611 | MB21D2 | Mab-21 domain containing 2 [Source:HGNC Symbol;Acc:HGNC:30438] | -0.64 | 7.80E-12 |
| ENSG00000266401 | NA | novel transcript, antisense to DLGAP1 | 2.23 | 1.17E-11 |
| ENSG00000168917 | SLC35G2 | solute carrier family 35 member G2 [Source:HGNC Symbol;Acc:HGNC:28480] | 0.59 | 2.53E-11 |
| ENSG00000213465 | ARL 2,00 | ADP ribosylation factor like GTPase 2 [Source:HGNC Symbol;Acc:HGNC:693] | -0.51 | 2.90E-11 |
| ENSG00000054356 | PTPRN | protein tyrosine phosphatase receptor type N [Source:HGNC Symbol;Acc:HGNC:9676] | 2.41 | 3.03E-11 |
| ENSG00000153132 | CLGN | calmegin [Source:HGNC Symbol;Acc:HGNC:2060] | 0.61 | 3.03E-11 |
| ENSG00000053918 | KCNQ1 | potassium voltage-gated channel subfamily Q member 1 [Source:HGNC Symbol;Acc:HGNC:6294] | 2.35 | 4.06E-11 |
| ENSG00000123496 | IL13RA2 | interleukin 13 receptor subunit alpha 2 [Source:HGNC Symbol;Acc:HGNC:5975] | 0.67 | 4.06E-11 |
| ENSG00000102265 | TIMP1 | TIMP metallopeptidase inhibitor 1 [Source:HGNC Symbol;Acc:HGNC:11820] | 0.78 | 5.43E-11 |
| ENSG00000164733 | CTSB | cathepsin B [Source:HGNC Symbol;Acc:HGNC:2527] | 0.68 | 7.34E-11 |
| ENSG00000133800 | LYVE1 | lymphatic vessel endothelial hyaluronan receptor 1 [Source:HGNC Symbol;Acc:HGNC:14687] | 1.66 | 7.48E-11 |
| ENSG00000169508 | GPR183 | G protein-coupled receptor 183 [Source:HGNC Symbol;Acc:HGNC:3128] | -1.69 | 2.20E-10 |
| ENSG00000131015 | ULBP2 | UL16 binding protein 2 [Source:HGNC Symbol;Acc:HGNC:14894] | 1.83 | 2.91E-10 |
| ENSG00000170421 | KRT8 | keratin 8 [Source:HGNC Symbol;Acc:HGNC:6446] | 1.51 | 3.73E-10 |
| ENSG00000102962 | CCL22 | C-C motif chemokine ligand 22 [Source:HGNC Symbol;Acc:HGNC:10621] | 0.98 | 5.50E-10 |
| ENSG00000136928 | GABBR2 | gamma-aminobutyric acid type B receptor subunit 2 [Source:HGNC Symbol;Acc:HGNC:4507] | -0.77 | 6.06E-10 |
| ENSG00000156298 | TSPAN7 | tetraspanin 7 [Source:HGNC Symbol;Acc:HGNC:11854] | -0.56 | 7.03E-10 |
| ENSG00000105825 | TFPI2 | tissue factor pathway inhibitor 2 [Source:HGNC Symbol;Acc:HGNC:11761] | 1.94 | 7.56E-10 |
| ENSG00000261037 | NA | novel transcript | 1.73 | 8.79E-10 |
| ENSG00000101210 | EEF1A2 | eukaryotic translation elongation factor 1 alpha 2 [Source:HGNC Symbol;Acc:HGNC:3192] | 1.05 | 1.27E-09 |
| ENSG00000106624 | AEBP1 | AE binding protein 1 [Source:HGNC Symbol;Acc:HGNC:303] | 0.62 | 1.55E-09 |
| ENSG00000170624 | SGCD | sarcoglycan delta [Source:HGNC Symbol;Acc:HGNC:10807] | 0.72 | 1.55E-09 |
| ENSG00000076770 | MBNL3 | muscleblind like splicing regulator 3 [Source:HGNC Symbol;Acc:HGNC:20564] | -0.65 | 1.63E-09 |
| ENSG00000231852 | CYP21A2 | cytochrome P450 family 21 subfamily A member 2 [Source:HGNC Symbol;Acc:HGNC:2600] | 1.14 | 2.66E-09 |
| ENSG00000122034 | GTF3A | general transcription factor IIIA [Source:HGNC Symbol;Acc:HGNC:4662] | -0.57 | 4.85E-09 |

| | | | | |
|---|---|---|---|---|
| ENSG00000132970 | WASF3 | WASP family member 3 [Source:HGNC Symbol;Acc:HGNC:12734] | -0.64 | 6.85E-09 |
| ENSG00000148408 | CACNA1B | calcium voltage-gated channel subunit alpha1 B [Source:HGNC Symbol;Acc:HGNC:1389] | -0.66 | 8.02E-09 |
| ENSG00000168477 | TNXB | tenascin XB [Source:HGNC Symbol;Acc:HGNC:11976] | 1.26 | 8.02E-09 |
| ENSG00000237943 | PRKCQ-AS1 | PRKCQ antisense RNA 1 [Source:HGNC Symbol;Acc:HGNC:44689] | 0.75 | 8.84E-09 |
| ENSG00000272405 | NA | novel transcript, antisense to BCAN | 1.18 | 9.89E-09 |
| ENSG00000101000 | PROCR | protein C receptor [Source:HGNC Symbol;Acc:HGNC:9452] | 1.26 | 9.96E-09 |
| ENSG00000155265 | GOLGA7B | golgin A7 family member B [Source:HGNC Symbol;Acc:HGNC:31668] | 1.08 | 1.37E-08 |
| ENSG00000104341 | LAPTM4B | lysosomal protein transmembrane 4 beta [Source:HGNC Symbol;Acc:HGNC:13646] | 1.64 | 1.88E-08 |
| ENSG00000103855 | CD276 | CD276 molecule [Source:HGNC Symbol;Acc:HGNC:19137] | 0.63 | 2.33E-08 |
| ENSG00000006210 | CX3CL1 | C-X3-C motif chemokine ligand 1 [Source:HGNC Symbol;Acc:HGNC:10647] | 0.71 | 2.38E-08 |
| ENSG00000132357 | CARD6 | caspase recruitment domain family member 6 [Source:HGNC Symbol;Acc:HGNC:16394] | 0.72 | 2.57E-08 |
| ENSG00000179855 | GIPC3 | GIPC PDZ domain containing family member 3 [Source:HGNC Symbol;Acc:HGNC:18183] | 1.50 | 2.57E-08 |
| ENSG00000173641 | HSPB7 | heat shock protein family B (small) member 7 [Source:HGNC Symbol;Acc:HGNC:5249] | 0.69 | 3.07E-08 |
| ENSG00000110697 | PITPNM1 | phosphatidylinositol transfer protein membrane associated 1 [Source:HGNC Symbol;Acc:HGNC:9003] | 0.69 | 3.53E-08 |
| ENSG00000171159 | C9orf16 | chromosome 9 open reading frame 16 [Source:HGNC Symbol;Acc:HGNC:17823] | 0.73 | 3.74E-08 |
| ENSG00000183023 | SLC8A1 | solute carrier family 8 member A1 [Source:HGNC Symbol;Acc:HGNC:11068] | 1.84 | 3.74E-08 |
| ENSG00000197635 | DPP4 | dipeptidyl peptidase 4 [Source:HGNC Symbol;Acc:HGNC:3009] | 0.77 | 5.24E-08 |
| ENSG00000132561 | MATN2 | matrilin 2 [Source:HGNC Symbol;Acc:HGNC:6908] | 2.23 | 6.36E-08 |
| ENSG00000198753 | PLXNB3 | plexin B3 [Source:HGNC Symbol;Acc:HGNC:9105] | 1.04 | 7.06E-08 |
| ENSG00000185518 | SV2B | synaptic vesicle glycoprotein 2B [Source:HGNC Symbol;Acc:HGNC:16874] | -0.76 | 7.13E-08 |
| ENSG00000087842 | PIR | pirin [Source:HGNC Symbol;Acc:HGNC:30048] | -0.61 | 7.64E-08 |
| ENSG00000147889 | CDKN2A | cyclin dependent kinase inhibitor 2A [Source:HGNC Symbol;Acc:HGNC:1787] | -0.53 | 7.64E-08 |
| ENSG00000182534 | MXRA7 | matrix remodeling associated 7 [Source:HGNC Symbol;Acc:HGNC:7541] | 0.76 | 1.01E-07 |
| ENSG00000154639 | CXADR | CXADR Ig-like cell adhesion molecule [Source:HGNC Symbol;Acc:HGNC:2559] | 0.93 | 1.04E-07 |
| ENSG00000021300 | PLEKHB1 | pleckstrin homology domain containing B1 [Source:HGNC Symbol;Acc:HGNC:19079] | -0.59 | 1.27E-07 |
| ENSG00000007255 | TRAPPC6A | trafficking protein particle complex 6A [Source:HGNC Symbol;Acc:HGNC:23069] | -1.11 | 1.43E-07 |
| ENSG00000133710 | SPINK5 | serine peptidase inhibitor Kazal type 5 [Source:HGNC Symbol;Acc:HGNC:15464] | -1.03 | 1.57E-07 |
| ENSG00000105409 | ATP1A3 | ATPase Na+/K+ transporting subunit alpha 3 [Source:HGNC Symbol;Acc:HGNC:801] | -0.72 | 1.79E-07 |
| ENSG00000162337 | LRP5 | LDL receptor related protein 5 [Source:HGNC Symbol;Acc:HGNC:6697] | 0.56 | 1.82E-07 |
| ENSG00000174469 | CNTNAP2 | contactin associated protein 2 [Source:HGNC Symbol;Acc:HGNC:13830] | -1.16 | 1.83E-07 |
| ENSG00000113368 | LMNB1 | lamin B1 [Source:HGNC Symbol;Acc:HGNC:6637] | -0.54 | 2.32E-07 |
| ENSG00000196368 | NUDT11 | nudix hydrolase 11 [Source:HGNC Symbol;Acc:HGNC:18011] | 0.75 | 2.42E-07 |
| ENSG00000079257 | LXN | latexin [Source:HGNC Symbol;Acc:HGNC:13347] | 1.04 | 2.44E-07 |
| ENSG00000115457 | IGFBP2 | insulin like growth factor binding protein 2 [Source:HGNC Symbol;Acc:HGNC:5471] | 0.77 | 2.44E-07 |
| ENSG00000171724 | VAT1L | vesicle amine transport 1 like [Source:HGNC Symbol;Acc:HGNC:29315] | 1.23 | 2.85E-07 |
| ENSG00000158813 | EDA | ectodysplasin A [Source:HGNC Symbol;Acc:HGNC:3157] | 0.68 | 3.02E-07 |
| ENSG00000089639 | GMIP | GEM interacting protein [Source:HGNC Symbol;Acc:HGNC:24852] | 1.13 | 3.18E-07 |
| ENSG00000134215 | VAV3 | vav guanine nucleotide exchange factor 3 [Source:HGNC Symbol;Acc:HGNC:12659] | -0.75 | 3.40E-07 |
| ENSG00000139514 | SLC7A1 | solute carrier family 7 member 1 [Source:HGNC Symbol;Acc:HGNC:11057] | -0.54 | 3.73E-07 |
| ENSG00000149218 | ENDOD1 | endonuclease domain containing 1 [Source:HGNC Symbol;Acc:HGNC:29129] | 0.70 | 3.86E-07 |
| ENSG00000150995 | ITPR1 | inositol 1,4,5-trisphosphate receptor type 1 [Source:HGNC Symbol;Acc:HGNC:6180] | -0.81 | 5.34E-07 |
| ENSG00000139194 | RBP5 | retinol binding protein 5 [Source:HGNC Symbol;Acc:HGNC:15847] | 0.96 | 5.68E-07 |
| ENSG00000145623 | OSMR | oncostatin M receptor [Source:HGNC Symbol;Acc:HGNC:8507] | 0.86 | 6.05E-07 |
| ENSG00000157514 | TSC22D3 | TSC22 domain family member 3 [Source:HGNC Symbol;Acc:HGNC:3051] | -0.61 | 6.21E-07 |
| ENSG00000146648 | EGFR | epidermal growth factor receptor [Source:HGNC Symbol;Acc:HGNC:3236] | 2.34 | 6.55E-07 |
| ENSG00000144535 | DIS3L2 | DIS3 like 3'-5' exoribonuclease 2 [Source:HGNC Symbol;Acc:HGNC:28648] | 0.52 | 7.36E-07 |
| ENSG00000133083 | DCLK1 | doublecortin like kinase 1 [Source:HGNC Symbol;Acc:HGNC:2700] | -0.58 | 9.12E-07 |
| ENSG00000165124 | SVEP1 | sushi, von Willebrand factor type A, EGF and pentraxin domain containing 1 [Source:HGNC Symbol;Acc:HGNC:15985] | -1.22 | 9.67E-07 |

| | | | | |
|---|---|---|---|---|
| ENSG00000170915 | PAQR8 | progestin and adipoQ receptor family member 8 [Source:HGNC Symbol;Acc:HGNC:15708] | -0.63 | 1.06E-06 |
| ENSG00000146122 | DAAM2 | dishevelled associated activator of morphogenesis 2 [Source:HGNC Symbol;Acc:HGNC:18143] | -0.61 | 1.42E-06 |
| ENSG00000167693 | NXN | nucleoredoxin [Source:HGNC Symbol;Acc:HGNC:18008] | 1.21 | 1.57E-06 |
| ENSG00000168765 | GSTM4 | glutathione S-transferase mu 4 [Source:HGNC Symbol;Acc:HGNC:4636] | -0.75 | 1.59E-06 |
| ENSG00000116815 | CD58 | CD58 molecule [Source:HGNC Symbol;Acc:HGNC:1688] | 0.91 | 1.89E-06 |
| ENSG00000136628 | EPRS1 | glutamyl-prolyl-tRNA synthetase 1 [Source:HGNC Symbol;Acc:HGNC:3418] | -0.79 | 1.90E-06 |
| ENSG00000100426 | ZBED4 | zinc finger BED-type containing 4 [Source:HGNC Symbol;Acc:HGNC:20721] | -0.52 | 2.04E-06 |
| ENSG00000164904 | ALDH7A1 | aldehyde dehydrogenase 7 family member A1 [Source:HGNC Symbol;Acc:HGNC:877] | -0.51 | 2.26E-06 |
| ENSG00000136999 | CCN3 | cellular communication network factor 3 [Source:HGNC Symbol;Acc:HGNC:7885] | 0.75 | 3.01E-06 |
| ENSG00000103257 | SLC7A5 | solute carrier family 7 member 5 [Source:HGNC Symbol;Acc:HGNC:11063] | -0.86 | 3.01E-06 |
| ENSG00000178397 | FAM220A | family with sequence similarity 220 member A [Source:HGNC Symbol;Acc:HGNC:22422] | -19.33 | 3.04E-06 |
| ENSG00000120594 | PLXDC2 | plexin domain containing 2 [Source:HGNC Symbol;Acc:HGNC:21013] | -0.60 | 3.33E-06 |
| ENSG00000164050 | PLXNB1 | plexin B1 [Source:HGNC Symbol;Acc:HGNC:9103] | 0.71 | 3.40E-06 |
| ENSG00000101347 | SAMHD1 | SAM and HD domain containing deoxynucleoside triphosphate triphosphohydrolase 1 [Source:HGNC Symbol;Acc:HGNC:15925] | -0.68 | 3.60E-06 |
| ENSG00000110852 | CLEC2B | C-type lectin domain family 2 member B [Source:HGNC Symbol;Acc:HGNC:2053] | 0.67 | 3.63E-06 |
| ENSG00000064547 | LPAR2 | lysophosphatidic acid receptor 2 [Source:HGNC Symbol;Acc:HGNC:3168] | 1.19 | 3.65E-06 |
| ENSG00000221968 | FADS3 | fatty acid desaturase 3 [Source:HGNC Symbol;Acc:HGNC:3576] | 0.66 | 3.91E-06 |
| ENSG00000272068 | NA | novel transcript | 1.09 | 3.91E-06 |
| ENSG00000168938 | PPIC | peptidylprolyl isomerase C [Source:HGNC Symbol;Acc:HGNC:9256] | 1.05 | 4.38E-06 |
| ENSG00000139289 | PHLDA1 | pleckstrin homology like domain family A member 1 [Source:HGNC Symbol;Acc:HGNC:8933] | 0.57 | 4.53E-06 |
| ENSG00000155827 | RNF20 | ring finger protein 20 [Source:HGNC Symbol;Acc:HGNC:10062] | -0.62 | 4.60E-06 |
| ENSG00000005884 | ITGA3 | integrin subunit alpha 3 [Source:HGNC Symbol;Acc:HGNC:6139] | 1.19 | 4.78E-06 |
| ENSG00000132639 | SNAP25 | synaptosome associated protein 25 [Source:HGNC Symbol;Acc:HGNC:11132] | -0.92 | 5.11E-06 |
| ENSG00000105849 | TWISTNB | TWIST neighbor [Source:HGNC Symbol;Acc:HGNC:18027] | 0.90 | 5.49E-06 |
| ENSG00000166401 | SERPINB8 | serpin family B member 8 [Source:HGNC Symbol;Acc:HGNC:8952] | 1.16 | 6.22E-06 |
| ENSG00000175899 | A2M | alpha-2-macroglobulin [Source:HGNC Symbol;Acc:HGNC:7] | 0.50 | 6.48E-06 |
| ENSG00000117385 | P3H1 | prolyl 3-hydroxylase 1 [Source:HGNC Symbol;Acc:HGNC:19316] | 0.51 | 6.76E-06 |
| ENSG00000006327 | TNFRSF12A | TNF receptor superfamily member 12A [Source:HGNC Symbol;Acc:HGNC:18152] | 1.52 | 7.11E-06 |
| ENSG00000090530 | P3H2 | prolyl 3-hydroxylase 2 [Source:HGNC Symbol;Acc:HGNC:19317] | 0.65 | 7.11E-06 |
| ENSG00000167578 | RAB4B | RAB4B, member RAS oncogene family [Source:HGNC Symbol;Acc:HGNC:9782] | 0.60 | 7.11E-06 |
| ENSG00000186862 | PDZD7 | PDZ domain containing 7 [Source:HGNC Symbol;Acc:HGNC:26257] | 1.26 | 7.11E-06 |
| ENSG00000107738 | VSIR | V-set immunoregulatory receptor [Source:HGNC Symbol;Acc:HGNC:30085] | 2.31 | 7.51E-06 |
| ENSG00000069702 | TGFBR3 | transforming growth factor beta receptor 3 [Source:HGNC Symbol;Acc:HGNC:11774] | -0.67 | 7.70E-06 |
| ENSG00000139629 | GALNT6 | polypeptide N-acetylgalactosaminyltransferase 6 [Source:HGNC Symbol;Acc:HGNC:4128] | 0.94 | 7.81E-06 |
| ENSG00000164877 | MICALL2 | MICAL like 2 [Source:HGNC Symbol;Acc:HGNC:29672] | 1.27 | 7.94E-06 |
| ENSG00000138166 | DUSP5 | dual specificity phosphatase 5 [Source:HGNC Symbol;Acc:HGNC:3071] | 0.87 | 8.26E-06 |
| ENSG00000152620 | NADK2 | NAD kinase 2, mitochondrial [Source:HGNC Symbol;Acc:HGNC:26404] | -0.53 | 8.29E-06 |
| ENSG00000154217 | PITPNC1 | phosphatidylinositol transfer protein cytoplasmic 1 [Source:HGNC Symbol;Acc:HGNC:21045] | 0.92 | 8.53E-06 |
| ENSG00000242247 | ARFGAP3 | ADP ribosylation factor GTPase activating protein 3 [Source:HGNC Symbol;Acc:HGNC:661] | 0.64 | 8.65E-06 |
| ENSG00000118804 | STBD1 | starch binding domain 1 [Source:HGNC Symbol;Acc:HGNC:24854] | 0.58 | 8.70E-06 |
| ENSG00000077150 | NFKB2 | nuclear factor kappa B subunit 2 [Source:HGNC Symbol;Acc:HGNC:7795] | 0.60 | 1.01E-05 |
| ENSG00000102109 | PCSK1N | proprotein convertase subtilisin/kexin type 1 inhibitor [Source:HGNC Symbol;Acc:HGNC:17301] | -0.56 | 1.07E-05 |
| ENSG00000196557 | CACNA1H | calcium voltage-gated channel subunit alpha1 H [Source:HGNC Symbol;Acc:HGNC:1395] | 0.57 | 1.07E-05 |
| ENSG00000185100 | ADSS1 | adenylosuccinate synthase 1 [Source:HGNC Symbol;Acc:HGNC:20093] | 0.76 | 1.13E-05 |
| ENSG00000175600 | SUGCT | succinyl-CoA:glutarate-CoA transferase [Source:HGNC Symbol;Acc:HGNC:16001] | 2.19 | 1.16E-05 |
| ENSG00000286169 | PDCD6-AHRR | PDCD6-AHRR readthrough (NMD candidate) [Source:HGNC Symbol;Acc:HGNC:54724] | 0.56 | 1.16E-05 |
| ENSG00000189337 | KAZN | kazrin, periplakin interacting protein [Source:HGNC Symbol;Acc:HGNC:29173] | 0.75 | 1.21E-05 |
| ENSG00000139182 | CLSTN3 | calsyntenin 3 [Source:HGNC Symbol;Acc:HGNC:18371] | 0.79 | 1.23E-05 |

| | | | | |
|---|---|---|---|---|
| ENSG00000152580 | IGSF10 | immunoglobulin superfamily member 10 [Source:HGNC Symbol;Acc:HGNC:26384] | 1.41 | 1.42E-05 |
| ENSG00000111057 | KRT18 | keratin 18 [Source:HGNC Symbol;Acc:HGNC:6430] | 0.55 | 1.46E-05 |
| ENSG00000184500 | PROS1 | protein S [Source:HGNC Symbol;Acc:HGNC:9456] | 1.12 | 1.49E-05 |
| ENSG00000127824 | TUBA4A | tubulin alpha 4a [Source:HGNC Symbol;Acc:HGNC:12407] | 1.14 | 1.54E-05 |
| ENSG00000286638 | NA | novel transcript | -0.99 | 1.54E-05 |
| ENSG00000107798 | LIPA | lipase A, lysosomal acid type [Source:HGNC Symbol;Acc:HGNC:6617] | 0.56 | 1.55E-05 |
| ENSG00000174460 | ZCCHC12 | zinc finger CCHC-type containing 12 [Source:HGNC Symbol;Acc:HGNC:27273] | 0.96 | 1.72E-05 |
| ENSG00000100196 | KDELR3 | KDEL endoplasmic reticulum protein retention receptor 3 [Source:HGNC Symbol;Acc:HGNC:6306] | 0.86 | 1.85E-05 |
| ENSG00000130052 | STARD8 | StAR related lipid transfer domain containing 8 [Source:HGNC Symbol;Acc:HGNC:19161] | 0.52 | 1.87E-05 |
| ENSG00000106049 | HIBADH | 3-hydroxyisobutyrate dehydrogenase [Source:HGNC Symbol;Acc:HGNC:4907] | -0.57 | 1.91E-05 |
| ENSG00000131724 | IL13RA1 | interleukin 13 receptor subunit alpha 1 [Source:HGNC Symbol;Acc:HGNC:5974] | 0.52 | 2.11E-05 |
| ENSG00000260019 | LINC01992 | long intergenic non-protein coding RNA 1992 [Source:HGNC Symbol;Acc:HGNC:52824] | 1.45 | 2.30E-05 |
| ENSG00000054793 | ATP9A | ATPase phospholipid transporting 9A (putative) [Source:HGNC Symbol;Acc:HGNC:13540] | 0.50 | 2.51E-05 |
| ENSG00000233581 | NA | novel transcript | 1.59 | 2.57E-05 |
| ENSG00000072422 | RHOBTB1 | Rho related BTB domain containing 1 [Source:HGNC Symbol;Acc:HGNC:18738] | -0.57 | 2.58E-05 |
| ENSG00000102678 | FGF9 | fibroblast growth factor 9 [Source:HGNC Symbol;Acc:HGNC:3687] | -1.12 | 2.61E-05 |
| ENSG00000130429 | ARPC1B | actin related protein 2/3 complex subunit 1B [Source:HGNC Symbol;Acc:HGNC:704] | 0.73 | 3.06E-05 |
| ENSG00000130589 | HELZ2 | helicase with zinc finger 2 [Source:HGNC Symbol;Acc:HGNC:30021] | 1.24 | 3.11E-05 |
| ENSG00000158710 | TAGLN2 | transgelin 2 [Source:HGNC Symbol;Acc:HGNC:11554] | 0.55 | 3.30E-05 |
| ENSG00000104998 | IL27RA | interleukin 27 receptor subunit alpha [Source:HGNC Symbol;Acc:HGNC:17290] | 1.39 | 3.62E-05 |
| ENSG00000174010 | KLHL15 | kelch like family member 15 [Source:HGNC Symbol;Acc:HGNC:29347] | -0.53 | 3.62E-05 |
| ENSG00000137962 | ARHGAP29 | Rho GTPase activating protein 29 [Source:HGNC Symbol;Acc:HGNC:30207] | 1.91 | 4.31E-05 |
| ENSG00000161640 | SIGLEC11 | sialic acid binding Ig like lectin 11 [Source:HGNC Symbol;Acc:HGNC:15622] | -0.68 | 4.74E-05 |
| ENSG00000085831 | TTC39A | tetratricopeptide repeat domain 39A [Source:HGNC Symbol;Acc:HGNC:18657] | 1.39 | 4.80E-05 |
| ENSG00000124942 | AHNAK | AHNAK nucleoprotein [Source:HGNC Symbol;Acc:HGNC:347] | 2.05 | 4.96E-05 |
| ENSG00000136842 | TMOD1 | tropomodulin 1 [Source:HGNC Symbol;Acc:HGNC:11871] | 0.97 | 5.08E-05 |
| ENSG00000116717 | GADD45A | growth arrest and DNA damage inducible alpha [Source:HGNC Symbol;Acc:HGNC:4095] | 0.81 | 5.47E-05 |
| ENSG00000197948 | FCHSD1 | FCH and double SH3 domains 1 [Source:HGNC Symbol;Acc:HGNC:25463] | 0.57 | 6.35E-05 |
| ENSG00000147378 | FATE1 | fetal and adult testis expressed 1 [Source:HGNC Symbol;Acc:HGNC:24683] | -0.99 | 6.84E-05 |
| ENSG00000104043 | ATP8B4 | ATPase phospholipid transporting 8B4 (putative) [Source:HGNC Symbol;Acc:HGNC:13536] | 1.14 | 7.64E-05 |
| ENSG00000196739 | COL27A1 | collagen type XXVII alpha 1 chain [Source:HGNC Symbol;Acc:HGNC:22986] | 0.72 | 8.53E-05 |
| ENSG00000135074 | ADAM19 | ADAM metallopeptidase domain 19 [Source:HGNC Symbol;Acc:HGNC:197] | 1.62 | 8.59E-05 |
| ENSG00000117707 | PROX1 | prospero homeobox 1 [Source:HGNC Symbol;Acc:HGNC:9459] | 0.86 | 8.86E-05 |
| ENSG00000125730 | C3 | complement C3 [Source:HGNC Symbol;Acc:HGNC:1318] | 2.58 | 9.00E-05 |
| ENSG00000204386 | NEU1 | neuraminidase 1 [Source:HGNC Symbol;Acc:HGNC:7758] | 0.59 | 9.68E-05 |
| ENSG00000026025 | VIM | vimentin [Source:HGNC Symbol;Acc:HGNC:12692] | -0.50 | 0.000120 |
| ENSG00000088970 | KIZ | kizuna centrosomal protein [Source:HGNC Symbol;Acc:HGNC:15865] | 0.73 | 0.000131 |
| ENSG00000018236 | CNTN1 | contactin 1 [Source:HGNC Symbol;Acc:HGNC:2171] | 1.05 | 0.000131 |
| ENSG00000160211 | G6PD | glucose-6-phosphate dehydrogenase [Source:HGNC Symbol;Acc:HGNC:4057] | 0.52 | 0.000138 |
| ENSG00000198417 | MT1F | metallothionein 1F [Source:HGNC Symbol;Acc:HGNC:7398] | -0.71 | 0.000139 |
| ENSG00000087074 | PPP1R15A | protein phosphatase 1 regulatory subunit 15A [Source:HGNC Symbol;Acc:HGNC:14375] | 0.52 | 0.000147 |
| ENSG00000123080 | CDKN2C | cyclin dependent kinase inhibitor 2C [Source:HGNC Symbol;Acc:HGNC:1789] | -0.50 | 0.000149 |
| ENSG00000148123 | PLPPR1 | phospholipid phosphatase related 1 [Source:HGNC Symbol;Acc:HGNC:25993] | -0.63 | 0.000160 |
| ENSG00000176533 | GNG7 | G protein subunit gamma 7 [Source:HGNC Symbol;Acc:HGNC:4410] | 1.65 | 0.000160 |
| ENSG00000147174 | GCNA | germ cell nuclear acidic peptidase [Source:HGNC Symbol;Acc:HGNC:15805] | 1.90 | 0.000161 |
| ENSG00000278530 | CHMP1B2P | charged multivesicular body protein 1B2, pseudogene [Source:HGNC Symbol;Acc:HGNC:49380] | 0.83 | 0.000171 |
| ENSG00000135519 | KCNH3 | potassium voltage-gated channel subfamily H member 3 [Source:HGNC Symbol;Acc:HGNC:6252] | 0.57 | 0.000182 |
| ENSG00000111674 | ENO2 | enolase 2 [Source:HGNC Symbol;Acc:HGNC:3353] | 0.61 | 0.000194 |
| ENSG00000115956 | PLEK | pleckstrin [Source:HGNC Symbol;Acc:HGNC:9070] | 2.07 | 0.000205 |

| | | | | |
|---|---|---|---|---|
| ENSG00000258056 | NA | novel transcript, antisense to CD63 | -0.54 | 0.000213 |
| ENSG00000182704 | TSKU | tsukushi, small leucine rich proteoglycan [Source:HGNC Symbol;Acc:HGNC:28850] | 0.57 | 0.000229 |
| ENSG00000145685 | LHFPL2 | LHFPL tetraspan subfamily member 2 [Source:HGNC Symbol;Acc:HGNC:6588] | 0.53 | 0.000233 |
| ENSG00000022267 | FHL1 | four and a half LIM domains 1 [Source:HGNC Symbol;Acc:HGNC:3702] | 0.54 | 0.000248 |
| ENSG00000067057 | PFKP | phosphofructokinase, platelet [Source:HGNC Symbol;Acc:HGNC:8878] | 0.82 | 0.000249 |
| ENSG00000146250 | PRSS35 | serine protease 35 [Source:HGNC Symbol;Acc:HGNC:21387] | 0.95 | 0.000257 |
| ENSG00000225697 | SLC26A6 | solute carrier family 26 member 6 [Source:HGNC Symbol;Acc:HGNC:14472] | 0.62 | 0.000257 |
| ENSG00000234956 | LINC02539 | long intergenic non-protein coding RNA 2539 [Source:HGNC Symbol;Acc:HGNC:53572] | -0.92 | 0.000257 |
| ENSG00000100097 | LGALS1 | galectin 1 [Source:HGNC Symbol;Acc:HGNC:6561] | 0.70 | 0.000266 |
| ENSG00000091592 | NLRP1 | NLR family pyrin domain containing 1 [Source:HGNC Symbol;Acc:HGNC:14374] | 0.58 | 0.000276 |
| ENSG00000141639 | MAPK4 | mitogen-activated protein kinase 4 [Source:HGNC Symbol;Acc:HGNC:6878] | -0.82 | 0.000284 |
| ENSG00000145730 | PAM | peptidylglycine alpha-amidating monooxygenase [Source:HGNC Symbol;Acc:HGNC:8596] | 0.76 | 0.000297 |
| ENSG00000228536 | LYPLAL1-AS1 | LYPLAL1 antisense RNA 1 [Source:HGNC Symbol;Acc:HGNC:54054] | 0.80 | 0.000328 |
| ENSG00000139178 | C1RL | complement C1r subcomponent like [Source:HGNC Symbol;Acc:HGNC:21265] | 0.70 | 0.000328 |
| ENSG00000215218 | UBE2QL1 | ubiquitin conjugating enzyme E2 Q family like 1 [Source:HGNC Symbol;Acc:HGNC:37269] | 0.67 | 0.000334 |
| ENSG00000148090 | AUH | AU RNA binding methylglutaconyl-CoA hydratase [Source:HGNC Symbol;Acc:HGNC:890] | 0.50 | 0.000339 |
| ENSG00000139278 | GLIPR1 | GLI pathogenesis related 1 [Source:HGNC Symbol;Acc:HGNC:17001] | 0.85 | 0.000342 |
| ENSG00000082781 | ITGB5 | integrin subunit beta 5 [Source:HGNC Symbol;Acc:HGNC:6160] | 0.99 | 0.000346 |
| ENSG00000279561 | NA | uncharacterized LOC100132249 [Source:NCBI gene (formerly Entrezgene);Acc:100132249] | -0.56 | 0.000373 |
| ENSG00000179954 | SSC5D | scavenger receptor cysteine rich family member with 5 domains [Source:HGNC Symbol;Acc:HGNC:26641] | 1.69 | 0.000374 |
| ENSG00000166963 | MAP1A | microtubule associated protein 1A [Source:HGNC Symbol;Acc:HGNC:6835] | 0.70 | 0.000376 |
| ENSG00000249471 | ZNF324B | zinc finger protein 324B [Source:HGNC Symbol;Acc:HGNC:33107] | -0.57 | 0.000383 |
| ENSG00000184838 | PRR16 | proline rich 16 [Source:HGNC Symbol;Acc:HGNC:29654] | -0.63 | 0.000404 |
| ENSG00000090020 | SLC9A1 | solute carrier family 9 member A1 [Source:HGNC Symbol;Acc:HGNC:11071] | 0.59 | 0.000414 |
| ENSG00000161638 | ITGA5 | integrin subunit alpha 5 [Source:HGNC Symbol;Acc:HGNC:6141] | 1.42 | 0.000432 |
| ENSG00000136997 | MYC | MYC proto-oncogene, bHLH transcription factor [Source:HGNC Symbol;Acc:HGNC:7553] | -0.61 | 0.000439 |
| ENSG00000114378 | HYAL1 | hyaluronidase 1 [Source:HGNC Symbol;Acc:HGNC:5320] | 0.59 | 0.000441 |
| ENSG00000243766 | HOTTIP | HOXA distal transcript antisense RNA [Source:HGNC Symbol;Acc:HGNC:37461] | 0.70 | 0.000441 |
| ENSG00000085117 | CD82 | CD82 molecule [Source:HGNC Symbol;Acc:HGNC:6210] | 0.86 | 0.000458 |
| ENSG00000134121 | CHL1 | cell adhesion molecule L1 like [Source:HGNC Symbol;Acc:HGNC:1939] | 0.57 | 0.000467 |
| ENSG00000138162 | TACC2 | transforming acidic coiled-coil containing protein 2 [Source:HGNC Symbol;Acc:HGNC:11523] | 1.42 | 0.000471 |
| ENSG00000213949 | ITGA1 | integrin subunit alpha 1 [Source:HGNC Symbol;Acc:HGNC:6134] | 0.70 | 0.000471 |
| ENSG00000105711 | SCN1B | sodium voltage-gated channel beta subunit 1 [Source:HGNC Symbol;Acc:HGNC:10586] | 0.97 | 0.000475 |
| ENSG00000139428 | MMAB | metabolism of cobalamin associated B [Source:HGNC Symbol;Acc:HGNC:19331] | -0.59 | 0.000475 |
| ENSG00000247092 | SNHG10 | small nucleolar RNA host gene 10 [Source:HGNC Symbol;Acc:HGNC:27510] | -0.65 | 0.000476 |
| ENSG00000134871 | COL4A2 | collagen type IV alpha 2 chain [Source:HGNC Symbol;Acc:HGNC:2203] | 0.53 | 0.000481 |
| ENSG00000169330 | MINAR1 | membrane integral NOTCH2 associated receptor 1 [Source:HGNC Symbol;Acc:HGNC:29172] | 0.72 | 0.000481 |
| ENSG00000268350 | FAM156A | family with sequence similarity 156 member A [Source:HGNC Symbol;Acc:HGNC:30114] | 0.69 | 0.000494 |
| ENSG00000223802 | CERS1 | ceramide synthase 1 [Source:HGNC Symbol;Acc:HGNC:14253] | -0.60 | 0.000498 |
| ENSG00000070061 | ELP1 | elongator complex protein 1 [Source:HGNC Symbol;Acc:HGNC:5959] | 0.56 | 0.000507 |
| ENSG00000101977 | MCF 2,00 | MCF.2 cell line derived transforming sequence [Source:HGNC Symbol;Acc:HGNC:6940] | -0.85 | 0.000548 |
| ENSG00000179598 | PLD6 | phospholipase D family member 6 [Source:HGNC Symbol;Acc:HGNC:30447] | -0.82 | 0.000550 |
| ENSG00000100065 | CARD10 | caspase recruitment domain family member 10 [Source:HGNC Symbol;Acc:HGNC:16422] | 0.77 | 0.000561 |
| ENSG00000140479 | PCSK6 | proprotein convertase subtilisin/kexin type 6 [Source:HGNC Symbol;Acc:HGNC:8569] | 0.74 | 0.000577 |
| ENSG00000248115 | NA | novel transcript | -1.84 | 0.000590 |
| ENSG00000108797 | CNTNAP1 | contactin associated protein 1 [Source:HGNC Symbol;Acc:HGNC:8011] | 0.68 | 0.000592 |
| ENSG00000163191 | S100A11 | S100 calcium binding protein A11 [Source:HGNC Symbol;Acc:HGNC:10488] | 0.76 | 0.000600 |
| ENSG00000197467 | COL13A1 | collagen type XIII alpha 1 chain [Source:HGNC Symbol;Acc:HGNC:2190] | 2.05 | 0.000600 |

| | | | | |
|---|---|---|---|---|
| ENSG00000205208 | C4orf46 | chromosome 4 open reading frame 46 [Source:HGNC Symbol;Acc:HGNC:27320] | -0.58 | 0.000600 |
| ENSG00000079215 | SLC1A3 | solute carrier family 1 member 3 [Source:HGNC Symbol;Acc:HGNC:10941] | -0.65 | 0.000607 |
| ENSG00000087510 | TFAP2C | transcription factor AP-2 gamma [Source:HGNC Symbol;Acc:HGNC:11744] | 0.98 | 0.000625 |
| ENSG00000168394 | TAP1 | transporter 1, ATP binding cassette subfamily B member [Source:HGNC Symbol;Acc:HGNC:43] | 0.56 | 0.000649 |
| ENSG00000171004 | HS6ST2 | heparan sulfate 6-O-sulfotransferase 2 [Source:HGNC Symbol;Acc:HGNC:19133] | 1.04 | 0.000649 |
| ENSG00000173698 | ADGRG2 | adhesion G protein-coupled receptor G2 [Source:HGNC Symbol;Acc:HGNC:4516] | -1.03 | 0.000669 |
| ENSG00000170779 | CDCA4 | cell division cycle associated 4 [Source:HGNC Symbol;Acc:HGNC:14625] | -0.54 | 0.000688 |
| ENSG00000147459 | DOCK5 | dedicator of cytokinesis 5 [Source:HGNC Symbol;Acc:HGNC:23476] | -0.71 | 0.000695 |
| ENSG00000224389 | C4B | complement C4B (Chido blood group) [Source:HGNC Symbol;Acc:HGNC:1324] | 0.92 | 0.000696 |
| ENSG00000058799 | YIPF1 | Yip1 domain family member 1 [Source:HGNC Symbol;Acc:HGNC:25231] | 0.60 | 0.000707 |
| ENSG00000108387 | SEPTIN4 | septin 4 [Source:HGNC Symbol;Acc:HGNC:9165] | 0.71 | 0.000743 |
| ENSG00000189227 | C15orf61 | chromosome 15 open reading frame 61 [Source:HGNC Symbol;Acc:HGNC:34453] | -0.77 | 0.000756 |
| ENSG00000248008 | NRAV | negative regulator of antiviral response [Source:HGNC Symbol;Acc:HGNC:48588] | 0.54 | 0.000759 |
| ENSG00000158769 | F11R | F11 receptor [Source:HGNC Symbol;Acc:HGNC:14685] | 0.74 | 0.000772 |
| ENSG00000106546 | AHR | aryl hydrocarbon receptor [Source:HGNC Symbol;Acc:HGNC:348] | 0.78 | 0.000784 |
| ENSG00000139636 | LMBR1L | limb development membrane protein 1 like [Source:HGNC Symbol;Acc:HGNC:18268] | 0.54 | 0.000798 |
| ENSG00000175395 | ZNF25 | zinc finger protein 25 [Source:HGNC Symbol;Acc:HGNC:13043] | -0.51 | 0.000804 |
| ENSG00000108352 | RAPGEFL1 | Rap guanine nucleotide exchange factor like 1 [Source:HGNC Symbol;Acc:HGNC:17428] | 0.63 | 0.000826 |
| ENSG00000204876 | NA | uncharacterized LOC389602 [Source:NCBI gene (formerly Entrezgene);Acc:389602] | 0.92 | 0.000860 |
| ENSG00000130158 | DOCK6 | dedicator of cytokinesis 6 [Source:HGNC Symbol;Acc:HGNC:19189] | 1.17 | 0.000864 |
| ENSG00000187957 | DNER | delta/notch like EGF repeat containing [Source:HGNC Symbol;Acc:HGNC:24456] | -0.51 | 0.000864 |
| ENSG00000100949 | RABGGTA | Rab geranylgeranyltransferase subunit alpha [Source:HGNC Symbol;Acc:HGNC:9795] | 0.58 | 0.000976 |
| ENSG00000055118 | KCNH2 | potassium voltage-gated channel subfamily H member 2 [Source:HGNC Symbol;Acc:HGNC:6251] | 0.54 | 0.000991 |
| ENSG00000007516 | BAIAP3 | BAI1 associated protein 3 [Source:HGNC Symbol;Acc:HGNC:948] | 0.83 | 0.001003 |
| ENSG00000184009 | ACTG1 | actin gamma 1 [Source:HGNC Symbol;Acc:HGNC:144] | -0.56 | 0.001014 |
| ENSG00000186642 | PDE2A | phosphodiesterase 2A [Source:HGNC Symbol;Acc:HGNC:8777] | 0.62 | 0.001040 |
| ENSG00000144724 | PTPRG | protein tyrosine phosphatase receptor type G [Source:HGNC Symbol;Acc:HGNC:9671] | 0.75 | 0.001044 |
| ENSG00000008283 | CYB561 | cytochrome b561 [Source:HGNC Symbol;Acc:HGNC:2571] | 0.57 | 0.001076 |
| ENSG00000204516 | MICB | MHC class I polypeptide-related sequence B [Source:HGNC Symbol;Acc:HGNC:7091] | 0.79 | 0.001084 |
| ENSG00000116584 | ARHGEF2 | Rho/Rac guanine nucleotide exchange factor 2 [Source:HGNC Symbol;Acc:HGNC:682] | 0.62 | 0.001113 |
| ENSG00000235431 | NA | novel transcript | 0.69 | 0.001130 |
| ENSG00000128596 | CCDC136 | coiled-coil domain containing 136 [Source:HGNC Symbol;Acc:HGNC:22225] | 0.64 | 0.001167 |
| ENSG00000136859 | ANGPTL2 | angiopoietin like 2 [Source:HGNC Symbol;Acc:HGNC:490] | 0.75 | 0.001175 |
| ENSG00000112541 | PDE10A | phosphodiesterase 10A [Source:HGNC Symbol;Acc:HGNC:8772] | -0.84 | 0.001178 |
| ENSG00000133315 | MACROD1 | mono-ADP ribosylhydrolase 1 [Source:HGNC Symbol;Acc:HGNC:29598] | -0.54 | 0.001205 |
| ENSG00000155011 | DKK 2,00 | dickkopf WNT signaling pathway inhibitor 2 [Source:HGNC Symbol;Acc:HGNC:2892] | 1.52 | 0.001212 |
| ENSG00000250208 | FZD10-AS1 | FZD10 antisense divergent transcript [Source:HGNC Symbol;Acc:HGNC:48632] | 1.49 | 0.001221 |
| ENSG00000213366 | GSTM2 | glutathione S-transferase mu 2 [Source:HGNC Symbol;Acc:HGNC:4634] | -0.60 | 0.001236 |
| ENSG00000157110 | RBPMS | RNA binding protein, mRNA processing factor [Source:HGNC Symbol;Acc:HGNC:19097] | -0.61 | 0.001236 |
| ENSG00000169715 | MT1E | metallothionein 1E [Source:HGNC Symbol;Acc:HGNC:7397] | -0.75 | 0.001260 |
| ENSG00000152669 | CCNO | cyclin O [Source:HGNC Symbol;Acc:HGNC:18576] | 0.89 | 0.001267 |
| ENSG00000118777 | ABCG2 | ATP binding cassette subfamily G member 2 (Junior blood group) [Source:HGNC Symbol;Acc:HGNC:74] | 2.98 | 0.001284 |
| ENSG00000166831 | RBPMS2 | RNA binding protein, mRNA processing factor 2 [Source:HGNC Symbol;Acc:HGNC:19098] | 0.53 | 0.001343 |
| ENSG00000281832 | LINC00602 | long intergenic non-protein coding RNA 602 [Source:HGNC Symbol;Acc:HGNC:43917] | -1.76 | 0.001353 |
| ENSG00000162545 | CAMK2N1 | calcium/calmodulin dependent protein kinase II inhibitor 1 [Source:HGNC Symbol;Acc:HGNC:24190] | -0.85 | 0.001379 |
| ENSG00000136944 | LMX1B | LIM homeobox transcription factor 1 beta [Source:HGNC Symbol;Acc:HGNC:6654] | 0.62 | 0.001385 |
| ENSG00000163958 | ZDHHC19 | zinc finger DHHC-type palmitoyltransferase 19 [Source:HGNC Symbol;Acc:HGNC:20713] | -0.78 | 0.001437 |
| ENSG00000130768 | SMPDL3B | sphingomyelin phosphodiesterase acid like 3B [Source:HGNC Symbol;Acc:HGNC:21416] | 0.87 | 0.001540 |
| ENSG00000229953 | NA | novel transcript | 1.13 | 0.001546 |

| | | | | |
|---|---|---|---|---|
| ENSG00000272275 | NA | novel transcript | 1.12 | 0.001598 |
| ENSG00000167323 | STIM1 | stromal interaction molecule 1 [Source:HGNC Symbol;Acc:HGNC:11386] | 0.52 | 0.001785 |
| ENSG00000271614 | ATP2B1-AS1 | ATP2B1 antisense RNA 1 [Source:HGNC Symbol;Acc:HGNC:27883] | -0.91 | 0.001968 |
| ENSG00000006283 | CACNA1G | calcium voltage-gated channel subunit alpha1 G [Source:HGNC Symbol;Acc:HGNC:1394] | 0.56 | 0.002025 |
| ENSG00000105464 | GRIN2D | glutamate ionotropic receptor NMDA type subunit 2D [Source:HGNC Symbol;Acc:HGNC:4588] | 0.74 | 0.002032 |
| ENSG00000197320 | NA | C3 and PZP-like, alpha-2-macroglobulin domain containing 8 (CPAMD8) pseudogene | 0.94 | 0.002086 |
| ENSG00000092621 | PHGDH | phosphoglycerate dehydrogenase [Source:HGNC Symbol;Acc:HGNC:8923] | -0.50 | 0.002118 |
| ENSG00000144278 | GALNT13 | polypeptide N-acetylgalactosaminyltransferase 13 [Source:HGNC Symbol;Acc:HGNC:23242] | -0.61 | 0.002131 |
| ENSG00000185551 | NR2F2 | nuclear receptor subfamily 2 group F member 2 [Source:HGNC Symbol;Acc:HGNC:7976] | -0.54 | 0.002132 |
| ENSG00000166979 | EVA1C | eva-1 homolog C [Source:HGNC Symbol;Acc:HGNC:13239] | 1.02 | 0.002135 |
| ENSG00000170989 | S1PR1 | sphingosine-1-phosphate receptor 1 [Source:HGNC Symbol;Acc:HGNC:3165] | -1.27 | 0.002142 |
| ENSG00000124588 | NQO2 | N-ribosyldihydronicotinamide:quinone reductase 2 [Source:HGNC Symbol;Acc:HGNC:7856] | 0.55 | 0.002198 |
| ENSG00000233058 | LINC00884 | long intergenic non-protein coding RNA 884 [Source:HGNC Symbol;Acc:HGNC:48570] | 0.94 | 0.002198 |
| ENSG00000146386 | ABRACL | ABRA C-terminal like [Source:HGNC Symbol;Acc:HGNC:21230] | -0.55 | 0.002221 |
| ENSG00000267009 | NA | novel transcript | -1.41 | 0.002278 |
| ENSG00000010278 | CD9 | CD9 molecule [Source:HGNC Symbol;Acc:HGNC:1709] | 0.70 | 0.002312 |
| ENSG00000111087 | GLI1 | GLI family zinc finger 1 [Source:HGNC Symbol;Acc:HGNC:4317] | 1.13 | 0.002319 |
| ENSG00000116661 | FBXO2 | F-box protein 2 [Source:HGNC Symbol;Acc:HGNC:13581] | 0.78 | 0.002319 |
| ENSG00000067992 | PDK3 | pyruvate dehydrogenase kinase 3 [Source:HGNC Symbol;Acc:HGNC:8811] | -0.56 | 0.002328 |
| ENSG00000272414 | FAM47E-STBD1 | FAM47E-STBD1 readthrough [Source:HGNC Symbol;Acc:HGNC:44667] | 0.94 | 0.002328 |
| ENSG00000163040 | CCDC74A | coiled-coil domain containing 74A [Source:HGNC Symbol;Acc:HGNC:25197] | 0.64 | 0.002377 |
| ENSG00000092969 | TGFB2 | transforming growth factor beta 2 [Source:HGNC Symbol;Acc:HGNC:11768] | 0.84 | 0.002479 |
| ENSG00000115902 | SLC1A4 | solute carrier family 1 member 4 [Source:HGNC Symbol;Acc:HGNC:10942] | 0.55 | 0.002492 |
| ENSG00000237149 | ZNF503-AS2 | ZNF503 antisense RNA 2 [Source:HGNC Symbol;Acc:HGNC:23525] | 0.67 | 0.002620 |
| ENSG00000102755 | FLT1 | fms related receptor tyrosine kinase 1 [Source:HGNC Symbol;Acc:HGNC:3763] | 3.30 | 0.002621 |
| ENSG00000184349 | EFNA5 | ephrin A5 [Source:HGNC Symbol;Acc:HGNC:3225] | 0.89 | 0.002658 |
| ENSG00000103034 | NDRG4 | NDRG family member 4 [Source:HGNC Symbol;Acc:HGNC:14466] | 1.08 | 0.002687 |
| ENSG00000197566 | ZNF624 | zinc finger protein 624 [Source:HGNC Symbol;Acc:HGNC:29254] | -0.72 | 0.002898 |
| ENSG00000140853 | NLRC5 | NLR family CARD domain containing 5 [Source:HGNC Symbol;Acc:HGNC:29933] | 0.91 | 0.002900 |
| ENSG00000100979 | PLTP | phospholipid transfer protein [Source:HGNC Symbol;Acc:HGNC:9093] | 0.62 | 0.002925 |
| ENSG00000065054 | SLC9A3R2 | SLC9A3 regulator 2 [Source:HGNC Symbol;Acc:HGNC:11076] | 0.98 | 0.002931 |
| ENSG00000166311 | SMPD1 | sphingomyelin phosphodiesterase 1 [Source:HGNC Symbol;Acc:HGNC:11120] | 0.53 | 0.003128 |
| ENSG00000180155 | LYNX1 | Ly6/neurotoxin 1 [Source:HGNC Symbol;Acc:HGNC:29604] | 0.61 | 0.003128 |
| ENSG00000260807 | CEROX1 | cytoplasmic endogenous regulator of oxidative phosphorylation 1 [Source:HGNC Symbol;Acc:HGNC:53928] | 0.56 | 0.003128 |
| ENSG00000186469 | GNG2 | G protein subunit gamma 2 [Source:HGNC Symbol;Acc:HGNC:4404] | -0.51 | 0.003166 |
| ENSG00000145012 | LPP | LIM domain containing preferred translocation partner in lipoma [Source:HGNC Symbol;Acc:HGNC:6679] | 0.91 | 0.003169 |
| ENSG00000197696 | NMB | neuromedin B [Source:HGNC Symbol;Acc:HGNC:7842] | 0.93 | 0.003315 |
| ENSG00000136378 | ADAMTS7 | ADAM metallopeptidase with thrombospondin type 1 motif 7 [Source:HGNC Symbol;Acc:HGNC:223] | 0.85 | 0.003694 |
| ENSG00000249992 | TMEM158 | transmembrane protein 158 (gene/pseudogene) [Source:HGNC Symbol;Acc:HGNC:30293] | -0.62 | 0.003749 |
| ENSG00000091986 | CCDC80 | coiled-coil domain containing 80 [Source:HGNC Symbol;Acc:HGNC:30649] | 0.55 | 0.003766 |
| ENSG00000104774 | MAN2B1 | mannosidase alpha class 2B member 1 [Source:HGNC Symbol;Acc:HGNC:6826] | 0.58 | 0.003785 |
| ENSG00000164690 | SHH | sonic hedgehog signaling molecule [Source:HGNC Symbol;Acc:HGNC:10848] | -0.69 | 0.003852 |
| ENSG00000286478 | NA | novel transcript, antisense to HIBADH | 0.99 | 0.003853 |
| ENSG00000087085 | ACHE | acetylcholinesterase (Cartwright blood group) [Source:HGNC Symbol;Acc:HGNC:108] | 0.93 | 0.003872 |
| ENSG00000269293 | ZSCAN16-AS1 | ZSCAN16 antisense RNA 1 [Source:HGNC Symbol;Acc:HGNC:48982] | -0.59 | 0.004046 |
| ENSG00000112378 | PERP | p53 apoptosis effector related to PMP22 [Source:HGNC Symbol;Acc:HGNC:17637] | 0.57 | 0.004093 |
| ENSG00000114315 | HES1 | hes family bHLH transcription factor 1 [Source:HGNC Symbol;Acc:HGNC:5192] | 0.75 | 0.004093 |

| | | | | |
|---|---|---|---|---|
| ENSG00000130558 | OLFM1 | olfactomedin 1 [Source:HGNC Symbol;Acc:HGNC:17187] | 0.57 | 0.004093 |
| ENSG00000115226 | FNDC4 | fibronectin type III domain containing 4 [Source:HGNC Symbol;Acc:HGNC:20239] | 0.66 | 0.004150 |
| ENSG00000131831 | RAI2 | retinoic acid induced 2 [Source:HGNC Symbol;Acc:HGNC:9835] | -0.55 | 0.004308 |
| ENSG00000287151 | NA | chromosome 2 open reading frame 27A [Source:NCBI gene (formerly Entrezgene);Acc:29798] | 0.71 | 0.004341 |
| ENSG00000141314 | RHBDL3 | rhomboid like 3 [Source:HGNC Symbol;Acc:HGNC:16502] | 0.66 | 0.004342 |
| ENSG00000167861 | HID1 | HID1 domain containing [Source:HGNC Symbol;Acc:HGNC:15736] | 0.62 | 0.004395 |
| ENSG00000105877 | DNAH11 | dynein axonemal heavy chain 11 [Source:HGNC Symbol;Acc:HGNC:2942] | 0.80 | 0.004431 |
| ENSG00000174307 | PHLDA3 | pleckstrin homology like domain family A member 3 [Source:HGNC Symbol;Acc:HGNC:8934] | 1.13 | 0.004511 |
| ENSG00000119888 | EPCAM | epithelial cell adhesion molecule [Source:HGNC Symbol;Acc:HGNC:11529] | 0.89 | 0.004713 |
| ENSG00000135547 | HEY2 | hes related family bHLH transcription factor with YRPW motif 2 [Source:HGNC Symbol;Acc:HGNC:4881] | -0.70 | 0.004906 |
| ENSG00000155966 | AFF2 | AF4/FMR2 family member 2 [Source:HGNC Symbol;Acc:HGNC:3776] | -0.52 | 0.004906 |
| ENSG00000158715 | SLC45A3 | solute carrier family 45 member 3 [Source:HGNC Symbol;Acc:HGNC:8642] | 1.27 | 0.004906 |
| ENSG00000107819 | SFXN3 | sideroflexin 3 [Source:HGNC Symbol;Acc:HGNC:16087] | 0.75 | 0.005092 |
| ENSG00000123146 | ADGRE5 | adhesion G protein-coupled receptor E5 [Source:HGNC Symbol;Acc:HGNC:1711] | 1.42 | 0.005102 |
| ENSG00000118971 | CCND2 | cyclin D2 [Source:HGNC Symbol;Acc:HGNC:1583] | -0.52 | 0.005216 |
| ENSG00000142173 | COL6A2 | collagen type VI alpha 2 chain [Source:HGNC Symbol;Acc:HGNC:2212] | 0.79 | 0.005364 |
| ENSG00000132199 | ENOSF1 | enolase superfamily member 1 [Source:HGNC Symbol;Acc:HGNC:30365] | -0.69 | 0.005387 |
| ENSG00000176490 | DIRAS1 | DIRAS family GTPase 1 [Source:HGNC Symbol;Acc:HGNC:19127] | 0.57 | 0.005435 |
| ENSG00000175344 | CHRNA7 | cholinergic receptor nicotinic alpha 7 subunit [Source:HGNC Symbol;Acc:HGNC:1960] | -0.80 | 0.005436 |
| ENSG00000150594 | ADRA2A | adrenoceptor alpha 2A [Source:HGNC Symbol;Acc:HGNC:281] | -0.65 | 0.005464 |
| ENSG00000286190 | NA | uncharacterized LOC728392 [Source:NCBI gene (formerly Entrezgene);Acc:728392] | 0.66 | 0.005491 |
| ENSG00000144485 | HES6 | hes family bHLH transcription factor 6 [Source:HGNC Symbol;Acc:HGNC:18254] | 0.76 | 0.005547 |
| ENSG00000268471 | MIR4453HG | MIR4453 host gene [Source:HGNC Symbol;Acc:HGNC:25288] | -0.52 | 0.005705 |
| ENSG00000167100 | SAMD14 | sterile alpha motif domain containing 14 [Source:HGNC Symbol;Acc:HGNC:27312] | 0.62 | 0.005881 |
| ENSG00000126733 | DACH2 | dachshund family transcription factor 2 [Source:HGNC Symbol;Acc:HGNC:16814] | 0.77 | 0.005883 |
| ENSG00000119915 | ELOVL3 | ELOVL fatty acid elongase 3 [Source:HGNC Symbol;Acc:HGNC:18047] | 0.70 | 0.006102 |
| ENSG00000153029 | MR1 | major histocompatibility complex, class I-related [Source:HGNC Symbol;Acc:HGNC:4975] | 0.55 | 0.006115 |
| ENSG00000224081 | SLC44A3-AS1 | SLC44A3 antisense RNA 1 [Source:HGNC Symbol;Acc:HGNC:49057] | 1.00 | 0.006197 |
| ENSG00000163377 | TAFA4 | TAFA chemokine like family member 4 [Source:HGNC Symbol;Acc:HGNC:21591] | -0.60 | 0.006219 |
| ENSG00000164761 | TNFRSF11B | TNF receptor superfamily member 11b [Source:HGNC Symbol;Acc:HGNC:11909] | 0.61 | 0.006313 |
| ENSG00000146350 | TBC1D32 | TBC1 domain family member 32 [Source:HGNC Symbol;Acc:HGNC:21485] | -0.64 | 0.006462 |
| ENSG00000110811 | P3H3 | prolyl 3-hydroxylase 3 [Source:HGNC Symbol;Acc:HGNC:19318] | 0.79 | 0.006499 |
| ENSG00000168993 | CPLX1 | complexin 1 [Source:HGNC Symbol;Acc:HGNC:2309] | -0.76 | 0.006631 |
| ENSG00000139352 | ASCL1 | achaete-scute family bHLH transcription factor 1 [Source:HGNC Symbol;Acc:HGNC:738] | -0.98 | 0.006684 |
| ENSG00000070961 | ATP2B1 | ATPase plasma membrane Ca2+ transporting 1 [Source:HGNC Symbol;Acc:HGNC:814] | -0.53 | 0.007050 |
| ENSG00000118785 | SPP1 | secreted phosphoprotein 1 [Source:HGNC Symbol;Acc:HGNC:11255] | 0.84 | 0.007073 |
| ENSG00000165474 | GJB2 | gap junction protein beta 2 [Source:HGNC Symbol;Acc:HGNC:4284] | 1.06 | 0.007367 |
| ENSG00000244067 | GSTA2 | glutathione S-transferase alpha 2 [Source:HGNC Symbol;Acc:HGNC:4627] | -1.33 | 0.007367 |
| ENSG00000108960 | MMD | monocyte to macrophage differentiation associated [Source:HGNC Symbol;Acc:HGNC:7153] | -0.60 | 0.007373 |
| ENSG00000130598 | TNNI2 | troponin I2, fast skeletal type [Source:HGNC Symbol;Acc:HGNC:11946] | 0.75 | 0.007433 |
| ENSG00000227906 | SNAP25-AS1 | SNAP25 antisense RNA 1 [Source:HGNC Symbol;Acc:HGNC:44312] | -1.11 | 0.007433 |
| ENSG00000112320 | SOBP | sine oculis binding protein homolog [Source:HGNC Symbol;Acc:HGNC:29256] | -0.52 | 0.007483 |
| ENSG00000119508 | NR4A3 | nuclear receptor subfamily 4 group A member 3 [Source:HGNC Symbol;Acc:HGNC:7982] | -0.99 | 0.007483 |
| ENSG00000121236 | TRIM6 | tripartite motif containing 6 [Source:HGNC Symbol;Acc:HGNC:16277] | 0.75 | 0.007483 |
| ENSG00000153823 | PID1 | phosphotyrosine interaction domain containing 1 [Source:HGNC Symbol;Acc:HGNC:26084] | -0.58 | 0.007563 |
| ENSG00000244731 | C4A | complement C4A (Rodgers blood group) [Source:HGNC Symbol;Acc:HGNC:1323] | 0.79 | 0.007635 |
| ENSG00000183688 | RFLNB | refilin B [Source:HGNC Symbol;Acc:HGNC:28705] | 0.94 | 0.007657 |
| ENSG00000268996 | MAN1B1-DT | MAN1B1 divergent transcript [Source:HGNC Symbol;Acc:HGNC:48715] | -0.55 | 0.007658 |
| ENSG00000138678 | GPAT3 | glycerol-3-phosphate acyltransferase 3 [Source:HGNC Symbol;Acc:HGNC:28157] | 0.60 | 0.007832 |

| | | | | |
|---|---|---|---|---|
| ENSG00000060709 | RIMBP2 | RIMS binding protein 2 [Source:HGNC Symbol;Acc:HGNC:30339] | 0.84 | 0.007980 |
| ENSG00000131650 | KREMEN2 | kringle containing transmembrane protein 2 [Source:HGNC Symbol;Acc:HGNC:18797] | 0.52 | 0.008256 |
| ENSG00000120820 | GLT8D2 | glycosyltransferase 8 domain containing 2 [Source:HGNC Symbol;Acc:HGNC:24890] | 0.72 | 0.008352 |
| ENSG00000108813 | DLX4 | distal-less homeobox 4 [Source:HGNC Symbol;Acc:HGNC:2917] | 0.60 | 0.008455 |
| ENSG00000284776 | NA | novel protein | -0.58 | 0.008489 |
| ENSG00000155974 | GRIP1 | glutamate receptor interacting protein 1 [Source:HGNC Symbol;Acc:HGNC:18708] | -0.69 | 0.008696 |
| ENSG00000124507 | PACSIN1 | protein kinase C and casein kinase substrate in neurons 1 [Source:HGNC Symbol;Acc:HGNC:8570] | 1.20 | 0.008742 |
| ENSG00000137869 | CYP19A1 | cytochrome P450 family 19 subfamily A member 1 [Source:HGNC Symbol;Acc:HGNC:2594] | 1.58 | 0.008783 |
| ENSG00000149212 | SESN3 | sestrin 3 [Source:HGNC Symbol;Acc:HGNC:23060] | -0.52 | 0.008783 |
| ENSG00000197859 | ADAMTSL2 | ADAMTS like 2 [Source:HGNC Symbol;Acc:HGNC:14631] | 1.61 | 0.008831 |
| ENSG00000003436 | TFPI | tissue factor pathway inhibitor [Source:HGNC Symbol;Acc:HGNC:11760] | 0.73 | 0.008863 |
| ENSG00000100003 | SEC14L2 | SEC14 like lipid binding 2 [Source:HGNC Symbol;Acc:HGNC:10699] | 0.52 | 0.008908 |
| ENSG00000073060 | SCARB1 | scavenger receptor class B member 1 [Source:HGNC Symbol;Acc:HGNC:1664] | 0.57 | 0.008926 |
| ENSG00000111981 | ULBP1 | UL16 binding protein 1 [Source:HGNC Symbol;Acc:HGNC:14893] | 0.62 | 0.009655 |
| ENSG00000240445 | FOXO3B | forkhead box O3B [Source:HGNC Symbol;Acc:HGNC:3822] | -0.60 | 0.009698 |
| ENSG00000105717 | PBX4 | PBX homeobox 4 [Source:HGNC Symbol;Acc:HGNC:13403] | 0.84 | 0.009732 |
| ENSG00000128510 | CPA4 | carboxypeptidase A4 [Source:HGNC Symbol;Acc:HGNC:15740] | 1.81 | 0.009852 |
| ENSG00000182255 | KCNA4 | potassium voltage-gated channel subfamily A member 4 [Source:HGNC Symbol;Acc:HGNC:6222] | -0.77 | 0.009889 |
| ENSG00000153214 | TMEM87B | transmembrane protein 87B [Source:HGNC Symbol;Acc:HGNC:25913] | 0.69 | 0.009894 |
| ENSG00000137261 | KIAA0319 | KIAA0319 [Source:HGNC Symbol;Acc:HGNC:21580] | -0.63 | 0.010035 |
| ENSG00000124406 | ATP8A1 | ATPase phospholipid transporting 8A1 [Source:HGNC Symbol;Acc:HGNC:13531] | 0.67 | 0.010132 |
| ENSG00000173281 | PPP1R3B | protein phosphatase 1 regulatory subunit 3B [Source:HGNC Symbol;Acc:HGNC:14942] | -0.75 | 0.010545 |
| ENSG00000133818 | RRAS2 | RAS related 2 [Source:HGNC Symbol;Acc:HGNC:17271] | -0.53 | 0.010573 |
| ENSG00000104419 | NDRG1 | N-myc downstream regulated 1 [Source:HGNC Symbol;Acc:HGNC:7679] | 0.83 | 0.010653 |
| ENSG00000272145 | NFYC-AS1 | NFYC antisense RNA 1 [Source:HGNC Symbol;Acc:HGNC:49451] | -0.55 | 0.010653 |
| ENSG00000100311 | PDGFB | platelet derived growth factor subunit B [Source:HGNC Symbol;Acc:HGNC:8800] | 0.97 | 0.010944 |
| ENSG00000121361 | KCNJ8 | potassium inwardly rectifying channel subfamily J member 8 [Source:HGNC Symbol;Acc:HGNC:6269] | -0.64 | 0.011550 |
| ENSG00000073849 | ST6GAL1 | ST6 beta-galactoside alpha-2,6-sialyltransferase 1 [Source:HGNC Symbol;Acc:HGNC:10860] | -0.56 | 0.011568 |
| ENSG00000072274 | TFRC | transferrin receptor [Source:HGNC Symbol;Acc:HGNC:11763] | -0.58 | 0.011979 |
| ENSG00000164520 | RAET1E | retinoic acid early transcript 1E [Source:HGNC Symbol;Acc:HGNC:16793] | 1.41 | 0.012104 |
| ENSG00000144959 | NCEH1 | neutral cholesterol ester hydrolase 1 [Source:HGNC Symbol;Acc:HGNC:29260] | 0.53 | 0.012110 |
| ENSG00000140511 | HAPLN3 | hyaluronan and proteoglycan link protein 3 [Source:HGNC Symbol;Acc:HGNC:21446] | 0.82 | 0.012128 |
| ENSG00000137727 | ARHGAP20 | Rho GTPase activating protein 20 [Source:HGNC Symbol;Acc:HGNC:18357] | 0.60 | 0.012277 |
| ENSG00000196337 | CGB7 | chorionic gonadotropin subunit beta 7 [Source:HGNC Symbol;Acc:HGNC:16451] | 1.04 | 0.012305 |
| ENSG00000261594 | TPBGL | trophoblast glycoprotein like [Source:HGNC Symbol;Acc:HGNC:44159] | 0.91 | 0.012305 |
| ENSG00000274718 | NA | novel transcript | -0.86 | 0.012343 |
| ENSG00000138411 | HECW2 | HECT, C2 and WW domain containing E3 ubiquitin protein ligase 2 [Source:HGNC Symbol;Acc:HGNC:29853] | 0.80 | 0.012699 |
| ENSG00000065600 | PACC1 | proton activated chloride channel 1 [Source:HGNC Symbol;Acc:HGNC:25593] | 0.59 | 0.013077 |
| ENSG00000079156 | OSBPL6 | oxysterol binding protein like 6 [Source:HGNC Symbol;Acc:HGNC:16388] | -0.71 | 0.013226 |
| ENSG00000066629 | EML1 | EMAP like 1 [Source:HGNC Symbol;Acc:HGNC:3330] | 0.65 | 0.013301 |
| ENSG00000171951 | SCG2 | secretogranin II [Source:HGNC Symbol;Acc:HGNC:10575] | 1.19 | 0.013501 |
| ENSG00000205809 | KLRC2 | killer cell lectin like receptor C2 [Source:HGNC Symbol;Acc:HGNC:6375] | 3.21 | 0.013508 |
| ENSG00000077943 | ITGA8 | integrin subunit alpha 8 [Source:HGNC Symbol;Acc:HGNC:6144] | 0.59 | 0.013531 |
| ENSG00000077092 | RARB | retinoic acid receptor beta [Source:HGNC Symbol;Acc:HGNC:9865] | -0.80 | 0.013685 |
| ENSG00000229847 | EMX2OS | EMX2 opposite strand/antisense RNA [Source:HGNC Symbol;Acc:HGNC:18511] | 1.00 | 0.013689 |
| ENSG00000182165 | TP53TG1 | TP53 target 1 [Source:HGNC Symbol;Acc:HGNC:17026] | -0.55 | 0.013756 |
| ENSG00000186594 | MIR22HG | MIR22 host gene [Source:HGNC Symbol;Acc:HGNC:28219] | 0.58 | 0.013922 |
| ENSG00000135502 | SLC26A10 | solute carrier family 26 member 10 [Source:HGNC Symbol;Acc:HGNC:14470] | 0.85 | 0.014111 |

| | | | | |
|---|---|---|---|---|
| ENSG00000100036 | SLC35E4 | solute carrier family 35 member E4 [Source:HGNC Symbol;Acc:HGNC:17058] | 0.68 | 0.014361 |
| ENSG00000171812 | COL8A2 | collagen type VIII alpha 2 chain [Source:HGNC Symbol;Acc:HGNC:2216] | 0.98 | 0.014388 |
| ENSG00000174791 | RIN1 | Ras and Rab interactor 1 [Source:HGNC Symbol;Acc:HGNC:18749] | 1.05 | 0.014986 |
| ENSG00000107807 | TLX1 | T cell leukemia homeobox 1 [Source:HGNC Symbol;Acc:HGNC:5056] | 1.15 | 0.015031 |
| ENSG00000176597 | B3GNT5 | UDP-GlcNAc:betaGal beta-1,3-N-acetylglucosaminyltransferase 5 [Source:HGNC Symbol;Acc:HGNC:15684] | 1.48 | 0.015135 |
| ENSG00000172159 | FRMD3 | FERM domain containing 3 [Source:HGNC Symbol;Acc:HGNC:24125] | 0.60 | 0.015293 |
| ENSG00000130294 | KIF1A | kinesin family member 1A [Source:HGNC Symbol;Acc:HGNC:888] | -0.71 | 0.015716 |
| ENSG00000167191 | GPRC5B | G protein-coupled receptor class C group 5 member B [Source:HGNC Symbol;Acc:HGNC:13308] | 0.67 | 0.015793 |
| ENSG00000168970 | JMJD7-PLA2G4B | JMJD7-PLA2G4B readthrough [Source:HGNC Symbol;Acc:HGNC:34449] | 0.73 | 0.015818 |
| ENSG00000179142 | CYP11B2 | cytochrome P450 family 11 subfamily B member 2 [Source:HGNC Symbol;Acc:HGNC:2592] | 1.53 | 0.016265 |
| ENSG00000023902 | PLEKHO1 | pleckstrin homology domain containing O1 [Source:HGNC Symbol;Acc:HGNC:24310] | 0.66 | 0.016344 |
| ENSG00000139549 | DHH | desert hedgehog signaling molecule [Source:HGNC Symbol;Acc:HGNC:2865] | 1.14 | 0.016445 |
| ENSG00000119922 | IFIT2 | interferon induced protein with tetratricopeptide repeats 2 [Source:HGNC Symbol;Acc:HGNC:5409] | 0.58 | 0.016461 |
| ENSG00000229422 | NA | novel transcript | -0.53 | 0.017001 |
| ENSG00000214237 | MINDY4B | MINDY family member 4B [Source:HGNC Symbol;Acc:HGNC:35475] | -0.53 | 0.017091 |
| ENSG00000237651 | C2orf74 | chromosome 2 open reading frame 74 [Source:HGNC Symbol;Acc:HGNC:34439] | -0.62 | 0.017424 |
| ENSG00000227036 | LINC00511 | long intergenic non-protein coding RNA 511 [Source:HGNC Symbol;Acc:HGNC:43564] | 0.91 | 0.017756 |
| ENSG00000167434 | CA4 | carbonic anhydrase 4 [Source:HGNC Symbol;Acc:HGNC:1375] | 0.68 | 0.018332 |
| ENSG00000237499 | WAKMAR2 | wound and keratinocyte migration associated lncRNA 2 [Source:HGNC Symbol;Acc:HGNC:53754] | -0.63 | 0.018353 |
| ENSG00000170092 | SPDYE5 | speedy/RINGO cell cycle regulator family member E5 [Source:HGNC Symbol;Acc:HGNC:35464] | 1.26 | 0.018398 |
| ENSG00000072952 | IRAG1 | inositol 1,4,5-triphosphate receptor associated 1 [Source:HGNC Symbol;Acc:HGNC:7237] | 0.68 | 0.018514 |
| ENSG00000142178 | SIK1 | salt inducible kinase 1 [Source:HGNC Symbol;Acc:HGNC:11142] | -1.51 | 0.018725 |
| ENSG00000204442 | FAM155A | family with sequence similarity 155 member A [Source:HGNC Symbol;Acc:HGNC:33877] | -0.53 | 0.019183 |
| ENSG00000226686 | LINC01535 | long intergenic non-protein coding RNA 1535 [Source:HGNC Symbol;Acc:HGNC:51282] | -0.73 | 0.019183 |
| ENSG00000149260 | CAPN5 | calpain 5 [Source:HGNC Symbol;Acc:HGNC:1482] | 0.56 | 0.019286 |
| ENSG00000166349 | RAG1 | recombination activating 1 [Source:HGNC Symbol;Acc:HGNC:9831] | 0.90 | 0.019452 |
| ENSG00000116962 | NID1 | nidogen 1 [Source:HGNC Symbol;Acc:HGNC:7821] | 0.92 | 0.019592 |
| ENSG00000157064 | NMNAT2 | nicotinamide nucleotide adenylyltransferase 2 [Source:HGNC Symbol;Acc:HGNC:16789] | -1.74 | 0.019592 |
| ENSG00000175093 | SPSB4 | splA/ryanodine receptor domain and SOCS box containing 4 [Source:HGNC Symbol;Acc:HGNC:30630] | -0.54 | 0.019592 |
| ENSG00000075651 | PLD1 | phospholipase D1 [Source:HGNC Symbol;Acc:HGNC:9067] | -0.51 | 0.020376 |
| ENSG00000166886 | NAB2 | NGFI-A binding protein 2 [Source:HGNC Symbol;Acc:HGNC:7627] | 0.69 | 0.020391 |
| ENSG00000285860 | NA | novel transcript | -0.74 | 0.020585 |
| ENSG00000110427 | KIAA1549L | KIAA1549 like [Source:HGNC Symbol;Acc:HGNC:24836] | -0.75 | 0.021693 |
| ENSG00000205403 | CFI | complement factor I [Source:HGNC Symbol;Acc:HGNC:5394] | 0.80 | 0.021866 |
| ENSG00000215492 | HNRNPA1P7 | heterogeneous nuclear ribonucleoprotein A1 pseudogene 7 [Source:HGNC Symbol;Acc:HGNC:31015] | -0.55 | 0.021971 |
| ENSG00000104368 | PLAT | plasminogen activator, tissue type [Source:HGNC Symbol;Acc:HGNC:9051] | 0.55 | 0.022128 |
| ENSG00000273259 | NA | novel protein | 0.51 | 0.022297 |
| ENSG00000099282 | TSPAN15 | tetraspanin 15 [Source:HGNC Symbol;Acc:HGNC:23298] | 0.83 | 0.022441 |
| ENSG00000157335 | CLEC18C | C-type lectin domain family 18 member C [Source:HGNC Symbol;Acc:HGNC:28538] | 0.78 | 0.022627 |
| ENSG00000119408 | NEK6 | NIMA related kinase 6 [Source:HGNC Symbol;Acc:HGNC:7749] | 0.62 | 0.022807 |
| ENSG00000163808 | KIF15 | kinesin family member 15 [Source:HGNC Symbol;Acc:HGNC:17273] | -0.56 | 0.022862 |
| ENSG00000118515 | SGK1 | serum/glucocorticoid regulated kinase 1 [Source:HGNC Symbol;Acc:HGNC:10810] | -0.67 | 0.022985 |
| ENSG00000141750 | STAC2 | SH3 and cysteine rich domain 2 [Source:HGNC Symbol;Acc:HGNC:23990] | 0.62 | 0.023306 |
| ENSG00000244405 | ETV5 | ETS variant transcription factor 5 [Source:HGNC Symbol;Acc:HGNC:3494] | 0.81 | 0.023471 |
| ENSG00000167680 | SEMA6B | semaphorin 6B [Source:HGNC Symbol;Acc:HGNC:10739] | 0.52 | 0.023729 |
| ENSG00000117533 | VAMP4 | vesicle associated membrane protein 4 [Source:HGNC Symbol;Acc:HGNC:12645] | 0.51 | 0.023758 |
| ENSG00000135454 | B4GALNT1 | beta-1,4-N-acetyl-galactosaminyltransferase 1 [Source:HGNC Symbol;Acc:HGNC:4117] | 0.54 | 0.023758 |
| ENSG00000223658 | C1GALT1C1L | C1GALT1 specific chaperone 1 like [Source:HGNC Symbol;Acc:HGNC:51617] | 0.55 | 0.023771 |

| | | | | |
|---|---|---|---|---|
| ENSG00000277399 | GPR179 | G protein-coupled receptor 179 [Source:HGNC Symbol;Acc:HGNC:31371] | -1.05 | 0.023878 |
| ENSG00000095539 | SEMA4G | semaphorin 4G [Source:HGNC Symbol;Acc:HGNC:10735] | 0.59 | 0.024021 |
| ENSG00000100604 | CHGA | chromogranin A [Source:HGNC Symbol;Acc:HGNC:1929] | -0.75 | 0.024039 |
| ENSG00000236393 | NA | novel transcript | 0.67 | 0.024174 |
| ENSG00000232022 | FAAHP1 | fatty acid amide hydrolase pseudogene 1 [Source:HGNC Symbol;Acc:HGNC:50679] | 1.15 | 0.024503 |
| ENSG00000223711 | NA | novel transcript | 1.05 | 0.024878 |
| ENSG00000113100 | CDH9 | cadherin 9 [Source:HGNC Symbol;Acc:HGNC:1768] | -0.67 | 0.024892 |
| ENSG00000188573 | FBLL1 | fibrillarin like 1 [Source:HGNC Symbol;Acc:HGNC:35458] | -0.65 | 0.025001 |
| ENSG00000111186 | WNT5B | Wnt family member 5B [Source:HGNC Symbol;Acc:HGNC:16265] | 1.10 | 0.025268 |
| ENSG00000169862 | CTNND2 | catenin delta 2 [Source:HGNC Symbol;Acc:HGNC:2516] | 1.93 | 0.026113 |
| ENSG00000219665 | ZNF433-AS1 | ZNF433 and ZNF878 antisense RNA 1 [Source:HGNC Symbol;Acc:HGNC:53776] | 0.52 | 0.026346 |
| ENSG00000128578 | STRIP2 | striatin interacting protein 2 [Source:HGNC Symbol;Acc:HGNC:22209] | 0.68 | 0.026816 |
| ENSG00000160712 | IL6R | interleukin 6 receptor [Source:HGNC Symbol;Acc:HGNC:6019] | 2.25 | 0.026816 |
| ENSG00000169570 | DTWD2 | DTW domain containing 2 [Source:HGNC Symbol;Acc:HGNC:19334] | -0.60 | 0.026933 |
| ENSG00000059573 | ALDH18A1 | aldehyde dehydrogenase 18 family member A1 [Source:HGNC Symbol;Acc:HGNC:9722] | -0.50 | 0.027112 |
| ENSG00000236753 | MKLN1-AS | MKLN1 antisense RNA [Source:HGNC Symbol;Acc:HGNC:40374] | 0.59 | 0.027136 |
| ENSG00000149403 | GRIK4 | glutamate ionotropic receptor kainate type subunit 4 [Source:HGNC Symbol;Acc:HGNC:4582] | 1.10 | 0.027311 |
| ENSG00000083807 | SLC27A5 | solute carrier family 27 member 5 [Source:HGNC Symbol;Acc:HGNC:10999] | -0.61 | 0.027541 |
| ENSG00000134013 | LOXL2 | lysyl oxidase like 2 [Source:HGNC Symbol;Acc:HGNC:6666] | 0.62 | 0.027564 |
| ENSG00000272256 | NA | novel transcript, antisense to TMEM66 | 1.29 | 0.027564 |
| ENSG00000146072 | TNFRSF21 | TNF receptor superfamily member 21 [Source:HGNC Symbol;Acc:HGNC:13469] | 1.28 | 0.027773 |
| ENSG00000251623 | NA | centrosomal protein 192 (CEP192), pseudogene | 0.90 | 0.028133 |
| ENSG00000165434 | PGM2L1 | phosphoglucomutase 2 like 1 [Source:HGNC Symbol;Acc:HGNC:20898] | 0.60 | 0.028257 |
| ENSG00000180096 | SEPTIN1 | septin 1 [Source:HGNC Symbol;Acc:HGNC:2879] | -0.56 | 0.028506 |
| ENSG00000120254 | MTHFD1L | methylenetetrahydrofolate dehydrogenase (NADP+ dependent) 1 like [Source:HGNC Symbol;Acc:HGNC:21055] | -0.66 | 0.028593 |
| ENSG00000074590 | NUAK1 | NUAK family kinase 1 [Source:HGNC Symbol;Acc:HGNC:14311] | 0.67 | 0.028846 |
| ENSG00000128641 | MYO1B | myosin IB [Source:HGNC Symbol;Acc:HGNC:7596] | 1.83 | 0.028940 |
| ENSG00000285799 | NA | MHC class I polypeptide-related sequence F pseudogene | -2.48 | 0.029484 |
| ENSG00000120693 | SMAD9 | SMAD family member 9 [Source:HGNC Symbol;Acc:HGNC:6774] | 0.55 | 0.029845 |
| ENSG00000135144 | DTX1 | deltex E3 ubiquitin ligase 1 [Source:HGNC Symbol;Acc:HGNC:3060] | 0.80 | 0.029873 |
| ENSG00000117226 | GBP3 | guanylate binding protein 3 [Source:HGNC Symbol;Acc:HGNC:4184] | 2.29 | 0.030196 |
| ENSG00000176697 | BDNF | brain derived neurotrophic factor [Source:HGNC Symbol;Acc:HGNC:1033] | 0.69 | 0.030463 |
| ENSG00000006025 | OSBPL7 | oxysterol binding protein like 7 [Source:HGNC Symbol;Acc:HGNC:16387] | 0.51 | 0.030605 |
| ENSG00000078814 | MYH7B | myosin heavy chain 7B [Source:HGNC Symbol;Acc:HGNC:15906] | 0.52 | 0.030954 |
| ENSG00000105639 | JAK3 | Janus kinase 3 [Source:HGNC Symbol;Acc:HGNC:6193] | 0.70 | 0.031007 |
| ENSG00000175785 | PRIMA1 | proline rich membrane anchor 1 [Source:HGNC Symbol;Acc:HGNC:18319] | 0.56 | 0.031224 |
| ENSG00000230257 | NFE4 | nuclear factor, erythroid 4 [Source:HGNC Symbol;Acc:HGNC:29902] | 0.58 | 0.031327 |
| ENSG00000058404 | CAMK2B | calcium/calmodulin dependent protein kinase II beta [Source:HGNC Symbol;Acc:HGNC:1461] | 0.52 | 0.031778 |
| ENSG00000135709 | KIAA0513 | KIAA0513 [Source:HGNC Symbol;Acc:HGNC:29058] | 0.57 | 0.031879 |
| ENSG00000272047 | GTF2H5 | general transcription factor IIH subunit 5 [Source:HGNC Symbol;Acc:HGNC:21157] | -0.51 | 0.031879 |
| ENSG00000162692 | VCAM1 | vascular cell adhesion molecule 1 [Source:HGNC Symbol;Acc:HGNC:12663] | 0.51 | 0.031993 |
| ENSG00000163874 | ZC3H12A | zinc finger CCCH-type containing 12A [Source:HGNC Symbol;Acc:HGNC:26259] | 0.74 | 0.032661 |
| ENSG00000188833 | ENTPD8 | ectonucleoside triphosphate diphosphohydrolase 8 [Source:HGNC Symbol;Acc:HGNC:24860] | 0.68 | 0.032661 |
| ENSG00000183117 | CSMD1 | CUB and Sushi multiple domains 1 [Source:HGNC Symbol;Acc:HGNC:14026] | -0.61 | 0.032672 |
| ENSG00000174721 | FGFBP3 | fibroblast growth factor binding protein 3 [Source:HGNC Symbol;Acc:HGNC:23428] | -0.54 | 0.032950 |
| ENSG00000131242 | RAB11FIP4 | RAB11 family interacting protein 4 [Source:HGNC Symbol;Acc:HGNC:30267] | 0.80 | 0.033169 |
| ENSG00000168542 | COL3A1 | collagen type III alpha 1 chain [Source:HGNC Symbol;Acc:HGNC:2201] | 1.53 | 0.033669 |
| ENSG00000196878 | LAMB3 | laminin subunit beta 3 [Source:HGNC Symbol;Acc:HGNC:6490] | 2.20 | 0.033774 |

| | | | | |
|---|---|---|---|---|
| ENSG00000277957 | SENP3-EIF4A1 | SENP3-EIF4A1 readthrough (NMD candidate) [Source:HGNC Symbol;Acc:HGNC:49182] | 0.83 | 0.033774 |
| ENSG00000167074 | TEF | TEF transcription factor, PAR bZIP family member [Source:HGNC Symbol;Acc:HGNC:11722] | 0.72 | 0.034002 |
| ENSG00000258733 | LINC02328 | long intergenic non-protein coding RNA 2328 [Source:HGNC Symbol;Acc:HGNC:53248] | 0.88 | 0.034144 |
| ENSG00000196843 | ARID5A | AT-rich interaction domain 5A [Source:HGNC Symbol;Acc:HGNC:17361] | 1.06 | 0.034544 |
| ENSG00000213080 | NA | spermine synthase (SMS) pseudogene | -0.50 | 0.034544 |
| ENSG00000133392 | MYH11 | myosin heavy chain 11 [Source:HGNC Symbol;Acc:HGNC:7569] | 1.05 | 0.034757 |
| ENSG00000278916 | CEP83-DT | CEP83 divergent transcript [Source:HGNC Symbol;Acc:HGNC:27055] | -0.65 | 0.035103 |
| ENSG00000111215 | PRR4 | proline rich 4 [Source:HGNC Symbol;Acc:HGNC:18020] | 0.66 | 0.035729 |
| ENSG00000248498 | ASNSP1 | asparagine synthetase pseudogene 1 [Source:HGNC Symbol;Acc:HGNC:754] | -0.66 | 0.035975 |
| ENSG00000254995 | STX16-NPEPL1 | STX16-NPEPL1 readthrough (NMD candidate) [Source:HGNC Symbol;Acc:HGNC:41993] | 1.14 | 0.036333 |
| ENSG00000203859 | HSD3B2 | hydroxy-delta-5-steroid dehydrogenase, 3 beta- and steroid delta-isomerase 2 [Source:HGNC Symbol;Acc:HGNC:5218] | 1.45 | 0.036352 |
| ENSG00000072682 | P4HA2 | prolyl 4-hydroxylase subunit alpha 2 [Source:HGNC Symbol;Acc:HGNC:8547] | 0.67 | 0.036589 |
| ENSG00000115107 | STEAP3 | STEAP3 metalloreductase [Source:HGNC Symbol;Acc:HGNC:24592] | 0.88 | 0.036589 |
| ENSG00000151689 | INPP1 | inositol polyphosphate-1-phosphatase [Source:HGNC Symbol;Acc:HGNC:6071] | 0.54 | 0.036589 |
| ENSG00000157570 | TSPAN18 | tetraspanin 18 [Source:HGNC Symbol;Acc:HGNC:20660] | 0.64 | 0.036589 |
| ENSG00000173221 | GLRX | glutaredoxin [Source:HGNC Symbol;Acc:HGNC:4330] | 0.59 | 0.037212 |
| ENSG00000163531 | NFASC | neurofascin [Source:HGNC Symbol;Acc:HGNC:29866] | 0.58 | 0.037215 |
| ENSG00000248869 | LINC02511 | long intergenic non-protein coding RNA 2511 [Source:HGNC Symbol;Acc:HGNC:53500] | 1.17 | 0.037296 |
| ENSG00000053747 | LAMA3 | laminin subunit alpha 3 [Source:HGNC Symbol;Acc:HGNC:6483] | -0.72 | 0.037986 |
| ENSG00000286614 | NA | novel pseudogene | 2.16 | 0.038555 |
| ENSG00000136205 | TNS3 | tensin 3 [Source:HGNC Symbol;Acc:HGNC:21616] | 1.02 | 0.038884 |
| ENSG00000133794 | ARNTL | aryl hydrocarbon receptor nuclear translocator like [Source:HGNC Symbol;Acc:HGNC:701] | -0.57 | 0.039016 |
| ENSG00000179071 | CCDC89 | coiled-coil domain containing 89 [Source:HGNC Symbol;Acc:HGNC:26762] | -0.60 | 0.039019 |
| ENSG00000259985 | NA | novel transcript, antisense to B4GALT6 | -0.70 | 0.039239 |
| ENSG00000262769 | NA | novel transcript, antisense to SLC47A1 | 0.69 | 0.039478 |
| ENSG00000139914 | FITM1 | fat storage inducing transmembrane protein 1 [Source:HGNC Symbol;Acc:HGNC:33714] | 0.53 | 0.039949 |
| ENSG00000153976 | HS3ST3A1 | heparan sulfate-glucosamine 3-sulfotransferase 3A1 [Source:HGNC Symbol;Acc:HGNC:5196] | -1.38 | 0.040235 |
| ENSG00000151006 | PRSS53 | serine protease 53 [Source:HGNC Symbol;Acc:HGNC:34407] | 1.29 | 0.040359 |
| ENSG00000036448 | MYOM2 | myomesin 2 [Source:HGNC Symbol;Acc:HGNC:7614] | -0.82 | 0.040486 |
| ENSG00000143036 | SLC44A3 | solute carrier family 44 member 3 [Source:HGNC Symbol;Acc:HGNC:28689] | 0.68 | 0.040718 |
| ENSG00000109819 | PPARGC1A | PPARG coactivator 1 alpha [Source:HGNC Symbol;Acc:HGNC:9237] | -0.52 | 0.040875 |
| ENSG00000262155 | LINC02175 | long intergenic non-protein coding RNA 2175 [Source:HGNC Symbol;Acc:HGNC:27550] | -0.55 | 0.041024 |
| ENSG00000100092 | SH3BP1 | SH3 domain binding protein 1 [Source:HGNC Symbol;Acc:HGNC:10824] | 0.69 | 0.041160 |
| ENSG00000123213 | NLN | neurolysin [Source:HGNC Symbol;Acc:HGNC:16058] | -0.51 | 0.041906 |
| ENSG00000175866 | BAIAP2 | BAR/IMD domain containing adaptor protein 2 [Source:HGNC Symbol;Acc:HGNC:947] | 0.51 | 0.041999 |
| ENSG00000106633 | GCK | glucokinase [Source:HGNC Symbol;Acc:HGNC:4195] | 0.60 | 0.042050 |
| ENSG00000158258 | CLSTN2 | calsyntenin 2 [Source:HGNC Symbol;Acc:HGNC:17448] | 0.88 | 0.042131 |
| ENSG00000280434 | NA | novel transcript, sense overlapping PARVB | 0.76 | 0.043436 |
| ENSG00000260552 | NA | novel transcript, antisense to MOCOS | -0.76 | 0.043540 |
| ENSG00000027075 | PRKCH | protein kinase C eta [Source:HGNC Symbol;Acc:HGNC:9403] | 0.73 | 0.044316 |
| ENSG00000064309 | CDON | cell adhesion associated, oncogene regulated [Source:HGNC Symbol;Acc:HGNC:17104] | -0.57 | 0.044397 |
| ENSG00000088836 | SLC4A11 | solute carrier family 4 member 11 [Source:HGNC Symbol;Acc:HGNC:16438] | 0.82 | 0.044682 |
| ENSG00000167580 | AQP2 | aquaporin 2 [Source:HGNC Symbol;Acc:HGNC:634] | 0.59 | 0.044682 |
| ENSG00000175426 | PCSK1 | proprotein convertase subtilisin/kexin type 1 [Source:HGNC Symbol;Acc:HGNC:8743] | 1.07 | 0.044724 |
| ENSG00000142102 | PGGHG | protein-glucosylgalactosylhydroxylysine glucosidase [Source:HGNC Symbol;Acc:HGNC:26210] | 0.77 | 0.045325 |
| ENSG00000204758 | NA | novel transcript, antisense to RPL26L1 | -1.04 | 0.045531 |
| ENSG00000125968 | ID1 | inhibitor of DNA binding 1, HLH protein [Source:HGNC Symbol;Acc:HGNC:5360] | 0.87 | 0.045649 |
| ENSG00000137494 | ANKRD42 | ankyrin repeat domain 42 [Source:HGNC Symbol;Acc:HGNC:26752] | 0.58 | 0.045779 |

| | | | | |
|---|---|---|---|---|
| ENSG00000080224 | EPHA6 | EPH receptor A6 [Source:HGNC Symbol;Acc:HGNC:19296] | 0.88 | 0.046723 |
| ENSG00000074527 | NTN4 | netrin 4 [Source:HGNC Symbol;Acc:HGNC:13658] | 0.59 | 0.046897 |
| ENSG00000287409 | NA | novel transcript | 0.76 | 0.047393 |
| ENSG00000270885 | RASL10B | RAS like family 10 member B [Source:HGNC Symbol;Acc:HGNC:30295] | -0.82 | 0.047565 |
| ENSG00000267302 | RNFT1-DT | RNFT1 divergent transcript [Source:HGNC Symbol;Acc:HGNC:51346] | 1.09 | 0.047657 |
| ENSG00000173267 | SNCG | synuclein gamma [Source:HGNC Symbol;Acc:HGNC:11141] | -0.51 | 0.048201 |
| ENSG00000089127 | OAS1 | 2'-5'-oligoadenylate synthetase 1 [Source:HGNC Symbol;Acc:HGNC:8086] | 1.00 | 0.048366 |
| ENSG00000168528 | SERINC2 | serine incorporator 2 [Source:HGNC Symbol;Acc:HGNC:23231] | 0.61 | 0.048519 |
| ENSG00000053438 | NNAT | neuronatin [Source:HGNC Symbol;Acc:HGNC:7860] | 1.20 | 0.048535 |
| ENSG00000181625 | SLX1B | SLX1 homolog B, structure-specific endonuclease subunit [Source:HGNC Symbol;Acc:HGNC:28748] | 2.15 | 0.048821 |
| ENSG00000266916 | ZNF793-AS1 | ZNF793 antisense RNA 1 (head to head) [Source:HGNC Symbol;Acc:HGNC:51303] | 0.72 | 0.049017 |
| ENSG00000277801 | NA | novel transcript | -0.63 | 0.049930 |
| ENSG00000181392 | SYNE4 | spectrin repeat containing nuclear envelope family member 4 [Source:HGNC Symbol;Acc:HGNC:26703] | 0.83 | 0.049953 |
