## Supplemental Table 9 for "Modulation of calcium signaling on demand to decipher the molecular mechanisms of primary aldosteronism"

**Supplementary Table 9. List of significantly enriched pathways in H295R_S2 cells expressing the _7-5HT3 receptor in response to 24h treatment with 10-8M AngII**

| Pathway | Fold Enrichment | PValue |
|---|---|---|
| GO:0098609~cell-cell adhesion | 5.44 | 1.37E-11 |
| GO:0007155~cell adhesion | 2.61 | 4.79E-07 |
| GO:0030198~extracellular matrix organization | 3.73 | 1.43E-05 |
| hsa04151:PI3K-Akt signaling pathway | 2.47 | 2.15E-05 |
| hsa04512:ECM-receptor interaction | 4.56 | 2.31E-05 |
| hsa05200:Pathways in cancer | 2.06 | 7.08E-05 |
| GO:0007160~cell-matrix adhesion | 4.49 | 7.44E-05 |
| GO:0007420~brain development | 2.97 | 8.21E-05 |
| GO:0071356~cellular response to tumor necrosis factor | 3.87 | 1.40E-04 |
| GO:0046718~viral entry into host cell | 4.40 | 2.00E-04 |
| GO:0050892~intestinal absorption | 14.57 | 2.91E-04 |
| GO:0060038~cardiac muscle cell proliferation | 9.79 | 3.04E-04 |
| GO:0030199~collagen fibril organization | 4.83 | 5.32E-04 |
| hsa04510:Focal adhesion | 2.62 | 7.54E-04 |
| GO:0048812~neuron projection morphogenesis | 5.18 | 8.51E-04 |
| GO:0006006~glucose metabolic process | 5.10 | 9.36E-04 |
| GO:0071347~cellular response to interleukin-1 | 4.37 | 0.00104 |
| hsa04925:Aldosterone synthesis and secretion | 3.51 | 0.00109 |
| hsa04934:Cushing syndrome | 2.82 | 0.00133 |
| GO:0070633~transepithelial transport | 16.31 | 0.00154 |
| hsa04610:Complement and coagulation cascades | 3.63 | 0.00160 |
| GO:0032355~response to estradiol | 4.03 | 0.00174 |
| GO:0015914~phospholipid transport | 6.27 | 0.00249 |
| GO:0048661~positive regulation of smooth muscle cell proliferation | 4.92 | 0.00287 |
| GO:0071346~cellular response to interferon-gamma | 3.71 | 0.00297 |
| GO:0070509~calcium ion import | 8.16 | 0.00298 |
| GO:0033627~cell adhesion mediated by integrin | 5.97 | 0.00311 |
| GO:0035910~ascending aorta morphogenesis | 30.59 | 0.00347 |
| GO:0008284~positive regulation of cell proliferation | 1.88 | 0.00383 |
| GO:0009410~response to xenobiotic stimulus | 2.42 | 0.00394 |
| GO:0030324~lung development | 3.93 | 0.00421 |
| GO:0001525~angiogenesis | 2.40 | 0.00424 |
| GO:0032963~collagen metabolic process | 11.65 | 0.00433 |
| GO:0034383~low-density lipoprotein particle clearance | 11.65 | 0.00433 |

| | | |
|---|---|---|
| GO:0060317~cardiac epithelial to mesenchymal transition | 11.65 | 0.00433 |
| hsa04927:Cortisol synthesis and secretion | 3.84 | 0.00445 |
| GO:0030335~positive regulation of cell migration | 2.34 | 0.00536 |
| hsa04974:Protein digestion and absorption | 3.03 | 0.00548 |
| GO:0042102~positive regulation of T cell proliferation | 4.26 | 0.00589 |
| GO:0007229~integrin-mediated signaling pathway | 3.31 | 0.00595 |
| GO:0007507~heart development | 2.50 | 0.00615 |
| hsa04810:Regulation of actin cytoskeleton | 2.18 | 0.00642 |
| GO:0002486~antigen processing and presentation of endogenous peptide antigen via MHC class I via ER pathway, TAP-independent | 10.20 | 0.00643 |
| GO:0002021~response to dietary excess | 10.20 | 0.00643 |
| GO:0035987~endodermal cell differentiation | 6.58 | 0.00660 |
| GO:0002476~antigen processing and presentation of endogenous peptide antigen via MHC class Ib | 9.60 | 0.00767 |
| GO:0016486~peptide hormone processing | 9.60 | 0.00767 |
| GO:0070528~protein kinase C signaling | 9.60 | 0.00767 |
| hsa05410:Hypertrophic cardiomyopathy | 3.12 | 0.00782 |
| GO:0010976~positive regulation of neuron projection development | 3.14 | 0.00810 |
| GO:0016477~cell migration | 2.22 | 0.00813 |
| GO:2000427~positive regulation of apoptotic cell clearance | 20.39 | 0.00839 |
| GO:0035556~intracellular signal transduction | 1.87 | 0.00965 |
| GO:0071526~semaphorin-plexin signaling pathway | 5.83 | 0.01017 |
| GO:0001764~neuron migration | 3.01 | 0.01030 |
| GO:0048167~regulation of synaptic plasticity | 4.45 | 0.01094 |
| hsa05412:Arrhythmogenic right ventricular cardiomyopathy | 3.25 | 0.01115 |
| GO:0010811~positive regulation of cell-substrate adhesion | 5.66 | 0.01122 |
| hsa05414:Dilated cardiomyopathy | 2.93 | 0.01137 |
| GO:0007267~cell-cell signaling | 2.31 | 0.01138 |
| GO:0002062~chondrocyte differentiation | 4.37 | 0.01178 |
| GO:0060412~ventricular septum morphogenesis | 5.51 | 0.01235 |
| GO:0022900~electron transport chain | 4.29 | 0.01266 |
| GO:0007584~response to nutrient | 4.22 | 0.01358 |
| GO:0043651~linoleic acid metabolic process | 7.77 | 0.01395 |
| GO:0048513~animal organ development | 5.23 | 0.01480 |
| GO:1901687~glutathione derivative biosynthetic process | 15.29 | 0.01517 |
| GO:0070307~lens fiber cell development | 15.29 | 0.01517 |
| GO:0016125~sterol metabolic process | 7.42 | 0.01586 |

| | | |
|---|---|---|
| GO:0045737~positive regulation of cyclin-dependent protein serine/threonine kinase activity | 5.10 | 0.01614 |
| GO:0050767~regulation of neurogenesis | 5.10 | 0.01614 |
| GO:0007405~neuroblast proliferation | 4.97 | 0.01754 |
| GO:0001666~response to hypoxia | 2.53 | 0.01777 |
| hsa05225:Hepatocellular carcinoma | 2.23 | 0.01848 |
| GO:0006954~inflammatory response | 1.81 | 0.01862 |
| GO:0007411~axon guidance | 2.36 | 0.01882 |
| GO:0060048~cardiac muscle contraction | 4.86 | 0.01903 |
| GO:0006865~amino acid transport | 4.86 | 0.01903 |
| GO:0044320~cellular response to leptin stimulus | 13.60 | 0.01919 |
| GO:0007613~memory | 3.28 | 0.01988 |
| GO:1904754~positive regulation of vascular associated smooth muscle cell migration | 6.80 | 0.02011 |
| GO:0006883~cellular sodium ion homeostasis | 6.80 | 0.02011 |
| GO:0042098~T cell proliferation | 4.74 | 0.02058 |
| GO:0035249~synaptic transmission, glutamatergic | 4.63 | 0.02222 |
| GO:0055013~cardiac muscle cell development | 6.53 | 0.02245 |
| GO:0002548~monocyte chemotaxis | 4.53 | 0.02393 |
| GO:0071466~cellular response to xenobiotic stimulus | 3.65 | 0.02399 |
| GO:0007596~blood coagulation | 3.14 | 0.02421 |
| GO:0050731~positive regulation of peptidyl-tyrosine phosphorylation | 3.14 | 0.02421 |
| GO:0042127~regulation of cell proliferation | 2.39 | 0.02512 |
| GO:0030168~platelet activation | 3.60 | 0.02539 |
| GO:0007568~aging | 2.53 | 0.02641 |
| GO:0042593~glucose homeostasis | 2.74 | 0.02708 |
| hsa05222:Small cell lung cancer | 2.72 | 0.02724 |
| GO:0010628~positive regulation of gene expression | 1.67 | 0.02762 |
| GO:0032956~regulation of actin cytoskeleton organization | 3.04 | 0.02785 |
| GO:1990535~neuron projection maintenance | 11.12 | 0.02839 |
| GO:0071492~cellular response to UV-A | 11.12 | 0.02839 |
| GO:0034097~response to cytokine | 4.25 | 0.02953 |
| GO:0006749~glutathione metabolic process | 4.25 | 0.02953 |
| GO:0048839~inner ear development | 4.25 | 0.02953 |
| GO:0009611~response to wounding | 3.45 | 0.02991 |
| hsa04115:p53 signaling pathway | 2.95 | 0.03025 |
| GO:0033344~cholesterol efflux | 5.83 | 0.03031 |

| | | |
|---|---|---|
| GO:0031016~pancreas development | 5.83 | 0.03031 |
| GO:0010468~regulation of gene expression | 2.00 | 0.03041 |
| hsa05205:Proteoglycans in cancer | 1.98 | 0.03101 |
| GO:0050679~positive regulation of epithelial cell proliferation | 3.40 | 0.03152 |
| GO:0032526~response to retinoic acid | 4.16 | 0.03155 |
| hsa04010:MAPK signaling pathway | 1.76 | 0.03191 |
| GO:0007520~myoblast fusion | 5.63 | 0.03321 |
| GO:0034113~heterotypic cell-cell adhesion | 5.63 | 0.03321 |
| GO:0097264~self proteolysis | 10.20 | 0.03352 |
| GO:0048711~positive regulation of astrocyte differentiation | 10.20 | 0.03352 |
| GO:0048251~elastic fiber assembly | 10.20 | 0.03352 |
| GO:0030154~cell differentiation | 1.54 | 0.03368 |
| GO:0007165~signal transduction | 1.37 | 0.03382 |
| GO:0030308~negative regulation of cell growth | 2.59 | 0.03539 |
| hsa05165:Human papillomavirus infection | 1.70 | 0.03542 |
| GO:0099505~regulation of presynaptic membrane potential | 5.44 | 0.03625 |
| GO:0007179~transforming growth factor beta receptor signaling pathway | 2.83 | 0.03768 |
| GO:0008344~adult locomotory behavior | 3.92 | 0.03810 |
| GO:0098915~membrane repolarization during ventricular cardiac muscle cell action potential | 9.41 | 0.03899 |
| GO:0032233~positive regulation of actin filament bundle assembly | 9.41 | 0.03899 |
| GO:0016322~neuron remodeling | 9.41 | 0.03899 |
| GO:0060070~canonical Wnt signaling pathway | 2.80 | 0.03925 |
| GO:0060045~positive regulation of cardiac muscle cell proliferation | 5.26 | 0.03942 |
| GO:0090102~cochlea development | 5.26 | 0.03942 |
| GO:0030522~intracellular receptor signaling pathway | 5.26 | 0.03942 |
| hsa04630:JAK-STAT signaling pathway | 2.07 | 0.03953 |
| GO:0006897~endocytosis | 2.08 | 0.04035 |
| GO:0021987~cerebral cortex development | 3.18 | 0.04037 |
| GO:0008217~regulation of blood pressure | 3.18 | 0.04037 |
| hsa04978:Mineral absorption | 3.12 | 0.04183 |
| hsa05202:Transcriptional misregulation in cancer | 1.94 | 0.04460 |
| GO:0046688~response to copper ion | 8.74 | 0.04476 |
| GO:0030595~leukocyte chemotaxis | 8.74 | 0.04476 |
| GO:0030195~negative regulation of blood coagulation | 8.74 | 0.04476 |
| GO:0071805~potassium ion transmembrane transport | 2.45 | 0.04524 |

| | | |
|---|---|---|
| GO:0000079~regulation of cyclin-dependent protein serine/threonine kinase activity | 3.71 | 0.04538 |
| GO:0043410~positive regulation of MAPK cascade | 2.27 | 0.04611 |
| GO:2000379~positive regulation of reactive oxygen species metabolic process | 4.94 | 0.04619 |
| GO:0034765~regulation of ion transmembrane transport | 2.43 | 0.04678 |
| GO:0010951~negative regulation of endopeptidase activity | 3.64 | 0.04796 |
| GO:2000866~positive regulation of estradiol secretion | 40.79 | 0.04834 |
| GO:0061102~stomach neuroendocrine cell differentiation | 40.79 | 0.04834 |
| GO:0003431~growth plate cartilage chondrocyte development | 40.79 | 0.04834 |
| GO:0006705~mineralocorticoid biosynthetic process | 40.79 | 0.04834 |
