## Supplemental Table 10 for "Modulation of calcium signaling on demand to decipher the molecular mechanisms of primary aldosteronism"

**Supplementary Table 10. List of differentially expressed genes in H295R_S2 cells expressing the α7-5HT3 receptor in response to 24h treatment with 12 mM K+**

| Ensembl Gene ID | Gene Symbol | Gene Name | Log2 Fold Chang | Padj |
|---|---|---|---|---|
| ENSG00000197635 | DPP4 | dipeptidyl peptidase 4 [Source:HGNC Symbol;Acc:HGNC:3009] | 1.77 | 4.11E-44 |
| ENSG00000101605 | MYOM1 | myomesin 1 [Source:HGNC Symbol;Acc:HGNC:7613] | 2.20 | 2.26E-29 |
| ENSG00000143153 | ATP1B1 | ATPase Na+/K+ transporting subunit beta 1 [Source:HGNC Symbol;Acc:HGNC:804] | 1.26 | 2.23E-26 |
| ENSG00000196586 | MYO6 | myosin VI [Source:HGNC Symbol;Acc:HGNC:7605] | 1.44 | 4.80E-24 |
| ENSG00000138759 | FRAS1 | Fraser extracellular matrix complex subunit 1 [Source:HGNC Symbol;Acc:HGNC:19185] | 1.78 | 3.45E-22 |
| ENSG00000155827 | RNF20 | ring finger protein 20 [Source:HGNC Symbol;Acc:HGNC:10062] | 1.14 | 3.38E-21 |
| ENSG00000076716 | GPC4 | glypican 4 [Source:HGNC Symbol;Acc:HGNC:4452] | 1.32 | 5.83E-21 |
| ENSG00000196352 | CD55 | CD55 molecule (Cromer blood group) [Source:HGNC Symbol;Acc:HGNC:2665] | 1.19 | 5.00E-20 |
| ENSG00000065183 | WDR3 | WD repeat domain 3 [Source:HGNC Symbol;Acc:HGNC:12755] | -0.85 | 5.07E-20 |
| ENSG00000150893 | FREM2 | FRAS1 related extracellular matrix 2 [Source:HGNC Symbol;Acc:HGNC:25396] | 1.40 | 2.97E-19 |
| ENSG00000087250 | MT3 | metallothionein 3 [Source:HGNC Symbol;Acc:HGNC:7408] | 1.00 | 3.77E-19 |
| ENSG00000111424 | VDR | vitamin D receptor [Source:HGNC Symbol;Acc:HGNC:12679] | 1.37 | 3.77E-19 |
| ENSG00000154217 | PITPNC1 | phosphatidylinositol transfer protein cytoplasmic 1 [Source:HGNC Symbol;Acc:HGNC:21045 | 1.58 | 2.60E-17 |
| ENSG00000077943 | ITGA8 | integrin subunit alpha 8 [Source:HGNC Symbol;Acc:HGNC:6144] | 1.56 | 4.18E-17 |
| ENSG00000081181 | ARG2 | arginase 2 [Source:HGNC Symbol;Acc:HGNC:664] | 1.28 | 3.17E-16 |
| ENSG00000119698 | PPP4R4 | protein phosphatase 4 regulatory subunit 4 [Source:HGNC Symbol;Acc:HGNC:23788] | 1.65 | 4.38E-16 |
| ENSG00000074370 | ATP2A3 | ATPase sarcoplasmic/endoplasmic reticulum Ca2+ transporting 3 [Source:HGNC Symbol;Ac | 1.22 | 5.57E-16 |
| ENSG00000101230 | ISM1 | isthmin 1 [Source:HGNC Symbol;Acc:HGNC:16213] | -1.19 | 5.57E-16 |
| ENSG00000113231 | PDE8B | phosphodiesterase 8B [Source:HGNC Symbol;Acc:HGNC:8794] | 1.64 | 5.57E-16 |
| ENSG00000115758 | ODC1 | ornithine decarboxylase 1 [Source:HGNC Symbol;Acc:HGNC:8109] | -0.78 | 5.57E-16 |
| ENSG00000176697 | BDNF | brain derived neurotrophic factor [Source:HGNC Symbol;Acc:HGNC:1033] | 1.90 | 1.08E-15 |
| ENSG00000184349 | EFNA5 | ephrin A5 [Source:HGNC Symbol;Acc:HGNC:3225] | 1.97 | 1.40E-15 |
| ENSG00000102172 | SMS | spermine synthase [Source:HGNC Symbol;Acc:HGNC:11123] | -0.67 | 1.64E-15 |
| ENSG00000109689 | STIM2 | stromal interaction molecule 2 [Source:HGNC Symbol;Acc:HGNC:19205] | 1.27 | 6.62E-15 |
| ENSG00000165092 | ALDH1A1 | aldehyde dehydrogenase 1 family member A1 [Source:HGNC Symbol;Acc:HGNC:402] | -0.72 | 7.79E-15 |
| ENSG00000116661 | FBXO2 | F-box protein 2 [Source:HGNC Symbol;Acc:HGNC:13581] | 1.60 | 1.44E-14 |
| ENSG00000139998 | RAB15 | RAB15, member RAS oncogene family [Source:HGNC Symbol;Acc:HGNC:20150] | 1.01 | 4.03E-14 |
| ENSG00000244405 | ETV5 | ETS variant transcription factor 5 [Source:HGNC Symbol;Acc:HGNC:3494] | 2.11 | 4.51E-14 |
| ENSG00000172915 | NBEA | neurobeachin [Source:HGNC Symbol;Acc:HGNC:7648] | 0.58 | 9.06E-14 |
| ENSG00000134508 | CABLES1 | Cdk5 and Abl enzyme substrate 1 [Source:HGNC Symbol;Acc:HGNC:25097] | 1.87 | 1.32E-13 |
| ENSG00000106025 | TSPAN12 | tetraspanin 12 [Source:HGNC Symbol;Acc:HGNC:21641] | 1.00 | 1.48E-13 |
| ENSG00000128050 | PAICS | phosphoribosylaminoimidazole carboxylase and phosphoribosylaminoimidazolesuccinocar | -0.57 | 1.78E-13 |
| ENSG00000188766 | SPRED3 | sprouty related EVH1 domain containing 3 [Source:HGNC Symbol;Acc:HGNC:31041] | 1.86 | 2.50E-13 |
| ENSG00000113389 | NPR3 | natriuretic peptide receptor 3 [Source:HGNC Symbol;Acc:HGNC:7945] | 1.42 | 2.79E-13 |
| ENSG00000101445 | PPP1R16B | protein phosphatase 1 regulatory subunit 16B [Source:HGNC Symbol;Acc:HGNC:15850] | 0.87 | 3.56E-13 |
| ENSG00000182054 | IDH2 | isocitrate dehydrogenase (NADP(+)) 2 [Source:HGNC Symbol;Acc:HGNC:5383] | 0.99 | 3.56E-13 |
| ENSG00000187231 | SESTD1 | SEC14 and spectrin domain containing 1 [Source:HGNC Symbol;Acc:HGNC:18379] | 0.74 | 3.99E-13 |

| | | | | |
|---|---|---|---|---|
| ENSG00000091428 | RAPGEF4 | Rap guanine nucleotide exchange factor 4 [Source:HGNC Symbol;Acc:HGNC:16626] | 2.06 | 6.90E-13 |
| ENSG00000122786 | CALD1 | caldesmon 1 [Source:HGNC Symbol;Acc:HGNC:1441] | 1.56 | 7.57E-13 |
| ENSG00000188517 | COL25A1 | collagen type XXV alpha 1 chain [Source:HGNC Symbol;Acc:HGNC:18603] | 0.91 | 1.81E-12 |
| ENSG00000157851 | DPYSL5 | dihydropyrimidinase like 5 [Source:HGNC Symbol;Acc:HGNC:20637] | 0.53 | 2.12E-12 |
| ENSG00000153162 | BMP6 | bone morphogenetic protein 6 [Source:HGNC Symbol;Acc:HGNC:1073] | 1.83 | 5.63E-12 |
| ENSG00000151835 | SACS | sacsin molecular chaperone [Source:HGNC Symbol;Acc:HGNC:10519] | 0.75 | 5.95E-12 |
| ENSG00000006468 | ETV1 | ETS variant transcription factor 1 [Source:HGNC Symbol;Acc:HGNC:3490] | 0.85 | 7.80E-12 |
| ENSG00000139973 | SYT16 | synaptotagmin 16 [Source:HGNC Symbol;Acc:HGNC:23142] | 2.40 | 8.95E-12 |
| ENSG00000179142 | CYP11B2 | cytochrome P450 family 11 subfamily B member 2 [Source:HGNC Symbol;Acc:HGNC:2592] | 3.48 | 1.15E-11 |
| ENSG00000171130 | ATP6V0E2 | ATPase H+ transporting V0 subunit e2 [Source:HGNC Symbol;Acc:HGNC:21723] | 0.54 | 2.26E-11 |
| ENSG00000070061 | ELP1 | elongator complex protein 1 [Source:HGNC Symbol;Acc:HGNC:5959] | 0.96 | 2.26E-11 |
| ENSG00000196782 | MAML3 | mastermind like transcriptional coactivator 3 [Source:HGNC Symbol;Acc:HGNC:16272] | 0.67 | 2.89E-11 |
| ENSG00000084710 | EFR3B | EFR3 homolog B [Source:HGNC Symbol;Acc:HGNC:29155] | 1.37 | 3.53E-11 |
| ENSG00000272398 | CD24 | CD24 molecule [Source:HGNC Symbol;Acc:HGNC:1645] | 2.14 | 3.53E-11 |
| ENSG00000165152 | PGAP4 | post-GPI attachment to proteins GalNAc transferase 4 [Source:HGNC Symbol;Acc:HGNC:28 | 0.61 | 3.77E-11 |
| ENSG00000134602 | STK26 | serine/threonine kinase 26 [Source:HGNC Symbol;Acc:HGNC:18174] | 1.08 | 4.64E-11 |
| ENSG00000183023 | SLC8A1 | solute carrier family 8 member A1 [Source:HGNC Symbol;Acc:HGNC:11068] | 2.12 | 5.30E-11 |
| ENSG00000160882 | CYP11B1 | cytochrome P450 family 11 subfamily B member 1 [Source:HGNC Symbol;Acc:HGNC:2591] | 2.27 | 5.73E-11 |
| ENSG00000078098 | FAP | fibroblast activation protein alpha [Source:HGNC Symbol;Acc:HGNC:3590] | 0.94 | 7.10E-11 |
| ENSG00000054793 | ATP9A | ATPase phospholipid transporting 9A (putative) [Source:HGNC Symbol;Acc:HGNC:13540] | 0.72 | 7.86E-11 |
| ENSG00000137575 | SDCBP | syndecan binding protein [Source:HGNC Symbol;Acc:HGNC:10662] | 0.80 | 9.17E-11 |
| ENSG00000058866 | DGKG | diacylglycerol kinase gamma [Source:HGNC Symbol;Acc:HGNC:2853] | 0.70 | 1.09E-10 |
| ENSG00000110697 | PITPNM1 | phosphatidylinositol transfer protein membrane associated 1 [Source:HGNC Symbol;Acc:HG | 0.78 | 1.13E-10 |
| ENSG00000266401 | NA | novel transcript, antisense to DLGAP1 | 2.10 | 1.18E-10 |
| ENSG00000058404 | CAMK2B | calcium/calmodulin dependent protein kinase II beta [Source:HGNC Symbol;Acc:HGNC:146 | 1.18 | 1.23E-10 |
| ENSG00000136997 | MYC | MYC proto-oncogene, bHLH transcription factor [Source:HGNC Symbol;Acc:HGNC:7553] | -0.99 | 1.88E-10 |
| ENSG00000133083 | DCLK1 | doublecortin like kinase 1 [Source:HGNC Symbol;Acc:HGNC:2700] | -0.72 | 2.04E-10 |
| ENSG00000153132 | CLGN | calmegin [Source:HGNC Symbol;Acc:HGNC:2060] | 0.58 | 2.04E-10 |
| ENSG00000179314 | WSCD1 | WSC domain containing 1 [Source:HGNC Symbol;Acc:HGNC:29060] | 1.96 | 2.06E-10 |
| ENSG00000158966 | CACHD1 | cache domain containing 1 [Source:HGNC Symbol;Acc:HGNC:29314] | 0.62 | 2.82E-10 |
| ENSG00000133612 | AGAP3 | ArfGAP with GTPase domain, ankyrin repeat and PH domain 3 [Source:HGNC Symbol;Acc:H | 0.71 | 5.41E-10 |
| ENSG00000139988 | RDH12 | retinol dehydrogenase 12 [Source:HGNC Symbol;Acc:HGNC:19977] | 1.45 | 5.41E-10 |
| ENSG00000151790 | TDO2 | tryptophan 2,3-dioxygenase [Source:HGNC Symbol;Acc:HGNC:11708] | 1.29 | 8.31E-10 |
| ENSG00000178966 | RMI1 | RecQ mediated genome instability 1 [Source:HGNC Symbol;Acc:HGNC:25764] | -0.52 | 1.14E-09 |
| ENSG00000108515 | ENO3 | enolase 3 [Source:HGNC Symbol;Acc:HGNC:3354] | 0.60 | 1.33E-09 |
| ENSG00000280061 | NA | TEC | 0.98 | 1.43E-09 |
| ENSG00000117586 | TNFSF4 | TNF superfamily member 4 [Source:HGNC Symbol;Acc:HGNC:11934] | 1.02 | 1.49E-09 |
| ENSG00000171408 | PDE7B | phosphodiesterase 7B [Source:HGNC Symbol;Acc:HGNC:8792] | 0.80 | 1.81E-09 |
| ENSG00000149557 | FEZ1 | fasciculation and elongation protein zeta 1 [Source:HGNC Symbol;Acc:HGNC:3659] | 1.10 | 2.05E-09 |
| ENSG00000167528 | ZNF641 | zinc finger protein 641 [Source:HGNC Symbol;Acc:HGNC:31834] | 1.06 | 2.05E-09 |

| | | | | |
|---|---|---|---|---|
| ENSG00000155011 | DKK2 | dickkopf WNT signaling pathway inhibitor 2 [Source:HGNC Symbol;Acc:HGNC:2892] | 2.45 | 2.11E-09 |
| ENSG00000118513 | MYB | MYB proto-oncogene, transcription factor [Source:HGNC Symbol;Acc:HGNC:7545] | 1.44 | 3.64E-09 |
| ENSG00000010818 | HIVEP2 | HIVEP zinc finger 2 [Source:HGNC Symbol;Acc:HGNC:4921] | 0.85 | 3.70E-09 |
| ENSG00000104043 | ATP8B4 | ATPase phospholipid transporting 8B4 (putative) [Source:HGNC Symbol;Acc:HGNC:13536] | 1.56 | 3.70E-09 |
| ENSG00000185231 | MC2R | melanocortin 2 receptor [Source:HGNC Symbol;Acc:HGNC:6930] | 2.32 | 4.09E-09 |
| ENSG00000146250 | PRSS35 | serine protease 35 [Source:HGNC Symbol;Acc:HGNC:21387] | 1.36 | 4.17E-09 |
| ENSG00000177875 | CCDC184 | coiled-coil domain containing 184 [Source:HGNC Symbol;Acc:HGNC:33749] | 1.06 | 5.87E-09 |
| ENSG00000053747 | LAMA3 | laminin subunit alpha 3 [Source:HGNC Symbol;Acc:HGNC:6483] | 1.53 | 5.90E-09 |
| ENSG00000121966 | CXCR4 | C-X-C motif chemokine receptor 4 [Source:HGNC Symbol;Acc:HGNC:2561] | 0.84 | 5.92E-09 |
| ENSG00000132970 | WASF3 | WASP family member 3 [Source:HGNC Symbol;Acc:HGNC:12734] | -0.62 | 1.09E-08 |
| ENSG00000018236 | CNTN1 | contactin 1 [Source:HGNC Symbol;Acc:HGNC:2171] | 1.44 | 1.17E-08 |
| ENSG00000228536 | LYPLAL1-AS1 | LYPLAL1 antisense RNA 1 [Source:HGNC Symbol;Acc:HGNC:54054] | 1.15 | 1.23E-08 |
| ENSG00000140848 | CPNE2 | copine 2 [Source:HGNC Symbol;Acc:HGNC:2315] | 0.82 | 1.43E-08 |
| ENSG00000215218 | UBE2QL1 | ubiquitin conjugating enzyme E2 Q family like 1 [Source:HGNC Symbol;Acc:HGNC:37269] | 0.96 | 1.72E-08 |
| ENSG00000198597 | ZNF536 | zinc finger protein 536 [Source:HGNC Symbol;Acc:HGNC:29025] | 0.81 | 1.83E-08 |
| ENSG00000111674 | ENO2 | enolase 2 [Source:HGNC Symbol;Acc:HGNC:3353] | 0.84 | 2.47E-08 |
| ENSG00000179855 | GIPC3 | GIPC PDZ domain containing family member 3 [Source:HGNC Symbol;Acc:HGNC:18183] | 1.48 | 2.53E-08 |
| ENSG00000053372 | MRTO4 | MRT4 homolog, ribosome maturation factor [Source:HGNC Symbol;Acc:HGNC:18477] | -0.57 | 2.94E-08 |
| ENSG00000168539 | CHRM1 | cholinergic receptor muscarinic 1 [Source:HGNC Symbol;Acc:HGNC:1950] | 2.93 | 3.17E-08 |
| ENSG00000149571 | KIRREL3 | kirre like nephrin family adhesion molecule 3 [Source:HGNC Symbol;Acc:HGNC:23204] | 1.45 | 3.41E-08 |
| ENSG00000180616 | SSTR2 | somatostatin receptor 2 [Source:HGNC Symbol;Acc:HGNC:11331] | 1.32 | 3.41E-08 |
| ENSG00000090975 | PITPNM2 | phosphatidylinositol transfer protein membrane associated 2 [Source:HGNC Symbol;Acc:HG | 0.60 | 3.68E-08 |
| ENSG00000173641 | HSPB7 | heat shock protein family B (small) member 7 [Source:HGNC Symbol;Acc:HGNC:5249] | 0.68 | 3.70E-08 |
| ENSG00000106624 | AEBP1 | AE binding protein 1 [Source:HGNC Symbol;Acc:HGNC:303] | 0.57 | 3.94E-08 |
| ENSG00000102780 | DGKH | diacylglycerol kinase eta [Source:HGNC Symbol;Acc:HGNC:2854] | 0.89 | 4.77E-08 |
| ENSG00000173548 | SNX33 | sorting nexin 33 [Source:HGNC Symbol;Acc:HGNC:28468] | 1.21 | 5.12E-08 |
| ENSG00000072110 | ACTN1 | actinin alpha 1 [Source:HGNC Symbol;Acc:HGNC:163] | 0.77 | 5.16E-08 |
| ENSG00000102962 | CCL22 | C-C motif chemokine ligand 22 [Source:HGNC Symbol;Acc:HGNC:10621] | 0.87 | 5.46E-08 |
| ENSG00000131791 | PRKAB2 | protein kinase AMP-activated non-catalytic subunit beta 2 [Source:HGNC Symbol;Acc:HGNC | -0.58 | 5.81E-08 |
| ENSG00000092969 | TGFB2 | transforming growth factor beta 2 [Source:HGNC Symbol;Acc:HGNC:11768] | 1.32 | 5.87E-08 |
| ENSG00000181045 | SLC26A11 | solute carrier family 26 member 11 [Source:HGNC Symbol;Acc:HGNC:14471] | 0.59 | 5.92E-08 |
| ENSG00000253882 | NA | family with sequence similarity 115, member C (FAM115C) pseudogene | -1.60 | 6.23E-08 |
| ENSG00000124479 | NDP | norrin cystine knot growth factor NDP [Source:HGNC Symbol;Acc:HGNC:7678] | -1.58 | 1.15E-07 |
| ENSG00000115255 | REEP6 | receptor accessory protein 6 [Source:HGNC Symbol;Acc:HGNC:30078] | -0.55 | 1.17E-07 |
| ENSG00000184838 | PRR16 | proline rich 16 [Source:HGNC Symbol;Acc:HGNC:29654] | -0.88 | 1.19E-07 |
| ENSG00000141639 | MAPK4 | mitogen-activated protein kinase 4 [Source:HGNC Symbol;Acc:HGNC:6878] | 1.05 | 1.41E-07 |
| ENSG00000198369 | SPRED2 | sprouty related EVH1 domain containing 2 [Source:HGNC Symbol;Acc:HGNC:17722] | 1.21 | 1.52E-07 |
| ENSG00000254615 | NA | novel transcript | 1.91 | 1.54E-07 |
| ENSG00000152932 | RAB3C | RAB3C, member RAS oncogene family [Source:HGNC Symbol;Acc:HGNC:30269] | 1.00 | 2.02E-07 |
| ENSG00000086300 | SNX10 | sorting nexin 10 [Source:HGNC Symbol;Acc:HGNC:14974] | 0.58 | 2.05E-07 |

| | | | | |
|---|---|---|---|---|
| ENSG00000116514 | RNF19B | ring finger protein 19B [Source:HGNC Symbol;Acc:HGNC:26886] | 0.91 | 2.13E-07 |
| ENSG00000137713 | PPP2R1B | protein phosphatase 2 scaffold subunit Abeta [Source:HGNC Symbol;Acc:HGNC:9303] | 0.91 | 2.20E-07 |
| ENSG00000124588 | NQO2 | N-ribosyldihydronicotinamide:quinone reductase 2 [Source:HGNC Symbol;Acc:HGNC:7856] | 0.82 | 2.26E-07 |
| ENSG00000187479 | C11orf96 | chromosome 11 open reading frame 96 [Source:HGNC Symbol;Acc:HGNC:38675] | 1.36 | 2.51E-07 |
| ENSG00000090530 | P3H2 | prolyl 3-hydroxylase 2 [Source:HGNC Symbol;Acc:HGNC:19317] | 0.72 | 3.01E-07 |
| ENSG00000148841 | ITPRIP | inositol 1,4,5-trisphosphate receptor interacting protein [Source:HGNC Symbol;Acc:HGNC:2 | 0.89 | 3.06E-07 |
| ENSG00000122012 | SV2C | synaptic vesicle glycoprotein 2C [Source:HGNC Symbol;Acc:HGNC:30670] | 0.96 | 3.09E-07 |
| ENSG00000152049 | KCNE4 | potassium voltage-gated channel subfamily E regulatory subunit 4 [Source:HGNC Symbol;A | 1.09 | 3.09E-07 |
| ENSG00000155265 | GOLGA7B | golgin A7 family member B [Source:HGNC Symbol;Acc:HGNC:31668] | 0.98 | 3.09E-07 |
| ENSG00000084674 | APOB | apolipoprotein B [Source:HGNC Symbol;Acc:HGNC:603] | 0.94 | 3.13E-07 |
| ENSG00000236008 | LINC01814 | long intergenic non-protein coding RNA 1814 [Source:HGNC Symbol;Acc:HGNC:52618] | -0.83 | 3.89E-07 |
| ENSG00000006210 | CX3CL1 | C-X3-C motif chemokine ligand 1 [Source:HGNC Symbol;Acc:HGNC:10647] | 0.65 | 4.03E-07 |
| ENSG00000137124 | ALDH1B1 | aldehyde dehydrogenase 1 family member B1 [Source:HGNC Symbol;Acc:HGNC:407] | -0.51 | 4.30E-07 |
| ENSG00000119927 | GPAM | glycerol-3-phosphate acyltransferase, mitochondrial [Source:HGNC Symbol;Acc:HGNC:248 | -0.66 | 5.38E-07 |
| ENSG00000124787 | RPP40 | ribonuclease P/MRP subunit p40 [Source:HGNC Symbol;Acc:HGNC:20992] | -0.81 | 5.42E-07 |
| ENSG00000136928 | GABBR2 | gamma-aminobutyric acid type B receptor subunit 2 [Source:HGNC Symbol;Acc:HGNC:4507 | -0.63 | 5.63E-07 |
| ENSG00000197852 | INKA2 | inka box actin regulator 2 [Source:HGNC Symbol;Acc:HGNC:28045] | 0.73 | 6.15E-07 |
| ENSG00000168671 | UGT3A2 | UDP glycosyltransferase family 3 member A2 [Source:HGNC Symbol;Acc:HGNC:27266] | 1.01 | 6.25E-07 |
| ENSG00000072422 | RHOBTB1 | Rho related BTB domain containing 1 [Source:HGNC Symbol;Acc:HGNC:18738] | -0.65 | 6.42E-07 |
| ENSG00000140859 | KIFC3 | kinesin family member C3 [Source:HGNC Symbol;Acc:HGNC:6326] | 0.83 | 6.96E-07 |
| ENSG00000107738 | VSIR | V-set immunoregulatory receptor [Source:HGNC Symbol;Acc:HGNC:30085] | 2.47 | 7.43E-07 |
| ENSG00000163958 | ZDHHC19 | zinc finger DHHC-type palmitoyltransferase 19 [Source:HGNC Symbol;Acc:HGNC:20713] | -1.10 | 7.90E-07 |
| ENSG00000187720 | THSD4 | thrombospondin type 1 domain containing 4 [Source:HGNC Symbol;Acc:HGNC:25835] | 0.81 | 8.93E-07 |
| ENSG00000136155 | SCEL | sciellin [Source:HGNC Symbol;Acc:HGNC:10573] | -1.98 | 9.03E-07 |
| ENSG00000168675 | LDLRAD4 | low density lipoprotein receptor class A domain containing 4 [Source:HGNC Symbol;Acc:HG | 1.20 | 9.03E-07 |
| ENSG00000181773 | GPR3 | G protein-coupled receptor 3 [Source:HGNC Symbol;Acc:HGNC:4484] | 2.25 | 9.16E-07 |
| ENSG00000169330 | MINAR1 | membrane integral NOTCH2 associated receptor 1 [Source:HGNC Symbol;Acc:HGNC:2917 | 0.94 | 9.20E-07 |
| ENSG00000225663 | MCRIP1 | MAPK regulated corepressor interacting protein 1 [Source:HGNC Symbol;Acc:HGNC:28007 | -0.63 | 1.00E-06 |
| ENSG00000087510 | TFAP2C | transcription factor AP-2 gamma [Source:HGNC Symbol;Acc:HGNC:11744] | 1.26 | 1.02E-06 |
| ENSG00000095303 | PTGS1 | prostaglandin-endoperoxide synthase 1 [Source:HGNC Symbol;Acc:HGNC:9604] | 1.42 | 1.12E-06 |
| ENSG00000127920 | GNG11 | G protein subunit gamma 11 [Source:HGNC Symbol;Acc:HGNC:4403] | 0.97 | 1.15E-06 |
| ENSG00000204262 | COL5A2 | collagen type V alpha 2 chain [Source:HGNC Symbol;Acc:HGNC:2210] | 0.68 | 1.15E-06 |
| ENSG00000112186 | CAP2 | cyclase associated actin cytoskeleton regulatory protein 2 [Source:HGNC Symbol;Acc:HGN | 0.61 | 1.30E-06 |
| ENSG00000186862 | PDZD7 | PDZ domain containing 7 [Source:HGNC Symbol;Acc:HGNC:26257] | 1.32 | 1.30E-06 |
| ENSG00000162552 | WNT4 | Wnt family member 4 [Source:HGNC Symbol;Acc:HGNC:12783] | -0.86 | 1.38E-06 |
| ENSG00000174156 | GSTA3 | glutathione S-transferase alpha 3 [Source:HGNC Symbol;Acc:HGNC:4628] | -0.80 | 1.38E-06 |
| ENSG00000170485 | NPAS2 | neuronal PAS domain protein 2 [Source:HGNC Symbol;Acc:HGNC:7895] | 1.00 | 1.46E-06 |
| ENSG00000120068 | HOXB8 | homeobox B8 [Source:HGNC Symbol;Acc:HGNC:5119] | 0.92 | 1.66E-06 |
| ENSG00000168394 | TAP1 | transporter 1, ATP binding cassette subfamily B member [Source:HGNC Symbol;Acc:HGNC: | 0.71 | 1.94E-06 |
| ENSG00000135439 | AGAP2 | ArfGAP with GTPase domain, ankyrin repeat and PH domain 2 [Source:HGNC Symbol;Acc:H | 0.87 | 2.01E-06 |

| | | | | |
|---|---|---|---|---|
| ENSG00000117707 | PROX1 | prospero homeobox 1 [Source:HGNC Symbol;Acc:HGNC:9459] | 0.99 | 2.10E-06 |
| ENSG00000145246 | ATP10D | ATPase phospholipid transporting 10D (putative) [Source:HGNC Symbol;Acc:HGNC:13549] | 0.91 | 2.20E-06 |
| ENSG00000204516 | MICB | MHC class I polypeptide-related sequence B [Source:HGNC Symbol;Acc:HGNC:7091] | 1.03 | 2.31E-06 |
| ENSG00000151229 | SLC2A13 | solute carrier family 2 member 13 [Source:HGNC Symbol;Acc:HGNC:15956] | 0.99 | 2.54E-06 |
| ENSG00000188582 | PAQR9 | progestin and adipoQ receptor family member 9 [Source:HGNC Symbol;Acc:HGNC:30131] | 1.62 | 2.99E-06 |
| ENSG00000261325 | LINC02192 | long intergenic non-protein coding RNA 2192 [Source:HGNC Symbol;Acc:HGNC:53054] | 1.21 | 3.36E-06 |
| ENSG00000143641 | GALNT2 | polypeptide N-acetylgalactosaminyltransferase 2 [Source:HGNC Symbol;Acc:HGNC:4124] | 0.58 | 3.68E-06 |
| ENSG00000145779 | TNFAIP8 | TNF alpha induced protein 8 [Source:HGNC Symbol;Acc:HGNC:17260] | 0.99 | 3.68E-06 |
| ENSG00000099625 | CBARP | CACN subunit beta associated regulatory protein [Source:HGNC Symbol;Acc:HGNC:28617] | 1.78 | 3.91E-06 |
| ENSG00000255036 | STRA6LP | STRA6 like, pseudogene [Source:HGNC Symbol;Acc:HGNC:53830] | 0.70 | 4.06E-06 |
| ENSG00000108176 | DNAJC12 | DnaJ heat shock protein family (Hsp40) member C12 [Source:HGNC Symbol;Acc:HGNC:28 | 0.95 | 4.79E-06 |
| ENSG00000185742 | C11orf87 | chromosome 11 open reading frame 87 [Source:HGNC Symbol;Acc:HGNC:33788] | 1.78 | 5.39E-06 |
| ENSG00000157680 | DGKI | diacylglycerol kinase iota [Source:HGNC Symbol;Acc:HGNC:2855] | 1.13 | 5.70E-06 |
| ENSG00000175832 | ETV4 | ETS variant transcription factor 4 [Source:HGNC Symbol;Acc:HGNC:3493] | 3.51 | 6.05E-06 |
| ENSG00000104332 | SFRP1 | secreted frizzled related protein 1 [Source:HGNC Symbol;Acc:HGNC:10776] | 2.41 | 6.50E-06 |
| ENSG00000155090 | KLF10 | Kruppel like factor 10 [Source:HGNC Symbol;Acc:HGNC:11810] | -1.12 | 6.86E-06 |
| ENSG00000101638 | ST8SIA5 | ST8 alpha-N-acetyl-neuraminide alpha-2,8-sialyltransferase 5 [Source:HGNC Symbol;Acc:H | -0.60 | 8.15E-06 |
| ENSG00000235026 | DPP10-AS1 | DPP10 antisense RNA 1 [Source:HGNC Symbol;Acc:HGNC:40941] | -0.73 | 8.74E-06 |
| ENSG00000106689 | LHX2 | LIM homeobox 2 [Source:HGNC Symbol;Acc:HGNC:6594] | 1.19 | 9.64E-06 |
| ENSG00000135541 | AHI1 | Abelson helper integration site 1 [Source:HGNC Symbol;Acc:HGNC:21575] | 0.63 | 9.83E-06 |
| ENSG00000010278 | CD9 | CD9 molecule [Source:HGNC Symbol;Acc:HGNC:1709] | 0.92 | 9.95E-06 |
| ENSG00000167580 | AQP2 | aquaporin 2 [Source:HGNC Symbol;Acc:HGNC:634] | 1.04 | 9.95E-06 |
| ENSG00000170915 | PAQR8 | progestin and adipoQ receptor family member 8 [Source:HGNC Symbol;Acc:HGNC:15708] | 0.55 | 9.96E-06 |
| ENSG00000184675 | AMER1 | APC membrane recruitment protein 1 [Source:HGNC Symbol;Acc:HGNC:26837] | -0.65 | 1.07E-05 |
| ENSG00000172551 | MUCL1 | mucin like 1 [Source:HGNC Symbol;Acc:HGNC:30588] | -0.71 | 1.09E-05 |
| ENSG00000129158 | SERGEF | secretion regulating guanine nucleotide exchange factor [Source:HGNC Symbol;Acc:HGNC | 1.00 | 1.13E-05 |
| ENSG00000120306 | CYSTM1 | cysteine rich transmembrane module containing 1 [Source:HGNC Symbol;Acc:HGNC:3023 | 0.58 | 1.14E-05 |
| ENSG00000068971 | PPP2R5B | protein phosphatase 2 regulatory subunit B'beta [Source:HGNC Symbol;Acc:HGNC:9310] | 0.51 | 1.15E-05 |
| ENSG00000170624 | SGCD | sarcoglycan delta [Source:HGNC Symbol;Acc:HGNC:10807] | 0.54 | 1.17E-05 |
| ENSG00000145632 | PLK2 | polo like kinase 2 [Source:HGNC Symbol;Acc:HGNC:19699] | 1.75 | 1.22E-05 |
| ENSG00000147894 | C9orf72 | C9orf72-SMCR8 complex subunit [Source:HGNC Symbol;Acc:HGNC:28337] | 0.62 | 1.22E-05 |
| ENSG00000130052 | STARD8 | StAR related lipid transfer domain containing 8 [Source:HGNC Symbol;Acc:HGNC:19161] | 0.52 | 1.27E-05 |
| ENSG00000167693 | NXN | nucleoredoxin [Source:HGNC Symbol;Acc:HGNC:18008] | 1.11 | 1.28E-05 |
| ENSG00000136933 | RABEPK | Rab9 effector protein with kelch motifs [Source:HGNC Symbol;Acc:HGNC:16896] | -0.51 | 1.29E-05 |
| ENSG00000125520 | SLC2A4RG | SLC2A4 regulator [Source:HGNC Symbol;Acc:HGNC:15930] | -0.55 | 1.33E-05 |
| ENSG00000167100 | SAMD14 | sterile alpha motif domain containing 14 [Source:HGNC Symbol;Acc:HGNC:27312] | 0.86 | 1.36E-05 |
| ENSG00000099260 | PALMD | palmdelphin [Source:HGNC Symbol;Acc:HGNC:15846] | -0.81 | 1.38E-05 |
| ENSG00000105825 | TFPI2 | tissue factor pathway inhibitor 2 [Source:HGNC Symbol;Acc:HGNC:11761] | 1.43 | 1.54E-05 |
| ENSG00000259417 | CTXND1 | cortexin domain containing 1 [Source:HGNC Symbol;Acc:HGNC:50507] | 1.61 | 1.54E-05 |
| ENSG00000166979 | EVA1C | eva-1 homolog C [Source:HGNC Symbol;Acc:HGNC:13239] | 1.31 | 1.55E-05 |

| | | | | |
|---|---|---|---|---|
| ENSG00000107937 | GTPBP4 | GTP binding protein 4 [Source:HGNC Symbol;Acc:HGNC:21535] | -0.76 | 1.57E-05 |
| ENSG00000128641 | MYO1B | myosin IB [Source:HGNC Symbol;Acc:HGNC:7596] | 3.00 | 1.57E-05 |
| ENSG00000243955 | GSTA1 | glutathione S-transferase alpha 1 [Source:HGNC Symbol;Acc:HGNC:4626] | -0.82 | 1.59E-05 |
| ENSG00000167861 | HID1 | HID1 domain containing [Source:HGNC Symbol;Acc:HGNC:15736] | 0.84 | 1.72E-05 |
| ENSG00000144891 | AGTR1 | angiotensin II receptor type 1 [Source:HGNC Symbol;Acc:HGNC:336] | -0.54 | 1.81E-05 |
| ENSG00000121297 | TSHZ3 | teashirt zinc finger homeobox 3 [Source:HGNC Symbol;Acc:HGNC:30700] | 0.73 | 1.83E-05 |
| ENSG00000165698 | SPACA9 | sperm acrosome associated 9 [Source:HGNC Symbol;Acc:HGNC:1367] | -0.73 | 1.91E-05 |
| ENSG00000173334 | TRIB1 | tribbles pseudokinase 1 [Source:HGNC Symbol;Acc:HGNC:16891] | 1.86 | 2.03E-05 |
| ENSG00000129682 | FGF13 | fibroblast growth factor 13 [Source:HGNC Symbol;Acc:HGNC:3670] | -0.61 | 2.07E-05 |
| ENSG00000151914 | DST | dystonin [Source:HGNC Symbol;Acc:HGNC:1090] | 0.52 | 2.07E-05 |
| ENSG00000164237 | CMBL | carboxymethylenebutenolidase homolog [Source:HGNC Symbol;Acc:HGNC:25090] | -0.63 | 2.07E-05 |
| ENSG00000261594 | TPBGL | trophoblast glycoprotein like [Source:HGNC Symbol;Acc:HGNC:44159] | 1.33 | 2.10E-05 |
| ENSG00000230918 | DPP4-DT | DPP4 divergent transcript [Source:HGNC Symbol;Acc:HGNC:40191] | 1.53 | 2.13E-05 |
| ENSG00000087303 | NID2 | nidogen 2 [Source:HGNC Symbol;Acc:HGNC:13389] | 1.30 | 2.14E-05 |
| ENSG00000125744 | RTN2 | reticulon 2 [Source:HGNC Symbol;Acc:HGNC:10468] | 0.94 | 2.14E-05 |
| ENSG00000137959 | IFI44L | interferon induced protein 44 like [Source:HGNC Symbol;Acc:HGNC:17817] | -0.95 | 2.31E-05 |
| ENSG00000161682 | FAM171A2 | family with sequence similarity 171 member A2 [Source:HGNC Symbol;Acc:HGNC:30480] | 0.96 | 2.51E-05 |
| ENSG00000053918 | KCNQ1 | potassium voltage-gated channel subfamily Q member 1 [Source:HGNC Symbol;Acc:HGNC | 1.60 | 2.63E-05 |
| ENSG00000164089 | ETNPPL | ethanolamine-phosphate phospho-lyase [Source:HGNC Symbol;Acc:HGNC:14404] | 1.07 | 2.68E-05 |
| ENSG00000071246 | VASH1 | vasohibin 1 [Source:HGNC Symbol;Acc:HGNC:19964] | 0.50 | 2.87E-05 |
| ENSG00000127824 | TUBA4A | tubulin alpha 4a [Source:HGNC Symbol;Acc:HGNC:12407] | 1.09 | 2.93E-05 |
| ENSG00000157514 | TSC22D3 | TSC22 domain family member 3 [Source:HGNC Symbol;Acc:HGNC:3051] | 0.52 | 2.93E-05 |
| ENSG00000178233 | TMEM151B | transmembrane protein 151B [Source:HGNC Symbol;Acc:HGNC:21315] | 1.09 | 3.30E-05 |
| ENSG00000250208 | FZD10-AS1 | FZD10 antisense divergent transcript [Source:HGNC Symbol;Acc:HGNC:48632] | 1.78 | 3.43E-05 |
| ENSG00000164050 | PLXNB1 | plexin B1 [Source:HGNC Symbol;Acc:HGNC:9103] | 0.64 | 3.61E-05 |
| ENSG00000137872 | SEMA6D | semaphorin 6D [Source:HGNC Symbol;Acc:HGNC:16770] | 1.00 | 3.61E-05 |
| ENSG00000134569 | LRP4 | LDL receptor related protein 4 [Source:HGNC Symbol;Acc:HGNC:6696] | 1.51 | 3.65E-05 |
| ENSG00000151067 | CACNA1C | calcium voltage-gated channel subunit alpha1 C [Source:HGNC Symbol;Acc:HGNC:1390] | 0.81 | 3.69E-05 |
| ENSG00000019549 | SNAI2 | snail family transcriptional repressor 2 [Source:HGNC Symbol;Acc:HGNC:11094] | -0.62 | 3.70E-05 |
| ENSG00000120875 | DUSP4 | dual specificity phosphatase 4 [Source:HGNC Symbol;Acc:HGNC:3070] | 1.00 | 3.70E-05 |
| ENSG00000127561 | SYNGR3 | synaptogyrin 3 [Source:HGNC Symbol;Acc:HGNC:11501] | 0.71 | 3.70E-05 |
| ENSG00000108244 | KRT23 | keratin 23 [Source:HGNC Symbol;Acc:HGNC:6438] | 1.72 | 4.00E-05 |
| ENSG00000141258 | SGSM2 | small G protein signaling modulator 2 [Source:HGNC Symbol;Acc:HGNC:29026] | 0.65 | 4.05E-05 |
| ENSG00000146233 | CYP39A1 | cytochrome P450 family 39 subfamily A member 1 [Source:HGNC Symbol;Acc:HGNC:1744[illegible] | 0.84 | 4.16E-05 |
| ENSG00000111057 | KRT18 | keratin 18 [Source:HGNC Symbol;Acc:HGNC:6430] | 0.51 | 4.52E-05 |
| ENSG00000225376 | TMEM246-AS1 | TMEM246 antisense RNA 1 [Source:HGNC Symbol;Acc:HGNC:51191] | 1.77 | 4.53E-05 |
| ENSG00000157827 | FMNL2 | formin like 2 [Source:HGNC Symbol;Acc:HGNC:18267] | 0.59 | 4.58E-05 |
| ENSG00000139182 | CLSTN3 | calsyntenin 3 [Source:HGNC Symbol;Acc:HGNC:18371] | 0.73 | 4.79E-05 |
| ENSG00000105711 | SCN1B | sodium voltage-gated channel beta subunit 1 [Source:HGNC Symbol;Acc:HGNC:10586] | 1.07 | 5.06E-05 |
| ENSG00000186806 | VSIG10L | V-set and immunoglobulin domain containing 10 like [Source:HGNC Symbol;Acc:HGNC:271 | -0.61 | 5.07E-05 |

| | | | | |
|---|---|---|---|---|
| ENSG00000187243 | MAGED4B | MAGE family member D4B [Source:HGNC Symbol;Acc:HGNC:22880] | 0.61 | 5.24E-05 |
| ENSG00000141854 | MISP3 | MISP family member 3 [Source:HGNC Symbol;Acc:HGNC:26963] | 1.14 | 5.37E-05 |
| ENSG00000131018 | SYNE1 | spectrin repeat containing nuclear envelope protein 1 [Source:HGNC Symbol;Acc:HGNC:17 | 0.57 | 5.40E-05 |
| ENSG00000164761 | TNFRSF11B | TNF receptor superfamily member 11b [Source:HGNC Symbol;Acc:HGNC:11909] | 0.81 | 5.40E-05 |
| ENSG00000100003 | SEC14L2 | SEC14 like lipid binding 2 [Source:HGNC Symbol;Acc:HGNC:10699] | 0.71 | 5.57E-05 |
| ENSG00000115963 | RND3 | Rho family GTPase 3 [Source:HGNC Symbol;Acc:HGNC:671] | 1.68 | 5.58E-05 |
| ENSG00000087085 | ACHE | acetylcholinesterase (Cartwright blood group) [Source:HGNC Symbol;Acc:HGNC:108] | 1.17 | 5.63E-05 |
| ENSG00000140284 | SLC27A2 | solute carrier family 27 member 2 [Source:HGNC Symbol;Acc:HGNC:10996] | 2.54 | 5.74E-05 |
| ENSG00000175928 | LRRN1 | leucine rich repeat neuronal 1 [Source:HGNC Symbol;Acc:HGNC:20980] | 1.33 | 6.11E-05 |
| ENSG00000245864 | MEF2C-AS2 | MEF2C antisense RNA 2 [Source:HGNC Symbol;Acc:HGNC:53115] | 0.54 | 6.55E-05 |
| ENSG00000064547 | LPAR2 | lysophosphatidic acid receptor 2 [Source:HGNC Symbol;Acc:HGNC:3168] | 1.05 | 6.68E-05 |
| ENSG00000229847 | EMX2OS | EMX2 opposite strand/antisense RNA [Source:HGNC Symbol;Acc:HGNC:18511] | 1.41 | 6.71E-05 |
| ENSG00000070182 | SPTB | spectrin beta, erythrocytic [Source:HGNC Symbol;Acc:HGNC:11274] | 0.67 | 6.90E-05 |
| ENSG00000142634 | EFHD2 | EF-hand domain family member D2 [Source:HGNC Symbol;Acc:HGNC:28670] | -0.59 | 6.97E-05 |
| ENSG00000139428 | MMAB | metabolism of cobalamin associated B [Source:HGNC Symbol;Acc:HGNC:19331] | -0.64 | 7.22E-05 |
| ENSG00000179041 | RRS1 | ribosome biogenesis regulator 1 homolog [Source:HGNC Symbol;Acc:HGNC:17083] | -0.52 | 7.40E-05 |
| ENSG00000189046 | ALKBH2 | alkB homolog 2, alpha-ketoglutarate dependent dioxygenase [Source:HGNC Symbol;Acc:H | -0.69 | 7.44E-05 |
| ENSG00000007402 | CACNA2D2 | calcium voltage-gated channel auxiliary subunit alpha2delta 2 [Source:HGNC Symbol;Acc:H | 0.66 | 7.60E-05 |
| ENSG00000147027 | TMEM47 | transmembrane protein 47 [Source:HGNC Symbol;Acc:HGNC:18515] | 0.66 | 7.60E-05 |
| ENSG00000168214 | RBPJ | recombination signal binding protein for immunoglobulin kappa J region [Source:HGNC Syn | 0.52 | 7.83E-05 |
| ENSG00000110880 | CORO1C | coronin 1C [Source:HGNC Symbol;Acc:HGNC:2254] | 0.58 | 7.92E-05 |
| ENSG00000123080 | CDKN2C | cyclin dependent kinase inhibitor 2C [Source:HGNC Symbol;Acc:HGNC:1789] | -0.51 | 8.05E-05 |
| ENSG00000127528 | KLF2 | Kruppel like factor 2 [Source:HGNC Symbol;Acc:HGNC:6347] | 1.17 | 8.05E-05 |
| ENSG00000144560 | VGLL4 | vestigial like family member 4 [Source:HGNC Symbol;Acc:HGNC:28966] | 0.68 | 8.05E-05 |
| ENSG00000182575 | NXPH3 | neurexophilin 3 [Source:HGNC Symbol;Acc:HGNC:8077] | 0.93 | 8.05E-05 |
| ENSG00000185875 | THNSL1 | threonine synthase like 1 [Source:HGNC Symbol;Acc:HGNC:26160] | -0.65 | 8.05E-05 |
| ENSG00000154640 | BTG3 | BTG anti-proliferation factor 3 [Source:HGNC Symbol;Acc:HGNC:1132] | 0.73 | 8.07E-05 |
| ENSG00000186642 | PDE2A | phosphodiesterase 2A [Source:HGNC Symbol;Acc:HGNC:8777] | -0.70 | 8.23E-05 |
| ENSG00000165434 | PGM2L1 | phosphoglucomutase 2 like 1 [Source:HGNC Symbol;Acc:HGNC:20898] | 0.91 | 8.40E-05 |
| ENSG00000261971 | MMP25-AS1 | MMP25 antisense RNA 1 [Source:HGNC Symbol;Acc:HGNC:51372] | 0.76 | 8.78E-05 |
| ENSG00000187957 | DNER | delta/notch like EGF repeat containing [Source:HGNC Symbol;Acc:HGNC:24456] | 0.56 | 8.82E-05 |
| ENSG00000168542 | COL3A1 | collagen type III alpha 1 chain [Source:HGNC Symbol;Acc:HGNC:2201] | 2.32 | 9.31E-05 |
| ENSG00000027075 | PRKCH | protein kinase C eta [Source:HGNC Symbol;Acc:HGNC:9403] | 1.17 | 9.50E-05 |
| ENSG00000198915 | RASGEF1A | RasGEF domain family member 1A [Source:HGNC Symbol;Acc:HGNC:24246] | 1.75 | 0.00010 |
| ENSG00000235431 | NA | novel transcript | 0.78 | 0.00010 |
| ENSG00000196843 | ARID5A | AT-rich interaction domain 5A [Source:HGNC Symbol;Acc:HGNC:17361] | 1.63 | 0.00011 |
| ENSG00000183688 | RFLNB | refilin B [Source:HGNC Symbol;Acc:HGNC:28705] | 1.23 | 0.00011 |
| ENSG00000256664 | NA | ribosomal L24 domain containing 1 (RSL24D1) pseudogene | 0.61 | 0.00012 |
| ENSG00000140563 | MCTP2 | multiple C2 and transmembrane domain containing 2 [Source:HGNC Symbol;Acc:HGNC:25 | 0.75 | 0.00012 |
| ENSG00000108813 | DLX4 | distal-less homeobox 4 [Source:HGNC Symbol;Acc:HGNC:2917] | 0.79 | 0.00013 |

| | | | | |
|---|---|---|---|---|
| ENSG00000103356 | EARS2 | glutamyl-tRNA synthetase 2, mitochondrial [Source:HGNC Symbol;Acc:HGNC:29419] | -0.61 | 0.00013 |
| ENSG00000004660 | CAMKK1 | calcium/calmodulin dependent protein kinase kinase 1 [Source:HGNC Symbol;Acc:HGNC:1 | -0.77 | 0.00014 |
| ENSG00000160867 | FGFR4 | fibroblast growth factor receptor 4 [Source:HGNC Symbol;Acc:HGNC:3691] | -0.69 | 0.00015 |
| ENSG00000073670 | ADAM11 | ADAM metallopeptidase domain 11 [Source:HGNC Symbol;Acc:HGNC:189] | 0.95 | 0.00015 |
| ENSG00000233369 | GTF2IP4 | general transcription factor IIi pseudogene 4 [Source:HGNC Symbol;Acc:HGNC:51716] | 0.52 | 0.00015 |
| ENSG00000239887 | C1orf226 | chromosome 1 open reading frame 226 [Source:HGNC Symbol;Acc:HGNC:34351] | -0.62 | 0.00015 |
| ENSG00000005981 | ASB4 | ankyrin repeat and SOCS box containing 4 [Source:HGNC Symbol;Acc:HGNC:16009] | 0.61 | 0.00015 |
| ENSG00000049883 | PTCD2 | pentatricopeptide repeat domain 2 [Source:HGNC Symbol;Acc:HGNC:25734] | -0.51 | 0.00015 |
| ENSG00000145730 | PAM | peptidylglycine alpha-amidating monooxygenase [Source:HGNC Symbol;Acc:HGNC:8596] | 0.78 | 0.00015 |
| ENSG00000143344 | RGL1 | ral guanine nucleotide dissociation stimulator like 1 [Source:HGNC Symbol;Acc:HGNC:3028 | 0.70 | 0.00015 |
| ENSG00000137142 | IGFBPL1 | insulin like growth factor binding protein like 1 [Source:HGNC Symbol;Acc:HGNC:20081] | 1.08 | 0.00016 |
| ENSG00000073969 | NSF | N-ethylmaleimide sensitive factor, vesicle fusing ATPase [Source:HGNC Symbol;Acc:HGNC | 0.52 | 0.00016 |
| ENSG00000157600 | TMEM164 | transmembrane protein 164 [Source:HGNC Symbol;Acc:HGNC:26217] | -0.60 | 0.00016 |
| ENSG00000080608 | PUM3 | pumilio RNA binding family member 3 [Source:HGNC Symbol;Acc:HGNC:29676] | -0.59 | 0.00017 |
| ENSG00000165194 | PCDH19 | protocadherin 19 [Source:HGNC Symbol;Acc:HGNC:14270] | 1.61 | 0.00017 |
| ENSG00000269378 | NA | integrin beta 1 pseudogene 1 | 0.86 | 0.00017 |
| ENSG00000117245 | KIF17 | kinesin family member 17 [Source:HGNC Symbol;Acc:HGNC:19167] | 0.52 | 0.00017 |
| ENSG00000104341 | LAPTM4B | lysosomal protein transmembrane 4 beta [Source:HGNC Symbol;Acc:HGNC:13646] | 1.15 | 0.00019 |
| ENSG00000162494 | LRRC38 | leucine rich repeat containing 38 [Source:HGNC Symbol;Acc:HGNC:27005] | 1.00 | 0.00019 |
| ENSG00000116717 | GADD45A | growth arrest and DNA damage inducible alpha [Source:HGNC Symbol;Acc:HGNC:4095] | 0.75 | 0.00019 |
| ENSG00000182492 | BGN | biglycan [Source:HGNC Symbol;Acc:HGNC:1044] | 0.75 | 0.00020 |
| ENSG00000125534 | PPDPF | pancreatic progenitor cell differentiation and proliferation factor [Source:HGNC Symbol;Acc:H | -0.59 | 0.00021 |
| ENSG00000183654 | MARCHF11 | membrane associated ring-CH-type finger 11 [Source:HGNC Symbol;Acc:HGNC:33609] | 0.68 | 0.00021 |
| ENSG00000135097 | MSI1 | musashi RNA binding protein 1 [Source:HGNC Symbol;Acc:HGNC:7330] | 0.86 | 0.00021 |
| ENSG00000068137 | PLEKHH3 | pleckstrin homology, MyTH4 and FERM domain containing H3 [Source:HGNC Symbol;Acc:H | 0.66 | 0.00022 |
| ENSG00000204291 | COL15A1 | collagen type XV alpha 1 chain [Source:HGNC Symbol;Acc:HGNC:2192] | 0.72 | 0.00022 |
| ENSG00000174672 | BRSK2 | BR serine/threonine kinase 2 [Source:HGNC Symbol;Acc:HGNC:11405] | 0.63 | 0.00022 |
| ENSG00000026103 | FAS | Fas cell surface death receptor [Source:HGNC Symbol;Acc:HGNC:11920] | 0.69 | 0.00022 |
| ENSG00000068383 | INPP5A | inositol polyphosphate-5-phosphatase A [Source:HGNC Symbol;Acc:HGNC:6076] | 0.61 | 0.00023 |
| ENSG00000184588 | PDE4B | phosphodiesterase 4B [Source:HGNC Symbol;Acc:HGNC:8781] | 0.67 | 0.00024 |
| ENSG00000095397 | WHRN | whirlin [Source:HGNC Symbol;Acc:HGNC:16361] | 0.58 | 0.00024 |
| ENSG00000185338 | SOCS1 | suppressor of cytokine signaling 1 [Source:HGNC Symbol;Acc:HGNC:19383] | 0.77 | 0.00024 |
| ENSG00000115738 | ID2 | inhibitor of DNA binding 2 [Source:HGNC Symbol;Acc:HGNC:5361] | -0.51 | 0.00025 |
| ENSG00000119900 | OGFRL1 | opioid growth factor receptor like 1 [Source:HGNC Symbol;Acc:HGNC:21378] | 0.72 | 0.00026 |
| ENSG00000101236 | RNF24 | ring finger protein 24 [Source:HGNC Symbol;Acc:HGNC:13779] | 0.52 | 0.00026 |
| ENSG00000115226 | FNDC4 | fibronectin type III domain containing 4 [Source:HGNC Symbol;Acc:HGNC:20239] | 0.77 | 0.00028 |
| ENSG00000171951 | SCG2 | secretogranin II [Source:HGNC Symbol;Acc:HGNC:10575] | 1.55 | 0.00028 |
| ENSG00000049246 | PER3 | period circadian regulator 3 [Source:HGNC Symbol;Acc:HGNC:8847] | -1.12 | 0.00028 |
| ENSG00000120254 | MTHFD1L | methylenetetrahydrofolate dehydrogenase (NADP+ dependent) 1 like [Source:HGNC Symbo | -0.95 | 0.00029 |
| ENSG00000104381 | GDAP1 | ganglioside induced differentiation associated protein 1 [Source:HGNC Symbol;Acc:HGNC: | -0.54 | 0.00029 |

| | | | | |
|---|---|---|---|---|
| ENSG00000144485 | HES6 | hes family bHLH transcription factor 6 [Source:HGNC Symbol;Acc:HGNC:18254] | 0.92 | 0.00030 |
| ENSG00000198797 | BRINP2 | BMP/retinoic acid inducible neural specific 2 [Source:HGNC Symbol;Acc:HGNC:13746] | 0.65 | 0.00030 |
| ENSG00000148154 | UGCG | UDP-glucose ceramide glucosyltransferase [Source:HGNC Symbol;Acc:HGNC:12524] | 0.51 | 0.00031 |
| ENSG00000162769 | FLVCR1 | FLVCR heme transporter 1 [Source:HGNC Symbol;Acc:HGNC:24682] | 0.50 | 0.00031 |
| ENSG00000261340 | LINC01616 | long intergenic non-protein coding RNA 1616 [Source:HGNC Symbol;Acc:HGNC:51900] | 1.40 | 0.00032 |
| ENSG00000197948 | FCHSD1 | FCH and double SH3 domains 1 [Source:HGNC Symbol;Acc:HGNC:25463] | 0.52 | 0.00032 |
| ENSG00000145555 | MYO10 | myosin X [Source:HGNC Symbol;Acc:HGNC:7593] | -0.54 | 0.00034 |
| ENSG00000234745 | HLA-B | major histocompatibility complex, class I, B [Source:HGNC Symbol;Acc:HGNC:4932] | 0.68 | 0.00034 |
| ENSG00000078804 | TP53INP2 | tumor protein p53 inducible nuclear protein 2 [Source:HGNC Symbol;Acc:HGNC:16104] | 0.59 | 0.00034 |
| ENSG00000111087 | GLI1 | GLI family zinc finger 1 [Source:HGNC Symbol;Acc:HGNC:4317] | 1.26 | 0.00034 |
| ENSG00000166557 | TMED3 | transmembrane p24 trafficking protein 3 [Source:HGNC Symbol;Acc:HGNC:28889] | 0.52 | 0.00034 |
| ENSG00000171105 | INSR | insulin receptor [Source:HGNC Symbol;Acc:HGNC:6091] | 0.75 | 0.00036 |
| ENSG00000080644 | CHRNA3 | cholinergic receptor nicotinic alpha 3 subunit [Source:HGNC Symbol;Acc:HGNC:1957] | 1.06 | 0.00036 |
| ENSG00000165702 | GFI1B | growth factor independent 1B transcriptional repressor [Source:HGNC Symbol;Acc:HGNC:4[illegible] | -1.13 | 0.00037 |
| ENSG00000140853 | NLRC5 | NLR family CARD domain containing 5 [Source:HGNC Symbol;Acc:HGNC:29933] | 1.02 | 0.00037 |
| ENSG00000141448 | GATA6 | GATA binding protein 6 [Source:HGNC Symbol;Acc:HGNC:4174] | 0.77 | 0.00038 |
| ENSG00000139200 | PIANP | PILR alpha associated neural protein [Source:HGNC Symbol;Acc:HGNC:25338] | 0.75 | 0.00040 |
| ENSG00000213694 | S1PR3 | sphingosine-1-phosphate receptor 3 [Source:HGNC Symbol;Acc:HGNC:3167] | 0.51 | 0.00042 |
| ENSG00000134758 | RNF138 | ring finger protein 138 [Source:HGNC Symbol;Acc:HGNC:17765] | -0.51 | 0.00043 |
| ENSG00000182534 | MXRA7 | matrix remodeling associated 7 [Source:HGNC Symbol;Acc:HGNC:7541] | 0.53 | 0.00043 |
| ENSG00000183010 | PYCR1 | pyrroline-5-carboxylate reductase 1 [Source:HGNC Symbol;Acc:HGNC:9721] | -0.66 | 0.00045 |
| ENSG00000136197 | C7orf25 | chromosome 7 open reading frame 25 [Source:HGNC Symbol;Acc:HGNC:21703] | -0.74 | 0.00046 |
| ENSG00000176658 | MYO1D | myosin ID [Source:HGNC Symbol;Acc:HGNC:7598] | 0.51 | 0.00046 |
| ENSG00000139211 | AMIGO2 | adhesion molecule with Ig like domain 2 [Source:HGNC Symbol;Acc:HGNC:24073] | 0.99 | 0.00046 |
| ENSG00000104419 | NDRG1 | N-myc downstream regulated 1 [Source:HGNC Symbol;Acc:HGNC:7679] | 1.03 | 0.00046 |
| ENSG00000135919 | SERPINE2 | serpin family E member 2 [Source:HGNC Symbol;Acc:HGNC:8951] | 0.83 | 0.00047 |
| ENSG00000263513 | FAM72C | family with sequence similarity 72 member C [Source:HGNC Symbol;Acc:HGNC:30602] | -0.66 | 0.00049 |
| ENSG00000109452 | INPP4B | inositol polyphosphate-4-phosphatase type II B [Source:HGNC Symbol;Acc:HGNC:6075] | 0.51 | 0.00049 |
| ENSG00000064666 | CNN2 | calponin 2 [Source:HGNC Symbol;Acc:HGNC:2156] | 0.74 | 0.00051 |
| ENSG00000132561 | MATN2 | matrilin 2 [Source:HGNC Symbol;Acc:HGNC:6908] | 1.53 | 0.00053 |
| ENSG00000104368 | PLAT | plasminogen activator, tissue type [Source:HGNC Symbol;Acc:HGNC:9051] | 0.73 | 0.00054 |
| ENSG00000104998 | IL27RA | interleukin 27 receptor subunit alpha [Source:HGNC Symbol;Acc:HGNC:17290] | 1.19 | 0.00055 |
| ENSG00000137727 | ARHGAP20 | Rho GTPase activating protein 20 [Source:HGNC Symbol;Acc:HGNC:18357] | -0.76 | 0.00055 |
| ENSG00000114923 | SLC4A3 | solute carrier family 4 member 3 [Source:HGNC Symbol;Acc:HGNC:11029] | 0.61 | 0.00058 |
| ENSG00000171724 | VAT1L | vesicle amine transport 1 like [Source:HGNC Symbol;Acc:HGNC:29315] | 0.87 | 0.00058 |
| ENSG00000253304 | TMEM200B | transmembrane protein 200B [Source:HGNC Symbol;Acc:HGNC:33785] | 0.87 | 0.00064 |
| ENSG00000127955 | GNAI1 | G protein subunit alpha i1 [Source:HGNC Symbol;Acc:HGNC:4384] | 0.59 | 0.00065 |
| ENSG00000148123 | PLPPR1 | phospholipid phosphatase related 1 [Source:HGNC Symbol;Acc:HGNC:25993] | -0.57 | 0.00066 |
| ENSG00000274180 | NATD1 | N-acetyltransferase domain containing 1 [Source:HGNC Symbol;Acc:HGNC:30770] | 0.54 | 0.00067 |
| ENSG00000171812 | COL8A2 | collagen type VIII alpha 2 chain [Source:HGNC Symbol;Acc:HGNC:2216] | 1.22 | 0.00068 |

| | | | | |
|---|---|---|---|---|
| ENSG00000123600 | METTL8 | methyltransferase like 8 [Source:HGNC Symbol;Acc:HGNC:25856] | -0.72 | 0.00068 |
| ENSG00000164742 | ADCY1 | adenylate cyclase 1 [Source:HGNC Symbol;Acc:HGNC:232] | 1.06 | 0.00070 |
| ENSG00000166917 | MIR202HG | MIR202 host gene [Source:HGNC Symbol;Acc:HGNC:49402] | -1.21 | 0.00071 |
| ENSG00000280734 | LINC01232 | long intergenic non-protein coding RNA 1232 [Source:HGNC Symbol;Acc:HGNC:49755] | -0.56 | 0.00072 |
| ENSG00000060709 | RIMBP2 | RIMS binding protein 2 [Source:HGNC Symbol;Acc:HGNC:30339] | 0.99 | 0.00073 |
| ENSG00000136378 | ADAMTS7 | ADAM metallopeptidase with thrombospondin type 1 motif 7 [Source:HGNC Symbol;Acc:HG | 0.92 | 0.00073 |
| ENSG00000165449 | SLC16A9 | solute carrier family 16 member 9 [Source:HGNC Symbol;Acc:HGNC:23520] | -1.21 | 0.00073 |
| ENSG00000274070 | CASTOR2 | cytosolic arginine sensor for mTORC1 subunit 2 [Source:HGNC Symbol;Acc:HGNC:37073] | 0.68 | 0.00075 |
| ENSG00000167434 | CA4 | carbonic anhydrase 4 [Source:HGNC Symbol;Acc:HGNC:1375] | -0.92 | 0.00077 |
| ENSG00000213080 | NA | spermine synthase (SMS) pseudogene | -0.70 | 0.00078 |
| ENSG00000095794 | CREM | cAMP responsive element modulator [Source:HGNC Symbol;Acc:HGNC:2352] | 0.51 | 0.00078 |
| ENSG00000148082 | SHC3 | SHC adaptor protein 3 [Source:HGNC Symbol;Acc:HGNC:18181] | 1.09 | 0.00084 |
| ENSG00000119599 | DCAF4 | DDB1 and CUL4 associated factor 4 [Source:HGNC Symbol;Acc:HGNC:20229] | -0.69 | 0.00084 |
| ENSG00000099250 | NRP1 | neuropilin 1 [Source:HGNC Symbol;Acc:HGNC:8004] | 0.85 | 0.00084 |
| ENSG00000155754 | C2CD6 | C2 calcium dependent domain containing 6 [Source:HGNC Symbol;Acc:HGNC:14438] | -0.84 | 0.00084 |
| ENSG00000089639 | GMIP | GEM interacting protein [Source:HGNC Symbol;Acc:HGNC:24852] | 0.79 | 0.00084 |
| ENSG00000101210 | EEF1A2 | eukaryotic translation elongation factor 1 alpha 2 [Source:HGNC Symbol;Acc:HGNC:3192] | 0.64 | 0.00085 |
| ENSG00000172379 | ARNT2 | aryl hydrocarbon receptor nuclear translocator 2 [Source:HGNC Symbol;Acc:HGNC:16876] | 0.58 | 0.00087 |
| ENSG00000082269 | FAM135A | family with sequence similarity 135 member A [Source:HGNC Symbol;Acc:HGNC:21084] | 0.59 | 0.00088 |
| ENSG00000182197 | EXT1 | exostosin glycosyltransferase 1 [Source:HGNC Symbol;Acc:HGNC:3512] | 0.64 | 0.00094 |
| ENSG00000116649 | SRM | spermidine synthase [Source:HGNC Symbol;Acc:HGNC:11296] | -0.52 | 0.00095 |
| ENSG00000118407 | FILIP1 | filamin A interacting protein 1 [Source:HGNC Symbol;Acc:HGNC:21015] | -0.68 | 0.00095 |
| ENSG00000083535 | PIBF1 | progesterone immunomodulatory binding factor 1 [Source:HGNC Symbol;Acc:HGNC:23352 | -0.65 | 0.00101 |
| ENSG00000180875 | GREM2 | gremlin 2, DAN family BMP antagonist [Source:HGNC Symbol;Acc:HGNC:17655] | 1.29 | 0.00101 |
| ENSG00000111432 | FZD10 | frizzled class receptor 10 [Source:HGNC Symbol;Acc:HGNC:4039] | 1.49 | 0.00105 |
| ENSG00000157782 | CABP1 | calcium binding protein 1 [Source:HGNC Symbol;Acc:HGNC:1384] | 0.58 | 0.00105 |
| ENSG00000164442 | CITED2 | Cbp/p300 interacting transactivator with Glu/Asp rich carboxy-terminal domain 2 [Source:HG | 0.65 | 0.00109 |
| ENSG00000118777 | ABCG2 | ATP binding cassette subfamily G member 2 (Junior blood group) [Source:HGNC Symbol;Ac | 2.97 | 0.00109 |
| ENSG00000091409 | ITGA6 | integrin subunit alpha 6 [Source:HGNC Symbol;Acc:HGNC:6142] | 0.66 | 0.00111 |
| ENSG00000152669 | CCNO | cyclin O [Source:HGNC Symbol;Acc:HGNC:18576] | 0.88 | 0.00112 |
| ENSG00000065600 | PACC1 | proton activated chloride channel 1 [Source:HGNC Symbol;Acc:HGNC:25593] | 0.71 | 0.00113 |
| ENSG00000133800 | LYVE1 | lymphatic vessel endothelial hyaluronan receptor 1 [Source:HGNC Symbol;Acc:HGNC:1468 | 0.94 | 0.00116 |
| ENSG00000172889 | EGFL7 | EGF like domain multiple 7 [Source:HGNC Symbol;Acc:HGNC:20594] | 0.72 | 0.00118 |
| ENSG00000233485 | FHAD1-AS1 | FHAD1 antisense RNA 1 [Source:HGNC Symbol;Acc:HGNC:41241] | -0.84 | 0.00120 |
| ENSG00000173281 | PPP1R3B | protein phosphatase 1 regulatory subunit 3B [Source:HGNC Symbol;Acc:HGNC:14942] | -0.88 | 0.00121 |
| ENSG00000067533 | RRP15 | ribosomal RNA processing 15 homolog [Source:HGNC Symbol;Acc:HGNC:24255] | -0.62 | 0.00121 |
| ENSG00000110723 | EXPH5 | exophilin 5 [Source:HGNC Symbol;Acc:HGNC:30578] | 0.53 | 0.00122 |
| ENSG00000151623 | NR3C2 | nuclear receptor subfamily 3 group C member 2 [Source:HGNC Symbol;Acc:HGNC:7979] | -1.72 | 0.00122 |
| ENSG00000169508 | GPR183 | G protein-coupled receptor 183 [Source:HGNC Symbol;Acc:HGNC:3128] | -0.94 | 0.00125 |
| ENSG00000105875 | WDR91 | WD repeat domain 91 [Source:HGNC Symbol;Acc:HGNC:24997] | 0.72 | 0.00128 |

| | | | | |
|---|---|---|---|---|
| ENSG00000153930 | ANKFN1 | ankyrin repeat and fibronectin type III domain containing 1 [Source:HGNC Symbol;Acc:HGN( | -0.76 | 0.00128 |
| ENSG00000140006 | WDR89 | WD repeat domain 89 [Source:HGNC Symbol;Acc:HGNC:20489] | -0.53 | 0.00132 |
| ENSG00000145626 | UGT3A1 | UDP glycosyltransferase family 3 member A1 [Source:HGNC Symbol;Acc:HGNC:26625] | 1.34 | 0.00132 |
| ENSG00000140279 | DUOX2 | dual oxidase 2 [Source:HGNC Symbol;Acc:HGNC:13273] | 1.17 | 0.00133 |
| ENSG00000130158 | DOCK6 | dedicator of cytokinesis 6 [Source:HGNC Symbol;Acc:HGNC:19189] | 1.12 | 0.00134 |
| ENSG00000281398 | SNHG4 | small nucleolar RNA host gene 4 [Source:HGNC Symbol;Acc:HGNC:32964] | -0.93 | 0.00135 |
| ENSG00000165474 | GJB2 | gap junction protein beta 2 [Source:HGNC Symbol;Acc:HGNC:4284] | 1.18 | 0.00137 |
| ENSG00000074935 | TUBE1 | tubulin epsilon 1 [Source:HGNC Symbol;Acc:HGNC:20775] | -0.50 | 0.00138 |
| ENSG00000168765 | GSTM4 | glutathione S-transferase mu 4 [Source:HGNC Symbol;Acc:HGNC:4636] | -0.53 | 0.00142 |
| ENSG00000175600 | SUGCT | succinyl-CoA:glutarate-CoA transferase [Source:HGNC Symbol;Acc:HGNC:16001] | 1.69 | 0.00143 |
| ENSG00000122550 | KLHL7 | kelch like family member 7 [Source:HGNC Symbol;Acc:HGNC:15646] | 0.57 | 0.00145 |
| ENSG00000170779 | CDCA4 | cell division cycle associated 4 [Source:HGNC Symbol;Acc:HGNC:14625] | -0.50 | 0.00146 |
| ENSG00000136542 | GALNT5 | polypeptide N-acetylgalactosaminyltransferase 5 [Source:HGNC Symbol;Acc:HGNC:4127] | 2.21 | 0.00148 |
| ENSG00000158792 | SPATA2L | spermatogenesis associated 2 like [Source:HGNC Symbol;Acc:HGNC:28393] | 0.52 | 0.00148 |
| ENSG00000273419 | NA | novel transcript, antisense to ZNF862 | 0.84 | 0.00148 |
| ENSG00000108797 | CNTNAP1 | contactin associated protein 1 [Source:HGNC Symbol;Acc:HGNC:8011] | 0.62 | 0.00151 |
| ENSG00000103034 | NDRG4 | NDRG family member 4 [Source:HGNC Symbol;Acc:HGNC:14466] | 1.11 | 0.00154 |
| ENSG00000104774 | MAN2B1 | mannosidase alpha class 2B member 1 [Source:HGNC Symbol;Acc:HGNC:6826] | 0.60 | 0.00154 |
| ENSG00000101384 | JAG1 | jagged canonical Notch ligand 1 [Source:HGNC Symbol;Acc:HGNC:6188] | 1.00 | 0.00155 |
| ENSG00000008283 | CYB561 | cytochrome b561 [Source:HGNC Symbol;Acc:HGNC:2571] | 0.55 | 0.00155 |
| ENSG00000131668 | BARX1 | BARX homeobox 1 [Source:HGNC Symbol;Acc:HGNC:955] | 1.10 | 0.00155 |
| ENSG00000117877 | CD3EAP | CD3e molecule associated protein [Source:HGNC Symbol;Acc:HGNC:24219] | -0.59 | 0.00158 |
| ENSG00000223947 | NA | novel transcript | 1.20 | 0.00159 |
| ENSG00000132938 | MTUS2 | microtubule associated scaffold protein 2 [Source:HGNC Symbol;Acc:HGNC:20595] | 1.11 | 0.00162 |
| ENSG00000128596 | CCDC136 | coiled-coil domain containing 136 [Source:HGNC Symbol;Acc:HGNC:22225] | 0.61 | 0.00163 |
| ENSG00000131979 | GCH1 | GTP cyclohydrolase 1 [Source:HGNC Symbol;Acc:HGNC:4193] | 0.57 | 0.00163 |
| ENSG00000159387 | IRX6 | iroquois homeobox 6 [Source:HGNC Symbol;Acc:HGNC:14675] | 0.52 | 0.00166 |
| ENSG00000187140 | FOXD3 | forkhead box D3 [Source:HGNC Symbol;Acc:HGNC:3804] | 1.03 | 0.00172 |
| ENSG00000158106 | RHPN1 | rhophilin Rho GTPase binding protein 1 [Source:HGNC Symbol;Acc:HGNC:19973] | 0.63 | 0.00176 |
| ENSG00000120868 | APAF1 | apoptotic peptidase activating factor 1 [Source:HGNC Symbol;Acc:HGNC:576] | 0.53 | 0.00178 |
| ENSG00000158292 | GPR153 | G protein-coupled receptor 153 [Source:HGNC Symbol;Acc:HGNC:23618] | 0.58 | 0.00179 |
| ENSG00000255561 | FDXACB1 | ferredoxin-fold anticodon binding domain containing 1 [Source:HGNC Symbol;Acc:HGNC:2[illegible] | -1.04 | 0.00180 |
| ENSG00000136943 | CTSV | cathepsin V [Source:HGNC Symbol;Acc:HGNC:2538] | 0.89 | 0.00181 |
| ENSG00000135119 | RNFT2 | ring finger protein, transmembrane 2 [Source:HGNC Symbol;Acc:HGNC:25905] | 0.61 | 0.00182 |
| ENSG00000147378 | FATE1 | fetal and adult testis expressed 1 [Source:HGNC Symbol;Acc:HGNC:24683] | -0.79 | 0.00182 |
| ENSG00000183779 | ZNF703 | zinc finger protein 703 [Source:HGNC Symbol;Acc:HGNC:25883] | -0.51 | 0.00183 |
| ENSG00000147174 | GCNA | germ cell nuclear acidic peptidase [Source:HGNC Symbol;Acc:HGNC:15805] | 1.61 | 0.00185 |
| ENSG00000139344 | AMDHD1 | amidohydrolase domain containing 1 [Source:HGNC Symbol;Acc:HGNC:28577] | 0.66 | 0.00185 |
| ENSG00000184408 | KCND2 | potassium voltage-gated channel subfamily D member 2 [Source:HGNC Symbol;Acc:HGNC | 3.72 | 0.00188 |
| ENSG00000197329 | PELI1 | pellino E3 ubiquitin protein ligase 1 [Source:HGNC Symbol;Acc:HGNC:8827] | 1.01 | 0.00192 |

| | | | | |
|---|---|---|---|---|
| ENSG00000196562 | SULF2 | sulfatase 2 [Source:HGNC Symbol;Acc:HGNC:20392] | -1.08 | 0.00198 |
| ENSG00000112972 | HMGCS1 | 3-hydroxy-3-methylglutaryl-CoA synthase 1 [Source:HGNC Symbol;Acc:HGNC:5007] | -0.51 | 0.00199 |
| ENSG00000267432 | DNAH17-AS1 | DNAH17 antisense RNA 1 [Source:HGNC Symbol;Acc:HGNC:48594] | 1.03 | 0.00201 |
| ENSG00000169862 | CTNND2 | catenin delta 2 [Source:HGNC Symbol;Acc:HGNC:2516] | 2.40 | 0.00207 |
| ENSG00000121753 | ADGRB2 | adhesion G protein-coupled receptor B2 [Source:HGNC Symbol;Acc:HGNC:944] | 0.62 | 0.00211 |
| ENSG00000119771 | KLHL29 | kelch like family member 29 [Source:HGNC Symbol;Acc:HGNC:29404] | 0.55 | 0.00215 |
| ENSG00000286190 | NA | uncharacterized LOC728392 [Source:NCBI gene (formerly Entrezgene);Acc:728392] | 0.69 | 0.00218 |
| ENSG00000131951 | LRRC9 | leucine rich repeat containing 9 [Source:HGNC Symbol;Acc:HGNC:19848] | -0.98 | 0.00221 |
| ENSG00000204934 | ATP6V0E2-AS1 | ATP6V0E2 antisense RNA 1 [Source:HGNC Symbol;Acc:HGNC:44180] | 0.58 | 0.00223 |
| ENSG00000262769 | NA | novel transcript, antisense to SLC47A1 | 0.89 | 0.00225 |
| ENSG00000140993 | TIGD7 | tigger transposable element derived 7 [Source:HGNC Symbol;Acc:HGNC:18331] | -0.53 | 0.00234 |
| ENSG00000142494 | SLC47A1 | solute carrier family 47 member 1 [Source:HGNC Symbol;Acc:HGNC:25588] | 0.53 | 0.00237 |
| ENSG00000118971 | CCND2 | cyclin D2 [Source:HGNC Symbol;Acc:HGNC:1583] | 0.53 | 0.00237 |
| ENSG00000151090 | THRB | thyroid hormone receptor beta [Source:HGNC Symbol;Acc:HGNC:11799] | 0.52 | 0.00238 |
| ENSG00000105613 | MAST1 | microtubule associated serine/threonine kinase 1 [Source:HGNC Symbol;Acc:HGNC:19034] | 0.65 | 0.00242 |
| ENSG00000204967 | PCDHA4 | protocadherin alpha 4 [Source:HGNC Symbol;Acc:HGNC:8670] | 0.59 | 0.00243 |
| ENSG00000117152 | RGS4 | regulator of G protein signaling 4 [Source:HGNC Symbol;Acc:HGNC:10000] | 2.73 | 0.00248 |
| ENSG00000196611 | MMP1 | matrix metallopeptidase 1 [Source:HGNC Symbol;Acc:HGNC:7155] | 2.61 | 0.00248 |
| ENSG00000267278 | MAP3K14-AS1 | MAP3K14 antisense RNA 1 [Source:HGNC Symbol;Acc:HGNC:44359] | 0.60 | 0.00248 |
| ENSG00000183508 | TENT5C | terminal nucleotidyltransferase 5C [Source:HGNC Symbol;Acc:HGNC:24712] | 0.93 | 0.00256 |
| ENSG00000150594 | ADRA2A | adrenoceptor alpha 2A [Source:HGNC Symbol;Acc:HGNC:281] | -0.67 | 0.00258 |
| ENSG00000086200 | IPO11 | importin 11 [Source:HGNC Symbol;Acc:HGNC:20628] | -0.54 | 0.00259 |
| ENSG00000182901 | RGS7 | regulator of G protein signaling 7 [Source:HGNC Symbol;Acc:HGNC:10003] | 0.83 | 0.00265 |
| ENSG00000162520 | SYNC | syncoilin, intermediate filament protein [Source:HGNC Symbol;Acc:HGNC:28897] | -0.95 | 0.00271 |
| ENSG00000119125 | GDA | guanine deaminase [Source:HGNC Symbol;Acc:HGNC:4212] | 1.15 | 0.00272 |
| ENSG00000230359 | TPI1P2 | triosephosphate isomerase 1 pseudogene 2 [Source:HGNC Symbol;Acc:HGNC:38069] | -0.87 | 0.00273 |
| ENSG00000178031 | ADAMTSL1 | ADAMTS like 1 [Source:HGNC Symbol;Acc:HGNC:14632] | 0.63 | 0.00277 |
| ENSG00000172331 | BPGM | bisphosphoglycerate mutase [Source:HGNC Symbol;Acc:HGNC:1093] | 0.52 | 0.00282 |
| ENSG00000197498 | RPF2 | ribosome production factor 2 homolog [Source:HGNC Symbol;Acc:HGNC:20870] | -0.59 | 0.00285 |
| ENSG00000164188 | RANBP3L | RAN binding protein 3 like [Source:HGNC Symbol;Acc:HGNC:26353] | 0.58 | 0.00290 |
| ENSG00000130558 | OLFM1 | olfactomedin 1 [Source:HGNC Symbol;Acc:HGNC:17187] | 0.57 | 0.00292 |
| ENSG00000101665 | SMAD7 | SMAD family member 7 [Source:HGNC Symbol;Acc:HGNC:6773] | -0.85 | 0.00294 |
| ENSG00000162929 | KIAA1841 | KIAA1841 [Source:HGNC Symbol;Acc:HGNC:29387] | 0.63 | 0.00298 |
| ENSG00000153064 | BANK1 | B cell scaffold protein with ankyrin repeats 1 [Source:HGNC Symbol;Acc:HGNC:18233] | 1.56 | 0.00299 |
| ENSG00000261037 | NA | novel transcript | 0.96 | 0.00299 |
| ENSG00000170500 | LONRF2 | LON peptidase N-terminal domain and ring finger 2 [Source:HGNC Symbol;Acc:HGNC:2478 | 0.51 | 0.00302 |
| ENSG00000104524 | PYCR3 | pyrroline-5-carboxylate reductase 3 [Source:HGNC Symbol;Acc:HGNC:25846] | -0.53 | 0.00302 |
| ENSG00000163884 | KLF15 | Kruppel like factor 15 [Source:HGNC Symbol;Acc:HGNC:14536] | 0.78 | 0.00325 |
| ENSG00000031081 | ARHGAP31 | Rho GTPase activating protein 31 [Source:HGNC Symbol;Acc:HGNC:29216] | 0.84 | 0.00327 |
| ENSG00000105894 | PTN | pleiotrophin [Source:HGNC Symbol;Acc:HGNC:9630] | 1.41 | 0.00335 |

| | | | | |
|---|---|---|---|---|
| ENSG00000100604 | CHGA | chromogranin A [Source:HGNC Symbol;Acc:HGNC:1929] | -0.89 | 0.00341 |
| ENSG00000151693 | ASAP2 | ArfGAP with SH3 domain, ankyrin repeat and PH domain 2 [Source:HGNC Symbol;Acc:HGN | 0.57 | 0.00348 |
| ENSG00000170370 | EMX2 | empty spiracles homeobox 2 [Source:HGNC Symbol;Acc:HGNC:3341] | 1.36 | 0.00350 |
| ENSG00000182255 | KCNA4 | potassium voltage-gated channel subfamily A member 4 [Source:HGNC Symbol;Acc:HGNC: | 0.81 | 0.00353 |
| ENSG00000118257 | NRP2 | neuropilin 2 [Source:HGNC Symbol;Acc:HGNC:8005] | 0.63 | 0.00354 |
| ENSG00000179111 | HES7 | hes family bHLH transcription factor 7 [Source:HGNC Symbol;Acc:HGNC:15977] | 1.46 | 0.00355 |
| ENSG00000124701 | APOBEC2 | apolipoprotein B mRNA editing enzyme catalytic subunit 2 [Source:HGNC Symbol;Acc:HGN | 1.49 | 0.00361 |
| ENSG00000115159 | GPD2 | glycerol-3-phosphate dehydrogenase 2 [Source:HGNC Symbol;Acc:HGNC:4456] | 0.53 | 0.00361 |
| ENSG00000079257 | LXN | latexin [Source:HGNC Symbol;Acc:HGNC:13347] | 0.65 | 0.00365 |
| ENSG00000169689 | CENPX | centromere protein X [Source:HGNC Symbol;Acc:HGNC:11422] | -0.58 | 0.00369 |
| ENSG00000100949 | RABGGTA | Rab geranylgeranyltransferase subunit alpha [Source:HGNC Symbol;Acc:HGNC:9795] | 0.51 | 0.00372 |
| ENSG00000163359 | COL6A3 | collagen type VI alpha 3 chain [Source:HGNC Symbol;Acc:HGNC:2213] | -2.14 | 0.00374 |
| ENSG00000285508 | NA | novel protein | -7.28 | 0.00376 |
| ENSG00000159212 | CLIC6 | chloride intracellular channel 6 [Source:HGNC Symbol;Acc:HGNC:2065] | 1.07 | 0.00378 |
| ENSG00000167995 | BEST1 | bestrophin 1 [Source:HGNC Symbol;Acc:HGNC:12703] | -0.88 | 0.00382 |
| ENSG00000272275 | NA | novel transcript | 1.02 | 0.00388 |
| ENSG00000180287 | PLD5 | phospholipase D family member 5 [Source:HGNC Symbol;Acc:HGNC:26879] | 0.65 | 0.00388 |
| ENSG00000198846 | TOX | thymocyte selection associated high mobility group box [Source:HGNC Symbol;Acc:HGNC:1 | 0.87 | 0.00404 |
| ENSG00000123213 | NLN | neurolysin [Source:HGNC Symbol;Acc:HGNC:16058] | -0.65 | 0.00405 |
| ENSG00000166793 | YPEL4 | yippee like 4 [Source:HGNC Symbol;Acc:HGNC:18328] | 0.86 | 0.00406 |
| ENSG00000259291 | ZNF710-AS1 | ZNF710 antisense RNA 1 [Source:HGNC Symbol;Acc:HGNC:53141] | 1.07 | 0.00411 |
| ENSG00000285476 | NA | novel transcript | 0.67 | 0.00414 |
| ENSG00000125148 | MT2A | metallothionein 2A [Source:HGNC Symbol;Acc:HGNC:7406] | 0.55 | 0.00420 |
| ENSG00000139354 | GAS2L3 | growth arrest specific 2 like 3 [Source:HGNC Symbol;Acc:HGNC:27475] | -0.50 | 0.00422 |
| ENSG00000154639 | CXADR | CXADR Ig-like cell adhesion molecule [Source:HGNC Symbol;Acc:HGNC:2559] | 0.56 | 0.00429 |
| ENSG00000181322 | NME9 | NME/NM23 family member 9 [Source:HGNC Symbol;Acc:HGNC:21343] | 0.73 | 0.00445 |
| ENSG00000182747 | SLC35D3 | solute carrier family 35 member D3 [Source:HGNC Symbol;Acc:HGNC:15621] | 1.41 | 0.00446 |
| ENSG00000168135 | KCNJ4 | potassium inwardly rectifying channel subfamily J member 4 [Source:HGNC Symbol;Acc:HG | 0.68 | 0.00459 |
| ENSG00000171877 | FRMD5 | FERM domain containing 5 [Source:HGNC Symbol;Acc:HGNC:28214] | 1.15 | 0.00459 |
| ENSG00000165030 | NFIL3 | nuclear factor, interleukin 3 regulated [Source:HGNC Symbol;Acc:HGNC:7787] | 0.74 | 0.00483 |
| ENSG00000135502 | SLC26A10 | solute carrier family 26 member 10 [Source:HGNC Symbol;Acc:HGNC:14470] | 0.90 | 0.00484 |
| ENSG00000110900 | TSPAN11 | tetraspanin 11 [Source:HGNC Symbol;Acc:HGNC:30795] | -0.53 | 0.00499 |
| ENSG00000007866 | TEAD3 | TEA domain transcription factor 3 [Source:HGNC Symbol;Acc:HGNC:11716] | 0.89 | 0.00502 |
| ENSG00000136531 | SCN2A | sodium voltage-gated channel alpha subunit 2 [Source:HGNC Symbol;Acc:HGNC:10588] | 0.55 | 0.00515 |
| ENSG00000099860 | GADD45B | growth arrest and DNA damage inducible beta [Source:HGNC Symbol;Acc:HGNC:4096] | -0.57 | 0.00527 |
| ENSG00000077009 | NMRK2 | nicotinamide riboside kinase 2 [Source:HGNC Symbol;Acc:HGNC:17871] | 1.04 | 0.00528 |
| ENSG00000182463 | TSHZ2 | teashirt zinc finger homeobox 2 [Source:HGNC Symbol;Acc:HGNC:13010] | 0.76 | 0.00543 |
| ENSG00000143416 | SELENBP1 | selenium binding protein 1 [Source:HGNC Symbol;Acc:HGNC:10719] | 0.56 | 0.00572 |
| ENSG00000173846 | PLK3 | polo like kinase 3 [Source:HGNC Symbol;Acc:HGNC:2154] | 0.84 | 0.00576 |
| ENSG00000244968 | LIFR-AS1 | LIFR antisense RNA 1 [Source:HGNC Symbol;Acc:HGNC:43600] | -0.55 | 0.00583 |

| | | | | |
|---|---|---|---|---|
| ENSG00000119915 | ELOVL3 | ELOVL fatty acid elongase 3 [Source:HGNC Symbol;Acc:HGNC:18047] | -0.73 | 0.00597 |
| ENSG00000152377 | SPOCK1 | SPARC (osteonectin), cwcv and kazal like domains proteoglycan 1 [Source:HGNC Symbol;A | -0.70 | 0.00613 |
| ENSG00000145920 | CPLX2 | complexin 2 [Source:HGNC Symbol;Acc:HGNC:2310] | -1.52 | 0.00618 |
| ENSG00000118514 | ALDH8A1 | aldehyde dehydrogenase 8 family member A1 [Source:HGNC Symbol;Acc:HGNC:15471] | 0.67 | 0.00623 |
| ENSG00000180447 | GAS1 | growth arrest specific 1 [Source:HGNC Symbol;Acc:HGNC:4165] | 0.90 | 0.00625 |
| ENSG00000215440 | NPEPL1 | aminopeptidase like 1 [Source:HGNC Symbol;Acc:HGNC:16244] | 0.63 | 0.00630 |
| ENSG00000106034 | CPED1 | cadherin like and PC-esterase domain containing 1 [Source:HGNC Symbol;Acc:HGNC:2615 | 1.10 | 0.00636 |
| ENSG00000268996 | MAN1B1-DT | MAN1B1 divergent transcript [Source:HGNC Symbol;Acc:HGNC:48715] | -0.54 | 0.00636 |
| ENSG00000100196 | KDELR3 | KDEL endoplasmic reticulum protein retention receptor 3 [Source:HGNC Symbol;Acc:HGNC | 0.60 | 0.00638 |
| ENSG00000132481 | TRIM47 | tripartite motif containing 47 [Source:HGNC Symbol;Acc:HGNC:19020] | -0.50 | 0.00647 |
| ENSG00000142173 | COL6A2 | collagen type VI alpha 2 chain [Source:HGNC Symbol;Acc:HGNC:2212] | 0.76 | 0.00649 |
| ENSG00000163624 | CDS1 | CDP-diacylglycerol synthase 1 [Source:HGNC Symbol;Acc:HGNC:1800] | 0.50 | 0.00662 |
| ENSG00000250565 | ATP6V1E2 | ATPase H+ transporting V1 subunit E2 [Source:HGNC Symbol;Acc:HGNC:18125] | -0.78 | 0.00669 |
| ENSG00000133687 | TMTC1 | transmembrane O-mannosyltransferase targeting cadherins 1 [Source:HGNC Symbol;Acc:H | 0.51 | 0.00693 |
| ENSG00000173114 | LRRN3 | leucine rich repeat neuronal 3 [Source:HGNC Symbol;Acc:HGNC:17200] | -1.90 | 0.00696 |
| ENSG00000112378 | PERP | p53 apoptosis effector related to PMP22 [Source:HGNC Symbol;Acc:HGNC:17637] | 0.53 | 0.00696 |
| ENSG00000237651 | C2orf74 | chromosome 2 open reading frame 74 [Source:HGNC Symbol;Acc:HGNC:34439] | -0.66 | 0.00707 |
| ENSG00000273038 | NA | novel transcript | 0.96 | 0.00725 |
| ENSG00000147234 | FRMPD3 | FERM and PDZ domain containing 3 [Source:HGNC Symbol;Acc:HGNC:29382] | 0.97 | 0.00725 |
| ENSG00000164076 | CAMKV | CaM kinase like vesicle associated [Source:HGNC Symbol;Acc:HGNC:28788] | 0.98 | 0.00725 |
| ENSG00000265972 | TXNIP | thioredoxin interacting protein [Source:HGNC Symbol;Acc:HGNC:16952] | -1.13 | 0.00744 |
| ENSG00000163644 | PPM1K | protein phosphatase, Mg2+/Mn2+ dependent 1K [Source:HGNC Symbol;Acc:HGNC:25415] | 0.56 | 0.00751 |
| ENSG00000198598 | MMP17 | matrix metallopeptidase 17 [Source:HGNC Symbol;Acc:HGNC:7163] | 1.20 | 0.00756 |
| ENSG00000168389 | MFSD2A | major facilitator superfamily domain containing 2A [Source:HGNC Symbol;Acc:HGNC:25897 | 1.18 | 0.00757 |
| ENSG00000146674 | IGFBP3 | insulin like growth factor binding protein 3 [Source:HGNC Symbol;Acc:HGNC:5472] | 1.10 | 0.00771 |
| ENSG00000166068 | SPRED1 | sprouty related EVH1 domain containing 1 [Source:HGNC Symbol;Acc:HGNC:20249] | 0.68 | 0.00771 |
| ENSG00000105609 | LILRB5 | leukocyte immunoglobulin like receptor B5 [Source:HGNC Symbol;Acc:HGNC:6609] | 0.96 | 0.00772 |
| ENSG00000078725 | BRINP1 | BMP/retinoic acid inducible neural specific 1 [Source:HGNC Symbol;Acc:HGNC:2687] | 1.05 | 0.00772 |
| ENSG00000106546 | AHR | aryl hydrocarbon receptor [Source:HGNC Symbol;Acc:HGNC:348] | 0.64 | 0.00775 |
| ENSG00000134013 | LOXL2 | lysyl oxidase like 2 [Source:HGNC Symbol;Acc:HGNC:6666] | 0.69 | 0.00775 |
| ENSG00000261046 | C2orf69P1 | chromosome 2 open reading frame 69 pseudogene 1 [Source:HGNC Symbol;Acc:HGNC:51 | -1.15 | 0.00787 |
| ENSG00000135454 | B4GALNT1 | beta-1,4-N-acetyl-galactosaminyltransferase 1 [Source:HGNC Symbol;Acc:HGNC:4117] | 0.59 | 0.00787 |
| ENSG00000149212 | SESN3 | sestrin 3 [Source:HGNC Symbol;Acc:HGNC:23060] | 0.51 | 0.00787 |
| ENSG00000165895 | ARHGAP42 | Rho GTPase activating protein 42 [Source:HGNC Symbol;Acc:HGNC:26545] | 0.52 | 0.00790 |
| ENSG00000128203 | ASPHD2 | aspartate beta-hydroxylase domain containing 2 [Source:HGNC Symbol;Acc:HGNC:30437] | 0.82 | 0.00801 |
| ENSG00000233175 | NA | novel transcript, antisense to FMNL1 | 0.93 | 0.00802 |
| ENSG00000005961 | ITGA2B | integrin subunit alpha 2b [Source:HGNC Symbol;Acc:HGNC:6138] | 1.16 | 0.00803 |
| ENSG00000091490 | SEL1L3 | SEL1L family member 3 [Source:HGNC Symbol;Acc:HGNC:29108] | 0.94 | 0.00807 |
| ENSG00000129946 | SHC2 | SHC adaptor protein 2 [Source:HGNC Symbol;Acc:HGNC:29869] | 0.71 | 0.00817 |
| ENSG00000100290 | BIK | BCL2 interacting killer [Source:HGNC Symbol;Acc:HGNC:1051] | 0.74 | 0.00817 |

| | | | | |
|---|---|---|---|---|
| ENSG00000126814 | TRMT5 | tRNA methyltransferase 5 [Source:HGNC Symbol;Acc:HGNC:23141] | -0.80 | 0.00827 |
| ENSG00000229320 | KRT8P12 | keratin 8 pseudogene 12 [Source:HGNC Symbol;Acc:HGNC:28057] | -0.58 | 0.00845 |
| ENSG00000223749 | MIR503HG | MIR503 host gene [Source:HGNC Symbol;Acc:HGNC:28258] | 0.56 | 0.00845 |
| ENSG00000236501 | NA | novel transcript | 0.83 | 0.00887 |
| ENSG00000093217 | XYLB | xylulokinase [Source:HGNC Symbol;Acc:HGNC:12839] | -0.79 | 0.00890 |
| ENSG00000130813 | SHFL | shiftless antiviral inhibitor of ribosomal frameshifting [Source:HGNC Symbol;Acc:HGNC:256 | 0.84 | 0.00896 |
| ENSG00000160961 | ZNF333 | zinc finger protein 333 [Source:HGNC Symbol;Acc:HGNC:15624] | -0.58 | 0.00907 |
| ENSG00000233058 | LINC00884 | long intergenic non-protein coding RNA 884 [Source:HGNC Symbol;Acc:HGNC:48570] | 0.81 | 0.00910 |
| ENSG00000171488 | LRRC8C | leucine rich repeat containing 8 VRAC subunit C [Source:HGNC Symbol;Acc:HGNC:25075] | 0.53 | 0.00923 |
| ENSG00000224945 | NA | novel transcript | 1.43 | 0.00934 |
| ENSG00000287575 | NA | novel transcript | 1.43 | 0.00942 |
| ENSG00000067141 | NEO1 | neogenin 1 [Source:HGNC Symbol;Acc:HGNC:7754] | 0.69 | 0.00964 |
| ENSG00000137486 | ARRB1 | arrestin beta 1 [Source:HGNC Symbol;Acc:HGNC:711] | 1.38 | 0.00964 |
| ENSG00000170153 | RNF150 | ring finger protein 150 [Source:HGNC Symbol;Acc:HGNC:23138] | 0.77 | 0.00968 |
| ENSG00000137449 | CPEB2 | cytoplasmic polyadenylation element binding protein 2 [Source:HGNC Symbol;Acc:HGNC:2 | 0.75 | 0.00972 |
| ENSG00000148985 | PGAP2 | post-GPI attachment to proteins 2 [Source:HGNC Symbol;Acc:HGNC:17893] | -0.78 | 0.00973 |
| ENSG00000102755 | FLT1 | fms related receptor tyrosine kinase 1 [Source:HGNC Symbol;Acc:HGNC:3763] | 2.87 | 0.00980 |
| ENSG00000080709 | KCNN2 | potassium calcium-activated channel subfamily N member 2 [Source:HGNC Symbol;Acc:HG | -0.66 | 0.01002 |
| ENSG00000137714 | FDX1 | ferredoxin 1 [Source:HGNC Symbol;Acc:HGNC:3638] | 0.51 | 0.01002 |
| ENSG00000182919 | C11orf54 | chromosome 11 open reading frame 54 [Source:HGNC Symbol;Acc:HGNC:30204] | 0.53 | 0.01016 |
| ENSG00000105649 | RAB3A | RAB3A, member RAS oncogene family [Source:HGNC Symbol;Acc:HGNC:9777] | 0.73 | 0.01056 |
| ENSG00000169744 | LDB2 | LIM domain binding 2 [Source:HGNC Symbol;Acc:HGNC:6533] | -0.62 | 0.01057 |
| ENSG00000166289 | PLEKHF1 | pleckstrin homology and FYVE domain containing 1 [Source:HGNC Symbol;Acc:HGNC:207 | 0.68 | 0.01086 |
| ENSG00000137501 | SYTL2 | synaptotagmin like 2 [Source:HGNC Symbol;Acc:HGNC:15585] | -0.56 | 0.01088 |
| ENSG00000180758 | GPR157 | G protein-coupled receptor 157 [Source:HGNC Symbol;Acc:HGNC:23687] | -0.77 | 0.01113 |
| ENSG00000176490 | DIRAS1 | DIRAS family GTPase 1 [Source:HGNC Symbol;Acc:HGNC:19127] | 0.51 | 0.01132 |
| ENSG00000185519 | FAM131C | family with sequence similarity 131 member C [Source:HGNC Symbol;Acc:HGNC:26717] | 0.54 | 0.01132 |
| ENSG00000128965 | CHAC1 | ChaC glutathione specific gamma-glutamylcyclotransferase 1 [Source:HGNC Symbol;Acc:H | -0.63 | 0.01133 |
| ENSG00000187902 | SHISA7 | shisa family member 7 [Source:HGNC Symbol;Acc:HGNC:35409] | 0.76 | 0.01161 |
| ENSG00000075651 | PLD1 | phospholipase D1 [Source:HGNC Symbol;Acc:HGNC:9067] | -0.53 | 0.01183 |
| ENSG00000152580 | IGSF10 | immunoglobulin superfamily member 10 [Source:HGNC Symbol;Acc:HGNC:26384] | 0.91 | 0.01193 |
| ENSG00000112541 | PDE10A | phosphodiesterase 10A [Source:HGNC Symbol;Acc:HGNC:8772] | 0.66 | 0.01209 |
| ENSG00000187678 | SPRY4 | sprouty RTK signaling antagonist 4 [Source:HGNC Symbol;Acc:HGNC:15533] | 2.24 | 0.01212 |
| ENSG00000165757 | JCAD | junctional cadherin 5 associated [Source:HGNC Symbol;Acc:HGNC:29283] | -0.86 | 0.01219 |
| ENSG00000158258 | CLSTN2 | calsyntenin 2 [Source:HGNC Symbol;Acc:HGNC:17448] | 0.99 | 0.01226 |
| ENSG00000146411 | SLC2A12 | solute carrier family 2 member 12 [Source:HGNC Symbol;Acc:HGNC:18067] | 0.70 | 0.01228 |
| ENSG00000187024 | PTRH1 | peptidyl-tRNA hydrolase 1 homolog [Source:HGNC Symbol;Acc:HGNC:27039] | -0.51 | 0.01229 |
| ENSG00000222009 | BTBD19 | BTB domain containing 19 [Source:HGNC Symbol;Acc:HGNC:27145] | 0.61 | 0.01235 |
| ENSG00000174469 | CNTNAP2 | contactin associated protein 2 [Source:HGNC Symbol;Acc:HGNC:13830] | -0.61 | 0.01241 |
| ENSG00000123364 | HOXC13 | homeobox C13 [Source:HGNC Symbol;Acc:HGNC:5125] | 0.62 | 0.01244 |

| | | | | |
|---|---|---|---|---|
| ENSG00000236841 | NA | novel transcript | 0.93 | 0.01261 |
| ENSG00000007314 | SCN4A | sodium voltage-gated channel alpha subunit 4 [Source:HGNC Symbol;Acc:HGNC:10591] | 0.79 | 0.01281 |
| ENSG00000180822 | PSMG4 | proteasome assembly chaperone 4 [Source:HGNC Symbol;Acc:HGNC:21108] | -0.51 | 0.01292 |
| ENSG00000286478 | NA | novel transcript, antisense to HIBADH | 0.86 | 0.01308 |
| ENSG00000151883 | PARP8 | poly(ADP-ribose) polymerase family member 8 [Source:HGNC Symbol;Acc:HGNC:26124] | 1.41 | 0.01331 |
| ENSG00000277027 | RMRP | RNA component of mitochondrial RNA processing endoribonuclease [Source:HGNC Symbc | -7.32 | 0.01353 |
| ENSG00000153574 | RPIA | ribose 5-phosphate isomerase A [Source:HGNC Symbol;Acc:HGNC:10297] | -0.52 | 0.01353 |
| ENSG00000128973 | CLN6 | CLN6 transmembrane ER protein [Source:HGNC Symbol;Acc:HGNC:2077] | -0.51 | 0.01376 |
| ENSG00000144290 | SLC4A10 | solute carrier family 4 member 10 [Source:HGNC Symbol;Acc:HGNC:13811] | 0.70 | 0.01378 |
| ENSG00000054356 | PTPRN | protein tyrosine phosphatase receptor type N [Source:HGNC Symbol;Acc:HGNC:9676] | 1.07 | 0.01380 |
| ENSG00000037897 | METTL1 | methyltransferase like 1 [Source:HGNC Symbol;Acc:HGNC:7030] | -0.53 | 0.01386 |
| ENSG00000100418 | DESI1 | desumoylating isopeptidase 1 [Source:HGNC Symbol;Acc:HGNC:24577] | -0.51 | 0.01386 |
| ENSG00000145147 | SLIT2 | slit guidance ligand 2 [Source:HGNC Symbol;Acc:HGNC:11086] | 0.54 | 0.01388 |
| ENSG00000105289 | TJP3 | tight junction protein 3 [Source:HGNC Symbol;Acc:HGNC:11829] | 0.76 | 0.01415 |
| ENSG00000165810 | BTNL9 | butyrophilin like 9 [Source:HGNC Symbol;Acc:HGNC:24176] | 0.78 | 0.01415 |
| ENSG00000163431 | LMOD1 | leiomodin 1 [Source:HGNC Symbol;Acc:HGNC:6647] | -0.56 | 0.01429 |
| ENSG00000279875 | NA | TEC | 1.26 | 0.01449 |
| ENSG00000267302 | RNFT1-DT | RNFT1 divergent transcript [Source:HGNC Symbol;Acc:HGNC:51346] | 1.25 | 0.01461 |
| ENSG00000101349 | PAK5 | p21 (RAC1) activated kinase 5 [Source:HGNC Symbol;Acc:HGNC:15916] | -0.75 | 0.01465 |
| ENSG00000139318 | DUSP6 | dual specificity phosphatase 6 [Source:HGNC Symbol;Acc:HGNC:3072] | 0.83 | 0.01467 |
| ENSG00000254692 | NA | novel protein | -8.32 | 0.01467 |
| ENSG00000224389 | C4B | complement C4B (Chido blood group) [Source:HGNC Symbol;Acc:HGNC:1324] | 0.70 | 0.01473 |
| ENSG00000168993 | CPLX1 | complexin 1 [Source:HGNC Symbol;Acc:HGNC:2309] | -0.67 | 0.01488 |
| ENSG00000167535 | CACNB3 | calcium voltage-gated channel auxiliary subunit beta 3 [Source:HGNC Symbol;Acc:HGNC:1 | 0.52 | 0.01494 |
| ENSG00000204677 | FAM153CP | family with sequence similarity 153 member C, pseudogene [Source:HGNC Symbol;Acc:HGI | 0.61 | 0.01515 |
| ENSG00000187210 | GCNT1 | glucosaminyl (N-acetyl) transferase 1 [Source:HGNC Symbol;Acc:HGNC:4203] | 1.22 | 0.01533 |
| ENSG00000105717 | PBX4 | PBX homeobox 4 [Source:HGNC Symbol;Acc:HGNC:13403] | 0.78 | 0.01538 |
| ENSG00000248538 | NA | novel transcript | -0.54 | 0.01540 |
| ENSG00000198873 | GRK5 | G protein-coupled receptor kinase 5 [Source:HGNC Symbol;Acc:HGNC:4544] | 1.04 | 0.01550 |
| ENSG00000107807 | TLX1 | T cell leukemia homeobox 1 [Source:HGNC Symbol;Acc:HGNC:5056] | 1.11 | 0.01592 |
| ENSG00000166105 | GLB1L3 | galactosidase beta 1 like 3 [Source:HGNC Symbol;Acc:HGNC:25147] | 0.60 | 0.01592 |
| ENSG00000185561 | TLCD2 | TLC domain containing 2 [Source:HGNC Symbol;Acc:HGNC:33522] | 0.55 | 0.01592 |
| ENSG00000112276 | BVES | blood vessel epicardial substance [Source:HGNC Symbol;Acc:HGNC:1152] | 1.04 | 0.01655 |
| ENSG00000139549 | DHH | desert hedgehog signaling molecule [Source:HGNC Symbol;Acc:HGNC:2865] | 1.11 | 0.01673 |
| ENSG00000197320 | NA | C3 and PZP-like, alpha-2-macroglobulin domain containing 8 (CPAMD8) pseudogene | 0.76 | 0.01688 |
| ENSG00000198208 | RPS6KL1 | ribosomal protein S6 kinase like 1 [Source:HGNC Symbol;Acc:HGNC:20222] | 0.89 | 0.01688 |
| ENSG00000160796 | NBEAL2 | neurobeachin like 2 [Source:HGNC Symbol;Acc:HGNC:31928] | 0.54 | 0.01698 |
| ENSG00000139629 | GALNT6 | polypeptide N-acetylgalactosaminyltransferase 6 [Source:HGNC Symbol;Acc:HGNC:4128] | 0.57 | 0.01704 |
| ENSG00000158201 | ABHD3 | abhydrolase domain containing 3, phospholipase [Source:HGNC Symbol;Acc:HGNC:18718 | 0.56 | 0.01704 |
| ENSG00000179598 | PLD6 | phospholipase D family member 6 [Source:HGNC Symbol;Acc:HGNC:30447] | -0.60 | 0.01704 |

| | | | | |
|---|---|---|---|---|
| ENSG00000153234 | NR4A2 | nuclear receptor subfamily 4 group A member 2 [Source:HGNC Symbol;Acc:HGNC:7981] | 1.65 | 0.01710 |
| ENSG00000177606 | JUN | Jun proto-oncogene, AP-1 transcription factor subunit [Source:HGNC Symbol;Acc:HGNC:62 | -0.67 | 0.01754 |
| ENSG00000176531 | PHLDB3 | pleckstrin homology like domain family B member 3 [Source:HGNC Symbol;Acc:HGNC:304 | 0.69 | 0.01768 |
| ENSG00000162745 | OLFML2B | olfactomedin like 2B [Source:HGNC Symbol;Acc:HGNC:24558] | -0.76 | 0.01794 |
| ENSG00000168938 | PPIC | peptidylprolyl isomerase C [Source:HGNC Symbol;Acc:HGNC:9256] | 0.62 | 0.01820 |
| ENSG00000075340 | ADD2 | adducin 2 [Source:HGNC Symbol;Acc:HGNC:244] | 0.79 | 0.01825 |
| ENSG00000163131 | CTSS | cathepsin S [Source:HGNC Symbol;Acc:HGNC:2545] | -0.66 | 0.01825 |
| ENSG00000142621 | FHAD1 | forkhead associated phosphopeptide binding domain 1 [Source:HGNC Symbol;Acc:HGNC:2 | -0.61 | 0.01826 |
| ENSG00000130768 | SMPDL3B | sphingomyelin phosphodiesterase acid like 3B [Source:HGNC Symbol;Acc:HGNC:21416] | 0.68 | 0.01834 |
| ENSG00000236675 | MTX1P1 | metaxin 1 pseudogene 1 [Source:HGNC Symbol;Acc:HGNC:7505] | 1.55 | 0.01834 |
| ENSG00000251562 | MALAT1 | metastasis associated lung adenocarcinoma transcript 1 [Source:HGNC Symbol;Acc:HGNC: | 0.57 | 0.01842 |
| ENSG00000130589 | HELZ2 | helicase with zinc finger 2 [Source:HGNC Symbol;Acc:HGNC:30021] | 0.79 | 0.01844 |
| ENSG00000111452 | ADGRD1 | adhesion G protein-coupled receptor D1 [Source:HGNC Symbol;Acc:HGNC:19893] | 1.07 | 0.01860 |
| ENSG00000076706 | MCAM | melanoma cell adhesion molecule [Source:HGNC Symbol;Acc:HGNC:6934] | 0.88 | 0.01867 |
| ENSG00000186594 | MIR22HG | MIR22 host gene [Source:HGNC Symbol;Acc:HGNC:28219] | 0.55 | 0.01867 |
| ENSG00000213949 | ITGA1 | integrin subunit alpha 1 [Source:HGNC Symbol;Acc:HGNC:6134] | 0.50 | 0.01886 |
| ENSG00000126583 | PRKCG | protein kinase C gamma [Source:HGNC Symbol;Acc:HGNC:9402] | 0.66 | 0.01918 |
| ENSG00000099282 | TSPAN15 | tetraspanin 15 [Source:HGNC Symbol;Acc:HGNC:23298] | 0.82 | 0.01924 |
| ENSG00000198825 | INPP5F | inositol polyphosphate-5-phosphatase F [Source:HGNC Symbol;Acc:HGNC:17054] | -0.63 | 0.02034 |
| ENSG00000176092 | CRYBG2 | crystallin beta-gamma domain containing 2 [Source:HGNC Symbol;Acc:HGNC:17295] | -1.24 | 0.02046 |
| ENSG00000204020 | LIPN | lipase family member N [Source:HGNC Symbol;Acc:HGNC:23452] | 1.76 | 0.02051 |
| ENSG00000204876 | NA | uncharacterized LOC389602 [Source:NCBI gene (formerly Entrezgene);Acc:389602] | 0.68 | 0.02058 |
| ENSG00000110427 | KIAA1549L | KIAA1549 like [Source:HGNC Symbol;Acc:HGNC:24836] | 0.70 | 0.02065 |
| ENSG00000106852 | LHX6 | LIM homeobox 6 [Source:HGNC Symbol;Acc:HGNC:21735] | -0.65 | 0.02078 |
| ENSG00000132688 | NES | nestin [Source:HGNC Symbol;Acc:HGNC:7756] | 1.13 | 0.02089 |
| ENSG00000197566 | ZNF624 | zinc finger protein 624 [Source:HGNC Symbol;Acc:HGNC:29254] | -0.57 | 0.02089 |
| ENSG00000244731 | C4A | complement C4A (Rodgers blood group) [Source:HGNC Symbol;Acc:HGNC:1323] | 0.69 | 0.02140 |
| ENSG00000278921 | EPB41L4A-DT | EPB41L4A divergent transcript [Source:HGNC Symbol;Acc:HGNC:25643] | 0.53 | 0.02169 |
| ENSG00000140015 | KCNH5 | potassium voltage-gated channel subfamily H member 5 [Source:HGNC Symbol;Acc:HGNC | 1.25 | 0.02179 |
| ENSG00000280649 | NA | TEC | 0.64 | 0.02193 |
| ENSG00000064763 | FAR2 | fatty acyl-CoA reductase 2 [Source:HGNC Symbol;Acc:HGNC:25531] | 0.53 | 0.02245 |
| ENSG00000181982 | CCDC149 | coiled-coil domain containing 149 [Source:HGNC Symbol;Acc:HGNC:25405] | 0.98 | 0.02255 |
| ENSG00000231852 | CYP21A2 | cytochrome P450 family 21 subfamily A member 2 [Source:HGNC Symbol;Acc:HGNC:2600] | 0.52 | 0.02256 |
| ENSG00000100024 | UPB1 | beta-ureidopropionase 1 [Source:HGNC Symbol;Acc:HGNC:16297] | 1.30 | 0.02262 |
| ENSG00000285799 | NA | MHC class I polypeptide-related sequence F pseudogene | -2.49 | 0.02271 |
| ENSG00000105516 | DBP | D-box binding PAR bZIP transcription factor [Source:HGNC Symbol;Acc:HGNC:2697] | -0.54 | 0.02320 |
| ENSG00000149292 | TTC12 | tetratricopeptide repeat domain 12 [Source:HGNC Symbol;Acc:HGNC:23700] | -0.60 | 0.02320 |
| ENSG00000144649 | GASK1A | golgi associated kinase 1A [Source:HGNC Symbol;Acc:HGNC:24485] | 0.60 | 0.02328 |
| ENSG00000273044 | NA | novel transcript | 1.00 | 0.02334 |
| ENSG00000160469 | BRSK1 | BR serine/threonine kinase 1 [Source:HGNC Symbol;Acc:HGNC:18994] | 0.53 | 0.02348 |

| | | | | |
|---|---|---|---|---|
| ENSG00000187801 | ZFP69B | ZFP69 zinc finger protein B [Source:HGNC Symbol;Acc:HGNC:28053] | -0.75 | 0.02353 |
| ENSG00000109738 | GLRB | glycine receptor beta [Source:HGNC Symbol;Acc:HGNC:4329] | 0.57 | 0.02371 |
| ENSG00000287861 | NA | novel transcript | -0.59 | 0.02371 |
| ENSG00000161921 | CXCL16 | C-X-C motif chemokine ligand 16 [Source:HGNC Symbol;Acc:HGNC:16642] | 1.01 | 0.02406 |
| ENSG00000100036 | SLC35E4 | solute carrier family 35 member E4 [Source:HGNC Symbol;Acc:HGNC:17058] | 0.62 | 0.02423 |
| ENSG00000165695 | AK8 | adenylate kinase 8 [Source:HGNC Symbol;Acc:HGNC:26526] | -0.70 | 0.02427 |
| ENSG00000273599 | NA | novel transcript, antisense to CTBP2 | 0.64 | 0.02433 |
| ENSG00000116815 | CD58 | CD58 molecule [Source:HGNC Symbol;Acc:HGNC:1688] | 0.50 | 0.02440 |
| ENSG00000168874 | ATOH8 | atonal bHLH transcription factor 8 [Source:HGNC Symbol;Acc:HGNC:24126] | -1.80 | 0.02459 |
| ENSG00000196497 | IPO4 | importin 4 [Source:HGNC Symbol;Acc:HGNC:19426] | -0.50 | 0.02473 |
| ENSG00000107819 | SFXN3 | sideroflexin 3 [Source:HGNC Symbol;Acc:HGNC:16087] | 0.62 | 0.02497 |
| ENSG00000197121 | PGAP1 | post-GPI attachment to proteins inositol deacylase 1 [Source:HGNC Symbol;Acc:HGNC:257 | 0.55 | 0.02498 |
| ENSG00000135074 | ADAM19 | ADAM metallopeptidase domain 19 [Source:HGNC Symbol;Acc:HGNC:197] | 1.04 | 0.02502 |
| ENSG00000005884 | ITGA3 | integrin subunit alpha 3 [Source:HGNC Symbol;Acc:HGNC:6139] | 0.67 | 0.02511 |
| ENSG00000196090 | PTPRT | protein tyrosine phosphatase receptor type T [Source:HGNC Symbol;Acc:HGNC:9682] | 1.80 | 0.02532 |
| ENSG00000080224 | EPHA6 | EPH receptor A6 [Source:HGNC Symbol;Acc:HGNC:19296] | 0.93 | 0.02558 |
| ENSG00000112379 | ARFGEF3 | ARFGEF family member 3 [Source:HGNC Symbol;Acc:HGNC:21213] | 1.87 | 0.02558 |
| ENSG00000165071 | TMEM71 | transmembrane protein 71 [Source:HGNC Symbol;Acc:HGNC:26572] | 1.23 | 0.02567 |
| ENSG00000261420 | NA | novel transcript, antisense to BRP44L | -1.09 | 0.02567 |
| ENSG00000054690 | PLEKHH1 | pleckstrin homology, MyTH4 and FERM domain containing H1 [Source:HGNC Symbol;Acc:H | 0.56 | 0.02590 |
| ENSG00000088836 | SLC4A11 | solute carrier family 4 member 11 [Source:HGNC Symbol;Acc:HGNC:16438] | 0.86 | 0.02641 |
| ENSG00000228725 | MTND2P12 | MT-ND2 pseudogene 12 [Source:HGNC Symbol;Acc:HGNC:42113] | 1.43 | 0.02667 |
| ENSG00000103335 | PIEZO1 | piezo type mechanosensitive ion channel component 1 [Source:HGNC Symbol;Acc:HGNC:2 | 1.14 | 0.02725 |
| ENSG00000099365 | STX1B | syntaxin 1B [Source:HGNC Symbol;Acc:HGNC:18539] | 0.71 | 0.02740 |
| ENSG00000050030 | NEXMIF | neurite extension and migration factor [Source:HGNC Symbol;Acc:HGNC:29433] | -0.65 | 0.02829 |
| ENSG00000181690 | PLAG1 | PLAG1 zinc finger [Source:HGNC Symbol;Acc:HGNC:9045] | 1.05 | 0.02836 |
| ENSG00000125730 | C3 | complement C3 [Source:HGNC Symbol;Acc:HGNC:1318] | 1.63 | 0.02847 |
| ENSG00000142149 | HUNK | hormonally up-regulated Neu-associated kinase [Source:HGNC Symbol;Acc:HGNC:13326] | 1.54 | 0.02857 |
| ENSG00000112837 | TBX18 | T-box transcription factor 18 [Source:HGNC Symbol;Acc:HGNC:11595] | 1.13 | 0.02879 |
| ENSG00000267169 | NA | novel transcript, antisense to LPHN1 | 0.56 | 0.02879 |
| ENSG00000257181 | NA | novel transcript | 0.50 | 0.02896 |
| ENSG00000230316 | FEZF1-AS1 | FEZF1 antisense RNA 1 [Source:HGNC Symbol;Acc:HGNC:41001] | 0.87 | 0.02916 |
| ENSG00000134343 | ANO3 | anoctamin 3 [Source:HGNC Symbol;Acc:HGNC:14004] | 0.51 | 0.02956 |
| ENSG00000165105 | RASEF | RAS and EF-hand domain containing [Source:HGNC Symbol;Acc:HGNC:26464] | 0.72 | 0.02956 |
| ENSG00000206337 | HCP5 | HLA complex P5 [Source:HGNC Symbol;Acc:HGNC:21659] | 0.61 | 0.03008 |
| ENSG00000146054 | TRIM7 | tripartite motif containing 7 [Source:HGNC Symbol;Acc:HGNC:16278] | 0.60 | 0.03020 |
| ENSG00000063127 | SLC6A16 | solute carrier family 6 member 16 [Source:HGNC Symbol;Acc:HGNC:13622] | 0.98 | 0.03036 |
| ENSG00000134363 | FST | follistatin [Source:HGNC Symbol;Acc:HGNC:3971] | 0.79 | 0.03039 |
| ENSG00000120756 | PLS1 | plastin 1 [Source:HGNC Symbol;Acc:HGNC:9090] | 0.53 | 0.03063 |
| ENSG00000136490 | LIMD2 | LIM domain containing 2 [Source:HGNC Symbol;Acc:HGNC:28142] | 0.75 | 0.03071 |

| | | | | |
|---|---|---|---|---|
| ENSG00000164949 | GEM | GTP binding protein overexpressed in skeletal muscle [Source:HGNC Symbol;Acc:HGNC:42 | 0.61 | 0.03078 |
| ENSG00000189056 | RELN | reelin [Source:HGNC Symbol;Acc:HGNC:9957] | 0.51 | 0.03098 |
| ENSG00000161653 | NAGS | N-acetylglutamate synthase [Source:HGNC Symbol;Acc:HGNC:17996] | 0.77 | 0.03124 |
| ENSG00000178860 | MSC | musculin [Source:HGNC Symbol;Acc:HGNC:7321] | 0.97 | 0.03138 |
| ENSG00000130653 | PNPLA7 | patatin like phospholipase domain containing 7 [Source:HGNC Symbol;Acc:HGNC:24768] | 0.70 | 0.03148 |
| ENSG00000251623 | NA | centrosomal protein 192 (CEP192), pseudogene | 0.85 | 0.03203 |
| ENSG00000285412 | NA | novel transcript, antisense to CPVL | 1.22 | 0.03226 |
| ENSG00000173267 | SNCG | synuclein gamma [Source:HGNC Symbol;Acc:HGNC:11141] | -0.52 | 0.03232 |
| ENSG00000165886 | UBTD1 | ubiquitin domain containing 1 [Source:HGNC Symbol;Acc:HGNC:25683] | 0.62 | 0.03243 |
| ENSG00000188649 | CC2D2B | coiled-coil and C2 domain containing 2B [Source:HGNC Symbol;Acc:HGNC:31666] | 0.97 | 0.03283 |
| ENSG00000266010 | GATA6-AS1 | GATA6 antisense RNA 1 (head to head) [Source:HGNC Symbol;Acc:HGNC:48840] | 0.76 | 0.03308 |
| ENSG00000054967 | RELT | RELT TNF receptor [Source:HGNC Symbol;Acc:HGNC:13764] | 0.64 | 0.03324 |
| ENSG00000179630 | LACC1 | laccase domain containing 1 [Source:HGNC Symbol;Acc:HGNC:26789] | -0.91 | 0.03356 |
| ENSG00000162913 | OBSCN-AS1 | OBSCN antisense RNA 1 [Source:HGNC Symbol;Acc:HGNC:32047] | 0.84 | 0.03382 |
| ENSG00000128567 | PODXL | podocalyxin like [Source:HGNC Symbol;Acc:HGNC:9171] | 0.61 | 0.03384 |
| ENSG00000144749 | LRIG1 | leucine rich repeats and immunoglobulin like domains 1 [Source:HGNC Symbol;Acc:HGNC: | 1.25 | 0.03408 |
| ENSG00000168477 | TNXB | tenascin XB [Source:HGNC Symbol;Acc:HGNC:11976] | 0.56 | 0.03412 |
| ENSG00000113971 | NPHP3 | nephrocystin 3 [Source:HGNC Symbol;Acc:HGNC:7907] | 0.58 | 0.03423 |
| ENSG00000173221 | GLRX | glutaredoxin [Source:HGNC Symbol;Acc:HGNC:4330] | 0.58 | 0.03428 |
| ENSG00000131471 | AOC3 | amine oxidase copper containing 3 [Source:HGNC Symbol;Acc:HGNC:550] | 0.70 | 0.03437 |
| ENSG00000064489 | BORCS8-MEF2B | BORCS8-MEF2B readthrough [Source:HGNC Symbol;Acc:HGNC:39979] | 1.16 | 0.03488 |
| ENSG00000130598 | TNNI2 | troponin I2, fast skeletal type [Source:HGNC Symbol;Acc:HGNC:11946] | 0.61 | 0.03505 |
| ENSG00000216937 | CCDC7 | coiled-coil domain containing 7 [Source:HGNC Symbol;Acc:HGNC:26533] | -1.00 | 0.03568 |
| ENSG00000123146 | ADGRE5 | adhesion G protein-coupled receptor E5 [Source:HGNC Symbol;Acc:HGNC:1711] | 1.12 | 0.03576 |
| ENSG00000110328 | GALNT18 | polypeptide N-acetylgalactosaminyltransferase 18 [Source:HGNC Symbol;Acc:HGNC:3048[illegible] | -0.93 | 0.03609 |
| ENSG00000178922 | HYI | hydroxypyruvate isomerase (putative) [Source:HGNC Symbol;Acc:HGNC:26948] | 1.25 | 0.03611 |
| ENSG00000177108 | ZDHHC22 | zinc finger DHHC-type palmitoyltransferase 22 [Source:HGNC Symbol;Acc:HGNC:20106] | 0.73 | 0.03637 |
| ENSG00000006016 | CRLF1 | cytokine receptor like factor 1 [Source:HGNC Symbol;Acc:HGNC:2364] | 0.64 | 0.03638 |
| ENSG00000121361 | KCNJ8 | potassium inwardly rectifying channel subfamily J member 8 [Source:HGNC Symbol;Acc:HG | 0.53 | 0.03692 |
| ENSG00000151689 | INPP1 | inositol polyphosphate-1-phosphatase [Source:HGNC Symbol;Acc:HGNC:6071] | 0.52 | 0.03692 |
| ENSG00000233581 | NA | novel transcript | 0.93 | 0.03736 |
| ENSG00000130635 | COL5A1 | collagen type V alpha 1 chain [Source:HGNC Symbol;Acc:HGNC:2209] | 1.42 | 0.03741 |
| ENSG00000159712 | ANKRD18CP | ankyrin repeat domain 18C, pseudogene [Source:HGNC Symbol;Acc:HGNC:43601] | 1.15 | 0.03750 |
| ENSG00000196154 | S100A4 | S100 calcium binding protein A4 [Source:HGNC Symbol;Acc:HGNC:10494] | 0.94 | 0.03750 |
| ENSG00000163703 | CRELD1 | cysteine rich with EGF like domains 1 [Source:HGNC Symbol;Acc:HGNC:14630] | 0.52 | 0.03779 |
| ENSG00000225580 | NA | proliferation-associated 2G4, 38kD (PA2G4) pseudogene | -0.98 | 0.03867 |
| ENSG00000175592 | FOSL1 | FOS like 1, AP-1 transcription factor subunit [Source:HGNC Symbol;Acc:HGNC:13718] | 1.90 | 0.03902 |
| ENSG00000260077 | NA | novel transcript | -0.87 | 0.03906 |
| ENSG00000145949 | MYLK4 | myosin light chain kinase family member 4 [Source:HGNC Symbol;Acc:HGNC:27972] | 0.84 | 0.03942 |
| ENSG00000177640 | CASC2 | cancer susceptibility 2 [Source:HGNC Symbol;Acc:HGNC:22933] | -0.63 | 0.04000 |

| | | | | |
|---|---|---|---|---|
| ENSG00000273142 | LINC02604 | long intergenic non-protein coding RNA 2604 [Source:HGNC Symbol;Acc:HGNC:53972] | -0.53 | 0.04001 |
| ENSG00000254860 | TMEM9B-AS1 | TMEM9B antisense RNA 1 [Source:HGNC Symbol;Acc:HGNC:19230] | -0.97 | 0.04037 |
| ENSG00000105357 | MYH14 | myosin heavy chain 14 [Source:HGNC Symbol;Acc:HGNC:23212] | 0.58 | 0.04045 |
| ENSG00000126785 | RHOJ | ras homolog family member J [Source:HGNC Symbol;Acc:HGNC:688] | 0.53 | 0.04045 |
| ENSG00000197122 | SRC | SRC proto-oncogene, non-receptor tyrosine kinase [Source:HGNC Symbol;Acc:HGNC:1128 | 0.86 | 0.04045 |
| ENSG00000133103 | COG6 | component of oligomeric golgi complex 6 [Source:HGNC Symbol;Acc:HGNC:18621] | 0.64 | 0.04052 |
| ENSG00000057704 | TMCC3 | transmembrane and coiled-coil domain family 3 [Source:HGNC Symbol;Acc:HGNC:29199] | -0.82 | 0.04102 |
| ENSG00000091622 | PITPNM3 | PITPNM family member 3 [Source:HGNC Symbol;Acc:HGNC:21043] | 0.53 | 0.04123 |
| ENSG00000166106 | ADAMTS15 | ADAM metallopeptidase with thrombospondin type 1 motif 15 [Source:HGNC Symbol;Acc:H( | -1.02 | 0.04138 |
| ENSG00000100065 | CARD10 | caspase recruitment domain family member 10 [Source:HGNC Symbol;Acc:HGNC:16422] | 0.50 | 0.04153 |
| ENSG00000225177 | NA | novel transcript | 1.27 | 0.04153 |
| ENSG00000185022 | MAFF | MAF bZIP transcription factor F [Source:HGNC Symbol;Acc:HGNC:6780] | 1.12 | 0.04168 |
| ENSG00000229589 | ACVR2B-AS1 | ACVR2B antisense RNA 1 [Source:HGNC Symbol;Acc:HGNC:44161] | -0.53 | 0.04168 |
| ENSG00000080200 | CRYBG3 | crystallin beta-gamma domain containing 3 [Source:HGNC Symbol;Acc:HGNC:34427] | 0.77 | 0.04220 |
| ENSG00000111728 | ST8SIA1 | ST8 alpha-N-acetyl-neuraminide alpha-2,8-sialyltransferase 1 [Source:HGNC Symbol;Acc:H | 1.01 | 0.04220 |
| ENSG00000122756 | CNTFR | ciliary neurotrophic factor receptor [Source:HGNC Symbol;Acc:HGNC:2170] | 0.91 | 0.04220 |
| ENSG00000075461 | CACNG4 | calcium voltage-gated channel auxiliary subunit gamma 4 [Source:HGNC Symbol;Acc:HGN( | 0.69 | 0.04247 |
| ENSG00000160447 | PKN3 | protein kinase N3 [Source:HGNC Symbol;Acc:HGNC:17999] | 0.53 | 0.04275 |
| ENSG00000072682 | P4HA2 | prolyl 4-hydroxylase subunit alpha 2 [Source:HGNC Symbol;Acc:HGNC:8547] | 0.63 | 0.04283 |
| ENSG00000129244 | ATP1B2 | ATPase Na+/K+ transporting subunit beta 2 [Source:HGNC Symbol;Acc:HGNC:805] | 0.54 | 0.04292 |
| ENSG00000188641 | DPYD | dihydropyrimidine dehydrogenase [Source:HGNC Symbol;Acc:HGNC:3012] | -1.10 | 0.04292 |
| ENSG00000230876 | LINC00486 | long intergenic non-protein coding RNA 486 [Source:HGNC Symbol;Acc:HGNC:42946] | -1.33 | 0.04301 |
| ENSG00000146648 | EGFR | epidermal growth factor receptor [Source:HGNC Symbol;Acc:HGNC:3236] | 1.14 | 0.04323 |
| ENSG00000197321 | SVIL | supervillin [Source:HGNC Symbol;Acc:HGNC:11480] | -0.84 | 0.04361 |
| ENSG00000146373 | RNF217 | ring finger protein 217 [Source:HGNC Symbol;Acc:HGNC:21487] | 2.04 | 0.04370 |
| ENSG00000067057 | PFKP | phosphofructokinase, platelet [Source:HGNC Symbol;Acc:HGNC:8878] | 0.51 | 0.04376 |
| ENSG00000111186 | WNT5B | Wnt family member 5B [Source:HGNC Symbol;Acc:HGNC:16265] | 0.98 | 0.04397 |
| ENSG00000162817 | C1orf115 | chromosome 1 open reading frame 115 [Source:HGNC Symbol;Acc:HGNC:25873] | 0.65 | 0.04400 |
| ENSG00000121905 | HPCA | hippocalcin [Source:HGNC Symbol;Acc:HGNC:5144] | 1.53 | 0.04435 |
| ENSG00000205809 | KLRC2 | killer cell lectin like receptor C2 [Source:HGNC Symbol;Acc:HGNC:6375] | 2.69 | 0.04440 |
| ENSG00000124406 | ATP8A1 | ATPase phospholipid transporting 8A1 [Source:HGNC Symbol;Acc:HGNC:13531] | 0.54 | 0.04441 |
| ENSG00000078114 | NEBL | nebulette [Source:HGNC Symbol;Acc:HGNC:16932] | 0.93 | 0.04465 |
| ENSG00000170989 | S1PR1 | sphingosine-1-phosphate receptor 1 [Source:HGNC Symbol;Acc:HGNC:3165] | -0.87 | 0.04540 |
| ENSG00000284461 | NA | novel transcript | 1.80 | 0.04664 |
| ENSG00000244026 | FAM86DP | family with sequence similarity 86 member D, pseudogene [Source:HGNC Symbol;Acc:HGN | -0.51 | 0.04686 |
| ENSG00000136010 | ALDH1L2 | aldehyde dehydrogenase 1 family member L2 [Source:HGNC Symbol;Acc:HGNC:26777] | -0.66 | 0.04690 |
| ENSG00000165181 | SHOC1 | shortage in chiasmata 1 [Source:HGNC Symbol;Acc:HGNC:26535] | 0.52 | 0.04702 |
| ENSG00000135373 | EHF | ETS homologous factor [Source:HGNC Symbol;Acc:HGNC:3246] | -1.25 | 0.04711 |
| ENSG00000112769 | LAMA4 | laminin subunit alpha 4 [Source:HGNC Symbol;Acc:HGNC:6484] | -0.54 | 0.04744 |
| ENSG00000127530 | OR7C1 | olfactory receptor family 7 subfamily C member 1 [Source:HGNC Symbol;Acc:HGNC:8373] | 1.24 | 0.04768 |

| | | | | |
|---|---|---|---|---|
| ENSG00000231528 | FAM225A | family with sequence similarity 225 member A [Source:HGNC Symbol;Acc:HGNC:27855] | -0.50 | 0.04775 |
| ENSG00000165617 | DACT1 | dishevelled binding antagonist of beta catenin 1 [Source:HGNC Symbol;Acc:HGNC:17748] | 0.52 | 0.04790 |
| ENSG00000172456 | FGGY | FGGY carbohydrate kinase domain containing [Source:HGNC Symbol;Acc:HGNC:25610] | -0.88 | 0.04790 |
| ENSG00000231625 | SLC47A1P2 | SLC47A1 pseudogene 2 [Source:HGNC Symbol;Acc:HGNC:53866] | 0.94 | 0.04826 |
| ENSG00000232160 | RAP2C-AS1 | RAP2C antisense RNA 1 [Source:HGNC Symbol;Acc:HGNC:40957] | -0.61 | 0.04846 |
| ENSG00000204060 | FOXO6 | forkhead box O6 [Source:HGNC Symbol;Acc:HGNC:24814] | -0.55 | 0.04901 |
| ENSG00000213261 | EEF1B2P6 | eukaryotic translation elongation factor 1 beta 2 pseudogene 6 [Source:HGNC Symbol;Acc:[illegible] | -1.08 | 0.04902 |
| ENSG00000286257 | NA | novel transcript | 0.53 | 0.04902 |
| ENSG00000228742 | LINC02577 | long intergenic non-protein coding RNA 2577 [Source:HGNC Symbol;Acc:HGNC:53749] | -1.42 | 0.04931 |
| ENSG00000126010 | GRPR | gastrin releasing peptide receptor [Source:HGNC Symbol;Acc:HGNC:4609] | -0.85 | 0.04933 |
| ENSG00000280893 | NA | novel transcript | -1.42 | 0.04933 |
| ENSG00000143842 | SOX13 | SRY-box transcription factor 13 [Source:HGNC Symbol;Acc:HGNC:11192] | -0.60 | 0.04954 |
| ENSG00000276740 | NA | novel transcript, sense intronic to FAM155A | 0.95 | 0.04981 |
