## Supplemental Table 11 for "Modulation of calcium signaling on demand to decipher the molecular mechanisms of primary aldosteronism"

**Supplementary Table 11. List of significantly enriched pathways in H295R_S2 cells expressing the α7-5HT3 receptor in response to 24h treatment with 12 mM K+**

| Pathway | Fold Enrichment | PValue |
|---|---|---|
| GO:0007155~cell adhesion | 2.49 | 2.43E-08 |
| GO:0098609~cell-cell adhesion | 3.22 | 9.08E-06 |
| hsa04261:Adrenergic signaling in cardiomyocytes | 3.15 | 2.98E-05 |
| GO:0007399~nervous system development | 2.29 | 3.22E-05 |
| hsa04512:ECM-receptor interaction | 4.01 | 3.78E-05 |
| GO:0001822~kidney development | 3.8 | 4.01E-05 |
| hsa04510:Focal adhesion | 2.76 | 4.21E-05 |
| hsa05410:Hypertrophic cardiomyopathy | 3.97 | 4.27E-05 |
| GO:0030198~extracellular matrix organization | 3.05 | 5.46E-05 |
| hsa05414:Dilated cardiomyopathy | 3.72 | 8.48E-05 |
| GO:0010976~positive regulation of neuron projection development | 3.57 | 1.47E-04 |
| hsa05412:Arrhythmogenic right ventricular cardiomyopathy | 3.97 | 1.83E-04 |
| GO:0017157~regulation of exocytosis | 6.46 | 1.94E-04 |
| GO:0007420~brain development | 2.52 | 1.96E-04 |
| GO:0007411~axon guidance | 2.83 | 2.27E-04 |
| GO:0030168~platelet activation | 4.39 | 4.18E-04 |
| hsa05165:Human papillomavirus infection | 2.08 | 5.28E-04 |
| GO:0043547~positive regulation of GTPase activity | 2.69 | 6.33E-04 |
| GO:0007613~memory | 3.78 | 6.42E-04 |
| hsa04974:Protein digestion and absorption | 3.22 | 6.52E-04 |
| GO:2001135~regulation of endocytic recycling | 19.91 | 6.90E-04 |
| GO:0014912~negative regulation of smooth muscle cell migration | 11.49 | 6.95E-04 |
| hsa05032:Morphine addiction | 3.36 | 7.96E-04 |
| GO:0001764~neuron migration | 3.18 | 8.01E-04 |
| GO:1904322~cellular response to forskolin | 10.67 | 9.48E-04 |
| GO:0030199~collagen fibril organization | 3.93 | 9.54E-04 |
| GO:0006904~vesicle docking involved in exocytosis | 5.97 | 9.67E-04 |
| GO:0001525~angiogenesis | 2.34 | 9.90E-04 |
| hsa04934:Cushing syndrome | 2.63 | 0.0011 |
| GO:0007229~integrin-mediated signaling pathway | 3.23 | 0.0012 |
| GO:0007165~signal transduction | 1.49 | 0.0014 |
| hsa04925:Aldosterone synthesis and secretion | 3.12 | 0.0015 |
| GO:0045446~endothelial cell differentiation | 9.33 | 0.0016 |
| GO:0009410~response to xenobiotic stimulus | 2.24 | 0.0022 |
| GO:0006198~cAMP catabolic process | 13.27 | 0.0027 |
| hsa04151:PI3K-Akt signaling pathway | 1.87 | 0.0030 |
| hsa00330:Arginine and proline metabolism | 4.08 | 0.0031 |
| hsa04024:cAMP signaling pathway | 2.15 | 0.0031 |
| GO:0061744~motor behavior | 7.86 | 0.0032 |
| hsa01100:Metabolic pathways | 1.32 | 0.0050 |
| GO:0007189~adenylate cyclase-activating G-protein coupled receptor signalir | 2.69 | 0.0051 |
| hsa04068:FoxO signaling pathway | 2.53 | 0.0051 |
| hsa05200:Pathways in cancer | 1.63 | 0.0053 |
| GO:0034765~regulation of ion transmembrane transport | 2.67 | 0.0054 |
| GO:0071456~cellular response to hypoxia | 2.67 | 0.0054 |
| GO:0043542~endothelial cell migration | 5.12 | 0.0058 |
| GO:0016055~Wnt signaling pathway | 2.30 | 0.0060 |
| hsa00480:Glutathione metabolism | 3.58 | 0.0065 |
| GO:0022407~regulation of cell-cell adhesion | 9.96 | 0.0065 |
| GO:0001837~epithelial to mesenchymal transition | 4.10 | 0.0068 |
| h_agrPathway:Agrin in Postsynaptic Differentiation | 3.89 | 0.0072 |
| hsa04921:Oxytocin signaling pathway | 2.32 | 0.0072 |
| GO:0030335~positive regulation of cell migration | 2.05 | 0.0072 |
| GO:0060412~ventricular septum morphogenesis | 4.84 | 0.0074 |
| GO:0051017~actin filament bundle assembly | 4.84 | 0.0074 |
| GO:0001666~response to hypoxia | 2.41 | 0.0080 |
| GO:1902895~positive regulation of pri-miRNA transcription from RNA polymer | 3.94 | 0.0082 |
| GO:0086064~cell communication by electrical coupling involved in cardiac coı | 9.19 | 0.0083 |
| hsa01230:Biosynthesis of amino acids | 3.06 | 0.0087 |

| | | |
|---|---|---|
| hsa05207:Chemical carcinogenesis - receptor activation | 2.04 | 0.0089 |
| GO:0019222~regulation of metabolic process | 5.97 | 0.0090 |
| GO:0015914~phospholipid transport | 4.60 | 0.0092 |
| hsa04971:Gastric acid secretion | 3.02 | 0.0094 |
| GO:0009611~response to wounding | 3.37 | 0.0094 |
| GO:0010719~negative regulation of epithelial to mesenchymal transition | 4.48 | 0.0103 |
| GO:0046697~decidualization | 5.74 | 0.0103 |
| GO:0006595~polyamine metabolic process | 17.92 | 0.0104 |
| GO:0007417~central nervous system development | 2.42 | 0.0110 |
| GO:0030036~actin cytoskeleton organization | 2.21 | 0.0112 |
| GO:0033627~cell adhesion mediated by integrin | 4.37 | 0.0114 |
| GO:0007193~adenylate cyclase-inhibiting G-protein coupled receptor signalin | 3.67 | 0.0117 |
| hsa04360:Axon guidance | 2.10 | 0.0119 |
| GO:0035556~intracellular signal transduction | 1.69 | 0.0119 |
| GO:0001938~positive regulation of endothelial cell proliferation | 3.19 | 0.0126 |
| GO:0060048~cardiac muscle contraction | 4.27 | 0.0126 |
| GO:0001942~hair follicle development | 4.27 | 0.0126 |
| hsa04725:Cholinergic synapse | 2.48 | 0.0128 |
| hsa04927:Cortisol synthesis and secretion | 3.14 | 0.0130 |
| GO:0050900~leukocyte migration | 5.33 | 0.0134 |
| hsa04022:cGMP-PKG signaling pathway | 2.14 | 0.0137 |
| GO:0043409~negative regulation of MAPK cascade | 4.17 | 0.0139 |
| hsa05225:Hepatocellular carcinoma | 2.12 | 0.0144 |
| GO:0007015~actin filament organization | 2.33 | 0.0145 |
| GO:0007612~learning | 3.48 | 0.0148 |
| GO:0051965~positive regulation of synapse assembly | 3.48 | 0.0148 |
| GO:0071625~vocalization behavior | 7.47 | 0.0150 |
| hsa00564:Glycerophospholipid metabolism | 2.58 | 0.0151 |
| GO:0006704~glucocorticoid biosynthetic process | 14.93 | 0.0153 |
| GO:2000427~positive regulation of apoptotic cell clearance | 14.93 | 0.0153 |
| GO:0061299~retina vasculature morphogenesis in camera-type eye | 14.93 | 0.0153 |
| GO:1902747~negative regulation of lens fiber cell differentiation | 14.93 | 0.0153 |
| GO:0031630~regulation of synaptic vesicle fusion to presynaptic membrane | 14.93 | 0.0153 |
| GO:0007169~transmembrane receptor protein tyrosine kinase signaling pathw | 2.43 | 0.0153 |
| GO:0019933~cAMP-mediated signaling | 4.98 | 0.0170 |
| GO:0007219~Notch signaling pathway | 2.53 | 0.0175 |
| hsa04012:ErbB signaling pathway | 2.70 | 0.0177 |
| GO:0021675~nerve development | 7.03 | 0.0178 |
| GO:0060384~innervation | 7.03 | 0.0178 |
| GO:0006661~phosphatidylinositol biosynthetic process | 3.90 | 0.0182 |
| GO:0048812~neuron projection morphogenesis | 3.32 | 0.0185 |
| hsa04911:Insulin secretion | 2.67 | 0.0189 |
| hsa05202:Transcriptional misregulation in cancer | 1.98 | 0.0190 |
| GO:0009653~anatomical structure morphogenesis | 2.33 | 0.0201 |
| GO:0051145~smooth muscle cell differentiation | 6.64 | 0.0209 |
| GO:0048839~inner ear development | 3.73 | 0.0215 |
| GO:0000122~negative regulation of transcription from RNA polymerase II pror | 1.39 | 0.0217 |
| GO:0045944~positive regulation of transcription from RNA polymerase II prom | 1.34 | 0.0218 |
| GO:0030336~negative regulation of cell migration | 2.10 | 0.0219 |
| hsa04928:Parathyroid hormone synthesis, secretion and action | 2.41 | 0.0226 |
| GO:0016477~cell migration | 1.85 | 0.0229 |
| GO:0030154~cell differentiation | 1.47 | 0.0250 |
| hsa04115:p53 signaling pathway | 2.76 | 0.0251 |
| hsa04350:TGF-beta signaling pathway | 2.36 | 0.0252 |
| GO:0007507~heart development | 1.97 | 0.0264 |
| hsa05222:Small cell lung cancer | 2.49 | 0.0271 |
| hsa04010:MAPK signaling pathway | 1.69 | 0.0274 |
| GO:0021591~ventricular system development | 5.97 | 0.0278 |
| GO:2000179~positive regulation of neural precursor cell proliferation | 5.97 | 0.0278 |
| GO:0031290~retinal ganglion cell axon guidance | 5.97 | 0.0278 |
| GO:0030279~negative regulation of ossification | 5.97 | 0.0278 |
| GO:0030182~neuron differentiation | 2.11 | 0.0279 |

| | | |
|---|---|---|
| GO:0015701~bicarbonate transport | 4.27 | 0.0285 |
| hsa04926:Relaxin signaling pathway | 2.17 | 0.0295 |
| GO:0007160~cell-matrix adhesion | 2.47 | 0.0298 |
| hsa04072:Phospholipase D signaling pathway | 2.07 | 0.0302 |
| GO:0009615~response to virus | 2.44 | 0.0312 |
| GO:0001657~ureteric bud development | 4.15 | 0.0313 |
| GO:0007274~neuromuscular synaptic transmission | 5.69 | 0.0316 |
| GO:0050731~positive regulation of peptidyl-tyrosine phosphorylation | 2.63 | 0.0327 |
| GO:0010718~positive regulation of epithelial to mesenchymal transition | 3.32 | 0.0339 |
| GO:2001237~negative regulation of extrinsic apoptotic signaling pathway | 4.04 | 0.0342 |
| GO:0045787~positive regulation of cell cycle | 4.04 | 0.0342 |
| GO:0043524~negative regulation of neuron apoptotic process | 2.13 | 0.0343 |
| GO:0061337~cardiac conduction | 9.96 | 0.0344 |
| GO:0042416~dopamine biosynthetic process | 9.96 | 0.0344 |
| GO:0060856~establishment of blood-brain barrier | 9.96 | 0.0344 |
| GO:0035264~multicellular organism growth | 2.60 | 0.0344 |
| hsa04070:Phosphatidylinositol signaling system | 2.37 | 0.0357 |
| GO:0016125~sterol metabolic process | 5.43 | 0.0357 |
| GO:0014911~positive regulation of smooth muscle cell migration | 5.43 | 0.0357 |
| GO:0007265~Ras protein signal transduction | 2.57 | 0.0362 |
| GO:0042327~positive regulation of phosphorylation | 3.93 | 0.0372 |
| GO:0042311~vasodilation | 3.93 | 0.0372 |
| hsa01522:Endocrine resistance | 2.34 | 0.0376 |
| GO:0009966~regulation of signal transduction | 3.20 | 0.0388 |
| GO:0032700~negative regulation of interleukin-17 production | 5.19 | 0.0401 |
| GO:0050919~negative chemotaxis | 3.83 | 0.0404 |
| GO:2000300~regulation of synaptic vesicle exocytosis | 3.14 | 0.0414 |
| GO:0031397~negative regulation of protein ubiquitination | 3.14 | 0.0414 |
| GO:0006813~potassium ion transport | 2.75 | 0.0416 |
| GO:0072112~glomerular visceral epithelial cell differentiation | 8.96 | 0.0420 |
| hsa04710:Circadian rhythm | 3.75 | 0.0423 |
| hsa04390:Hippo signaling pathway | 1.95 | 0.0436 |
| GO:0045165~cell fate commitment | 3.09 | 0.0441 |
| GO:0051924~regulation of calcium ion transport | 4.98 | 0.0447 |
| GO:0046928~regulation of neurotransmitter secretion | 4.98 | 0.0447 |
| hsa04371:Apelin signaling pathway | 2.02 | 0.0456 |
| GO:0055085~transmembrane transport | 1.76 | 0.0482 |
| GO:0001933~negative regulation of protein phosphorylation | 2.65 | 0.0487 |
| GO:0060038~cardiac muscle cell proliferation | 4.78 | 0.0496 |
| GO:0003148~outflow tract septum morphogenesis | 4.78 | 0.0496 |
